# New analysis framework incorporating mixed mutual information and scalable Bayesian networks for multimodal high dimensional genomic and epigenomic cancer data

**DOI:** 10.1101/812446

**Authors:** Xichun Wang, Sergio Branciamore, Grigoriy Gogoshin, Shuyu Ding, Andrei S Rodin

## Abstract

We propose a novel two-stage analysis strategy to discover candidate genes associated with the particular cancer outcomes in large multimodal genomic cancers databases, such as The Cancer Genome Atlas (TCGA). During the first stage, we use mixed mutual information to perform variable selection; during the second stage, we use scalable Bayesian network (BN) modeling to identify candidate genes and their interactions. Two crucial features of the proposed approach are (i) the ability to handle mixed data types (continuous and discrete, genomic, epigenomic, etc.), and (ii) a flexible boundary between the variable selection and network modeling stages --- the boundary that can be adjusted in accordance with the investigators’ BN software scalability and hardware implementation. These two aspects result in high generalizability of the proposed analytical framework. We apply the above strategy to three different TCGA datasets (LGG, Brain Lower Grade Glioma; HNSC, Head and Neck Squamous Cell Carcinoma; STES, Stomach and Esophageal Carcinoma), linking multimodal molecular information (SNPs, mRNA expression, DNA methylation) to two clinical outcome variables (tumor status and patient survival). We identify 11 candidate genes, of which 6 have already been directly implicated in the cancer literature. One novel LGG prognostic factor suggested by our analysis, methylation of TMPRSS11F type II transmembrane serine protease, presents intriguing direction for the follow-up studies.

## Introduction

The Cancer Genome Atlas (TCGA) resource contains genomic data compiled for more than 30 different types/subtypes of cancer [1]. For each type, clinical outcome/progression data (e.g., tumor status and patient survival) for a considerable number of patients is matched to the large-scale molecular data. The latter is multimodal, ranging from genetic (e.g., somatic mutations) to expression (e.g., RNA-seq gene expression) to epigenetic (e.g., promoter methylation) data. Not surprisingly, there is substantial enthusiasm for causally linking the latter to the former using various modeling and secondary data analysis techniques [2–6]. The ultimate goals of these analyses are (i) to gain better mechanistic understanding of the underlying molecular biology of cancer, primarily by identifying important genes and their interactions; (ii) to construct compact and efficient clinical predictors (e.g., prognostic scores, indices and signatures); (iii) to associate the latter with the particular patient groups and subgroups, in the context of personalized/precision medicine. One of the more attractive and popular methods for such multivariate analysis is Bayesian networks (BNs) [7], a well-established fixture in computational systems biology [8]. Among the BN advantages are their probabilistic nature, model flexibility, ability to handle non-additive, higher-order, interactions, and ease of the result interpretation. However, applications of BNs to the TCGA (and TCGA-like) data [9–16] face two principal difficulties: combining mixed data types in a single analysis framework, *and* achieving sufficient (for genomic data) scalability, *simultaneously*. (These, of course, are the two fundamental, and interconnected, BN modeling challenges in general, not just in the TCGA application.) The latest developments in addressing these two challenges encompass more efficient computational approaches [17, 18], and mathematically rigorous and robust methods for handling mixed data, such as mixed local probability models and/or adaptive discretization [18–20]. Nevertheless, resolving both difficulties simultaneously in a generalizable toolkit (seamlessly applicable to, for example, across the individual TCGA datasets) remains elusive. A promising approach to devising such a toolkit would be to precede the comparatively exhaustive NP-hard BN modeling with a variable selection procedure [21, for example], where the full dataset is pared down to a subset of variables most relevant to a particular clinical outcome or phenotype. While alleviating the scalability issue, this, however, could potentially “throw away the wheat with the chaff”, especially if the variable selection process [22, 23] is of a simplistic and overly too restrictive kind (e.g., a statistically conservative univariate filter). There are three possible ways to address this, namely: (i) increase the scalability of the BN modeling to genomic data levels (possible, but impractical for frequent/serial analyses), (ii) incorporate higher-order interactions into the variable selection step (thus “upgrading” it from the simple filter to the wrapper [23–25] --- this is the solution implemented in [21]), or (iii) adjust the transition boundary between the variable selection step and the BN modeling step, depending on the investigators’ computational resources and the nature (dimensionality, sparseness, heterogeneity) of the actual data. It is the third analytical strategy that we propose in this study, with the goal to achieve the optimal compromise between the computational practicality and modeling exhaustiveness.

In our analysis pipeline, we start with the variable selection procedure based on the mixed-type Mixed Mutual Information (MMI) forward selection filter. We compute the MMI values for all available gene-outcome (specifically, tumor status and patient survival) pairs, and use the MMI frequency distribution to select top variables/genes (or, alternatively, to remove bottom variables/genes) before moving on to the BN modeling. This mixed-type measure-based approach to gene selection is the principal innovation of this paper. We then use the maximum entropy (ME) – based discretization to construct the mixed-type BNs using our previously reported scalable BN modeling algorithm and software [18]. Subsequently, we concentrate on the sub-networks centered around the clinical outcome variables of interest, and identify the molecular gene components belonging to these sub-networks.

The proposed analysis strategy has been applied by us to 12 different TCGA cancer datasets. This allowed us to check for robustness, scalability and generalizability. Here, we present the results for the Brain Lower Grade Glioma (LGG), Head and Neck Squamous Cell Carcinoma (HNSC) and Stomach and Esophageal Carcinoma (STES) datasets (all three datasets being reasonably well-populated and proportionally balanced across the different outcomes and molecular data types). For the purposes of this particular analysis, we decided to concentrate on three types of molecular data, one discrete (somatic mutations) and two – continuous (RNA-seq gene expression, and promoter methylation). This selection is reflective of the recent trends in multimodal cancer data analyses [21, 26], makes sense in the broad cancer genetics context [27–32], and underscores the comparative importance of the methylation molecular data [29].

We conclude by identifying a compact list of genes potentially associated with cancer-related clinical phenotypes (tumor status and patient survival), scrutinizing these genes in light of the current literature, and discussing the generalizability of our approach to the different datasets, diseases and molecular data types.

## Materials and Methods

### Data preprocessing

TCGA LGG, HNSC and STES datasets were downloaded for the clinical data (“Clinical_Pick_Tier1 (MD5)”), SNP data (“Mutation_Packager_Calls (MD5)”), expression data (“mRNAseq_Preprocess (MD5)”) and promoter-centric methylation data (“Methylation_Preprocess (MD5)”). Patients were further subdivided into (i) two disease progression categories (according to the “tumor status” variable), and (ii) two patient survival categories (high death risk, with survival less than 2 years, and low death risk, with survival more than 2 years, which is a common cutoff point in recent cancer literature). We further excluded patients with ambiguous or missing outcome variable values (e.g., no survival status, survival status as “living” with survival time less than 2 years, tumor status neither “tumor-free” nor “with tumor”, etc.). These clinical variables (“tumor status” and “2 year survival”) were subsequently used for the variable selection purposes, and, eventually, to extract “tumor status” and “survival” – centered sub-networks from the full BNs. Expression data and methylation data (designated by “E” and “M” below, for brevity) were not discretized at this stage, as both variable selection and BN construction tools in our computational pipeline can, by design, accept mixed (continuous and discreet) variable types. SNP (somatic mutation) data (designated by “S” below) were compressed into a binary variable (presence or absence of at least one non-synonymous mutation in at least one sample of the particular gene).

After filtering out patient records with incomplete, partially missing, or ambiguously labeled data, the final datasets consisted of 4782 genes (LGG), 12516 genes (HNSC) and 16164 genes (STES). 273 patient records were available for LGG/tumor status analysis (140 patients with tumor, 133 without); 213 patients – for LGG/survival (120 patients with survival less than 2 years, 93 with long-term survival). Similarly, 260 patient records were available for HNSC/tumor status analysis (94 patients with tumor, 166 without); 139 patients – for HNSC/survival (40 patients with survival less than 2 years, 99 with long-term survival). Finally, 403 patient records were available for STES/tumor status analysis (147 patients with tumor, 256 without); 258 patients – for STES/survival (191 patients with survival less than 2 years, 67 with long-term survival).

(It is possible to include other different molecular data types and outcome variables, both continuous and discrete, into the proposed framework without substantial alterations to the analysis pipeline, except for some rudimentary data preprocessing.)

### Variable selection

There are very few BN algorithms/software solutions that scale up to (epi)genomic levels (tens to hundreds of thousands of variables) [17, 18]. Even with these, exhaustive analyses require dedicated hardware and weeks of processing time. This might be acceptable for a one-off, “final” analysis, but is clearly impractical for the exploratory research. This is why it is a common practice to carry out variable selection (or feature selection, or feature set reduction) in order to generate a comparatively compact subset of variables to be subsequently fed into the network modeling algorithm/software [23]. Variable selection approaches range from the very simple (univariate filters) to increasingly more sophisticated; at some point, the latter become essentially inseparable from the multivariate modeling methods *per se*. Depending on the dataset to be analyzed, different “couplings” of variable selection and multivariate modeling methods might prove to be more or less effective, and it is difficult to devise *a priori* the objectively optimal combination for each new dataset. For a principally network-centric data analysis approach (innate to the systems biology), it would make sense to feed as many variables into the network-building module as possible, thus “delegating” the resolution of the higher-order / non-additive interactions and conditional independence relationships to the BN algorithm itself. Therefore, for the exploratory research, we suggest that the investigators first define the upper BN scalability limit that they are comfortable with (given the available software/hardware), and then adjust the variable selection cutoff point accordingly. For more “finalized” analysis, that limit should be raised higher (and the variable selection process, consequently, be made less restrictive).

In TCGA dataset (and other similar (epi)genomic resources), there are tens of thousands of potentially predictive/relevant variables (roughly proportional to the number of genes in the human genome). The “hand off” point between the variable selection and BN analysis steps should therefore vary between 100s of variables (for the exploratory and preliminary analyses) and 1,000s of variables (for the final analyses). The actual number might also depend on the shape of the variable selection curve, or on the statistical significance criteria --- we stop adding increasingly less significant variables during the forward variable selection process (or stop removing increasingly more significant variables during the backward variable elimination process) when a certain statistical significance cutoff point is reached [33]. The above considerations were taken into account in the course of this study, as detailed in the Results section below.

It is difficult to integrate the multimodal, mixed-type, data into the variable selection process (filter or wrapper) as, until recently, there has been a paucity of the usable mixed-type metrics. In this study, a recently developed measure, Mixed Mutual Information (MMI) [34], was used to link the gene information (a mixed-type vector consisting of the S, E and M molecular data components for each gene) to the clinical variable (tumor status or 2 year survival) in a “forward-selection-filter” variable selection procedure. MMI is a non-parametric and distribution-free measure (which makes it more attractive than the alternatives, such as linear correlation --- especially in the biological networks context [35, 36]) that is based on the entropy estimates from k-nearest neighbor (k-NN) distances [37]. It is, therefore, sensitive to the choice of the *k* parameter. Lower values of *k* (1-4) tend to lead to higher dispersion, while much higher values (>20) are associated with unnecessarily increased computational complexity and possible overfitting [34, personal communication from Weihao Gao]. We have evaluated different values of *k* on the actual TCGA datasets by measuring the Jaccard index for the pairs of consecutive (in *k*) post-selection variable sets as a function of *k*. The index appeared to stabilize in the 8-20 range in 12 different TCGA datasets analyzed (see Results section below); therefore, *k* was set at 15 throughout this study.

### Bayesian networks modeling

BN modeling, in its basic form, reconstructs a sparse graphical representation of a joint multivariate probability distribution of random variables from a “flat” dataset. Nodes in the network represent random variables, edges --- dependencies. Absence of an edge between the two nodes indicates conditional independence between them. Recent work in BN methodology refinement led to significant progress in scalability --- our latest BN modeling software implementation [18] easily processes datasets up to ∼ 1mln variables x 1mln datapoints. Handling mixed variable types (both continuous and discrete, in a typical application) is still not entirely seamless; it was recently suggested [18, 19] that adaptive discretization (of continuous variables) might be preferable to forcing mixed local probability models. Consequently, we were using maximum entropy – based three-bin discretization throughout this study --- expression data (“E” molecular data component) and methylation data (“M” molecular data component) were discretized into three bins --- which has attractive mathematical properties, and has been shown by us earlier to maintain near-optimal over/under-fitting balance [18].

Detailed description of the BN methodology in general and of our implementation (including applications to other types of high-dimensional biological data) in particular can be found in [18, 38]; here we will only note that (1) our BN implementation uses a hybrid “sparse candidates” + “search-and-score” graduate descent algorithm coupled with various model scoring metrics and maximum entropy-based adaptive discretization; (2) in the resulting BN visualizations, numbers next to the edges and edge “thickness” indicate relative edge strengths (the numbers are the model scores’ ratios for the models with/without corresponding edges, which are proportional to the marginal likelihood ratios); (3) directionality in the network (arrow points attached to the edges, when present) does not necessarily imply the causality flow, and is used predominantly for the mathematical convenience (to avoid cyclic dependencies); (4) when deciphering conditional dependence and independence patterns, it is useful to concentrate on the immediate Markov neighborhood (MN) of a particular variable of interest (such as a clinical outcome). This neighborhood can be roughly defined as all the nodes that are in immediate contact with (“one degree of separation” from) the node representing the aforementioned variable of interest. Under certain conditions, given its MN, the variable of interest is conditionally independent of the remaining variables (rest of the network). Therefore, deriving a MN for a variable of interest is analogous to the variable selection activity, specifically of the embedded variety [23]. The central step in our computational analysis pipeline is using full BN reconstruction to generate the MN for the clinical outcome variable, and then ascertaining the interplay of the (small number of) gene-related variables (S, E and M molecular data components) within that MN. (It should be noted that MN is a simplification of the more rigorous concept of Markov Blanket --- meaning, for our purposes, that sometimes “two degrees of separation” are needed for encapsulating a variable/node of interest.)

## Results

Figure 1 depicts the variable selection process for six possible combinations of two clinical variables (“tumor status” and “survival”) and three TCGA cancer datasets (LGG, HNSC, STES). MMI (mixed mutual information) between (S, E, M) and tumor status/survival was computed for 4782 genes (LGG), 12516 genes (HNSC) and 16164 genes (STES). (All six gene lists, with corresponding MMI values, are available in Supplementary Tables 1-6.). The histogram representation of the MMI distribution, as shown in Figure 1, is convenient, as it allows to evaluate (both visually and quantitatively) the relative predictive values of the top-ranking genes with respect to the outcome variable classification. For the purposes of this study, and to make the resulting full BNs “observable”, we have chosen the “top genes” cutoff value of 99.5% MMI CDF (cumulative distribution function), which leads to the selection of 24 genes (72 future BN nodes/variables in total, comprising 24 S, 24 E, and 24 M components) out of 4728 for two LGG networks, 63 genes (189 nodes/variables) out of 12516 for two HNSC networks, and 81 genes (243 nodes/variables) out of 16164 for two STES networks. Note that the S, E, and M components of each gene vector were considered as the separate nodes/variables in the subsequent BN construction, as at this time we do not have a BN scoring function that can incorporate mixed multivariate distance measures. It should also be noted that although MMI, intuitively, should not be negative, due to the way it is computed it can get into the negative range when (i) continuous variables are involved, and (ii) the number of dimensions is more than two (four, in our case). This said, all the negative MMI values in Figure 1 reside well within the allowed algorithmic negative deviation range, and should not influence the variable rankings [34, personal communication from Weihao Gao].

**Figure 1.**
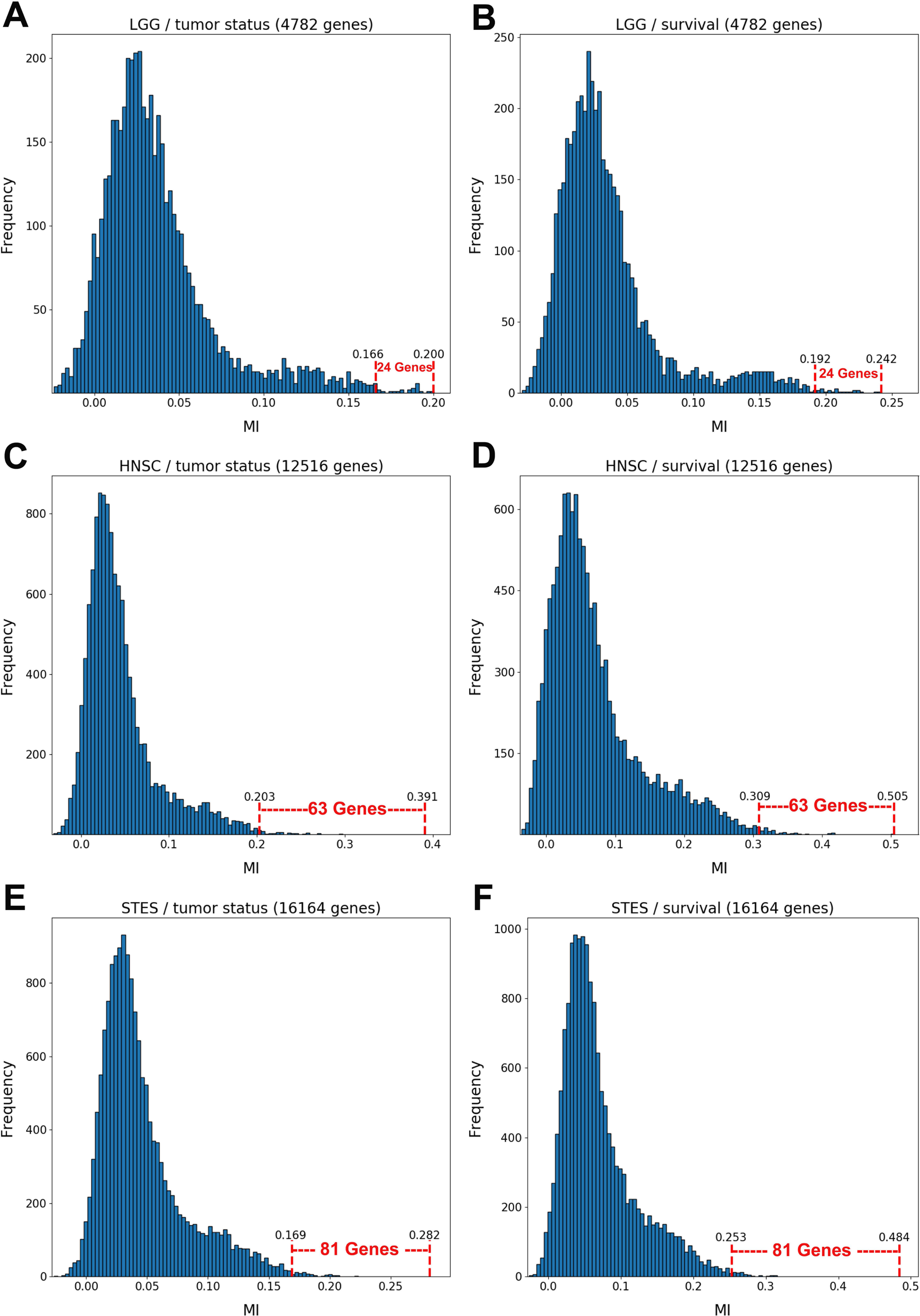
Variable selection process for six combinations of two clinical outcome variables (“tumor status” and “survival”) and three TCGA cancer datasets (LGG, HNSC, STES). MMI between the (S, E, M) molecular data vector and tumor status/survival was computed for 4782 genes (LGG), 12516 genes (HNSC) and 16164 genes (STES). The histogram representation of the MMI distribution is shown with the selection of “top” (i.e., with the MMI CDF > 99.5%) genes superimposed on the right tail of the MMI frequency distribution. (A) LGG/tumor status; (B) LGG/survival; (C) HNSC/tumor status; (D) HNSC/survival; (E) STES/tumor status; (F) STES/survival.

Interestingly, every histogram in Figure 1 has a heavy right tail, which sometimes appears to follow a clear “knee point” --- for example, at MMI ∼= 0.08 in Figures 1ABC. This suggests that MMI > 0.08 could also be used as a “natural” cutoff value, at least in these three datasets.

The variable selection distributions shown in Figure 1 were derived with the MMI parameter *k* set at 15. Figure 2 illustrates the motivation behind that choice, using the LGG/survival dataset example. Shown is the plot of the Jaccard index (JI, a.k.a. set “Intersection over Union”, which is a common measure of sample set similarity) comparing the gene/variable sets resulting from the above variable selection procedure, with cutoff set at 99.5% MMI CDF, where JI(*k*) compares the sets obtained with *k* and *k+1*. It is clear that as k reaches ∼15, the set composition somewhat stabilizes; further increase in k does not seem to offer any advantages. (JI plots for the other datasets exhibit a similar pattern).

**Figure 2.**
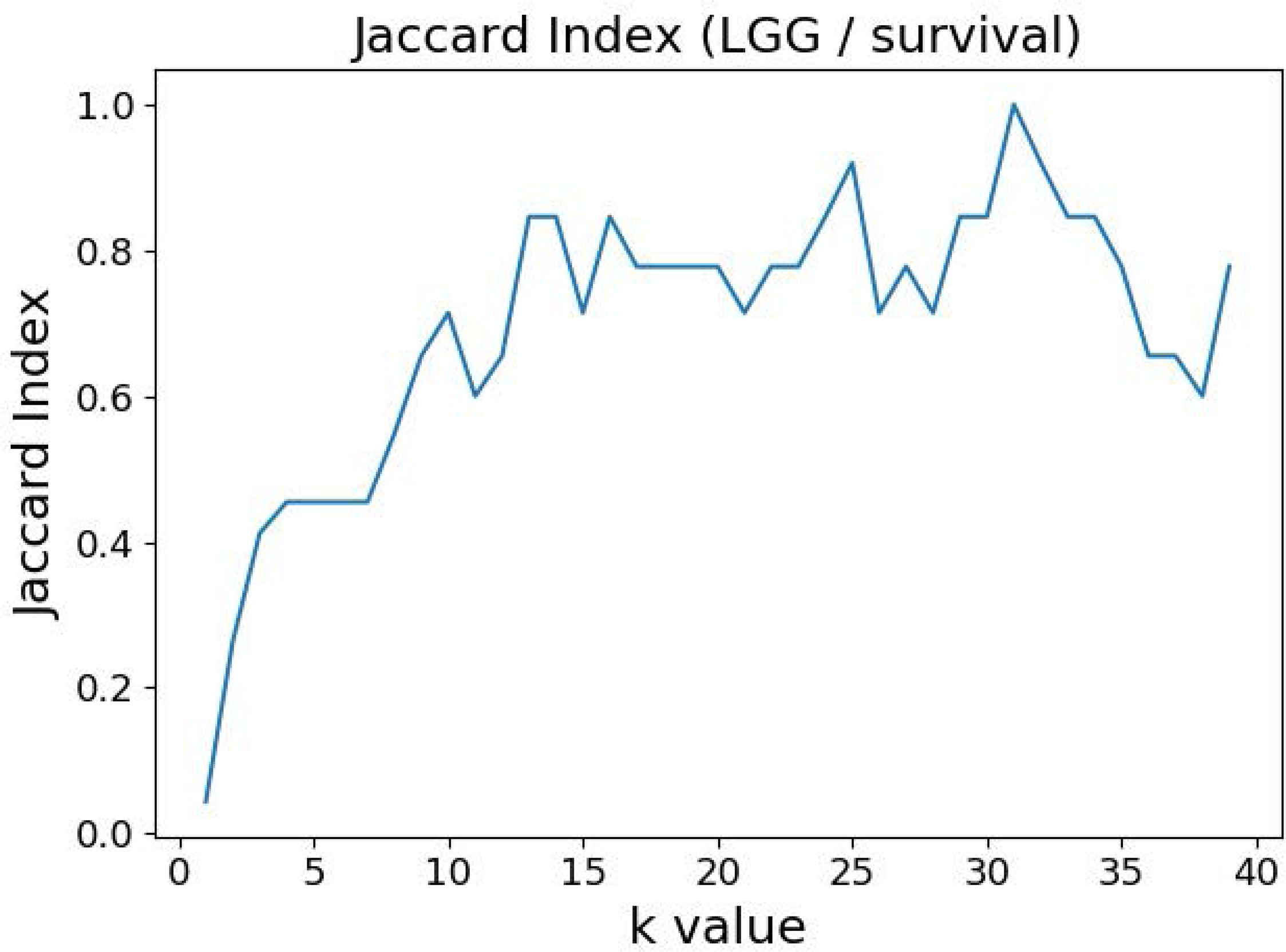
Jaccard index (JI) (“Intersection over Union”) comparing the gene/variable sets resulting from the LGG/survival dataset variable selection with the cutoff set at 99.5% MMI CDF. JI(*k*) compares the sets obtained with *k* and *k+1*.

Figures 3 and 4 depict the full BNs obtained from the LGG/tumor status and LGG/survival datasets. Supplementary Data Sheets 1-6 depict, in PDF format, the full BNs obtained from the LGG/tumor status, LGG/survival, HNSC/tumor status, HNSC/survival, STES/tumor status, and STES/survival datasets, respectively. Six corresponding DOT (standard network / causal graphical models format) files can be found in the Supplementary Tables 7-12.

**Figure 3.**
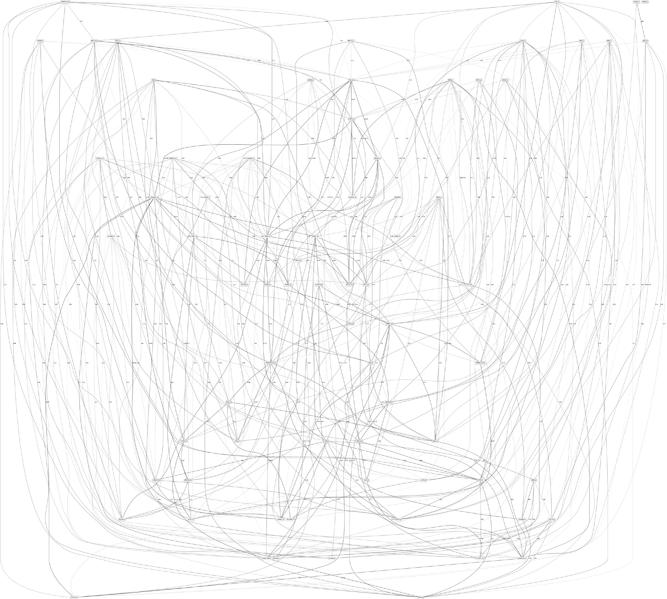
Full BN derived from the LGG/tumor status data. “Tumor_Status” node in the BN is self-explanatory. Other nodes in the networks correspond to the genes / molecular components (gene name_S/E/M). Edges in the network correspond to the dependencies between the nodes. Directionality of the edge (arrow) is for mathematical convenience only and does not imply causation. “Boldness” of the edge is proportional to the dependency strength, also indicated by the number shown next to the edge.

**Figure 4.**
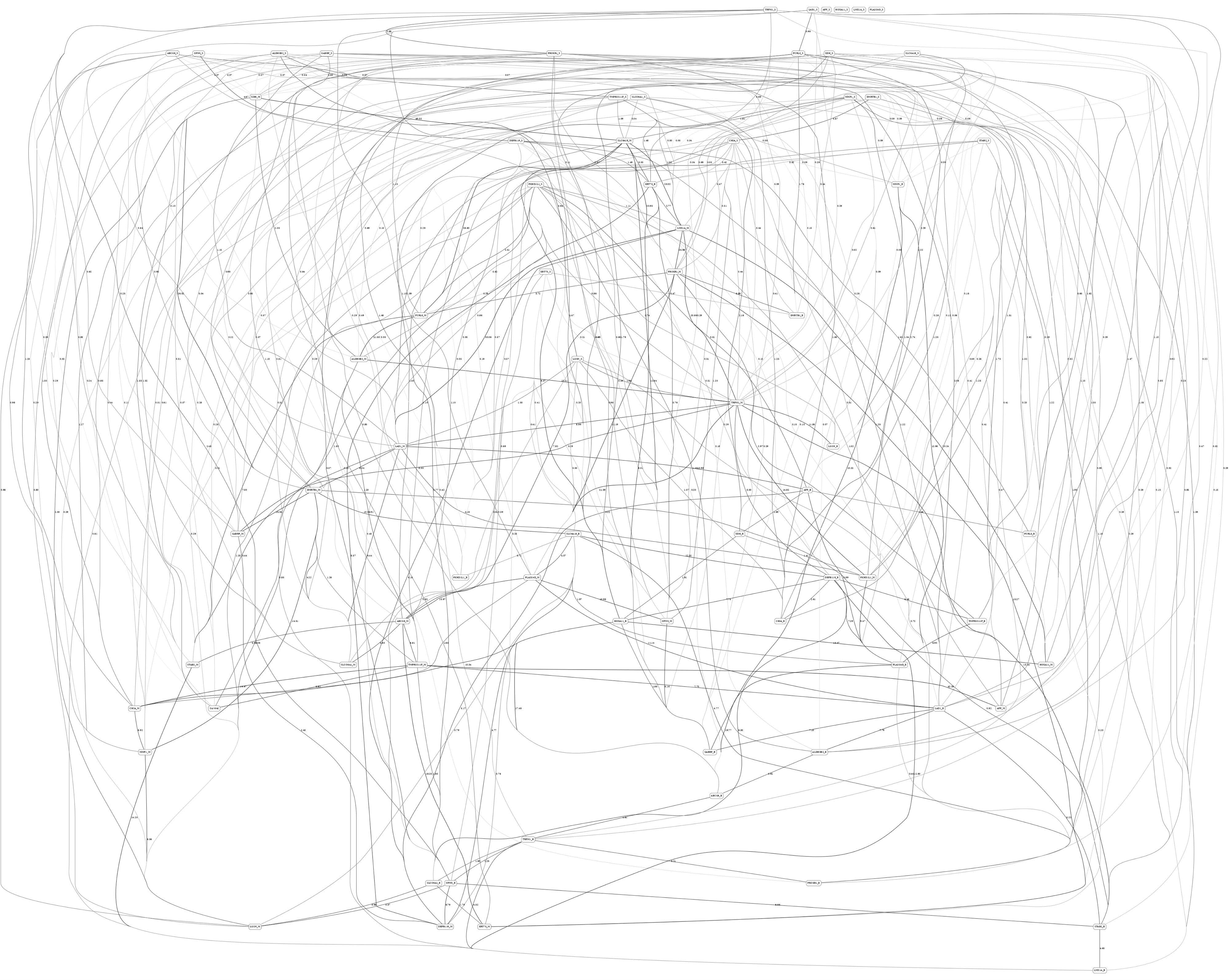
Full BN derived from the LGG/survival data. “Survival” node in the BN is self-explanatory. Other designations are as in Figure 3.

While the resulting full BNs, in PDF format, are zoom-able and searchable, and the DOT files can be exported into the specialized network-oriented software, the full BNs tend to be visually overwhelming for the number of variables/nodes > 100. Consequently, Figures 5 - 10 depict the immediate MNs of the clinical variables/nodes in the corresponding six BNs: LGG/tumor status (Figure 5), LGG/survival (Figure 6), HNSC/tumor status (Figure 7), HNSC/survival (Figure 8), STES/tumor status (Figure 9) and STES/survival (Figure 10).

**Figure 5.**
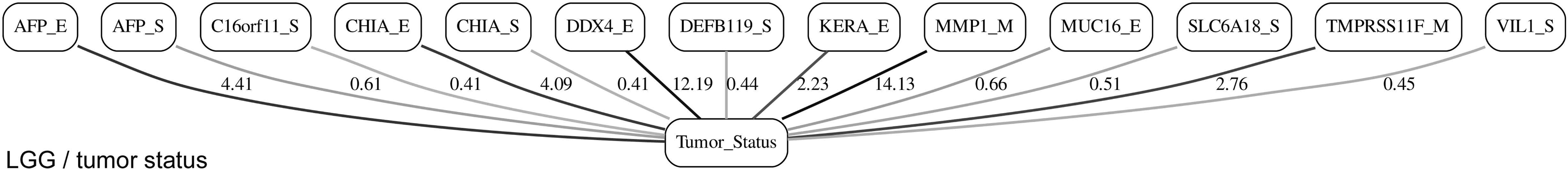
MN of the “Tumor_Status” node in the LGG/tumor status BN.

**Figure 6.**
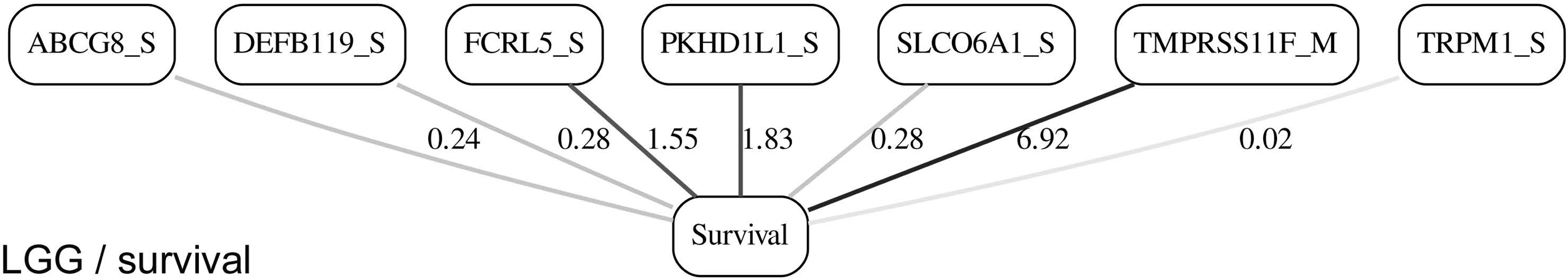
MN of the “Survival” node in the LGG/survival BN.

**Figure 7.**
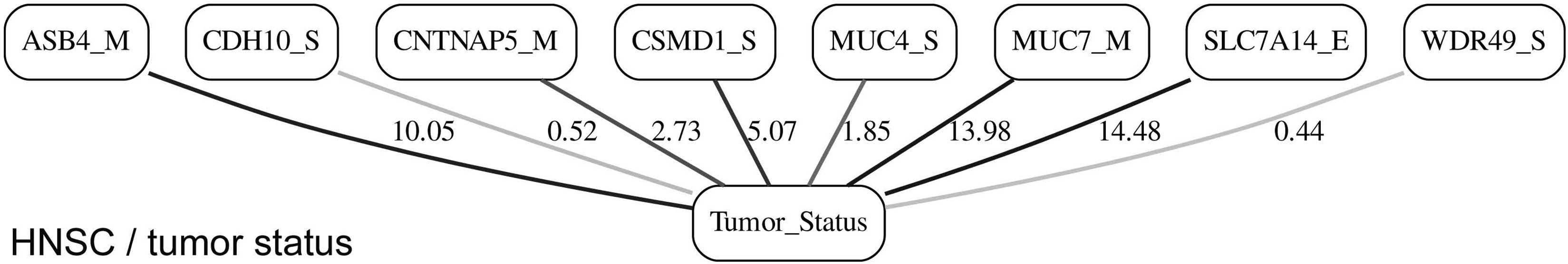
MN of the “Tumor_Status” node in the HNSC/tumor status BN.

**Figure 8.**
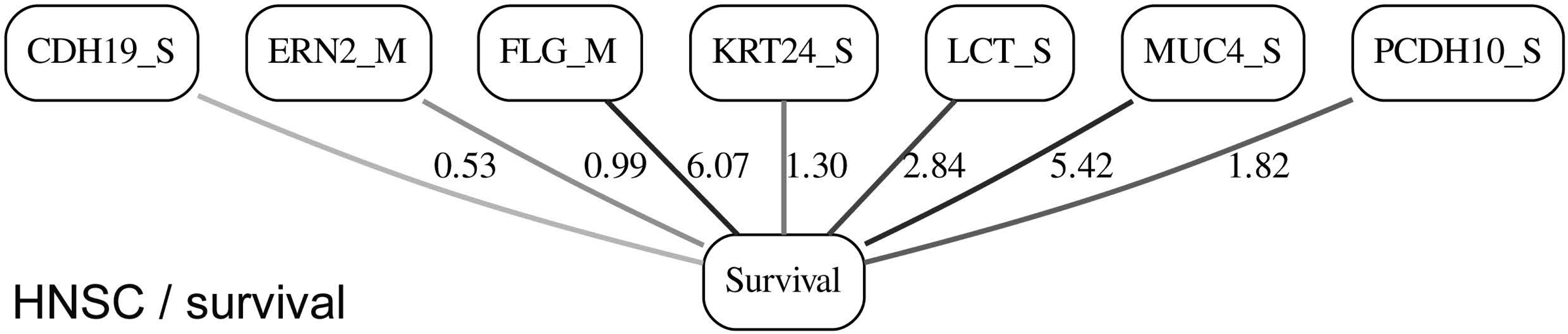
MN of the “Survival” node in the HNSC/survival BN.

**Figure 9.**
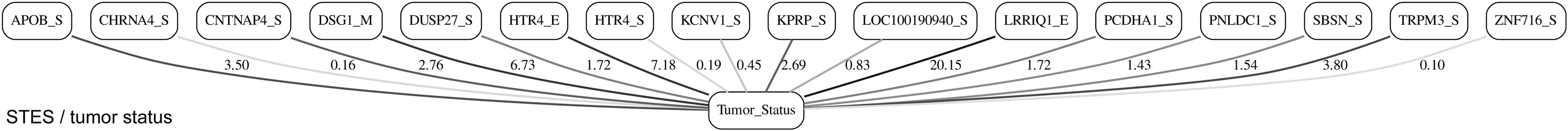
MN of the “Tumor_Status” node in the STES/tumor status BN.

**Figure 10.**
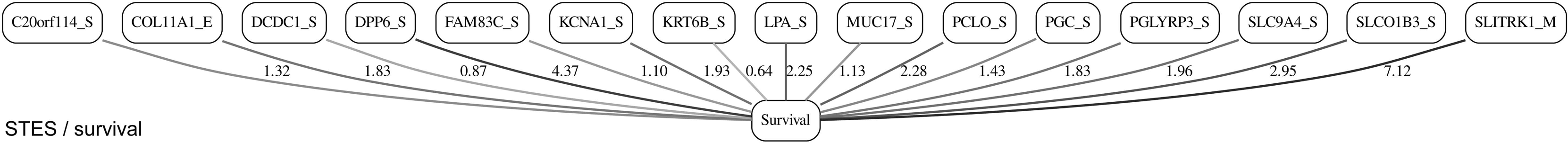
MN of the “Survival” node in the STES/survival BN.

It is noticeable in Figures 5 - 10 that all three molecular data components (S, E and M) are represented in the MNs. This testifies to the efficacy and proportionality of both the MMI measure (during the variable selection stage) and the maximum entropy - based discretization (during the BN construction stage). Also of note, for some genes, more than one component is present (HTR4 E and S for STES/tumor status, CHIA E and S for LGG/tumor status, AFP E and S for LGG/tumor status). Conversely, some genes are associated with both tumor status and survival (MUC4 for HNSC, TMPRSS11F, SLC6A18 and DEFB119 for LGG).

The performance of our BN reconstruction algorithm / software is discussed in general terms in [18 and references therein]; here, we will evaluate the statistical significance of the resulting MNs. While the edge strength estimates in Figures 5 – 10 are useful in the relative sense, they do not immediately translate into the statistical significance measurements (such as *p-*values). Therefore, we have augmented the edge strengths with the *p-*values obtained *via* two-sample Kolmogorov-Smirnov (KS) probability distribution equality test (for continuous E and M molecular component variables) and two-sided Fisher’s exact test (for discrete S molecular component variable). To illustrate the KS test application, Figure 11 shows CDFs, separately for two “tumor status” groups, for seven continuous variables present in the MN depicted in Figure 5 (LGG/tumor status), in order of decreasing edge strength (Figure 11A, MMP1_M; Figure 11B, DDX4_E; Figure 11C, AFP_E; Figure 11D, CHIA_E; Figure 11E, TMPRSS11F_M; Figure 11F, KERA_E; Figure 11G, MUC16_E). Only MMP1_M and DDX4_E appear to be statistically highly significant, with TMPRSS11F_M being arguably a borderline case.

**Figure 11.**
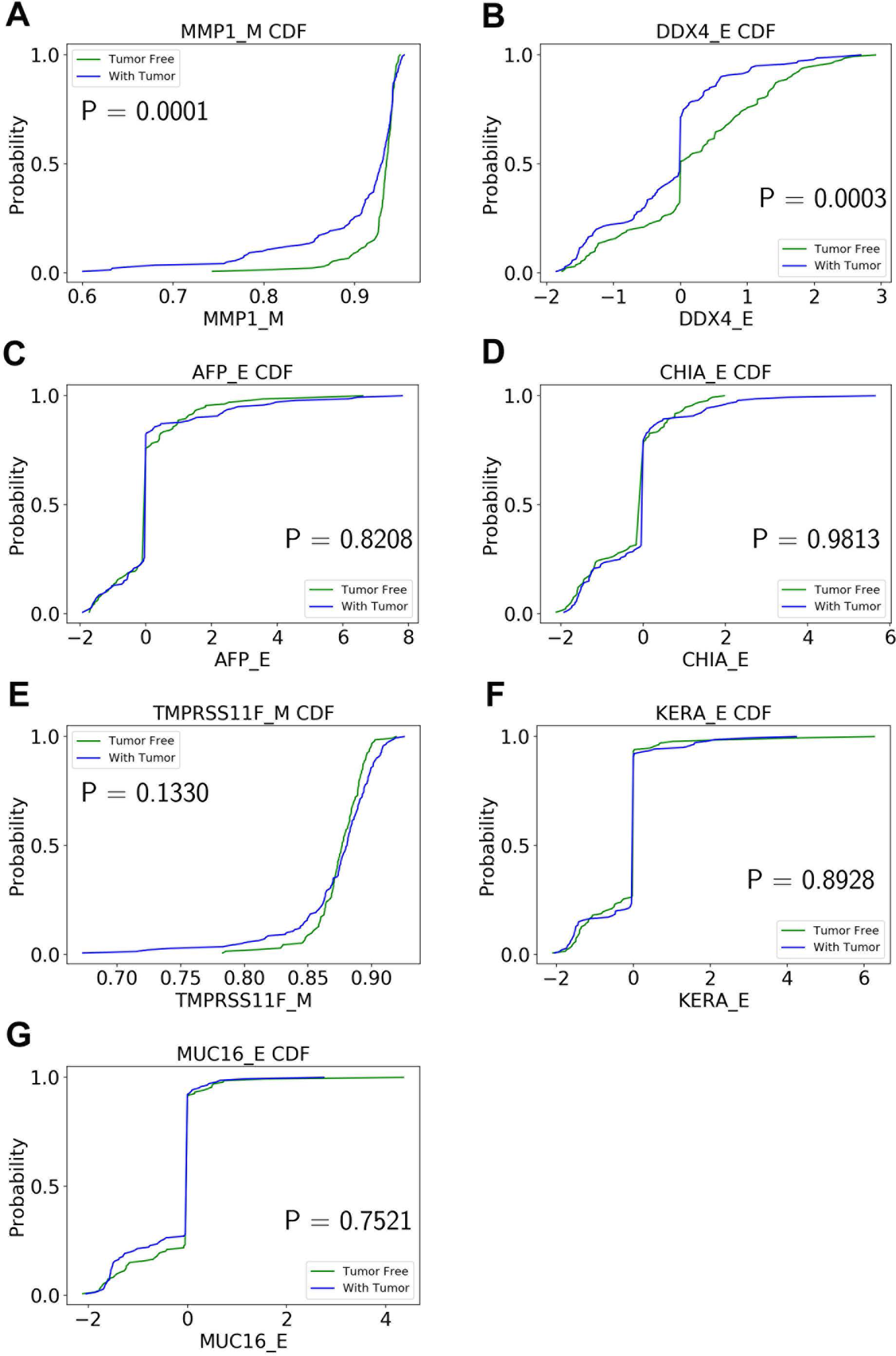
CDFs, shown separately for two “tumor status” groups, for seven continuous variables present in the MN depicted in Figure 5 (LGG/tumor status), in order of decreasing edge strength. (A) MMP1_M; (B) DDX4_E; (C) AFP_E; (D) CHIA_E; (E) TMPRSS11F_M; (F) KERA_E; (G) MUC16_E. *P-*values for the two-sample Kolmogorov-Smirnov test are shown in each chart.

Table 1 lists the *p-*values for all 55 potentially predictive molecular gene components present in six MNs depicted in Figures 5 - 10, in order of decreasing edge strength for each network / MN. 12 gene components were found to be statistically significant (marked with an asterisk in Table 1); however, we decide to exclude LCT_S (marked with ** in Table 1) from further scrutiny because of the very low mutation counts in both survival groups.

Subsequently, we performed manual literature / database search to ascertain if any of the remaining 11 genes were previously reported in the cancer context. The following resources were used: GeneCards [39] and DisGeNET [40] databases, PubMed, and Google Scholar. Six genes were found to be implicated in cancer etiology / progression / clinical outcomes with high degree of certainty: MMP1, DDX4, TRPM3, DPP6, KCNA1 and MUC17 [41–46]. Four genes (SLC7A14, LRRIQ, SLCO1B3, SLC9A4) were supported by weaker, circumstantial evidence [47–50]. One gene, TMPRSS11F, has not been discussed in the cancer context before, to the best of our knowledge (see also [51]). However, increased expression levels of a similar type II transmembrane serine protease, TMPRSS11D, were found to be a significant non-small cell lung cancer survival predictor [52]. Therefore, we suggest that TMPRSS11F should be further investigated as a strong predictive factor playing a role in LGG patients’ clinical characteristics --- survival, especially. Lower TMPRSS11F methylation values correspond to a poorer long-term (2-year) survival. One possible mechanism is *via* the proteolysis of extracellular matrix which, in turn, is linked to the metastatic processes [52].

In summary, our analysis framework confirmed six well-known cancer-related genes, supplied additional evidence to support four other suspected cancer-related genes, and identified one novel potentially strongly predictive factor, methylation of TMPRSS11F.

## Discussion

Systems biology approach to the complex genetic and epigenetic cancer data analysis is arguably superior to the simpler single-gene (or even single-data type) alternatives. However, it is intrinsically linked to the fundamental, interrelated, challenges --- scalability, “curse of dimensionality”, accounting for non-additive, higher-order interactions, and visualization of the results (i.e., translation of the massive network graphs into concrete biomedical insights). In this study we propose a flexible and generalizable approach to the BN-based systems biology analysis of the multi-modal cancer data, using the TCGA database as an example. It consists of the variable selection step (which is not computationally demanding) and the BN reconstruction step (which is substantially computationally demanding). Ideally, the investigators would simply feed the complete dataset (all variables) into the BN software, obtain the full graphical model (no matter how large and complex), and then “zoom in” on the MN of the variable(s) of interest, such as a clinical outcome or a cancer phenotype. However, this is impractical for most real datasets and available hardware configurations.

Consequently, we propose starting with the variable selection step to select a (relatively) small subset of genes that are associated with the variable(s) of interest (tumor status and 2-year survival in the present study). The principal novelty of our approach lies in using the MMI measure for the variable/gene selection, in which all possible types of molecular information (discrete and continuous, genetic and epigenetic) are considered simultaneously. The other innovative aspect of our approach lies in the adjustability of the “hand-off” point between the variable selection and BN modeling steps. This hand-off point can depend on the investigators’ computational resources, the shape of the variable selection curves, or the predefined statistical cutoff points. For example, ∼20K genes can be reduced to 100-200 genes for the subsequent BNs construction, in which case the complete analysis takes less than an hour on a mid-level PC. When feeding the complete datasets (10,000-15,000 genes, in case of TCGA and similar genomic resources) into our BN software [18], without the preliminary variable selection step, it takes about three days to build a full BN on a dedicated multi-core workstation. Therefore, the investigators can choose the appropriate balance depending on whether they are interested in a quick, exploratory analysis or a finalized, exhaustive one.

Our computational pipeline is inherently generalizable, as it can be directly applied to any large multimodal genetic/epigenetic dataset with minimal preprocessing. The only two changeable parameters are the variable selection / BN modeling hand-off point, and the BN discretization mechanism. The latter is currently set as the 3-bin maximum entropy-based discretization coupled with the multinomial local probability model [18]. This is not the most elegant, or universally applicable, solution. In future, we plan to develop a novel BN model scoring function derived from a mixed distance measure (such as the MMI), or a similar metric that expresses divergence between the current network model and the data *via* mixed-type distances. The resulting two-stage analytical strategy will thus fully automatically deal with the mixed variables, in both of its stages. This has not been done before, so we plan to implement and test the MMI-based BN algorithm alongside the more established mixed-type BN solutions (hybrid local probability models, adaptive discretization), and use both real and simulated data to investigate which method is preferable.

Another limitation of the present study has to do with its primary focus on the clinical outcomes / phenotypes; at this time, we decided to largely concentrate on the MNs of the clinical variables/nodes. In future, we intend to analyze the resulting full BNs more “holistically”, paying attention to the general network topological properties, gene clusters, hub and bottleneck genes, etc.

Application of our pipeline to TCGA data resulted in the identification of a number of candidate genes for the different clinical cancer characteristics, *via* varied molecular components. It is well known that epigenetic processes / DNA methylation play an important role in many cancers’ diagnosis, progression, and outcome; our results support that notion, as many of the most statistically significant predictors generated in the present study were in fact the methylation molecular components (Table 1). Notably, the one novel candidate gene pinpointed in this study, TMPRSS11F, likely would not have been identified *via* any other (non-epigenetic) modality. Our results, therefore, underscore the essentiality of the simultaneous analysis of different molecular modalities, including the epigenetic ones, for the precision / personalized medicine to be effective in cancer treatment.

**Table 1.**
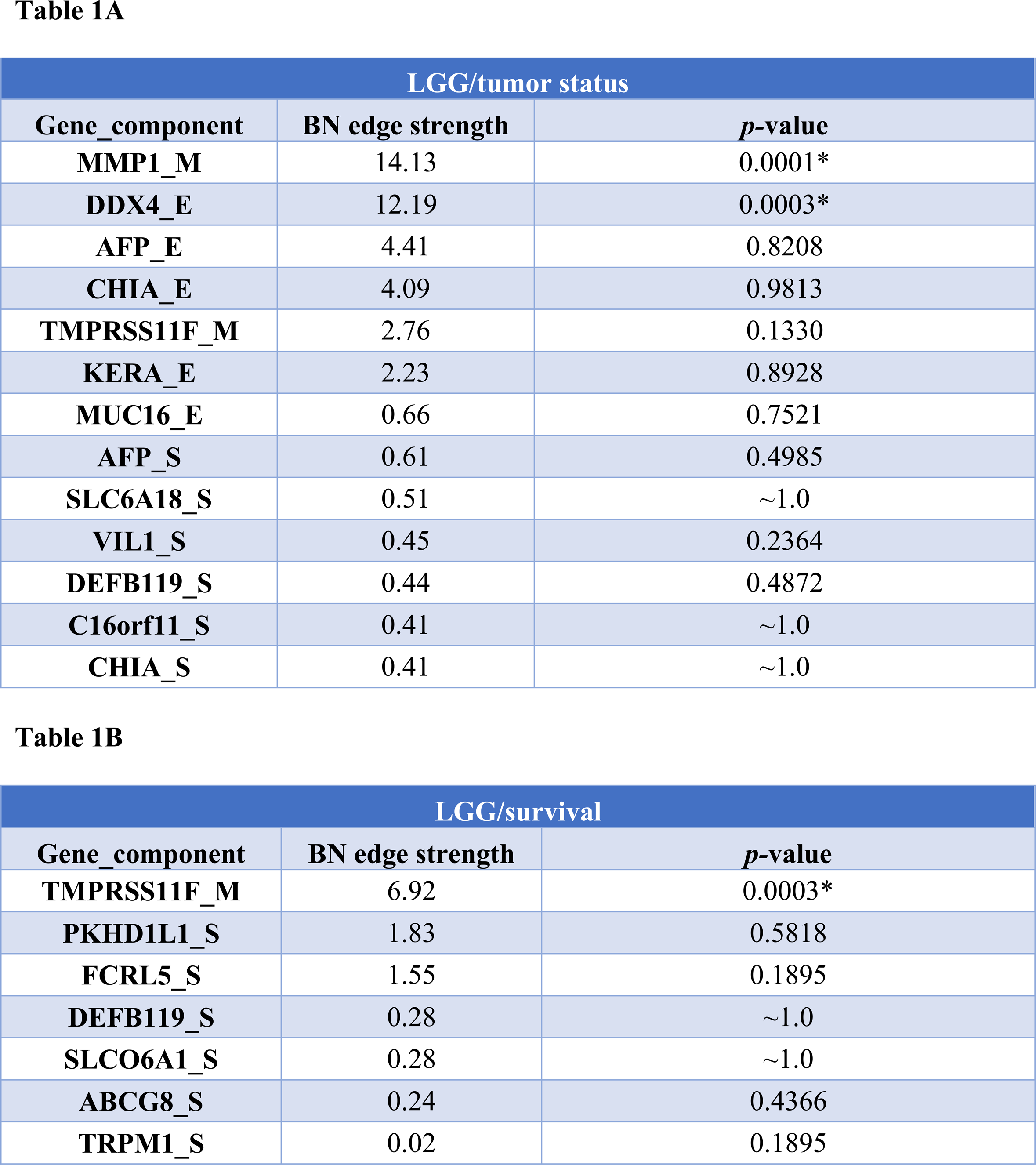
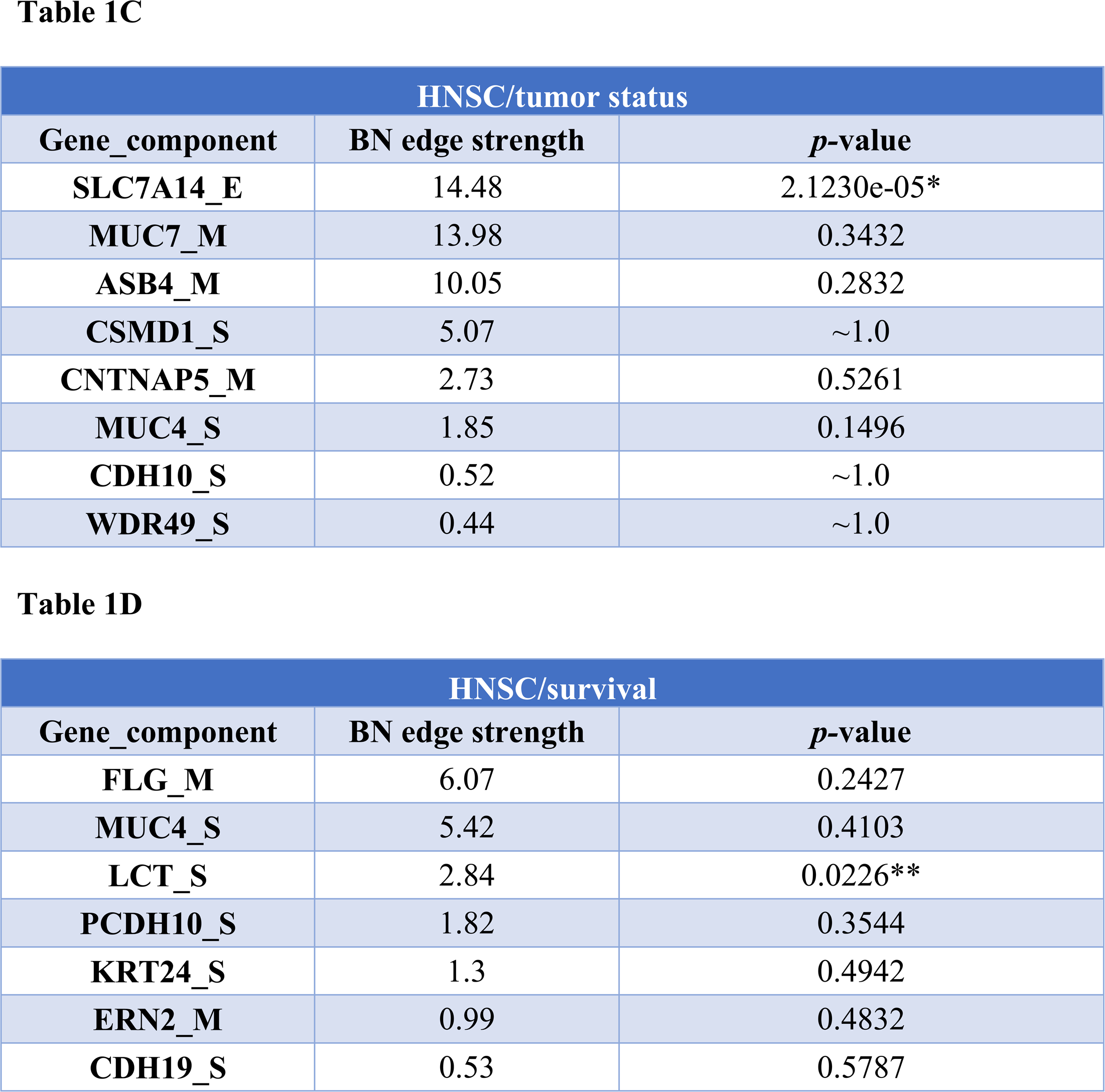
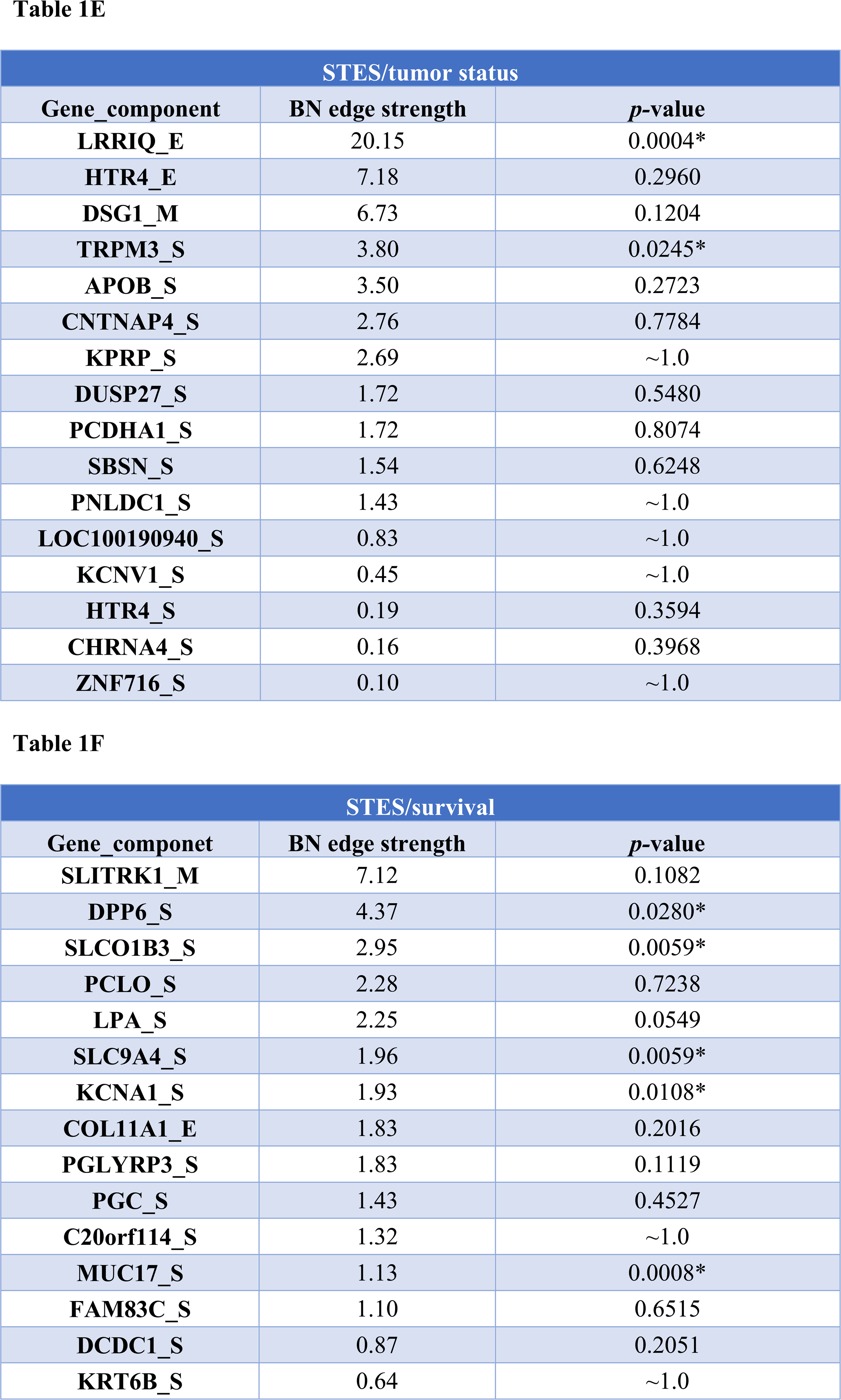
*P-*values for 55 potentially predictive molecular gene components present in six MNs depicted in Figures 5 - 10, subdivided by six datasets, in order of decreasing edge strength for each dataset/MN. 12 gene components were found to be statistically significant (marked with an asterisk); LCT_S (marked with **) was excluded from further analysis because of the very low mutation counts (zero mutations in > 2-year survival group, three mutations in < 2-year survival group).

## Supporting information

Suppl Data Sheet 1

Suppl Data Sheet 2

Suppl Data Sheet 3

Suppl Data Sheet 4

Suppl Data Sheet 5

Suppl Data Sheet 6

## Supplementary Material

**Supplementary Data Sheets 1-6**

Full Bayesian networks, in PDF format, for the six datasets in this study --- (1) LGG/tumor status, (2) LGG/survival, (3) HNSC/tumor status, (4) HNSC/survival, (5) STES/tumor status, (6) STES/survival. Designations are as in main text Figure 3.

**Supplementary Tables 1-6**

Data files, in Excel format, listing the genes analyzed in this study, together with the corresponding MMI values, in order of decreasing MMI values. (1) LGG/tumor status, (2) LGG/survival, (3) HNSC/tumor status, (4) HNSC/survival, (5) STES/tumor status, (6) STES/survival. Genes with the MMI values > 99.5% MMI CDF (i.e., genes selected for further BN analyses) are shown in bold.

**Supplementary Tables 7-12**

Full Bayesian networks, in Word / DOT format, for the six datasets in this study --- (1) LGG/tumor status, (2) LGG/survival, (3) HNSC/tumor status, (4) HNSC/survival, (5) STES/tumor status, (6) STES/survival.

## Acknowledgements

The authors are grateful to Arthur D. Riggs, Russell Rockne, Dustin Schones, Wendong Huang and Weihao Gao for many stimulating discussions and useful suggestions.

## Author Contributions Statement

X.W. conceptualized the study, carried out the analyses, and wrote the manuscript. S.B. conceptualized the study, contributed to carrying out the analyses, and contributed to writing the manuscript. G.G. contributed to carrying out the analyses, and contributed to writing the manuscript. S.D. contributed to carrying out the analyses, and contributed to writing the manuscript. A.S.R. conceptualized the study, contributed to carrying out the analyses, and wrote the manuscript.

## Conflict of Interest Statement

The authors declare that they have no competing interests.

## Funding Disclosure Statement

This work was supported by the Susumu Ohno Chair in Theoretical and Computational Biology (held by A.S.R.), a Susumo Ohno Distinguished Investigator fellowship (to G.G.), and City of Hope funds (to X.W., S.B., G.G. and A.S.R.). The funders had no role in study design, data collection and analysis, decision to publish, or preparation of the manuscript.

## Data Availability Statement

Principal datasets analyzed in this study are included in the manuscript and the supplementary material files. All intermediate/auxiliary datasets will be made available by the authors, without undue reservation, to any qualified researcher.

