## Supplementary figures and images for "New analysis framework incorporating mixed mutual information and scalable Bayesian networks for multimodal high dimensional genomic and epigenomic cancer data"

### Suppl Data Sheet 1

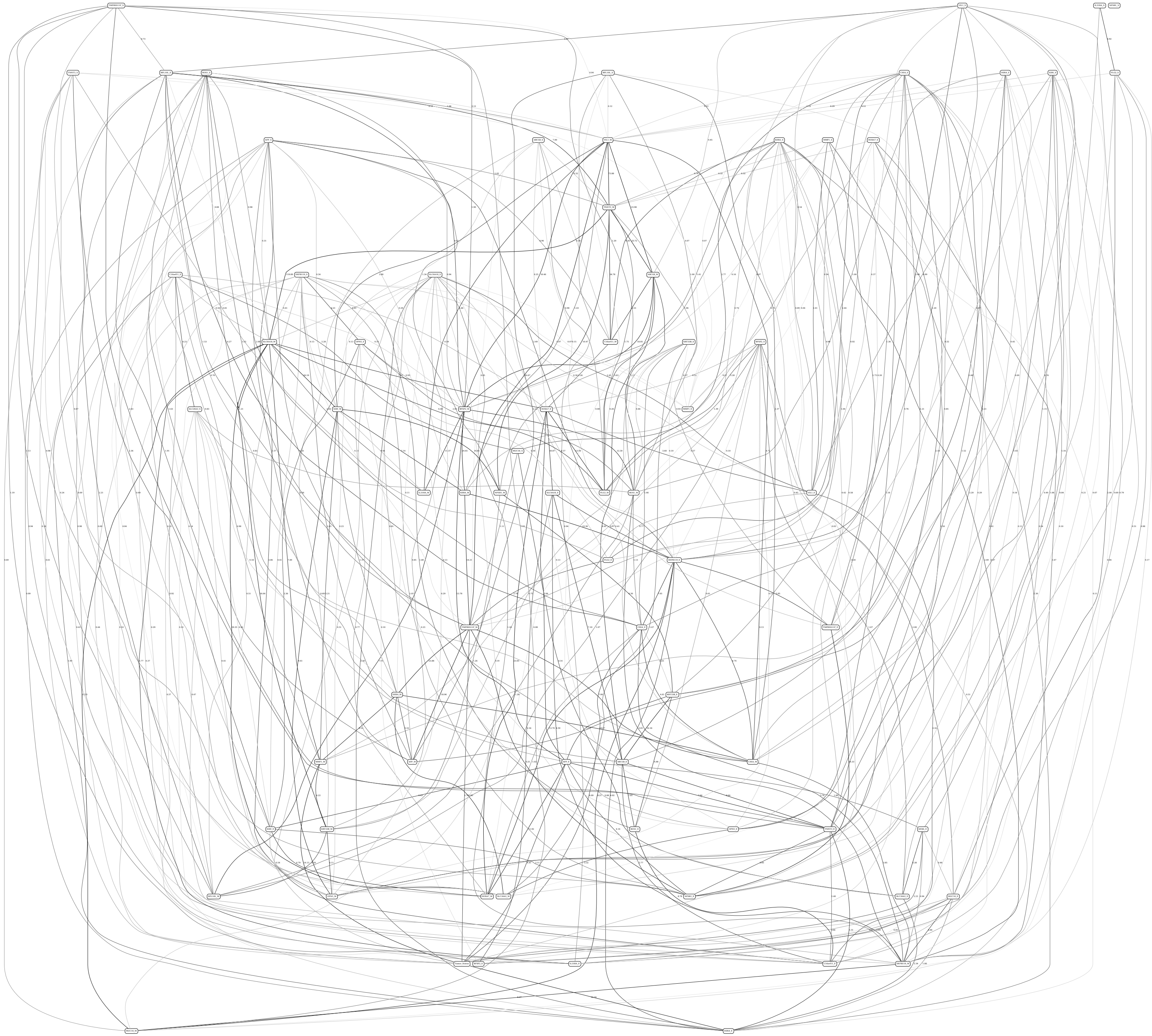

### Suppl Data Sheet 2

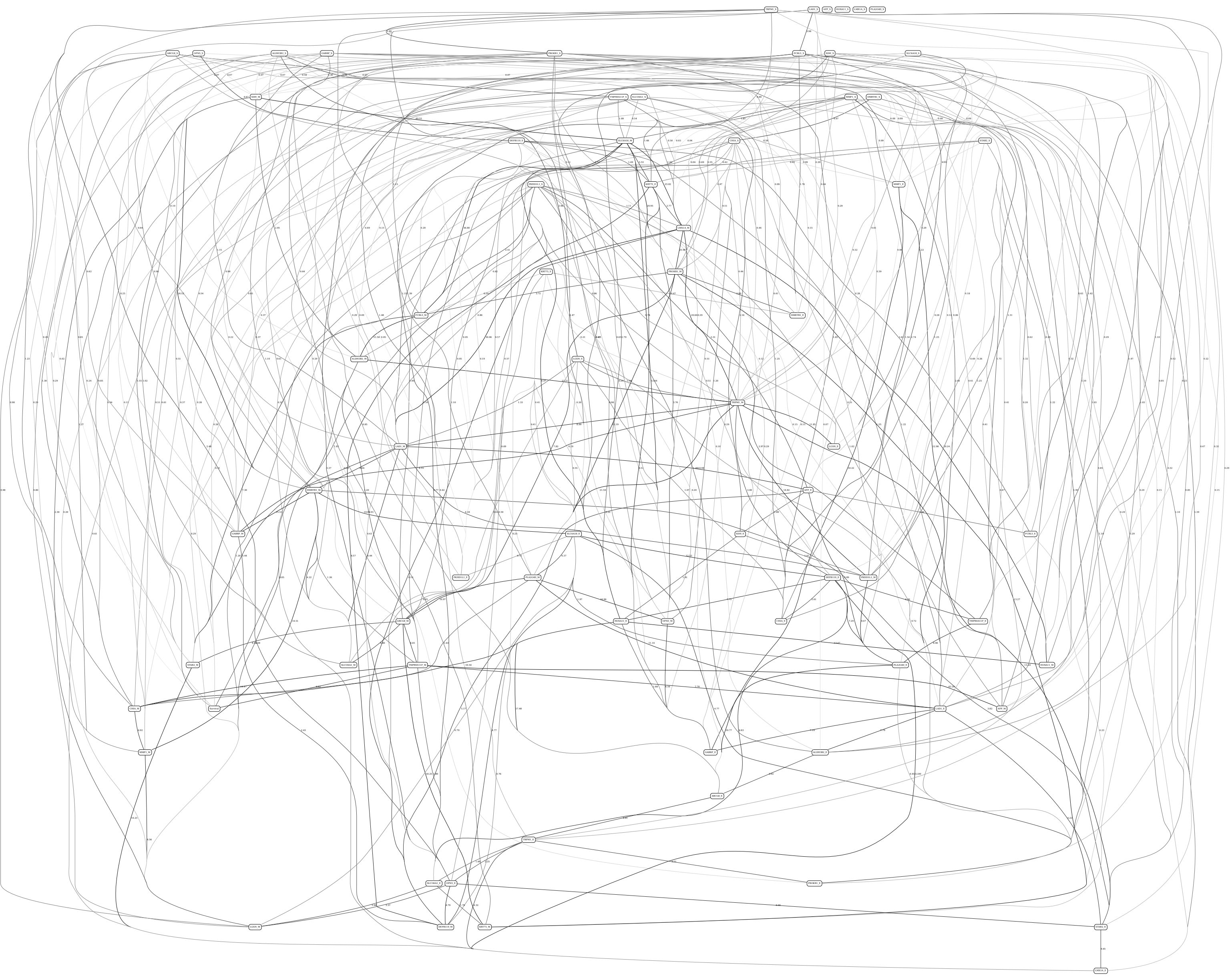

### Suppl Data Sheet 3

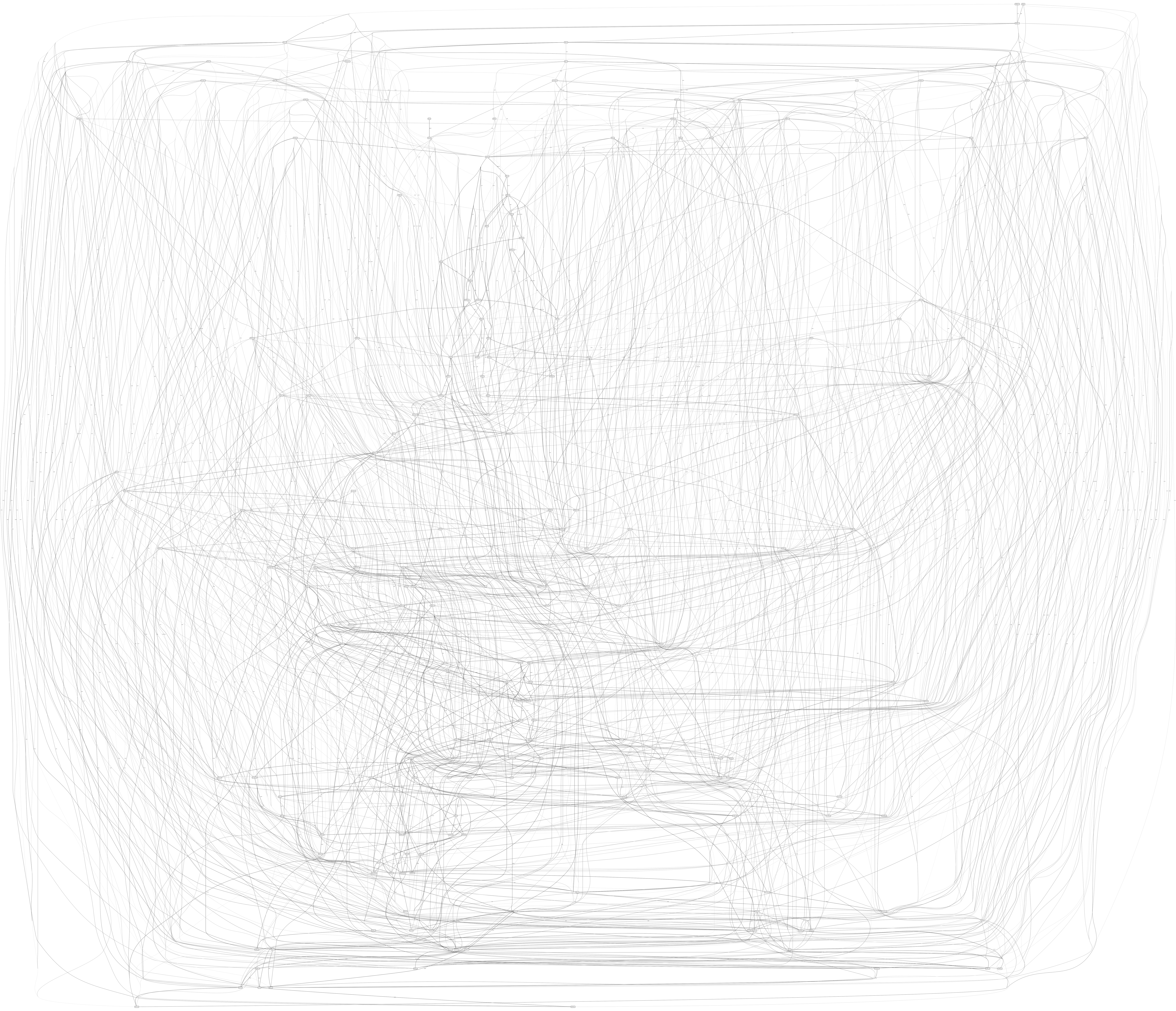

### Suppl Data Sheet 4

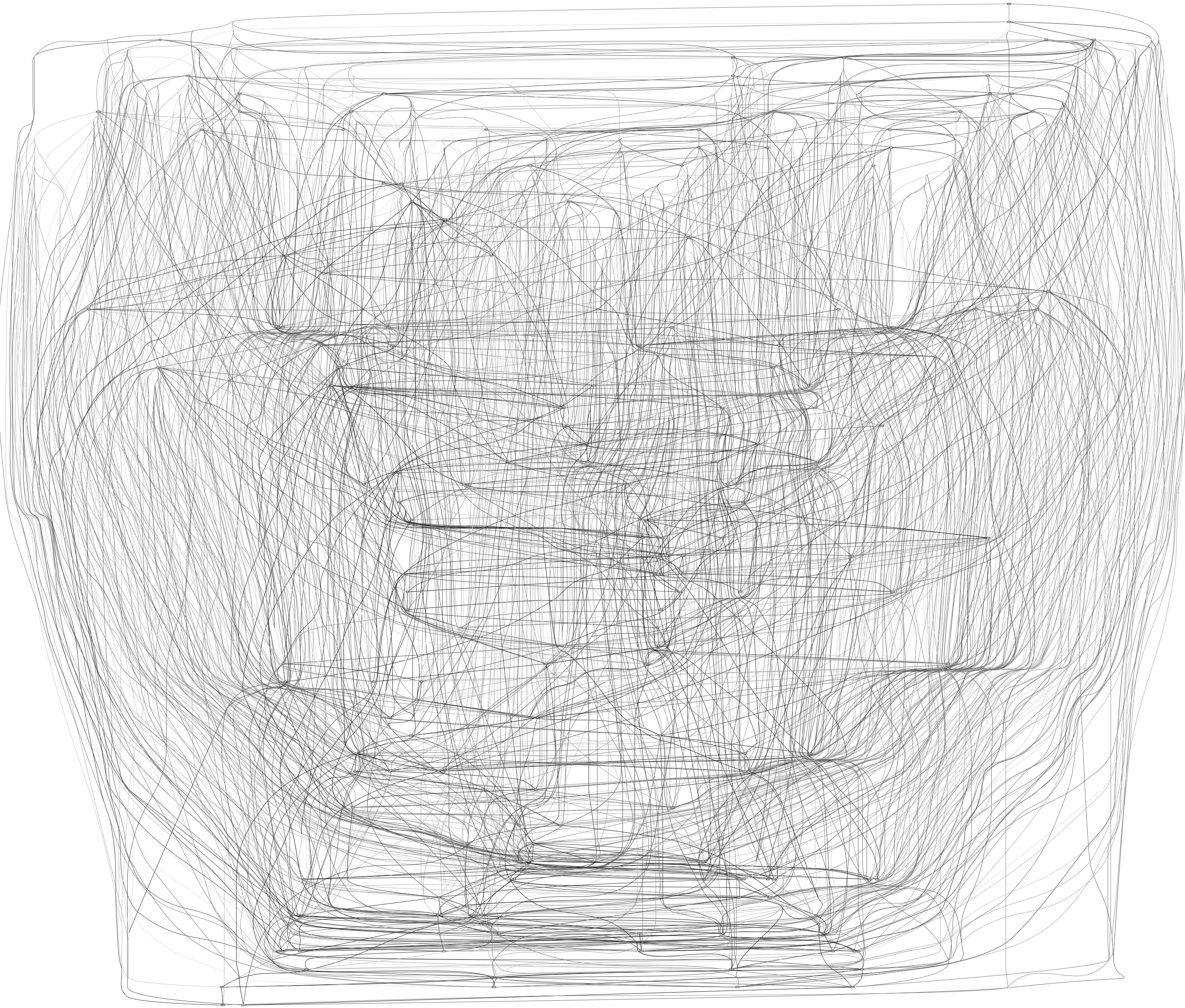

### Suppl Data Sheet 5

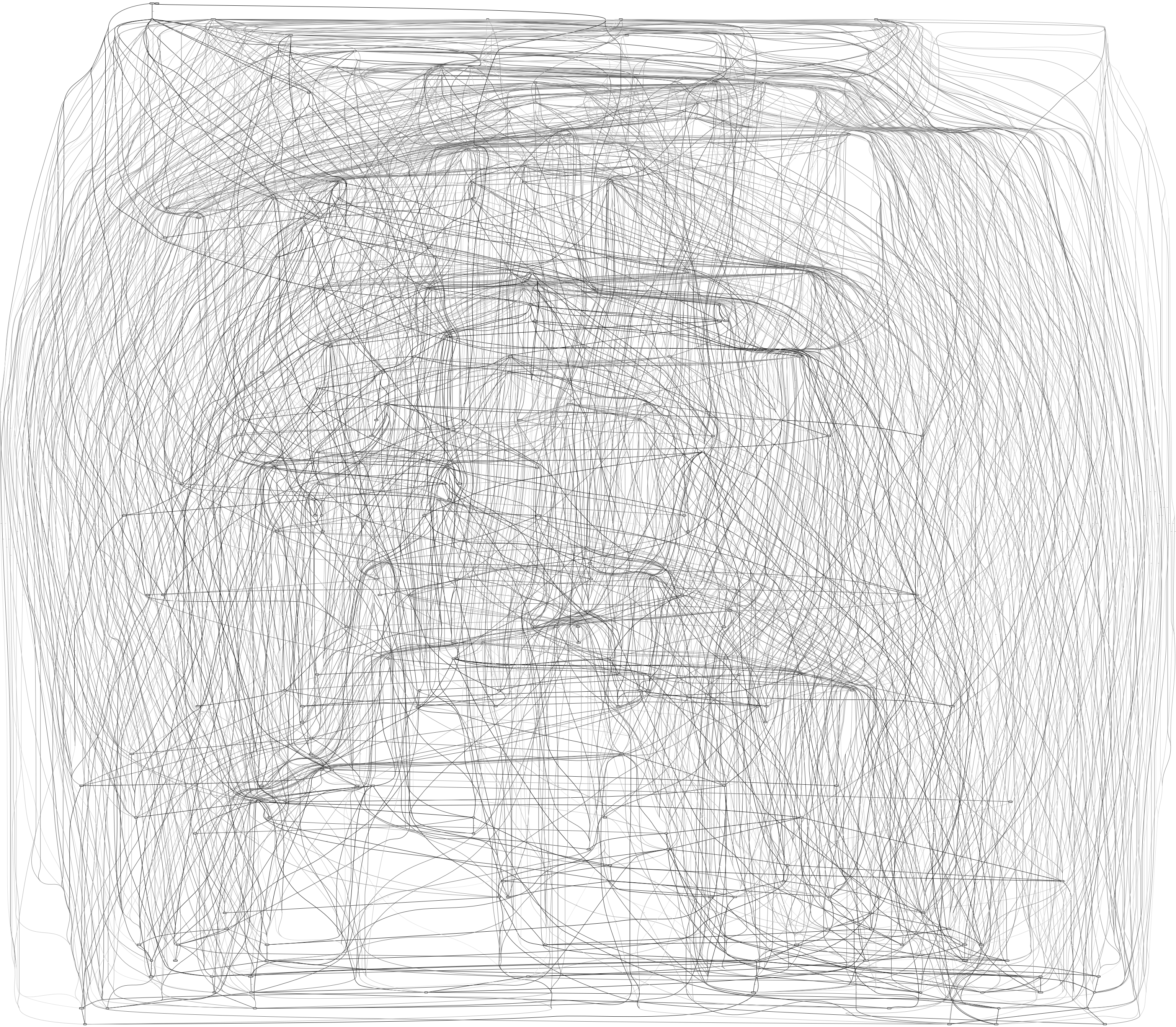

### Suppl Data Sheet 6

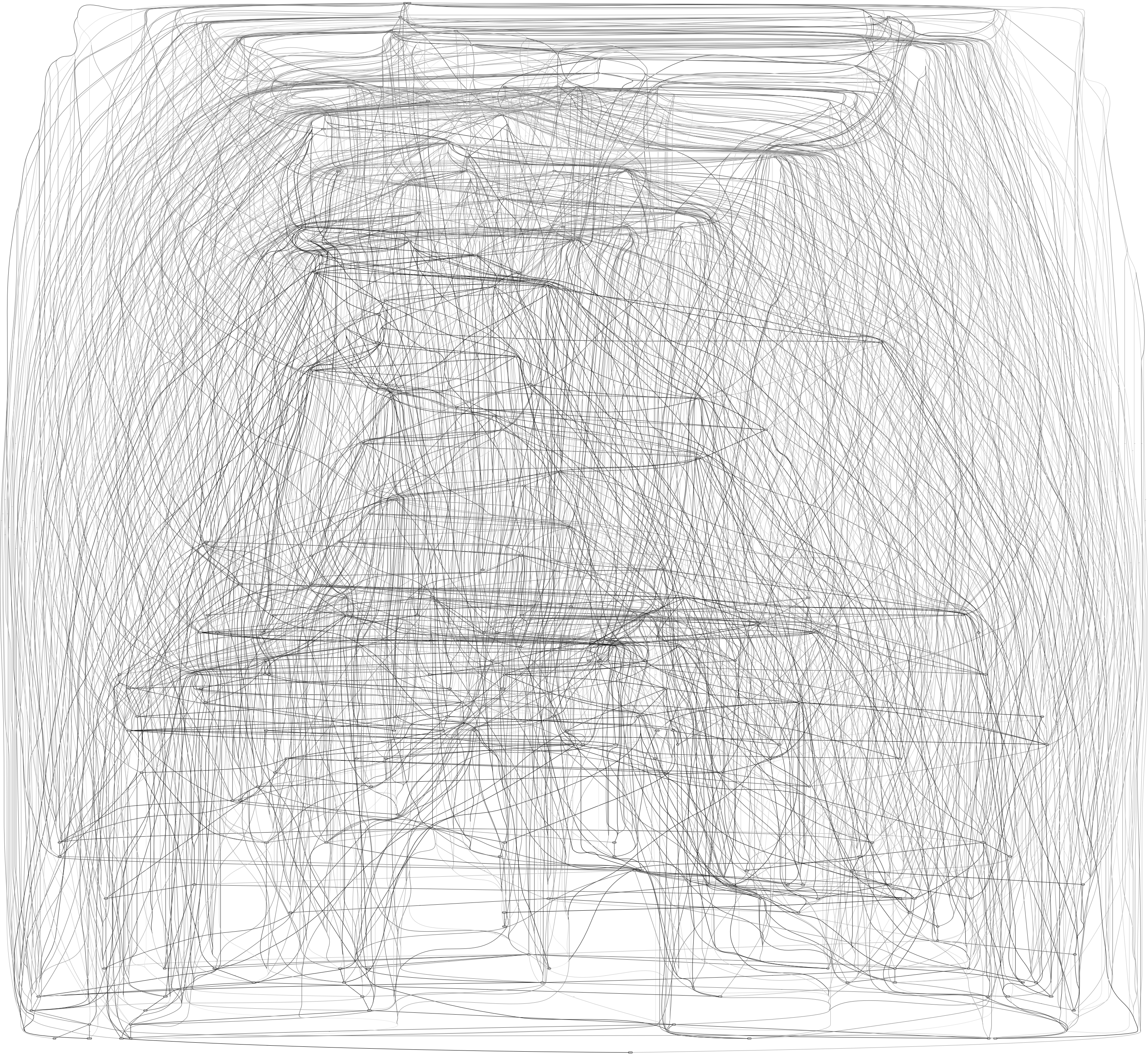
