## Supplementary material for "New analysis framework incorporating mixed mutual information and scalable Bayesian networks for multimodal high dimensional genomic and epigenomic cancer data": Suppl Table 7

```
digraph G{
 ratio=fill;
 node [shape=box, style=rounded];
 edge [arrowhead=none];
"ABCG8_S";
"ABCG8_E";
"ABCG8_M";
"AFP_S";
"AFP_E";
"AFP_M";
"BFSP2_S";
"BFSP2_E";
"BFSP2_M";
"C16orf11_S";
"C16orf11_E";
"C16orf11_M";
"CHIA_S";
"CHIA_E";
"CHIA_M";
"DDX4_S";
"DDX4_E";
"DDX4_M";
"DEFB119_S";
"DEFB119_E";
"DEFB119_M";
"FLG2_S";
"FLG2_E";
"FLG2_M";
"HOXA7_S";
"HOXA7_E";
"HOXA7_M";
"IL31RA_S";
"IL31RA_E";
"IL31RA_M";
"KERA_S";
"KERA_E";
"KERA_M";
"KRT33B_S";
"KRT33B_E";
"KRT33B_M";
"MMP1_S";
"MMP1_E";
"MMP1_M";
"MUC16_S";
"MUC16_E";
"MUC16_M";
"NPSR1_S";
"NPSR1_E";
"NPSR1_M";
"OPN5_S";
"OPN5_E";
"OPN5_M";
"ROS1_S";
"ROS1_E";
"ROS1_M";
"RPL10L_S";
"RPL10L_E";
"RPL10L_M";
"SLC26A3_S";
"SLC26A3_E";
"SLC26A3_M";
"SLC6A18_S";
"SLC6A18_E";
"SLC6A18_M";
"TMPRSS11F_S";
"TMPRSS11F_E";
"TMPRSS11F_M";
"TNNT3_S";
"TNNT3_E";
"TNNT3_M";
"VIL1_S";
"VIL1_E";
"VIL1_M";
"XDH_S";
"XDH_E";
"XDH_M";
"Tumor_Status";
"KERA_S" -> "SLC26A3_M" [color="0 0 0.9", label="0.03",style=bold];
"AFP_S" -> "RPL10L_M" [color="0 0 0.897398843931", label="0.04",style=bold];
"HOXA7_S" -> "ROS1_M" [color="0 0 0.894797687861", label="0.04",style=bold];
"DDX4_S" -> "MUC16_M" [color="0 0 0.892196531792", label="0.04",style=bold];
"TMPRSS11F_S" -> "VIL1_M" [color="0 0 0.889595375723", label="0.04",style=bold];
"MUC16_E" -> "C16orf11_E" [color="0 0 0.886994219653", label="0.05",style=bold];
"ABCG8_S" -> "TMPRSS11F_M" [color="0 0 0.884393063584", label="0.05",style=bold];
"MMP1_S" -> "MMP1_E" [color="0 0 0.881791907514", label="0.07",style=bold];
"KRT33B_S" -> "MMP1_E" [color="0 0 0.879190751445", label="0.07",style=bold];
"TNNT3_S" -> "MMP1_E" [color="0 0 0.876589595376", label="0.07",style=bold];
"SLC6A18_S" -> "MMP1_E" [color="0 0 0.873988439306", label="0.07",style=bold];
"DEFB119_S" -> "MMP1_E" [color="0 0 0.871387283237", label="0.07",style=bold];
"BFSP2_S" -> "MMP1_E" [color="0 0 0.868786127168", label="0.07",style=bold];
"CHIA_S" -> "MMP1_E" [color="0 0 0.866184971098", label="0.07",style=bold];
"VIL1_S" -> "IL31RA_E" [color="0 0 0.863583815029", label="0.07",style=bold];
"RPL10L_S" -> "AFP_E" [color="0 0 0.86098265896", label="0.08",style=bold];
"KRT33B_S" -> "HOXA7_M" [color="0 0 0.85838150289", label="0.08",style=bold];
"RPL10L_E" -> "SLC6A18_M" [color="0 0 0.855780346821", label="0.08",style=bold];
"KRT33B_S" -> "DEFB119_E" [color="0 0 0.853179190751", label="0.10",style=bold];
"BFSP2_S" -> "DEFB119_E" [color="0 0 0.850578034682", label="0.10",style=bold];
"SLC6A18_S" -> "BFSP2_E" [color="0 0 0.847976878613", label="0.10",style=bold];
"DEFB119_S" -> "BFSP2_E" [color="0 0 0.845375722543", label="0.10",style=bold];
"C16orf11_S" -> "BFSP2_E" [color="0 0 0.842774566474", label="0.10",style=bold];
"DDX4_S" -> "BFSP2_E" [color="0 0 0.840173410405", label="0.10",style=bold];
"SLC6A18_S" -> "DDX4_M" [color="0 0 0.837572254335", label="0.11",style=bold];
"MMP1_S" -> "DDX4_E" [color="0 0 0.834971098266", label="0.11",style=bold];
"RPL10L_S" -> "DEFB119_M" [color="0 0 0.832369942197", label="0.11",style=bold];
"DEFB119_S" -> "TMPRSS11F_M" [color="0 0 0.829768786127", label="0.11",style=bold];
"ROS1_S" -> "VIL1_M" [color="0 0 0.827167630058", label="0.12",style=bold];
"FLG2_S" -> "VIL1_M" [color="0 0 0.824566473988", label="0.12",style=bold];
"TNNT3_S" -> "VIL1_M" [color="0 0 0.821965317919", label="0.12",style=bold];
"RPL10L_S" -> "VIL1_M" [color="0 0 0.81936416185", label="0.12",style=bold];
"CHIA_S" -> "VIL1_M" [color="0 0 0.81676300578", label="0.12",style=bold];
"DDX4_S" -> "VIL1_M" [color="0 0 0.814161849711", label="0.12",style=bold];
"BFSP2_S" -> "OPN5_M" [color="0 0 0.811560693642", label="0.13",style=bold];
"DEFB119_S" -> "XDH_M" [color="0 0 0.808959537572", label="0.13",style=bold];
"SLC6A18_S" -> "XDH_M" [color="0 0 0.806358381503", label="0.13",style=bold];
"SLC6A18_S" -> "OPN5_E" [color="0 0 0.803757225434", label="0.13",style=bold];
"BFSP2_S" -> "OPN5_E" [color="0 0 0.801156069364", label="0.13",style=bold];
"DEFB119_S" -> "OPN5_E" [color="0 0 0.798554913295", label="0.13",style=bold];
"AFP_S" -> "SLC26A3_M" [color="0 0 0.795953757225", label="0.14",style=bold];
"ABCG8_S" -> "DEFB119_E" [color="0 0 0.793352601156", label="0.15",style=bold];
"SLC26A3_S" -> "XDH_E" [color="0 0 0.790751445087", label="0.16",style=bold];
"C16orf11_S" -> "BFSP2_M" [color="0 0 0.788150289017", label="0.16",style=bold];
"MMP1_S" -> "BFSP2_M" [color="0 0 0.785549132948", label="0.16",style=bold];
"FLG2_S" -> "MUC16_M" [color="0 0 0.782947976879", label="0.17",style=bold];
"SLC26A3_S" -> "DDX4_M" [color="0 0 0.780346820809", label="0.18",style=bold];
"DDX4_S" -> "NPSR1_E" [color="0 0 0.77774566474", label="0.18",style=bold];
"SLC6A18_S" -> "MUC16_M" [color="0 0 0.775144508671", label="0.19",style=bold];
"KERA_S" -> "BFSP2_E" [color="0 0 0.772543352601", label="0.20",style=bold];
"HOXA7_S" -> "TMPRSS11F_M" [color="0 0 0.769942196532", label="0.20",style=bold];
"XDH_S" -> "MUC16_E" [color="0 0 0.767341040462", label="0.21",style=bold];
"OPN5_S" -> "SLC26A3_E" [color="0 0 0.764739884393", label="0.23",style=bold];
"ABCG8_S" -> "OPN5_M" [color="0 0 0.762138728324", label="0.23",style=bold];
"TNNT3_S" -> "SLC26A3_M" [color="0 0 0.759537572254", label="0.24",style=bold];
"CHIA_E" -> "KRT33B_E" [color="0 0 0.756936416185", label="0.25",style=bold];
"ABCG8_S" -> "HOXA7_E" [color="0 0 0.754335260116", label="0.25",style=bold];
"KRT33B_S" -> "FLG2_M" [color="0 0 0.751734104046", label="0.26",style=bold];
"CHIA_S" -> "SLC26A3_M" [color="0 0 0.749132947977", label="0.29",style=bold];
"SLC6A18_S" -> "OPN5_M" [color="0 0 0.746531791908", label="0.29",style=bold];
"C16orf11_S" -> "C16orf11_E" [color="0 0 0.743930635838", label="0.29",style=bold];
"AFP_S" -> "KERA_M" [color="0 0 0.741329479769", label="0.29",style=bold];
"XDH_S" -> "VIL1_M" [color="0 0 0.738728323699", label="0.29",style=bold];
"DEFB119_S" -> "VIL1_E" [color="0 0 0.73612716763", label="0.30",style=bold];
"AFP_S" -> "SLC6A18_M" [color="0 0 0.733526011561", label="0.31",style=bold];
"DDX4_S" -> "TMPRSS11F_E" [color="0 0 0.730924855491", label="0.32",style=bold];
"ABCG8_S" -> "TNNT3_M" [color="0 0 0.728323699422", label="0.32",style=bold];
"HOXA7_S" -> "TNNT3_M" [color="0 0 0.725722543353", label="0.32",style=bold];
"MMP1_S" -> "TNNT3_M" [color="0 0 0.723121387283", label="0.32",style=bold];
"AFP_S" -> "XDH_M" [color="0 0 0.720520231214", label="0.34",style=bold];
"RPL10L_E" -> "DEFB119_M" [color="0 0 0.717919075145", label="0.36",style=bold];
"OPN5_S" -> "FLG2_E" [color="0 0 0.715317919075", label="0.36",style=bold];
"DEFB119_S" -> "DDX4_E" [color="0 0 0.712716763006", label="0.37",style=bold];
"SLC26A3_S" -> "IL31RA_E" [color="0 0 0.710115606936", label="0.37",style=bold];
"VIL1_S" -> "FLG2_E" [color="0 0 0.707514450867", label="0.37",style=bold];
"SLC6A18_S" -> "TMPRSS11F_E" [color="0 0 0.704913294798", label="0.37",style=bold];
"ABCG8_S" -> "C16orf11_M" [color="0 0 0.702312138728", label="0.38",style=bold];
"C16orf11_S" -> "Tumor_Status" [color="0 0 0.699710982659", label="0.41",style=bold];
"DEFB119_S" -> "XDH_E" [color="0 0 0.69710982659", label="0.41",style=bold];
"CHIA_S" -> "Tumor_Status" [color="0 0 0.69450867052", label="0.41",style=bold];
"SLC6A18_S" -> "CHIA_M" [color="0 0 0.691907514451", label="0.41",style=bold];
"SLC6A18_S" -> "SLC6A18_M" [color="0 0 0.689306358382", label="0.41",style=bold];
"DEFB119_S" -> "SLC6A18_M" [color="0 0 0.686705202312", label="0.41",style=bold];
"SLC26A3_S" -> "OPN5_M" [color="0 0 0.684104046243", label="0.41",style=bold];
"KERA_S" -> "OPN5_E" [color="0 0 0.681502890173", label="0.42",style=bold];
"DEFB119_S" -> "Tumor_Status" [color="0 0 0.678901734104", label="0.44",style=bold];
"XDH_M" -> "IL31RA_M" [color="0 0 0.676300578035", label="0.44",style=bold];
"VIL1_S" -> "Tumor_Status" [color="0 0 0.673699421965", label="0.45",style=bold];
"VIL1_S" -> "KRT33B_E" [color="0 0 0.671098265896", label="0.45",style=bold];
"VIL1_S" -> "ABCG8_M" [color="0 0 0.668497109827", label="0.45",style=bold];
"KRT33B_S" -> "ROS1_M" [color="0 0 0.665895953757", label="0.46",style=bold];
"SLC26A3_S" -> "C16orf11_E" [color="0 0 0.663294797688", label="0.47",style=bold];
"HOXA7_S" -> "C16orf11_E" [color="0 0 0.660693641618", label="0.47",style=bold];
"XDH_S" -> "DEFB119_E" [color="0 0 0.658092485549", label="0.47",style=bold];
"KERA_E" -> "MUC16_E" [color="0 0 0.65549132948", label="0.48",style=bold];
"FLG2_S" -> "OPN5_M" [color="0 0 0.65289017341", label="0.49",style=bold];
"ROS1_S" -> "OPN5_M" [color="0 0 0.650289017341", label="0.49",style=bold];
"C16orf11_S" -> "SLC6A18_M" [color="0 0 0.647687861272", label="0.50",style=bold];
"SLC6A18_S" -> "Tumor_Status" [color="0 0 0.645086705202", label="0.51",style=bold];
"BFSP2_S" -> "DDX4_E" [color="0 0 0.642485549133", label="0.51",style=bold];
"XDH_S" -> "DEFB119_M" [color="0 0 0.639884393064", label="0.51",style=bold];
"BFSP2_S" -> "HOXA7_E" [color="0 0 0.637283236994", label="0.51",style=bold];
"DEFB119_S" -> "RPL10L_M" [color="0 0 0.634682080925", label="0.51",style=bold];
"VIL1_S" -> "MMP1_E" [color="0 0 0.632080924855", label="0.54",style=bold];
"XDH_E" -> "OPN5_M" [color="0 0 0.629479768786", label="0.54",style=bold];
"ABCG8_S" -> "KRT33B_M" [color="0 0 0.626878612717", label="0.56",style=bold];
"KERA_S" -> "MMP1_M" [color="0 0 0.624277456647", label="0.57",style=bold];
"KERA_S" -> "TNNT3_E" [color="0 0 0.621676300578", label="0.58",style=bold];
"C16orf11_S" -> "HOXA7_E" [color="0 0 0.619075144509", label="0.59",style=bold];
"BFSP2_S" -> "ROS1_E" [color="0 0 0.616473988439", label="0.61",style=bold];
"AFP_S" -> "Tumor_Status" [color="0 0 0.61387283237", label="0.61",style=bold];
"MUC16_E" -> "Tumor_Status" [color="0 0 0.611271676301", label="0.66",style=bold];
"TMPRSS11F_S" -> "XDH_E" [color="0 0 0.608670520231", label="0.67",style=bold];
"ROS1_S" -> "DEFB119_M" [color="0 0 0.606069364162", label="0.69",style=bold];
"SLC26A3_S" -> "CHIA_M" [color="0 0 0.603468208092", label="0.69",style=bold];
"VIL1_S" -> "CHIA_M" [color="0 0 0.600867052023", label="0.69",style=bold];
"FLG2_S" -> "NPSR1_E" [color="0 0 0.598265895954", label="0.69",style=bold];
"TNNT3_S" -> "MUC16_M" [color="0 0 0.595664739884", label="0.69",style=bold];
"TNNT3_S" -> "HOXA7_M" [color="0 0 0.593063583815", label="0.69",style=bold];
"TMPRSS11F_S" -> "RPL10L_E" [color="0 0 0.590462427746", label="0.73",style=bold];
"KERA_S" -> "ROS1_M" [color="0 0 0.587861271676", label="0.74",style=bold];
"KRT33B_S" -> "KRT33B_M" [color="0 0 0.585260115607", label="0.75",style=bold];
"HOXA7_S" -> "KERA_M" [color="0 0 0.582658959538", label="0.75",style=bold];
"IL31RA_S" -> "CHIA_M" [color="0 0 0.580057803468", label="0.76",style=bold];
"RPL10L_E" -> "DDX4_M" [color="0 0 0.577456647399", label="0.76",style=bold];
"CHIA_S" -> "MMP1_M" [color="0 0 0.574855491329", label="0.76",style=bold];
"KERA_S" -> "TNNT3_M" [color="0 0 0.57225433526", label="0.76",style=bold];
"VIL1_E" -> "FLG2_E" [color="0 0 0.569653179191", label="0.77",style=bold];
"OPN5_S" -> "HOXA7_M" [color="0 0 0.567052023121", label="0.77",style=bold];
"VIL1_S" -> "C16orf11_E" [color="0 0 0.564450867052", label="0.79",style=bold];
"SLC6A18_S" -> "AFP_M" [color="0 0 0.561849710983", label="0.81",style=bold];
"TNNT3_S" -> "FLG2_E" [color="0 0 0.559248554913", label="0.81",style=bold];
"SLC26A3_S" -> "ROS1_M" [color="0 0 0.556647398844", label="0.81",style=bold];
"AFP_S" -> "OPN5_M" [color="0 0 0.554046242775", label="0.82",style=bold];
"ROS1_S" -> "AFP_E" [color="0 0 0.551445086705", label="0.83",style=bold];
"XDH_S" -> "SLC26A3_E" [color="0 0 0.548843930636", label="0.84",style=bold];
"CHIA_S" -> "CHIA_M" [color="0 0 0.546242774566", label="0.85",style=bold];
"SLC6A18_S" -> "KRT33B_M" [color="0 0 0.543641618497", label="0.85",style=bold];
"MUC16_E" -> "BFSP2_E" [color="0 0 0.541040462428", label="0.87",style=bold];
"AFP_E" -> "OPN5_E" [color="0 0 0.538439306358", label="0.88",style=bold];
"KERA_S" -> "VIL1_E" [color="0 0 0.535838150289", label="0.89",style=bold];
"KERA_S" -> "DEFB119_E" [color="0 0 0.53323699422", label="0.90",style=bold];
"CHIA_S" -> "TMPRSS11F_M" [color="0 0 0.53063583815", label="0.92",style=bold];
"C16orf11_S" -> "RPL10L_M" [color="0 0 0.528034682081", label="0.92",style=bold];
"OPN5_S" -> "BFSP2_M" [color="0 0 0.525433526012", label="0.93",style=bold];
"TNNT3_M" -> "TMPRSS11F_E" [color="0 0 0.522832369942", label="0.93",style=bold];
"AFP_S" -> "BFSP2_M" [color="0 0 0.520231213873", label="0.94",style=bold];
"CHIA_S" -> "VIL1_E" [color="0 0 0.517630057803", label="0.94",style=bold];
"KERA_E" -> "DEFB119_M" [color="0 0 0.515028901734", label="0.94",style=bold];
"DDX4_S" -> "IL31RA_E" [color="0 0 0.512427745665", label="0.94",style=bold];
"RPL10L_S" -> "IL31RA_E" [color="0 0 0.509826589595", label="0.94",style=bold];
"TNNT3_S" -> "IL31RA_E" [color="0 0 0.507225433526", label="0.94",style=bold];
"AFP_S" -> "C16orf11_M" [color="0 0 0.504624277457", label="0.96",style=bold];
"C16orf11_S" -> "MMP1_M" [color="0 0 0.502023121387", label="0.98",style=bold];
"ROS1_S" -> "SLC6A18_M" [color="0 0 0.499421965318", label="0.98",style=bold];
"TMPRSS11F_S" -> "NPSR1_E" [color="0 0 0.496820809249", label="0.98",style=bold];
"ABCG8_S" -> "FLG2_M" [color="0 0 0.494219653179", label="0.98",style=bold];
"ROS1_S" -> "C16orf11_E" [color="0 0 0.49161849711", label="0.98",style=bold];
"FLG2_S" -> "BFSP2_E" [color="0 0 0.48901734104", label="0.98",style=bold];
"XDH_S" -> "VIL1_E" [color="0 0 0.486416184971", label="0.99",style=bold];
"AFP_S" -> "DDX4_E" [color="0 0 0.483815028902", label="0.99",style=bold];
"MUC16_E" -> "DDX4_E" [color="0 0 0.481213872832", label="1.00",style=bold];
"RPL10L_S" -> "ROS1_M" [color="0 0 0.478612716763", label="1.00",style=bold];
"DDX4_S" -> "ROS1_M" [color="0 0 0.476011560694", label="1.00",style=bold];
"DEFB119_S" -> "NPSR1_E" [color="0 0 0.473410404624", label="1.01",style=bold];
"AFP_S" -> "TNNT3_M" [color="0 0 0.470809248555", label="1.02",style=bold];
"CHIA_E" -> "HOXA7_M" [color="0 0 0.468208092486", label="1.02",style=bold];
"TMPRSS11F_S" -> "MUC16_M" [color="0 0 0.465606936416", label="1.03",style=bold];
"KERA_S" -> "KRT33B_E" [color="0 0 0.463005780347", label="1.04",style=bold];
"KERA_S" -> "NPSR1_M" [color="0 0 0.460404624277", label="1.06",style=bold];
"BFSP2_S" -> "RPL10L_M" [color="0 0 0.457803468208", label="1.07",style=bold];
"XDH_S" -> "NPSR1_E" [color="0 0 0.455202312139", label="1.08",style=bold];
"RPL10L_E" -> "VIL1_M" [color="0 0 0.452601156069", label="1.08",style=bold];
"MUC16_S" -> "TMPRSS11F_M" [color="0 0 0.45", label="1.11",style=bold];
"ROS1_M" -> "CHIA_E" [color="0 0 0.447398843931", label="1.13",style=bold];
"TMPRSS11F_S" -> "RPL10L_M" [color="0 0 0.444797687861", label="1.13",style=bold];
"VIL1_S" -> "TNNT3_E" [color="0 0 0.442196531792", label="1.15",style=bold];
"XDH_S" -> "KERA_E" [color="0 0 0.439595375723", label="1.16",style=bold];
"TMPRSS11F_S" -> "BFSP2_E" [color="0 0 0.436994219653", label="1.19",style=bold];
"SLC6A18_S" -> "DEFB119_M" [color="0 0 0.434393063584", label="1.20",style=bold];
"VIL1_S" -> "CHIA_E" [color="0 0 0.431791907514", label="1.20",style=bold];
"TMPRSS11F_S" -> "ROS1_M" [color="0 0 0.429190751445", label="1.20",style=bold];
"CHIA_S" -> "ABCG8_E" [color="0 0 0.426589595376", label="1.21",style=bold];
"ROS1_S" -> "ABCG8_E" [color="0 0 0.423988439306", label="1.21",style=bold];
"AFP_S" -> "ABCG8_E" [color="0 0 0.421387283237", label="1.25",style=bold];
"NPSR1_M" -> "KRT33B_M" [color="0 0 0.418786127168", label="1.26",style=bold];
"ROS1_M" -> "RPL10L_M" [color="0 0 0.416184971098", label="1.28",style=bold];
"KERA_S" -> "DDX4_E" [color="0 0 0.413583815029", label="1.30",style=bold];
"ROS1_S" -> "NPSR1_M" [color="0 0 0.41098265896", label="1.34",style=bold];
"CHIA_S" -> "TMPRSS11F_E" [color="0 0 0.40838150289", label="1.36",style=bold];
"VIL1_S" -> "RPL10L_E" [color="0 0 0.405780346821", label="1.37",style=bold];
"XDH_M" -> "DDX4_M" [color="0 0 0.403179190751", label="1.37",style=bold];
"BFSP2_S" -> "FLG2_M" [color="0 0 0.400578034682", label="1.39",style=bold];
"TNNT3_E" -> "C16orf11_E" [color="0 0 0.397976878613", label="1.40",style=bold];
"KERA_S" -> "BFSP2_M" [color="0 0 0.395375722543", label="1.41",style=bold];
"VIL1_M" -> "KERA_M" [color="0 0 0.392774566474", label="1.43",style=bold];
"KRT33B_S" -> "CHIA_E" [color="0 0 0.390173410405", label="1.46",style=bold];
"HOXA7_S" -> "OPN5_M" [color="0 0 0.387572254335", label="1.50",style=bold];
"CHIA_S" -> "TNNT3_E" [color="0 0 0.384971098266", label="1.52",style=bold];
"RPL10L_E" -> "TMPRSS11F_M" [color="0 0 0.382369942197", label="1.53",style=bold];
"AFP_S" -> "AFP_M" [color="0 0 0.379768786127", label="1.57",style=bold];
"ROS1_S" -> "AFP_M" [color="0 0 0.377167630058", label="1.63",style=bold];
"DEFB119_S" -> "AFP_M" [color="0 0 0.374566473988", label="1.63",style=bold];
"MUC16_E" -> "IL31RA_E" [color="0 0 0.371965317919", label="1.65",style=bold];
"ROS1_S" -> "TMPRSS11F_M" [color="0 0 0.36936416185", label="1.65",style=bold];
"BFSP2_S" -> "SLC26A3_E" [color="0 0 0.36676300578", label="1.67",style=bold];
"KRT33B_S" -> "SLC26A3_M" [color="0 0 0.364161849711", label="1.67",style=bold];
"HOXA7_S" -> "DEFB119_M" [color="0 0 0.361560693642", label="1.67",style=bold];
"MMP1_S" -> "TMPRSS11F_E" [color="0 0 0.358959537572", label="1.73",style=bold];
"TNNT3_M" -> "FLG2_M" [color="0 0 0.356358381503", label="1.75",style=bold];
"DEFB119_S" -> "ROS1_E" [color="0 0 0.353757225434", label="1.80",style=bold];
"XDH_S" -> "TNNT3_E" [color="0 0 0.351156069364", label="1.83",style=bold];
"AFP_E" -> "KERA_E" [color="0 0 0.348554913295", label="1.89",style=bold];
"TMPRSS11F_S" -> "BFSP2_M" [color="0 0 0.345953757225", label="1.90",style=bold];
"C16orf11_S" -> "DDX4_E" [color="0 0 0.343352601156", label="1.90",style=bold];
"RPL10L_E" -> "TNNT3_E" [color="0 0 0.340751445087", label="1.91",style=bold];
"BFSP2_S" -> "MUC16_E" [color="0 0 0.338150289017", label="2.00",style=bold];
"FLG2_S" -> "MUC16_E" [color="0 0 0.335549132948", label="2.00",style=bold];
"ROS1_S" -> "CHIA_E" [color="0 0 0.332947976879", label="2.00",style=bold];
"OPN5_S" -> "ROS1_M" [color="0 0 0.330346820809", label="2.05",style=bold];
"XDH_E" -> "NPSR1_E" [color="0 0 0.32774566474", label="2.11",style=bold];
"CHIA_S" -> "OPN5_E" [color="0 0 0.325144508671", label="2.18",style=bold];
"MMP1_S" -> "OPN5_E" [color="0 0 0.322543352601", label="2.18",style=bold];
"CHIA_S" -> "OPN5_M" [color="0 0 0.319942196532", label="2.20",style=bold];
"RPL10L_E" -> "RPL10L_M" [color="0 0 0.317341040462", label="2.21",style=bold];
"SLC6A18_M" -> "IL31RA_M" [color="0 0 0.314739884393", label="2.23",style=bold];
"KERA_E" -> "Tumor_Status" [color="0 0 0.312138728324", label="2.23",style=bold];
"DDX4_S" -> "AFP_M" [color="0 0 0.309537572254", label="2.23",style=bold];
"RPL10L_S" -> "AFP_M" [color="0 0 0.306936416185", label="2.23",style=bold];
"C16orf11_M" -> "HOXA7_E" [color="0 0 0.304335260116", label="2.26",style=bold];
"BFSP2_S" -> "MUC16_S" [color="0 0 0.301734104046", label="2.28",style=bold];
"C16orf11_S" -> "MUC16_S" [color="0 0 0.299132947977", label="2.28",style=bold];
"SLC6A18_S" -> "DDX4_E" [color="0 0 0.296531791908", label="2.30",style=bold];
"VIL1_E" -> "C16orf11_E" [color="0 0 0.293930635838", label="2.32",style=bold];
"SLC26A3_E" -> "DDX4_E" [color="0 0 0.291329479769", label="2.34",style=bold];
"OPN5_S" -> "XDH_E" [color="0 0 0.288728323699", label="2.36",style=bold];
"TNNT3_S" -> "XDH_E" [color="0 0 0.28612716763", label="2.36",style=bold];
"RPL10L_S" -> "CHIA_M" [color="0 0 0.283526011561", label="2.37",style=bold];
"MMP1_S" -> "VIL1_E" [color="0 0 0.280924855491", label="2.41",style=bold];
"SLC6A18_S" -> "VIL1_E" [color="0 0 0.278323699422", label="2.41",style=bold];
"TMPRSS11F_S" -> "TNNT3_E" [color="0 0 0.275722543353", label="2.43",style=bold];
"CHIA_S" -> "FLG2_E" [color="0 0 0.273121387283", label="2.44",style=bold];
"DDX4_S" -> "KRT33B_E" [color="0 0 0.270520231214", label="2.48",style=bold];
"VIL1_S" -> "DEFB119_E" [color="0 0 0.267919075145", label="2.49",style=bold];
"AFP_S" -> "DEFB119_E" [color="0 0 0.265317919075", label="2.49",style=bold];
"KERA_S" -> "DEFB119_M" [color="0 0 0.262716763006", label="2.49",style=bold];
"SLC6A18_E" -> "BFSP2_E" [color="0 0 0.260115606936", label="2.54",style=bold];
"FLG2_E" -> "TMPRSS11F_M" [color="0 0 0.257514450867", label="2.70",style=bold];
"TMPRSS11F_M" -> "Tumor_Status" [color="0 0 0.254913294798", label="2.76",style=bold];
"CHIA_S" -> "FLG2_M" [color="0 0 0.252312138728", label="2.77",style=bold];
"ROS1_M" -> "DEFB119_M" [color="0 0 0.249710982659", label="3.01",style=bold];
"TNNT3_E" -> "NPSR1_E" [color="0 0 0.24710982659", label="3.05",style=bold];
"AFP_S" -> "C16orf11_E" [color="0 0 0.24450867052", label="3.06",style=bold];
"XDH_M" -> "MMP1_M" [color="0 0 0.241907514451", label="3.21",style=bold];
"OPN5_E" -> "SLC26A3_M" [color="0 0 0.239306358382", label="3.32",style=bold];
"KRT33B_M" -> "RPL10L_M" [color="0 0 0.236705202312", label="3.59",style=bold];
"HOXA7_E" -> "VIL1_E" [color="0 0 0.234104046243", label="3.60",style=bold];
"ABCG8_M" -> "FLG2_M" [color="0 0 0.231502890173", label="3.63",style=bold];
"VIL1_M" -> "XDH_M" [color="0 0 0.228901734104", label="3.68",style=bold];
"CHIA_E" -> "DEFB119_M" [color="0 0 0.226300578035", label="3.76",style=bold];
"ABCG8_E" -> "C16orf11_E" [color="0 0 0.223699421965", label="3.77",style=bold];
"CHIA_M" -> "DEFB119_M" [color="0 0 0.221098265896", label="3.85",style=bold];
"DEFB119_E" -> "DDX4_E" [color="0 0 0.218497109827", label="4.02",style=bold];
"CHIA_E" -> "Tumor_Status" [color="0 0 0.215895953757", label="4.09",style=bold];
"MUC16_S" -> "SLC26A3_E" [color="0 0 0.213294797688", label="4.14",style=bold];
"AFP_E" -> "NPSR1_E" [color="0 0 0.210693641618", label="4.14",style=bold];
"DEFB119_S" -> "OPN5_S" [color="0 0 0.208092485549", label="4.23",style=bold];
"C16orf11_S" -> "SLC26A3_S" [color="0 0 0.20549132948", label="4.23",style=bold];
"KERA_S" -> "C16orf11_M" [color="0 0 0.20289017341", label="4.26",style=bold];
"RPL10L_E" -> "CHIA_E" [color="0 0 0.200289017341", label="4.27",style=bold];
"RPL10L_E" -> "BFSP2_M" [color="0 0 0.197687861272", label="4.41",style=bold];
"AFP_E" -> "Tumor_Status" [color="0 0 0.195086705202", label="4.41",style=bold];
"SLC6A18_E" -> "DEFB119_M" [color="0 0 0.192485549133", label="4.59",style=bold];
"CHIA_E" -> "CHIA_M" [color="0 0 0.189884393064", label="4.61",style=bold];
"MUC16_E" -> "DEFB119_M" [color="0 0 0.187283236994", label="4.66",style=bold];
"VIL1_M" -> "CHIA_M" [color="0 0 0.184682080925", label="4.71",style=bold];
"DEFB119_E" -> "CHIA_M" [color="0 0 0.182080924855", label="4.74",style=bold];
"XDH_E" -> "HOXA7_M" [color="0 0 0.179479768786", label="4.79",style=bold];
"IL31RA_S" -> "FLG2_S" [color="0 0 0.176878612717", label="4.92",style=bold];
"KRT33B_E" -> "ROS1_E" [color="0 0 0.174277456647", label="4.98",style=bold];
"KRT33B_E" -> "AFP_E" [color="0 0 0.171676300578", label="5.10",style=bold];
"ABCG8_M" -> "ROS1_M" [color="0 0 0.169075144509", label="5.12",style=bold];
"TNNT3_E" -> "DDX4_E" [color="0 0 0.166473988439", label="5.31",style=bold];
"AFP_E" -> "XDH_E" [color="0 0 0.16387283237", label="5.43",style=bold];
"VIL1_M" -> "IL31RA_M" [color="0 0 0.161271676301", label="5.46",style=bold];
"DDX4_M" -> "AFP_M" [color="0 0 0.158670520231", label="5.52",style=bold];
"SLC6A18_M" -> "IL31RA_E" [color="0 0 0.156069364162", label="5.77",style=bold];
"DEFB119_E" -> "TMPRSS11F_E" [color="0 0 0.153468208092", label="5.95",style=bold];
"KRT33B_M" -> "OPN5_M" [color="0 0 0.150867052023", label="6.11",style=bold];
"MMP1_M" -> "KRT33B_M" [color="0 0 0.148265895954", label="6.22",style=bold];
"NPSR1_M" -> "KRT33B_E" [color="0 0 0.145664739884", label="6.30",style=bold];
"KERA_E" -> "SLC26A3_E" [color="0 0 0.143063583815", label="6.38",style=bold];
"SLC6A18_M" -> "FLG2_M" [color="0 0 0.140462427746", label="6.40",style=bold];
"ABCG8_M" -> "C16orf11_M" [color="0 0 0.137861271676", label="6.53",style=bold];
"ABCG8_M" -> "ABCG8_E" [color="0 0 0.135260115607", label="6.92",style=bold];
"SLC6A18_E" -> "DEFB119_E" [color="0 0 0.132658959538", label="6.97",style=bold];
"RPL10L_E" -> "TNNT3_M" [color="0 0 0.130057803468", label="7.00",style=bold];
"TMPRSS11F_M" -> "CHIA_M" [color="0 0 0.127456647399", label="7.00",style=bold];
"DEFB119_E" -> "CHIA_E" [color="0 0 0.124855491329", label="7.05",style=bold];
"SLC6A18_E" -> "ABCG8_E" [color="0 0 0.12225433526", label="7.10",style=bold];
"SLC6A18_M" -> "HOXA7_E" [color="0 0 0.119653179191", label="7.13",style=bold];
"XDH_M" -> "KERA_M" [color="0 0 0.117052023121", label="7.14",style=bold];
"SLC6A18_M" -> "MMP1_M" [color="0 0 0.114450867052", label="7.49",style=bold];
"AFP_E" -> "TNNT3_E" [color="0 0 0.111849710983", label="7.60",style=bold];
"ABCG8_E" -> "ROS1_E" [color="0 0 0.109248554913", label="7.81",style=bold];
"HOXA7_E" -> "AFP_E" [color="0 0 0.106647398844", label="7.84",style=bold];
"BFSP2_M" -> "RPL10L_M" [color="0 0 0.104046242775", label="7.95",style=bold];
"DEFB119_E" -> "MUC16_M" [color="0 0 0.101445086705", label="8.21",style=bold];
"DEFB119_M" -> "MUC16_M" [color="0 0 0.0988439306358", label="8.47",style=bold];
"ABCG8_E" -> "TNNT3_E" [color="0 0 0.0962427745665", label="8.64",style=bold];
"ROS1_E" -> "C16orf11_E" [color="0 0 0.0936416184971", label="8.78",style=bold];
"TMPRSS11F_M" -> "NPSR1_E" [color="0 0 0.0910404624277", label="9.25",style=bold];
"XDH_M" -> "OPN5_M" [color="0 0 0.0884393063584", label="9.41",style=bold];
"DDX4_M" -> "MMP1_M" [color="0 0 0.085838150289", label="9.52",style=bold];
"DDX4_M" -> "SLC26A3_M" [color="0 0 0.0832369942197", label="9.85",style=bold];
"VIL1_M" -> "BFSP2_M" [color="0 0 0.0806358381503", label="10.46",style=bold];
"TNNT3_M" -> "NPSR1_M" [color="0 0 0.0780346820809", label="10.47",style=bold];
"DDX4_M" -> "CHIA_M" [color="0 0 0.0754335260116", label="10.67",style=bold];
"HOXA7_E" -> "SLC6A18_E" [color="0 0 0.0728323699422", label="10.87",style=bold];
"AFP_E" -> "HOXA7_M" [color="0 0 0.0702312138728", label="11.92",style=bold];
"Tumor_Status" -> "DDX4_E" [color="0 0 0.0676300578035", label="12.19",style=bold];
"BFSP2_M" -> "ROS1_M" [color="0 0 0.0650289017341", label="12.30",style=bold];
"SLC6A18_M" -> "MUC16_M" [color="0 0 0.0624277456647", label="12.35",style=bold];
"DEFB119_E" -> "HOXA7_M" [color="0 0 0.0598265895954", label="13.70",style=bold];
"BFSP2_M" -> "AFP_E" [color="0 0 0.057225433526", label="13.78",style=bold];
"HOXA7_E" -> "FLG2_M" [color="0 0 0.0546242774566", label="13.96",style=bold];
"MMP1_M" -> "Tumor_Status" [color="0 0 0.0520231213873", label="14.13",style=bold];
"KERA_M" -> "TMPRSS11F_M" [color="0 0 0.0494219653179", label="14.21",style=bold];
"TMPRSS11F_E" -> "NPSR1_E" [color="0 0 0.0468208092486", label="14.33",style=bold];
"BFSP2_M" -> "IL31RA_M" [color="0 0 0.0442196531792", label="15.37",style=bold];
"SLC6A18_M" -> "KRT33B_M" [color="0 0 0.0416184971098", label="16.64",style=bold];
"XDH_M" -> "NPSR1_M" [color="0 0 0.0390173410405", label="16.65",style=bold];
"VIL1_M" -> "TMPRSS11F_M" [color="0 0 0.0364161849711", label="16.67",style=bold];
"ABCG8_M" -> "BFSP2_M" [color="0 0 0.0338150289017", label="18.05",style=bold];
"TMPRSS11F_M" -> "AFP_M" [color="0 0 0.0312138728324", label="18.96",style=bold];
"HOXA7_E" -> "HOXA7_M" [color="0 0 0.028612716763", label="23.05",style=bold];
"KERA_M" -> "DEFB119_E" [color="0 0 0.0260115606936", label="23.56",style=bold];
"TNNT3_M" -> "ABCG8_M" [color="0 0 0.0234104046243", label="23.72",style=bold];
"TMPRSS11F_M" -> "DDX4_M" [color="0 0 0.0208092485549", label="23.88",style=bold];
"VIL1_M" -> "ABCG8_M" [color="0 0 0.0182080924855", label="23.98",style=bold];
"BFSP2_M" -> "KERA_M" [color="0 0 0.0156069364162", label="24.65",style=bold];
"SLC6A18_M" -> "XDH_M" [color="0 0 0.0130057803468", label="26.19",style=bold];
"SLC6A18_M" -> "SLC26A3_M" [color="0 0 0.0104046242775", label="29.35",style=bold];
"KRT33B_E" -> "ABCG8_E" [color="0 0 0.00780346820809", label="32.16",style=bold];
"TNNT3_M" -> "C16orf11_M" [color="0 0 0.00520231213873", label="34.78",style=bold];
"VIL1_M" -> "TNNT3_M" [color="0 0 0.00260115606936", label="73.88",style=bold];
"TNNT3_M" -> "SLC6A18_M" [color="0 0 0.0", label="119.89",style=bold];
}
```
