## Supplementary material for "New analysis framework incorporating mixed mutual information and scalable Bayesian networks for multimodal high dimensional genomic and epigenomic cancer data": Suppl Table 8

```
digraph G{
 ratio=fill;
 node [shape=box, style=rounded];
"ABCG8_S";
"ABCG8_E";
"ABCG8_M";
"AFP_S";
"AFP_E";
"AFP_M";
"ALDH3B2_S";
"ALDH3B2_E";
"ALDH3B2_M";
"CHIA_S";
"CHIA_E";
"CHIA_M";
"DEFB119_S";
"DEFB119_E";
"DEFB119_M";
"DMRTB1_S";
"DMRTB1_E";
"DMRTB1_M";
"FCRL5_S";
"FCRL5_E";
"FCRL5_M";
"GABRP_S";
"GABRP_E";
"GABRP_M";
"HOXA11_S";
"HOXA11_E";
"HOXA11_M";
"KRT75_S";
"KRT75_E";
"KRT75_M";
"LAD1_S";
"LAD1_E";
"LAD1_M";
"LGSN_S";
"LGSN_E";
"LGSN_M";
"LMX1A_S";
"LMX1A_E";
"LMX1A_M";
"MMP1_S";
"MMP1_E";
"MMP1_M";
"OPN5_S";
"OPN5_E";
"OPN5_M";
"PKHD1L1_S";
"PKHD1L1_E";
"PKHD1L1_M";
"PLA2G4D_S";
"PLA2G4D_E";
"PLA2G4D_M";
"PROKR1_S";
"PROKR1_E";
"PROKR1_M";
"SLC6A18_S";
"SLC6A18_E";
"SLC6A18_M";
"SLCO6A1_S";
"SLCO6A1_E";
"SLCO6A1_M";
"STAB2_S";
"STAB2_E";
"STAB2_M";
"TMPRSS11F_S";
"TMPRSS11F_E";
"TMPRSS11F_M";
"TRPM1_S";
"TRPM1_E";
"TRPM1_M";
"XDH_S";
"XDH_E";
"XDH_M";
"Survival";
"TRPM1_S" -> "Survival" [color="0 0 0.9", label="0.02",style=bold];
"PROKR1_S" -> "LMX1A_M" [color="0 0 0.89738372093", label="0.03",style=bold];
"MMP1_S" -> "LMX1A_M" [color="0 0 0.89476744186", label="0.04",style=bold];
"FCRL5_S" -> "LMX1A_M" [color="0 0 0.892151162791", label="0.04",style=bold];
"TMPRSS11F_S" -> "PKHD1L1_E" [color="0 0 0.889534883721", label="0.05",style=bold];
"DEFB119_S" -> "PKHD1L1_E" [color="0 0 0.886918604651", label="0.05",style=bold];
"SLCO6A1_S" -> "PKHD1L1_E" [color="0 0 0.884302325581", label="0.05",style=bold];
"SLC6A18_S" -> "DEFB119_E" [color="0 0 0.881686046512", label="0.06",style=bold];
"SLCO6A1_S" -> "ALDH3B2_E" [color="0 0 0.879069767442", label="0.07",style=bold];
"OPN5_S" -> "XDH_M" [color="0 0 0.876453488372", label="0.07",style=bold];
"ALDH3B2_S" -> "XDH_M" [color="0 0 0.873837209302", label="0.07",style=bold];
"XDH_S" -> "XDH_M" [color="0 0 0.871220930233", label="0.07",style=bold];
"ABCG8_S" -> "XDH_M" [color="0 0 0.868604651163", label="0.07",style=bold];
"SLC6A18_S" -> "XDH_M" [color="0 0 0.865988372093", label="0.07",style=bold];
"GABRP_S" -> "XDH_M" [color="0 0 0.863372093023", label="0.07",style=bold];
"FCRL5_S" -> "LGSN_E" [color="0 0 0.860755813953", label="0.08",style=bold];
"ABCG8_S" -> "DMRTB1_E" [color="0 0 0.858139534884", label="0.09",style=bold];
"SLC6A18_S" -> "MMP1_E" [color="0 0 0.855523255814", label="0.09",style=bold];
"MMP1_S" -> "MMP1_E" [color="0 0 0.852906976744", label="0.09",style=bold];
"OPN5_S" -> "MMP1_E" [color="0 0 0.850290697674", label="0.09",style=bold];
"GABRP_S" -> "MMP1_E" [color="0 0 0.847674418605", label="0.09",style=bold];
"CHIA_S" -> "MMP1_E" [color="0 0 0.845058139535", label="0.09",style=bold];
"XDH_S" -> "MMP1_E" [color="0 0 0.842441860465", label="0.09",style=bold];
"STAB2_S" -> "MMP1_E" [color="0 0 0.839825581395", label="0.09",style=bold];
"ABCG8_S" -> "ALDH3B2_M" [color="0 0 0.837209302326", label="0.10",style=bold];
"FCRL5_S" -> "MMP1_M" [color="0 0 0.834593023256", label="0.11",style=bold];
"ALDH3B2_S" -> "PROKR1_M" [color="0 0 0.831976744186", label="0.11",style=bold];
"ABCG8_S" -> "PROKR1_E" [color="0 0 0.829360465116", label="0.12",style=bold];
"XDH_S" -> "DEFB119_E" [color="0 0 0.826744186047", label="0.12",style=bold];
"DEFB119_S" -> "DEFB119_E" [color="0 0 0.824127906977", label="0.12",style=bold];
"SLC6A18_S" -> "STAB2_E" [color="0 0 0.821511627907", label="0.15",style=bold];
"DEFB119_S" -> "LAD1_E" [color="0 0 0.818895348837", label="0.15",style=bold];
"MMP1_S" -> "ALDH3B2_M" [color="0 0 0.816279069767", label="0.15",style=bold];
"MMP1_S" -> "GABRP_E" [color="0 0 0.813662790698", label="0.15",style=bold];
"PROKR1_S" -> "STAB2_E" [color="0 0 0.811046511628", label="0.15",style=bold];
"ABCG8_S" -> "TRPM1_M" [color="0 0 0.808430232558", label="0.15",style=bold];
"DEFB119_S" -> "ALDH3B2_E" [color="0 0 0.805813953488", label="0.18",style=bold];
"TRPM1_S" -> "DEFB119_E" [color="0 0 0.803197674419", label="0.18",style=bold];
"DEFB119_S" -> "PLA2G4D_M" [color="0 0 0.800581395349", label="0.19",style=bold];
"STAB2_S" -> "HOXA11_M" [color="0 0 0.797965116279", label="0.20",style=bold];
"PKHD1L1_S" -> "ABCG8_E" [color="0 0 0.795348837209", label="0.20",style=bold];
"PKHD1L1_S" -> "HOXA11_E" [color="0 0 0.79273255814", label="0.20",style=bold];
"KRT75_S" -> "TRPM1_M" [color="0 0 0.79011627907", label="0.21",style=bold];
"LAD1_S" -> "AFP_M" [color="0 0 0.7875", label="0.22",style=bold];
"KRT75_S" -> "GABRP_E" [color="0 0 0.78488372093", label="0.22",style=bold];
"GABRP_S" -> "ABCG8_M" [color="0 0 0.78226744186", label="0.22",style=bold];
"FCRL5_E" -> "KRT75_M" [color="0 0 0.779651162791", label="0.23",style=bold];
"SLC6A18_S" -> "AFP_M" [color="0 0 0.777034883721", label="0.23",style=bold];
"ABCG8_S" -> "Survival" [color="0 0 0.774418604651", label="0.24",style=bold];
"KRT75_S" -> "ABCG8_E" [color="0 0 0.771802325581", label="0.25",style=bold];
"ABCG8_S" -> "GABRP_M" [color="0 0 0.769186046512", label="0.25",style=bold];
"PROKR1_S" -> "PLA2G4D_E" [color="0 0 0.766569767442", label="0.26",style=bold];
"SLCO6A1_S" -> "Survival" [color="0 0 0.763953488372", label="0.28",style=bold];
"DEFB119_S" -> "Survival" [color="0 0 0.761337209302", label="0.28",style=bold];
"SLCO6A1_S" -> "AFP_E" [color="0 0 0.758720930233", label="0.28",style=bold];
"STAB2_S" -> "AFP_E" [color="0 0 0.756104651163", label="0.28",style=bold];
"XDH_S" -> "TMPRSS11F_E" [color="0 0 0.753488372093", label="0.29",style=bold];
"SLC6A18_S" -> "TMPRSS11F_E" [color="0 0 0.750872093023", label="0.29",style=bold];
"XDH_S" -> "TMPRSS11F_M" [color="0 0 0.748255813953", label="0.29",style=bold];
"MMP1_S" -> "STAB2_E" [color="0 0 0.745639534884", label="0.29",style=bold];
"KRT75_S" -> "FCRL5_M" [color="0 0 0.743023255814", label="0.29",style=bold];
"DEFB119_S" -> "FCRL5_M" [color="0 0 0.740406976744", label="0.29",style=bold];
"TRPM1_S" -> "STAB2_M" [color="0 0 0.737790697674", label="0.29",style=bold];
"ALDH3B2_S" -> "MMP1_M" [color="0 0 0.735174418605", label="0.29",style=bold];
"CHIA_S" -> "GABRP_E" [color="0 0 0.732558139535", label="0.29",style=bold];
"SLCO6A1_S" -> "GABRP_E" [color="0 0 0.729941860465", label="0.29",style=bold];
"LAD1_S" -> "DMRTB1_E" [color="0 0 0.727325581395", label="0.29",style=bold];
"TRPM1_S" -> "STAB2_E" [color="0 0 0.724709302326", label="0.29",style=bold];
"TRPM1_S" -> "OPN5_M" [color="0 0 0.722093023256", label="0.29",style=bold];
"TRPM1_S" -> "TRPM1_M" [color="0 0 0.719476744186", label="0.29",style=bold];
"MMP1_S" -> "LMX1A_E" [color="0 0 0.716860465116", label="0.29",style=bold];
"KRT75_S" -> "OPN5_M" [color="0 0 0.714244186047", label="0.29",style=bold];
"DEFB119_S" -> "OPN5_M" [color="0 0 0.711627906977", label="0.29",style=bold];
"CHIA_S" -> "LGSN_M" [color="0 0 0.709011627907", label="0.29",style=bold];
"SLC6A18_S" -> "TRPM1_M" [color="0 0 0.706395348837", label="0.29",style=bold];
"GABRP_S" -> "KRT75_M" [color="0 0 0.703779069767", label="0.29",style=bold];
"ALDH3B2_S" -> "PKHD1L1_E" [color="0 0 0.701162790698", label="0.29",style=bold];
"SLCO6A1_S" -> "LGSN_M" [color="0 0 0.698546511628", label="0.29",style=bold];
"ALDH3B2_S" -> "CHIA_M" [color="0 0 0.695930232558", label="0.29",style=bold];
"PKHD1L1_S" -> "MMP1_M" [color="0 0 0.693313953488", label="0.31",style=bold];
"TRPM1_S" -> "PROKR1_E" [color="0 0 0.690697674419", label="0.32",style=bold];
"XDH_S" -> "PROKR1_E" [color="0 0 0.688081395349", label="0.32",style=bold];
"GABRP_S" -> "HOXA11_M" [color="0 0 0.685465116279", label="0.32",style=bold];
"MMP1_S" -> "TRPM1_M" [color="0 0 0.682848837209", label="0.32",style=bold];
"FCRL5_S" -> "XDH_M" [color="0 0 0.68023255814", label="0.36",style=bold];
"DEFB119_S" -> "STAB2_M" [color="0 0 0.67761627907", label="0.37",style=bold];
"DEFB119_S" -> "GABRP_M" [color="0 0 0.675", label="0.37",style=bold];
"MMP1_S" -> "GABRP_M" [color="0 0 0.67238372093", label="0.37",style=bold];
"MMP1_S" -> "LGSN_E" [color="0 0 0.66976744186", label="0.39",style=bold];
"KRT75_S" -> "SLC6A18_E" [color="0 0 0.667151162791", label="0.41",style=bold];
"KRT75_S" -> "PLA2G4D_M" [color="0 0 0.664534883721", label="0.41",style=bold];
"SLCO6A1_S" -> "LAD1_E" [color="0 0 0.661918604651", label="0.41",style=bold];
"ALDH3B2_S" -> "LAD1_E" [color="0 0 0.659302325581", label="0.41",style=bold];
"ALDH3B2_S" -> "XDH_E" [color="0 0 0.656686046512", label="0.41",style=bold];
"STAB2_S" -> "AFP_M" [color="0 0 0.654069767442", label="0.41",style=bold];
"DEFB119_S" -> "TRPM1_E" [color="0 0 0.651453488372", label="0.42",style=bold];
"CHIA_S" -> "KRT75_E" [color="0 0 0.648837209302", label="0.42",style=bold];
"STAB2_S" -> "KRT75_E" [color="0 0 0.646220930233", label="0.42",style=bold];
"PKHD1L1_S" -> "SLCO6A1_E" [color="0 0 0.643604651163", label="0.43",style=bold];
"GABRP_S" -> "MMP1_M" [color="0 0 0.640988372093", label="0.44",style=bold];
"XDH_S" -> "DMRTB1_E" [color="0 0 0.638372093023", label="0.44",style=bold];
"CHIA_S" -> "DMRTB1_E" [color="0 0 0.635755813953", label="0.44",style=bold];
"CHIA_S" -> "TRPM1_M" [color="0 0 0.633139534884", label="0.44",style=bold];
"PLA2G4D_E" -> "PROKR1_E" [color="0 0 0.630523255814", label="0.45",style=bold];
"CHIA_S" -> "LMX1A_M" [color="0 0 0.627906976744", label="0.47",style=bold];
"KRT75_S" -> "DMRTB1_E" [color="0 0 0.625290697674", label="0.47",style=bold];
"STAB2_S" -> "TRPM1_E" [color="0 0 0.622674418605", label="0.47",style=bold];
"SLC6A18_S" -> "TRPM1_E" [color="0 0 0.620058139535", label="0.47",style=bold];
"PROKR1_S" -> "PKHD1L1_E" [color="0 0 0.617441860465", label="0.49",style=bold];
"ABCG8_S" -> "LMX1A_M" [color="0 0 0.614825581395", label="0.50",style=bold];
"PKHD1L1_S" -> "DEFB119_E" [color="0 0 0.612209302326", label="0.51",style=bold];
"GABRP_S" -> "SLCO6A1_M" [color="0 0 0.609593023256", label="0.51",style=bold];
"MMP1_S" -> "OPN5_M" [color="0 0 0.606976744186", label="0.51",style=bold];
"CHIA_S" -> "PROKR1_M" [color="0 0 0.604360465116", label="0.51",style=bold];
"XDH_S" -> "MMP1_M" [color="0 0 0.601744186047", label="0.51",style=bold];
"CHIA_S" -> "PLA2G4D_E" [color="0 0 0.599127906977", label="0.51",style=bold];
"DEFB119_S" -> "SLCO6A1_M" [color="0 0 0.596511627907", label="0.51",style=bold];
"PROKR1_S" -> "XDH_M" [color="0 0 0.593895348837", label="0.54",style=bold];
"SLCO6A1_S" -> "SLC6A18_M" [color="0 0 0.591279069767", label="0.54",style=bold];
"SLC6A18_S" -> "GABRP_M" [color="0 0 0.588662790698", label="0.54",style=bold];
"SLC6A18_S" -> "PROKR1_M" [color="0 0 0.586046511628", label="0.55",style=bold];
"LGSN_S" -> "PLA2G4D_M" [color="0 0 0.583430232558", label="0.55",style=bold];
"PROKR1_S" -> "PROKR1_M" [color="0 0 0.580813953488", label="0.57",style=bold];
"DMRTB1_S" -> "PKHD1L1_E" [color="0 0 0.578197674419", label="0.57",style=bold];
"STAB2_S" -> "PKHD1L1_E" [color="0 0 0.575581395349", label="0.57",style=bold];
"TMPRSS11F_S" -> "MMP1_E" [color="0 0 0.572965116279", label="0.58",style=bold];
"CHIA_S" -> "OPN5_E" [color="0 0 0.570348837209", label="0.59",style=bold];
"LGSN_S" -> "SLCO6A1_M" [color="0 0 0.56773255814", label="0.61",style=bold];
"TMPRSS11F_S" -> "LMX1A_E" [color="0 0 0.56511627907", label="0.61",style=bold];
"SLC6A18_S" -> "SLCO6A1_M" [color="0 0 0.5625", label="0.61",style=bold];
"SLCO6A1_S" -> "CHIA_M" [color="0 0 0.55988372093", label="0.61",style=bold];
"PROKR1_S" -> "TRPM1_M" [color="0 0 0.55726744186", label="0.61",style=bold];
"TRPM1_S" -> "GABRP_M" [color="0 0 0.554651162791", label="0.62",style=bold];
"STAB2_S" -> "FCRL5_E" [color="0 0 0.552034883721", label="0.62",style=bold];
"PROKR1_S" -> "FCRL5_E" [color="0 0 0.549418604651", label="0.63",style=bold];
"LAD1_S" -> "LAD1_M" [color="0 0 0.546802325581", label="0.64",style=bold];
"TMPRSS11F_S" -> "LAD1_M" [color="0 0 0.544186046512", label="0.64",style=bold];
"SLCO6A1_S" -> "DMRTB1_M" [color="0 0 0.541569767442", label="0.65",style=bold];
"ABCG8_S" -> "OPN5_M" [color="0 0 0.538953488372", label="0.65",style=bold];
"PROKR1_S" -> "OPN5_M" [color="0 0 0.536337209302", label="0.65",style=bold];
"SLCO6A1_S" -> "ALDH3B2_M" [color="0 0 0.533720930233", label="0.69",style=bold];
"KRT75_S" -> "ABCG8_M" [color="0 0 0.531104651163", label="0.69",style=bold];
"STAB2_S" -> "PKHD1L1_M" [color="0 0 0.528488372093", label="0.69",style=bold];
"XDH_S" -> "PROKR1_M" [color="0 0 0.525872093023", label="0.69",style=bold];
"SLC6A18_E" -> "PKHD1L1_E" [color="0 0 0.523255813953", label="0.71",style=bold];
"LGSN_S" -> "PROKR1_E" [color="0 0 0.520639534884", label="0.72",style=bold];
"PKHD1L1_S" -> "LGSN_E" [color="0 0 0.518023255814", label="0.74",style=bold];
"HOXA11_E" -> "DEFB119_M" [color="0 0 0.515406976744", label="0.76",style=bold];
"SLC6A18_E" -> "LGSN_M" [color="0 0 0.512790697674", label="0.79",style=bold];
"PROKR1_S" -> "LGSN_M" [color="0 0 0.510174418605", label="0.80",style=bold];
"PKHD1L1_S" -> "ALDH3B2_M" [color="0 0 0.507558139535", label="0.82",style=bold];
"XDH_S" -> "STAB2_E" [color="0 0 0.504941860465", label="0.85",style=bold];
"SLC6A18_S" -> "LAD1_E" [color="0 0 0.502325581395", label="0.85",style=bold];
"KRT75_S" -> "DEFB119_M" [color="0 0 0.499709302326", label="0.85",style=bold];
"PROKR1_S" -> "ABCG8_E" [color="0 0 0.497093023256", label="0.85",style=bold];
"LAD1_S" -> "STAB2_M" [color="0 0 0.494476744186", label="0.85",style=bold];
"PKHD1L1_S" -> "LMX1A_E" [color="0 0 0.491860465116", label="0.85",style=bold];
"PROKR1_S" -> "PROKR1_E" [color="0 0 0.489244186047", label="0.85",style=bold];
"XDH_S" -> "CHIA_M" [color="0 0 0.486627906977", label="0.85",style=bold];
"PKHD1L1_S" -> "TRPM1_M" [color="0 0 0.484011627907", label="0.86",style=bold];
"PKHD1L1_S" -> "LAD1_M" [color="0 0 0.481395348837", label="0.86",style=bold];
"TRPM1_S" -> "KRT75_E" [color="0 0 0.478779069767", label="0.89",style=bold];
"LAD1_S" -> "LAD1_E" [color="0 0 0.476162790698", label="0.92",style=bold];
"ABCG8_S" -> "MMP1_M" [color="0 0 0.473546511628", label="0.98",style=bold];
"ALDH3B2_S" -> "DMRTB1_M" [color="0 0 0.470930232558", label="0.98",style=bold];
"XDH_S" -> "DMRTB1_M" [color="0 0 0.468313953488", label="0.98",style=bold];
"PROKR1_S" -> "GABRP_E" [color="0 0 0.465697674419", label="0.98",style=bold];
"ABCG8_S" -> "LGSN_M" [color="0 0 0.463081395349", label="0.98",style=bold];
"PKHD1L1_S" -> "AFP_M" [color="0 0 0.460465116279", label="1.02",style=bold];
"TMPRSS11F_S" -> "LMX1A_M" [color="0 0 0.457848837209", label="1.02",style=bold];
"XDH_M" -> "SLCO6A1_M" [color="0 0 0.45523255814", label="1.02",style=bold];
"GABRP_S" -> "ALDH3B2_E" [color="0 0 0.45261627907", label="1.03",style=bold];
"XDH_S" -> "ALDH3B2_E" [color="0 0 0.45", label="1.03",style=bold];
"LGSN_S" -> "CHIA_E" [color="0 0 0.44738372093", label="1.07",style=bold];
"TMPRSS11F_S" -> "SLC6A18_M" [color="0 0 0.44476744186", label="1.09",style=bold];
"LAD1_S" -> "HOXA11_M" [color="0 0 0.442151162791", label="1.10",style=bold];
"STAB2_S" -> "STAB2_E" [color="0 0 0.439534883721", label="1.10",style=bold];
"CHIA_S" -> "TMPRSS11F_M" [color="0 0 0.436918604651", label="1.10",style=bold];
"GABRP_S" -> "LMX1A_E" [color="0 0 0.434302325581", label="1.10",style=bold];
"XDH_S" -> "LAD1_E" [color="0 0 0.431686046512", label="1.10",style=bold];
"GABRP_S" -> "TMPRSS11F_M" [color="0 0 0.429069767442", label="1.10",style=bold];
"STAB2_S" -> "TMPRSS11F_M" [color="0 0 0.426453488372", label="1.10",style=bold];
"PKHD1L1_S" -> "LMX1A_M" [color="0 0 0.423837209302", label="1.11",style=bold];
"TRPM1_S" -> "PLA2G4D_M" [color="0 0 0.421220930233", label="1.15",style=bold];
"XDH_S" -> "FCRL5_M" [color="0 0 0.418604651163", label="1.15",style=bold];
"GABRP_S" -> "LAD1_M" [color="0 0 0.415988372093", label="1.15",style=bold];
"PROKR1_S" -> "KRT75_M" [color="0 0 0.413372093023", label="1.20",style=bold];
"CHIA_S" -> "OPN5_M" [color="0 0 0.410755813953", label="1.20",style=bold];
"PKHD1L1_S" -> "LGSN_M" [color="0 0 0.408139534884", label="1.20",style=bold];
"DEFB119_S" -> "AFP_M" [color="0 0 0.405523255814", label="1.22",style=bold];
"SLCO6A1_S" -> "AFP_M" [color="0 0 0.402906976744", label="1.22",style=bold];
"XDH_S" -> "LGSN_E" [color="0 0 0.400290697674", label="1.22",style=bold];
"TRPM1_S" -> "CHIA_M" [color="0 0 0.397674418605", label="1.23",style=bold];
"STAB2_S" -> "CHIA_E" [color="0 0 0.395058139535", label="1.25",style=bold];
"XDH_S" -> "CHIA_E" [color="0 0 0.392441860465", label="1.25",style=bold];
"SLCO6A1_S" -> "CHIA_E" [color="0 0 0.389825581395", label="1.25",style=bold];
"CHIA_S" -> "LAD1_E" [color="0 0 0.387209302326", label="1.25",style=bold];
"DMRTB1_M" -> "TMPRSS11F_M" [color="0 0 0.384593023256", label="1.26",style=bold];
"XDH_M" -> "MMP1_M" [color="0 0 0.381976744186", label="1.27",style=bold];
"OPN5_S" -> "CHIA_M" [color="0 0 0.379360465116", label="1.30",style=bold];
"FCRL5_S" -> "PKHD1L1_M" [color="0 0 0.376744186047", label="1.31",style=bold];
"DMRTB1_S" -> "TMPRSS11F_E" [color="0 0 0.374127906977", label="1.32",style=bold];
"LGSN_S" -> "LAD1_M" [color="0 0 0.371511627907", label="1.33",style=bold];
"FCRL5_S" -> "LMX1A_E" [color="0 0 0.368895348837", label="1.39",style=bold];
"CHIA_S" -> "LAD1_M" [color="0 0 0.366279069767", label="1.39",style=bold];
"SLC6A18_M" -> "ABCG8_M" [color="0 0 0.363662790698", label="1.40",style=bold];
"XDH_E" -> "PKHD1L1_M" [color="0 0 0.361046511628", label="1.41",style=bold];
"AFP_E" -> "FCRL5_E" [color="0 0 0.358430232558", label="1.46",style=bold];
"LAD1_S" -> "HOXA11_E" [color="0 0 0.355813953488", label="1.47",style=bold];
"SLC6A18_S" -> "HOXA11_M" [color="0 0 0.353197674419", label="1.47",style=bold];
"XDH_S" -> "HOXA11_E" [color="0 0 0.350581395349", label="1.47",style=bold];
"FCRL5_S" -> "PKHD1L1_E" [color="0 0 0.347965116279", label="1.48",style=bold];
"PKHD1L1_S" -> "ALDH3B2_E" [color="0 0 0.345348837209", label="1.48",style=bold];
"TMPRSS11F_S" -> "KRT75_E" [color="0 0 0.34273255814", label="1.48",style=bold];
"DEFB119_S" -> "KRT75_E" [color="0 0 0.34011627907", label="1.49",style=bold];
"MMP1_S" -> "PKHD1L1_M" [color="0 0 0.3375", label="1.54",style=bold];
"FCRL5_S" -> "Survival" [color="0 0 0.33488372093", label="1.55",style=bold];
"TRPM1_E" -> "SLCO6A1_E" [color="0 0 0.33226744186", label="1.60",style=bold];
"DEFB119_S" -> "FCRL5_E" [color="0 0 0.329651162791", label="1.62",style=bold];
"XDH_S" -> "FCRL5_E" [color="0 0 0.327034883721", label="1.62",style=bold];
"PLA2G4D_M" -> "CHIA_M" [color="0 0 0.324418604651", label="1.63",style=bold];
"PROKR1_S" -> "CHIA_E" [color="0 0 0.321802325581", label="1.65",style=bold];
"MMP1_S" -> "TMPRSS11F_E" [color="0 0 0.319186046512", label="1.73",style=bold];
"FCRL5_S" -> "DMRTB1_E" [color="0 0 0.316569767442", label="1.76",style=bold];
"SLCO6A1_E" -> "KRT75_M" [color="0 0 0.313953488372", label="1.79",style=bold];
"SLC6A18_S" -> "HOXA11_E" [color="0 0 0.311337209302", label="1.79",style=bold];
"TMPRSS11F_M" -> "OPN5_E" [color="0 0 0.308720930233", label="1.80",style=bold];
"MMP1_S" -> "SLC6A18_M" [color="0 0 0.306104651163", label="1.81",style=bold];
"PKHD1L1_S" -> "Survival" [color="0 0 0.303488372093", label="1.83",style=bold];
"HOXA11_E" -> "GABRP_E" [color="0 0 0.300872093023", label="1.88",style=bold];
"AFP_E" -> "XDH_E" [color="0 0 0.298255813953", label="1.89",style=bold];
"XDH_E" -> "HOXA11_E" [color="0 0 0.295639534884", label="1.91",style=bold];
"DMRTB1_S" -> "PROKR1_E" [color="0 0 0.293023255814", label="1.95",style=bold];
"LGSN_S" -> "TRPM1_M" [color="0 0 0.290406976744", label="1.96",style=bold];
"SLC6A18_E" -> "PLA2G4D_E" [color="0 0 0.287790697674", label="1.97",style=bold];
"XDH_M" -> "ALDH3B2_M" [color="0 0 0.285174418605", label="2.05",style=bold];
"ABCG8_S" -> "LAD1_E" [color="0 0 0.282558139535", label="2.08",style=bold];
"ALDH3B2_M" -> "LGSN_M" [color="0 0 0.279941860465", label="2.08",style=bold];
"TMPRSS11F_S" -> "XDH_E" [color="0 0 0.277325581395", label="2.16",style=bold];
"TRPM1_E" -> "OPN5_E" [color="0 0 0.274709302326", label="2.25",style=bold];
"XDH_M" -> "LGSN_M" [color="0 0 0.272093023256", label="2.30",style=bold];
"GABRP_M" -> "DEFB119_M" [color="0 0 0.269476744186", label="2.43",style=bold];
"TRPM1_S" -> "OPN5_E" [color="0 0 0.266860465116", label="2.48",style=bold];
"PKHD1L1_S" -> "FCRL5_M" [color="0 0 0.264244186047", label="2.51",style=bold];
"DEFB119_E" -> "CHIA_E" [color="0 0 0.261627906977", label="2.61",style=bold];
"FCRL5_S" -> "CHIA_E" [color="0 0 0.259011627907", label="2.74",style=bold];
"KRT75_E" -> "LMX1A_M" [color="0 0 0.256395348837", label="2.77",style=bold];
"MMP1_S" -> "OPN5_E" [color="0 0 0.253779069767", label="2.77",style=bold];
"TRPM1_S" -> "PROKR1_S" [color="0 0 0.251162790698", label="2.89",style=bold];
"ALDH3B2_M" -> "TMPRSS11F_M" [color="0 0 0.248546511628", label="2.91",style=bold];
"PROKR1_M" -> "TMPRSS11F_E" [color="0 0 0.245930232558", label="2.97",style=bold];
"TRPM1_M" -> "DEFB119_E" [color="0 0 0.243313953488", label="3.00",style=bold];
"TRPM1_E" -> "PROKR1_E" [color="0 0 0.240697674419", label="3.21",style=bold];
"PKHD1L1_S" -> "AFP_E" [color="0 0 0.238081395349", label="3.31",style=bold];
"SLC6A18_S" -> "SLCO6A1_E" [color="0 0 0.235465116279", label="3.37",style=bold];
"OPN5_E" -> "LGSN_M" [color="0 0 0.232848837209", label="3.37",style=bold];
"PROKR1_M" -> "FCRL5_M" [color="0 0 0.23023255814", label="3.72",style=bold];
"KRT75_E" -> "OPN5_M" [color="0 0 0.22761627907", label="3.76",style=bold];
"DMRTB1_M" -> "STAB2_E" [color="0 0 0.225", label="3.82",style=bold];
"PROKR1_M" -> "DMRTB1_E" [color="0 0 0.22238372093", label="3.89",style=bold];
"ALDH3B2_E" -> "ABCG8_E" [color="0 0 0.21976744186", label="3.92",style=bold];
"SLCO6A1_E" -> "LGSN_M" [color="0 0 0.217151162791", label="3.98",style=bold];
"LAD1_S" -> "FCRL5_S" [color="0 0 0.214534883721", label="3.98",style=bold];
"ALDH3B2_S" -> "LGSN_S" [color="0 0 0.211918604651", label="3.98",style=bold];
"ALDH3B2_S" -> "TMPRSS11F_S" [color="0 0 0.209302325581", label="3.98",style=bold];
"LAD1_M" -> "PKHD1L1_M" [color="0 0 0.206686046512", label="4.18",style=bold];
"DMRTB1_M" -> "SLCO6A1_M" [color="0 0 0.204069767442", label="4.22",style=bold];
"TRPM1_E" -> "LMX1A_E" [color="0 0 0.201453488372", label="4.32",style=bold];
"SLC6A18_E" -> "PLA2G4D_M" [color="0 0 0.198837209302", label="4.37",style=bold];
"STAB2_E" -> "LMX1A_E" [color="0 0 0.196220930233", label="4.65",style=bold];
"OPN5_S" -> "DEFB119_S" [color="0 0 0.193604651163", label="4.67",style=bold];
"DMRTB1_S" -> "CHIA_S" [color="0 0 0.190988372093", label="4.67",style=bold];
"SLC6A18_E" -> "PROKR1_E" [color="0 0 0.188372093023", label="4.77",style=bold];
"SLC6A18_E" -> "DEFB119_M" [color="0 0 0.185755813953", label="4.77",style=bold];
"ABCG8_M" -> "SLCO6A1_M" [color="0 0 0.183139534884", label="4.80",style=bold];
"ABCG8_E" -> "TRPM1_E" [color="0 0 0.180523255814", label="4.81",style=bold];
"ABCG8_M" -> "STAB2_M" [color="0 0 0.177906976744", label="4.93",style=bold];
"AFP_E" -> "DEFB119_M" [color="0 0 0.175290697674", label="5.17",style=bold];
"LAD1_E" -> "STAB2_E" [color="0 0 0.172674418605", label="5.22",style=bold];
"LAD1_M" -> "STAB2_M" [color="0 0 0.170058139535", label="5.44",style=bold];
"SLC6A18_M" -> "KRT75_E" [color="0 0 0.167441860465", label="5.53",style=bold];
"OPN5_M" -> "GABRP_E" [color="0 0 0.164825581395", label="6.16",style=bold];
"LMX1A_M" -> "DEFB119_M" [color="0 0 0.162209302326", label="6.16",style=bold];
"LAD1_M" -> "SLCO6A1_M" [color="0 0 0.159593023256", label="6.44",style=bold];
"DEFB119_E" -> "TMPRSS11F_E" [color="0 0 0.156976744186", label="6.46",style=bold];
"OPN5_E" -> "STAB2_E" [color="0 0 0.154360465116", label="6.66",style=bold];
"ABCG8_M" -> "TMPRSS11F_M" [color="0 0 0.151744186047", label="6.81",style=bold];
"DEFB119_E" -> "SLCO6A1_E" [color="0 0 0.149127906977", label="6.83",style=bold];
"AFP_E" -> "PLA2G4D_E" [color="0 0 0.146511627907", label="6.89",style=bold];
"TMPRSS11F_E" -> "PLA2G4D_E" [color="0 0 0.143895348837", label="6.90",style=bold];
"CHIA_M" -> "MMP1_M" [color="0 0 0.141279069767", label="6.92",style=bold];
"TMPRSS11F_M" -> "Survival" [color="0 0 0.138662790698", label="6.92",style=bold];
"LAD1_E" -> "GABRP_E" [color="0 0 0.136046511628", label="7.10",style=bold];
"DEFB119_E" -> "PLA2G4D_E" [color="0 0 0.133430232558", label="7.29",style=bold];
"FCRL5_M" -> "STAB2_M" [color="0 0 0.130813953488", label="7.30",style=bold];
"TMPRSS11F_M" -> "LAD1_E" [color="0 0 0.128197674419", label="7.70",style=bold];
"DEFB119_E" -> "HOXA11_E" [color="0 0 0.125581395349", label="7.75",style=bold];
"LAD1_E" -> "ALDH3B2_E" [color="0 0 0.122965116279", label="7.79",style=bold];
"PROKR1_M" -> "ABCG8_M" [color="0 0 0.120348837209", label="7.82",style=bold];
"LAD1_M" -> "DMRTB1_M" [color="0 0 0.11773255814", label="8.04",style=bold];
"KRT75_E" -> "GABRP_E" [color="0 0 0.11511627907", label="8.51",style=bold];
"TRPM1_M" -> "GABRP_M" [color="0 0 0.1125", label="8.55",style=bold];
"MMP1_M" -> "LGSN_M" [color="0 0 0.10988372093", label="8.56",style=bold];
"KRT75_E" -> "DEFB119_M" [color="0 0 0.10726744186", label="8.57",style=bold];
"OPN5_E" -> "DEFB119_M" [color="0 0 0.104651162791", label="8.79",style=bold];
"TRPM1_M" -> "LAD1_M" [color="0 0 0.102034883721", label="8.99",style=bold];
"SLC6A18_M" -> "HOXA11_E" [color="0 0 0.0994186046512", label="9.37",style=bold];
"PKHD1L1_M" -> "PLA2G4D_E" [color="0 0 0.0968023255814", label="9.47",style=bold];
"DMRTB1_M" -> "SLC6A18_E" [color="0 0 0.0941860465116", label="10.04",style=bold];
"MMP1_E" -> "GABRP_E" [color="0 0 0.0915697674419", label="10.22",style=bold];
"HOXA11_E" -> "CHIA_M" [color="0 0 0.0889534883721", label="10.54",style=bold];
"PLA2G4D_M" -> "OPN5_M" [color="0 0 0.0863372093023", label="10.96",style=bold];
"PLA2G4D_M" -> "ABCG8_M" [color="0 0 0.0837209302326", label="10.97",style=bold];
"LAD1_M" -> "AFP_E" [color="0 0 0.0811046511628", label="10.98",style=bold];
"PLA2G4D_M" -> "LAD1_E" [color="0 0 0.078488372093", label="11.14",style=bold];
"PROKR1_M" -> "PLA2G4D_M" [color="0 0 0.0758720930233", label="11.19",style=bold];
"TRPM1_M" -> "LGSN_E" [color="0 0 0.0732558139535", label="11.85",style=bold];
"SLC6A18_E" -> "DEFB119_E" [color="0 0 0.0706395348837", label="12.53",style=bold];
"SLC6A18_M" -> "OPN5_M" [color="0 0 0.068023255814", label="12.64",style=bold];
"DEFB119_E" -> "LMX1A_E" [color="0 0 0.0654069767442", label="12.80",style=bold];
"MMP1_E" -> "LAD1_E" [color="0 0 0.0627906976744", label="12.98",style=bold];
"PROKR1_M" -> "KRT75_M" [color="0 0 0.0601744186047", label="13.27",style=bold];
"ALDH3B2_M" -> "TRPM1_M" [color="0 0 0.0575581395349", label="13.31",style=bold];
"TRPM1_M" -> "STAB2_E" [color="0 0 0.0549418604651", label="13.44",style=bold];
"HOXA11_E" -> "HOXA11_M" [color="0 0 0.0523255813953", label="13.47",style=bold];
"STAB2_M" -> "LMX1A_E" [color="0 0 0.0497093023256", label="14.23",style=bold];
"DMRTB1_M" -> "MMP1_M" [color="0 0 0.0470930232558", label="14.51",style=bold];
"TRPM1_M" -> "PKHD1L1_M" [color="0 0 0.044476744186", label="14.82",style=bold];
"TMPRSS11F_M" -> "CHIA_M" [color="0 0 0.0418604651163", label="14.97",style=bold];
"DMRTB1_M" -> "GABRP_M" [color="0 0 0.0392441860465", label="15.53",style=bold];
"HOXA11_E" -> "ABCG8_E" [color="0 0 0.0366279069767", label="17.48",style=bold];
"ABCG8_M" -> "KRT75_M" [color="0 0 0.034011627907", label="18.25",style=bold];
"PLA2G4D_E" -> "ABCG8_E" [color="0 0 0.0313953488372", label="19.77",style=bold];
"LMX1A_M" -> "HOXA11_M" [color="0 0 0.0287790697674", label="20.24",style=bold];
"LMX1A_M" -> "TRPM1_M" [color="0 0 0.0261627906977", label="20.66",style=bold];
"SLC6A18_M" -> "PROKR1_M" [color="0 0 0.0235465116279", label="20.95",style=bold];
"FCRL5_M" -> "ALDH3B2_M" [color="0 0 0.0209302325581", label="21.58",style=bold];
"TRPM1_M" -> "PLA2G4D_M" [color="0 0 0.0183139534884", label="21.59",style=bold];
"SLC6A18_M" -> "LMX1A_M" [color="0 0 0.0156976744186", label="23.02",style=bold];
"LMX1A_M" -> "PROKR1_M" [color="0 0 0.0130813953488", label="25.98",style=bold];
"XDH_M" -> "DMRTB1_M" [color="0 0 0.0104651162791", label="26.52",style=bold];
"LMX1A_M" -> "LAD1_M" [color="0 0 0.0078488372093", label="30.06",style=bold];
"SLC6A18_M" -> "FCRL5_M" [color="0 0 0.00523255813953", label="36.80",style=bold];
"TMPRSS11F_M" -> "AFP_M" [color="0 0 0.00261627906977", label="42.26",style=bold];
"XDH_M" -> "SLC6A18_M" [color="0 0 0.0", label="49.55",style=bold];
}
```
