## Supplementary material for "New analysis framework incorporating mixed mutual information and scalable Bayesian networks for multimodal high dimensional genomic and epigenomic cancer data": Suppl Table 9

```
digraph G{
 ratio=fill;
 node [shape=box, style=rounded];
 edge [arrowhead=none];
"ASB4_S";
"ASB4_E";
"ASB4_M";
"ASB5_S";
"ASB5_E";
"ASB5_M";
"C6_S";
"C6_E";
"C6_M";
"CACNA1S_S";
"CACNA1S_E";
"CACNA1S_M";
"CCDC129_S";
"CCDC129_E";
"CCDC129_M";
"CDH10_S";
"CDH10_E";
"CDH10_M";
"CHRNA4_S";
"CHRNA4_E";
"CHRNA4_M";
"CHRND_S";
"CHRND_E";
"CHRND_M";
"CNTN5_S";
"CNTN5_E";
"CNTN5_M";
"CNTNAP5_S";
"CNTNAP5_E";
"CNTNAP5_M";
"COL11A1_S";
"COL11A1_E";
"COL11A1_M";
"CSMD1_S";
"CSMD1_E";
"CSMD1_M";
"CSMD3_S";
"CSMD3_E";
"CSMD3_M";
"CSRP3_S";
"CSRP3_E";
"CSRP3_M";
"DUSP27_S";
"DUSP27_E";
"DUSP27_M";
"FBN3_S";
"FBN3_E";
"FBN3_M";
"FLG_S";
"FLG_E";
"FLG_M";
"FLG2_S";
"FLG2_E";
"FLG2_M";
"FREM2_S";
"FREM2_E";
"FREM2_M";
"GABRA2_S";
"GABRA2_E";
"GABRA2_M";
"GRIN2B_S";
"GRIN2B_E";
"GRIN2B_M";
"KRT38_S";
"KRT38_E";
"KRT38_M";
"LCT_S";
"LCT_E";
"LCT_M";
"LRP1B_S";
"LRP1B_E";
"LRP1B_M";
"MPPED1_S";
"MPPED1_E";
"MPPED1_M";
"MUC16_S";
"MUC16_E";
"MUC16_M";
"MUC4_S";
"MUC4_E";
"MUC4_M";
"MUC7_S";
"MUC7_E";
"MUC7_M";
"MYBPC1_S";
"MYBPC1_E";
"MYBPC1_M";
"MYH1_S";
"MYH1_E";
"MYH1_M";
"MYH13_S";
"MYH13_E";
"MYH13_M";
"MYH2_S";
"MYH2_E";
"MYH2_M";
"MYH4_S";
"MYH4_E";
"MYH4_M";
"MYH6_S";
"MYH6_E";
"MYH6_M";
"MYH7_S";
"MYH7_E";
"MYH7_M";
"MYH8_S";
"MYH8_E";
"MYH8_M";
"MYL1_S";
"MYL1_E";
"MYL1_M";
"MYOG_S";
"MYOG_E";
"MYOG_M";
"NEB_S";
"NEB_E";
"NEB_M";
"NLRP11_S";
"NLRP11_E";
"NLRP11_M";
"NRAP_S";
"NRAP_E";
"NRAP_M";
"PAK7_S";
"PAK7_E";
"PAK7_M";
"PCLO_S";
"PCLO_E";
"PCLO_M";
"PKHD1_S";
"PKHD1_E";
"PKHD1_M";
"PKHD1L1_S";
"PKHD1L1_E";
"PKHD1L1_M";
"PLD5_S";
"PLD5_E";
"PLD5_M";
"PPP1R3A_S";
"PPP1R3A_E";
"PPP1R3A_M";
"REG1A_S";
"REG1A_E";
"REG1A_M";
"SALL3_S";
"SALL3_E";
"SALL3_M";
"SIX6_S";
"SIX6_E";
"SIX6_M";
"SLC18A3_S";
"SLC18A3_E";
"SLC18A3_M";
"SLC7A14_S";
"SLC7A14_E";
"SLC7A14_M";
"SMYD1_S";
"SMYD1_E";
"SMYD1_M";
"TEX15_S";
"TEX15_E";
"TEX15_M";
"TMEM132D_S";
"TMEM132D_E";
"TMEM132D_M";
"TRDN_S";
"TRDN_E";
"TRDN_M";
"TTN_S";
"TTN_E";
"TTN_M";
"WDR49_S";
"WDR49_E";
"WDR49_M";
"XIRP2_S";
"XIRP2_E";
"XIRP2_M";
"ZIC1_S";
"ZIC1_E";
"ZIC1_M";
"ZIC4_S";
"ZIC4_E";
"ZIC4_M";
"ZNF280A_S";
"ZNF280A_E";
"ZNF280A_M";
"ZNF536_S";
"ZNF536_E";
"ZNF536_M";
"Tumor_Status";
"PKHD1_S" -> "MYH13_S" [color="0 0 0.9", label="0.00",style=bold];
"DUSP27_S" -> "MYH2_S" [color="0 0 0.8992518703241895", label="0.00",style=bold];
"ASB4_S" -> "MYBPC1_M" [color="0 0 0.8985037406483791", label="0.00",style=bold];
"LCT_S" -> "LCT_M" [color="0 0 0.8977556109725686", label="0.01",style=bold];
"CSRP3_S" -> "CSRP3_M" [color="0 0 0.8970074812967581", label="0.01",style=bold];
"NRAP_S" -> "SLC18A3_E" [color="0 0 0.8962593516209476", label="0.01",style=bold];
"NRAP_S" -> "ZNF280A_E" [color="0 0 0.8955112219451372", label="0.02",style=bold];
"ASB5_S" -> "DUSP27_M" [color="0 0 0.8947630922693267", label="0.02",style=bold];
"TEX15_S" -> "WDR49_E" [color="0 0 0.8940149625935162", label="0.02",style=bold];
"MUC7_S" -> "PAK7_E" [color="0 0 0.8932668329177058", label="0.02",style=bold];
"ASB5_S" -> "PKHD1_E" [color="0 0 0.8925187032418953", label="0.02",style=bold];
"WDR49_S" -> "PPP1R3A_E" [color="0 0 0.8917705735660848", label="0.02",style=bold];
"MYH2_S" -> "MYH7_M" [color="0 0 0.8910224438902743", label="0.02",style=bold];
"CHRND_S" -> "PLD5_E" [color="0 0 0.8902743142144639", label="0.03",style=bold];
"MYBPC1_S" -> "CCDC129_E" [color="0 0 0.8895261845386534", label="0.03",style=bold];
"MPPED1_S" -> "MYH8_E" [color="0 0 0.8887780548628429", label="0.03",style=bold];
"SLC7A14_S" -> "SALL3_M" [color="0 0 0.8880299251870324", label="0.03",style=bold];
"MYL1_S" -> "NRAP_E" [color="0 0 0.887281795511222", label="0.03",style=bold];
"MYL1_S" -> "MYBPC1_M" [color="0 0 0.8865336658354115", label="0.03",style=bold];
"PLD5_S" -> "ASB4_E" [color="0 0 0.885785536159601", label="0.04",style=bold];
"MPPED1_S" -> "SMYD1_E" [color="0 0 0.8850374064837906", label="0.04",style=bold];
"MYL1_S" -> "CACNA1S_M" [color="0 0 0.8842892768079801", label="0.04",style=bold];
"ZNF280A_S" -> "MUC4_M" [color="0 0 0.8835411471321696", label="0.04",style=bold];
"PLD5_S" -> "CDH10_S" [color="0 0 0.8827930174563591", label="0.04",style=bold];
"SIX6_S" -> "REG1A_E" [color="0 0 0.8820448877805487", label="0.04",style=bold];
"PAK7_S" -> "PCLO_E" [color="0 0 0.8812967581047382", label="0.04",style=bold];
"CHRNA4_S" -> "MYL1_E" [color="0 0 0.8805486284289277", label="0.04",style=bold];
"CSRP3_S" -> "PKHD1_E" [color="0 0 0.8798004987531173", label="0.05",style=bold];
"ASB4_S" -> "FLG_M" [color="0 0 0.8790523690773068", label="0.05",style=bold];
"SLC18A3_S" -> "PKHD1L1_M" [color="0 0 0.8783042394014963", label="0.05",style=bold];
"MYBPC1_S" -> "KRT38_E" [color="0 0 0.8775561097256858", label="0.05",style=bold];
"CSRP3_S" -> "LRP1B_E" [color="0 0 0.8768079800498754", label="0.06",style=bold];
"SMYD1_S" -> "CSRP3_M" [color="0 0 0.8760598503740649", label="0.06",style=bold];
"MYH1_S" -> "CSRP3_E" [color="0 0 0.8753117206982544", label="0.06",style=bold];
"MPPED1_S" -> "MUC16_E" [color="0 0 0.874563591022444", label="0.06",style=bold];
"SMYD1_S" -> "ASB5_M" [color="0 0 0.8738154613466335", label="0.06",style=bold];
"CSRP3_S" -> "MYOG_E" [color="0 0 0.873067331670823", label="0.07",style=bold];
"CCDC129_S" -> "GRIN2B_E" [color="0 0 0.8723192019950124", label="0.07",style=bold];
"CACNA1S_S" -> "SALL3_E" [color="0 0 0.8715710723192021", label="0.07",style=bold];
"SIX6_S" -> "WDR49_E" [color="0 0 0.8708229426433916", label="0.07",style=bold];
"MYOG_S" -> "ZNF536_M" [color="0 0 0.870074812967581", label="0.07",style=bold];
"PAK7_S" -> "MYH13_E" [color="0 0 0.8693266832917705", label="0.08",style=bold];
"CHRNA4_S" -> "CSMD3_E" [color="0 0 0.8685785536159601", label="0.08",style=bold];
"MYL1_S" -> "CHRND_M" [color="0 0 0.8678304239401496", label="0.08",style=bold];
"MYOG_S" -> "SMYD1_M" [color="0 0 0.8670822942643391", label="0.08",style=bold];
"SALL3_S" -> "WDR49_E" [color="0 0 0.8663341645885287", label="0.08",style=bold];
"MYH1_S" -> "FBN3_E" [color="0 0 0.8655860349127182", label="0.09",style=bold];
"SIX6_S" -> "NRAP_E" [color="0 0 0.8648379052369077", label="0.09",style=bold];
"SMYD1_S" -> "CDH10_M" [color="0 0 0.8640897755610972", label="0.10",style=bold];
"FBN3_S" -> "TMEM132D_S" [color="0 0 0.8633416458852868", label="0.10",style=bold];
"FLG2_S" -> "C6_M" [color="0 0 0.8625935162094763", label="0.10",style=bold];
"CHRND_S" -> "NLRP11_M" [color="0 0 0.8618453865336658", label="0.10",style=bold];
"ZNF280A_S" -> "ZNF536_E" [color="0 0 0.8610972568578554", label="0.10",style=bold];
"MYBPC1_S" -> "MUC7_E" [color="0 0 0.8603491271820449", label="0.10",style=bold];
"CACNA1S_S" -> "ZIC1_E" [color="0 0 0.8596009975062344", label="0.11",style=bold];
"GABRA2_S" -> "SLC7A14_M" [color="0 0 0.8588528678304239", label="0.11",style=bold];
"NRAP_S" -> "FREM2_M" [color="0 0 0.8581047381546135", label="0.11",style=bold];
"ZIC4_S" -> "GRIN2B_S" [color="0 0 0.857356608478803", label="0.11",style=bold];
"SIX6_S" -> "LCT_E" [color="0 0 0.8566084788029925", label="0.11",style=bold];
"SIX6_S" -> "CSMD3_E" [color="0 0 0.8558603491271821", label="0.12",style=bold];
"CACNA1S_S" -> "CSMD3_S" [color="0 0 0.8551122194513716", label="0.12",style=bold];
"LCT_S" -> "CHRNA4_M" [color="0 0 0.8543640897755611", label="0.12",style=bold];
"MUC7_S" -> "CSRP3_E" [color="0 0 0.8536159600997506", label="0.12",style=bold];
"MUC7_S" -> "MYOG_M" [color="0 0 0.8528678304239402", label="0.12",style=bold];
"ASB5_S" -> "CSMD1_E" [color="0 0 0.8521197007481297", label="0.13",style=bold];
"TEX15_S" -> "SLC7A14_S" [color="0 0 0.8513715710723192", label="0.14",style=bold];
"CNTN5_S" -> "ZIC4_M" [color="0 0 0.8506234413965088", label="0.14",style=bold];
"SIX6_S" -> "ASB4_M" [color="0 0 0.8498753117206983", label="0.14",style=bold];
"CSRP3_S" -> "ZNF536_M" [color="0 0 0.8491271820448878", label="0.14",style=bold];
"ZNF280A_S" -> "COL11A1_E" [color="0 0 0.8483790523690773", label="0.14",style=bold];
"SIX6_S" -> "CNTN5_M" [color="0 0 0.8476309226932669", label="0.14",style=bold];
"FBN3_S" -> "GABRA2_M" [color="0 0 0.8468827930174564", label="0.15",style=bold];
"LCT_S" -> "PCLO_S" [color="0 0 0.8461346633416459", label="0.15",style=bold];
"TRDN_S" -> "MYH4_S" [color="0 0 0.8453865336658354", label="0.15",style=bold];
"NRAP_S" -> "GABRA2_S" [color="0 0 0.844638403990025", label="0.15",style=bold];
"MYL1_S" -> "COL11A1_E" [color="0 0 0.8438902743142145", label="0.15",style=bold];
"MYL1_S" -> "XIRP2_S" [color="0 0 0.843142144638404", label="0.15",style=bold];
"CHRNA4_S" -> "PKHD1L1_M" [color="0 0 0.8423940149625936", label="0.15",style=bold];
"CSRP3_S" -> "COL11A1_E" [color="0 0 0.8416458852867831", label="0.16",style=bold];
"CACNA1S_S" -> "MUC4_S" [color="0 0 0.8408977556109726", label="0.16",style=bold];
"WDR49_S" -> "ZNF536_E" [color="0 0 0.8401496259351621", label="0.16",style=bold];
"CHRND_S" -> "FBN3_M" [color="0 0 0.8394014962593517", label="0.16",style=bold];
"DUSP27_S" -> "ASB4_M" [color="0 0 0.8386533665835412", label="0.16",style=bold];
"ASB5_S" -> "SALL3_M" [color="0 0 0.8379052369077307", label="0.17",style=bold];
"CHRNA4_S" -> "FBN3_M" [color="0 0 0.8371571072319202", label="0.17",style=bold];
"TEX15_S" -> "COL11A1_E" [color="0 0 0.8364089775561098", label="0.18",style=bold];
"SIX6_S" -> "TMEM132D_M" [color="0 0 0.8356608478802993", label="0.18",style=bold];
"ASB5_S" -> "NEB_S" [color="0 0 0.8349127182044888", label="0.18",style=bold];
"CHRNA4_S" -> "SIX6_M" [color="0 0 0.8341645885286784", label="0.19",style=bold];
"PLD5_S" -> "GABRA2_E" [color="0 0 0.8334164588528679", label="0.19",style=bold];
"WDR49_S" -> "ZNF280A_E" [color="0 0 0.8326683291770574", label="0.19",style=bold];
"CCDC129_S" -> "FLG_M" [color="0 0 0.8319201995012468", label="0.19",style=bold];
"ASB5_S" -> "MYH1_S" [color="0 0 0.8311720698254365", label="0.19",style=bold];
"ZNF280A_S" -> "PLD5_E" [color="0 0 0.830423940149626", label="0.20",style=bold];
"MYL1_S" -> "SALL3_E" [color="0 0 0.8296758104738154", label="0.20",style=bold];
"MYL1_S" -> "TTN_M" [color="0 0 0.828927680798005", label="0.20",style=bold];
"CNTN5_S" -> "DUSP27_S" [color="0 0 0.8281795511221945", label="0.20",style=bold];
"GABRA2_S" -> "GRIN2B_E" [color="0 0 0.827431421446384", label="0.20",style=bold];
"MYH2_S" -> "C6_E" [color="0 0 0.8266832917705735", label="0.21",style=bold];
"SIX6_S" -> "MPPED1_M" [color="0 0 0.8259351620947631", label="0.21",style=bold];
"TEX15_S" -> "ZNF280A_E" [color="0 0 0.8251870324189526", label="0.21",style=bold];
"DUSP27_S" -> "REG1A_M" [color="0 0 0.8244389027431421", label="0.21",style=bold];
"NRAP_E" -> "MYH2_E" [color="0 0 0.8236907730673317", label="0.21",style=bold];
"CNTN5_S" -> "FLG_M" [color="0 0 0.8229426433915212", label="0.21",style=bold];
"SIX6_S" -> "PKHD1_M" [color="0 0 0.8221945137157107", label="0.21",style=bold];
"SMYD1_S" -> "ASB4_E" [color="0 0 0.8214463840399002", label="0.21",style=bold];
"ASB4_S" -> "MUC7_E" [color="0 0 0.8206982543640898", label="0.21",style=bold];
"ASB4_S" -> "KRT38_M" [color="0 0 0.8199501246882793", label="0.22",style=bold];
"MYOG_S" -> "CCDC129_E" [color="0 0 0.8192019950124688", label="0.22",style=bold];
"MYOG_S" -> "CHRNA4_E" [color="0 0 0.8184538653366584", label="0.22",style=bold];
"CHRNA4_S" -> "CHRNA4_E" [color="0 0 0.8177057356608479", label="0.22",style=bold];
"MYOG_S" -> "MYH7_S" [color="0 0 0.8169576059850374", label="0.22",style=bold];
"ZIC4_S" -> "NLRP11_E" [color="0 0 0.8162094763092269", label="0.22",style=bold];
"CHRND_S" -> "MYH13_S" [color="0 0 0.8154613466334165", label="0.24",style=bold];
"CCDC129_S" -> "PLD5_S" [color="0 0 0.814713216957606", label="0.24",style=bold];
"SLC18A3_S" -> "FBN3_E" [color="0 0 0.8139650872817955", label="0.25",style=bold];
"SLC18A3_S" -> "PAK7_M" [color="0 0 0.8132169576059851", label="0.25",style=bold];
"CHRNA4_S" -> "SALL3_E" [color="0 0 0.8124688279301746", label="0.25",style=bold];
"CHRND_S" -> "ZNF536_E" [color="0 0 0.8117206982543641", label="0.26",style=bold];
"MYBPC1_S" -> "SIX6_M" [color="0 0 0.8109725685785536", label="0.26",style=bold];
"CSRP3_S" -> "DUSP27_M" [color="0 0 0.8102244389027432", label="0.26",style=bold];
"FLG2_S" -> "FLG2_E" [color="0 0 0.8094763092269327", label="0.26",style=bold];
"TRDN_S" -> "LRP1B_S" [color="0 0 0.8087281795511222", label="0.26",style=bold];
"MYH1_S" -> "CDH10_E" [color="0 0 0.8079800498753118", label="0.26",style=bold];
"WDR49_S" -> "COL11A1_M" [color="0 0 0.8072319201995013", label="0.27",style=bold];
"FLG2_S" -> "ASB5_M" [color="0 0 0.8064837905236908", label="0.27",style=bold];
"MPPED1_S" -> "XIRP2_S" [color="0 0 0.8057356608478803", label="0.27",style=bold];
"SIX6_S" -> "CHRND_E" [color="0 0 0.8049875311720699", label="0.27",style=bold];
"REG1A_S" -> "CNTN5_E" [color="0 0 0.8042394014962594", label="0.28",style=bold];
"NLRP11_S" -> "KRT38_M" [color="0 0 0.8034912718204489", label="0.28",style=bold];
"CHRNA4_S" -> "NLRP11_M" [color="0 0 0.8027431421446384", label="0.29",style=bold];
"SIX6_S" -> "KRT38_M" [color="0 0 0.801995012468828", label="0.29",style=bold];
"MYL1_S" -> "COL11A1_M" [color="0 0 0.8012468827930175", label="0.29",style=bold];
"MYOG_S" -> "KRT38_M" [color="0 0 0.800498753117207", label="0.29",style=bold];
"MYL1_S" -> "XIRP2_E" [color="0 0 0.7997506234413965", label="0.29",style=bold];
"MYOG_S" -> "MYH7_E" [color="0 0 0.7990024937655861", label="0.29",style=bold];
"MPPED1_S" -> "PKHD1L1_M" [color="0 0 0.7982543640897756", label="0.29",style=bold];
"MYOG_S" -> "MYBPC1_M" [color="0 0 0.7975062344139651", label="0.29",style=bold];
"CHRNA4_S" -> "XIRP2_M" [color="0 0 0.7967581047381547", label="0.29",style=bold];
"MYL1_S" -> "PPP1R3A_M" [color="0 0 0.7960099750623442", label="0.29",style=bold];
"KRT38_S" -> "MYL1_M" [color="0 0 0.7952618453865337", label="0.29",style=bold];
"FLG2_S" -> "MUC4_S" [color="0 0 0.7945137157107232", label="0.29",style=bold];
"WDR49_S" -> "MYH1_M" [color="0 0 0.7937655860349128", label="0.29",style=bold];
"SIX6_S" -> "NLRP11_E" [color="0 0 0.7930174563591023", label="0.29",style=bold];
"ZNF280A_S" -> "LRP1B_S" [color="0 0 0.7922693266832918", label="0.29",style=bold];
"SMYD1_S" -> "LCT_E" [color="0 0 0.7915211970074814", label="0.29",style=bold];
"GABRA2_S" -> "CDH10_M" [color="0 0 0.7907730673316709", label="0.29",style=bold];
"SLC18A3_S" -> "C6_M" [color="0 0 0.7900249376558603", label="0.29",style=bold];
"LCT_S" -> "PPP1R3A_S" [color="0 0 0.7892768079800498", label="0.29",style=bold];
"PPP1R3A_S" -> "TRDN_E" [color="0 0 0.7885286783042394", label="0.29",style=bold];
"SIX6_S" -> "FLG2_M" [color="0 0 0.787780548628429", label="0.31",style=bold];
"ASB4_S" -> "FLG2_M" [color="0 0 0.7870324189526184", label="0.31",style=bold];
"SALL3_S" -> "C6_S" [color="0 0 0.786284289276808", label="0.31",style=bold];
"LCT_S" -> "CNTNAP5_S" [color="0 0 0.7855361596009975", label="0.31",style=bold];
"SLC18A3_S" -> "MUC7_E" [color="0 0 0.784788029925187", label="0.32",style=bold];
"CNTN5_S" -> "XIRP2_S" [color="0 0 0.7840399002493765", label="0.32",style=bold];
"SLC7A14_S" -> "KRT38_E" [color="0 0 0.7832917705735661", label="0.32",style=bold];
"PPP1R3A_S" -> "PLD5_M" [color="0 0 0.7825436408977556", label="0.33",style=bold];
"MUC7_S" -> "MYH6_E" [color="0 0 0.7817955112219451", label="0.33",style=bold];
"SLC7A14_S" -> "GABRA2_E" [color="0 0 0.7810473815461347", label="0.33",style=bold];
"SIX6_S" -> "MYOG_E" [color="0 0 0.7802992518703242", label="0.33",style=bold];
"GRIN2B_S" -> "GRIN2B_E" [color="0 0 0.7795511221945137", label="0.33",style=bold];
"ZNF280A_S" -> "GRIN2B_E" [color="0 0 0.7788029925187032", label="0.34",style=bold];
"PPP1R3A_S" -> "COL11A1_E" [color="0 0 0.7780548628428928", label="0.34",style=bold];
"LCT_S" -> "SMYD1_E" [color="0 0 0.7773067331670823", label="0.35",style=bold];
"ASB4_S" -> "CHRND_E" [color="0 0 0.7765586034912718", label="0.35",style=bold];
"MYH7_S" -> "TEX15_M" [color="0 0 0.7758104738154614", label="0.35",style=bold];
"PLD5_S" -> "NRAP_E" [color="0 0 0.7750623441396509", label="0.36",style=bold];
"TEX15_S" -> "CSMD1_E" [color="0 0 0.7743142144638404", label="0.36",style=bold];
"MYL1_S" -> "TTN_S" [color="0 0 0.7735660847880299", label="0.36",style=bold];
"SIX6_S" -> "DUSP27_M" [color="0 0 0.7728179551122195", label="0.36",style=bold];
"MUC7_S" -> "WDR49_M" [color="0 0 0.772069825436409", label="0.36",style=bold];
"SALL3_S" -> "MYH13_S" [color="0 0 0.7713216957605985", label="0.36",style=bold];
"MYL1_S" -> "SMYD1_M" [color="0 0 0.7705735660847881", label="0.37",style=bold];
"MYL1_S" -> "SLC7A14_M" [color="0 0 0.7698254364089776", label="0.37",style=bold];
"PAK7_S" -> "MYL1_E" [color="0 0 0.7690773067331671", label="0.38",style=bold];
"CCDC129_S" -> "NRAP_S" [color="0 0 0.7683291770573566", label="0.38",style=bold];
"FLG2_S" -> "NRAP_S" [color="0 0 0.7675810473815461", label="0.38",style=bold];
"MYH7_S" -> "MYH13_S" [color="0 0 0.7668329177057357", label="0.38",style=bold];
"MYL1_S" -> "NLRP11_M" [color="0 0 0.7660847880299252", label="0.38",style=bold];
"SIX6_S" -> "SIX6_E" [color="0 0 0.7653366583541148", label="0.39",style=bold];
"SIX6_S" -> "PCLO_E" [color="0 0 0.7645885286783043", label="0.39",style=bold];
"MYL1_S" -> "MYH8_E" [color="0 0 0.7638403990024938", label="0.39",style=bold];
"GABRA2_S" -> "PKHD1_S" [color="0 0 0.7630922693266833", label="0.40",style=bold];
"MYH4_S" -> "MPPED1_M" [color="0 0 0.7623441396508728", label="0.40",style=bold];
"MYOG_S" -> "NEB_M" [color="0 0 0.7615960099750624", label="0.41",style=bold];
"NRAP_S" -> "SLC18A3_M" [color="0 0 0.7608478802992519", label="0.41",style=bold];
"SLC18A3_S" -> "PCLO_S" [color="0 0 0.7600997506234415", label="0.41",style=bold];
"SLC18A3_S" -> "ZNF536_E" [color="0 0 0.759351620947631", label="0.41",style=bold];
"FLG2_S" -> "SLC18A3_M" [color="0 0 0.7586034912718205", label="0.42",style=bold];
"SLC7A14_S" -> "MYBPC1_S" [color="0 0 0.75785536159601", label="0.42",style=bold];
"REG1A_M" -> "FLG_S" [color="0 0 0.7571072319201995", label="0.42",style=bold];
"NRAP_S" -> "CSRP3_M" [color="0 0 0.7563591022443891", label="0.43",style=bold];
"MYL1_S" -> "FLG2_E" [color="0 0 0.7556109725685786", label="0.43",style=bold];
"MYL1_S" -> "C6_E" [color="0 0 0.7548628428927681", label="0.43",style=bold];
"CSRP3_S" -> "MYH1_E" [color="0 0 0.7541147132169577", label="0.43",style=bold];
"SLC18A3_S" -> "CSRP3_M" [color="0 0 0.7533665835411472", label="0.43",style=bold];
"NLRP11_S" -> "MUC16_E" [color="0 0 0.7526184538653367", label="0.43",style=bold];
"CSMD1_S" -> "FLG_S" [color="0 0 0.7518703241895262", label="0.43",style=bold];
"PAK7_S" -> "FREM2_E" [color="0 0 0.7511221945137158", label="0.44",style=bold];
"MYH7_S" -> "PCLO_S" [color="0 0 0.7503740648379053", label="0.44",style=bold];
"WDR49_S" -> "Tumor_Status" [color="0 0 0.7496259351620947", label="0.44",style=bold];
"SIX6_S" -> "C6_M" [color="0 0 0.7488778054862844", label="0.44",style=bold];
"ZNF280A_S" -> "PKHD1L1_M" [color="0 0 0.7481296758104738", label="0.44",style=bold];
"ZNF280A_S" -> "MUC7_E" [color="0 0 0.7473815461346633", label="0.44",style=bold];
"ZNF280A_S" -> "SLC7A14_S" [color="0 0 0.7466334164588528", label="0.44",style=bold];
"PAK7_S" -> "NEB_M" [color="0 0 0.7458852867830424", label="0.45",style=bold];
"SALL3_S" -> "CSMD3_E" [color="0 0 0.7451371571072319", label="0.45",style=bold];
"CCDC129_S" -> "PKHD1_E" [color="0 0 0.7443890274314214", label="0.45",style=bold];
"SALL3_S" -> "FBN3_S" [color="0 0 0.743640897755611", label="0.46",style=bold];
"MUC7_S" -> "TTN_S" [color="0 0 0.7428927680798005", label="0.46",style=bold];
"ZNF280A_S" -> "ZIC4_S" [color="0 0 0.74214463840399", label="0.46",style=bold];
"MYBPC1_S" -> "MYL1_M" [color="0 0 0.7413965087281795", label="0.46",style=bold];
"CSRP3_S" -> "MUC16_M" [color="0 0 0.7406483790523691", label="0.46",style=bold];
"MYH4_S" -> "ZNF536_E" [color="0 0 0.7399002493765586", label="0.46",style=bold];
"PAK7_S" -> "SMYD1_M" [color="0 0 0.7391521197007481", label="0.46",style=bold];
"CSRP3_S" -> "WDR49_M" [color="0 0 0.7384039900249377", label="0.46",style=bold];
"SIX6_S" -> "PKHD1L1_E" [color="0 0 0.7376558603491272", label="0.47",style=bold];
"CHRNA4_S" -> "PKHD1L1_E" [color="0 0 0.7369077306733167", label="0.47",style=bold];
"SALL3_S" -> "CHRND_M" [color="0 0 0.7361596009975062", label="0.47",style=bold];
"CNTNAP5_S" -> "KRT38_E" [color="0 0 0.7354114713216958", label="0.47",style=bold];
"ASB4_S" -> "SIX6_E" [color="0 0 0.7346633416458853", label="0.47",style=bold];
"MYOG_S" -> "MYH6_M" [color="0 0 0.7339152119700748", label="0.47",style=bold];
"ZNF280A_S" -> "ZNF536_S" [color="0 0 0.7331670822942644", label="0.47",style=bold];
"ASB5_S" -> "PAK7_M" [color="0 0 0.7324189526184539", label="0.48",style=bold];
"NRAP_S" -> "SLC7A14_E" [color="0 0 0.7316708229426434", label="0.48",style=bold];
"CCDC129_S" -> "MYH8_S" [color="0 0 0.7309226932668329", label="0.48",style=bold];
"MYBPC1_S" -> "MYH8_S" [color="0 0 0.7301745635910224", label="0.48",style=bold];
"SLC7A14_S" -> "LCT_M" [color="0 0 0.729426433915212", label="0.48",style=bold];
"SMYD1_S" -> "SALL3_E" [color="0 0 0.7286783042394015", label="0.49",style=bold];
"NLRP11_S" -> "ZNF280A_E" [color="0 0 0.7279301745635911", label="0.49",style=bold];
"ZIC1_S" -> "ZNF280A_M" [color="0 0 0.7271820448877806", label="0.49",style=bold];
"MYH7_S" -> "MYH13_E" [color="0 0 0.7264339152119701", label="0.49",style=bold];
"GABRA2_S" -> "NEB_E" [color="0 0 0.7256857855361596", label="0.49",style=bold];
"ZIC1_S" -> "PKHD1L1_S" [color="0 0 0.7249376558603491", label="0.49",style=bold];
"REG1A_S" -> "PKHD1L1_S" [color="0 0 0.7241895261845387", label="0.50",style=bold];
"PAK7_S" -> "WDR49_M" [color="0 0 0.7234413965087282", label="0.50",style=bold];
"ASB5_S" -> "PAK7_E" [color="0 0 0.7226932668329178", label="0.50",style=bold];
"MPPED1_S" -> "TRDN_M" [color="0 0 0.7219451371571073", label="0.51",style=bold];
"CHRND_S" -> "CNTNAP5_M" [color="0 0 0.7211970074812968", label="0.52",style=bold];
"SIX6_S" -> "TTN_M" [color="0 0 0.7204488778054863", label="0.52",style=bold];
"CDH10_S" -> "Tumor_Status" [color="0 0 0.7197007481296758", label="0.52",style=bold];
"WDR49_S" -> "TEX15_E" [color="0 0 0.7189526184538654", label="0.52",style=bold];
"ASB5_S" -> "FLG_S" [color="0 0 0.7182044887780549", label="0.52",style=bold];
"ZIC4_S" -> "MYH8_M" [color="0 0 0.7174563591022445", label="0.52",style=bold];
"TEX15_S" -> "CSRP3_E" [color="0 0 0.716708229426434", label="0.53",style=bold];
"ASB4_S" -> "CCDC129_M" [color="0 0 0.7159600997506235", label="0.53",style=bold];
"CHRND_S" -> "FBN3_E" [color="0 0 0.715211970074813", label="0.53",style=bold];
"CNTNAP5_S" -> "ZNF280A_E" [color="0 0 0.7144638403990025", label="0.54",style=bold];
"TEX15_S" -> "ZIC1_E" [color="0 0 0.7137157107231921", label="0.55",style=bold];
"SALL3_S" -> "MYH7_E" [color="0 0 0.7129675810473816", label="0.55",style=bold];
"MYH8_S" -> "SLC18A3_E" [color="0 0 0.712219451371571", label="0.55",style=bold];
"ZNF280A_S" -> "XIRP2_M" [color="0 0 0.7114713216957607", label="0.55",style=bold];
"PPP1R3A_S" -> "PKHD1L1_M" [color="0 0 0.7107231920199502", label="0.56",style=bold];
"CNTNAP5_M" -> "CSMD1_M" [color="0 0 0.7099750623441397", label="0.56",style=bold];
"MYH7_M" -> "TTN_M" [color="0 0 0.7092269326683291", label="0.56",style=bold];
"CHRND_S" -> "FREM2_E" [color="0 0 0.7084788029925188", label="0.57",style=bold];
"MYH7_S" -> "MYBPC1_M" [color="0 0 0.7077306733167082", label="0.57",style=bold];
"MUC7_S" -> "FLG2_E" [color="0 0 0.7069825436408977", label="0.57",style=bold];
"ZNF280A_S" -> "PAK7_E" [color="0 0 0.7062344139650873", label="0.57",style=bold];
"MPPED1_S" -> "COL11A1_M" [color="0 0 0.7054862842892768", label="0.58",style=bold];
"DUSP27_S" -> "TEX15_E" [color="0 0 0.7047381546134663", label="0.58",style=bold];
"CHRNA4_S" -> "MYH7_E" [color="0 0 0.7039900249376558", label="0.58",style=bold];
"SIX6_S" -> "GABRA2_M" [color="0 0 0.7032418952618454", label="0.58",style=bold];
"SLC18A3_S" -> "TEX15_M" [color="0 0 0.7024937655860349", label="0.58",style=bold];
"MYH4_S" -> "CSMD1_E" [color="0 0 0.7017456359102244", label="0.59",style=bold];
"CACNA1S_S" -> "FBN3_S" [color="0 0 0.700997506234414", label="0.59",style=bold];
"SLC18A3_S" -> "C6_E" [color="0 0 0.7002493765586035", label="0.59",style=bold];
"CACNA1S_S" -> "MPPED1_M" [color="0 0 0.699501246882793", label="0.59",style=bold];
"MYH6_S" -> "ZNF280A_S" [color="0 0 0.6987531172069825", label="0.60",style=bold];
"CSRP3_S" -> "REG1A_E" [color="0 0 0.6980049875311721", label="0.60",style=bold];
"CNTNAP5_S" -> "MPPED1_E" [color="0 0 0.6972568578553616", label="0.60",style=bold];
"DUSP27_S" -> "TTN_S" [color="0 0 0.6965087281795511", label="0.61",style=bold];
"CHRND_S" -> "CSRP3_E" [color="0 0 0.6957605985037407", label="0.61",style=bold];
"DUSP27_S" -> "NEB_M" [color="0 0 0.6950124688279302", label="0.62",style=bold];
"ZIC4_S" -> "PKHD1_S" [color="0 0 0.6942643391521197", label="0.62",style=bold];
"KRT38_S" -> "SLC7A14_E" [color="0 0 0.6935162094763092", label="0.63",style=bold];
"MPPED1_S" -> "GABRA2_E" [color="0 0 0.6927680798004988", label="0.63",style=bold];
"MPPED1_S" -> "MYH1_S" [color="0 0 0.6920199501246883", label="0.63",style=bold];
"CHRND_S" -> "ZNF280A_E" [color="0 0 0.6912718204488778", label="0.63",style=bold];
"MYH6_S" -> "TEX15_E" [color="0 0 0.6905236907730674", label="0.63",style=bold];
"LCT_S" -> "CSRP3_M" [color="0 0 0.6897755610972569", label="0.63",style=bold];
"MUC7_S" -> "REG1A_M" [color="0 0 0.6890274314214464", label="0.63",style=bold];
"SLC7A14_S" -> "SALL3_E" [color="0 0 0.6882793017456359", label="0.64",style=bold];
"ZNF536_S" -> "CNTN5_M" [color="0 0 0.6875311720698254", label="0.64",style=bold];
"CHRND_S" -> "NLRP11_E" [color="0 0 0.686783042394015", label="0.65",style=bold];
"DUSP27_S" -> "WDR49_M" [color="0 0 0.6860349127182045", label="0.65",style=bold];
"ASB4_S" -> "PAK7_E" [color="0 0 0.6852867830423941", label="0.65",style=bold];
"SIX6_S" -> "NLRP11_M" [color="0 0 0.6845386533665836", label="0.66",style=bold];
"PLD5_S" -> "ZIC1_E" [color="0 0 0.6837905236907731", label="0.66",style=bold];
"SIX6_S" -> "SMYD1_M" [color="0 0 0.6830423940149626", label="0.66",style=bold];
"ASB4_S" -> "REG1A_E" [color="0 0 0.6822942643391521", label="0.66",style=bold];
"SMYD1_S" -> "MYH13_E" [color="0 0 0.6815461346633417", label="0.66",style=bold];
"SMYD1_S" -> "MYOG_M" [color="0 0 0.6807980049875312", label="0.66",style=bold];
"FBN3_S" -> "KRT38_M" [color="0 0 0.6800498753117208", label="0.66",style=bold];
"MYL1_S" -> "CSRP3_M" [color="0 0 0.6793017456359103", label="0.66",style=bold];
"ASB5_S" -> "PCLO_E" [color="0 0 0.6785536159600998", label="0.67",style=bold];
"MPPED1_S" -> "LCT_E" [color="0 0 0.6778054862842893", label="0.67",style=bold];
"FREM2_S" -> "CSMD1_M" [color="0 0 0.6770573566084788", label="0.67",style=bold];
"LCT_S" -> "COL11A1_S" [color="0 0 0.6763092269326684", label="0.68",style=bold];
"C6_S" -> "SALL3_M" [color="0 0 0.6755610972568579", label="0.68",style=bold];
"MYH7_S" -> "SLC7A14_M" [color="0 0 0.6748129675810474", label="0.68",style=bold];
"FBN3_S" -> "MYOG_M" [color="0 0 0.674064837905237", label="0.68",style=bold];
"ASB5_S" -> "MUC4_S" [color="0 0 0.6733167082294265", label="0.68",style=bold];
"GRIN2B_S" -> "MYL1_M" [color="0 0 0.672568578553616", label="0.68",style=bold];
"FREM2_S" -> "PCLO_M" [color="0 0 0.6718204488778055", label="0.69",style=bold];
"ZIC4_S" -> "C6_M" [color="0 0 0.6710723192019951", label="0.69",style=bold];
"MYOG_S" -> "GRIN2B_S" [color="0 0 0.6703241895261846", label="0.69",style=bold];
"MYL1_S" -> "MYH2_M" [color="0 0 0.669576059850374", label="0.69",style=bold];
"SIX6_S" -> "FLG_M" [color="0 0 0.6688279301745637", label="0.69",style=bold];
"SIX6_S" -> "MYH6_M" [color="0 0 0.6680798004987532", label="0.70",style=bold];
"FREM2_S" -> "MPPED1_E" [color="0 0 0.6673316708229426", label="0.70",style=bold];
"SMYD1_S" -> "TTN_M" [color="0 0 0.6665835411471321", label="0.71",style=bold];
"TRDN_S" -> "ZIC1_M" [color="0 0 0.6658354114713217", label="0.71",style=bold];
"FBN3_S" -> "LCT_M" [color="0 0 0.6650872817955112", label="0.71",style=bold];
"SIX6_S" -> "ZNF280A_M" [color="0 0 0.6643391521197007", label="0.72",style=bold];
"ZIC4_S" -> "KRT38_E" [color="0 0 0.6635910224438903", label="0.72",style=bold];
"SIX6_S" -> "MYBPC1_M" [color="0 0 0.6628428927680798", label="0.72",style=bold];
"C6_S" -> "SLC18A3_E" [color="0 0 0.6620947630922693", label="0.72",style=bold];
"CHRND_S" -> "TEX15_E" [color="0 0 0.6613466334164588", label="0.73",style=bold];
"ZIC4_S" -> "CSMD3_E" [color="0 0 0.6605985037406484", label="0.73",style=bold];
"CNTNAP5_S" -> "PPP1R3A_M" [color="0 0 0.6598503740648379", label="0.73",style=bold];
"SIX6_S" -> "C6_E" [color="0 0 0.6591022443890274", label="0.73",style=bold];
"MYL1_S" -> "PAK7_M" [color="0 0 0.658354114713217", label="0.73",style=bold];
"CSRP3_S" -> "CSMD3_S" [color="0 0 0.6576059850374065", label="0.74",style=bold];
"KRT38_S" -> "XIRP2_M" [color="0 0 0.656857855361596", label="0.74",style=bold];
"LCT_S" -> "PLD5_S" [color="0 0 0.6561097256857855", label="0.75",style=bold];
"FLG2_S" -> "MUC16_S" [color="0 0 0.6553615960099751", label="0.75",style=bold];
"MYH6_S" -> "CHRNA4_E" [color="0 0 0.6546134663341646", label="0.75",style=bold];
"PLD5_S" -> "CSRP3_E" [color="0 0 0.6538653366583541", label="0.76",style=bold];
"SIX6_S" -> "REG1A_M" [color="0 0 0.6531172069825437", label="0.76",style=bold];
"DUSP27_S" -> "PKHD1L1_M" [color="0 0 0.6523690773067332", label="0.76",style=bold];
"ASB4_S" -> "LRP1B_S" [color="0 0 0.6516209476309227", label="0.76",style=bold];
"CNTNAP5_S" -> "MYH4_S" [color="0 0 0.6508728179551122", label="0.76",style=bold];
"CACNA1S_S" -> "PKHD1_E" [color="0 0 0.6501246882793017", label="0.77",style=bold];
"SALL3_S" -> "TTN_S" [color="0 0 0.6493765586034913", label="0.77",style=bold];
"PCLO_S" -> "CNTNAP5_S" [color="0 0 0.6486284289276808", label="0.77",style=bold];
"FBN3_S" -> "LCT_E" [color="0 0 0.6478802992518704", label="0.77",style=bold];
"DUSP27_S" -> "C6_M" [color="0 0 0.6471321695760599", label="0.77",style=bold];
"PKHD1L1_M" -> "NEB_S" [color="0 0 0.6463840399002494", label="0.78",style=bold];
"MYH1_S" -> "PLD5_E" [color="0 0 0.6456359102244389", label="0.78",style=bold];
"CHRND_S" -> "MYH13_E" [color="0 0 0.6448877805486284", label="0.78",style=bold];
"FBN3_S" -> "SLC7A14_E" [color="0 0 0.644139650872818", label="0.78",style=bold];
"SMYD1_S" -> "CSMD3_E" [color="0 0 0.6433915211970075", label="0.78",style=bold];
"ASB5_S" -> "MUC16_E" [color="0 0 0.6426433915211971", label="0.78",style=bold];
"TRDN_S" -> "CCDC129_E" [color="0 0 0.6418952618453866", label="0.78",style=bold];
"DUSP27_S" -> "FBN3_E" [color="0 0 0.6411471321695761", label="0.79",style=bold];
"CSRP3_S" -> "MUC4_E" [color="0 0 0.6403990024937656", label="0.79",style=bold];
"MYOG_S" -> "MUC16_S" [color="0 0 0.6396508728179551", label="0.79",style=bold];
"MYL1_S" -> "MUC16_S" [color="0 0 0.6389027431421447", label="0.79",style=bold];
"TRDN_S" -> "GRIN2B_S" [color="0 0 0.6381546134663342", label="0.79",style=bold];
"DUSP27_S" -> "PAK7_E" [color="0 0 0.6374064837905238", label="0.79",style=bold];
"TEX15_S" -> "MUC7_M" [color="0 0 0.6366583541147133", label="0.80",style=bold];
"TTN_M" -> "PPP1R3A_M" [color="0 0 0.6359102244389028", label="0.80",style=bold];
"DUSP27_S" -> "COL11A1_M" [color="0 0 0.6351620947630923", label="0.80",style=bold];
"GABRA2_S" -> "ZIC1_E" [color="0 0 0.6344139650872818", label="0.80",style=bold];
"SMYD1_S" -> "ZIC4_M" [color="0 0 0.6336658354114714", label="0.80",style=bold];
"LCT_S" -> "LRP1B_M" [color="0 0 0.6329177057356609", label="0.80",style=bold];
"NRAP_S" -> "REG1A_M" [color="0 0 0.6321695760598505", label="0.80",style=bold];
"ZNF280A_S" -> "MYH4_M" [color="0 0 0.63142144638404", label="0.81",style=bold];
"GRIN2B_S" -> "ZIC4_E" [color="0 0 0.6306733167082295", label="0.81",style=bold];
"LCT_S" -> "CNTN5_E" [color="0 0 0.629925187032419", label="0.81",style=bold];
"MYL1_S" -> "ZNF280A_E" [color="0 0 0.6291770573566084", label="0.81",style=bold];
"SLC18A3_S" -> "PCLO_M" [color="0 0 0.6284289276807979", label="0.82",style=bold];
"ZNF280A_S" -> "CSRP3_E" [color="0 0 0.6276807980049876", label="0.82",style=bold];
"TRDN_S" -> "SLC7A14_E" [color="0 0 0.6269326683291772", label="0.82",style=bold];
"MYH7_S" -> "CCDC129_E" [color="0 0 0.6261845386533667", label="0.83",style=bold];
"ASB5_S" -> "FLG2_E" [color="0 0 0.6254364089775561", label="0.83",style=bold];
"MYH4_S" -> "ZNF280A_E" [color="0 0 0.6246882793017456", label="0.83",style=bold];
"REG1A_S" -> "SLC7A14_E" [color="0 0 0.6239401496259351", label="0.84",style=bold];
"TRDN_S" -> "PAK7_S" [color="0 0 0.6231920199501246", label="0.84",style=bold];
"TEX15_S" -> "MUC4_M" [color="0 0 0.6224438902743142", label="0.84",style=bold];
"PAK7_S" -> "SIX6_E" [color="0 0 0.6216957605985038", label="0.84",style=bold];
"GRIN2B_S" -> "PCLO_M" [color="0 0 0.6209476309226933", label="0.85",style=bold];
"PLD5_S" -> "LRP1B_M" [color="0 0 0.6201995012468828", label="0.85",style=bold];
"PAK7_S" -> "CACNA1S_S" [color="0 0 0.6194513715710723", label="0.85",style=bold];
"MYH7_S" -> "ZNF280A_M" [color="0 0 0.6187032418952618", label="0.85",style=bold];
"SMYD1_S" -> "WDR49_E" [color="0 0 0.6179551122194513", label="0.85",style=bold];
"CNTNAP5_S" -> "CHRNA4_M" [color="0 0 0.6172069825436409", label="0.85",style=bold];
"LCT_S" -> "ZNF280A_E" [color="0 0 0.6164588528678304", label="0.85",style=bold];
"MYH7_S" -> "PKHD1L1_M" [color="0 0 0.61571072319202", label="0.86",style=bold];
"SMYD1_S" -> "CCDC129_M" [color="0 0 0.6149625935162095", label="0.86",style=bold];
"NLRP11_S" -> "MYH4_S" [color="0 0 0.614214463840399", label="0.86",style=bold];
"FREM2_S" -> "LCT_M" [color="0 0 0.6134663341645885", label="0.86",style=bold];
"MUC7_S" -> "FBN3_E" [color="0 0 0.612718204488778", label="0.86",style=bold];
"PAK7_S" -> "CACNA1S_M" [color="0 0 0.6119700748129676", label="0.86",style=bold];
"PAK7_S" -> "PKHD1L1_E" [color="0 0 0.6112219451371571", label="0.86",style=bold];
"MYBPC1_S" -> "ZNF536_E" [color="0 0 0.6104738154613467", label="0.87",style=bold];
"MYH13_S" -> "LRP1B_S" [color="0 0 0.6097256857855362", label="0.87",style=bold];
"SLC7A14_S" -> "FBN3_E" [color="0 0 0.6089775561097257", label="0.87",style=bold];
"MYH2_S" -> "PCLO_E" [color="0 0 0.6082294264339152", label="0.87",style=bold];
"MUC4_S" -> "WDR49_E" [color="0 0 0.6074812967581047", label="0.87",style=bold];
"CNTN5_S" -> "TTN_S" [color="0 0 0.6067331670822943", label="0.88",style=bold];
"SMYD1_S" -> "NRAP_M" [color="0 0 0.6059850374064838", label="0.88",style=bold];
"MUC7_S" -> "XIRP2_S" [color="0 0 0.6052369077306734", label="0.88",style=bold];
"CHRNA4_S" -> "MYH13_S" [color="0 0 0.6044887780548629", label="0.88",style=bold];
"ASB4_S" -> "PKHD1L1_E" [color="0 0 0.6037406483790524", label="0.88",style=bold];
"DUSP27_S" -> "MUC16_E" [color="0 0 0.6029925187032419", label="0.89",style=bold];
"MYH6_S" -> "CSMD1_S" [color="0 0 0.6022443890274314", label="0.89",style=bold];
"CSRP3_S" -> "FREM2_E" [color="0 0 0.601496259351621", label="0.89",style=bold];
"CHRNA4_S" -> "SLC18A3_M" [color="0 0 0.6007481296758105", label="0.89",style=bold];
"NRAP_S" -> "CNTN5_M" [color="0 0 0.6000000000000001", label="0.90",style=bold];
"ZIC4_S" -> "ZNF280A_E" [color="0 0 0.5992518703241896", label="0.90",style=bold];
"ZNF280A_S" -> "CNTNAP5_S" [color="0 0 0.5985037406483791", label="0.90",style=bold];
"MYH4_S" -> "MYH4_M" [color="0 0 0.5977556109725686", label="0.90",style=bold];
"CHRND_S" -> "DUSP27_M" [color="0 0 0.5970074812967581", label="0.90",style=bold];
"MYH7_S" -> "XIRP2_M" [color="0 0 0.5962593516209477", label="0.91",style=bold];
"SMYD1_S" -> "PKHD1L1_E" [color="0 0 0.5955112219451372", label="0.91",style=bold];
"MYL1_S" -> "NEB_M" [color="0 0 0.5947630922693268", label="0.92",style=bold];
"SMYD1_S" -> "PCLO_S" [color="0 0 0.5940149625935163", label="0.92",style=bold];
"CDH10_S" -> "MUC7_E" [color="0 0 0.5932668329177058", label="0.92",style=bold];
"FBN3_S" -> "PAK7_E" [color="0 0 0.5925187032418953", label="0.93",style=bold];
"ASB5_S" -> "PKHD1L1_M" [color="0 0 0.5917705735660848", label="0.93",style=bold];
"MYH13_S" -> "ZNF536_E" [color="0 0 0.5910224438902744", label="0.93",style=bold];
"ZIC1_S" -> "CDH10_S" [color="0 0 0.5902743142144639", label="0.94",style=bold];
"ZIC4_S" -> "REG1A_M" [color="0 0 0.5895261845386535", label="0.94",style=bold];
"MUC4_S" -> "PLD5_E" [color="0 0 0.588778054862843", label="0.94",style=bold];
"REG1A_S" -> "GABRA2_E" [color="0 0 0.5880299251870325", label="0.95",style=bold];
"MYL1_S" -> "SLC18A3_E" [color="0 0 0.587281795511222", label="0.95",style=bold];
"SIX6_S" -> "SLC18A3_E" [color="0 0 0.5865336658354114", label="0.95",style=bold];
"WDR49_S" -> "PLD5_E" [color="0 0 0.5857855361596009", label="0.95",style=bold];
"DUSP27_S" -> "LCT_M" [color="0 0 0.5850374064837905", label="0.96",style=bold];
"SLC18A3_S" -> "CDH10_M" [color="0 0 0.5842892768079802", label="0.96",style=bold];
"SMYD1_S" -> "FREM2_M" [color="0 0 0.5835411471321696", label="0.97",style=bold];
"MUC7_S" -> "ASB4_E" [color="0 0 0.5827930174563591", label="0.97",style=bold];
"WDR49_S" -> "CSMD3_S" [color="0 0 0.5820448877805486", label="0.97",style=bold];
"TEX15_S" -> "SLC7A14_E" [color="0 0 0.5812967581047381", label="0.97",style=bold];
"MYL1_S" -> "MYH1_E" [color="0 0 0.5805486284289276", label="0.97",style=bold];
"SIX6_S" -> "MUC16_S" [color="0 0 0.5798004987531172", label="0.97",style=bold];
"ZNF280A_S" -> "MUC16_S" [color="0 0 0.5790523690773067", label="0.97",style=bold];
"GRIN2B_S" -> "SALL3_M" [color="0 0 0.5783042394014963", label="0.97",style=bold];
"CNTN5_S" -> "FREM2_M" [color="0 0 0.5775561097256858", label="0.98",style=bold];
"SMYD1_S" -> "KRT38_M" [color="0 0 0.5768079800498753", label="0.98",style=bold];
"MYOG_S" -> "CSRP3_E" [color="0 0 0.5760598503740648", label="0.98",style=bold];
"CHRND_S" -> "NRAP_M" [color="0 0 0.5753117206982543", label="0.98",style=bold];
"NRAP_S" -> "LRP1B_S" [color="0 0 0.5745635910224439", label="0.99",style=bold];
"NEB_S" -> "ZNF536_S" [color="0 0 0.5738154613466334", label="1.00",style=bold];
"FLG2_S" -> "SALL3_M" [color="0 0 0.573067331670823", label="1.00",style=bold];
"WDR49_S" -> "TTN_S" [color="0 0 0.5723192019950125", label="1.00",style=bold];
"MYH2_S" -> "WDR49_E" [color="0 0 0.571571072319202", label="1.00",style=bold];
"SLC18A3_S" -> "LRP1B_S" [color="0 0 0.5708229426433915", label="1.01",style=bold];
"SLC18A3_S" -> "CHRNA4_E" [color="0 0 0.570074812967581", label="1.01",style=bold];
"MYOG_S" -> "CDH10_E" [color="0 0 0.5693266832917706", label="1.01",style=bold];
"PLD5_S" -> "C6_S" [color="0 0 0.5685785536159601", label="1.01",style=bold];
"PLD5_S" -> "CCDC129_E" [color="0 0 0.5678304239401497", label="1.01",style=bold];
"PLD5_S" -> "COL11A1_M" [color="0 0 0.5670822942643392", label="1.01",style=bold];
"SLC18A3_S" -> "MYOG_M" [color="0 0 0.5663341645885287", label="1.01",style=bold];
"PAK7_S" -> "ASB5_S" [color="0 0 0.5655860349127182", label="1.02",style=bold];
"MYH7_S" -> "NLRP11_M" [color="0 0 0.5648379052369077", label="1.02",style=bold];
"MYH8_S" -> "MYH4_S" [color="0 0 0.5640897755610973", label="1.02",style=bold];
"FLG2_S" -> "PAK7_E" [color="0 0 0.5633416458852868", label="1.02",style=bold];
"CHRND_S" -> "TTN_E" [color="0 0 0.5625935162094764", label="1.02",style=bold];
"PKHD1L1_S" -> "LRP1B_S" [color="0 0 0.5618453865336659", label="1.03",style=bold];
"MYH6_S" -> "GABRA2_E" [color="0 0 0.5610972568578554", label="1.03",style=bold];
"MYH8_S" -> "COL11A1_E" [color="0 0 0.5603491271820449", label="1.03",style=bold];
"ZIC4_S" -> "ZIC1_E" [color="0 0 0.5596009975062344", label="1.04",style=bold];
"SLC18A3_S" -> "SLC18A3_E" [color="0 0 0.558852867830424", label="1.04",style=bold];
"ZNF280A_S" -> "CHRNA4_E" [color="0 0 0.5581047381546135", label="1.05",style=bold];
"PLD5_S" -> "CSRP3_M" [color="0 0 0.5573566084788031", label="1.06",style=bold];
"MPPED1_S" -> "MYBPC1_E" [color="0 0 0.5566084788029926", label="1.06",style=bold];
"MYH7_S" -> "TRDN_M" [color="0 0 0.5558603491271821", label="1.06",style=bold];
"PKHD1L1_S" -> "C6_S" [color="0 0 0.5551122194513716", label="1.06",style=bold];
"KRT38_S" -> "MYBPC1_S" [color="0 0 0.5543640897755611", label="1.06",style=bold];
"SLC18A3_S" -> "FLG2_M" [color="0 0 0.5536159600997507", label="1.07",style=bold];
"CNTN5_S" -> "FREM2_E" [color="0 0 0.5528678304239402", label="1.07",style=bold];
"MYH13_S" -> "MUC16_M" [color="0 0 0.5521197007481298", label="1.08",style=bold];
"KRT38_S" -> "TTN_S" [color="0 0 0.5513715710723193", label="1.08",style=bold];
"LCT_S" -> "CDH10_M" [color="0 0 0.5506234413965088", label="1.08",style=bold];
"PLD5_S" -> "XIRP2_M" [color="0 0 0.5498753117206983", label="1.09",style=bold];
"TEX15_S" -> "NRAP_S" [color="0 0 0.5491271820448878", label="1.10",style=bold];
"MYBPC1_S" -> "PPP1R3A_M" [color="0 0 0.5483790523690772", label="1.10",style=bold];
"MYL1_S" -> "TEX15_E" [color="0 0 0.5476309226932669", label="1.10",style=bold];
"MYL1_S" -> "DUSP27_M" [color="0 0 0.5468827930174565", label="1.10",style=bold];
"MYL1_S" -> "LRP1B_M" [color="0 0 0.546134663341646", label="1.10",style=bold];
"LCT_S" -> "CSMD1_S" [color="0 0 0.5453865336658354", label="1.11",style=bold];
"GABRA2_S" -> "ZNF280A_S" [color="0 0 0.5446384039900249", label="1.12",style=bold];
"NEB_S" -> "NLRP11_S" [color="0 0 0.5438902743142144", label="1.12",style=bold];
"DUSP27_E" -> "CDH10_S" [color="0 0 0.5431421446384039", label="1.13",style=bold];
"MYBPC1_S" -> "SLC18A3_M" [color="0 0 0.5423940149625935", label="1.13",style=bold];
"PPP1R3A_S" -> "PLD5_E" [color="0 0 0.5416458852867831", label="1.13",style=bold];
"PLD5_S" -> "KRT38_M" [color="0 0 0.5408977556109726", label="1.13",style=bold];
"SMYD1_S" -> "ZIC1_E" [color="0 0 0.5401496259351621", label="1.13",style=bold];
"PKHD1L1_S" -> "MUC4_M" [color="0 0 0.5394014962593516", label="1.14",style=bold];
"GRIN2B_S" -> "CNTNAP5_E" [color="0 0 0.5386533665835411", label="1.15",style=bold];
"CACNA1S_S" -> "MYL1_M" [color="0 0 0.5379052369077306", label="1.15",style=bold];
"GABRA2_S" -> "LCT_M" [color="0 0 0.5371571072319202", label="1.15",style=bold];
"MYL1_S" -> "MYH8_M" [color="0 0 0.5364089775561097", label="1.15",style=bold];
"SLC18A3_S" -> "MUC4_S" [color="0 0 0.5356608478802993", label="1.16",style=bold];
"KRT38_S" -> "ZIC1_S" [color="0 0 0.5349127182044888", label="1.17",style=bold];
"NRAP_S" -> "WDR49_M" [color="0 0 0.5341645885286783", label="1.17",style=bold];
"WDR49_S" -> "CSMD3_E" [color="0 0 0.5334164588528678", label="1.18",style=bold];
"FBN3_S" -> "MUC7_E" [color="0 0 0.5326683291770573", label="1.18",style=bold];
"SLC18A3_S" -> "ZNF536_S" [color="0 0 0.5319201995012469", label="1.18",style=bold];
"MUC7_S" -> "ZIC1_M" [color="0 0 0.5311720698254364", label="1.18",style=bold];
"ASB5_S" -> "MUC7_E" [color="0 0 0.530423940149626", label="1.18",style=bold];
"SMYD1_S" -> "CACNA1S_M" [color="0 0 0.5296758104738155", label="1.18",style=bold];
"WDR49_S" -> "MYH6_S" [color="0 0 0.528927680798005", label="1.18",style=bold];
"CHRND_S" -> "SALL3_S" [color="0 0 0.5281795511221945", label="1.19",style=bold];
"MYH1_E" -> "XIRP2_E" [color="0 0 0.527431421446384", label="1.19",style=bold];
"MUC7_S" -> "CSMD3_S" [color="0 0 0.5266832917705736", label="1.19",style=bold];
"SALL3_S" -> "LRP1B_E" [color="0 0 0.5259351620947631", label="1.19",style=bold];
"SLC7A14_S" -> "MUC16_S" [color="0 0 0.5251870324189527", label="1.19",style=bold];
"CNTN5_S" -> "ASB5_M" [color="0 0 0.5244389027431422", label="1.20",style=bold];
"SLC18A3_S" -> "CSMD3_E" [color="0 0 0.5236907730673317", label="1.20",style=bold];
"MYL1_S" -> "REG1A_M" [color="0 0 0.5229426433915212", label="1.20",style=bold];
"SLC7A14_S" -> "PLD5_M" [color="0 0 0.5221945137157107", label="1.20",style=bold];
"PPP1R3A_S" -> "ASB4_E" [color="0 0 0.5214463840399003", label="1.21",style=bold];
"MYH13_S" -> "ZIC1_M" [color="0 0 0.5206982543640898", label="1.22",style=bold];
"MYL1_S" -> "ZNF536_M" [color="0 0 0.5199501246882794", label="1.22",style=bold];
"CHRND_S" -> "TMEM132D_S" [color="0 0 0.5192019950124689", label="1.22",style=bold];
"CSRP3_S" -> "PAK7_M" [color="0 0 0.5184538653366584", label="1.23",style=bold];
"MYH13_S" -> "COL11A1_E" [color="0 0 0.5177057356608479", label="1.24",style=bold];
"SLC18A3_S" -> "CSMD1_S" [color="0 0 0.5169576059850374", label="1.24",style=bold];
"NRAP_S" -> "MUC16_S" [color="0 0 0.516209476309227", label="1.24",style=bold];
"TEX15_E" -> "COL11A1_M" [color="0 0 0.5154613466334165", label="1.24",style=bold];
"ZIC4_S" -> "LRP1B_S" [color="0 0 0.5147132169576061", label="1.25",style=bold];
"FLG2_S" -> "MYBPC1_M" [color="0 0 0.5139650872817956", label="1.25",style=bold];
"CNTNAP5_S" -> "MYH7_M" [color="0 0 0.5132169576059851", label="1.25",style=bold];
"ZIC4_S" -> "PKHD1L1_E" [color="0 0 0.5124688279301746", label="1.26",style=bold];
"ZNF536_S" -> "MPPED1_E" [color="0 0 0.5117206982543641", label="1.26",style=bold];
"NRAP_S" -> "GABRA2_E" [color="0 0 0.5109725685785536", label="1.27",style=bold];
"MYH6_S" -> "PKHD1L1_M" [color="0 0 0.5102244389027432", label="1.27",style=bold];
"ASB5_S" -> "C6_S" [color="0 0 0.5094763092269328", label="1.28",style=bold];
"ASB4_S" -> "CSRP3_M" [color="0 0 0.5087281795511223", label="1.28",style=bold];
"SLC18A3_S" -> "XIRP2_S" [color="0 0 0.5079800498753118", label="1.29",style=bold];
"COL11A1_S" -> "SLC18A3_E" [color="0 0 0.5072319201995013", label="1.29",style=bold];
"CNTN5_S" -> "MUC16_S" [color="0 0 0.5064837905236907", label="1.29",style=bold];
"SLC18A3_S" -> "CSRP3_E" [color="0 0 0.5057356608478802", label="1.29",style=bold];
"PAK7_S" -> "FBN3_S" [color="0 0 0.5049875311720698", label="1.29",style=bold];
"DUSP27_S" -> "MYH8_S" [color="0 0 0.5042394014962595", label="1.30",style=bold];
"GABRA2_S" -> "MYH8_S" [color="0 0 0.503491271820449", label="1.30",style=bold];
"ZNF280A_S" -> "LCT_M" [color="0 0 0.5027431421446384", label="1.30",style=bold];
"TEX15_S" -> "PKHD1_E" [color="0 0 0.5019950124688279", label="1.30",style=bold];
"SALL3_S" -> "SALL3_M" [color="0 0 0.5012468827930174", label="1.31",style=bold];
"SMYD1_S" -> "REG1A_M" [color="0 0 0.5004987531172069", label="1.31",style=bold];
"TRDN_S" -> "MYH7_S" [color="0 0 0.49975062344139654", label="1.32",style=bold];
"CHRND_S" -> "ZNF280A_M" [color="0 0 0.49900249376558603", label="1.33",style=bold];
"SMYD1_S" -> "ZIC1_M" [color="0 0 0.4982543640897756", label="1.33",style=bold];
"MUC7_S" -> "WDR49_E" [color="0 0 0.49750623441396513", label="1.33",style=bold];
"NLRP11_S" -> "PKHD1_S" [color="0 0 0.4967581047381546", label="1.34",style=bold];
"MYBPC1_S" -> "SALL3_M" [color="0 0 0.4960099750623442", label="1.35",style=bold];
"MYH4_S" -> "MUC16_S" [color="0 0 0.4952618453865337", label="1.36",style=bold];
"DUSP27_S" -> "CSMD1_S" [color="0 0 0.4945137157107232", label="1.36",style=bold];
"ZIC4_S" -> "CSRP3_M" [color="0 0 0.4937655860349127", label="1.36",style=bold];
"TRDN_S" -> "NLRP11_E" [color="0 0 0.49301745635910227", label="1.37",style=bold];
"CACNA1S_S" -> "CNTN5_E" [color="0 0 0.49226932668329176", label="1.37",style=bold];
"TRDN_S" -> "CHRNA4_E" [color="0 0 0.4915211970074813", label="1.37",style=bold];
"ZIC4_S" -> "SIX6_E" [color="0 0 0.49077306733167086", label="1.37",style=bold];
"CACNA1S_S" -> "PKHD1L1_M" [color="0 0 0.49002493765586036", label="1.38",style=bold];
"PLD5_S" -> "FLG_M" [color="0 0 0.4892768079800499", label="1.38",style=bold];
"GABRA2_S" -> "PKHD1L1_E" [color="0 0 0.4885286783042394", label="1.38",style=bold];
"ASB4_S" -> "MYL1_E" [color="0 0 0.48778054862842896", label="1.39",style=bold];
"MYL1_S" -> "CSMD1_M" [color="0 0 0.48703241895261845", label="1.39",style=bold];
"CHRNA4_S" -> "LRP1B_M" [color="0 0 0.486284289276808", label="1.39",style=bold];
"LCT_S" -> "MUC16_M" [color="0 0 0.48553615960099755", label="1.39",style=bold];
"ZIC4_S" -> "MUC16_S" [color="0 0 0.48478802992518705", label="1.40",style=bold];
"MYH6_S" -> "LRP1B_E" [color="0 0 0.4840399002493766", label="1.40",style=bold];
"PAK7_S" -> "MUC7_E" [color="0 0 0.4832917705735661", label="1.40",style=bold];
"SLC18A3_S" -> "ASB4_M" [color="0 0 0.48254364089775564", label="1.40",style=bold];
"REG1A_S" -> "WDR49_E" [color="0 0 0.48179551122194514", label="1.41",style=bold];
"ZIC4_S" -> "PCLO_E" [color="0 0 0.4810473815461347", label="1.41",style=bold];
"SIX6_S" -> "TEX15_M" [color="0 0 0.4802992518703242", label="1.43",style=bold];
"ASB5_S" -> "MUC16_M" [color="0 0 0.47955112219451373", label="1.43",style=bold];
"ASB5_S" -> "LRP1B_E" [color="0 0 0.4788029925187033", label="1.43",style=bold];
"SLC18A3_S" -> "SIX6_M" [color="0 0 0.4780548628428928", label="1.45",style=bold];
"MYH13_E" -> "CSMD1_S" [color="0 0 0.47730673316708233", label="1.45",style=bold];
"PLD5_S" -> "LRP1B_S" [color="0 0 0.4765586034912718", label="1.45",style=bold];
"CHRNA4_S" -> "NEB_S" [color="0 0 0.4758104738154614", label="1.45",style=bold];
"GABRA2_M" -> "GABRA2_E" [color="0 0 0.47506234413965087", label="1.45",style=bold];
"WDR49_S" -> "PKHD1L1_E" [color="0 0 0.4743142144638404", label="1.46",style=bold];
"MYH1_S" -> "MPPED1_E" [color="0 0 0.47356608478803", label="1.47",style=bold];
"TRDN_S" -> "CSMD3_E" [color="0 0 0.47281795511221947", label="1.48",style=bold];
"SALL3_S" -> "TEX15_M" [color="0 0 0.472069825436409", label="1.48",style=bold];
"MYH6_S" -> "SLC18A3_E" [color="0 0 0.4713216957605985", label="1.48",style=bold];
"SMYD1_S" -> "CSMD3_M" [color="0 0 0.47057356608478806", label="1.49",style=bold];
"MYH1_S" -> "CNTNAP5_M" [color="0 0 0.46982543640897756", label="1.49",style=bold];
"ZIC1_S" -> "MYBPC1_M" [color="0 0 0.4690773067331671", label="1.49",style=bold];
"MYH1_S" -> "SIX6_M" [color="0 0 0.4683291770573566", label="1.49",style=bold];
"MUC4_S" -> "GABRA2_E" [color="0 0 0.46758104738154616", label="1.50",style=bold];
"MYL1_S" -> "MYH1_M" [color="0 0 0.4668329177057357", label="1.50",style=bold];
"MYL1_S" -> "C6_M" [color="0 0 0.4660847880299252", label="1.52",style=bold];
"PLD5_S" -> "KRT38_E" [color="0 0 0.46533665835411475", label="1.52",style=bold];
"CHRND_S" -> "KRT38_E" [color="0 0 0.46458852867830425", label="1.52",style=bold];
"SALL3_S" -> "C6_M" [color="0 0 0.4638403990024938", label="1.53",style=bold];
"CCDC129_S" -> "SIX6_M" [color="0 0 0.4630922693266833", label="1.53",style=bold];
"PLD5_S" -> "PKHD1_E" [color="0 0 0.46234413965087284", label="1.53",style=bold];
"CSMD1_E" -> "NLRP11_E" [color="0 0 0.46159600997506234", label="1.53",style=bold];
"PAK7_S" -> "PKHD1L1_M" [color="0 0 0.4608478802992519", label="1.54",style=bold];
"SMYD1_S" -> "GRIN2B_M" [color="0 0 0.46009975062344144", label="1.54",style=bold];
"ZIC4_S" -> "TRDN_M" [color="0 0 0.45935162094763093", label="1.54",style=bold];
"ASB4_S" -> "MUC16_S" [color="0 0 0.4586034912718205", label="1.54",style=bold];
"WDR49_S" -> "MYH7_E" [color="0 0 0.45785536159601", label="1.54",style=bold];
"PLD5_S" -> "WDR49_M" [color="0 0 0.45710723192019953", label="1.55",style=bold];
"MYL1_S" -> "SIX6_E" [color="0 0 0.456359102244389", label="1.55",style=bold];
"CNTNAP5_S" -> "FREM2_S" [color="0 0 0.4556109725685786", label="1.57",style=bold];
"FLG2_S" -> "CNTN5_E" [color="0 0 0.4548628428927681", label="1.58",style=bold];
"NLRP11_S" -> "GRIN2B_E" [color="0 0 0.4541147132169576", label="1.59",style=bold];
"ZNF536_M" -> "MYH2_M" [color="0 0 0.4533665835411472", label="1.59",style=bold];
"MYH7_S" -> "CSRP3_E" [color="0 0 0.45261845386533667", label="1.59",style=bold];
"ZNF280A_S" -> "KRT38_E" [color="0 0 0.4518703241895262", label="1.60",style=bold];
"TEX15_S" -> "TEX15_E" [color="0 0 0.4511221945137157", label="1.60",style=bold];
"REG1A_S" -> "ZIC1_M" [color="0 0 0.45037406483790526", label="1.61",style=bold];
"ZNF536_S" -> "LRP1B_S" [color="0 0 0.44962593516209476", label="1.61",style=bold];
"ZNF280A_S" -> "NLRP11_E" [color="0 0 0.4488778054862843", label="1.61",style=bold];
"SLC7A14_S" -> "SMYD1_M" [color="0 0 0.44812967581047386", label="1.61",style=bold];
"CHRNA4_S" -> "FLG_S" [color="0 0 0.44738154613466335", label="1.61",style=bold];
"DUSP27_S" -> "MUC16_M" [color="0 0 0.4466334164588529", label="1.61",style=bold];
"MYH6_S" -> "CSMD3_S" [color="0 0 0.4458852867830424", label="1.62",style=bold];
"CDH10_S" -> "ZIC1_E" [color="0 0 0.44513715710723195", label="1.62",style=bold];
"MYBPC1_M" -> "TEX15_E" [color="0 0 0.44438902743142145", label="1.63",style=bold];
"MYH4_S" -> "LRP1B_E" [color="0 0 0.443640897755611", label="1.64",style=bold];
"TRDN_E" -> "FREM2_E" [color="0 0 0.4428927680798005", label="1.64",style=bold];
"ZIC4_S" -> "NEB_M" [color="0 0 0.44214463840399004", label="1.64",style=bold];
"CNTN5_S" -> "KRT38_E" [color="0 0 0.4413965087281796", label="1.64",style=bold];
"KRT38_S" -> "PKHD1_S" [color="0 0 0.4406483790523691", label="1.65",style=bold];
"CSRP3_E" -> "CACNA1S_E" [color="0 0 0.43990024937655864", label="1.66",style=bold];
"DUSP27_S" -> "TMEM132D_S" [color="0 0 0.43915211970074813", label="1.66",style=bold];
"KRT38_S" -> "SMYD1_M" [color="0 0 0.4384039900249377", label="1.66",style=bold];
"CHRND_S" -> "ZIC1_S" [color="0 0 0.4376558603491272", label="1.66",style=bold];
"FLG_S" -> "XIRP2_S" [color="0 0 0.43690773067331673", label="1.67",style=bold];
"MYH6_S" -> "WDR49_E" [color="0 0 0.4361596009975063", label="1.67",style=bold];
"CHRND_S" -> "MYH6_S" [color="0 0 0.4354114713216958", label="1.67",style=bold];
"SLC18A3_S" -> "MYH6_S" [color="0 0 0.4346633416458853", label="1.67",style=bold];
"MYH6_S" -> "ZIC1_E" [color="0 0 0.4339152119700748", label="1.68",style=bold];
"ZNF536_S" -> "KRT38_E" [color="0 0 0.43316708229426437", label="1.69",style=bold];
"SMYD1_E" -> "CACNA1S_E" [color="0 0 0.43241895261845387", label="1.69",style=bold];
"SMYD1_S" -> "SIX6_M" [color="0 0 0.4316708229426434", label="1.70",style=bold];
"GABRA2_S" -> "PAK7_E" [color="0 0 0.4309226932668329", label="1.70",style=bold];
"PLD5_S" -> "PLD5_E" [color="0 0 0.43017456359102246", label="1.70",style=bold];
"CHRNA4_S" -> "NLRP11_S" [color="0 0 0.429426433915212", label="1.70",style=bold];
"FBN3_S" -> "TEX15_E" [color="0 0 0.4286783042394015", label="1.71",style=bold];
"MUC7_S" -> "TMEM132D_S" [color="0 0 0.42793017456359106", label="1.71",style=bold];
"MYH8_S" -> "GABRA2_M" [color="0 0 0.42718204488778055", label="1.71",style=bold];
"PAK7_S" -> "MYH4_M" [color="0 0 0.4264339152119701", label="1.71",style=bold];
"CACNA1S_S" -> "FLG2_E" [color="0 0 0.4256857855361596", label="1.71",style=bold];
"MYH1_S" -> "ZIC1_E" [color="0 0 0.42493765586034915", label="1.71",style=bold];
"CNTNAP5_S" -> "SALL3_M" [color="0 0 0.4241895261845387", label="1.72",style=bold];
"PAK7_S" -> "MYH8_S" [color="0 0 0.4234413965087282", label="1.73",style=bold];
"MYL1_S" -> "MYH6_E" [color="0 0 0.42269326683291775", label="1.73",style=bold];
"SIX6_S" -> "CHRND_M" [color="0 0 0.42194513715710724", label="1.75",style=bold];
"ZIC1_S" -> "PLD5_E" [color="0 0 0.4211970074812968", label="1.76",style=bold];
"SIX6_S" -> "FLG_S" [color="0 0 0.4204488778054863", label="1.76",style=bold];
"CDH10_S" -> "MUC4_S" [color="0 0 0.41970074812967584", label="1.77",style=bold];
"NRAP_S" -> "PCLO_S" [color="0 0 0.41895261845386533", label="1.77",style=bold];
"CACNA1S_S" -> "MUC16_M" [color="0 0 0.4182044887780549", label="1.78",style=bold];
"REG1A_S" -> "TTN_E" [color="0 0 0.41745635910224443", label="1.80",style=bold];
"WDR49_S" -> "MYH7_S" [color="0 0 0.41670822942643393", label="1.80",style=bold];
"MPPED1_S" -> "SIX6_E" [color="0 0 0.4159600997506235", label="1.82",style=bold];
"CNTN5_S" -> "PCLO_E" [color="0 0 0.415211970074813", label="1.82",style=bold];
"ASB5_S" -> "COL11A1_M" [color="0 0 0.4144638403990025", label="1.82",style=bold];
"CSRP3_S" -> "PCLO_S" [color="0 0 0.413715710723192", label="1.82",style=bold];
"MUC7_S" -> "PLD5_E" [color="0 0 0.41296758104738157", label="1.83",style=bold];
"XIRP2_E" -> "MYH2_E" [color="0 0 0.41221945137157107", label="1.84",style=bold];
"ZNF536_S" -> "MUC16_S" [color="0 0 0.4114713216957606", label="1.84",style=bold];
"FLG2_S" -> "KRT38_E" [color="0 0 0.41072319201995017", label="1.84",style=bold];
"FLG2_S" -> "FREM2_E" [color="0 0 0.40997506234413966", label="1.84",style=bold];
"SALL3_S" -> "CNTN5_E" [color="0 0 0.4092269326683292", label="1.84",style=bold];
"DUSP27_S" -> "TEX15_M" [color="0 0 0.4084788029925187", label="1.84",style=bold];
"CNTNAP5_S" -> "SIX6_M" [color="0 0 0.40773067331670826", label="1.85",style=bold];
"ASB4_S" -> "CCDC129_E" [color="0 0 0.40698254364089775", label="1.85",style=bold];
"SMYD1_S" -> "MYH13_S" [color="0 0 0.4062344139650873", label="1.85",style=bold];
"MUC4_S" -> "Tumor_Status" [color="0 0 0.40548628428927685", label="1.85",style=bold];
"MYH6_S" -> "LCT_E" [color="0 0 0.40473815461346635", label="1.86",style=bold];
"MYL1_S" -> "MUC7_E" [color="0 0 0.4039900249376559", label="1.87",style=bold];
"SLC18A3_S" -> "MUC16_E" [color="0 0 0.4032418952618454", label="1.87",style=bold];
"DUSP27_S" -> "PCLO_M" [color="0 0 0.40249376558603495", label="1.88",style=bold];
"SLC18A3_S" -> "CNTNAP5_S" [color="0 0 0.40174563591022444", label="1.89",style=bold];
"PKHD1_S" -> "CSMD3_S" [color="0 0 0.400997506234414", label="1.89",style=bold];
"GABRA2_S" -> "PKHD1L1_M" [color="0 0 0.4002493765586035", label="1.90",style=bold];
"CHRNA4_S" -> "CNTN5_M" [color="0 0 0.3995012468827931", label="1.90",style=bold];
"FLG_S" -> "PCLO_E" [color="0 0 0.3987531172069826", label="1.90",style=bold];
"TRDN_S" -> "WDR49_S" [color="0 0 0.3980049875311721", label="1.90",style=bold];
"CACNA1S_S" -> "PKHD1L1_S" [color="0 0 0.3972568578553616", label="1.91",style=bold];
"PKHD1_S" -> "PCLO_E" [color="0 0 0.3965087281795512", label="1.91",style=bold];
"CHRND_S" -> "FBN3_S" [color="0 0 0.3957605985037407", label="1.91",style=bold];
"SIX6_S" -> "COL11A1_M" [color="0 0 0.3950124688279302", label="1.92",style=bold];
"WDR49_S" -> "MUC16_M" [color="0 0 0.39426433915211967", label="1.92",style=bold];
"MPPED1_S" -> "SLC7A14_E" [color="0 0 0.3935162094763093", label="1.92",style=bold];
"CSRP3_S" -> "MYH2_S" [color="0 0 0.39276807980049877", label="1.93",style=bold];
"FBN3_S" -> "PPP1R3A_M" [color="0 0 0.39201995012468827", label="1.93",style=bold];
"LCT_S" -> "PKHD1L1_M" [color="0 0 0.39127182044887787", label="1.94",style=bold];
"MYH7_S" -> "PKHD1L1_S" [color="0 0 0.39052369077306737", label="1.95",style=bold];
"MYH6_S" -> "ASB4_M" [color="0 0 0.38977556109725686", label="1.95",style=bold];
"GRIN2B_S" -> "PAK7_E" [color="0 0 0.38902743142144636", label="1.95",style=bold];
"ASB5_S" -> "MUC4_E" [color="0 0 0.38827930174563596", label="1.96",style=bold];
"CHRNA4_S" -> "PAK7_M" [color="0 0 0.38753117206982546", label="1.97",style=bold];
"MYBPC1_S" -> "GABRA2_E" [color="0 0 0.38678304239401495", label="1.97",style=bold];
"FREM2_S" -> "CCDC129_E" [color="0 0 0.38603491271820456", label="1.98",style=bold];
"ZNF280A_S" -> "PLD5_M" [color="0 0 0.38528678304239405", label="1.98",style=bold];
"MYOG_S" -> "ZIC4_S" [color="0 0 0.38453865336658355", label="1.99",style=bold];
"CNTN5_S" -> "ZNF536_E" [color="0 0 0.38379052369077304", label="1.99",style=bold];
"MYL1_S" -> "LRP1B_S" [color="0 0 0.38304239401496265", label="2.01",style=bold];
"CNTN5_S" -> "MUC16_M" [color="0 0 0.38229426433915215", label="2.01",style=bold];
"CSRP3_S" -> "CDH10_E" [color="0 0 0.38154613466334164", label="2.01",style=bold];
"SLC7A14_S" -> "MUC4_E" [color="0 0 0.38079800498753125", label="2.02",style=bold];
"PAK7_S" -> "TMEM132D_E" [color="0 0 0.38004987531172074", label="2.03",style=bold];
"MYH7_S" -> "DUSP27_M" [color="0 0 0.37930174563591024", label="2.05",style=bold];
"NLRP11_S" -> "LCT_E" [color="0 0 0.37855361596009973", label="2.05",style=bold];
"MYH13_S" -> "MUC4_S" [color="0 0 0.37780548628428934", label="2.07",style=bold];
"NRAP_S" -> "PKHD1L1_S" [color="0 0 0.37705735660847883", label="2.07",style=bold];
"SALL3_S" -> "KRT38_E" [color="0 0 0.37630922693266833", label="2.07",style=bold];
"NLRP11_S" -> "PCLO_S" [color="0 0 0.3755610972568578", label="2.07",style=bold];
"REG1A_S" -> "CSRP3_E" [color="0 0 0.37481296758104743", label="2.08",style=bold];
"SLC18A3_S" -> "MUC16_M" [color="0 0 0.3740648379052369", label="2.08",style=bold];
"MYH7_E" -> "MYH13_E" [color="0 0 0.3733167082294264", label="2.09",style=bold];
"MYOG_S" -> "FBN3_E" [color="0 0 0.372568578553616", label="2.09",style=bold];
"PLD5_S" -> "NEB_M" [color="0 0 0.3718204488778055", label="2.10",style=bold];
"MUC7_S" -> "CDH10_M" [color="0 0 0.371072319201995", label="2.10",style=bold];
"LCT_S" -> "PAK7_M" [color="0 0 0.3703241895261845", label="2.11",style=bold];
"MYBPC1_S" -> "LRP1B_E" [color="0 0 0.3695760598503741", label="2.12",style=bold];
"MYH13_S" -> "MYH4_S" [color="0 0 0.3688279301745636", label="2.12",style=bold];
"SLC18A3_S" -> "CCDC129_E" [color="0 0 0.3680798004987531", label="2.12",style=bold];
"MYH1_S" -> "FBN3_M" [color="0 0 0.3673316708229427", label="2.13",style=bold];
"LCT_S" -> "TRDN_M" [color="0 0 0.3665835411471322", label="2.14",style=bold];
"MYH7_S" -> "CSRP3_M" [color="0 0 0.3658354114713217", label="2.14",style=bold];
"GABRA2_S" -> "ZNF536_E" [color="0 0 0.3650872817955112", label="2.14",style=bold];
"WDR49_S" -> "PAK7_E" [color="0 0 0.3643391521197008", label="2.14",style=bold];
"FBN3_S" -> "ZIC1_E" [color="0 0 0.3635910224438903", label="2.15",style=bold];
"PAK7_S" -> "C6_S" [color="0 0 0.3628428927680798", label="2.16",style=bold];
"ZNF280A_S" -> "TTN_S" [color="0 0 0.3620947630922694", label="2.16",style=bold];
"ASB5_S" -> "ZNF536_E" [color="0 0 0.3613466334164589", label="2.17",style=bold];
"CHRNA4_S" -> "MYOG_E" [color="0 0 0.3605985037406484", label="2.17",style=bold];
"ZIC1_S" -> "ZNF280A_E" [color="0 0 0.3598503740648379", label="2.17",style=bold];
"MPPED1_S" -> "TRDN_S" [color="0 0 0.3591022443890275", label="2.18",style=bold];
"MYBPC1_S" -> "LCT_E" [color="0 0 0.358354114713217", label="2.20",style=bold];
"FBN3_S" -> "SALL3_M" [color="0 0 0.3576059850374065", label="2.21",style=bold];
"ASB5_S" -> "PLD5_E" [color="0 0 0.356857855361596", label="2.21",style=bold];
"GRIN2B_S" -> "MYBPC1_S" [color="0 0 0.3561097256857856", label="2.22",style=bold];
"MYBPC1_S" -> "FLG_M" [color="0 0 0.3553615960099751", label="2.23",style=bold];
"TEX15_S" -> "GABRA2_E" [color="0 0 0.3546134663341646", label="2.23",style=bold];
"MYH8_S" -> "CNTNAP5_E" [color="0 0 0.3538653366583542", label="2.24",style=bold];
"NRAP_S" -> "NEB_S" [color="0 0 0.3531172069825437", label="2.24",style=bold];
"MYH2_S" -> "MYBPC1_S" [color="0 0 0.35236907730673317", label="2.24",style=bold];
"FLG2_S" -> "FBN3_E" [color="0 0 0.35162094763092266", label="2.25",style=bold];
"SLC7A14_S" -> "ZNF536_S" [color="0 0 0.35087281795511227", label="2.25",style=bold];
"ASB5_S" -> "LCT_E" [color="0 0 0.35012468827930177", label="2.27",style=bold];
"ZNF280A_S" -> "REG1A_S" [color="0 0 0.34937655860349126", label="2.27",style=bold];
"ZIC4_S" -> "LCT_M" [color="0 0 0.34862842892768087", label="2.28",style=bold];
"CDH10_S" -> "TMEM132D_E" [color="0 0 0.34788029925187036", label="2.28",style=bold];
"CHRND_S" -> "MYBPC1_M" [color="0 0 0.34713216957605986", label="2.29",style=bold];
"DUSP27_S" -> "LRP1B_M" [color="0 0 0.34638403990024935", label="2.30",style=bold];
"MUC4_S" -> "CSRP3_M" [color="0 0 0.34563591022443896", label="2.31",style=bold];
"MUC7_S" -> "PCLO_M" [color="0 0 0.34488778054862845", label="2.31",style=bold];
"SALL3_S" -> "ZNF536_E" [color="0 0 0.34413965087281795", label="2.32",style=bold];
"MYH8_S" -> "TTN_S" [color="0 0 0.34339152119700755", label="2.34",style=bold];
"FBN3_S" -> "LRP1B_E" [color="0 0 0.34264339152119705", label="2.34",style=bold];
"FBN3_S" -> "TTN_M" [color="0 0 0.34189526184538654", label="2.35",style=bold];
"MYH7_S" -> "MUC16_M" [color="0 0 0.34114713216957604", label="2.35",style=bold];
"ZNF536_S" -> "MYH7_M" [color="0 0 0.34039900249376565", label="2.36",style=bold];
"ZIC1_S" -> "CSRP3_E" [color="0 0 0.33965087281795514", label="2.36",style=bold];
"NRAP_S" -> "ZIC4_M" [color="0 0 0.33890274314214464", label="2.36",style=bold];
"MYH7_S" -> "WDR49_M" [color="0 0 0.33815461346633413", label="2.37",style=bold];
"DUSP27_S" -> "CSMD3_S" [color="0 0 0.33740648379052374", label="2.37",style=bold];
"PAK7_S" -> "TTN_E" [color="0 0 0.33665835411471323", label="2.39",style=bold];
"MUC4_S" -> "FBN3_M" [color="0 0 0.3359102244389027", label="2.41",style=bold];
"C6_S" -> "MUC7_E" [color="0 0 0.33516209476309233", label="2.41",style=bold];
"MYH7_S" -> "MYH8_S" [color="0 0 0.33441396508728183", label="2.42",style=bold];
"SMYD1_E" -> "TRDN_E" [color="0 0 0.3336658354114713", label="2.42",style=bold];
"SALL3_S" -> "FREM2_E" [color="0 0 0.3329177057356608", label="2.43",style=bold];
"TRDN_S" -> "TTN_S" [color="0 0 0.3321695760598504", label="2.44",style=bold];
"GABRA2_S" -> "MUC16_S" [color="0 0 0.3314214463840399", label="2.44",style=bold];
"SIX6_S" -> "C6_S" [color="0 0 0.3306733167082294", label="2.45",style=bold];
"FLG2_S" -> "LRP1B_S" [color="0 0 0.329925187032419", label="2.48",style=bold];
"WDR49_S" -> "SLC7A14_E" [color="0 0 0.3291770573566085", label="2.48",style=bold];
"WDR49_S" -> "COL11A1_E" [color="0 0 0.328428927680798", label="2.48",style=bold];
"PKHD1L1_S" -> "CSMD1_E" [color="0 0 0.3276807980049875", label="2.49",style=bold];
"ASB4_S" -> "CHRNA4_M" [color="0 0 0.3269326683291771", label="2.49",style=bold];
"ZIC1_S" -> "CCDC129_M" [color="0 0 0.3261845386533666", label="2.52",style=bold];
"NLRP11_S" -> "TMEM132D_E" [color="0 0 0.3254364089775561", label="2.54",style=bold];
"FLG2_S" -> "SALL3_E" [color="0 0 0.3246882793017457", label="2.55",style=bold];
"CNTN5_S" -> "TTN_M" [color="0 0 0.3239401496259352", label="2.56",style=bold];
"PKHD1L1_S" -> "CSMD3_S" [color="0 0 0.3231920199501247", label="2.58",style=bold];
"CNTNAP5_E" -> "CDH10_E" [color="0 0 0.3224438902743142", label="2.59",style=bold];
"ZNF536_S" -> "MUC16_E" [color="0 0 0.3216957605985038", label="2.60",style=bold];
"ZNF280A_S" -> "ZNF280A_E" [color="0 0 0.3209476309226933", label="2.60",style=bold];
"FLG2_S" -> "FLG_S" [color="0 0 0.3201995012468828", label="2.61",style=bold];
"CHRND_S" -> "COL11A1_S" [color="0 0 0.3194513715710723", label="2.62",style=bold];
"MYOG_S" -> "PLD5_S" [color="0 0 0.3187032418952619", label="2.63",style=bold];
"FBN3_S" -> "ZIC1_M" [color="0 0 0.3179551122194514", label="2.64",style=bold];
"MUC4_S" -> "FBN3_E" [color="0 0 0.3172069825436409", label="2.64",style=bold];
"GRIN2B_S" -> "CNTN5_E" [color="0 0 0.3164588528678305", label="2.64",style=bold];
"MYH7_S" -> "ZIC1_E" [color="0 0 0.31571072319202", label="2.64",style=bold];
"SMYD1_S" -> "CNTNAP5_S" [color="0 0 0.3149625935162095", label="2.66",style=bold];
"ZIC1_S" -> "PPP1R3A_M" [color="0 0 0.31421446384039897", label="2.67",style=bold];
"C6_S" -> "C6_M" [color="0 0 0.3134663341645886", label="2.68",style=bold];
"MYH2_S" -> "MUC16_S" [color="0 0 0.3127182044887781", label="2.68",style=bold];
"FBN3_S" -> "MUC4_M" [color="0 0 0.31197007481296757", label="2.68",style=bold];
"ASB4_M" -> "MYH2_M" [color="0 0 0.3112219451371572", label="2.68",style=bold];
"DUSP27_S" -> "ASB4_E" [color="0 0 0.31047381546134667", label="2.68",style=bold];
"MPPED1_S" -> "SMYD1_S" [color="0 0 0.30972568578553616", label="2.69",style=bold];
"MUC7_S" -> "GRIN2B_E" [color="0 0 0.30897755610972566", label="2.70",style=bold];
"NLRP11_S" -> "MUC16_S" [color="0 0 0.30822942643391527", label="2.70",style=bold];
"ASB4_S" -> "FREM2_S" [color="0 0 0.30748129675810476", label="2.70",style=bold];
"CCDC129_S" -> "LRP1B_S" [color="0 0 0.30673316708229426", label="2.71",style=bold];
"MYH2_E" -> "ZIC1_E" [color="0 0 0.30598503740648386", label="2.71",style=bold];
"MYH6_M" -> "FLG2_M" [color="0 0 0.30523690773067336", label="2.71",style=bold];
"CHRNA4_M" -> "ZNF280A_M" [color="0 0 0.30448877805486285", label="2.72",style=bold];
"CNTNAP5_M" -> "Tumor_Status" [color="0 0 0.30374064837905235", label="2.73",style=bold];
"TRDN_S" -> "MUC7_E" [color="0 0 0.30299251870324195", label="2.73",style=bold];
"GABRA2_S" -> "GRIN2B_S" [color="0 0 0.30224438902743145", label="2.73",style=bold];
"TMEM132D_S" -> "C6_E" [color="0 0 0.30149625935162094", label="2.73",style=bold];
"SALL3_S" -> "ZNF280A_E" [color="0 0 0.30074812967581044", label="2.75",style=bold];
"MYOG_S" -> "MYH1_S" [color="0 0 0.30000000000000004", label="2.75",style=bold];
"MYH7_S" -> "CNTN5_M" [color="0 0 0.29925187032418954", label="2.76",style=bold];
"MUC16_S" -> "XIRP2_S" [color="0 0 0.29850374064837903", label="2.76",style=bold];
"SMYD1_S" -> "MUC16_S" [color="0 0 0.29775561097256864", label="2.77",style=bold];
"MUC4_S" -> "TMEM132D_E" [color="0 0 0.29700748129675814", label="2.78",style=bold];
"MYBPC1_S" -> "MUC4_E" [color="0 0 0.29625935162094763", label="2.82",style=bold];
"PLD5_S" -> "PKHD1_S" [color="0 0 0.2955112219451371", label="2.83",style=bold];
"ZNF536_S" -> "PCLO_E" [color="0 0 0.29476309226932673", label="2.84",style=bold];
"LCT_S" -> "KRT38_E" [color="0 0 0.2940149625935162", label="2.84",style=bold];
"WDR49_S" -> "C6_S" [color="0 0 0.2932668329177057", label="2.84",style=bold];
"CNTN5_S" -> "WDR49_M" [color="0 0 0.29251870324189533", label="2.85",style=bold];
"MYH6_S" -> "MUC16_S" [color="0 0 0.2917705735660848", label="2.85",style=bold];
"TMEM132D_S" -> "MUC16_M" [color="0 0 0.2910224438902743", label="2.86",style=bold];
"PPP1R3A_S" -> "PAK7_E" [color="0 0 0.2902743142144638", label="2.86",style=bold];
"PCLO_S" -> "TMEM132D_S" [color="0 0 0.2895261845386534", label="2.89",style=bold];
"MYH8_S" -> "LRP1B_E" [color="0 0 0.2887780548628429", label="2.90",style=bold];
"TRDN_S" -> "C6_S" [color="0 0 0.2880299251870324", label="2.91",style=bold];
"ZNF536_S" -> "LCT_E" [color="0 0 0.287281795511222", label="2.94",style=bold];
"MYOG_S" -> "PKHD1_M" [color="0 0 0.2865336658354115", label="2.95",style=bold];
"CNTN5_S" -> "FLG2_M" [color="0 0 0.285785536159601", label="2.95",style=bold];
"FLG2_S" -> "TTN_S" [color="0 0 0.2850374064837905", label="2.97",style=bold];
"CHRNA4_S" -> "GRIN2B_S" [color="0 0 0.2842892768079801", label="3.01",style=bold];
"ZIC1_S" -> "CSMD3_S" [color="0 0 0.2835411471321696", label="3.01",style=bold];
"LCT_S" -> "FREM2_M" [color="0 0 0.2827930174563591", label="3.02",style=bold];
"MYH1_S" -> "CSMD3_S" [color="0 0 0.2820448877805487", label="3.03",style=bold];
"DUSP27_S" -> "CDH10_S" [color="0 0 0.2812967581047382", label="3.05",style=bold];
"FLG_S" -> "FREM2_S" [color="0 0 0.2805486284289277", label="3.05",style=bold];
"ZIC1_S" -> "MUC16_E" [color="0 0 0.2798004987531172", label="3.06",style=bold];
"PAK7_S" -> "FLG_M" [color="0 0 0.2790523690773068", label="3.08",style=bold];
"ASB4_S" -> "MPPED1_S" [color="0 0 0.2783042394014963", label="3.08",style=bold];
"GRIN2B_S" -> "NLRP11_E" [color="0 0 0.2775561097256858", label="3.08",style=bold];
"MYH1_S" -> "MPPED1_M" [color="0 0 0.2768079800498753", label="3.09",style=bold];
"MPPED1_S" -> "MYOG_S" [color="0 0 0.2760598503740649", label="3.09",style=bold];
"PKHD1L1_M" -> "MYL1_M" [color="0 0 0.2753117206982544", label="3.09",style=bold];
"REG1A_S" -> "COL11A1_S" [color="0 0 0.2745635910224439", label="3.09",style=bold];
"C6_M" -> "CHRND_M" [color="0 0 0.2738154613466335", label="3.10",style=bold];
"CACNA1S_S" -> "PPP1R3A_S" [color="0 0 0.273067331670823", label="3.10",style=bold];
"ZIC1_S" -> "XIRP2_M" [color="0 0 0.27231920199501247", label="3.11",style=bold];
"GABRA2_S" -> "FREM2_E" [color="0 0 0.27157107231920197", label="3.11",style=bold];
"PPP1R3A_S" -> "FREM2_S" [color="0 0 0.2708229426433916", label="3.12",style=bold];
"CDH10_S" -> "MUC7_M" [color="0 0 0.27007481296758107", label="3.12",style=bold];
"WDR49_S" -> "GABRA2_E" [color="0 0 0.26932668329177056", label="3.12",style=bold];
"ZNF280A_S" -> "ZIC1_M" [color="0 0 0.26857855361596017", label="3.12",style=bold];
"MUC7_S" -> "ASB4_M" [color="0 0 0.26783042394014966", label="3.14",style=bold];
"KRT38_S" -> "MPPED1_M" [color="0 0 0.26708229426433916", label="3.15",style=bold];
"KRT38_S" -> "C6_S" [color="0 0 0.26633416458852865", label="3.15",style=bold];
"SLC7A14_S" -> "CNTNAP5_E" [color="0 0 0.26558603491271826", label="3.16",style=bold];
"MYOG_S" -> "MYH7_M" [color="0 0 0.26483790523690776", label="3.16",style=bold];
"PPP1R3A_S" -> "SIX6_E" [color="0 0 0.26408977556109725", label="3.18",style=bold];
"GABRA2_S" -> "CSMD1_E" [color="0 0 0.26334164588528686", label="3.21",style=bold];
"TRDN_E" -> "PAK7_E" [color="0 0 0.26259351620947635", label="3.26",style=bold];
"ZIC4_S" -> "MUC16_M" [color="0 0 0.26184538653366585", label="3.28",style=bold];
"MUC4_S" -> "C6_E" [color="0 0 0.26109725685785534", label="3.33",style=bold];
"PLD5_S" -> "FBN3_E" [color="0 0 0.26034912718204495", label="3.33",style=bold];
"CDH10_S" -> "PKHD1L1_E" [color="0 0 0.25960099750623444", label="3.36",style=bold];
"PKHD1L1_S" -> "NLRP11_M" [color="0 0 0.25885286783042394", label="3.39",style=bold];
"MYOG_S" -> "ZIC1_S" [color="0 0 0.25810473815461343", label="3.41",style=bold];
"GABRA2_S" -> "ZIC1_S" [color="0 0 0.25735660847880304", label="3.44",style=bold];
"MYBPC1_M" -> "TRDN_M" [color="0 0 0.25660847880299253", label="3.45",style=bold];
"CACNA1S_S" -> "MYH6_S" [color="0 0 0.25586034912718203", label="3.46",style=bold];
"MYL1_M" -> "PPP1R3A_M" [color="0 0 0.25511221945137164", label="3.46",style=bold];
"MYOG_S" -> "WDR49_S" [color="0 0 0.25436408977556113", label="3.47",style=bold];
"MYH7_S" -> "SMYD1_M" [color="0 0 0.2536159600997506", label="3.47",style=bold];
"MYH13_S" -> "CACNA1S_M" [color="0 0 0.2528678304239401", label="3.47",style=bold];
"TEX15_S" -> "FBN3_E" [color="0 0 0.2521197007481297", label="3.47",style=bold];
"NEB_M" -> "ZIC4_M" [color="0 0 0.2513715710723192", label="3.48",style=bold];
"SMYD1_E" -> "NRAP_E" [color="0 0 0.2506234413965087", label="3.48",style=bold];
"MYH6_S" -> "REG1A_S" [color="0 0 0.24987531172069832", label="3.49",style=bold];
"GRIN2B_S" -> "LRP1B_E" [color="0 0 0.24912718204488782", label="3.49",style=bold];
"FBN3_S" -> "CNTN5_E" [color="0 0 0.2483790523690773", label="3.51",style=bold];
"NEB_S" -> "MUC16_E" [color="0 0 0.2476309226932668", label="3.53",style=bold];
"FBN3_S" -> "SIX6_E" [color="0 0 0.24688279301745641", label="3.55",style=bold];
"LRP1B_M" -> "CNTNAP5_M" [color="0 0 0.2461346633416459", label="3.59",style=bold];
"CNTNAP5_S" -> "LRP1B_S" [color="0 0 0.2453865336658354", label="3.60",style=bold];
"MYH2_E" -> "MYH4_E" [color="0 0 0.244638403990025", label="3.62",style=bold];
"MYH4_S" -> "CHRNA4_M" [color="0 0 0.2438902743142145", label="3.65",style=bold];
"CNTN5_S" -> "ZNF536_S" [color="0 0 0.243142144638404", label="3.66",style=bold];
"MYL1_S" -> "REG1A_S" [color="0 0 0.2423940149625935", label="3.71",style=bold];
"LCT_M" -> "FLG_M" [color="0 0 0.2416458852867831", label="3.74",style=bold];
"MUC7_S" -> "FLG2_S" [color="0 0 0.2408977556109726", label="3.75",style=bold];
"CHRNA4_S" -> "MPPED1_S" [color="0 0 0.2401496259351621", label="3.77",style=bold];
"MYH6_S" -> "SALL3_E" [color="0 0 0.2394014962593516", label="3.77",style=bold];
"PLD5_S" -> "PCLO_S" [color="0 0 0.2386533665835412", label="3.82",style=bold];
"SLC18A3_S" -> "CACNA1S_M" [color="0 0 0.2379052369077307", label="3.85",style=bold];
"MYH8_S" -> "REG1A_M" [color="0 0 0.23715710723192018", label="3.86",style=bold];
"SMYD1_E" -> "MYH1_E" [color="0 0 0.2364089775561098", label="3.86",style=bold];
"NRAP_S" -> "REG1A_E" [color="0 0 0.23566084788029928", label="3.90",style=bold];
"MYH7_E" -> "XIRP2_E" [color="0 0 0.23491271820448878", label="3.91",style=bold];
"CSMD1_S" -> "SLC18A3_E" [color="0 0 0.23416458852867827", label="3.93",style=bold];
"PLD5_S" -> "FBN3_S" [color="0 0 0.23341645885286788", label="3.94",style=bold];
"ZIC1_S" -> "ZIC1_M" [color="0 0 0.23266832917705738", label="3.94",style=bold];
"MYH2_S" -> "PLD5_M" [color="0 0 0.23192019950124687", label="3.95",style=bold];
"KRT38_S" -> "MYOG_M" [color="0 0 0.23117206982543648", label="3.98",style=bold];
"SMYD1_E" -> "PKHD1L1_S" [color="0 0 0.23042394014962597", label="4.02",style=bold];
"NEB_S" -> "LRP1B_E" [color="0 0 0.22967581047381547", label="4.02",style=bold];
"MYH7_S" -> "LRP1B_M" [color="0 0 0.22892768079800496", label="4.05",style=bold];
"MPPED1_S" -> "NLRP11_S" [color="0 0 0.22817955112219457", label="4.06",style=bold];
"WDR49_S" -> "FLG_M" [color="0 0 0.22743142144638406", label="4.09",style=bold];
"CACNA1S_E" -> "PAK7_E" [color="0 0 0.22668329177057356", label="4.11",style=bold];
"MYH1_S" -> "ASB4_E" [color="0 0 0.22593516209476316", label="4.11",style=bold];
"GRIN2B_S" -> "CHRNA4_E" [color="0 0 0.22518703241895266", label="4.12",style=bold];
"PPP1R3A_S" -> "ZNF280A_S" [color="0 0 0.22443890274314215", label="4.14",style=bold];
"NLRP11_M" -> "CNTNAP5_M" [color="0 0 0.22369077306733165", label="4.15",style=bold];
"MYH7_S" -> "COL11A1_M" [color="0 0 0.22294264339152126", label="4.15",style=bold];
"MPPED1_S" -> "MUC7_S" [color="0 0 0.22219451371571075", label="4.17",style=bold];
"CHRNA4_S" -> "MUC7_S" [color="0 0 0.22144638403990025", label="4.17",style=bold];
"TMEM132D_S" -> "LRP1B_E" [color="0 0 0.22069825436408974", label="4.18",style=bold];
"FBN3_S" -> "FBN3_M" [color="0 0 0.21995012468827935", label="4.22",style=bold];
"NEB_E" -> "ZIC1_E" [color="0 0 0.21920199501246884", label="4.25",style=bold];
"CNTNAP5_S" -> "FREM2_M" [color="0 0 0.21845386533665834", label="4.25",style=bold];
"MYH2_M" -> "CHRNA4_E" [color="0 0 0.21770573566084794", label="4.27",style=bold];
"C6_S" -> "ASB4_E" [color="0 0 0.21695760598503744", label="4.29",style=bold];
"FBN3_S" -> "ZNF280A_E" [color="0 0 0.21620947630922693", label="4.30",style=bold];
"TMEM132D_S" -> "MYH8_E" [color="0 0 0.21546134663341643", label="4.31",style=bold];
"MUC7_S" -> "MYH1_S" [color="0 0 0.21471321695760603", label="4.33",style=bold];
"FLG2_S" -> "WDR49_M" [color="0 0 0.21396508728179553", label="4.35",style=bold];
"PKHD1_S" -> "MYH2_S" [color="0 0 0.21321695760598502", label="4.38",style=bold];
"FLG2_M" -> "PKHD1_E" [color="0 0 0.21246882793017463", label="4.39",style=bold];
"NRAP_E" -> "ASB4_E" [color="0 0 0.21172069825436413", label="4.46",style=bold];
"LCT_S" -> "NLRP11_S" [color="0 0 0.21097256857855362", label="4.53",style=bold];
"PKHD1L1_S" -> "MUC16_E" [color="0 0 0.21022443890274312", label="4.57",style=bold];
"PPP1R3A_S" -> "REG1A_S" [color="0 0 0.20947630922693272", label="4.59",style=bold];
"TEX15_S" -> "DUSP27_S" [color="0 0 0.20872817955112222", label="4.59",style=bold];
"SALL3_S" -> "MPPED1_E" [color="0 0 0.2079800498753117", label="4.63",style=bold];
"MYH13_S" -> "PPP1R3A_M" [color="0 0 0.20723192019950132", label="4.63",style=bold];
"TRDN_S" -> "ZIC1_S" [color="0 0 0.2064837905236908", label="4.64",style=bold];
"CACNA1S_S" -> "PKHD1_S" [color="0 0 0.2057356608478803", label="4.66",style=bold];
"MYH8_S" -> "WDR49_M" [color="0 0 0.2049875311720698", label="4.69",style=bold];
"CSRP3_E" -> "ASB5_E" [color="0 0 0.2042394014962594", label="4.73",style=bold];
"TMEM132D_M" -> "MUC7_M" [color="0 0 0.2034912718204489", label="4.76",style=bold];
"C6_S" -> "FLG_S" [color="0 0 0.2027431421446384", label="4.78",style=bold];
"MYH1_S" -> "COL11A1_S" [color="0 0 0.2019950124688279", label="4.78",style=bold];
"PCLO_E" -> "CSMD3_E" [color="0 0 0.2012468827930175", label="4.87",style=bold];
"MUC4_S" -> "ZIC1_M" [color="0 0 0.200498753117207", label="4.88",style=bold];
"FREM2_S" -> "ZNF280A_E" [color="0 0 0.1997506234413965", label="4.93",style=bold];
"MYBPC1_M" -> "CSMD1_M" [color="0 0 0.1990024937655861", label="4.95",style=bold];
"ZNF536_S" -> "PCLO_S" [color="0 0 0.1982543640897756", label="4.96",style=bold];
"ZNF536_S" -> "CSMD3_S" [color="0 0 0.1975062344139651", label="4.99",style=bold];
"GRIN2B_S" -> "FREM2_M" [color="0 0 0.19675810473815458", label="4.99",style=bold];
"CDH10_S" -> "LRP1B_E" [color="0 0 0.1960099750623442", label="5.04",style=bold];
"CSMD1_S" -> "Tumor_Status" [color="0 0 0.19526184538653368", label="5.07",style=bold];
"PPP1R3A_S" -> "MYH13_S" [color="0 0 0.19451371571072318", label="5.07",style=bold];
"FREM2_S" -> "CHRNA4_E" [color="0 0 0.19376558603491278", label="5.08",style=bold];
"MYBPC1_S" -> "MYH7_M" [color="0 0 0.19301745635910228", label="5.09",style=bold];
"TRDN_S" -> "NRAP_S" [color="0 0 0.19226932668329177", label="5.10",style=bold];
"MYH7_E" -> "ZIC1_E" [color="0 0 0.19152119700748127", label="5.12",style=bold];
"PLD5_S" -> "DUSP27_S" [color="0 0 0.19077306733167088", label="5.14",style=bold];
"ASB5_S" -> "PCLO_M" [color="0 0 0.19002493765586037", label="5.23",style=bold];
"NEB_S" -> "ZNF280A_E" [color="0 0 0.18927680798004987", label="5.27",style=bold];
"SALL3_S" -> "SIX6_E" [color="0 0 0.18852867830423947", label="5.29",style=bold];
"NLRP11_S" -> "NLRP11_E" [color="0 0 0.18778054862842897", label="5.30",style=bold];
"FLG2_S" -> "LRP1B_E" [color="0 0 0.18703241895261846", label="5.34",style=bold];
"ZIC1_M" -> "SLC18A3_M" [color="0 0 0.18628428927680796", label="5.41",style=bold];
"MUC7_M" -> "PCLO_M" [color="0 0 0.18553615960099756", label="5.46",style=bold];
"MYOG_E" -> "MYH8_E" [color="0 0 0.18478802992518706", label="5.47",style=bold];
"SIX6_M" -> "SALL3_M" [color="0 0 0.18403990024937655", label="5.50",style=bold];
"CSRP3_E" -> "NEB_E" [color="0 0 0.18329177057356616", label="5.53",style=bold];
"SIX6_M" -> "SALL3_E" [color="0 0 0.18254364089775565", label="5.56",style=bold];
"ZNF536_S" -> "CNTNAP5_E" [color="0 0 0.18179551122194515", label="5.61",style=bold];
"TMEM132D_M" -> "CSMD1_M" [color="0 0 0.18104738154613464", label="5.66",style=bold];
"MYH13_S" -> "CSMD1_S" [color="0 0 0.18029925187032425", label="5.71",style=bold];
"PKHD1L1_M" -> "GABRA2_M" [color="0 0 0.17955112219451375", label="5.73",style=bold];
"KRT38_M" -> "COL11A1_S" [color="0 0 0.17880299251870324", label="5.80",style=bold];
"FREM2_E" -> "SLC7A14_E" [color="0 0 0.17805486284289274", label="5.80",style=bold];
"CSRP3_E" -> "XIRP2_E" [color="0 0 0.17730673316708234", label="5.83",style=bold];
"PKHD1_M" -> "TRDN_M" [color="0 0 0.17655860349127184", label="5.91",style=bold];
"LRP1B_E" -> "CDH10_E" [color="0 0 0.17581047381546133", label="5.91",style=bold];
"PPP1R3A_S" -> "MYH8_S" [color="0 0 0.17506234413965094", label="6.00",style=bold];
"CDH10_S" -> "LRP1B_S" [color="0 0 0.17431421446384043", label="6.21",style=bold];
"MUC16_S" -> "FLG_S" [color="0 0 0.17356608478802993", label="6.27",style=bold];
"CNTN5_S" -> "NEB_M" [color="0 0 0.17281795511221942", label="6.36",style=bold];
"CSRP3_E" -> "TRDN_E" [color="0 0 0.17206982543640903", label="6.43",style=bold];
"MUC7_M" -> "MUC16_S" [color="0 0 0.17132169576059852", label="6.48",style=bold];
"REG1A_S" -> "FLG_S" [color="0 0 0.17057356608478802", label="6.48",style=bold];
"NRAP_M" -> "TTN_M" [color="0 0 0.16982543640897763", label="6.52",style=bold];
"ASB4_M" -> "PPP1R3A_M" [color="0 0 0.16907730673316712", label="6.54",style=bold];
"PLD5_S" -> "CNTNAP5_S" [color="0 0 0.16832917705735662", label="6.63",style=bold];
"PKHD1_M" -> "FLG_M" [color="0 0 0.1675810473815461", label="6.66",style=bold];
"MYOG_E" -> "CHRND_E" [color="0 0 0.16683291770573572", label="6.69",style=bold];
"MYBPC1_E" -> "MYH7_E" [color="0 0 0.1660847880299252", label="6.72",style=bold];
"GABRA2_M" -> "MYH4_M" [color="0 0 0.1653366583541147", label="6.74",style=bold];
"TEX15_M" -> "COL11A1_E" [color="0 0 0.1645885286783043", label="6.75",style=bold];
"MYH6_S" -> "PKHD1_S" [color="0 0 0.1638403990024938", label="6.75",style=bold];
"MYH8_S" -> "CSRP3_M" [color="0 0 0.1630922693266833", label="6.76",style=bold];
"MYH7_E" -> "PAK7_E" [color="0 0 0.1623441396508728", label="6.77",style=bold];
"SMYD1_M" -> "MYH6_M" [color="0 0 0.1615960099750624", label="6.87",style=bold];
"ZIC4_E" -> "PKHD1L1_M" [color="0 0 0.1608478802992519", label="6.91",style=bold];
"MYL1_M" -> "CHRNA4_M" [color="0 0 0.1600997506234414", label="6.99",style=bold];
"CSMD3_S" -> "LRP1B_S" [color="0 0 0.1593516209476309", label="7.00",style=bold];
"PKHD1_S" -> "TTN_S" [color="0 0 0.1586034912718205", label="7.03",style=bold];
"CHRNA4_S" -> "KRT38_S" [color="0 0 0.15785536159601", label="7.03",style=bold];
"FBN3_E" -> "MUC7_E" [color="0 0 0.15710723192019949", label="7.09",style=bold];
"CSRP3_E" -> "SMYD1_E" [color="0 0 0.1563591022443891", label="7.12",style=bold];
"MYH4_E" -> "ZIC1_E" [color="0 0 0.1556109725685786", label="7.15",style=bold];
"CSRP3_E" -> "MYOG_E" [color="0 0 0.15486284289276808", label="7.24",style=bold];
"FREM2_E" -> "ZIC1_E" [color="0 0 0.15411471321695758", label="7.27",style=bold];
"GABRA2_M" -> "CNTNAP5_M" [color="0 0 0.15336658354114718", label="7.29",style=bold];
"MYL1_M" -> "FLG2_M" [color="0 0 0.15261845386533668", label="7.29",style=bold];
"PLD5_M" -> "FREM2_M" [color="0 0 0.15187032418952617", label="7.40",style=bold];
"TTN_M" -> "KRT38_M" [color="0 0 0.15112219451371578", label="7.41",style=bold];
"CNTN5_S" -> "CSRP3_E" [color="0 0 0.15037406483790527", label="7.46",style=bold];
"NLRP11_M" -> "SALL3_M" [color="0 0 0.14962593516209477", label="7.55",style=bold];
"CHRND_M" -> "CACNA1S_M" [color="0 0 0.14887780548628426", label="7.57",style=bold];
"CACNA1S_M" -> "NEB_M" [color="0 0 0.14812967581047387", label="7.62",style=bold];
"MYH2_S" -> "LCT_E" [color="0 0 0.14738154613466337", label="7.65",style=bold];
"DUSP27_M" -> "LCT_M" [color="0 0 0.14663341645885286", label="7.66",style=bold];
"LRP1B_M" -> "PPP1R3A_M" [color="0 0 0.14588528678304247", label="7.67",style=bold];
"PKHD1L1_M" -> "CSRP3_M" [color="0 0 0.14513715710723196", label="7.69",style=bold];
"FBN3_E" -> "CHRNA4_E" [color="0 0 0.14438902743142146", label="7.73",style=bold];
"NRAP_E" -> "COL11A1_E" [color="0 0 0.14364089775561095", label="7.76",style=bold];
"PAK7_E" -> "ASB5_M" [color="0 0 0.14289276807980056", label="7.77",style=bold];
"ASB4_M" -> "MYL1_M" [color="0 0 0.14214463840399005", label="7.79",style=bold];
"CACNA1S_M" -> "ASB5_M" [color="0 0 0.14139650872817955", label="7.84",style=bold];
"PPP1R3A_E" -> "C6_E" [color="0 0 0.14064837905236904", label="7.94",style=bold];
"MYH2_M" -> "CCDC129_M" [color="0 0 0.13990024937655865", label="8.03",style=bold];
"PKHD1_M" -> "ASB4_M" [color="0 0 0.13915211970074814", label="8.13",style=bold];
"SLC7A14_E" -> "PKHD1L1_E" [color="0 0 0.13840399002493764", label="8.18",style=bold];
"MYOG_E" -> "SIX6_E" [color="0 0 0.13765586034912725", label="8.32",style=bold];
"CSMD3_S" -> "PCLO_S" [color="0 0 0.13690773067331674", label="8.46",style=bold];
"SMYD1_E" -> "ASB5_E" [color="0 0 0.13615960099750624", label="8.61",style=bold];
"MYH1_E" -> "PPP1R3A_E" [color="0 0 0.13541147132169573", label="8.66",style=bold];
"MYH4_M" -> "CCDC129_M" [color="0 0 0.13466334164588534", label="8.78",style=bold];
"CHRND_E" -> "CSMD3_S" [color="0 0 0.13391521197007483", label="8.79",style=bold];
"ASB5_E" -> "MYH8_E" [color="0 0 0.13316708229426433", label="8.83",style=bold];
"PAK7_E" -> "MUC4_E" [color="0 0 0.13241895261845393", label="8.89",style=bold];
"NLRP11_S" -> "PKHD1L1_S" [color="0 0 0.13167082294264343", label="8.90",style=bold];
"PKHD1L1_E" -> "GABRA2_E" [color="0 0 0.13092269326683292", label="8.93",style=bold];
"PLD5_M" -> "ZNF536_M" [color="0 0 0.13017456359102242", label="9.20",style=bold];
"MYH13_E" -> "CSMD3_S" [color="0 0 0.12942643391521202", label="9.26",style=bold];
"MYH1_E" -> "DUSP27_E" [color="0 0 0.12867830423940152", label="9.27",style=bold];
"XIRP2_E" -> "MYH6_E" [color="0 0 0.12793017456359101", label="9.35",style=bold];
"KRT38_M" -> "SLC7A14_M" [color="0 0 0.12718204488778062", label="9.41",style=bold];
"CSRP3_E" -> "MYH2_E" [color="0 0 0.12643391521197012", label="9.44",style=bold];
"PKHD1L1_E" -> "PKHD1L1_M" [color="0 0 0.1256857855361596", label="9.45",style=bold];
"MUC4_M" -> "CHRND_M" [color="0 0 0.1249376558603491", label="9.47",style=bold];
"PAK7_M" -> "ZNF280A_M" [color="0 0 0.12418952618453871", label="9.62",style=bold];
"TMEM132D_M" -> "LCT_M" [color="0 0 0.12344139650872821", label="9.64",style=bold];
"PKHD1L1_E" -> "NLRP11_E" [color="0 0 0.1226932668329177", label="9.75",style=bold];
"CSMD1_E" -> "KRT38_E" [color="0 0 0.1219451371571072", label="9.81",style=bold];
"ZNF536_M" -> "GRIN2B_M" [color="0 0 0.1211970074812968", label="9.85",style=bold];
"CNTNAP5_M" -> "GRIN2B_M" [color="0 0 0.1204488778054863", label="9.86",style=bold];
"TRDN_E" -> "CACNA1S_E" [color="0 0 0.1197007481296758", label="9.98",style=bold];
"ASB4_M" -> "Tumor_Status" [color="0 0 0.1189526184538654", label="10.05",style=bold];
"NLRP11_M" -> "MYH4_M" [color="0 0 0.1182044887780549", label="10.07",style=bold];
"GABRA2_M" -> "CDH10_M" [color="0 0 0.11745635910224439", label="10.17",style=bold];
"MYH7_M" -> "LRP1B_M" [color="0 0 0.11670822942643388", label="10.22",style=bold];
"PAK7_M" -> "CNTN5_M" [color="0 0 0.11596009975062349", label="10.23",style=bold];
"MYL1_M" -> "ZNF280A_M" [color="0 0 0.11521197007481299", label="10.29",style=bold];
"MYL1_M" -> "TRDN_M" [color="0 0 0.11446384039900248", label="10.41",style=bold];
"MYH7_M" -> "MPPED1_M" [color="0 0 0.11371571072319209", label="10.41",style=bold];
"TTN_M" -> "MYL1_M" [color="0 0 0.11296758104738158", label="10.57",style=bold];
"TTN_E" -> "ZNF536_E" [color="0 0 0.11221945137157108", label="10.62",style=bold];
"MYH7_M" -> "CSRP3_M" [color="0 0 0.11147132169576057", label="10.64",style=bold];
"CACNA1S_M" -> "DUSP27_M" [color="0 0 0.11072319201995018", label="10.79",style=bold];
"PAK7_M" -> "ZNF280A_E" [color="0 0 0.10997506234413967", label="10.98",style=bold];
"PKHD1_M" -> "C6_M" [color="0 0 0.10922693266832917", label="11.23",style=bold];
"MYL1_E" -> "SMYD1_E" [color="0 0 0.10847880299251877", label="11.26",style=bold];
"MYH13_E" -> "COL11A1_E" [color="0 0 0.10773067331670827", label="11.28",style=bold];
"ZNF536_E" -> "TMEM132D_E" [color="0 0 0.10698254364089776", label="11.45",style=bold];
"NEB_E" -> "PAK7_E" [color="0 0 0.10623441396508726", label="11.91",style=bold];
"CNTNAP5_M" -> "KRT38_M" [color="0 0 0.10548628428927687", label="12.13",style=bold];
"WDR49_E" -> "LCT_E" [color="0 0 0.10473815461346636", label="12.45",style=bold];
"SLC18A3_M" -> "SIX6_M" [color="0 0 0.10399002493765586", label="12.46",style=bold];
"FLG2_M" -> "CSMD3_E" [color="0 0 0.10324189526184535", label="12.49",style=bold];
"ZNF536_M" -> "MUC7_M" [color="0 0 0.10249376558603496", label="12.70",style=bold];
"REG1A_E" -> "FREM2_E" [color="0 0 0.10174563591022445", label="12.73",style=bold];
"MYH8_M" -> "FLG2_M" [color="0 0 0.10099750623441395", label="12.76",style=bold];
"MYH7_M" -> "NRAP_M" [color="0 0 0.10024937655860355", label="12.79",style=bold];
"MYH7_M" -> "LCT_M" [color="0 0 0.09950124688279305", label="13.00",style=bold];
"MYH1_M" -> "MYH2_M" [color="0 0 0.09875311720698254", label="13.05",style=bold];
"MYL1_M" -> "MYH4_M" [color="0 0 0.09800498753117204", label="13.09",style=bold];
"NEB_E" -> "TRDN_E" [color="0 0 0.09725685785536164", label="13.22",style=bold];
"PKHD1_M" -> "NEB_M" [color="0 0 0.09650872817955114", label="13.28",style=bold];
"NRAP_M" -> "MPPED1_M" [color="0 0 0.09576059850374063", label="13.41",style=bold];
"CSRP3_M" -> "KRT38_M" [color="0 0 0.09501246882793024", label="13.48",style=bold];
"CSMD1_M" -> "WDR49_M" [color="0 0 0.09426433915211974", label="13.68",style=bold];
"MPPED1_E" -> "CSMD1_E" [color="0 0 0.09351620947630923", label="13.68",style=bold];
"MYH1_M" -> "MUC16_M" [color="0 0 0.09276807980049873", label="13.79",style=bold];
"MUC7_M" -> "Tumor_Status" [color="0 0 0.09201995012468833", label="13.98",style=bold];
"SIX6_M" -> "COL11A1_M" [color="0 0 0.09127182044887783", label="14.00",style=bold];
"MYL1_E" -> "MYH1_E" [color="0 0 0.09052369077306732", label="14.27",style=bold];
"PPP1R3A_E" -> "NEB_E" [color="0 0 0.08977556109725693", label="14.32",style=bold];
"PKHD1_M" -> "PLD5_E" [color="0 0 0.08902743142144642", label="14.40",style=bold];
"MYBPC1_M" -> "DUSP27_M" [color="0 0 0.08827930174563592", label="14.45",style=bold];
"SLC7A14_E" -> "Tumor_Status" [color="0 0 0.08753117206982541", label="14.48",style=bold];
"CDH10_M" -> "MUC16_E" [color="0 0 0.08678304239401502", label="14.70",style=bold];
"MYH1_E" -> "COL11A1_E" [color="0 0 0.08603491271820451", label="14.93",style=bold];
"PKHD1_M" -> "NRAP_M" [color="0 0 0.08528678304239401", label="15.01",style=bold];
"TRDN_E" -> "MYBPC1_E" [color="0 0 0.0845386533665835", label="15.15",style=bold];
"PAK7_M" -> "FREM2_M" [color="0 0 0.08379052369077311", label="15.24",style=bold];
"PAK7_E" -> "PKHD1L1_M" [color="0 0 0.0830423940149626", label="15.27",style=bold];
"XIRP2_M" -> "MUC7_M" [color="0 0 0.0822942643391521", label="15.58",style=bold];
"CNTNAP5_E" -> "NLRP11_E" [color="0 0 0.08154613466334171", label="15.84",style=bold];
"MPPED1_M" -> "MUC4_M" [color="0 0 0.0807980049875312", label="15.96",style=bold];
"CSRP3_M" -> "PKHD1_M" [color="0 0 0.0800498753117207", label="16.01",style=bold];
"LCT_E" -> "GRIN2B_E" [color="0 0 0.07930174563591019", label="16.18",style=bold];
"PAK7_E" -> "REG1A_E" [color="0 0 0.0785536159600998", label="16.21",style=bold];
"NLRP11_M" -> "TMEM132D_M" [color="0 0 0.0778054862842893", label="16.24",style=bold];
"CNTNAP5_E" -> "ZNF280A_E" [color="0 0 0.07705735660847879", label="16.24",style=bold];
"SMYD1_M" -> "ASB5_M" [color="0 0 0.0763092269326684", label="16.27",style=bold];
"SMYD1_E" -> "PPP1R3A_E" [color="0 0 0.07556109725685789", label="16.38",style=bold];
"XIRP2_M" -> "REG1A_M" [color="0 0 0.07481296758104738", label="16.47",style=bold];
"CHRND_M" -> "SMYD1_M" [color="0 0 0.07406483790523688", label="16.94",style=bold];
"DUSP27_E" -> "TTN_E" [color="0 0 0.07331670822942649", label="16.95",style=bold];
"ASB4_M" -> "LRP1B_M" [color="0 0 0.07256857855361598", label="17.04",style=bold];
"CNTNAP5_E" -> "TEX15_E" [color="0 0 0.07182044887780548", label="17.11",style=bold];
"PPP1R3A_E" -> "ASB4_E" [color="0 0 0.07107231920199508", label="17.11",style=bold];
"MUC4_E" -> "MUC16_E" [color="0 0 0.07032418952618458", label="17.30",style=bold];
"CNTNAP5_E" -> "GABRA2_M" [color="0 0 0.06957605985037407", label="17.33",style=bold];
"PAK7_M" -> "DUSP27_M" [color="0 0 0.06882793017456357", label="17.35",style=bold];
"SMYD1_M" -> "MYH8_M" [color="0 0 0.06807980049875317", label="17.37",style=bold];
"WDR49_E" -> "MUC4_E" [color="0 0 0.06733167082294267", label="17.80",style=bold];
"PKHD1L1_E" -> "C6_E" [color="0 0 0.06658354114713216", label="17.83",style=bold];
"NRAP_M" -> "SMYD1_M" [color="0 0 0.06583541147132177", label="18.18",style=bold];
"CCDC129_E" -> "MUC7_E" [color="0 0 0.06508728179551126", label="18.18",style=bold];
"ASB4_M" -> "TTN_M" [color="0 0 0.06433915211970076", label="18.36",style=bold];
"LRP1B_M" -> "NLRP11_M" [color="0 0 0.06359102244389025", label="18.52",style=bold];
"SMYD1_M" -> "CACNA1S_M" [color="0 0 0.06284289276807986", label="18.67",style=bold];
"ZNF280A_E" -> "CHRNA4_E" [color="0 0 0.062094763092269356", label="18.72",style=bold];
"LRP1B_M" -> "C6_M" [color="0 0 0.06134663341645885", label="18.81",style=bold];
"MYH7_M" -> "MYOG_M" [color="0 0 0.060598503740648346", label="18.85",style=bold];
"GABRA2_M" -> "ZIC1_M" [color="0 0 0.05985037406483795", label="18.97",style=bold];
"PAK7_M" -> "PLD5_M" [color="0 0 0.05910224438902745", label="19.12",style=bold];
"PAK7_M" -> "PCLO_M" [color="0 0 0.05835411471321694", label="19.21",style=bold];
"CACNA1S_M" -> "KRT38_M" [color="0 0 0.05760598503740655", label="19.33",style=bold];
"MPPED1_M" -> "CHRND_M" [color="0 0 0.056857855361596044", label="19.43",style=bold];
"CACNA1S_E" -> "TTN_E" [color="0 0 0.05610972568578554", label="19.64",style=bold];
"ASB4_M" -> "MYBPC1_M" [color="0 0 0.055361596009975034", label="19.64",style=bold];
"MYH4_M" -> "MYH1_M" [color="0 0 0.05461346633416464", label="19.77",style=bold];
"MYBPC1_M" -> "NEB_M" [color="0 0 0.053865336658354135", label="19.92",style=bold];
"NEB_E" -> "FREM2_E" [color="0 0 0.05311720698254363", label="19.92",style=bold];
"PAK7_E" -> "SLC18A3_E" [color="0 0 0.052369077306733236", label="20.45",style=bold];
"CNTNAP5_E" -> "PKHD1_E" [color="0 0 0.05162094763092273", label="20.53",style=bold];
"MYBPC1_M" -> "MUC4_M" [color="0 0 0.050872817955112226", label="20.56",style=bold];
"PCLO_E" -> "PCLO_M" [color="0 0 0.05012468827930172", label="20.59",style=bold];
"MYH7_E" -> "MYH6_E" [color="0 0 0.04937655860349133", label="20.68",style=bold];
"MYH4_M" -> "XIRP2_M" [color="0 0 0.04862842892768082", label="20.71",style=bold];
"TTN_M" -> "LRP1B_M" [color="0 0 0.04788029925187032", label="20.81",style=bold];
"CSMD1_E" -> "LCT_E" [color="0 0 0.047132169576059924", label="20.84",style=bold];
"SMYD1_E" -> "DUSP27_E" [color="0 0 0.04638403990024942", label="21.09",style=bold];
"PAK7_E" -> "PLD5_E" [color="0 0 0.045635910224438914", label="21.10",style=bold];
"CSMD1_M" -> "MUC16_M" [color="0 0 0.04488778054862841", label="21.42",style=bold];
"MYBPC1_M" -> "PCLO_E" [color="0 0 0.044139650872818015", label="21.44",style=bold];
"ASB4_M" -> "CCDC129_M" [color="0 0 0.04339152119700751", label="21.76",style=bold];
"PKHD1_M" -> "WDR49_M" [color="0 0 0.042643391521197005", label="22.20",style=bold];
"MYH8_M" -> "MYH13_M" [color="0 0 0.0418952618453865", label="22.20",style=bold];
"PPP1R3A_E" -> "MYH4_E" [color="0 0 0.041147132169576106", label="22.49",style=bold];
"SALL3_M" -> "SALL3_E" [color="0 0 0.0403990024937656", label="22.66",style=bold];
"ASB5_E" -> "MYOG_E" [color="0 0 0.039650872817955096", label="22.76",style=bold];
"CHRNA4_M" -> "NLRP11_M" [color="0 0 0.0389027431421447", label="22.95",style=bold];
"TEX15_E" -> "TEX15_M" [color="0 0 0.0381546134663342", label="23.15",style=bold];
"SMYD1_E" -> "MYBPC1_E" [color="0 0 0.03740648379052369", label="23.16",style=bold];
"PAK7_E" -> "CNTNAP5_E" [color="0 0 0.03665835411471319", label="23.46",style=bold];
"PLD5_M" -> "CNTN5_M" [color="0 0 0.035910224438902794", label="23.52",style=bold];
"FLG2_M" -> "FLG_M" [color="0 0 0.03516209476309229", label="23.73",style=bold];
"CDH10_M" -> "PLD5_M" [color="0 0 0.034413965087281784", label="24.17",style=bold];
"CSMD1_E" -> "WDR49_E" [color="0 0 0.03366583541147139", label="24.39",style=bold];
"MPPED1_M" -> "FBN3_M" [color="0 0 0.032917705735660885", label="24.48",style=bold];
"GRIN2B_M" -> "CSMD3_M" [color="0 0 0.03216957605985038", label="24.84",style=bold];
"FBN3_E" -> "CCDC129_E" [color="0 0 0.031421446384039875", label="25.10",style=bold];
"PCLO_E" -> "CSMD1_E" [color="0 0 0.03067331670822948", label="25.47",style=bold];
"MYH4_E" -> "MYH13_E" [color="0 0 0.029925187032418976", label="25.85",style=bold];
"PKHD1L1_M" -> "CHRNA4_M" [color="0 0 0.02917705735660847", label="26.38",style=bold];
"CACNA1S_M" -> "MYOG_M" [color="0 0 0.028428927680798077", label="26.73",style=bold];
"PKHD1L1_E" -> "ZNF536_E" [color="0 0 0.027680798004987572", label="26.90",style=bold];
"GABRA2_M" -> "SLC18A3_M" [color="0 0 0.026932668329177067", label="27.70",style=bold];
"PKHD1L1_M" -> "PAK7_M" [color="0 0 0.026184538653366562", label="28.11",style=bold];
"SLC18A3_M" -> "SALL3_M" [color="0 0 0.02543640897755617", label="28.34",style=bold];
"PAK7_E" -> "PCLO_E" [color="0 0 0.024688279301745664", label="28.64",style=bold];
"SLC7A14_M" -> "COL11A1_M" [color="0 0 0.02394014962593516", label="29.35",style=bold];
"PCLO_E" -> "LRP1B_E" [color="0 0 0.023192019950124654", label="29.44",style=bold];
"CHRND_M" -> "FBN3_M" [color="0 0 0.02244389027431426", label="29.86",style=bold];
"MPPED1_E" -> "FBN3_E" [color="0 0 0.021695760598503755", label="31.48",style=bold];
"CNTNAP5_E" -> "CNTN5_E" [color="0 0 0.02094763092269325", label="32.51",style=bold];
"MYH4_M" -> "ZIC1_M" [color="0 0 0.020199501246882856", label="32.79",style=bold];
"CSRP3_M" -> "MYBPC1_M" [color="0 0 0.01945137157107235", label="33.10",style=bold];
"MYH1_M" -> "REG1A_M" [color="0 0 0.018703241895261846", label="33.46",style=bold];
"CDH10_M" -> "PAK7_M" [color="0 0 0.01795511221945134", label="33.84",style=bold];
"PKHD1L1_M" -> "ASB4_M" [color="0 0 0.017206982543640947", label="34.16",style=bold];
"TMEM132D_M" -> "ZNF536_M" [color="0 0 0.016458852867830442", label="35.18",style=bold];
"CSRP3_E" -> "MYH7_E" [color="0 0 0.015710723192019937", label="35.61",style=bold];
"GABRA2_M" -> "CSMD3_M" [color="0 0 0.014962593516209544", label="35.96",style=bold];
"NEB_E" -> "LRP1B_E" [color="0 0 0.014214463840399039", label="36.20",style=bold];
"PPP1R3A_E" -> "CHRND_E" [color="0 0 0.013466334164588534", label="38.13",style=bold];
"LRP1B_M" -> "XIRP2_M" [color="0 0 0.012718204488778029", label="39.50",style=bold];
"MYH4_M" -> "TEX15_M" [color="0 0 0.011970074812967635", label="40.22",style=bold];
"MYH7_M" -> "PKHD1_M" [color="0 0 0.01122194513715713", label="42.72",style=bold];
"ZIC4_M" -> "SIX6_M" [color="0 0 0.010473815461346625", label="43.57",style=bold];
"TMEM132D_M" -> "MYH1_M" [color="0 0 0.009725685785536231", label="44.74",style=bold];
"TMEM132D_M" -> "MYH13_M" [color="0 0 0.008977556109725726", label="44.84",style=bold];
"XIRP2_E" -> "NRAP_E" [color="0 0 0.008229426433915221", label="46.17",style=bold];
"CNTNAP5_M" -> "TMEM132D_M" [color="0 0 0.007481296758104716", label="46.48",style=bold];
"TMEM132D_M" -> "CDH10_M" [color="0 0 0.006733167082294322", label="47.80",style=bold];
"CSMD3_M" -> "SLC7A14_M" [color="0 0 0.0059850374064838174", label="48.07",style=bold];
"PAK7_E" -> "MPPED1_E" [color="0 0 0.0052369077306733125", label="48.15",style=bold];
"MYH4_M" -> "MYH8_M" [color="0 0 0.0044887780548628076", label="48.28",style=bold];
"PKHD1L1_M" -> "MYH7_M" [color="0 0 0.0037406483790524137", label="66.76",style=bold];
"ZIC1_E" -> "ZIC4_E" [color="0 0 0.0029925187032419087", label="78.79",style=bold];
"FLG_E" -> "FLG2_E" [color="0 0 0.0022443890274314038", label="80.85",style=bold];
"MYH7_M" -> "MYH6_M" [color="0 0 0.0014962593516210099", label="81.78",style=bold];
"ZIC1_M" -> "ZIC4_M" [color="0 0 0.0007481296758105049", label="98.45",style=bold];
"CSRP3_E" -> "MYL1_E" [color="0 0 0.0", label="200.33",style=bold];
}
```
