## Supplementary material for "New analysis framework incorporating mixed mutual information and scalable Bayesian networks for multimodal high dimensional genomic and epigenomic cancer data": Suppl Table 10

```
digraph G{
 ratio=fill;
 node [shape=box, style=rounded];
 edge [arrowhead=none];
"ABCA13_S";
"ABCA13_E";
"ABCA13_M";
"ADAMTS20_S";
"ADAMTS20_E";
"ADAMTS20_M";
"ATP13A4_S";
"ATP13A4_E";
"ATP13A4_M";
"BEST2_S";
"BEST2_E";
"BEST2_M";
"CCDC67_S";
"CCDC67_E";
"CCDC67_M";
"CDH19_S";
"CDH19_E";
"CDH19_M";
"CDKN2A_S";
"CDKN2A_E";
"CDKN2A_M";
"CNTN5_S";
"CNTN5_E";
"CNTN5_M";
"CNTNAP5_S";
"CNTNAP5_E";
"CNTNAP5_M";
"COL11A1_S";
"COL11A1_E";
"COL11A1_M";
"CPNE6_S";
"CPNE6_E";
"CPNE6_M";
"CR2_S";
"CR2_E";
"CR2_M";
"CSMD1_S";
"CSMD1_E";
"CSMD1_M";
"CSMD3_S";
"CSMD3_E";
"CSMD3_M";
"DNAH11_S";
"DNAH11_E";
"DNAH11_M";
"DSCAM_S";
"DSCAM_E";
"DSCAM_M";
"DUSP27_S";
"DUSP27_E";
"DUSP27_M";
"ERBB4_S";
"ERBB4_E";
"ERBB4_M";
"ERN2_S";
"ERN2_E";
"ERN2_M";
"FLG_S";
"FLG_E";
"FLG_M";
"FREM2_S";
"FREM2_E";
"FREM2_M";
"FSTL5_S";
"FSTL5_E";
"FSTL5_M";
"GABRB3_S";
"GABRB3_E";
"GABRB3_M";
"GALNTL6_S";
"GALNTL6_E";
"GALNTL6_M";
"HFM1_S";
"HFM1_E";
"HFM1_M";
"KRT1_S";
"KRT1_E";
"KRT1_M";
"KRT24_S";
"KRT24_E";
"KRT24_M";
"LCE2C_S";
"LCE2C_E";
"LCE2C_M";
"LCT_S";
"LCT_E";
"LCT_M";
"LINGO2_S";
"LINGO2_E";
"LINGO2_M";
"LPA_S";
"LPA_E";
"LPA_M";
"LPPR1_S";
"LPPR1_E";
"LPPR1_M";
"LRAT_S";
"LRAT_E";
"LRAT_M";
"LRP1B_S";
"LRP1B_E";
"LRP1B_M";
"LRRTM4_S";
"LRRTM4_E";
"LRRTM4_M";
"MAB21L2_S";
"MAB21L2_E";
"MAB21L2_M";
"MC5R_S";
"MC5R_E";
"MC5R_M";
"MRGPRX4_S";
"MRGPRX4_E";
"MRGPRX4_M";
"MUC16_S";
"MUC16_E";
"MUC16_M";
"MUC4_S";
"MUC4_E";
"MUC4_M";
"MYBPC2_S";
"MYBPC2_E";
"MYBPC2_M";
"MYH13_S";
"MYH13_E";
"MYH13_M";
"MYH6_S";
"MYH6_E";
"MYH6_M";
"MYH7_S";
"MYH7_E";
"MYH7_M";
"MYO3A_S";
"MYO3A_E";
"MYO3A_M";
"NEB_S";
"NEB_E";
"NEB_M";
"NELL1_S";
"NELL1_E";
"NELL1_M";
"NPFFR2_S";
"NPFFR2_E";
"NPFFR2_M";
"NTSR2_S";
"NTSR2_E";
"NTSR2_M";
"PCDH10_S";
"PCDH10_E";
"PCDH10_M";
"PCDHA10_S";
"PCDHA10_E";
"PCDHA10_M";
"PCDHA12_S";
"PCDHA12_E";
"PCDHA12_M";
"PCLO_S";
"PCLO_E";
"PCLO_M";
"PKHD1L1_S";
"PKHD1L1_E";
"PKHD1L1_M";
"PTPRT_S";
"PTPRT_E";
"PTPRT_M";
"RBP3_S";
"RBP3_E";
"RBP3_M";
"RP1_S";
"RP1_E";
"RP1_M";
"RYR2_S";
"RYR2_E";
"RYR2_M";
"TDRD5_S";
"TDRD5_E";
"TDRD5_M";
"TTN_S";
"TTN_E";
"TTN_M";
"TUBA3C_S";
"TUBA3C_E";
"TUBA3C_M";
"UGT2A1_S";
"UGT2A1_E";
"UGT2A1_M";
"ZIC1_S";
"ZIC1_E";
"ZIC1_M";
"Survival";
"UGT2A1_S" -> "PCDHA10_M" [color="0 0 0.9", label="0.00",style=bold];
"CCDC67_S" -> "ERBB4_M" [color="0 0 0.8993416239941477", label="0.01",style=bold];
"MAB21L2_S" -> "CR2_S" [color="0 0 0.8986832479882956", label="0.01",style=bold];
"KRT1_S" -> "LINGO2_E" [color="0 0 0.8980248719824433", label="0.01",style=bold];
"DUSP27_S" -> "NEB_M" [color="0 0 0.8973664959765911", label="0.01",style=bold];
"MYBPC2_S" -> "ERN2_E" [color="0 0 0.8967081199707388", label="0.02",style=bold];
"MYH7_S" -> "DSCAM_E" [color="0 0 0.8960497439648867", label="0.02",style=bold];
"ZIC1_S" -> "DNAH11_M" [color="0 0 0.8953913679590344", label="0.02",style=bold];
"PCDH10_S" -> "FLG_S" [color="0 0 0.8947329919531821", label="0.03",style=bold];
"MUC4_S" -> "FLG_S" [color="0 0 0.89407461594733", label="0.03",style=bold];
"LCE2C_S" -> "HFM1_S" [color="0 0 0.8934162399414777", label="0.03",style=bold];
"LCE2C_S" -> "CDKN2A_S" [color="0 0 0.8927578639356255", label="0.03",style=bold];
"CNTNAP5_S" -> "PCLO_E" [color="0 0 0.8920994879297732", label="0.03",style=bold];
"MYO3A_S" -> "RYR2_S" [color="0 0 0.8914411119239211", label="0.03",style=bold];
"RBP3_S" -> "PCDH10_M" [color="0 0 0.8907827359180688", label="0.03",style=bold];
"LCT_S" -> "ATP13A4_M" [color="0 0 0.8901243599122165", label="0.03",style=bold];
"GALNTL6_S" -> "DNAH11_S" [color="0 0 0.8894659839063643", label="0.03",style=bold];
"NELL1_S" -> "RP1_E" [color="0 0 0.888807607900512", label="0.04",style=bold];
"UGT2A1_S" -> "TUBA3C_M" [color="0 0 0.8881492318946599", label="0.04",style=bold];
"LCT_S" -> "LCT_M" [color="0 0 0.8874908558888076", label="0.04",style=bold];
"MAB21L2_S" -> "GABRB3_S" [color="0 0 0.8868324798829554", label="0.04",style=bold];
"ATP13A4_S" -> "FLG_E" [color="0 0 0.8861741038771032", label="0.04",style=bold];
"MYH7_S" -> "LPPR1_E" [color="0 0 0.885515727871251", label="0.04",style=bold];
"BEST2_S" -> "FLG_M" [color="0 0 0.8848573518653987", label="0.05",style=bold];
"NTSR2_S" -> "GALNTL6_M" [color="0 0 0.8841989758595464", label="0.05",style=bold];
"DUSP27_S" -> "MUC4_S" [color="0 0 0.8835405998536943", label="0.05",style=bold];
"PCDHA12_S" -> "PCDH10_M" [color="0 0 0.882882223847842", label="0.05",style=bold];
"MYBPC2_S" -> "MUC16_M" [color="0 0 0.8822238478419898", label="0.05",style=bold];
"CCDC67_S" -> "DNAH11_M" [color="0 0 0.8815654718361375", label="0.05",style=bold];
"ADAMTS20_S" -> "CNTN5_E" [color="0 0 0.8809070958302854", label="0.06",style=bold];
"CCDC67_S" -> "GABRB3_M" [color="0 0 0.8802487198244331", label="0.06",style=bold];
"FREM2_S" -> "CNTN5_M" [color="0 0 0.8795903438185808", label="0.06",style=bold];
"LCE2C_S" -> "LRRTM4_E" [color="0 0 0.8789319678127286", label="0.06",style=bold];
"LRRTM4_S" -> "NEB_E" [color="0 0 0.8782735918068764", label="0.06",style=bold];
"NPFFR2_S" -> "FLG_M" [color="0 0 0.8776152158010242", label="0.06",style=bold];
"MC5R_S" -> "MYH6_E" [color="0 0 0.8769568397951719", label="0.06",style=bold];
"CR2_S" -> "FLG_E" [color="0 0 0.8762984637893197", label="0.06",style=bold];
"BEST2_S" -> "TTN_E" [color="0 0 0.8756400877834675", label="0.06",style=bold];
"BEST2_S" -> "PTPRT_M" [color="0 0 0.8749817117776152", label="0.06",style=bold];
"LPPR1_S" -> "KRT24_E" [color="0 0 0.874323335771763", label="0.06",style=bold];
"MYH7_S" -> "CPNE6_M" [color="0 0 0.8736649597659107", label="0.07",style=bold];
"TUBA3C_S" -> "MYBPC2_E" [color="0 0 0.8730065837600586", label="0.07",style=bold];
"PCDH10_S" -> "MAB21L2_E" [color="0 0 0.8723482077542063", label="0.07",style=bold];
"PCDH10_S" -> "FLG_E" [color="0 0 0.8716898317483541", label="0.07",style=bold];
"ABCA13_S" -> "CSMD1_S" [color="0 0 0.8710314557425018", label="0.08",style=bold];
"FSTL5_S" -> "GALNTL6_E" [color="0 0 0.8703730797366496", label="0.08",style=bold];
"NTSR2_S" -> "NTSR2_E" [color="0 0 0.8697147037307974", label="0.08",style=bold];
"CPNE6_S" -> "NEB_E" [color="0 0 0.8690563277249451", label="0.08",style=bold];
"LRAT_S" -> "TTN_E" [color="0 0 0.868397951719093", label="0.09",style=bold];
"MYBPC2_S" -> "PTPRT_M" [color="0 0 0.8677395757132407", label="0.09",style=bold];
"GABRB3_S" -> "KRT1_M" [color="0 0 0.8670811997073885", label="0.09",style=bold];
"DUSP27_S" -> "CNTNAP5_E" [color="0 0 0.8664228237015362", label="0.10",style=bold];
"ATP13A4_S" -> "ERBB4_E" [color="0 0 0.8657644476956841", label="0.10",style=bold];
"CDH19_S" -> "MUC4_E" [color="0 0 0.8651060716898318", label="0.10",style=bold];
"DUSP27_S" -> "CDKN2A_M" [color="0 0 0.8644476956839795", label="0.11",style=bold];
"MYO3A_S" -> "CDKN2A_M" [color="0 0 0.8637893196781273", label="0.11",style=bold];
"UGT2A1_S" -> "GABRB3_E" [color="0 0 0.863130943672275", label="0.11",style=bold];
"GALNTL6_S" -> "MYO3A_E" [color="0 0 0.8624725676664229", label="0.11",style=bold];
"MYO3A_S" -> "NELL1_E" [color="0 0 0.8618141916605706", label="0.11",style=bold];
"NELL1_S" -> "MYH13_E" [color="0 0 0.8611558156547183", label="0.11",style=bold];
"CNTN5_S" -> "KRT24_E" [color="0 0 0.8604974396488662", label="0.11",style=bold];
"MYBPC2_S" -> "FLG_M" [color="0 0 0.8598390636430139", label="0.11",style=bold];
"ZIC1_S" -> "MUC4_M" [color="0 0 0.8591806876371617", label="0.12",style=bold];
"ABCA13_S" -> "COL11A1_E" [color="0 0 0.8585223116313094", label="0.12",style=bold];
"PCDH10_S" -> "HFM1_E" [color="0 0 0.8578639356254573", label="0.12",style=bold];
"ADAMTS20_S" -> "CSMD3_M" [color="0 0 0.857205559619605", label="0.13",style=bold];
"CPNE6_S" -> "CDH19_M" [color="0 0 0.8565471836137528", label="0.13",style=bold];
"TUBA3C_S" -> "DNAH11_E" [color="0 0 0.8558888076079005", label="0.13",style=bold];
"LCE2C_S" -> "LPPR1_M" [color="0 0 0.8552304316020483", label="0.14",style=bold];
"MRGPRX4_S" -> "LPPR1_M" [color="0 0 0.8545720555961961", label="0.14",style=bold];
"LCE2C_S" -> "CSMD1_M" [color="0 0 0.8539136795903438", label="0.14",style=bold];
"ADAMTS20_S" -> "UGT2A1_E" [color="0 0 0.8532553035844916", label="0.15",style=bold];
"LCT_S" -> "RYR2_M" [color="0 0 0.8525969275786394", label="0.15",style=bold];
"PCDH10_S" -> "RP1_E" [color="0 0 0.8519385515727872", label="0.15",style=bold];
"PCDHA12_S" -> "DSCAM_E" [color="0 0 0.8512801755669349", label="0.15",style=bold];
"CCDC67_S" -> "CSMD3_E" [color="0 0 0.8506217995610826", label="0.15",style=bold];
"MAB21L2_S" -> "ADAMTS20_E" [color="0 0 0.8499634235552305", label="0.15",style=bold];
"MAB21L2_S" -> "HFM1_S" [color="0 0 0.8493050475493782", label="0.15",style=bold];
"ADAMTS20_S" -> "BEST2_E" [color="0 0 0.848646671543526", label="0.15",style=bold];
"GALNTL6_S" -> "MUC4_E" [color="0 0 0.8479882955376737", label="0.15",style=bold];
"KRT1_S" -> "FLG_E" [color="0 0 0.8473299195318216", label="0.15",style=bold];
"CCDC67_S" -> "MYH7_M" [color="0 0 0.8466715435259693", label="0.15",style=bold];
"MRGPRX4_S" -> "GABRB3_E" [color="0 0 0.846013167520117", label="0.15",style=bold];
"PCDHA12_S" -> "BEST2_E" [color="0 0 0.8453547915142648", label="0.15",style=bold];
"LRAT_S" -> "CCDC67_E" [color="0 0 0.8446964155084126", label="0.16",style=bold];
"TDRD5_S" -> "ERN2_M" [color="0 0 0.8440380395025604", label="0.16",style=bold];
"KRT24_S" -> "CPNE6_E" [color="0 0 0.8433796634967081", label="0.16",style=bold];
"NEB_S" -> "CSMD3_E" [color="0 0 0.842721287490856", label="0.16",style=bold];
"MYBPC2_S" -> "LCE2C_M" [color="0 0 0.8420629114850037", label="0.17",style=bold];
"LCE2C_S" -> "KRT1_M" [color="0 0 0.8414045354791515", label="0.17",style=bold];
"LPPR1_S" -> "LINGO2_E" [color="0 0 0.8407461594732992", label="0.17",style=bold];
"MAB21L2_S" -> "CPNE6_M" [color="0 0 0.840087783467447", label="0.17",style=bold];
"MAB21L2_S" -> "NEB_E" [color="0 0 0.8394294074615948", label="0.18",style=bold];
"ADAMTS20_S" -> "CNTN5_M" [color="0 0 0.8387710314557425", label="0.19",style=bold];
"TUBA3C_S" -> "TTN_E" [color="0 0 0.8381126554498903", label="0.19",style=bold];
"MAB21L2_S" -> "HFM1_M" [color="0 0 0.837454279444038", label="0.19",style=bold];
"LPPR1_S" -> "NELL1_M" [color="0 0 0.8367959034381858", label="0.19",style=bold];
"ZIC1_S" -> "MYBPC2_E" [color="0 0 0.8361375274323336", label="0.20",style=bold];
"MYO3A_S" -> "PCLO_M" [color="0 0 0.8354791514264813", label="0.20",style=bold];
"LCE2C_S" -> "LCE2C_E" [color="0 0 0.8348207754206292", label="0.20",style=bold];
"MRGPRX4_S" -> "CPNE6_M" [color="0 0 0.8341623994147769", label="0.20",style=bold];
"LCE2C_S" -> "NELL1_M" [color="0 0 0.8335040234089247", label="0.20",style=bold];
"MRGPRX4_S" -> "NPFFR2_M" [color="0 0 0.8328456474030724", label="0.20",style=bold];
"MYO3A_S" -> "CDKN2A_E" [color="0 0 0.8321872713972203", label="0.21",style=bold];
"FREM2_S" -> "RYR2_E" [color="0 0 0.831528895391368", label="0.21",style=bold];
"PCDH10_S" -> "BEST2_E" [color="0 0 0.8308705193855157", label="0.21",style=bold];
"MUC4_S" -> "ERBB4_E" [color="0 0 0.8302121433796635", label="0.22",style=bold];
"TUBA3C_S" -> "PTPRT_M" [color="0 0 0.8295537673738113", label="0.22",style=bold];
"ZIC1_S" -> "MUC16_M" [color="0 0 0.8288953913679591", label="0.22",style=bold];
"MYH6_S" -> "HFM1_E" [color="0 0 0.8282370153621068", label="0.22",style=bold];
"KRT1_S" -> "PCDHA10_E" [color="0 0 0.8275786393562546", label="0.22",style=bold];
"UGT2A1_S" -> "LRRTM4_E" [color="0 0 0.8269202633504024", label="0.22",style=bold];
"CCDC67_S" -> "RBP3_E" [color="0 0 0.8262618873445502", label="0.22",style=bold];
"CDH19_S" -> "TTN_S" [color="0 0 0.8256035113386979", label="0.22",style=bold];
"GALNTL6_S" -> "PCLO_E" [color="0 0 0.8249451353328456", label="0.22",style=bold];
"KRT1_S" -> "ERBB4_S" [color="0 0 0.8242867593269935", label="0.22",style=bold];
"PCDHA12_S" -> "LPA_E" [color="0 0 0.8236283833211412", label="0.22",style=bold];
"BEST2_S" -> "NTSR2_E" [color="0 0 0.822970007315289", label="0.22",style=bold];
"NTSR2_S" -> "NTSR2_M" [color="0 0 0.8223116313094367", label="0.22",style=bold];
"CCDC67_S" -> "TUBA3C_E" [color="0 0 0.8216532553035845", label="0.22",style=bold];
"LINGO2_S" -> "FLG_M" [color="0 0 0.8209948792977323", label="0.23",style=bold];
"DSCAM_S" -> "DUSP27_E" [color="0 0 0.82033650329188", label="0.23",style=bold];
"LCT_S" -> "KRT1_E" [color="0 0 0.8196781272860278", label="0.23",style=bold];
"MAB21L2_S" -> "LRP1B_S" [color="0 0 0.8190197512801756", label="0.24",style=bold];
"ADAMTS20_S" -> "KRT24_E" [color="0 0 0.8183613752743234", label="0.24",style=bold];
"CPNE6_S" -> "MYH6_E" [color="0 0 0.8177029992684711", label="0.24",style=bold];
"LCT_S" -> "LPA_M" [color="0 0 0.817044623262619", label="0.24",style=bold];
"ADAMTS20_S" -> "ERN2_M" [color="0 0 0.8163862472567667", label="0.25",style=bold];
"ADAMTS20_S" -> "CCDC67_M" [color="0 0 0.8157278712509144", label="0.25",style=bold];
"MYO3A_S" -> "MC5R_M" [color="0 0 0.8150694952450622", label="0.25",style=bold];
"FREM2_S" -> "ABCA13_S" [color="0 0 0.81441111923921", label="0.25",style=bold];
"CPNE6_S" -> "NELL1_M" [color="0 0 0.8137527432333578", label="0.25",style=bold];
"LRAT_S" -> "CNTNAP5_E" [color="0 0 0.8130943672275055", label="0.25",style=bold];
"MYBPC2_S" -> "PKHD1L1_M" [color="0 0 0.8124359912216532", label="0.26",style=bold];
"TUBA3C_S" -> "DSCAM_E" [color="0 0 0.811777615215801", label="0.26",style=bold];
"MC5R_S" -> "TDRD5_M" [color="0 0 0.8111192392099488", label="0.26",style=bold];
"MC5R_S" -> "GABRB3_E" [color="0 0 0.8104608632040966", label="0.26",style=bold];
"ERBB4_S" -> "CDKN2A_E" [color="0 0 0.8098024871982443", label="0.26",style=bold];
"GABRB3_S" -> "CPNE6_E" [color="0 0 0.8091441111923922", label="0.26",style=bold];
"ZIC1_S" -> "CSMD1_M" [color="0 0 0.8084857351865399", label="0.27",style=bold];
"NELL1_S" -> "CR2_M" [color="0 0 0.8078273591806877", label="0.28",style=bold];
"ZIC1_S" -> "PCDHA12_M" [color="0 0 0.8071689831748354", label="0.28",style=bold];
"LCE2C_S" -> "UGT2A1_E" [color="0 0 0.8065106071689832", label="0.28",style=bold];
"LPPR1_S" -> "TUBA3C_M" [color="0 0 0.805852231163131", label="0.28",style=bold];
"MYO3A_S" -> "LPPR1_E" [color="0 0 0.8051938551572787", label="0.28",style=bold];
"ATP13A4_S" -> "CSMD3_S" [color="0 0 0.8045354791514265", label="0.28",style=bold];
"KRT1_S" -> "NEB_E" [color="0 0 0.8038771031455743", label="0.29",style=bold];
"NPFFR2_S" -> "NELL1_E" [color="0 0 0.8032187271397221", label="0.29",style=bold];
"GALNTL6_S" -> "MUC16_E" [color="0 0 0.8025603511338698", label="0.29",style=bold];
"KRT1_S" -> "LINGO2_M" [color="0 0 0.8019019751280176", label="0.29",style=bold];
"CCDC67_S" -> "NELL1_E" [color="0 0 0.8012435991221654", label="0.29",style=bold];
"BEST2_S" -> "LRP1B_M" [color="0 0 0.8005852231163131", label="0.29",style=bold];
"KRT1_S" -> "PCDH10_M" [color="0 0 0.7999268471104609", label="0.29",style=bold];
"CCDC67_S" -> "UGT2A1_M" [color="0 0 0.7992684711046086", label="0.29",style=bold];
"LCE2C_S" -> "NTSR2_M" [color="0 0 0.7986100950987565", label="0.29",style=bold];
"LPPR1_S" -> "DNAH11_E" [color="0 0 0.7979517190929042", label="0.29",style=bold];
"BEST2_S" -> "MYBPC2_M" [color="0 0 0.7972933430870519", label="0.29",style=bold];
"KRT1_S" -> "LRRTM4_S" [color="0 0 0.7966349670811997", label="0.29",style=bold];
"PCDHA12_S" -> "FSTL5_M" [color="0 0 0.7959765910753475", label="0.29",style=bold];
"KRT1_S" -> "LPPR1_M" [color="0 0 0.7953182150694953", label="0.29",style=bold];
"NELL1_S" -> "LCE2C_E" [color="0 0 0.794659839063643", label="0.29",style=bold];
"CPNE6_S" -> "MYH13_E" [color="0 0 0.7940014630577908", label="0.29",style=bold];
"KRT24_S" -> "MAB21L2_E" [color="0 0 0.7933430870519386", label="0.29",style=bold];
"PCDHA12_S" -> "LCE2C_M" [color="0 0 0.7926847110460864", label="0.29",style=bold];
"UGT2A1_S" -> "MUC4_E" [color="0 0 0.7920263350402341", label="0.29",style=bold];
"BEST2_S" -> "MYO3A_M" [color="0 0 0.7913679590343818", label="0.29",style=bold];
"KRT1_S" -> "NTSR2_M" [color="0 0 0.7907095830285297", label="0.29",style=bold];
"KRT1_S" -> "COL11A1_M" [color="0 0 0.7900512070226774", label="0.29",style=bold];
"LCT_S" -> "GABRB3_E" [color="0 0 0.7893928310168252", label="0.29",style=bold];
"UGT2A1_S" -> "CDKN2A_E" [color="0 0 0.788734455010973", label="0.29",style=bold];
"KRT24_S" -> "LCE2C_M" [color="0 0 0.7880760790051207", label="0.29",style=bold];
"CCDC67_S" -> "ZIC1_M" [color="0 0 0.7874177029992685", label="0.29",style=bold];
"BEST2_S" -> "CR2_E" [color="0 0 0.7867593269934162", label="0.29",style=bold];
"MYH6_S" -> "TDRD5_E" [color="0 0 0.786100950987564", label="0.29",style=bold];
"BEST2_S" -> "FSTL5_M" [color="0 0 0.7854425749817118", label="0.29",style=bold];
"KRT1_S" -> "MYO3A_E" [color="0 0 0.7847841989758596", label="0.29",style=bold];
"MYO3A_S" -> "LCE2C_E" [color="0 0 0.7841258229700073", label="0.29",style=bold];
"TDRD5_S" -> "ATP13A4_E" [color="0 0 0.7834674469641552", label="0.29",style=bold];
"ZIC1_S" -> "RYR2_S" [color="0 0 0.7828090709583029", label="0.29",style=bold];
"LINGO2_S" -> "CDKN2A_E" [color="0 0 0.7821506949524506", label="0.30",style=bold];
"CR2_S" -> "GABRB3_S" [color="0 0 0.7814923189465984", label="0.30",style=bold];
"LCE2C_S" -> "MUC16_S" [color="0 0 0.7808339429407462", label="0.31",style=bold];
"MYH6_S" -> "MAB21L2_M" [color="0 0 0.780175566934894", label="0.31",style=bold];
"MYBPC2_S" -> "LRP1B_E" [color="0 0 0.7795171909290417", label="0.31",style=bold];
"MYO3A_S" -> "ADAMTS20_E" [color="0 0 0.7788588149231895", label="0.31",style=bold];
"PCDHA12_S" -> "CDH19_E" [color="0 0 0.7782004389173373", label="0.31",style=bold];
"KRT1_S" -> "LPA_E" [color="0 0 0.7775420629114851", label="0.32",style=bold];
"CR2_S" -> "DSCAM_S" [color="0 0 0.7768836869056328", label="0.32",style=bold];
"MC5R_S" -> "ZIC1_E" [color="0 0 0.7762253108997805", label="0.32",style=bold];
"CCDC67_S" -> "LINGO2_M" [color="0 0 0.7755669348939284", label="0.33",style=bold];
"LPA_S" -> "GALNTL6_E" [color="0 0 0.7749085588880761", label="0.33",style=bold];
"RYR2_S" -> "PTPRT_S" [color="0 0 0.7742501828822239", label="0.33",style=bold];
"MUC4_S" -> "MYH7_M" [color="0 0 0.7735918068763716", label="0.34",style=bold];
"KRT1_S" -> "CDH19_M" [color="0 0 0.7729334308705194", label="0.34",style=bold];
"MYH7_S" -> "PKHD1L1_E" [color="0 0 0.7722750548646672", label="0.34",style=bold];
"ATP13A4_S" -> "CSMD1_E" [color="0 0 0.7716166788588149", label="0.34",style=bold];
"TDRD5_S" -> "NPFFR2_M" [color="0 0 0.7709583028529627", label="0.34",style=bold];
"MYBPC2_S" -> "NEB_E" [color="0 0 0.7702999268471105", label="0.34",style=bold];
"PCDH10_S" -> "ABCA13_E" [color="0 0 0.7696415508412583", label="0.35",style=bold];
"PCDHA12_S" -> "MUC4_E" [color="0 0 0.768983174835406", label="0.35",style=bold];
"MRGPRX4_S" -> "LPA_E" [color="0 0 0.7683247988295538", label="0.35",style=bold];
"LPPR1_S" -> "LPPR1_E" [color="0 0 0.7676664228237016", label="0.35",style=bold];
"TUBA3C_S" -> "MUC4_E" [color="0 0 0.7670080468178493", label="0.35",style=bold];
"NTSR2_S" -> "KRT24_M" [color="0 0 0.7663496708119971", label="0.35",style=bold];
"MYBPC2_S" -> "MUC4_E" [color="0 0 0.7656912948061448", label="0.35",style=bold];
"PCDHA12_S" -> "PTPRT_M" [color="0 0 0.7650329188002927", label="0.35",style=bold];
"LRRTM4_S" -> "CCDC67_M" [color="0 0 0.7643745427944404", label="0.35",style=bold];
"LINGO2_S" -> "MYH6_E" [color="0 0 0.7637161667885881", label="0.35",style=bold];
"MYBPC2_S" -> "NEB_M" [color="0 0 0.763057790782736", label="0.36",style=bold];
"LPPR1_S" -> "CSMD3_M" [color="0 0 0.7623994147768838", label="0.36",style=bold];
"DUSP27_S" -> "CSMD3_M" [color="0 0 0.7617410387710315", label="0.36",style=bold];
"NPFFR2_S" -> "DNAH11_E" [color="0 0 0.7610826627651792", label="0.37",style=bold];
"BEST2_S" -> "MUC4_M" [color="0 0 0.760424286759327", label="0.37",style=bold];
"LPPR1_S" -> "PTPRT_M" [color="0 0 0.7597659107534748", label="0.37",style=bold];
"PCDHA10_M" -> "NPFFR2_E" [color="0 0 0.7591075347476226", label="0.37",style=bold];
"MYO3A_S" -> "COL11A1_M" [color="0 0 0.7584491587417703", label="0.38",style=bold];
"GALNTL6_S" -> "CSMD3_E" [color="0 0 0.757790782735918", label="0.38",style=bold];
"LCT_S" -> "PTPRT_M" [color="0 0 0.7571324067300659", label="0.38",style=bold];
"CDH19_S" -> "MYO3A_E" [color="0 0 0.7564740307242136", label="0.39",style=bold];
"ADAMTS20_S" -> "HFM1_E" [color="0 0 0.7558156547183614", label="0.39",style=bold];
"CPNE6_S" -> "NELL1_E" [color="0 0 0.7551572787125092", label="0.39",style=bold];
"BEST2_S" -> "CSMD3_M" [color="0 0 0.7544989027066569", label="0.39",style=bold];
"DUSP27_S" -> "KRT24_M" [color="0 0 0.7538405267008047", label="0.39",style=bold];
"MYO3A_S" -> "GABRB3_M" [color="0 0 0.7531821506949525", label="0.40",style=bold];
"NTSR2_S" -> "DUSP27_M" [color="0 0 0.7525237746891003", label="0.40",style=bold];
"ATP13A4_S" -> "UGT2A1_M" [color="0 0 0.751865398683248", label="0.40",style=bold];
"CPNE6_S" -> "RYR2_E" [color="0 0 0.7512070226773958", label="0.41",style=bold];
"LCE2C_S" -> "MC5R_M" [color="0 0 0.7505486466715435", label="0.41",style=bold];
"CPNE6_S" -> "MYH7_E" [color="0 0 0.7498902706656914", label="0.41",style=bold];
"CPNE6_S" -> "CPNE6_E" [color="0 0 0.7492318946598391", label="0.41",style=bold];
"NTSR2_S" -> "MYH13_E" [color="0 0 0.7485735186539868", label="0.41",style=bold];
"CCDC67_S" -> "MAB21L2_M" [color="0 0 0.7479151426481346", label="0.41",style=bold];
"TDRD5_S" -> "PCLO_E" [color="0 0 0.7472567666422824", label="0.41",style=bold];
"ERN2_S" -> "CSMD3_S" [color="0 0 0.7465983906364302", label="0.41",style=bold];
"UGT2A1_S" -> "RP1_S" [color="0 0 0.7459400146305779", label="0.41",style=bold];
"CDH19_S" -> "PTPRT_E" [color="0 0 0.7452816386247257", label="0.41",style=bold];
"NTSR2_S" -> "TTN_S" [color="0 0 0.7446232626188735", label="0.41",style=bold];
"MUC4_S" -> "RP1_S" [color="0 0 0.7439648866130213", label="0.42",style=bold];
"KRT24_S" -> "FSTL5_E" [color="0 0 0.743306510607169", label="0.42",style=bold];
"ATP13A4_S" -> "NEB_M" [color="0 0 0.7426481346013167", label="0.42",style=bold];
"TDRD5_S" -> "PKHD1L1_S" [color="0 0 0.7419897585954646", label="0.43",style=bold];
"MYO3A_S" -> "UGT2A1_E" [color="0 0 0.7413313825896123", label="0.43",style=bold];
"CDH19_S" -> "FSTL5_M" [color="0 0 0.7406730065837601", label="0.43",style=bold];
"LRRTM4_S" -> "COL11A1_M" [color="0 0 0.7400146305779078", label="0.44",style=bold];
"FSTL5_S" -> "MYH7_E" [color="0 0 0.7393562545720556", label="0.44",style=bold];
"PCDHA12_S" -> "RP1_E" [color="0 0 0.7386978785662034", label="0.44",style=bold];
"DSCAM_S" -> "LCE2C_E" [color="0 0 0.7380395025603512", label="0.44",style=bold];
"LINGO2_S" -> "DNAH11_E" [color="0 0 0.737381126554499", label="0.44",style=bold];
"UGT2A1_S" -> "MAB21L2_E" [color="0 0 0.7367227505486467", label="0.44",style=bold];
"MYO3A_S" -> "MYH6_E" [color="0 0 0.7360643745427945", label="0.44",style=bold];
"LCE2C_S" -> "KRT24_E" [color="0 0 0.7354059985369422", label="0.44",style=bold];
"MAB21L2_S" -> "CNTNAP5_M" [color="0 0 0.73474762253109", label="0.44",style=bold];
"CR2_S" -> "LRP1B_E" [color="0 0 0.7340892465252378", label="0.44",style=bold];
"PTPRT_S" -> "MAB21L2_M" [color="0 0 0.7334308705193855", label="0.44",style=bold];
"GALNTL6_S" -> "CNTNAP5_E" [color="0 0 0.7327724945135333", label="0.45",style=bold];
"MUC4_S" -> "TUBA3C_E" [color="0 0 0.732114118507681", label="0.46",style=bold];
"LCE2C_S" -> "DNAH11_M" [color="0 0 0.7314557425018289", label="0.46",style=bold];
"MYBPC2_S" -> "LPPR1_E" [color="0 0 0.7307973664959766", label="0.46",style=bold];
"LCE2C_S" -> "CSMD3_E" [color="0 0 0.7301389904901243", label="0.46",style=bold];
"CR2_S" -> "DUSP27_E" [color="0 0 0.7294806144842721", label="0.47",style=bold];
"KRT24_S" -> "FLG_M" [color="0 0 0.72882223847842", label="0.47",style=bold];
"RBP3_S" -> "ERBB4_S" [color="0 0 0.7281638624725677", label="0.47",style=bold];
"NPFFR2_S" -> "MYBPC2_M" [color="0 0 0.7275054864667154", label="0.48",style=bold];
"MYO3A_S" -> "PTPRT_E" [color="0 0 0.7268471104608633", label="0.48",style=bold];
"PCDH10_S" -> "CSMD1_S" [color="0 0 0.726188734455011", label="0.49",style=bold];
"FSTL5_S" -> "COL11A1_M" [color="0 0 0.7255303584491588", label="0.49",style=bold];
"NPFFR2_S" -> "PCLO_S" [color="0 0 0.7248719824433065", label="0.49",style=bold];
"MYH6_S" -> "TTN_E" [color="0 0 0.7242136064374542", label="0.49",style=bold];
"UGT2A1_S" -> "PCLO_S" [color="0 0 0.7235552304316021", label="0.49",style=bold];
"PCDH10_S" -> "RBP3_E" [color="0 0 0.7228968544257498", label="0.50",style=bold];
"DNAH11_S" -> "LCT_M" [color="0 0 0.7222384784198976", label="0.50",style=bold];
"FSTL5_S" -> "PCDH10_E" [color="0 0 0.7215801024140454", label="0.51",style=bold];
"MYH7_S" -> "COL11A1_S" [color="0 0 0.7209217264081932", label="0.51",style=bold];
"LCT_S" -> "PKHD1L1_M" [color="0 0 0.7202633504023409", label="0.51",style=bold];
"LCE2C_S" -> "PCLO_E" [color="0 0 0.7196049743964887", label="0.51",style=bold];
"CCDC67_S" -> "GABRB3_E" [color="0 0 0.7189465983906365", label="0.51",style=bold];
"MRGPRX4_S" -> "CDKN2A_M" [color="0 0 0.7182882223847842", label="0.51",style=bold];
"LCE2C_S" -> "FSTL5_M" [color="0 0 0.717629846378932", label="0.51",style=bold];
"UGT2A1_S" -> "ABCA13_M" [color="0 0 0.7169714703730797", label="0.51",style=bold];
"MYH7_S" -> "FSTL5_M" [color="0 0 0.7163130943672276", label="0.51",style=bold];
"KRT1_S" -> "DSCAM_E" [color="0 0 0.7156547183613753", label="0.51",style=bold];
"MYBPC2_S" -> "BEST2_E" [color="0 0 0.714996342355523", label="0.51",style=bold];
"FREM2_S" -> "PTPRT_E" [color="0 0 0.7143379663496708", label="0.51",style=bold];
"MYH6_S" -> "NTSR2_E" [color="0 0 0.7136795903438187", label="0.51",style=bold];
"MYBPC2_S" -> "CDKN2A_S" [color="0 0 0.7130212143379664", label="0.51",style=bold];
"MYH6_S" -> "LINGO2_E" [color="0 0 0.7123628383321141", label="0.52",style=bold];
"CCDC67_S" -> "CDKN2A_S" [color="0 0 0.7117044623262619", label="0.52",style=bold];
"LPA_S" -> "CNTN5_E" [color="0 0 0.7110460863204097", label="0.52",style=bold];
"MYH6_S" -> "NPFFR2_E" [color="0 0 0.7103877103145575", label="0.52",style=bold];
"LINGO2_S" -> "CDKN2A_M" [color="0 0 0.7097293343087052", label="0.52",style=bold];
"MYO3A_S" -> "ERBB4_M" [color="0 0 0.7090709583028529", label="0.53",style=bold];
"NELL1_S" -> "CCDC67_M" [color="0 0 0.7084125822970008", label="0.53",style=bold];
"LRAT_S" -> "ERBB4_M" [color="0 0 0.7077542062911485", label="0.53",style=bold];
"CDH19_S" -> "Survival" [color="0 0 0.7070958302852963", label="0.53",style=bold];
"CPNE6_S" -> "PCDHA12_E" [color="0 0 0.706437454279444", label="0.53",style=bold];
"NPFFR2_S" -> "PKHD1L1_E" [color="0 0 0.7057790782735918", label="0.54",style=bold];
"LINGO2_S" -> "RBP3_E" [color="0 0 0.7051207022677396", label="0.54",style=bold];
"ABCA13_S" -> "PCDHA12_S" [color="0 0 0.7044623262618874", label="0.55",style=bold];
"CDH19_S" -> "TDRD5_E" [color="0 0 0.7038039502560351", label="0.55",style=bold];
"LPPR1_S" -> "TTN_S" [color="0 0 0.7031455742501829", label="0.55",style=bold];
"LINGO2_S" -> "UGT2A1_E" [color="0 0 0.7024871982443307", label="0.55",style=bold];
"UGT2A1_S" -> "CNTNAP5_E" [color="0 0 0.7018288222384784", label="0.56",style=bold];
"MYH6_S" -> "FSTL5_M" [color="0 0 0.7011704462326263", label="0.56",style=bold];
"PCDHA12_S" -> "NTSR2_M" [color="0 0 0.700512070226774", label="0.56",style=bold];
"ZIC1_S" -> "LRRTM4_E" [color="0 0 0.6998536942209217", label="0.56",style=bold];
"PCDH10_S" -> "NTSR2_M" [color="0 0 0.6991953182150695", label="0.57",style=bold];
"MAB21L2_S" -> "CDH19_M" [color="0 0 0.6985369422092174", label="0.57",style=bold];
"MYBPC2_S" -> "MAB21L2_E" [color="0 0 0.6978785662033651", label="0.58",style=bold];
"LRRTM4_S" -> "TDRD5_E" [color="0 0 0.6972201901975128", label="0.58",style=bold];
"MAB21L2_S" -> "KRT24_M" [color="0 0 0.6965618141916606", label="0.58",style=bold];
"MYO3A_S" -> "RP1_M" [color="0 0 0.6959034381858084", label="0.58",style=bold];
"MYH6_S" -> "LINGO2_M" [color="0 0 0.6952450621799562", label="0.58",style=bold];
"NTSR2_S" -> "LRAT_E" [color="0 0 0.6945866861741039", label="0.58",style=bold];
"MUC4_S" -> "NELL1_M" [color="0 0 0.6939283101682516", label="0.58",style=bold];
"NEB_S" -> "FLG_E" [color="0 0 0.6932699341623995", label="0.58",style=bold];
"CPNE6_S" -> "CNTN5_E" [color="0 0 0.6926115581565472", label="0.58",style=bold];
"MRGPRX4_S" -> "CNTN5_E" [color="0 0 0.691953182150695", label="0.58",style=bold];
"CDH19_S" -> "RYR2_E" [color="0 0 0.6912948061448427", label="0.59",style=bold];
"MYBPC2_S" -> "RP1_E" [color="0 0 0.6906364301389905", label="0.60",style=bold];
"LPA_S" -> "CNTNAP5_E" [color="0 0 0.6899780541331383", label="0.60",style=bold];
"LINGO2_S" -> "MUC16_E" [color="0 0 0.6893196781272861", label="0.61",style=bold];
"GALNTL6_S" -> "KRT1_M" [color="0 0 0.6886613021214338", label="0.61",style=bold];
"CPNE6_S" -> "MUC16_E" [color="0 0 0.6880029261155816", label="0.61",style=bold];
"CCDC67_S" -> "CNTN5_E" [color="0 0 0.6873445501097294", label="0.61",style=bold];
"UGT2A1_S" -> "CSMD1_E" [color="0 0 0.6866861741038771", label="0.61",style=bold];
"MAB21L2_S" -> "KRT1_M" [color="0 0 0.6860277980980249", label="0.61",style=bold];
"CNTN5_S" -> "MYH6_E" [color="0 0 0.6853694220921727", label="0.61",style=bold];
"BEST2_S" -> "KRT1_M" [color="0 0 0.6847110460863204", label="0.61",style=bold];
"MAB21L2_S" -> "ATP13A4_M" [color="0 0 0.6840526700804682", label="0.61",style=bold];
"GALNTL6_S" -> "CNTN5_S" [color="0 0 0.6833942940746159", label="0.61",style=bold];
"ZIC1_S" -> "GALNTL6_M" [color="0 0 0.6827359180687638", label="0.61",style=bold];
"KRT1_S" -> "NTSR2_E" [color="0 0 0.6820775420629115", label="0.61",style=bold];
"GABRB3_S" -> "CCDC67_M" [color="0 0 0.6814191660570593", label="0.61",style=bold];
"NELL1_S" -> "TTN_E" [color="0 0 0.680760790051207", label="0.62",style=bold];
"LPA_S" -> "NELL1_E" [color="0 0 0.6801024140453549", label="0.62",style=bold];
"MYBPC2_S" -> "CDKN2A_E" [color="0 0 0.6794440380395026", label="0.62",style=bold];
"MYO3A_S" -> "ATP13A4_S" [color="0 0 0.6787856620336503", label="0.63",style=bold];
"KRT24_S" -> "LRRTM4_E" [color="0 0 0.6781272860277981", label="0.63",style=bold];
"MYBPC2_S" -> "RBP3_M" [color="0 0 0.6774689100219459", label="0.63",style=bold];
"ADAMTS20_S" -> "TDRD5_M" [color="0 0 0.6768105340160937", label="0.63",style=bold];
"FREM2_S" -> "CNTNAP5_S" [color="0 0 0.6761521580102414", label="0.64",style=bold];
"ADAMTS20_S" -> "ABCA13_E" [color="0 0 0.6754937820043891", label="0.64",style=bold];
"LCE2C_S" -> "ERBB4_M" [color="0 0 0.674835405998537", label="0.64",style=bold];
"MYH6_S" -> "CNTNAP5_E" [color="0 0 0.6741770299926848", label="0.64",style=bold];
"RP1_S" -> "ABCA13_M" [color="0 0 0.6735186539868325", label="0.64",style=bold];
"MC5R_S" -> "CSMD1_E" [color="0 0 0.6728602779809802", label="0.64",style=bold];
"CCDC67_S" -> "CR2_E" [color="0 0 0.6722019019751281", label="0.64",style=bold];
"UGT2A1_S" -> "CR2_E" [color="0 0 0.6715435259692758", label="0.64",style=bold];
"LCE2C_S" -> "FREM2_E" [color="0 0 0.6708851499634236", label="0.64",style=bold];
"KRT1_S" -> "CR2_E" [color="0 0 0.6702267739575714", label="0.64",style=bold];
"MYH13_S" -> "LRRTM4_E" [color="0 0 0.6695683979517191", label="0.65",style=bold];
"ATP13A4_S" -> "LRAT_E" [color="0 0 0.6689100219458669", label="0.66",style=bold];
"PCDH10_S" -> "DSCAM_M" [color="0 0 0.6682516459400146", label="0.66",style=bold];
"ADAMTS20_S" -> "CSMD1_E" [color="0 0 0.6675932699341625", label="0.66",style=bold];
"ADAMTS20_S" -> "COL11A1_E" [color="0 0 0.6669348939283102", label="0.66",style=bold];
"FREM2_S" -> "LPA_M" [color="0 0 0.6662765179224579", label="0.66",style=bold];
"LRRTM4_S" -> "FSTL5_M" [color="0 0 0.6656181419166057", label="0.66",style=bold];
"CR2_S" -> "NELL1_M" [color="0 0 0.6649597659107536", label="0.67",style=bold];
"LPA_S" -> "FLG_S" [color="0 0 0.6643013899049013", label="0.67",style=bold];
"PCDH10_S" -> "GABRB3_E" [color="0 0 0.663643013899049", label="0.67",style=bold];
"LCE2C_S" -> "HFM1_E" [color="0 0 0.6629846378931968", label="0.68",style=bold];
"UGT2A1_S" -> "DUSP27_E" [color="0 0 0.6623262618873446", label="0.68",style=bold];
"CCDC67_S" -> "GALNTL6_M" [color="0 0 0.6616678858814924", label="0.68",style=bold];
"MYH7_S" -> "MYO3A_E" [color="0 0 0.6610095098756401", label="0.68",style=bold];
"HFM1_S" -> "PKHD1L1_E" [color="0 0 0.6603511338697878", label="0.69",style=bold];
"CCDC67_S" -> "FSTL5_E" [color="0 0 0.6596927578639357", label="0.69",style=bold];
"ERBB4_S" -> "FLG_M" [color="0 0 0.6590343818580834", label="0.69",style=bold];
"LCE2C_S" -> "CDH19_M" [color="0 0 0.6583760058522312", label="0.69",style=bold];
"MYBPC2_S" -> "TUBA3C_M" [color="0 0 0.6577176298463789", label="0.69",style=bold];
"LCE2C_S" -> "MUC4_M" [color="0 0 0.6570592538405268", label="0.69",style=bold];
"PCDHA12_S" -> "ZIC1_M" [color="0 0 0.6564008778346745", label="0.69",style=bold];
"BEST2_S" -> "CDH19_M" [color="0 0 0.6557425018288223", label="0.69",style=bold];
"UGT2A1_S" -> "CDH19_M" [color="0 0 0.65508412582297", label="0.69",style=bold];
"BEST2_S" -> "NPFFR2_E" [color="0 0 0.6544257498171178", label="0.69",style=bold];
"KRT1_S" -> "LCT_M" [color="0 0 0.6537673738112656", label="0.69",style=bold];
"KRT1_S" -> "PCLO_E" [color="0 0 0.6531089978054133", label="0.69",style=bold];
"MYH7_S" -> "PCLO_E" [color="0 0 0.6524506217995611", label="0.69",style=bold];
"PCDHA12_S" -> "CDKN2A_E" [color="0 0 0.6517922457937089", label="0.69",style=bold];
"MYO3A_S" -> "ZIC1_E" [color="0 0 0.6511338697878566", label="0.69",style=bold];
"MAB21L2_S" -> "LRAT_E" [color="0 0 0.6504754937820044", label="0.69",style=bold];
"LCT_S" -> "NTSR2_M" [color="0 0 0.6498171177761523", label="0.69",style=bold];
"MAB21L2_S" -> "MUC4_E" [color="0 0 0.6491587417703", label="0.69",style=bold];
"MC5R_S" -> "RBP3_M" [color="0 0 0.6485003657644477", label="0.69",style=bold];
"KRT1_S" -> "MRGPRX4_E" [color="0 0 0.6478419897585954", label="0.69",style=bold];
"KRT1_S" -> "LRAT_M" [color="0 0 0.6471836137527432", label="0.69",style=bold];
"KRT1_S" -> "ERN2_M" [color="0 0 0.6465252377468911", label="0.69",style=bold];
"KRT1_S" -> "PKHD1L1_M" [color="0 0 0.6458668617410388", label="0.69",style=bold];
"LCE2C_S" -> "MYO3A_E" [color="0 0 0.6452084857351865", label="0.69",style=bold];
"MRGPRX4_S" -> "KRT24_E" [color="0 0 0.6445501097293344", label="0.69",style=bold];
"ADAMTS20_S" -> "TUBA3C_E" [color="0 0 0.6438917337234822", label="0.69",style=bold];
"MYH7_S" -> "BEST2_M" [color="0 0 0.6432333577176299", label="0.70",style=bold];
"LCE2C_S" -> "PCDHA12_M" [color="0 0 0.6425749817117776", label="0.70",style=bold];
"MYH6_S" -> "MUC4_M" [color="0 0 0.6419166057059253", label="0.70",style=bold];
"MYBPC2_S" -> "UGT2A1_M" [color="0 0 0.6412582297000732", label="0.71",style=bold];
"NEB_S" -> "CDKN2A_S" [color="0 0 0.640599853694221", label="0.71",style=bold];
"MYBPC2_S" -> "TTN_S" [color="0 0 0.6399414776883687", label="0.72",style=bold];
"CCDC67_S" -> "FREM2_M" [color="0 0 0.6392831016825165", label="0.72",style=bold];
"LPA_S" -> "CSMD3_E" [color="0 0 0.6386247256766642", label="0.72",style=bold];
"PCDHA12_S" -> "CNTNAP5_E" [color="0 0 0.637966349670812", label="0.72",style=bold];
"NELL1_S" -> "MYH6_M" [color="0 0 0.6373079736649598", label="0.73",style=bold];
"PCDHA12_S" -> "CR2_E" [color="0 0 0.6366495976591076", label="0.73",style=bold];
"ADAMTS20_S" -> "LRP1B_E" [color="0 0 0.6359912216532553", label="0.73",style=bold];
"UGT2A1_S" -> "KRT1_E" [color="0 0 0.6353328456474031", label="0.73",style=bold];
"PCDHA12_S" -> "ERBB4_E" [color="0 0 0.6346744696415509", label="0.73",style=bold];
"ADAMTS20_S" -> "CDH19_M" [color="0 0 0.6340160936356987", label="0.73",style=bold];
"ATP13A4_S" -> "TUBA3C_E" [color="0 0 0.6333577176298464", label="0.73",style=bold];
"TUBA3C_S" -> "MYO3A_M" [color="0 0 0.6326993416239941", label="0.74",style=bold];
"TDRD5_S" -> "KRT24_M" [color="0 0 0.6320409656181419", label="0.75",style=bold];
"BEST2_S" -> "PCLO_S" [color="0 0 0.6313825896122898", label="0.75",style=bold];
"CCDC67_S" -> "LPA_E" [color="0 0 0.6307242136064375", label="0.75",style=bold];
"PCDHA12_S" -> "NELL1_M" [color="0 0 0.6300658376005852", label="0.75",style=bold];
"FSTL5_S" -> "ERN2_M" [color="0 0 0.629407461594733", label="0.76",style=bold];
"CNTNAP5_S" -> "TUBA3C_E" [color="0 0 0.6287490855888808", label="0.76",style=bold];
"ADAMTS20_S" -> "PTPRT_M" [color="0 0 0.6280907095830286", label="0.76",style=bold];
"LCE2C_S" -> "LINGO2_E" [color="0 0 0.6274323335771763", label="0.77",style=bold];
"CNTN5_S" -> "MAB21L2_E" [color="0 0 0.626773957571324", label="0.77",style=bold];
"PCDH10_S" -> "CDKN2A_E" [color="0 0 0.6261155815654719", label="0.77",style=bold];
"CNTN5_S" -> "DUSP27_S" [color="0 0 0.6254572055596197", label="0.77",style=bold];
"LPPR1_S" -> "GABRB3_E" [color="0 0 0.6247988295537674", label="0.77",style=bold];
"UGT2A1_S" -> "UGT2A1_M" [color="0 0 0.6241404535479151", label="0.77",style=bold];
"UGT2A1_S" -> "GALNTL6_E" [color="0 0 0.6234820775420629", label="0.77",style=bold];
"TUBA3C_S" -> "ERBB4_E" [color="0 0 0.6228237015362107", label="0.78",style=bold];
"ADAMTS20_S" -> "CNTNAP5_S" [color="0 0 0.6221653255303585", label="0.78",style=bold];
"PCDH10_S" -> "FLG_M" [color="0 0 0.6215069495245062", label="0.79",style=bold];
"FSTL5_S" -> "MYH6_E" [color="0 0 0.620848573518654", label="0.79",style=bold];
"MYH6_S" -> "MYH13_E" [color="0 0 0.6201901975128018", label="0.79",style=bold];
"ADAMTS20_S" -> "FLG_E" [color="0 0 0.6195318215069496", label="0.79",style=bold];
"LRRTM4_S" -> "MAB21L2_M" [color="0 0 0.6188734455010974", label="0.80",style=bold];
"TDRD5_S" -> "LRP1B_M" [color="0 0 0.6182150694952451", label="0.80",style=bold];
"NELL1_S" -> "MUC16_E" [color="0 0 0.6175566934893928", label="0.80",style=bold];
"MC5R_S" -> "COL11A1_E" [color="0 0 0.6168983174835406", label="0.80",style=bold];
"NTSR2_S" -> "ERBB4_E" [color="0 0 0.6162399414776885", label="0.80",style=bold];
"MC5R_S" -> "ATP13A4_E" [color="0 0 0.6155815654718362", label="0.80",style=bold];
"CNTNAP5_S" -> "CNTN5_E" [color="0 0 0.6149231894659839", label="0.80",style=bold];
"LCT_S" -> "FSTL5_E" [color="0 0 0.6142648134601317", label="0.80",style=bold];
"LPPR1_S" -> "LCT_E" [color="0 0 0.6136064374542795", label="0.81",style=bold];
"KRT24_S" -> "HFM1_E" [color="0 0 0.6129480614484273", label="0.81",style=bold];
"NELL1_S" -> "NELL1_E" [color="0 0 0.612289685442575", label="0.81",style=bold];
"ABCA13_S" -> "RP1_E" [color="0 0 0.6116313094367227", label="0.82",style=bold];
"ATP13A4_S" -> "DUSP27_E" [color="0 0 0.6109729334308706", label="0.82",style=bold];
"LPA_S" -> "TTN_S" [color="0 0 0.6103145574250184", label="0.82",style=bold];
"CCDC67_S" -> "CDH19_M" [color="0 0 0.6096561814191661", label="0.82",style=bold];
"KRT24_S" -> "CDH19_M" [color="0 0 0.6089978054133138", label="0.82",style=bold];
"ZIC1_S" -> "NTSR2_E" [color="0 0 0.6083394294074616", label="0.83",style=bold];
"TUBA3C_S" -> "CNTNAP5_M" [color="0 0 0.6076810534016094", label="0.83",style=bold];
"TUBA3C_S" -> "NPFFR2_M" [color="0 0 0.6070226773957572", label="0.84",style=bold];
"MYBPC2_S" -> "DSCAM_M" [color="0 0 0.6063643013899049", label="0.84",style=bold];
"ERBB4_S" -> "ZIC1_M" [color="0 0 0.6057059253840527", label="0.84",style=bold];
"ATP13A4_S" -> "CSMD1_S" [color="0 0 0.6050475493782005", label="0.85",style=bold];
"NELL1_S" -> "NTSR2_M" [color="0 0 0.6043891733723482", label="0.85",style=bold];
"CCDC67_S" -> "RYR2_M" [color="0 0 0.603730797366496", label="0.85",style=bold];
"LRAT_S" -> "CNTN5_E" [color="0 0 0.6030724213606438", label="0.85",style=bold];
"MRGPRX4_S" -> "ADAMTS20_E" [color="0 0 0.6024140453547915", label="0.85",style=bold];
"CR2_S" -> "MAB21L2_E" [color="0 0 0.6017556693489393", label="0.85",style=bold];
"KRT1_S" -> "TUBA3C_E" [color="0 0 0.6010972933430871", label="0.85",style=bold];
"TUBA3C_S" -> "MAB21L2_E" [color="0 0 0.6004389173372349", label="0.85",style=bold];
"TUBA3C_S" -> "LCE2C_E" [color="0 0 0.5997805413313826", label="0.85",style=bold];
"KRT24_S" -> "COL11A1_E" [color="0 0 0.5991221653255303", label="0.85",style=bold];
"ADAMTS20_S" -> "ZIC1_E" [color="0 0 0.5984637893196781", label="0.85",style=bold];
"COL11A1_S" -> "CDH19_M" [color="0 0 0.597805413313826", label="0.86",style=bold];
"ADAMTS20_S" -> "CR2_E" [color="0 0 0.5971470373079737", label="0.86",style=bold];
"RBP3_S" -> "GALNTL6_E" [color="0 0 0.5964886613021214", label="0.86",style=bold];
"DUSP27_S" -> "CR2_M" [color="0 0 0.5958302852962692", label="0.86",style=bold];
"MUC4_S" -> "CSMD1_E" [color="0 0 0.5951719092904171", label="0.86",style=bold];
"MYO3A_S" -> "LCT_M" [color="0 0 0.5945135332845648", label="0.86",style=bold];
"TUBA3C_S" -> "LRRTM4_M" [color="0 0 0.5938551572787125", label="0.87",style=bold];
"CNTN5_S" -> "MYH7_S" [color="0 0 0.5931967812728602", label="0.87",style=bold];
"ZIC1_S" -> "MYO3A_M" [color="0 0 0.5925384052670081", label="0.87",style=bold];
"MYO3A_S" -> "FREM2_S" [color="0 0 0.5918800292611559", label="0.87",style=bold];
"NPFFR2_S" -> "MUC4_S" [color="0 0 0.5912216532553036", label="0.87",style=bold];
"LINGO2_S" -> "LPPR1_E" [color="0 0 0.5905632772494513", label="0.88",style=bold];
"BEST2_S" -> "FREM2_M" [color="0 0 0.5899049012435992", label="0.88",style=bold];
"PTPRT_M" -> "PKHD1L1_S" [color="0 0 0.5892465252377469", label="0.88",style=bold];
"MUC4_S" -> "NEB_M" [color="0 0 0.5885881492318947", label="0.88",style=bold];
"MC5R_S" -> "MAB21L2_E" [color="0 0 0.5879297732260425", label="0.89",style=bold];
"PCDHA12_S" -> "KRT1_E" [color="0 0 0.5872713972201902", label="0.90",style=bold];
"PCDHA12_S" -> "LRP1B_M" [color="0 0 0.586613021214338", label="0.90",style=bold];
"LPPR1_S" -> "RBP3_M" [color="0 0 0.5859546452084858", label="0.90",style=bold];
"LCE2C_S" -> "TTN_M" [color="0 0 0.5852962692026336", label="0.92",style=bold];
"PCDHA12_S" -> "ADAMTS20_E" [color="0 0 0.5846378931967813", label="0.92",style=bold];
"CCDC67_S" -> "FSTL5_M" [color="0 0 0.583979517190929", label="0.92",style=bold];
"KRT1_S" -> "CPNE6_M" [color="0 0 0.5833211411850768", label="0.92",style=bold];
"UGT2A1_S" -> "CDKN2A_S" [color="0 0 0.5826627651792247", label="0.92",style=bold];
"MRGPRX4_S" -> "TTN_S" [color="0 0 0.5820043891733724", label="0.92",style=bold];
"LRAT_S" -> "LRRTM4_E" [color="0 0 0.5813460131675201", label="0.92",style=bold];
"BEST2_S" -> "LRRTM4_E" [color="0 0 0.5806876371616679", label="0.92",style=bold];
"MYO3A_S" -> "CNTNAP5_M" [color="0 0 0.5800292611558158", label="0.92",style=bold];
"CDH19_S" -> "CDKN2A_E" [color="0 0 0.5793708851499635", label="0.92",style=bold];
"CPNE6_S" -> "PKHD1L1_M" [color="0 0 0.5787125091441112", label="0.92",style=bold];
"UGT2A1_S" -> "LPA_M" [color="0 0 0.5780541331382589", label="0.93",style=bold];
"CR2_S" -> "GALNTL6_E" [color="0 0 0.5773957571324068", label="0.93",style=bold];
"PCDHA12_S" -> "TDRD5_E" [color="0 0 0.5767373811265546", label="0.93",style=bold];
"MYH6_S" -> "RYR2_S" [color="0 0 0.5760790051207023", label="0.93",style=bold];
"MYBPC2_S" -> "DUSP27_E" [color="0 0 0.57542062911485", label="0.93",style=bold];
"DUSP27_S" -> "NPFFR2_M" [color="0 0 0.5747622531089978", label="0.93",style=bold];
"PCDH10_S" -> "CPNE6_M" [color="0 0 0.5741038771031456", label="0.94",style=bold];
"MUC4_S" -> "ZIC1_M" [color="0 0 0.5734455010972934", label="0.95",style=bold];
"LCE2C_S" -> "CNTNAP5_E" [color="0 0 0.5727871250914411", label="0.95",style=bold];
"PCDHA12_S" -> "LINGO2_M" [color="0 0 0.5721287490855889", label="0.95",style=bold];
"CNTN5_S" -> "FREM2_E" [color="0 0 0.5714703730797367", label="0.97",style=bold];
"DNAH11_S" -> "ABCA13_M" [color="0 0 0.5708119970738845", label="0.97",style=bold];
"LRAT_S" -> "LINGO2_E" [color="0 0 0.5701536210680322", label="0.97",style=bold];
"DUSP27_S" -> "UGT2A1_E" [color="0 0 0.56949524506218", label="0.97",style=bold];
"MYO3A_S" -> "RP1_E" [color="0 0 0.5688368690563277", label="0.97",style=bold];
"FREM2_S" -> "TDRD5_M" [color="0 0 0.5681784930504755", label="0.97",style=bold];
"MYH6_S" -> "LPA_E" [color="0 0 0.5675201170446234", label="0.98",style=bold];
"TUBA3C_S" -> "RP1_E" [color="0 0 0.5668617410387711", label="0.98",style=bold];
"LPA_S" -> "CDKN2A_M" [color="0 0 0.5662033650329188", label="0.98",style=bold];
"MAB21L2_S" -> "CNTN5_E" [color="0 0 0.5655449890270666", label="0.98",style=bold];
"MAB21L2_S" -> "TTN_M" [color="0 0 0.5648866130212143", label="0.98",style=bold];
"MAB21L2_S" -> "UGT2A1_M" [color="0 0 0.5642282370153622", label="0.98",style=bold];
"MAB21L2_S" -> "PCDH10_M" [color="0 0 0.5635698610095099", label="0.98",style=bold];
"LPPR1_S" -> "LCE2C_E" [color="0 0 0.5629114850036576", label="0.98",style=bold];
"MAB21L2_S" -> "ZIC1_M" [color="0 0 0.5622531089978055", label="0.98",style=bold];
"MYH6_S" -> "COL11A1_E" [color="0 0 0.5615947329919533", label="0.98",style=bold];
"MYH7_S" -> "ZIC1_E" [color="0 0 0.560936356986101", label="0.98",style=bold];
"BEST2_S" -> "KRT1_E" [color="0 0 0.5602779809802487", label="0.98",style=bold];
"UGT2A1_S" -> "ATP13A4_E" [color="0 0 0.5596196049743964", label="0.98",style=bold];
"PCDHA12_S" -> "LPPR1_E" [color="0 0 0.5589612289685443", label="0.98",style=bold];
"MYH7_S" -> "MUC16_E" [color="0 0 0.5583028529626921", label="0.98",style=bold];
"NPFFR2_S" -> "MYO3A_E" [color="0 0 0.5576444769568398", label="0.98",style=bold];
"KRT1_S" -> "CR2_M" [color="0 0 0.5569861009509876", label="0.98",style=bold];
"COL11A1_S" -> "NEB_S" [color="0 0 0.5563277249451354", label="0.98",style=bold];
"CNTN5_S" -> "CSMD3_M" [color="0 0 0.5556693489392832", label="0.98",style=bold];
"ERN2_M" -> "Survival" [color="0 0 0.5550109729334309", label="0.99",style=bold];
"LPPR1_S" -> "LRP1B_E" [color="0 0 0.5543525969275787", label="1.00",style=bold];
"DSCAM_S" -> "NTSR2_M" [color="0 0 0.5536942209217264", label="1.00",style=bold];
"NTSR2_S" -> "PCDHA10_M" [color="0 0 0.5530358449158742", label="1.01",style=bold];
"FSTL5_S" -> "FLG_E" [color="0 0 0.552377468910022", label="1.01",style=bold];
"LINGO2_S" -> "MYH7_M" [color="0 0 0.5517190929041698", label="1.01",style=bold];
"LPPR1_S" -> "NELL1_E" [color="0 0 0.5510607168983175", label="1.01",style=bold];
"FREM2_S" -> "ADAMTS20_M" [color="0 0 0.5504023408924652", label="1.01",style=bold];
"MRGPRX4_S" -> "MYH6_M" [color="0 0 0.549743964886613", label="1.02",style=bold];
"UGT2A1_S" -> "UGT2A1_E" [color="0 0 0.5490855888807609", label="1.03",style=bold];
"NEB_S" -> "MYBPC2_E" [color="0 0 0.5484272128749086", label="1.03",style=bold];
"UGT2A1_S" -> "NEB_E" [color="0 0 0.5477688368690563", label="1.03",style=bold];
"UGT2A1_S" -> "NELL1_M" [color="0 0 0.5471104608632041", label="1.03",style=bold];
"PCDH10_S" -> "MYH13_E" [color="0 0 0.546452084857352", label="1.03",style=bold];
"DSCAM_M" -> "TTN_M" [color="0 0 0.5457937088514997", label="1.04",style=bold];
"MAB21L2_S" -> "ABCA13_E" [color="0 0 0.5451353328456474", label="1.04",style=bold];
"LCT_S" -> "UGT2A1_E" [color="0 0 0.5444769568397951", label="1.05",style=bold];
"NELL1_S" -> "PKHD1L1_E" [color="0 0 0.543818580833943", label="1.05",style=bold];
"LCT_E" -> "TUBA3C_E" [color="0 0 0.5431602048280908", label="1.06",style=bold];
"MYH7_S" -> "ERN2_M" [color="0 0 0.5425018288222385", label="1.06",style=bold];
"NEB_S" -> "HFM1_S" [color="0 0 0.5418434528163862", label="1.07",style=bold];
"PKHD1L1_S" -> "PCLO_M" [color="0 0 0.5411850768105341", label="1.07",style=bold];
"DUSP27_S" -> "CNTN5_E" [color="0 0 0.5405267008046818", label="1.07",style=bold];
"ADAMTS20_S" -> "CSMD3_E" [color="0 0 0.5398683247988296", label="1.07",style=bold];
"RP1_S" -> "NPFFR2_E" [color="0 0 0.5392099487929773", label="1.08",style=bold];
"GALNTL6_S" -> "DSCAM_M" [color="0 0 0.5385515727871251", label="1.08",style=bold];
"MUC4_S" -> "DSCAM_E" [color="0 0 0.5378931967812729", label="1.08",style=bold];
"MYO3A_S" -> "RYR2_M" [color="0 0 0.5372348207754207", label="1.09",style=bold];
"DUSP27_S" -> "GABRB3_E" [color="0 0 0.5365764447695685", label="1.09",style=bold];
"LCE2C_S" -> "PCDHA12_E" [color="0 0 0.5359180687637162", label="1.09",style=bold];
"MYH13_S" -> "NELL1_E" [color="0 0 0.5352596927578639", label="1.10",style=bold];
"NTSR2_S" -> "CDKN2A_M" [color="0 0 0.5346013167520117", label="1.10",style=bold];
"KRT1_S" -> "MYH13_E" [color="0 0 0.5339429407461596", label="1.10",style=bold];
"CCDC67_S" -> "PTPRT_E" [color="0 0 0.5332845647403073", label="1.10",style=bold];
"CPNE6_S" -> "BEST2_M" [color="0 0 0.532626188734455", label="1.10",style=bold];
"PTPRT_S" -> "HFM1_S" [color="0 0 0.5319678127286028", label="1.10",style=bold];
"ATP13A4_S" -> "MUC4_M" [color="0 0 0.5313094367227507", label="1.10",style=bold];
"NTSR2_S" -> "DNAH11_M" [color="0 0 0.5306510607168984", label="1.10",style=bold];
"ATP13A4_S" -> "RP1_S" [color="0 0 0.5299926847110461", label="1.11",style=bold];
"LCE2C_S" -> "CSMD3_M" [color="0 0 0.5293343087051938", label="1.11",style=bold];
"PCDHA10_S" -> "TDRD5_M" [color="0 0 0.5286759326993417", label="1.11",style=bold];
"ZIC1_S" -> "LCT_M" [color="0 0 0.5280175566934895", label="1.12",style=bold];
"NTSR2_S" -> "PCDHA12_M" [color="0 0 0.5273591806876372", label="1.12",style=bold];
"MYH7_S" -> "MC5R_M" [color="0 0 0.5267008046817849", label="1.12",style=bold];
"TDRD5_S" -> "DSCAM_E" [color="0 0 0.5260424286759326", label="1.12",style=bold];
"GABRB3_S" -> "PCDHA12_S" [color="0 0 0.5253840526700805", label="1.12",style=bold];
"LCT_S" -> "ADAMTS20_S" [color="0 0 0.5247256766642283", label="1.13",style=bold];
"TUBA3C_S" -> "TDRD5_S" [color="0 0 0.524067300658376", label="1.13",style=bold];
"MRGPRX4_S" -> "NTSR2_M" [color="0 0 0.5234089246525238", label="1.13",style=bold];
"UGT2A1_S" -> "LCT_M" [color="0 0 0.5227505486466716", label="1.13",style=bold];
"MRGPRX4_S" -> "MAB21L2_E" [color="0 0 0.5220921726408194", label="1.13",style=bold];
"LCT_S" -> "BEST2_M" [color="0 0 0.5214337966349671", label="1.13",style=bold];
"PCDHA12_S" -> "LCT_M" [color="0 0 0.5207754206291149", label="1.13",style=bold];
"LPA_S" -> "FREM2_E" [color="0 0 0.5201170446232626", label="1.14",style=bold];
"CPNE6_S" -> "UGT2A1_E" [color="0 0 0.5194586686174104", label="1.14",style=bold];
"PCDHA10_S" -> "ABCA13_S" [color="0 0 0.5188002926115582", label="1.14",style=bold];
"CCDC67_S" -> "CNTN5_M" [color="0 0 0.518141916605706", label="1.15",style=bold];
"NTSR2_S" -> "HFM1_M" [color="0 0 0.5174835405998537", label="1.15",style=bold];
"MYBPC2_S" -> "CDH19_E" [color="0 0 0.5168251645940015", label="1.15",style=bold];
"KRT1_S" -> "LCE2C_E" [color="0 0 0.5161667885881492", label="1.15",style=bold];
"PCDHA10_S" -> "RYR2_E" [color="0 0 0.5155084125822971", label="1.15",style=bold];
"FREM2_S" -> "ERN2_M" [color="0 0 0.5148500365764448", label="1.16",style=bold];
"PCDH10_S" -> "MYH6_E" [color="0 0 0.5141916605705925", label="1.16",style=bold];
"LINGO2_S" -> "MYBPC2_M" [color="0 0 0.5135332845647403", label="1.16",style=bold];
"HFM1_S" -> "MYH13_E" [color="0 0 0.5128749085588882", label="1.16",style=bold];
"CPNE6_S" -> "LPA_E" [color="0 0 0.5122165325530359", label="1.17",style=bold];
"BEST2_S" -> "DNAH11_M" [color="0 0 0.5115581565471836", label="1.17",style=bold];
"MYBPC2_S" -> "FREM2_M" [color="0 0 0.5108997805413313", label="1.17",style=bold];
"MYH13_S" -> "CNTN5_E" [color="0 0 0.5102414045354792", label="1.18",style=bold];
"NELL1_S" -> "LINGO2_S" [color="0 0 0.509583028529627", label="1.18",style=bold];
"LRAT_S" -> "LINGO2_S" [color="0 0 0.5089246525237747", label="1.18",style=bold];
"BEST2_S" -> "FSTL5_E" [color="0 0 0.5082662765179224", label="1.18",style=bold];
"MYH7_S" -> "UGT2A1_E" [color="0 0 0.5076079005120703", label="1.18",style=bold];
"MYO3A_S" -> "GALNTL6_E" [color="0 0 0.5069495245062181", label="1.18",style=bold];
"ABCA13_S" -> "LCE2C_E" [color="0 0 0.5062911485003658", label="1.18",style=bold];
"MAB21L2_S" -> "MYH13_M" [color="0 0 0.5056327724945135", label="1.19",style=bold];
"MYH6_S" -> "GALNTL6_E" [color="0 0 0.5049743964886613", label="1.19",style=bold];
"MYH13_S" -> "TUBA3C_E" [color="0 0 0.5043160204828091", label="1.19",style=bold];
"MYH13_S" -> "CSMD1_S" [color="0 0 0.5036576444769569", label="1.19",style=bold];
"PCDHA10_S" -> "MUC4_E" [color="0 0 0.5029992684711047", label="1.20",style=bold];
"ZIC1_S" -> "KRT1_M" [color="0 0 0.5023408924652524", label="1.20",style=bold];
"PCDH10_S" -> "GABRB3_M" [color="0 0 0.5016825164594001", label="1.20",style=bold];
"KRT1_S" -> "DNAH11_E" [color="0 0 0.5010241404535479", label="1.20",style=bold];
"LINGO2_S" -> "NELL1_E" [color="0 0 0.5003657644476958", label="1.20",style=bold];
"CCDC67_S" -> "MRGPRX4_M" [color="0 0 0.4997073884418435", label="1.20",style=bold];
"MYBPC2_S" -> "MRGPRX4_M" [color="0 0 0.49904901243599126", label="1.20",style=bold];
"CCDC67_S" -> "MYH13_M" [color="0 0 0.49839063643013903", label="1.20",style=bold];
"PCDHA12_S" -> "ATP13A4_E" [color="0 0 0.4977322604242868", label="1.20",style=bold];
"KRT1_S" -> "CNTNAP5_M" [color="0 0 0.4970738844184346", label="1.20",style=bold];
"MAB21L2_S" -> "ZIC1_E" [color="0 0 0.4964155084125823", label="1.20",style=bold];
"LPPR1_S" -> "PKHD1L1_E" [color="0 0 0.4957571324067301", label="1.20",style=bold];
"MC5R_S" -> "TDRD5_E" [color="0 0 0.49509875640087786", label="1.20",style=bold];
"MRGPRX4_S" -> "CSMD1_M" [color="0 0 0.49444038039502564", label="1.21",style=bold];
"PCDHA10_S" -> "ERBB4_S" [color="0 0 0.4937820043891734", label="1.21",style=bold];
"KRT1_S" -> "LRP1B_S" [color="0 0 0.4931236283833212", label="1.21",style=bold];
"RP1_S" -> "BEST2_E" [color="0 0 0.49246525237746896", label="1.21",style=bold];
"LPPR1_S" -> "PKHD1L1_M" [color="0 0 0.4918068763716167", label="1.21",style=bold];
"CDH19_S" -> "PCDH10_M" [color="0 0 0.49114850036576446", label="1.23",style=bold];
"CCDC67_S" -> "NPFFR2_E" [color="0 0 0.49049012435991224", label="1.23",style=bold];
"NPFFR2_S" -> "TUBA3C_E" [color="0 0 0.48983174835406", label="1.23",style=bold];
"MYH7_S" -> "CNTN5_M" [color="0 0 0.4891733723482078", label="1.24",style=bold];
"PCDHA10_S" -> "RP1_E" [color="0 0 0.48851499634235557", label="1.24",style=bold];
"GALNTL6_S" -> "PCDHA10_S" [color="0 0 0.48785662033650334", label="1.24",style=bold];
"MYBPC2_S" -> "RYR2_M" [color="0 0 0.4871982443306511", label="1.25",style=bold];
"ERN2_S" -> "ABCA13_E" [color="0 0 0.48653986832479884", label="1.25",style=bold];
"MRGPRX4_S" -> "KRT24_M" [color="0 0 0.4858814923189466", label="1.25",style=bold];
"ZIC1_S" -> "FSTL5_S" [color="0 0 0.4852231163130944", label="1.25",style=bold];
"MC5R_S" -> "ERN2_E" [color="0 0 0.48456474030724217", label="1.26",style=bold];
"NPFFR2_S" -> "RBP3_E" [color="0 0 0.48390636430138995", label="1.26",style=bold];
"DUSP27_S" -> "MYO3A_E" [color="0 0 0.4832479882955377", label="1.26",style=bold];
"ERBB4_S" -> "MYH13_M" [color="0 0 0.4825896122896855", label="1.27",style=bold];
"CDH19_S" -> "COL11A1_E" [color="0 0 0.4819312362838332", label="1.27",style=bold];
"MYH6_S" -> "CR2_M" [color="0 0 0.481272860277981", label="1.27",style=bold];
"MC5R_S" -> "DSCAM_M" [color="0 0 0.4806144842721288", label="1.27",style=bold];
"NTSR2_S" -> "ATP13A4_E" [color="0 0 0.47995610826627655", label="1.27",style=bold];
"LINGO2_S" -> "LPA_E" [color="0 0 0.4792977322604243", label="1.28",style=bold];
"MRGPRX4_S" -> "ZIC1_M" [color="0 0 0.4786393562545721", label="1.28",style=bold];
"PCDHA12_S" -> "GABRB3_M" [color="0 0 0.4779809802487199", label="1.29",style=bold];
"CCDC67_S" -> "CNTNAP5_E" [color="0 0 0.4773226042428676", label="1.30",style=bold];
"CCDC67_S" -> "CSMD1_M" [color="0 0 0.4766642282370154", label="1.30",style=bold];
"MRGPRX4_S" -> "MYH6_E" [color="0 0 0.47600585223116315", label="1.30",style=bold];
"LPPR1_S" -> "ADAMTS20_E" [color="0 0 0.4753474762253109", label="1.30",style=bold];
"MYH13_S" -> "PTPRT_E" [color="0 0 0.4746891002194587", label="1.30",style=bold];
"KRT24_S" -> "Survival" [color="0 0 0.4740307242136065", label="1.30",style=bold];
"ATP13A4_S" -> "MAB21L2_E" [color="0 0 0.47337234820775426", label="1.30",style=bold];
"LINGO2_S" -> "MYO3A_E" [color="0 0 0.47271397220190203", label="1.30",style=bold];
"MRGPRX4_S" -> "NTSR2_E" [color="0 0 0.47205559619604975", label="1.30",style=bold];
"DUSP27_S" -> "DNAH11_M" [color="0 0 0.47139722019019753", label="1.31",style=bold];
"MYBPC2_S" -> "PCDHA12_M" [color="0 0 0.4707388441843453", label="1.31",style=bold];
"MYH13_S" -> "MYO3A_E" [color="0 0 0.4700804681784931", label="1.32",style=bold];
"FSTL5_S" -> "MYO3A_E" [color="0 0 0.46942209217264086", label="1.32",style=bold];
"CNTNAP5_S" -> "COL11A1_M" [color="0 0 0.46876371616678864", label="1.32",style=bold];
"CR2_S" -> "LRRTM4_E" [color="0 0 0.4681053401609364", label="1.32",style=bold];
"DNAH11_S" -> "RYR2_S" [color="0 0 0.46744696415508413", label="1.33",style=bold];
"MUC4_S" -> "PKHD1L1_M" [color="0 0 0.4667885881492319", label="1.34",style=bold];
"ZIC1_S" -> "DUSP27_M" [color="0 0 0.4661302121433797", label="1.35",style=bold];
"LCT_S" -> "NEB_M" [color="0 0 0.46547183613752746", label="1.35",style=bold];
"LRAT_S" -> "FREM2_S" [color="0 0 0.46481346013167524", label="1.35",style=bold];
"ATP13A4_S" -> "COL11A1_E" [color="0 0 0.464155084125823", label="1.36",style=bold];
"DUSP27_S" -> "RYR2_E" [color="0 0 0.4634967081199708", label="1.36",style=bold];
"CCDC67_S" -> "CPNE6_M" [color="0 0 0.46283833211411857", label="1.36",style=bold];
"FSTL5_S" -> "ADAMTS20_E" [color="0 0 0.4621799561082663", label="1.36",style=bold];
"MC5R_S" -> "LPA_M" [color="0 0 0.46152158010241406", label="1.36",style=bold];
"CPNE6_S" -> "CPNE6_M" [color="0 0 0.46086320409656184", label="1.36",style=bold];
"MYH7_S" -> "CNTN5_E" [color="0 0 0.4602048280907096", label="1.36",style=bold];
"ATP13A4_S" -> "RYR2_E" [color="0 0 0.4595464520848574", label="1.36",style=bold];
"TUBA3C_S" -> "LRRTM4_E" [color="0 0 0.45888807607900517", label="1.36",style=bold];
"DNAH11_S" -> "CDKN2A_E" [color="0 0 0.45822970007315295", label="1.37",style=bold];
"NTSR2_S" -> "CSMD1_M" [color="0 0 0.45757132406730067", label="1.37",style=bold];
"MYO3A_S" -> "GALNTL6_M" [color="0 0 0.45691294806144844", label="1.38",style=bold];
"CCDC67_S" -> "MUC16_S" [color="0 0 0.4562545720555962", label="1.38",style=bold];
"NELL1_S" -> "MYBPC2_E" [color="0 0 0.455596196049744", label="1.38",style=bold];
"MYH7_S" -> "TTN_M" [color="0 0 0.45493782004389177", label="1.38",style=bold];
"TUBA3C_S" -> "CDH19_S" [color="0 0 0.45427944403803955", label="1.38",style=bold];
"NPFFR2_S" -> "CDH19_S" [color="0 0 0.4536210680321873", label="1.38",style=bold];
"GALNTL6_S" -> "MC5R_S" [color="0 0 0.45296269202633505", label="1.38",style=bold];
"CCDC67_S" -> "CPNE6_E" [color="0 0 0.4523043160204828", label="1.39",style=bold];
"CPNE6_S" -> "CDH19_E" [color="0 0 0.4516459400146306", label="1.39",style=bold];
"CR2_S" -> "NTSR2_M" [color="0 0 0.4509875640087784", label="1.39",style=bold];
"NPFFR2_S" -> "ZIC1_M" [color="0 0 0.45032918800292615", label="1.39",style=bold];
"MYH7_S" -> "ERBB4_M" [color="0 0 0.4496708119970739", label="1.39",style=bold];
"MRGPRX4_S" -> "COL11A1_M" [color="0 0 0.4490124359912217", label="1.39",style=bold];
"ATP13A4_S" -> "LCE2C_M" [color="0 0 0.4483540599853695", label="1.39",style=bold];
"ADAMTS20_S" -> "CDKN2A_E" [color="0 0 0.4476956839795172", label="1.39",style=bold];
"MAB21L2_S" -> "RYR2_S" [color="0 0 0.447037307973665", label="1.39",style=bold];
"LCT_S" -> "TDRD5_M" [color="0 0 0.44637893196781275", label="1.39",style=bold];
"CSMD1_S" -> "CSMD3_E" [color="0 0 0.44572055596196053", label="1.40",style=bold];
"LCT_S" -> "CSMD3_E" [color="0 0 0.4450621799561083", label="1.40",style=bold];
"LCT_S" -> "DUSP27_E" [color="0 0 0.4444038039502561", label="1.40",style=bold];
"CDH19_S" -> "RBP3_E" [color="0 0 0.44374542794440386", label="1.40",style=bold];
"TUBA3C_S" -> "CDKN2A_S" [color="0 0 0.4430870519385516", label="1.40",style=bold];
"PCDHA12_S" -> "CDKN2A_S" [color="0 0 0.44242867593269936", label="1.40",style=bold];
"MYH7_S" -> "MUC16_M" [color="0 0 0.44177029992684713", label="1.40",style=bold];
"LPPR1_S" -> "TTN_E" [color="0 0 0.4411119239209949", label="1.41",style=bold];
"LPA_S" -> "NELL1_M" [color="0 0 0.4404535479151427", label="1.42",style=bold];
"GABRB3_S" -> "BEST2_E" [color="0 0 0.43979517190929046", label="1.42",style=bold];
"PCDHA12_S" -> "CDH19_M" [color="0 0 0.43913679590343824", label="1.43",style=bold];
"MYH7_S" -> "DNAH11_M" [color="0 0 0.438478419897586", label="1.43",style=bold];
"CDH19_S" -> "TUBA3C_E" [color="0 0 0.43782004389173373", label="1.44",style=bold];
"CR2_S" -> "MYO3A_E" [color="0 0 0.4371616678858815", label="1.44",style=bold];
"CSMD1_S" -> "NELL1_S" [color="0 0 0.4365032918800293", label="1.44",style=bold];
"LPPR1_S" -> "MRGPRX4_E" [color="0 0 0.43584491587417706", label="1.45",style=bold];
"NPFFR2_S" -> "PCDH10_S" [color="0 0 0.43518653986832484", label="1.45",style=bold];
"PCDH10_S" -> "GALNTL6_E" [color="0 0 0.4345281638624726", label="1.45",style=bold];
"MYO3A_S" -> "DNAH11_E" [color="0 0 0.4338697878566204", label="1.46",style=bold];
"LCT_S" -> "ERN2_M" [color="0 0 0.4332114118507681", label="1.47",style=bold];
"MAB21L2_S" -> "NEB_M" [color="0 0 0.4325530358449159", label="1.47",style=bold];
"TUBA3C_S" -> "NEB_M" [color="0 0 0.43189465983906367", label="1.47",style=bold];
"NTSR2_S" -> "NEB_M" [color="0 0 0.43123628383321144", label="1.47",style=bold];
"TUBA3C_S" -> "CSMD3_M" [color="0 0 0.4305779078273592", label="1.47",style=bold];
"KRT1_S" -> "MUC4_E" [color="0 0 0.429919531821507", label="1.47",style=bold];
"PCDH10_S" -> "TDRD5_M" [color="0 0 0.42926115581565477", label="1.47",style=bold];
"DNAH11_S" -> "PCDHA12_E" [color="0 0 0.42860277980980255", label="1.47",style=bold];
"GALNTL6_S" -> "CNTNAP5_S" [color="0 0 0.42794440380395027", label="1.47",style=bold];
"DUSP27_S" -> "CSMD3_E" [color="0 0 0.42728602779809804", label="1.48",style=bold];
"MYH7_S" -> "GALNTL6_E" [color="0 0 0.4266276517922458", label="1.48",style=bold];
"MYH7_S" -> "HFM1_M" [color="0 0 0.4259692757863936", label="1.48",style=bold];
"ADAMTS20_S" -> "NELL1_E" [color="0 0 0.4253108997805414", label="1.49",style=bold];
"PCDH10_S" -> "ZIC1_E" [color="0 0 0.42465252377468915", label="1.49",style=bold];
"MYBPC2_S" -> "COL11A1_E" [color="0 0 0.4239941477688369", label="1.49",style=bold];
"NPFFR2_S" -> "LINGO2_M" [color="0 0 0.42333577176298465", label="1.49",style=bold];
"TUBA3C_S" -> "ABCA13_M" [color="0 0 0.4226773957571324", label="1.49",style=bold];
"ATP13A4_S" -> "ZIC1_E" [color="0 0 0.4220190197512802", label="1.49",style=bold];
"NTSR2_S" -> "TUBA3C_M" [color="0 0 0.421360643745428", label="1.49",style=bold];
"DUSP27_M" -> "RP1_M" [color="0 0 0.42070226773957575", label="1.50",style=bold];
"NTSR2_S" -> "ATP13A4_M" [color="0 0 0.42004389173372353", label="1.50",style=bold];
"ADAMTS20_S" -> "GABRB3_E" [color="0 0 0.4193855157278713", label="1.50",style=bold];
"CNTNAP5_S" -> "RYR2_E" [color="0 0 0.418727139722019", label="1.50",style=bold];
"RP1_S" -> "MUC16_E" [color="0 0 0.4180687637161668", label="1.51",style=bold];
"LRAT_S" -> "MUC4_M" [color="0 0 0.4174103877103146", label="1.52",style=bold];
"MAB21L2_S" -> "MUC4_S" [color="0 0 0.41675201170446236", label="1.52",style=bold];
"FREM2_S" -> "PCDHA12_E" [color="0 0 0.41609363569861013", label="1.53",style=bold];
"PCDHA10_S" -> "TDRD5_E" [color="0 0 0.4154352596927579", label="1.53",style=bold];
"PCDH10_S" -> "DNAH11_E" [color="0 0 0.4147768836869057", label="1.53",style=bold];
"LRAT_S" -> "LRP1B_S" [color="0 0 0.41411850768105346", label="1.54",style=bold];
"MRGPRX4_M" -> "LCE2C_M" [color="0 0 0.4134601316752012", label="1.54",style=bold];
"DUSP27_S" -> "CCDC67_M" [color="0 0 0.41280175566934896", label="1.54",style=bold];
"LCE2C_S" -> "CNTN5_E" [color="0 0 0.41214337966349673", label="1.54",style=bold];
"UGT2A1_S" -> "ERBB4_E" [color="0 0 0.4114850036576445", label="1.54",style=bold];
"CR2_S" -> "CSMD1_S" [color="0 0 0.4108266276517923", label="1.54",style=bold];
"NTSR2_S" -> "CDKN2A_E" [color="0 0 0.41016825164594006", label="1.54",style=bold];
"CSMD1_S" -> "ABCA13_E" [color="0 0 0.40950987564008784", label="1.55",style=bold];
"ATP13A4_S" -> "PTPRT_S" [color="0 0 0.40885149963423556", label="1.55",style=bold];
"LCT_S" -> "CSMD1_S" [color="0 0 0.40819312362838334", label="1.55",style=bold];
"RP1_S" -> "TDRD5_E" [color="0 0 0.4075347476225311", label="1.55",style=bold];
"MYBPC2_S" -> "RP1_M" [color="0 0 0.4068763716166789", label="1.56",style=bold];
"DUSP27_S" -> "KRT1_M" [color="0 0 0.40621799561082667", label="1.56",style=bold];
"ZIC1_S" -> "LRRTM4_M" [color="0 0 0.40555961960497444", label="1.56",style=bold];
"KRT1_S" -> "PCDHA12_E" [color="0 0 0.4049012435991222", label="1.56",style=bold];
"CDH19_S" -> "FSTL5_E" [color="0 0 0.40424286759327", label="1.56",style=bold];
"DNAH11_S" -> "BEST2_E" [color="0 0 0.4035844915874177", label="1.58",style=bold];
"TDRD5_S" -> "GABRB3_M" [color="0 0 0.4029261155815655", label="1.59",style=bold];
"LPPR1_S" -> "NEB_S" [color="0 0 0.40226773957571327", label="1.59",style=bold];
"CDH19_S" -> "LINGO2_M" [color="0 0 0.40160936356986104", label="1.59",style=bold];
"MRGPRX4_S" -> "LINGO2_M" [color="0 0 0.4009509875640088", label="1.59",style=bold];
"MAB21L2_S" -> "LINGO2_E" [color="0 0 0.4002926115581566", label="1.59",style=bold];
"MYBPC2_S" -> "RP1_S" [color="0 0 0.3996342355523044", label="1.59",style=bold];
"LCE2C_S" -> "MUC16_M" [color="0 0 0.3989758595464521", label="1.61",style=bold];
"TUBA3C_S" -> "CSMD3_E" [color="0 0 0.3983174835405999", label="1.61",style=bold];
"MYH6_S" -> "MYH6_E" [color="0 0 0.39765910753474765", label="1.61",style=bold];
"KRT1_S" -> "MYH6_E" [color="0 0 0.3970007315288955", label="1.61",style=bold];
"NPFFR2_S" -> "MYH13_E" [color="0 0 0.3963423555230432", label="1.61",style=bold];
"GALNTL6_S" -> "TDRD5_M" [color="0 0 0.3956839795171909", label="1.61",style=bold];
"CR2_S" -> "MYH6_E" [color="0 0 0.39502560351133875", label="1.61",style=bold];
"TUBA3C_S" -> "LRAT_M" [color="0 0 0.3943672275054865", label="1.62",style=bold];
"NTSR2_S" -> "DNAH11_E" [color="0 0 0.3937088514996343", label="1.62",style=bold];
"DSCAM_S" -> "RBP3_S" [color="0 0 0.393050475493782", label="1.62",style=bold];
"MYBPC2_S" -> "DSCAM_S" [color="0 0 0.39239209948792986", label="1.63",style=bold];
"RBP3_S" -> "ADAMTS20_E" [color="0 0 0.3917337234820776", label="1.63",style=bold];
"PCDHA10_S" -> "MYH6_E" [color="0 0 0.3910753474762253", label="1.64",style=bold];
"DSCAM_S" -> "BEST2_E" [color="0 0 0.39041697147037313", label="1.65",style=bold];
"ZIC1_S" -> "LPA_M" [color="0 0 0.38975859546452085", label="1.65",style=bold];
"PCDH10_S" -> "CDH19_E" [color="0 0 0.3891002194586687", label="1.66",style=bold];
"ERBB4_S" -> "CNTN5_E" [color="0 0 0.3884418434528164", label="1.66",style=bold];
"NTSR2_S" -> "RP1_M" [color="0 0 0.38778346744696424", label="1.67",style=bold];
"LRAT_S" -> "KRT1_M" [color="0 0 0.38712509144111196", label="1.67",style=bold];
"DUSP27_S" -> "LPPR1_M" [color="0 0 0.3864667154352597", label="1.68",style=bold];
"LINGO2_S" -> "MUC4_E" [color="0 0 0.3858083394294075", label="1.70",style=bold];
"LPA_S" -> "HFM1_E" [color="0 0 0.38514996342355523", label="1.70",style=bold];
"MC5R_S" -> "LPPR1_E" [color="0 0 0.38449158741770306", label="1.70",style=bold];
"BEST2_S" -> "CSMD3_E" [color="0 0 0.3838332114118508", label="1.70",style=bold];
"NTSR2_S" -> "COL11A1_M" [color="0 0 0.3831748354059986", label="1.70",style=bold];
"MYBPC2_S" -> "COL11A1_M" [color="0 0 0.38251645940014634", label="1.71",style=bold];
"TDRD5_S" -> "TUBA3C_E" [color="0 0 0.38185808339429406", label="1.72",style=bold];
"MYBPC2_S" -> "ADAMTS20_M" [color="0 0 0.3811997073884419", label="1.73",style=bold];
"NPFFR2_S" -> "MYH13_S" [color="0 0 0.3805413313825896", label="1.73",style=bold];
"PCDH10_S" -> "MC5R_E" [color="0 0 0.37988295537673744", label="1.73",style=bold];
"KRT24_S" -> "KRT24_M" [color="0 0 0.37922457937088516", label="1.73",style=bold];
"NPFFR2_S" -> "CDKN2A_E" [color="0 0 0.378566203365033", label="1.73",style=bold];
"NTSR2_S" -> "CNTN5_S" [color="0 0 0.3779078273591807", label="1.74",style=bold];
"UGT2A1_S" -> "FSTL5_S" [color="0 0 0.37724945135332844", label="1.74",style=bold];
"NELL1_S" -> "PCDH10_M" [color="0 0 0.37659107534747627", label="1.74",style=bold];
"TTN_E" -> "UGT2A1_E" [color="0 0 0.375932699341624", label="1.75",style=bold];
"NPFFR2_S" -> "KRT24_M" [color="0 0 0.3752743233357718", label="1.75",style=bold];
"LRRTM4_S" -> "MUC16_S" [color="0 0 0.37461594732991954", label="1.76",style=bold];
"CSMD1_S" -> "NEB_M" [color="0 0 0.3739575713240674", label="1.76",style=bold];
"RBP3_S" -> "CSMD3_S" [color="0 0 0.3732991953182151", label="1.77",style=bold];
"DUSP27_S" -> "ERBB4_S" [color="0 0 0.3726408193123629", label="1.77",style=bold];
"MYBPC2_S" -> "PCLO_M" [color="0 0 0.37198244330651065", label="1.77",style=bold];
"NELL1_S" -> "MUC4_M" [color="0 0 0.37132406730065837", label="1.77",style=bold];
"MYH6_M" -> "LCE2C_M" [color="0 0 0.3706656912948062", label="1.78",style=bold];
"CR2_S" -> "ADAMTS20_M" [color="0 0 0.3700073152889539", label="1.78",style=bold];
"PTPRT_S" -> "COL11A1_E" [color="0 0 0.36934893928310175", label="1.78",style=bold];
"MC5R_S" -> "PKHD1L1_S" [color="0 0 0.3686905632772495", label="1.78",style=bold];
"MAB21L2_S" -> "ERBB4_M" [color="0 0 0.3680321872713973", label="1.79",style=bold];
"DUSP27_S" -> "PCDHA10_E" [color="0 0 0.367373811265545", label="1.79",style=bold];
"CPNE6_S" -> "FSTL5_E" [color="0 0 0.36671543525969275", label="1.79",style=bold];
"NPFFR2_S" -> "FSTL5_M" [color="0 0 0.3660570592538406", label="1.79",style=bold];
"ADAMTS20_S" -> "PTPRT_E" [color="0 0 0.3653986832479883", label="1.79",style=bold];
"UGT2A1_S" -> "CCDC67_S" [color="0 0 0.36474030724213613", label="1.79",style=bold];
"LPA_S" -> "COL11A1_E" [color="0 0 0.36408193123628385", label="1.79",style=bold];
"PCDH10_S" -> "MYO3A_M" [color="0 0 0.3634235552304317", label="1.80",style=bold];
"MYBPC2_S" -> "MYO3A_M" [color="0 0 0.3627651792245794", label="1.80",style=bold];
"PCDH10_S" -> "Survival" [color="0 0 0.3621068032187271", label="1.82",style=bold];
"NTSR2_S" -> "CDH19_E" [color="0 0 0.36144842721287496", label="1.85",style=bold];
"ADAMTS20_S" -> "FSTL5_E" [color="0 0 0.3607900512070227", label="1.85",style=bold];
"CR2_S" -> "FLG_M" [color="0 0 0.3601316752011705", label="1.86",style=bold];
"MYH6_S" -> "MC5R_S" [color="0 0 0.35947329919531823", label="1.86",style=bold];
"ZIC1_S" -> "ATP13A4_M" [color="0 0 0.35881492318946606", label="1.86",style=bold];
"ADAMTS20_S" -> "ADAMTS20_M" [color="0 0 0.3581565471836138", label="1.86",style=bold];
"ERBB4_S" -> "PCDH10_S" [color="0 0 0.3574981711777615", label="1.87",style=bold];
"MYH7_S" -> "NPFFR2_M" [color="0 0 0.35683979517190934", label="1.87",style=bold];
"DUSP27_S" -> "DSCAM_E" [color="0 0 0.35618141916605706", label="1.87",style=bold];
"FSTL5_S" -> "CNTNAP5_E" [color="0 0 0.3555230431602049", label="1.88",style=bold];
"MYH7_S" -> "FLG_S" [color="0 0 0.3548646671543526", label="1.89",style=bold];
"CNTNAP5_S" -> "CDKN2A_M" [color="0 0 0.35420629114850044", label="1.90",style=bold];
"MRGPRX4_S" -> "LPPR1_E" [color="0 0 0.35354791514264816", label="1.90",style=bold];
"CPNE6_S" -> "HFM1_S" [color="0 0 0.352889539136796", label="1.91",style=bold];
"CR2_S" -> "CSMD3_S" [color="0 0 0.3522311631309437", label="1.91",style=bold];
"ABCA13_S" -> "TUBA3C_M" [color="0 0 0.35157278712509143", label="1.92",style=bold];
"GALNTL6_S" -> "MYH6_S" [color="0 0 0.35091441111923927", label="1.92",style=bold];
"GALNTL6_S" -> "LCT_E" [color="0 0 0.350256035113387", label="1.93",style=bold];
"ZIC1_S" -> "LPA_S" [color="0 0 0.3495976591075348", label="1.94",style=bold];
"MUC4_S" -> "FSTL5_M" [color="0 0 0.34893928310168254", label="1.94",style=bold];
"UGT2A1_E" -> "MUC4_E" [color="0 0 0.34828090709583037", label="1.94",style=bold];
"NEB_S" -> "FLG_S" [color="0 0 0.3476225310899781", label="1.94",style=bold];
"CDH19_S" -> "CCDC67_M" [color="0 0 0.3469641550841258", label="1.95",style=bold];
"TDRD5_S" -> "MAB21L2_S" [color="0 0 0.34630577907827365", label="1.95",style=bold];
"MYH6_S" -> "CSMD3_S" [color="0 0 0.34564740307242137", label="1.96",style=bold];
"MYBPC2_S" -> "MRGPRX4_E" [color="0 0 0.3449890270665692", label="1.96",style=bold];
"LPPR1_S" -> "CNTNAP5_M" [color="0 0 0.3443306510607169", label="1.99",style=bold];
"MRGPRX4_S" -> "CSMD3_S" [color="0 0 0.34367227505486475", label="1.99",style=bold];
"NTSR2_S" -> "FREM2_M" [color="0 0 0.34301389904901247", label="1.99",style=bold];
"MYBPC2_S" -> "ZIC1_M" [color="0 0 0.3423555230431602", label="1.99",style=bold];
"MAB21L2_S" -> "LPA_S" [color="0 0 0.341697147037308", label="2.01",style=bold];
"MAB21L2_S" -> "PTPRT_S" [color="0 0 0.34103877103145575", label="2.01",style=bold];
"PCDH10_S" -> "NEB_E" [color="0 0 0.3403803950256036", label="2.02",style=bold];
"CCDC67_S" -> "LINGO2_S" [color="0 0 0.3397220190197513", label="2.02",style=bold];
"CNTN5_S" -> "LINGO2_M" [color="0 0 0.33906364301389913", label="2.03",style=bold];
"ZIC1_S" -> "LRP1B_S" [color="0 0 0.33840526700804685", label="2.03",style=bold];
"PCDHA12_S" -> "TTN_E" [color="0 0 0.33774689100219457", label="2.04",style=bold];
"MYH7_S" -> "NEB_M" [color="0 0 0.3370885149963424", label="2.04",style=bold];
"CNTN5_S" -> "TTN_S" [color="0 0 0.3364301389904901", label="2.05",style=bold];
"MYH13_S" -> "ZIC1_E" [color="0 0 0.33577176298463796", label="2.05",style=bold];
"ERBB4_S" -> "LRAT_E" [color="0 0 0.3351133869787857", label="2.06",style=bold];
"DSCAM_S" -> "PCLO_S" [color="0 0 0.3344550109729335", label="2.06",style=bold];
"PCDHA10_S" -> "MAB21L2_M" [color="0 0 0.33379663496708123", label="2.07",style=bold];
"ZIC1_S" -> "CR2_M" [color="0 0 0.33313825896122895", label="2.09",style=bold];
"MYBPC2_S" -> "NPFFR2_M" [color="0 0 0.3324798829553768", label="2.09",style=bold];
"CNTNAP5_S" -> "ERN2_M" [color="0 0 0.3318215069495245", label="2.10",style=bold];
"DNAH11_S" -> "ERBB4_M" [color="0 0 0.33116313094367233", label="2.10",style=bold];
"TTN_M" -> "LCT_M" [color="0 0 0.33050475493782006", label="2.10",style=bold];
"CNTNAP5_S" -> "COL11A1_E" [color="0 0 0.3298463789319679", label="2.12",style=bold];
"NPFFR2_S" -> "LPPR1_E" [color="0 0 0.3291880029261156", label="2.12",style=bold];
"UGT2A1_S" -> "CNTNAP5_M" [color="0 0 0.32852962692026344", label="2.12",style=bold];
"MUC4_S" -> "CDH19_E" [color="0 0 0.32787125091441116", label="2.13",style=bold];
"MRGPRX4_S" -> "TUBA3C_M" [color="0 0 0.3272128749085589", label="2.13",style=bold];
"NELL1_S" -> "CDH19_E" [color="0 0 0.3265544989027067", label="2.14",style=bold];
"ZIC1_S" -> "LRRTM4_S" [color="0 0 0.32589612289685443", label="2.14",style=bold];
"PTPRT_S" -> "LPA_E" [color="0 0 0.32523774689100227", label="2.15",style=bold];
"RYR2_E" -> "TUBA3C_E" [color="0 0 0.32457937088515", label="2.15",style=bold];
"MYO3A_S" -> "MUC4_E" [color="0 0 0.3239209948792978", label="2.15",style=bold];
"PCDHA12_M" -> "MC5R_M" [color="0 0 0.32326261887344554", label="2.18",style=bold];
"CR2_S" -> "CR2_E" [color="0 0 0.32260424286759326", label="2.18",style=bold];
"ABCA13_S" -> "PTPRT_E" [color="0 0 0.3219458668617411", label="2.19",style=bold];
"DUSP27_S" -> "LINGO2_M" [color="0 0 0.3212874908558888", label="2.19",style=bold];
"NPFFR2_S" -> "FSTL5_E" [color="0 0 0.32062911485003665", label="2.20",style=bold];
"LPA_S" -> "LPPR1_E" [color="0 0 0.31997073884418437", label="2.20",style=bold];
"NTSR2_S" -> "MYH7_M" [color="0 0 0.3193123628383322", label="2.22",style=bold];
"GABRB3_S" -> "ABCA13_M" [color="0 0 0.3186539868324799", label="2.23",style=bold];
"MYH13_S" -> "CCDC67_E" [color="0 0 0.31799561082662764", label="2.23",style=bold];
"NTSR2_S" -> "MYBPC2_E" [color="0 0 0.31733723482077547", label="2.23",style=bold];
"TUBA3C_S" -> "FREM2_S" [color="0 0 0.3166788588149232", label="2.24",style=bold];
"MYH6_S" -> "CSMD1_S" [color="0 0 0.316020482809071", label="2.25",style=bold];
"RYR2_S" -> "HFM1_E" [color="0 0 0.31536210680321874", label="2.25",style=bold];
"LINGO2_S" -> "MRGPRX4_M" [color="0 0 0.3147037307973666", label="2.26",style=bold];
"MYH7_S" -> "ABCA13_E" [color="0 0 0.3140453547915143", label="2.27",style=bold];
"LCE2C_S" -> "PTPRT_E" [color="0 0 0.313386978785662", label="2.28",style=bold];
"TUBA3C_S" -> "PCLO_S" [color="0 0 0.31272860277980985", label="2.29",style=bold];
"LINGO2_S" -> "UGT2A1_M" [color="0 0 0.31207022677395757", label="2.29",style=bold];
"KRT24_S" -> "DSCAM_S" [color="0 0 0.3114118507681054", label="2.29",style=bold];
"MC5R_E" -> "CDH19_E" [color="0 0 0.3107534747622531", label="2.30",style=bold];
"PCDHA10_S" -> "CCDC67_M" [color="0 0 0.31009509875640096", label="2.30",style=bold];
"MRGPRX4_S" -> "MYBPC2_M" [color="0 0 0.3094367227505487", label="2.30",style=bold];
"CDH19_S" -> "LRAT_E" [color="0 0 0.3087783467446964", label="2.30",style=bold];
"LINGO2_S" -> "TUBA3C_E" [color="0 0 0.30811997073884423", label="2.30",style=bold];
"LPA_S" -> "TUBA3C_E" [color="0 0 0.30746159473299195", label="2.31",style=bold];
"LPA_S" -> "NEB_E" [color="0 0 0.3068032187271398", label="2.31",style=bold];
"DNAH11_M" -> "CPNE6_M" [color="0 0 0.3061448427212875", label="2.32",style=bold];
"NTSR2_S" -> "ERBB4_S" [color="0 0 0.30548646671543533", label="2.32",style=bold];
"MYH6_S" -> "COL11A1_S" [color="0 0 0.30482809070958305", label="2.32",style=bold];
"MC5R_S" -> "MUC16_S" [color="0 0 0.3041697147037309", label="2.32",style=bold];
"DUSP27_S" -> "LPA_M" [color="0 0 0.3035113386978786", label="2.38",style=bold];
"CNTNAP5_S" -> "ZIC1_S" [color="0 0 0.30285296269202633", label="2.38",style=bold];
"MRGPRX4_S" -> "MC5R_E" [color="0 0 0.30219458668617416", label="2.38",style=bold];
"MYH6_S" -> "CR2_S" [color="0 0 0.3015362106803219", label="2.38",style=bold];
"HFM1_S" -> "MYH7_M" [color="0 0 0.3008778346744697", label="2.39",style=bold];
"CPNE6_S" -> "DUSP27_S" [color="0 0 0.30021945866861743", label="2.40",style=bold];
"ATP13A4_S" -> "CDH19_E" [color="0 0 0.29956108266276527", label="2.41",style=bold];
"CPNE6_S" -> "FSTL5_S" [color="0 0 0.298902706656913", label="2.41",style=bold];
"LPPR1_S" -> "MYH6_S" [color="0 0 0.2982443306510607", label="2.42",style=bold];
"NPFFR2_S" -> "MYH6_S" [color="0 0 0.29758595464520854", label="2.42",style=bold];
"HFM1_S" -> "MYH6_M" [color="0 0 0.29692757863935626", label="2.43",style=bold];
"NELL1_S" -> "LRAT_E" [color="0 0 0.2962692026335041", label="2.43",style=bold];
"CNTNAP5_S" -> "FREM2_M" [color="0 0 0.2956108266276518", label="2.43",style=bold];
"LCT_S" -> "LRAT_S" [color="0 0 0.29495245062179964", label="2.43",style=bold];
"DUSP27_S" -> "MUC16_M" [color="0 0 0.29429407461594737", label="2.43",style=bold];
"ERN2_S" -> "MAB21L2_S" [color="0 0 0.2936356986100951", label="2.44",style=bold];
"FSTL5_S" -> "MYBPC2_E" [color="0 0 0.2929773226042429", label="2.45",style=bold];
"ADAMTS20_S" -> "MAB21L2_M" [color="0 0 0.29231894659839064", label="2.45",style=bold];
"TTN_S" -> "LRP1B_S" [color="0 0 0.29166057059253847", label="2.46",style=bold];
"GALNTL6_S" -> "UGT2A1_M" [color="0 0 0.2910021945866862", label="2.46",style=bold];
"PCDHA10_S" -> "DNAH11_S" [color="0 0 0.290343818580834", label="2.47",style=bold];
"LPA_S" -> "CNTN5_M" [color="0 0 0.28968544257498174", label="2.49",style=bold];
"CNTNAP5_S" -> "COL11A1_S" [color="0 0 0.28902706656912946", label="2.49",style=bold];
"NELL1_S" -> "CSMD1_E" [color="0 0 0.2883686905632773", label="2.50",style=bold];
"LRRTM4_S" -> "ABCA13_E" [color="0 0 0.287710314557425", label="2.51",style=bold];
"PCDHA10_S" -> "PCDHA10_E" [color="0 0 0.28705193855157285", label="2.51",style=bold];
"MYBPC2_S" -> "DNAH11_E" [color="0 0 0.28639356254572057", label="2.51",style=bold];
"ADAMTS20_S" -> "NTSR2_M" [color="0 0 0.2857351865398684", label="2.52",style=bold];
"CR2_S" -> "FREM2_E" [color="0 0 0.2850768105340161", label="2.52",style=bold];
"LCE2C_S" -> "FLG_S" [color="0 0 0.28441843452816384", label="2.54",style=bold];
"PCDHA12_S" -> "DNAH11_M" [color="0 0 0.2837600585223117", label="2.54",style=bold];
"GABRB3_S" -> "ZIC1_M" [color="0 0 0.2831016825164594", label="2.55",style=bold];
"MRGPRX4_S" -> "MAB21L2_M" [color="0 0 0.28244330651060723", label="2.56",style=bold];
"TUBA3C_S" -> "HFM1_S" [color="0 0 0.28178493050475495", label="2.57",style=bold];
"KRT1_S" -> "GABRB3_S" [color="0 0 0.2811265544989028", label="2.58",style=bold];
"LPA_S" -> "UGT2A1_E" [color="0 0 0.2804681784930505", label="2.58",style=bold];
"MYH13_S" -> "NELL1_M" [color="0 0 0.27980980248719833", label="2.58",style=bold];
"GABRB3_S" -> "RYR2_S" [color="0 0 0.27915142648134605", label="2.60",style=bold];
"FSTL5_S" -> "KRT24_E" [color="0 0 0.2784930504754938", label="2.60",style=bold];
"PCDHA12_S" -> "CSMD1_S" [color="0 0 0.2778346744696416", label="2.60",style=bold];
"UGT2A1_S" -> "RBP3_S" [color="0 0 0.27717629846378933", label="2.61",style=bold];
"DUSP27_S" -> "RYR2_S" [color="0 0 0.27651792245793716", label="2.62",style=bold];
"CNTN5_S" -> "NEB_M" [color="0 0 0.2758595464520849", label="2.63",style=bold];
"CNTN5_S" -> "CCDC67_M" [color="0 0 0.2752011704462327", label="2.63",style=bold];
"DNAH11_S" -> "MYH7_S" [color="0 0 0.27454279444038043", label="2.63",style=bold];
"RP1_S" -> "COL11A1_E" [color="0 0 0.27388441843452815", label="2.63",style=bold];
"CDH19_S" -> "LRRTM4_M" [color="0 0 0.273226042428676", label="2.64",style=bold];
"LINGO2_S" -> "ZIC1_M" [color="0 0 0.2725676664228237", label="2.64",style=bold];
"CSMD3_M" -> "RP1_M" [color="0 0 0.27190929041697154", label="2.65",style=bold];
"CCDC67_S" -> "PTPRT_S" [color="0 0 0.27125091441111926", label="2.65",style=bold];
"PCLO_S" -> "CDKN2A_E" [color="0 0 0.2705925384052671", label="2.65",style=bold];
"MYBPC2_S" -> "CR2_M" [color="0 0 0.2699341623994148", label="2.65",style=bold];
"MRGPRX4_S" -> "CR2_E" [color="0 0 0.26927578639356253", label="2.66",style=bold];
"BEST2_S" -> "COL11A1_S" [color="0 0 0.26861741038771036", label="2.68",style=bold];
"PCDHA10_S" -> "NEB_S" [color="0 0 0.2679590343818581", label="2.69",style=bold];
"MYH6_M" -> "BEST2_M" [color="0 0 0.2673006583760059", label="2.69",style=bold];
"LPPR1_S" -> "FREM2_M" [color="0 0 0.26664228237015364", label="2.71",style=bold];
"BEST2_S" -> "DNAH11_S" [color="0 0 0.26598390636430147", label="2.71",style=bold];
"MYH13_S" -> "CSMD1_E" [color="0 0 0.2653255303584492", label="2.71",style=bold];
"MC5R_S" -> "CCDC67_E" [color="0 0 0.2646671543525969", label="2.72",style=bold];
"HFM1_S" -> "MAB21L2_M" [color="0 0 0.26400877834674474", label="2.73",style=bold];
"ERBB4_M" -> "MUC16_S" [color="0 0 0.26335040234089246", label="2.75",style=bold];
"LRP1B_E" -> "LPA_E" [color="0 0 0.2626920263350403", label="2.75",style=bold];
"FSTL5_S" -> "CSMD3_S" [color="0 0 0.262033650329188", label="2.76",style=bold];
"GALNTL6_S" -> "CSMD3_S" [color="0 0 0.26137527432333585", label="2.76",style=bold];
"NTSR2_S" -> "MUC16_S" [color="0 0 0.26071689831748357", label="2.76",style=bold];
"LPPR1_S" -> "PCDHA10_E" [color="0 0 0.2600585223116314", label="2.79",style=bold];
"MYBPC2_S" -> "MYH6_E" [color="0 0 0.2594001463057791", label="2.80",style=bold];
"NEB_S" -> "LPPR1_E" [color="0 0 0.25874177029992684", label="2.82",style=bold];
"COL11A1_S" -> "LRP1B_E" [color="0 0 0.2580833942940747", label="2.82",style=bold];
"ERN2_S" -> "FLG_S" [color="0 0 0.2574250182882224", label="2.82",style=bold];
"LCT_S" -> "Survival" [color="0 0 0.2567666422823702", label="2.84",style=bold];
"PCDH10_S" -> "PCDH10_M" [color="0 0 0.25610826627651795", label="2.84",style=bold];
"PKHD1L1_S" -> "TTN_S" [color="0 0 0.2554498902706658", label="2.85",style=bold];
"CSMD3_M" -> "CNTNAP5_E" [color="0 0 0.2547915142648135", label="2.86",style=bold];
"HFM1_S" -> "PCDH10_E" [color="0 0 0.2541331382589612", label="2.86",style=bold];
"DNAH11_M" -> "LCE2C_M" [color="0 0 0.25347476225310905", label="2.89",style=bold];
"MYO3A_S" -> "RBP3_M" [color="0 0 0.2528163862472568", label="2.90",style=bold];
"LPA_S" -> "MYO3A_E" [color="0 0 0.2521580102414046", label="2.93",style=bold];
"MRGPRX4_S" -> "RYR2_S" [color="0 0 0.2514996342355523", label="2.94",style=bold];
"LRAT_S" -> "RP1_S" [color="0 0 0.25084125822970016", label="2.95",style=bold];
"PKHD1L1_S" -> "NTSR2_E" [color="0 0 0.2501828822238479", label="2.96",style=bold];
"MYO3A_S" -> "FLG_E" [color="0 0 0.2495245062179956", label="2.96",style=bold];
"GALNTL6_S" -> "LRP1B_M" [color="0 0 0.24886613021214343", label="2.97",style=bold];
"DSCAM_S" -> "COL11A1_E" [color="0 0 0.24820775420629115", label="2.98",style=bold];
"ERN2_S" -> "PKHD1L1_S" [color="0 0 0.24754937820043899", label="2.99",style=bold];
"ATP13A4_S" -> "MYBPC2_E" [color="0 0 0.2468910021945867", label="3.01",style=bold];
"DUSP27_M" -> "PCLO_S" [color="0 0 0.24623262618873454", label="3.01",style=bold];
"LCE2C_S" -> "NELL1_S" [color="0 0 0.24557425018288226", label="3.01",style=bold];
"NTSR2_S" -> "NELL1_S" [color="0 0 0.24491587417702998", label="3.01",style=bold];
"ADAMTS20_S" -> "DNAH11_M" [color="0 0 0.2442574981711778", label="3.02",style=bold];
"LRAT_S" -> "PTPRT_M" [color="0 0 0.24359912216532553", label="3.02",style=bold];
"LCE2C_S" -> "MYH7_S" [color="0 0 0.24294074615947336", label="3.02",style=bold];
"GALNTL6_S" -> "RP1_S" [color="0 0 0.24228237015362108", label="3.03",style=bold];
"CSMD3_M" -> "MC5R_M" [color="0 0 0.24162399414776892", label="3.04",style=bold];
"FREM2_S" -> "PCDHA10_S" [color="0 0 0.24096561814191664", label="3.04",style=bold];
"KRT1_S" -> "PCLO_S" [color="0 0 0.24030724213606436", label="3.04",style=bold];
"CNTNAP5_S" -> "MUC16_M" [color="0 0 0.2396488661302122", label="3.07",style=bold];
"RYR2_S" -> "BEST2_E" [color="0 0 0.2389904901243599", label="3.08",style=bold];
"DUSP27_S" -> "CSMD1_S" [color="0 0 0.23833211411850774", label="3.09",style=bold];
"PTPRT_S" -> "NPFFR2_M" [color="0 0 0.23767373811265546", label="3.09",style=bold];
"LPA_S" -> "LRAT_E" [color="0 0 0.2370153621068033", label="3.09",style=bold];
"LCE2C_S" -> "LRAT_S" [color="0 0 0.23635698610095102", label="3.11",style=bold];
"DNAH11_S" -> "GABRB3_E" [color="0 0 0.23569861009509885", label="3.11",style=bold];
"ABCA13_S" -> "PCLO_S" [color="0 0 0.23504023408924657", label="3.11",style=bold];
"LINGO2_S" -> "ATP13A4_E" [color="0 0 0.2343818580833943", label="3.13",style=bold];
"CPNE6_S" -> "KRT24_S" [color="0 0 0.23372348207754212", label="3.14",style=bold];
"NTSR2_S" -> "KRT24_S" [color="0 0 0.23306510607168984", label="3.14",style=bold];
"PCDH10_S" -> "CR2_M" [color="0 0 0.23240673006583767", label="3.17",style=bold];
"FLG_E" -> "RBP3_E" [color="0 0 0.2317483540599854", label="3.17",style=bold];
"CDKN2A_E" -> "LCE2C_E" [color="0 0 0.23108997805413323", label="3.19",style=bold];
"PCDHA10_S" -> "MYBPC2_E" [color="0 0 0.23043160204828095", label="3.19",style=bold];
"ATP13A4_S" -> "ATP13A4_M" [color="0 0 0.22977322604242867", label="3.20",style=bold];
"DSCAM_S" -> "DNAH11_S" [color="0 0 0.2291148500365765", label="3.21",style=bold];
"MYBPC2_S" -> "TTN_E" [color="0 0 0.22845647403072422", label="3.22",style=bold];
"ATP13A4_S" -> "ZIC1_M" [color="0 0 0.22779809802487205", label="3.23",style=bold];
"GABRB3_M" -> "RYR2_M" [color="0 0 0.22713972201901977", label="3.24",style=bold];
"MC5R_S" -> "ADAMTS20_M" [color="0 0 0.2264813460131676", label="3.25",style=bold];
"LCT_S" -> "LRRTM4_E" [color="0 0 0.22582297000731533", label="3.25",style=bold];
"HFM1_S" -> "KRT24_E" [color="0 0 0.22516459400146305", label="3.30",style=bold];
"MAB21L2_S" -> "RP1_S" [color="0 0 0.22450621799561088", label="3.33",style=bold];
"NELL1_S" -> "ZIC1_M" [color="0 0 0.2238478419897586", label="3.33",style=bold];
"LCT_S" -> "DNAH11_M" [color="0 0 0.22318946598390643", label="3.34",style=bold];
"MYH6_E" -> "MYH13_E" [color="0 0 0.22253108997805415", label="3.34",style=bold];
"RP1_M" -> "LCT_M" [color="0 0 0.22187271397220198", label="3.36",style=bold];
"ABCA13_S" -> "CR2_S" [color="0 0 0.2212143379663497", label="3.37",style=bold];
"CNTNAP5_S" -> "LPA_S" [color="0 0 0.22055596196049743", label="3.38",style=bold];
"FREM2_S" -> "MYH13_S" [color="0 0 0.21989758595464526", label="3.40",style=bold];
"NELL1_S" -> "MYH7_E" [color="0 0 0.21923920994879298", label="3.44",style=bold];
"NPFFR2_S" -> "PCDHA12_E" [color="0 0 0.2185808339429408", label="3.45",style=bold];
"CDH19_S" -> "CNTN5_S" [color="0 0 0.21792245793708853", label="3.45",style=bold];
"CSMD1_S" -> "LCE2C_E" [color="0 0 0.21726408193123636", label="3.45",style=bold];
"ZIC1_S" -> "PKHD1L1_S" [color="0 0 0.21660570592538408", label="3.46",style=bold];
"FSTL5_S" -> "CPNE6_E" [color="0 0 0.2159473299195318", label="3.47",style=bold];
"ATP13A4_M" -> "LINGO2_M" [color="0 0 0.21528895391367964", label="3.48",style=bold];
"BEST2_S" -> "FSTL5_S" [color="0 0 0.21463057790782736", label="3.49",style=bold];
"CSMD3_M" -> "RYR2_M" [color="0 0 0.2139722019019752", label="3.50",style=bold];
"MYH6_S" -> "FSTL5_E" [color="0 0 0.2133138258961229", label="3.50",style=bold];
"LRAT_E" -> "MRGPRX4_E" [color="0 0 0.21265544989027074", label="3.50",style=bold];
"PTPRT_S" -> "CDKN2A_E" [color="0 0 0.21199707388441846", label="3.50",style=bold];
"ATP13A4_S" -> "TDRD5_M" [color="0 0 0.2113386978785663", label="3.52",style=bold];
"CR2_E" -> "PTPRT_E" [color="0 0 0.21068032187271402", label="3.53",style=bold];
"ZIC1_S" -> "LCT_E" [color="0 0 0.21002194586686174", label="3.55",style=bold];
"MYH13_S" -> "MYBPC2_E" [color="0 0 0.20936356986100957", label="3.56",style=bold];
"BEST2_S" -> "MRGPRX4_S" [color="0 0 0.2087051938551573", label="3.56",style=bold];
"LPA_S" -> "CR2_E" [color="0 0 0.20804681784930512", label="3.56",style=bold];
"DNAH11_S" -> "GALNTL6_E" [color="0 0 0.20738844184345284", label="3.57",style=bold];
"ERBB4_S" -> "KRT24_M" [color="0 0 0.20673006583760067", label="3.59",style=bold];
"CNTNAP5_S" -> "PTPRT_S" [color="0 0 0.2060716898317484", label="3.63",style=bold];
"ERBB4_S" -> "NPFFR2_M" [color="0 0 0.20541331382589612", label="3.64",style=bold];
"DUSP27_E" -> "MYH13_E" [color="0 0 0.20475493782004395", label="3.69",style=bold];
"ABCA13_S" -> "KRT1_M" [color="0 0 0.20409656181419167", label="3.70",style=bold];
"FLG_E" -> "MRGPRX4_E" [color="0 0 0.2034381858083395", label="3.71",style=bold];
"FREM2_S" -> "CCDC67_M" [color="0 0 0.20277980980248722", label="3.74",style=bold];
"PCDH10_S" -> "GALNTL6_M" [color="0 0 0.20212143379663505", label="3.74",style=bold];
"CDKN2A_E" -> "KRT1_E" [color="0 0 0.20146305779078277", label="3.75",style=bold];
"LRRTM4_S" -> "UGT2A1_E" [color="0 0 0.2008046817849305", label="3.79",style=bold];
"MYH7_M" -> "NEB_M" [color="0 0 0.20014630577907833", label="3.81",style=bold];
"MYBPC2_S" -> "FLG_S" [color="0 0 0.19948792977322605", label="3.82",style=bold];
"RYR2_E" -> "PCDHA10_E" [color="0 0 0.19882955376737388", label="3.84",style=bold];
"CSMD3_M" -> "TDRD5_E" [color="0 0 0.1981711777615216", label="3.84",style=bold];
"PCDH10_S" -> "ERN2_M" [color="0 0 0.19751280175566943", label="3.86",style=bold];
"ADAMTS20_S" -> "DSCAM_E" [color="0 0 0.19685442574981715", label="3.87",style=bold];
"CR2_M" -> "LPPR1_E" [color="0 0 0.19619604974396487", label="3.88",style=bold];
"ZIC1_S" -> "CSMD3_S" [color="0 0 0.1955376737381127", label="3.89",style=bold];
"NPFFR2_S" -> "RYR2_E" [color="0 0 0.19487929773226043", label="3.89",style=bold];
"CR2_E" -> "CSMD3_S" [color="0 0 0.19422092172640826", label="3.92",style=bold];
"PCDH10_S" -> "MUC4_E" [color="0 0 0.19356254572055598", label="3.96",style=bold];
"LRRTM4_S" -> "PCLO_M" [color="0 0 0.1929041697147038", label="3.97",style=bold];
"PKHD1L1_S" -> "MUC4_M" [color="0 0 0.19224579370885153", label="3.99",style=bold];
"MRGPRX4_M" -> "ERN2_E" [color="0 0 0.19158741770299925", label="4.02",style=bold];
"MUC4_S" -> "KRT1_M" [color="0 0 0.19092904169714708", label="4.02",style=bold];
"CSMD1_S" -> "PCDH10_E" [color="0 0 0.1902706656912948", label="4.02",style=bold];
"TDRD5_S" -> "GALNTL6_S" [color="0 0 0.18961228968544264", label="4.03",style=bold];
"ATP13A4_S" -> "MYH6_E" [color="0 0 0.18895391367959036", label="4.06",style=bold];
"GABRB3_M" -> "FSTL5_M" [color="0 0 0.1882955376737382", label="4.07",style=bold];
"GABRB3_S" -> "LRRTM4_S" [color="0 0 0.1876371616678859", label="4.08",style=bold];
"ERBB4_S" -> "NEB_E" [color="0 0 0.18697878566203374", label="4.11",style=bold];
"RP1_S" -> "CDKN2A_S" [color="0 0 0.18632040965618146", label="4.11",style=bold];
"ATP13A4_S" -> "DNAH11_M" [color="0 0 0.18566203365032918", label="4.13",style=bold];
"MYBPC2_M" -> "MC5R_M" [color="0 0 0.18500365764447702", label="4.16",style=bold];
"LCE2C_E" -> "ATP13A4_E" [color="0 0 0.18434528163862474", label="4.20",style=bold];
"RBP3_M" -> "UGT2A1_M" [color="0 0 0.18368690563277257", label="4.22",style=bold];
"LRP1B_M" -> "CNTNAP5_M" [color="0 0 0.1830285296269203", label="4.25",style=bold];
"PCDHA12_S" -> "CSMD1_M" [color="0 0 0.18237015362106812", label="4.25",style=bold];
"FSTL5_S" -> "LINGO2_E" [color="0 0 0.18171177761521584", label="4.29",style=bold];
"NTSR2_E" -> "PKHD1L1_E" [color="0 0 0.18105340160936356", label="4.31",style=bold];
"RBP3_S" -> "RP1_S" [color="0 0 0.1803950256035114", label="4.32",style=bold];
"TTN_M" -> "KRT24_E" [color="0 0 0.17973664959765911", label="4.33",style=bold];
"ERBB4_M" -> "PCLO_M" [color="0 0 0.17907827359180695", label="4.36",style=bold];
"ERBB4_E" -> "CDH19_E" [color="0 0 0.17841989758595467", label="4.36",style=bold];
"ERN2_S" -> "ATP13A4_S" [color="0 0 0.1777615215801025", label="4.37",style=bold];
"TTN_S" -> "MAB21L2_E" [color="0 0 0.17710314557425022", label="4.37",style=bold];
"RYR2_M" -> "ABCA13_M" [color="0 0 0.17644476956839794", label="4.38",style=bold];
"LPA_M" -> "FLG_M" [color="0 0 0.17578639356254577", label="4.39",style=bold];
"CNTNAP5_M" -> "ADAMTS20_E" [color="0 0 0.1751280175566935", label="4.40",style=bold];
"PKHD1L1_S" -> "DSCAM_S" [color="0 0 0.17446964155084133", label="4.43",style=bold];
"DNAH11_S" -> "CNTN5_E" [color="0 0 0.17381126554498905", label="4.43",style=bold];
"ABCA13_S" -> "MUC16_S" [color="0 0 0.17315288953913688", label="4.49",style=bold];
"MUC4_S" -> "KRT24_M" [color="0 0 0.1724945135332846", label="4.50",style=bold];
"LINGO2_S" -> "LCT_E" [color="0 0 0.17183613752743232", label="4.52",style=bold];
"NPFFR2_S" -> "CSMD3_S" [color="0 0 0.17117776152158015", label="4.53",style=bold];
"RBP3_M" -> "RP1_M" [color="0 0 0.17051938551572787", label="4.60",style=bold];
"ATP13A4_S" -> "NEB_S" [color="0 0 0.1698610095098757", label="4.63",style=bold];
"CNTNAP5_S" -> "CSMD3_M" [color="0 0 0.16920263350402343", label="4.65",style=bold];
"PTPRT_M" -> "NPFFR2_E" [color="0 0 0.16854425749817126", label="4.66",style=bold];
"MYH13_S" -> "LRRTM4_S" [color="0 0 0.16788588149231898", label="4.71",style=bold];
"MYH7_M" -> "LPA_M" [color="0 0 0.1672275054864668", label="4.74",style=bold];
"TDRD5_E" -> "LINGO2_M" [color="0 0 0.16656912948061453", label="4.79",style=bold];
"DNAH11_S" -> "LRP1B_M" [color="0 0 0.16591075347476225", label="4.82",style=bold];
"RYR2_S" -> "CSMD1_E" [color="0 0 0.16525237746891008", label="4.82",style=bold];
"LCE2C_M" -> "GABRB3_E" [color="0 0 0.1645940014630578", label="4.83",style=bold];
"TDRD5_S" -> "DNAH11_S" [color="0 0 0.16393562545720564", label="4.91",style=bold];
"RYR2_M" -> "MUC16_M" [color="0 0 0.16327724945135336", label="4.98",style=bold];
"LRRTM4_M" -> "LCT_M" [color="0 0 0.1626188734455012", label="5.01",style=bold];
"GALNTL6_S" -> "DUSP27_S" [color="0 0 0.1619604974396489", label="5.08",style=bold];
"LCT_E" -> "MUC16_E" [color="0 0 0.16130212143379663", label="5.13",style=bold];
"NTSR2_S" -> "FLG_S" [color="0 0 0.16064374542794446", label="5.15",style=bold];
"KRT1_E" -> "NTSR2_M" [color="0 0 0.15998536942209218", label="5.15",style=bold];
"MAB21L2_M" -> "BEST2_E" [color="0 0 0.15932699341624001", label="5.21",style=bold];
"LCT_S" -> "DSCAM_S" [color="0 0 0.15866861741038774", label="5.27",style=bold];
"CSMD3_M" -> "CCDC67_E" [color="0 0 0.15801024140453557", label="5.29",style=bold];
"FREM2_M" -> "HFM1_M" [color="0 0 0.1573518653986833", label="5.29",style=bold];
"MYH6_E" -> "ERBB4_E" [color="0 0 0.156693489392831", label="5.32",style=bold];
"CSMD3_S" -> "CPNE6_E" [color="0 0 0.15603511338697884", label="5.33",style=bold];
"MUC4_S" -> "Survival" [color="0 0 0.15537673738112656", label="5.42",style=bold];
"COL11A1_S" -> "CDKN2A_M" [color="0 0 0.1547183613752744", label="5.42",style=bold];
"NPFFR2_S" -> "COL11A1_S" [color="0 0 0.15405998536942211", label="5.44",style=bold];
"LRRTM4_S" -> "ERBB4_S" [color="0 0 0.15340160936356995", label="5.44",style=bold];
"COL11A1_M" -> "PCDH10_M" [color="0 0 0.15274323335771767", label="5.45",style=bold];
"ATP13A4_S" -> "CPNE6_E" [color="0 0 0.1520848573518654", label="5.48",style=bold];
"UGT2A1_S" -> "MYH13_S" [color="0 0 0.15142648134601322", label="5.48",style=bold];
"ERBB4_E" -> "GALNTL6_E" [color="0 0 0.15076810534016094", label="5.50",style=bold];
"PKHD1L1_E" -> "PCDHA12_E" [color="0 0 0.15010972933430877", label="5.55",style=bold];
"GALNTL6_M" -> "DUSP27_M" [color="0 0 0.1494513533284565", label="5.55",style=bold];
"MUC16_S" -> "LRP1B_S" [color="0 0 0.14879297732260433", label="5.68",style=bold];
"LRAT_S" -> "HFM1_S" [color="0 0 0.14813460131675205", label="5.68",style=bold];
"PKHD1L1_S" -> "DSCAM_E" [color="0 0 0.14747622531089977", label="5.70",style=bold];
"GALNTL6_S" -> "LRAT_S" [color="0 0 0.1468178493050476", label="5.72",style=bold];
"MC5R_E" -> "PTPRT_E" [color="0 0 0.14615947329919532", label="5.76",style=bold];
"NEB_S" -> "PCDHA10_M" [color="0 0 0.14550109729334315", label="5.77",style=bold];
"NTSR2_M" -> "LCT_E" [color="0 0 0.14484272128749087", label="5.82",style=bold];
"MC5R_S" -> "FLG_S" [color="0 0 0.1441843452816387", label="5.84",style=bold];
"TDRD5_S" -> "FLG_S" [color="0 0 0.14352596927578642", label="5.84",style=bold];
"LCT_S" -> "COL11A1_S" [color="0 0 0.14286759326993426", label="5.84",style=bold];
"CNTN5_S" -> "MRGPRX4_E" [color="0 0 0.14220921726408198", label="5.89",style=bold];
"MYH7_M" -> "LRRTM4_M" [color="0 0 0.1415508412582297", label="5.98",style=bold];
"CNTNAP5_E" -> "CDKN2A_M" [color="0 0 0.14089246525237753", label="6.02",style=bold];
"FLG_M" -> "Survival" [color="0 0 0.14023408924652525", label="6.07",style=bold];
"PKHD1L1_S" -> "LINGO2_E" [color="0 0 0.13957571324067308", label="6.09",style=bold];
"MYH13_S" -> "ABCA13_S" [color="0 0 0.1389173372348208", label="6.10",style=bold];
"RBP3_M" -> "LCE2C_M" [color="0 0 0.13825896122896864", label="6.12",style=bold];
"CSMD3_M" -> "LRAT_M" [color="0 0 0.13760058522311636", label="6.16",style=bold];
"CNTNAP5_E" -> "RYR2_S" [color="0 0 0.13694220921726408", label="6.23",style=bold];
"LCE2C_M" -> "LRRTM4_E" [color="0 0 0.1362838332114119", label="6.25",style=bold];
"CR2_S" -> "RP1_S" [color="0 0 0.13562545720555963", label="6.27",style=bold];
"RP1_S" -> "LCT_E" [color="0 0 0.13496708119970746", label="6.36",style=bold];
"GABRB3_M" -> "LPPR1_M" [color="0 0 0.13430870519385518", label="6.37",style=bold];
"ABCA13_S" -> "PKHD1L1_S" [color="0 0 0.13365032918800301", label="6.43",style=bold];
"TDRD5_E" -> "RBP3_E" [color="0 0 0.13299195318215074", label="6.54",style=bold];
"NEB_E" -> "MYH13_E" [color="0 0 0.13233357717629846", label="6.59",style=bold];
"MUC16_S" -> "TTN_S" [color="0 0 0.1316752011704463", label="6.61",style=bold];
"CSMD1_E" -> "LRP1B_E" [color="0 0 0.131016825164594", label="6.66",style=bold];
"CSMD3_S" -> "HFM1_M" [color="0 0 0.13035844915874184", label="6.69",style=bold];
"ERBB4_M" -> "CR2_M" [color="0 0 0.12970007315288956", label="6.77",style=bold];
"LINGO2_M" -> "GABRB3_E" [color="0 0 0.1290416971470374", label="6.87",style=bold];
"KRT1_M" -> "LCE2C_E" [color="0 0 0.1283833211411851", label="6.91",style=bold];
"NTSR2_M" -> "TDRD5_M" [color="0 0 0.12772494513533283", label="6.97",style=bold];
"LRRTM4_M" -> "TTN_M" [color="0 0 0.12706656912948067", label="6.99",style=bold];
"DSCAM_E" -> "DSCAM_M" [color="0 0 0.1264081931236284", label="7.05",style=bold];
"RBP3_S" -> "COL11A1_S" [color="0 0 0.12574981711777622", label="7.06",style=bold];
"CPNE6_M" -> "MRGPRX4_M" [color="0 0 0.12509144111192394", label="7.06",style=bold];
"GABRB3_M" -> "MYO3A_M" [color="0 0 0.12443306510607177", label="7.08",style=bold];
"NEB_S" -> "HFM1_E" [color="0 0 0.12377468910021949", label="7.13",style=bold];
"CSMD1_M" -> "CDKN2A_M" [color="0 0 0.12311631309436721", label="7.14",style=bold];
"GABRB3_S" -> "TTN_M" [color="0 0 0.12245793708851505", label="7.15",style=bold];
"PCDHA10_M" -> "NELL1_E" [color="0 0 0.12179956108266277", label="7.19",style=bold];
"NELL1_E" -> "LRP1B_E" [color="0 0 0.1211411850768106", label="7.20",style=bold];
"MRGPRX4_M" -> "TUBA3C_M" [color="0 0 0.12048280907095832", label="7.23",style=bold];
"PCDHA12_M" -> "PTPRT_M" [color="0 0 0.11982443306510615", label="7.45",style=bold];
"UGT2A1_E" -> "TDRD5_E" [color="0 0 0.11916605705925387", label="7.46",style=bold];
"CSMD3_M" -> "MYO3A_M" [color="0 0 0.1185076810534017", label="7.53",style=bold];
"GALNTL6_S" -> "GABRB3_S" [color="0 0 0.11784930504754942", label="7.53",style=bold];
"ERBB4_M" -> "NELL1_E" [color="0 0 0.11719092904169714", label="7.59",style=bold];
"CSMD3_M" -> "PCDH10_M" [color="0 0 0.11653255303584498", label="7.62",style=bold];
"DNAH11_S" -> "COL11A1_E" [color="0 0 0.1158741770299927", label="7.76",style=bold];
"MYH6_M" -> "RBP3_M" [color="0 0 0.11521580102414053", label="7.83",style=bold];
"FSTL5_M" -> "PCLO_E" [color="0 0 0.11455742501828825", label="7.83",style=bold];
"CR2_E" -> "COL11A1_E" [color="0 0 0.11389904901243608", label="7.91",style=bold];
"DUSP27_E" -> "UGT2A1_E" [color="0 0 0.1132406730065838", label="7.94",style=bold];
"MYBPC2_M" -> "DNAH11_M" [color="0 0 0.11258229700073152", label="7.97",style=bold];
"CDKN2A_E" -> "CDKN2A_S" [color="0 0 0.11192392099487936", label="7.98",style=bold];
"RP1_S" -> "MUC16_S" [color="0 0 0.11126554498902708", label="8.01",style=bold];
"CSMD3_M" -> "DSCAM_E" [color="0 0 0.11060716898317491", label="8.02",style=bold];
"TTN_M" -> "LCE2C_M" [color="0 0 0.10994879297732263", label="8.02",style=bold];
"MYBPC2_M" -> "ATP13A4_M" [color="0 0 0.10929041697147046", label="8.07",style=bold];
"MYBPC2_E" -> "UGT2A1_E" [color="0 0 0.10863204096561818", label="8.10",style=bold];
"CNTNAP5_M" -> "RYR2_M" [color="0 0 0.1079736649597659", label="8.12",style=bold];
"HFM1_E" -> "NPFFR2_E" [color="0 0 0.10731528895391373", label="8.13",style=bold];
"LPPR1_E" -> "BEST2_E" [color="0 0 0.10665691294806146", label="8.16",style=bold];
"MYH6_E" -> "ZIC1_E" [color="0 0 0.10599853694220929", label="8.22",style=bold];
"CPNE6_E" -> "LRRTM4_E" [color="0 0 0.10534016093635701", label="8.25",style=bold];
"MUC16_S" -> "RP1_E" [color="0 0 0.10468178493050484", label="8.27",style=bold];
"MUC4_M" -> "PCDH10_E" [color="0 0 0.10402340892465256", label="8.42",style=bold];
"ERBB4_E" -> "PCDHA12_E" [color="0 0 0.10336503291880028", label="8.43",style=bold];
"CCDC67_E" -> "MAB21L2_M" [color="0 0 0.10270665691294811", label="8.50",style=bold];
"LRP1B_E" -> "MC5R_E" [color="0 0 0.10204828090709583", label="8.62",style=bold];
"MYBPC2_M" -> "BEST2_M" [color="0 0 0.10138990490124367", label="8.63",style=bold];
"LRRTM4_M" -> "CNTN5_M" [color="0 0 0.10073152889539139", label="8.64",style=bold];
"FLG_E" -> "ADAMTS20_E" [color="0 0 0.10007315288953922", label="8.66",style=bold];
"MYBPC2_E" -> "BEST2_E" [color="0 0 0.09941477688368694", label="8.67",style=bold];
"LRRTM4_M" -> "LRP1B_M" [color="0 0 0.09875640087783466", label="8.73",style=bold];
"PCDHA10_M" -> "LCE2C_E" [color="0 0 0.09809802487198249", label="8.83",style=bold];
"CNTN5_M" -> "FSTL5_M" [color="0 0 0.09743964886613021", label="8.84",style=bold];
"TTN_M" -> "PKHD1L1_M" [color="0 0 0.09678127286027804", label="8.87",style=bold];
"DNAH11_E" -> "TDRD5_E" [color="0 0 0.09612289685442577", label="9.09",style=bold];
"RP1_M" -> "MRGPRX4_M" [color="0 0 0.0954645208485736", label="9.12",style=bold];
"PCDHA10_M" -> "COL11A1_M" [color="0 0 0.09480614484272132", label="9.23",style=bold];
"CSMD1_E" -> "ABCA13_E" [color="0 0 0.09414776883686915", label="9.26",style=bold];
"PKHD1L1_M" -> "ATP13A4_M" [color="0 0 0.09348939283101687", label="9.28",style=bold];
"CNTN5_M" -> "PCLO_M" [color="0 0 0.09283101682516459", label="9.33",style=bold];
"PTPRT_M" -> "LRAT_M" [color="0 0 0.09217264081931242", label="9.34",style=bold];
"CSMD1_M" -> "ABCA13_M" [color="0 0 0.09151426481346014", label="9.52",style=bold];
"LCE2C_E" -> "MUC16_E" [color="0 0 0.09085588880760798", label="9.55",style=bold];
"ZIC1_E" -> "RYR2_E" [color="0 0 0.0901975128017557", label="9.61",style=bold];
"PTPRT_E" -> "FREM2_E" [color="0 0 0.08953913679590353", label="9.67",style=bold];
"LINGO2_E" -> "LRAT_E" [color="0 0 0.08888076079005125", label="9.79",style=bold];
"TTN_S" -> "RP1_E" [color="0 0 0.08822238478419897", label="10.02",style=bold];
"CSMD1_M" -> "ZIC1_E" [color="0 0 0.0875640087783468", label="10.08",style=bold];
"PTPRT_M" -> "KRT1_M" [color="0 0 0.08690563277249452", label="10.16",style=bold];
"LRP1B_M" -> "PKHD1L1_E" [color="0 0 0.08624725676664236", label="10.19",style=bold];
"DNAH11_M" -> "TDRD5_M" [color="0 0 0.08558888076079008", label="10.21",style=bold];
"MC5R_M" -> "DNAH11_E" [color="0 0 0.08493050475493791", label="10.24",style=bold];
"LCE2C_M" -> "BEST2_M" [color="0 0 0.08427212874908563", label="10.27",style=bold];
"PKHD1L1_E" -> "PTPRT_E" [color="0 0 0.08361375274323335", label="10.32",style=bold];
"NPFFR2_M" -> "COL11A1_M" [color="0 0 0.08295537673738118", label="10.35",style=bold];
"PCDH10_M" -> "ZIC1_M" [color="0 0 0.0822970007315289", label="10.41",style=bold];
"DSCAM_M" -> "MYBPC2_M" [color="0 0 0.08163862472567673", label="10.50",style=bold];
"LPPR1_M" -> "TTN_S" [color="0 0 0.08098024871982445", label="10.68",style=bold];
"GALNTL6_M" -> "RP1_E" [color="0 0 0.08032187271397229", label="10.74",style=bold];
"NPFFR2_M" -> "ADAMTS20_M" [color="0 0 0.07966349670812001", label="10.78",style=bold];
"MRGPRX4_M" -> "ATP13A4_E" [color="0 0 0.07900512070226773", label="10.79",style=bold];
"PKHD1L1_M" -> "LRP1B_M" [color="0 0 0.07834674469641556", label="10.83",style=bold];
"LPA_M" -> "LPA_E" [color="0 0 0.07768836869056328", label="10.83",style=bold];
"LCT_E" -> "DNAH11_E" [color="0 0 0.07702999268471111", label="10.90",style=bold];
"ZIC1_M" -> "DSCAM_E" [color="0 0 0.07637161667885883", label="10.97",style=bold];
"TDRD5_E" -> "MUC4_E" [color="0 0 0.07571324067300667", label="11.26",style=bold];
"MYH13_M" -> "UGT2A1_M" [color="0 0 0.07505486466715439", label="11.39",style=bold];
"FREM2_E" -> "ERBB4_E" [color="0 0 0.07439648866130222", label="11.45",style=bold];
"CR2_M" -> "HFM1_M" [color="0 0 0.07373811265544994", label="11.48",style=bold];
"UGT2A1_S" -> "ERN2_S" [color="0 0 0.07307973664959766", label="11.63",style=bold];
"COL11A1_M" -> "MYH6_E" [color="0 0 0.07242136064374549", label="11.66",style=bold];
"MYBPC2_M" -> "MYH13_M" [color="0 0 0.07176298463789321", label="11.69",style=bold];
"LRAT_E" -> "LRAT_M" [color="0 0 0.07110460863204104", label="11.79",style=bold];
"CSMD1_E" -> "PKHD1L1_E" [color="0 0 0.07044623262618877", label="11.83",style=bold];
"PCDH10_E" -> "MAB21L2_E" [color="0 0 0.0697878566203366", label="11.86",style=bold];
"NTSR2_M" -> "MYO3A_M" [color="0 0 0.06912948061448432", label="11.87",style=bold];
"CPNE6_M" -> "RBP3_M" [color="0 0 0.06847110460863204", label="11.88",style=bold];
"CNTNAP5_E" -> "PCDH10_E" [color="0 0 0.06781272860277987", label="11.89",style=bold];
"GABRB3_M" -> "KRT1_M" [color="0 0 0.06715435259692759", label="11.96",style=bold];
"CSMD1_M" -> "PCLO_M" [color="0 0 0.06649597659107542", label="12.14",style=bold];
"RBP3_M" -> "RP1_E" [color="0 0 0.06583760058522314", label="12.18",style=bold];
"TDRD5_E" -> "CSMD3_E" [color="0 0 0.06517922457937098", label="12.25",style=bold];
"CNTN5_E" -> "LRP1B_E" [color="0 0 0.0645208485735187", label="12.49",style=bold];
"CDKN2A_M" -> "CDKN2A_E" [color="0 0 0.06386247256766642", label="12.65",style=bold];
"MUC4_E" -> "MUC16_E" [color="0 0 0.06320409656181425", label="12.65",style=bold];
"TTN_E" -> "MAB21L2_E" [color="0 0 0.06254572055596197", label="12.67",style=bold];
"PCDHA10_M" -> "ERBB4_M" [color="0 0 0.0618873445501098", label="12.74",style=bold];
"MUC4_E" -> "KRT24_E" [color="0 0 0.06122896854425752", label="12.78",style=bold];
"MYH6_E" -> "PCDHA10_E" [color="0 0 0.060570592538405355", label="12.87",style=bold];
"PKHD1L1_E" -> "CDH19_E" [color="0 0 0.059912216532553075", label="12.92",style=bold];
"CNTNAP5_E" -> "GABRB3_E" [color="0 0 0.059253840526700796", label="12.97",style=bold];
"ATP13A4_M" -> "CPNE6_E" [color="0 0 0.05859546452084863", label="13.10",style=bold];
"DSCAM_M" -> "KRT24_M" [color="0 0 0.05793708851499635", label="13.39",style=bold];
"NPFFR2_M" -> "ZIC1_M" [color="0 0 0.05727871250914418", label="13.64",style=bold];
"MUC4_E" -> "ABCA13_M" [color="0 0 0.0566203365032919", label="13.74",style=bold];
"DSCAM_E" -> "CSMD3_E" [color="0 0 0.05596196049743973", label="13.77",style=bold];
"NELL1_M" -> "PTPRT_M" [color="0 0 0.055303584491587454", label="13.87",style=bold];
"NPFFR2_M" -> "CDKN2A_E" [color="0 0 0.054645208485735175", label="13.94",style=bold];
"ERBB4_E" -> "HFM1_E" [color="0 0 0.05398683247988301", label="13.99",style=bold];
"DSCAM_M" -> "PKHD1L1_M" [color="0 0 0.05332845647403073", label="14.01",style=bold];
"ZIC1_M" -> "ZIC1_E" [color="0 0 0.05267008046817856", label="14.08",style=bold];
"MUC4_E" -> "ERN2_E" [color="0 0 0.05201170446232628", label="14.49",style=bold];
"TDRD5_E" -> "TTN_S" [color="0 0 0.05135332845647411", label="14.66",style=bold];
"ATP13A4_M" -> "FLG_E" [color="0 0 0.05069495245062183", label="14.69",style=bold];
"CCDC67_E" -> "HFM1_E" [color="0 0 0.050036576444769665", label="14.92",style=bold];
"LPA_E" -> "LRAT_E" [color="0 0 0.049378200438917386", label="14.95",style=bold];
"CNTN5_M" -> "ADAMTS20_E" [color="0 0 0.048719824433065106", label="14.97",style=bold];
"MYH6_E" -> "GALNTL6_E" [color="0 0 0.04806144842721294", label="15.17",style=bold];
"MUC4_E" -> "ATP13A4_E" [color="0 0 0.04740307242136066", label="15.21",style=bold];
"PCLO_M" -> "PCLO_E" [color="0 0 0.04674469641550849", label="15.22",style=bold];
"KRT1_M" -> "MYH7_M" [color="0 0 0.04608632040965621", label="15.46",style=bold];
"PKHD1L1_M" -> "DNAH11_M" [color="0 0 0.045427944403804044", label="15.60",style=bold];
"KRT1_M" -> "LINGO2_M" [color="0 0 0.044769568397951764", label="15.73",style=bold];
"RYR2_E" -> "NELL1_E" [color="0 0 0.044111192392099485", label="15.77",style=bold];
"KRT1_E" -> "DNAH11_E" [color="0 0 0.04345281638624732", label="15.80",style=bold];
"LRRTM4_M" -> "MUC16_M" [color="0 0 0.04279444038039504", label="15.88",style=bold];
"CNTNAP5_E" -> "RYR2_E" [color="0 0 0.04213606437454287", label="15.91",style=bold];
"MYH13_M" -> "FLG_M" [color="0 0 0.04147768836869059", label="15.94",style=bold];
"TDRD5_M" -> "CR2_M" [color="0 0 0.04081931236283842", label="16.05",style=bold];
"TDRD5_E" -> "MYO3A_E" [color="0 0 0.04016093635698614", label="16.06",style=bold];
"NEB_E" -> "FREM2_E" [color="0 0 0.039502560351133864", label="16.29",style=bold];
"DNAH11_E" -> "MYH6_E" [color="0 0 0.038844184345281696", label="16.30",style=bold];
"MUC4_E" -> "ABCA13_E" [color="0 0 0.03818580833942942", label="16.39",style=bold];
"PKHD1L1_M" -> "LPPR1_M" [color="0 0 0.03752743233357725", label="16.44",style=bold];
"CPNE6_M" -> "CCDC67_M" [color="0 0 0.03686905632772497", label="16.92",style=bold];
"ERN2_E" -> "CCDC67_E" [color="0 0 0.0362106803218728", label="17.07",style=bold];
"ADAMTS20_M" -> "CDKN2A_M" [color="0 0 0.03555230431602052", label="17.25",style=bold];
"NEB_M" -> "LPA_M" [color="0 0 0.03489392831016824", label="17.62",style=bold];
"RBP3_M" -> "KRT24_E" [color="0 0 0.034235552304316075", label="17.69",style=bold];
"GABRB3_M" -> "MYH7_M" [color="0 0 0.033577176298463796", label="17.71",style=bold];
"MAB21L2_M" -> "COL11A1_E" [color="0 0 0.03291880029261163", label="17.86",style=bold];
"MYH7_M" -> "CPNE6_M" [color="0 0 0.03226042428675935", label="17.87",style=bold];
"GABRB3_M" -> "CNTNAP5_M" [color="0 0 0.03160204828090718", label="18.14",style=bold];
"ADAMTS20_M" -> "GALNTL6_M" [color="0 0 0.0309436722750549", label="18.21",style=bold];
"PCDHA12_E" -> "PCDHA10_E" [color="0 0 0.030285296269202622", label="18.26",style=bold];
"GALNTL6_M" -> "TUBA3C_M" [color="0 0 0.029626920263350454", label="18.26",style=bold];
"ZIC1_M" -> "ADAMTS20_M" [color="0 0 0.028968544257498174", label="18.31",style=bold];
"PCDHA10_M" -> "MYO3A_E" [color="0 0 0.028310168251646006", label="18.37",style=bold];
"LRP1B_E" -> "LPPR1_E" [color="0 0 0.027651792245793727", label="18.47",style=bold];
"LRP1B_E" -> "CCDC67_E" [color="0 0 0.02699341623994156", label="18.60",style=bold];
"PCDH10_M" -> "FSTL5_M" [color="0 0 0.02633504023408928", label="18.99",style=bold];
"MYH7_M" -> "TTN_M" [color="0 0 0.02567666422823711", label="19.22",style=bold];
"CPNE6_M" -> "KRT24_M" [color="0 0 0.025018288222384832", label="19.42",style=bold];
"LPPR1_M" -> "FREM2_M" [color="0 0 0.024359912216532553", label="19.45",style=bold];
"PCLO_E" -> "MUC4_M" [color="0 0 0.023701536210680385", label="19.60",style=bold];
"ADAMTS20_M" -> "ERN2_M" [color="0 0 0.023043160204828106", label="19.60",style=bold];
"ERBB4_M" -> "FREM2_M" [color="0 0 0.022384784198975938", label="20.06",style=bold];
"MYH13_E" -> "MAB21L2_M" [color="0 0 0.02172640819312366", label="20.57",style=bold];
"CR2_M" -> "CNTN5_M" [color="0 0 0.02106803218727149", label="20.82",style=bold];
"CSMD1_M" -> "LRRTM4_M" [color="0 0 0.02040965618141921", label="21.19",style=bold];
"MYH6_M" -> "MYBPC2_M" [color="0 0 0.019751280175566932", label="21.45",style=bold];
"LCE2C_E" -> "FLG_E" [color="0 0 0.019092904169714764", label="21.81",style=bold];
"CNTNAP5_M" -> "MYH13_M" [color="0 0 0.018434528163862485", label="21.90",style=bold];
"ABCA13_E" -> "PCLO_E" [color="0 0 0.017776152158010317", label="22.62",style=bold];
"PCDHA10_M" -> "ERN2_M" [color="0 0 0.017117776152158037", label="22.93",style=bold];
"CSMD3_M" -> "FSTL5_E" [color="0 0 0.01645940014630587", label="22.97",style=bold];
"CNTNAP5_E" -> "CNTN5_E" [color="0 0 0.01580102414045359", label="23.64",style=bold];
"RYR2_M" -> "GALNTL6_M" [color="0 0 0.015142648134601311", label="23.80",style=bold];
"KRT1_M" -> "NEB_M" [color="0 0 0.014484272128749143", label="26.80",style=bold];
"PKHD1L1_M" -> "ERBB4_M" [color="0 0 0.013825896122896864", label="27.69",style=bold];
"LRRTM4_M" -> "DSCAM_M" [color="0 0 0.013167520117044695", label="28.16",style=bold];
"MYO3A_E" -> "CSMD1_E" [color="0 0 0.012509144111192416", label="28.26",style=bold];
"PKHD1L1_M" -> "DUSP27_M" [color="0 0 0.011850768105340248", label="28.60",style=bold];
"CSMD3_M" -> "GABRB3_M" [color="0 0 0.011192392099487969", label="29.39",style=bold];
"DSCAM_M" -> "NTSR2_M" [color="0 0 0.01053401609363569", label="30.55",style=bold];
"MYBPC2_M" -> "MUC4_M" [color="0 0 0.009875640087783522", label="31.29",style=bold];
"CR2_M" -> "CCDC67_M" [color="0 0 0.009217264081931242", label="32.37",style=bold];
"LCE2C_E" -> "KRT1_E" [color="0 0 0.008558888076079074", label="33.41",style=bold];
"LRRTM4_M" -> "CDH19_M" [color="0 0 0.007900512070226795", label="34.66",style=bold];
"PCDHA10_M" -> "NPFFR2_M" [color="0 0 0.007242136064374627", label="37.51",style=bold];
"PKHD1L1_E" -> "CR2_E" [color="0 0 0.006583760058522348", label="41.46",style=bold];
"CSMD3_M" -> "PCDHA12_M" [color="0 0 0.0059253840526700685", label="41.68",style=bold];
"NELL1_M" -> "CSMD3_M" [color="0 0 0.0052670080468179", label="49.06",style=bold];
"GABRB3_M" -> "CSMD1_M" [color="0 0 0.004608632040965621", label="51.63",style=bold];
"TTN_E" -> "NEB_E" [color="0 0 0.003950256035113453", label="56.31",style=bold];
"MYH6_E" -> "MYBPC2_E" [color="0 0 0.003291880029261174", label="59.38",style=bold];
"MYH7_E" -> "DUSP27_E" [color="0 0 0.0026335040234090057", label="64.59",style=bold];
"DUSP27_E" -> "TTN_E" [color="0 0 0.0019751280175567265", label="65.03",style=bold];
"MYH6_E" -> "MYH7_E" [color="0 0 0.0013167520117045584", label="70.29",style=bold];
"MYH7_M" -> "MYH6_M" [color="0 0 0.0006583760058522792", label="80.12",style=bold];
"PCDHA12_M" -> "PCDHA10_M" [color="0 0 0.0", label="86.23",style=bold];
}
```
