## Supplementary material for "New analysis framework incorporating mixed mutual information and scalable Bayesian networks for multimodal high dimensional genomic and epigenomic cancer data": Suppl Table 11

```
digraph G{
 ratio=fill;
 node [shape=box, style=rounded];
 edge [arrowhead=none];
"A2ML1_S";
"A2ML1_E";
"A2ML1_M";
"ABCA13_S";
"ABCA13_E";
"ABCA13_M";
"ADAMTS19_S";
"ADAMTS19_E";
"ADAMTS19_M";
"ADAMTS20_S";
"ADAMTS20_E";
"ADAMTS20_M";
"ADH7_S";
"ADH7_E";
"ADH7_M";
"AKNAD1_S";
"AKNAD1_E";
"AKNAD1_M";
"APOB_S";
"APOB_E";
"APOB_M";
"BNC1_S";
"BNC1_E";
"BNC1_M";
"CA4_S";
"CA4_E";
"CA4_M";
"CFHR1_S";
"CFHR1_E";
"CFHR1_M";
"CHRNA4_S";
"CHRNA4_E";
"CHRNA4_M";
"CNTNAP4_S";
"CNTNAP4_E";
"CNTNAP4_M";
"CNTNAP5_S";
"CNTNAP5_E";
"CNTNAP5_M";
"CRCT1_S";
"CRCT1_E";
"CRCT1_M";
"CSMD1_S";
"CSMD1_E";
"CSMD1_M";
"CSMD3_S";
"CSMD3_E";
"CSMD3_M";
"CWH43_S";
"CWH43_E";
"CWH43_M";
"DCT_S";
"DCT_E";
"DCT_M";
"DMBT1_S";
"DMBT1_E";
"DMBT1_M";
"DPP6_S";
"DPP6_E";
"DPP6_M";
"DSG1_S";
"DSG1_E";
"DSG1_M";
"DUSP27_S";
"DUSP27_E";
"DUSP27_M";
"FBN3_S";
"FBN3_E";
"FBN3_M";
"FLG_S";
"FLG_E";
"FLG_M";
"FOXG1_S";
"FOXG1_E";
"FOXG1_M";
"FREM2_S";
"FREM2_E";
"FREM2_M";
"GABRA1_S";
"GABRA1_E";
"GABRA1_M";
"GABRG2_S";
"GABRG2_E";
"GABRG2_M";
"GPR98_S";
"GPR98_E";
"GPR98_M";
"HTR4_S";
"HTR4_E";
"HTR4_M";
"IRX4_S";
"IRX4_E";
"IRX4_M";
"IVL_S";
"IVL_E";
"IVL_M";
"KCNA10_S";
"KCNA10_E";
"KCNA10_M";
"KCNV1_S";
"KCNV1_E";
"KCNV1_M";
"KPRP_S";
"KPRP_E";
"KPRP_M";
"KRT1_S";
"KRT1_E";
"KRT1_M";
"KRT24_S";
"KRT24_E";
"KRT24_M";
"KRT6C_S";
"KRT6C_E";
"KRT6C_M";
"KRT78_S";
"KRT78_E";
"KRT78_M";
"LHCGR_S";
"LHCGR_E";
"LHCGR_M";
"LIN28B_S";
"LIN28B_E";
"LIN28B_M";
"LIPF_S";
"LIPF_E";
"LIPF_M";
"LMOD2_S";
"LMOD2_E";
"LMOD2_M";
"LOC100190940_S";
"LOC100190940_E";
"LOC100190940_M";
"LRRC7_S";
"LRRC7_E";
"LRRC7_M";
"LRRIQ1_S";
"LRRIQ1_E";
"LRRIQ1_M";
"MUC16_S";
"MUC16_E";
"MUC16_M";
"MUC17_S";
"MUC17_E";
"MUC17_M";
"MUC2_S";
"MUC2_E";
"MUC2_M";
"MUC4_S";
"MUC4_E";
"MUC4_M";
"MUC5B_S";
"MUC5B_E";
"MUC5B_M";
"MUC6_S";
"MUC6_E";
"MUC6_M";
"MYBPC1_S";
"MYBPC1_E";
"MYBPC1_M";
"MYO18B_S";
"MYO18B_E";
"MYO18B_M";
"NDST4_S";
"NDST4_E";
"NDST4_M";
"NELL1_S";
"NELL1_E";
"NELL1_M";
"PAX1_S";
"PAX1_E";
"PAX1_M";
"PCDHA1_S";
"PCDHA1_E";
"PCDHA1_M";
"PCDHA13_S";
"PCDHA13_E";
"PCDHA13_M";
"PGC_S";
"PGC_E";
"PGC_M";
"PNLDC1_S";
"PNLDC1_E";
"PNLDC1_M";
"PNLIPRP3_S";
"PNLIPRP3_E";
"PNLIPRP3_M";
"RIMS1_S";
"RIMS1_E";
"RIMS1_M";
"RPL10L_S";
"RPL10L_E";
"RPL10L_M";
"SBSN_S";
"SBSN_E";
"SBSN_M";
"SERPINB11_S";
"SERPINB11_E";
"SERPINB11_M";
"SERPINB13_S";
"SERPINB13_E";
"SERPINB13_M";
"SLC12A1_S";
"SLC12A1_E";
"SLC12A1_M";
"SLC1A6_S";
"SLC1A6_E";
"SLC1A6_M";
"SOX1_S";
"SOX1_E";
"SOX1_M";
"SPAG17_S";
"SPAG17_E";
"SPAG17_M";
"SPRR2A_S";
"SPRR2A_E";
"SPRR2A_M";
"SYT16_S";
"SYT16_E";
"SYT16_M";
"TMPRSS11A_S";
"TMPRSS11A_E";
"TMPRSS11A_M";
"TPTE_S";
"TPTE_E";
"TPTE_M";
"TRIML2_S";
"TRIML2_E";
"TRIML2_M";
"TRPM3_S";
"TRPM3_E";
"TRPM3_M";
"UNC5D_S";
"UNC5D_E";
"UNC5D_M";
"ZAN_S";
"ZAN_E";
"ZAN_M";
"ZIC4_S";
"ZIC4_E";
"ZIC4_M";
"ZNF716_S";
"ZNF716_E";
"ZNF716_M";
"Tumor_Status";
"CFHR1_S" -> "LHCGR_E" [color="0 0 0.9", label="0.00",style=bold];
"IRX4_S" -> "MUC2_E" [color="0 0 0.8994267515923567", label="0.00",style=bold];
"KRT24_S" -> "ABCA13_S" [color="0 0 0.8988535031847134", label="0.01",style=bold];
"ADH7_S" -> "TRPM3_M" [color="0 0 0.8982802547770701", label="0.01",style=bold];
"ZNF716_S" -> "TRPM3_E" [color="0 0 0.8977070063694268", label="0.01",style=bold];
"KCNA10_S" -> "ADH7_M" [color="0 0 0.8971337579617834", label="0.01",style=bold];
"NELL1_M" -> "RIMS1_M" [color="0 0 0.8965605095541401", label="0.01",style=bold];
"CRCT1_S" -> "KRT1_E" [color="0 0 0.8959872611464968", label="0.01",style=bold];
"A2ML1_S" -> "TRPM3_E" [color="0 0 0.8954140127388536", label="0.01",style=bold];
"KRT6C_S" -> "PNLIPRP3_S" [color="0 0 0.8948407643312102", label="0.01",style=bold];
"CRCT1_S" -> "GABRA1_S" [color="0 0 0.8942675159235669", label="0.01",style=bold];
"LHCGR_S" -> "CSMD1_E" [color="0 0 0.8936942675159236", label="0.01",style=bold];
"PGC_S" -> "TPTE_S" [color="0 0 0.8931210191082802", label="0.02",style=bold];
"SBSN_S" -> "CFHR1_E" [color="0 0 0.892547770700637", label="0.02",style=bold];
"KRT78_S" -> "MUC16_S" [color="0 0 0.8919745222929937", label="0.02",style=bold];
"LMOD2_S" -> "LMOD2_M" [color="0 0 0.8914012738853503", label="0.02",style=bold];
"LIPF_S" -> "NELL1_E" [color="0 0 0.890828025477707", label="0.02",style=bold];
"KRT78_S" -> "AKNAD1_M" [color="0 0 0.8902547770700637", label="0.02",style=bold];
"LIN28B_S" -> "CWH43_S" [color="0 0 0.8896815286624205", label="0.02",style=bold];
"ZNF716_S" -> "RIMS1_S" [color="0 0 0.8891082802547771", label="0.02",style=bold];
"SERPINB11_S" -> "CNTNAP5_S" [color="0 0 0.8885350318471338", label="0.03",style=bold];
"SERPINB11_S" -> "LMOD2_E" [color="0 0 0.8879617834394905", label="0.03",style=bold];
"TMPRSS11A_S" -> "ZAN_E" [color="0 0 0.8873885350318471", label="0.03",style=bold];
"CFHR1_S" -> "FLG_S" [color="0 0 0.8868152866242038", label="0.04",style=bold];
"PGC_S" -> "LRRIQ1_S" [color="0 0 0.8862420382165606", label="0.04",style=bold];
"SOX1_S" -> "PCDHA1_S" [color="0 0 0.8856687898089172", label="0.04",style=bold];
"ADH7_S" -> "MYO18B_E" [color="0 0 0.8850955414012739", label="0.04",style=bold];
"LHCGR_S" -> "FREM2_E" [color="0 0 0.8845222929936306", label="0.04",style=bold];
"LOC100190940_S" -> "GABRG2_S" [color="0 0 0.8839490445859873", label="0.04",style=bold];
"KRT78_S" -> "MUC2_M" [color="0 0 0.8833757961783439", label="0.04",style=bold];
"TRIML2_S" -> "PCDHA13_E" [color="0 0 0.8828025477707007", label="0.04",style=bold];
"MYBPC1_S" -> "ABCA13_S" [color="0 0 0.8822292993630574", label="0.04",style=bold];
"CRCT1_S" -> "RPL10L_E" [color="0 0 0.881656050955414", label="0.04",style=bold];
"HTR4_S" -> "FLG_E" [color="0 0 0.8810828025477707", label="0.05",style=bold];
"TRIML2_S" -> "TRPM3_M" [color="0 0 0.8805095541401274", label="0.05",style=bold];
"LOC100190940_S" -> "CSMD1_E" [color="0 0 0.879936305732484", label="0.05",style=bold];
"CRCT1_S" -> "MUC16_M" [color="0 0 0.8793630573248408", label="0.05",style=bold];
"CRCT1_S" -> "RIMS1_E" [color="0 0 0.8787898089171975", label="0.05",style=bold];
"DSG1_S" -> "CFHR1_M" [color="0 0 0.8782165605095542", label="0.06",style=bold];
"ADH7_S" -> "AKNAD1_E" [color="0 0 0.8776433121019108", label="0.06",style=bold];
"CRCT1_S" -> "SBSN_S" [color="0 0 0.8770700636942675", label="0.06",style=bold];
"KPRP_S" -> "PAX1_E" [color="0 0 0.8764968152866243", label="0.06",style=bold];
"SERPINB11_S" -> "KCNV1_S" [color="0 0 0.8759235668789809", label="0.06",style=bold];
"IRX4_S" -> "CSMD3_E" [color="0 0 0.8753503184713376", label="0.06",style=bold];
"CA4_S" -> "A2ML1_M" [color="0 0 0.8747770700636943", label="0.06",style=bold];
"KRT6C_S" -> "UNC5D_E" [color="0 0 0.8742038216560509", label="0.07",style=bold];
"HTR4_S" -> "TMPRSS11A_M" [color="0 0 0.8736305732484076", label="0.07",style=bold];
"DCT_S" -> "FREM2_M" [color="0 0 0.8730573248407644", label="0.07",style=bold];
"ADH7_S" -> "MUC6_M" [color="0 0 0.872484076433121", label="0.07",style=bold];
"KRT24_S" -> "A2ML1_E" [color="0 0 0.8719108280254777", label="0.07",style=bold];
"CFHR1_S" -> "GPR98_E" [color="0 0 0.8713375796178344", label="0.07",style=bold];
"LIPF_S" -> "SLC12A1_E" [color="0 0 0.8707643312101911", label="0.09",style=bold];
"CRCT1_S" -> "ADAMTS19_E" [color="0 0 0.8701910828025478", label="0.09",style=bold];
"LMOD2_S" -> "FBN3_M" [color="0 0 0.8696178343949045", label="0.09",style=bold];
"TRIML2_S" -> "UNC5D_S" [color="0 0 0.8690445859872612", label="0.09",style=bold];
"PNLDC1_S" -> "MUC5B_M" [color="0 0 0.8684713375796178", label="0.09",style=bold];
"ZNF716_S" -> "Tumor_Status" [color="0 0 0.8678980891719745", label="0.10",style=bold];
"DUSP27_S" -> "SERPINB11_M" [color="0 0 0.8673248407643313", label="0.10",style=bold];
"CRCT1_S" -> "CNTNAP4_M" [color="0 0 0.866751592356688", label="0.11",style=bold];
"ZNF716_S" -> "MUC17_S" [color="0 0 0.8661783439490446", label="0.11",style=bold];
"CFHR1_S" -> "APOB_M" [color="0 0 0.8656050955414013", label="0.11",style=bold];
"PNLDC1_S" -> "NELL1_E" [color="0 0 0.865031847133758", label="0.11",style=bold];
"TMPRSS11A_S" -> "RIMS1_E" [color="0 0 0.8644585987261146", label="0.12",style=bold];
"ZNF716_S" -> "ADH7_M" [color="0 0 0.8638853503184714", label="0.12",style=bold];
"KPRP_S" -> "SOX1_S" [color="0 0 0.8633121019108281", label="0.12",style=bold];
"DUSP27_S" -> "APOB_S" [color="0 0 0.8627388535031847", label="0.12",style=bold];
"LIPF_S" -> "APOB_E" [color="0 0 0.8621656050955414", label="0.12",style=bold];
"ADH7_S" -> "CSMD1_E" [color="0 0 0.8615923566878981", label="0.12",style=bold];
"HTR4_S" -> "LIPF_M" [color="0 0 0.8610191082802547", label="0.12",style=bold];
"PGC_S" -> "TRIML2_S" [color="0 0 0.8604458598726115", label="0.13",style=bold];
"CFHR1_S" -> "TRIML2_S" [color="0 0 0.8598726114649682", label="0.13",style=bold];
"CWH43_S" -> "FBN3_E" [color="0 0 0.8592993630573249", label="0.13",style=bold];
"PGC_S" -> "APOB_S" [color="0 0 0.8587261146496815", label="0.13",style=bold];
"GABRG2_S" -> "UNC5D_E" [color="0 0 0.8581528662420382", label="0.13",style=bold];
"LMOD2_S" -> "A2ML1_E" [color="0 0 0.857579617834395", label="0.13",style=bold];
"PGC_S" -> "KRT1_M" [color="0 0 0.8570063694267516", label="0.13",style=bold];
"HTR4_S" -> "DUSP27_M" [color="0 0 0.8564331210191083", label="0.14",style=bold];
"IVL_S" -> "GABRG2_S" [color="0 0 0.855859872611465", label="0.14",style=bold];
"CWH43_S" -> "CSMD1_S" [color="0 0 0.8552866242038216", label="0.14",style=bold];
"ADH7_S" -> "GPR98_S" [color="0 0 0.8547133757961783", label="0.14",style=bold];
"PGC_S" -> "SPAG17_M" [color="0 0 0.8541401273885351", label="0.14",style=bold];
"KCNA10_S" -> "MYBPC1_E" [color="0 0 0.8535668789808918", label="0.14",style=bold];
"KPRP_S" -> "GPR98_S" [color="0 0 0.8529936305732484", label="0.15",style=bold];
"CFHR1_S" -> "CHRNA4_M" [color="0 0 0.8524203821656051", label="0.15",style=bold];
"ZNF716_S" -> "SLC1A6_M" [color="0 0 0.8518471337579618", label="0.15",style=bold];
"LMOD2_S" -> "CSMD1_E" [color="0 0 0.8512738853503184", label="0.15",style=bold];
"ADH7_S" -> "ADH7_E" [color="0 0 0.8507006369426752", label="0.15",style=bold];
"KRT24_S" -> "PCDHA1_E" [color="0 0 0.8501273885350319", label="0.15",style=bold];
"LIN28B_S" -> "CSMD3_S" [color="0 0 0.8495541401273885", label="0.15",style=bold];
"SBSN_S" -> "FREM2_E" [color="0 0 0.8489808917197452", label="0.15",style=bold];
"LOC100190940_S" -> "SLC12A1_E" [color="0 0 0.8484076433121019", label="0.16",style=bold];
"CHRNA4_S" -> "Tumor_Status" [color="0 0 0.8478343949044587", label="0.16",style=bold];
"SBSN_S" -> "LRRC7_E" [color="0 0 0.8472611464968153", label="0.16",style=bold];
"PGC_S" -> "IRX4_M" [color="0 0 0.846687898089172", label="0.16",style=bold];
"SOX1_S" -> "CA4_E" [color="0 0 0.8461146496815287", label="0.17",style=bold];
"KRT78_S" -> "SPAG17_M" [color="0 0 0.8455414012738853", label="0.17",style=bold];
"PGC_S" -> "FOXG1_M" [color="0 0 0.844968152866242", label="0.17",style=bold];
"ZNF716_S" -> "ADH7_E" [color="0 0 0.8443949044585988", label="0.17",style=bold];
"CFHR1_S" -> "TPTE_E" [color="0 0 0.8438216560509554", label="0.17",style=bold];
"ZNF716_S" -> "ABCA13_M" [color="0 0 0.8432484076433121", label="0.17",style=bold];
"IRX4_S" -> "BNC1_S" [color="0 0 0.8426751592356688", label="0.18",style=bold];
"PCDHA1_S" -> "LRRIQ1_E" [color="0 0 0.8421019108280254", label="0.19",style=bold];
"PGC_S" -> "KCNA10_M" [color="0 0 0.8415286624203822", label="0.19",style=bold];
"KRT78_S" -> "DSG1_M" [color="0 0 0.8409554140127389", label="0.19",style=bold];
"SBSN_S" -> "TRPM3_S" [color="0 0 0.8403821656050956", label="0.19",style=bold];
"LIN28B_S" -> "DPP6_E" [color="0 0 0.8398089171974522", label="0.19",style=bold];
"HTR4_S" -> "Tumor_Status" [color="0 0 0.8392356687898089", label="0.19",style=bold];
"KRT6C_S" -> "PCDHA13_S" [color="0 0 0.8386624203821657", label="0.20",style=bold];
"ZNF716_S" -> "GABRG2_M" [color="0 0 0.8380891719745223", label="0.20",style=bold];
"CA4_S" -> "LIPF_M" [color="0 0 0.837515923566879", label="0.20",style=bold];
"SOX1_S" -> "SBSN_E" [color="0 0 0.8369426751592357", label="0.21",style=bold];
"CHRNA4_S" -> "DUSP27_M" [color="0 0 0.8363694267515924", label="0.22",style=bold];
"PCDHA13_S" -> "DMBT1_E" [color="0 0 0.835796178343949", label="0.22",style=bold];
"SOX1_S" -> "SPRR2A_E" [color="0 0 0.8352229299363058", label="0.22",style=bold];
"LMOD2_S" -> "IRX4_M" [color="0 0 0.8346496815286625", label="0.22",style=bold];
"HTR4_S" -> "ZAN_M" [color="0 0 0.8340764331210191", label="0.22",style=bold];
"LMOD2_S" -> "DMBT1_E" [color="0 0 0.8335031847133758", label="0.22",style=bold];
"CRCT1_S" -> "SLC12A1_E" [color="0 0 0.8329299363057325", label="0.22",style=bold];
"CRCT1_S" -> "CSMD3_S" [color="0 0 0.8323566878980891", label="0.22",style=bold];
"LIN28B_S" -> "ZAN_E" [color="0 0 0.8317834394904459", label="0.22",style=bold];
"SYT16_S" -> "PCDHA1_E" [color="0 0 0.8312101910828026", label="0.23",style=bold];
"CFHR1_S" -> "ABCA13_E" [color="0 0 0.8306369426751592", label="0.23",style=bold];
"ZNF716_S" -> "TRIML2_M" [color="0 0 0.8300636942675159", label="0.23",style=bold];
"CFHR1_S" -> "MUC16_S" [color="0 0 0.8294904458598726", label="0.23",style=bold];
"RPL10L_S" -> "CSMD1_E" [color="0 0 0.8289171974522294", label="0.23",style=bold];
"LMOD2_S" -> "DPP6_S" [color="0 0 0.828343949044586", label="0.23",style=bold];
"LMOD2_S" -> "MUC5B_S" [color="0 0 0.8277707006369427", label="0.24",style=bold];
"PGC_S" -> "MUC2_E" [color="0 0 0.8271974522292994", label="0.24",style=bold];
"CRCT1_S" -> "ADAMTS20_E" [color="0 0 0.826624203821656", label="0.24",style=bold];
"SLC1A6_S" -> "KRT6C_M" [color="0 0 0.8260509554140127", label="0.24",style=bold];
"LIPF_S" -> "HTR4_M" [color="0 0 0.8254777070063695", label="0.24",style=bold];
"KRT78_S" -> "LMOD2_E" [color="0 0 0.8249044585987262", label="0.24",style=bold];
"ZNF716_S" -> "PAX1_M" [color="0 0 0.8243312101910828", label="0.25",style=bold];
"KRT1_S" -> "AKNAD1_E" [color="0 0 0.8237579617834395", label="0.25",style=bold];
"HTR4_S" -> "DUSP27_S" [color="0 0 0.8231847133757962", label="0.25",style=bold];
"SBSN_S" -> "GABRA1_S" [color="0 0 0.8226114649681529", label="0.25",style=bold];
"CFHR1_S" -> "ZIC4_M" [color="0 0 0.8220382165605096", label="0.25",style=bold];
"SBSN_S" -> "CNTNAP4_E" [color="0 0 0.8214649681528663", label="0.26",style=bold];
"SERPINB11_S" -> "CHRNA4_S" [color="0 0 0.8208917197452229", label="0.26",style=bold];
"ADH7_S" -> "LMOD2_E" [color="0 0 0.8203184713375796", label="0.26",style=bold];
"SOX1_S" -> "NELL1_M" [color="0 0 0.8197452229299363", label="0.26",style=bold];
"ZNF716_S" -> "LRRC7_E" [color="0 0 0.819171974522293", label="0.26",style=bold];
"LOC100190940_S" -> "LIPF_M" [color="0 0 0.8185987261146497", label="0.26",style=bold];
"ZNF716_S" -> "PNLIPRP3_M" [color="0 0 0.8180254777070064", label="0.26",style=bold];
"HTR4_S" -> "PAX1_E" [color="0 0 0.8174522292993631", label="0.26",style=bold];
"SERPINB13_S" -> "IVL_S" [color="0 0 0.8168789808917197", label="0.27",style=bold];
"PNLDC1_S" -> "LRRC7_M" [color="0 0 0.8163057324840765", label="0.27",style=bold];
"CFHR1_S" -> "NDST4_S" [color="0 0 0.8157324840764332", label="0.27",style=bold];
"HTR4_S" -> "CA4_S" [color="0 0 0.8151592356687898", label="0.27",style=bold];
"KRT78_S" -> "BNC1_M" [color="0 0 0.8145859872611465", label="0.28",style=bold];
"ZNF716_S" -> "LHCGR_E" [color="0 0 0.8140127388535032", label="0.29",style=bold];
"LIN28B_S" -> "DUSP27_E" [color="0 0 0.8134394904458598", label="0.29",style=bold];
"LIN28B_S" -> "CNTNAP4_M" [color="0 0 0.8128662420382166", label="0.29",style=bold];
"LOC100190940_S" -> "MYO18B_E" [color="0 0 0.8122929936305733", label="0.29",style=bold];
"LIN28B_S" -> "MUC4_E" [color="0 0 0.8117197452229299", label="0.29",style=bold];
"LIN28B_S" -> "NDST4_M" [color="0 0 0.8111464968152866", label="0.29",style=bold];
"CRCT1_S" -> "PGC_E" [color="0 0 0.8105732484076433", label="0.29",style=bold];
"LIN28B_S" -> "DMBT1_M" [color="0 0 0.81", label="0.29",style=bold];
"CNTNAP4_S" -> "KRT6C_S" [color="0 0 0.8094267515923567", label="0.29",style=bold];
"HTR4_S" -> "AKNAD1_M" [color="0 0 0.8088535031847134", label="0.29",style=bold];
"TRIML2_S" -> "LRRIQ1_M" [color="0 0 0.8082802547770701", label="0.30",style=bold];
"PNLDC1_S" -> "NDST4_E" [color="0 0 0.8077070063694267", label="0.30",style=bold];
"CRCT1_S" -> "FREM2_S" [color="0 0 0.8071337579617834", label="0.30",style=bold];
"MUC2_S" -> "SLC1A6_S" [color="0 0 0.8065605095541402", label="0.30",style=bold];
"LOC100190940_S" -> "KCNA10_M" [color="0 0 0.8059872611464969", label="0.30",style=bold];
"CWH43_S" -> "ABCA13_M" [color="0 0 0.8054140127388535", label="0.31",style=bold];
"HTR4_S" -> "CFHR1_M" [color="0 0 0.8048407643312102", label="0.31",style=bold];
"ZNF716_S" -> "A2ML1_M" [color="0 0 0.8042675159235669", label="0.31",style=bold];
"ADAMTS19_S" -> "MUC16_M" [color="0 0 0.8036942675159235", label="0.31",style=bold];
"PNLDC1_S" -> "DSG1_S" [color="0 0 0.8031210191082803", label="0.32",style=bold];
"ZNF716_S" -> "LOC100190940_E" [color="0 0 0.802547770700637", label="0.32",style=bold];
"LIPF_S" -> "RPL10L_M" [color="0 0 0.8019745222929936", label="0.32",style=bold];
"CFHR1_S" -> "RIMS1_E" [color="0 0 0.8014012738853503", label="0.32",style=bold];
"SLC1A6_S" -> "MYO18B_E" [color="0 0 0.800828025477707", label="0.32",style=bold];
"SBSN_S" -> "DMBT1_M" [color="0 0 0.8002547770700636", label="0.33",style=bold];
"ZNF716_S" -> "SLC12A1_M" [color="0 0 0.7996815286624204", label="0.34",style=bold];
"GABRA1_S" -> "SPRR2A_E" [color="0 0 0.7991082802547771", label="0.34",style=bold];
"ADH7_S" -> "PCDHA13_M" [color="0 0 0.7985350318471338", label="0.35",style=bold];
"LIPF_S" -> "GPR98_S" [color="0 0 0.7979617834394904", label="0.35",style=bold];
"KRT78_S" -> "A2ML1_M" [color="0 0 0.7973885350318471", label="0.35",style=bold];
"NELL1_S" -> "CSMD1_S" [color="0 0 0.7968152866242039", label="0.35",style=bold];
"KRT78_S" -> "HTR4_M" [color="0 0 0.7962420382165605", label="0.36",style=bold];
"KRT24_S" -> "CNTNAP5_S" [color="0 0 0.7956687898089172", label="0.37",style=bold];
"KRT24_S" -> "APOB_M" [color="0 0 0.7950955414012739", label="0.37",style=bold];
"LOC100190940_S" -> "MUC4_E" [color="0 0 0.7945222929936306", label="0.37",style=bold];
"KCNV1_S" -> "SPAG17_S" [color="0 0 0.7939490445859873", label="0.37",style=bold];
"HTR4_S" -> "KCNV1_S" [color="0 0 0.793375796178344", label="0.37",style=bold];
"SERPINB13_S" -> "KCNV1_S" [color="0 0 0.7928025477707007", label="0.37",style=bold];
"SBSN_S" -> "KRT1_E" [color="0 0 0.7922292993630573", label="0.37",style=bold];
"KPRP_S" -> "PCDHA13_S" [color="0 0 0.791656050955414", label="0.37",style=bold];
"DMBT1_S" -> "IRX4_M" [color="0 0 0.7910828025477707", label="0.37",style=bold];
"CWH43_S" -> "CFHR1_E" [color="0 0 0.7905095541401274", label="0.37",style=bold];
"SERPINB11_S" -> "KRT78_M" [color="0 0 0.7899363057324841", label="0.38",style=bold];
"SYT16_S" -> "IVL_S" [color="0 0 0.7893630573248408", label="0.38",style=bold];
"LRRC7_S" -> "PNLIPRP3_S" [color="0 0 0.7887898089171974", label="0.38",style=bold];
"HTR4_S" -> "MUC4_E" [color="0 0 0.7882165605095541", label="0.38",style=bold];
"LMOD2_S" -> "DSG1_M" [color="0 0 0.7876433121019109", label="0.38",style=bold];
"LOC100190940_S" -> "DCT_E" [color="0 0 0.7870700636942676", label="0.38",style=bold];
"MYO18B_S" -> "LHCGR_S" [color="0 0 0.7864968152866242", label="0.38",style=bold];
"PGC_S" -> "AKNAD1_M" [color="0 0 0.7859235668789809", label="0.39",style=bold];
"CFHR1_S" -> "DCT_M" [color="0 0 0.7853503184713376", label="0.39",style=bold];
"CFHR1_S" -> "MYO18B_E" [color="0 0 0.7847770700636942", label="0.39",style=bold];
"KRT78_S" -> "DPP6_E" [color="0 0 0.784203821656051", label="0.39",style=bold];
"LMOD2_S" -> "CSMD1_S" [color="0 0 0.7836305732484077", label="0.40",style=bold];
"CRCT1_S" -> "LHCGR_E" [color="0 0 0.7830573248407644", label="0.41",style=bold];
"LIN28B_S" -> "CNTNAP5_S" [color="0 0 0.782484076433121", label="0.41",style=bold];
"CRCT1_S" -> "ZAN_S" [color="0 0 0.7819108280254777", label="0.41",style=bold];
"PNLDC1_S" -> "GPR98_M" [color="0 0 0.7813375796178343", label="0.41",style=bold];
"RPL10L_S" -> "MUC4_E" [color="0 0 0.7807643312101911", label="0.41",style=bold];
"SOX1_S" -> "FLG_S" [color="0 0 0.7801910828025478", label="0.41",style=bold];
"SBSN_S" -> "HTR4_E" [color="0 0 0.7796178343949045", label="0.41",style=bold];
"LIPF_S" -> "KRT1_E" [color="0 0 0.7790445859872611", label="0.41",style=bold];
"DCT_S" -> "ADAMTS19_M" [color="0 0 0.7784713375796178", label="0.41",style=bold];
"LIN28B_S" -> "KRT1_E" [color="0 0 0.7778980891719746", label="0.42",style=bold];
"DMBT1_S" -> "ABCA13_E" [color="0 0 0.7773248407643312", label="0.42",style=bold];
"TMPRSS11A_S" -> "SOX1_M" [color="0 0 0.7767515923566879", label="0.42",style=bold];
"FBN3_S" -> "MYBPC1_M" [color="0 0 0.7761783439490446", label="0.42",style=bold];
"CHRNA4_S" -> "DPP6_E" [color="0 0 0.7756050955414013", label="0.42",style=bold];
"PCDHA13_S" -> "DUSP27_M" [color="0 0 0.775031847133758", label="0.43",style=bold];
"AKNAD1_S" -> "MUC17_E" [color="0 0 0.7744585987261147", label="0.43",style=bold];
"LIN28B_S" -> "TPTE_E" [color="0 0 0.7738853503184713", label="0.43",style=bold];
"ADH7_S" -> "MUC2_M" [color="0 0 0.773312101910828", label="0.43",style=bold];
"PNLDC1_S" -> "DUSP27_E" [color="0 0 0.7727388535031847", label="0.43",style=bold];
"CFHR1_S" -> "MUC4_M" [color="0 0 0.7721656050955414", label="0.43",style=bold];
"LIPF_S" -> "FREM2_E" [color="0 0 0.7715923566878982", label="0.44",style=bold];
"KRT24_S" -> "LMOD2_E" [color="0 0 0.7710191082802548", label="0.44",style=bold];
"LIPF_S" -> "MUC4_E" [color="0 0 0.7704458598726115", label="0.45",style=bold];
"PGC_S" -> "FLG_E" [color="0 0 0.7698726114649681", label="0.45",style=bold];
"KRT6C_S" -> "LIPF_E" [color="0 0 0.7692993630573248", label="0.45",style=bold];
"KRT6C_S" -> "LIPF_M" [color="0 0 0.7687261146496815", label="0.45",style=bold];
"AKNAD1_S" -> "SBSN_E" [color="0 0 0.7681528662420383", label="0.45",style=bold];
"CRCT1_S" -> "KCNV1_M" [color="0 0 0.7675796178343949", label="0.45",style=bold];
"KCNV1_S" -> "Tumor_Status" [color="0 0 0.7670063694267516", label="0.45",style=bold];
"KPRP_S" -> "MUC6_E" [color="0 0 0.7664331210191083", label="0.45",style=bold];
"CA4_S" -> "KCNA10_E" [color="0 0 0.7658598726114649", label="0.46",style=bold];
"HTR4_S" -> "TPTE_M" [color="0 0 0.7652866242038217", label="0.46",style=bold];
"LIPF_S" -> "TRIML2_E" [color="0 0 0.7647133757961784", label="0.46",style=bold];
"LMOD2_S" -> "FLG_S" [color="0 0 0.7641401273885351", label="0.46",style=bold];
"ZNF716_S" -> "MYBPC1_M" [color="0 0 0.7635668789808917", label="0.47",style=bold];
"CHRNA4_S" -> "RIMS1_S" [color="0 0 0.7629936305732484", label="0.47",style=bold];
"LIN28B_S" -> "PAX1_S" [color="0 0 0.762420382165605", label="0.47",style=bold];
"DMBT1_S" -> "PCDHA13_E" [color="0 0 0.7618471337579618", label="0.47",style=bold];
"TRIML2_S" -> "MYBPC1_S" [color="0 0 0.7612738853503185", label="0.48",style=bold];
"LOC100190940_S" -> "GPR98_M" [color="0 0 0.7607006369426752", label="0.48",style=bold];
"ADH7_S" -> "RIMS1_M" [color="0 0 0.7601273885350318", label="0.48",style=bold];
"GABRG2_S" -> "SLC12A1_M" [color="0 0 0.7595541401273885", label="0.48",style=bold];
"KCNV1_S" -> "CSMD3_S" [color="0 0 0.7589808917197453", label="0.49",style=bold];
"LMOD2_S" -> "PNLIPRP3_M" [color="0 0 0.7584076433121019", label="0.49",style=bold];
"CRCT1_S" -> "DCT_S" [color="0 0 0.7578343949044586", label="0.49",style=bold];
"LMOD2_S" -> "GPR98_M" [color="0 0 0.7572611464968153", label="0.49",style=bold];
"DSG1_S" -> "TRIML2_M" [color="0 0 0.756687898089172", label="0.49",style=bold];
"TRIML2_S" -> "TRPM3_E" [color="0 0 0.7561146496815286", label="0.50",style=bold];
"SYT16_S" -> "NELL1_E" [color="0 0 0.7555414012738854", label="0.50",style=bold];
"LMOD2_S" -> "CNTNAP4_M" [color="0 0 0.7549681528662421", label="0.50",style=bold];
"PGC_S" -> "SPRR2A_M" [color="0 0 0.7543949044585987", label="0.50",style=bold];
"HTR4_S" -> "MUC17_S" [color="0 0 0.7538216560509554", label="0.50",style=bold];
"PNLDC1_S" -> "MUC17_S" [color="0 0 0.7532484076433121", label="0.50",style=bold];
"AKNAD1_S" -> "DSG1_M" [color="0 0 0.7526751592356689", label="0.50",style=bold];
"KPRP_S" -> "LMOD2_M" [color="0 0 0.7521019108280255", label="0.50",style=bold];
"CA4_S" -> "DCT_E" [color="0 0 0.7515286624203822", label="0.51",style=bold];
"KPRP_S" -> "PGC_E" [color="0 0 0.7509554140127388", label="0.51",style=bold];
"ZAN_S" -> "DSG1_M" [color="0 0 0.7503821656050955", label="0.51",style=bold];
"HTR4_S" -> "MUC16_M" [color="0 0 0.7498089171974522", label="0.51",style=bold];
"GABRA1_S" -> "CSMD1_E" [color="0 0 0.749235668789809", label="0.51",style=bold];
"SERPINB13_S" -> "RPL10L_M" [color="0 0 0.7486624203821656", label="0.52",style=bold];
"HTR4_S" -> "KPRP_E" [color="0 0 0.7480891719745223", label="0.53",style=bold];
"CWH43_S" -> "ZAN_E" [color="0 0 0.747515923566879", label="0.53",style=bold];
"KRT1_S" -> "MUC4_S" [color="0 0 0.7469426751592356", label="0.53",style=bold];
"LIN28B_S" -> "SERPINB13_M" [color="0 0 0.7463694267515923", label="0.53",style=bold];
"ZNF716_S" -> "FOXG1_E" [color="0 0 0.7457961783439491", label="0.54",style=bold];
"TPTE_S" -> "LIPF_S" [color="0 0 0.7452229299363058", label="0.54",style=bold];
"ZNF716_S" -> "SOX1_E" [color="0 0 0.7446496815286624", label="0.55",style=bold];
"PGC_S" -> "SYT16_M" [color="0 0 0.7440764331210191", label="0.55",style=bold];
"LIN28B_S" -> "KCNA10_E" [color="0 0 0.7435031847133757", label="0.55",style=bold];
"IRX4_S" -> "FBN3_E" [color="0 0 0.7429299363057325", label="0.55",style=bold];
"DPP6_S" -> "MUC16_M" [color="0 0 0.7423566878980892", label="0.55",style=bold];
"PNLDC1_S" -> "PCDHA1_E" [color="0 0 0.7417834394904459", label="0.56",style=bold];
"TMPRSS11A_S" -> "DUSP27_M" [color="0 0 0.7412101910828026", label="0.56",style=bold];
"KRT78_S" -> "FLG_S" [color="0 0 0.7406369426751592", label="0.56",style=bold];
"KRT78_S" -> "LIN28B_E" [color="0 0 0.740063694267516", label="0.56",style=bold];
"KCNV1_S" -> "NELL1_E" [color="0 0 0.7394904458598726", label="0.56",style=bold];
"DPP6_S" -> "DSG1_S" [color="0 0 0.7389171974522293", label="0.56",style=bold];
"LMOD2_S" -> "MYBPC1_E" [color="0 0 0.738343949044586", label="0.56",style=bold];
"ZIC4_S" -> "PNLIPRP3_S" [color="0 0 0.7377707006369427", label="0.57",style=bold];
"PCDHA13_S" -> "CNTNAP5_S" [color="0 0 0.7371974522292993", label="0.57",style=bold];
"SBSN_S" -> "HTR4_M" [color="0 0 0.736624203821656", label="0.57",style=bold];
"ZNF716_S" -> "SERPINB11_E" [color="0 0 0.7360509554140128", label="0.57",style=bold];
"DUSP27_S" -> "FBN3_M" [color="0 0 0.7354777070063694", label="0.57",style=bold];
"IRX4_S" -> "PNLDC1_M" [color="0 0 0.7349044585987261", label="0.58",style=bold];
"IRX4_S" -> "CWH43_S" [color="0 0 0.7343312101910828", label="0.58",style=bold];
"SYT16_S" -> "MUC5B_E" [color="0 0 0.7337579617834395", label="0.58",style=bold];
"SERPINB13_S" -> "A2ML1_S" [color="0 0 0.7331847133757962", label="0.58",style=bold];
"KPRP_S" -> "PCDHA1_E" [color="0 0 0.7326114649681529", label="0.58",style=bold];
"A2ML1_S" -> "NDST4_S" [color="0 0 0.7320382165605095", label="0.59",style=bold];
"CFHR1_S" -> "FBN3_S" [color="0 0 0.7314649681528662", label="0.59",style=bold];
"KPRP_S" -> "DPP6_E" [color="0 0 0.7308917197452229", label="0.59",style=bold];
"LHCGR_S" -> "DUSP27_E" [color="0 0 0.7303184713375797", label="0.59",style=bold];
"KRT1_S" -> "TMPRSS11A_E" [color="0 0 0.7297452229299364", label="0.60",style=bold];
"DUSP27_S" -> "CHRNA4_E" [color="0 0 0.729171974522293", label="0.61",style=bold];
"ZNF716_S" -> "AKNAD1_E" [color="0 0 0.7285987261146497", label="0.61",style=bold];
"SBSN_S" -> "UNC5D_M" [color="0 0 0.7280254777070063", label="0.61",style=bold];
"PGC_S" -> "LIN28B_E" [color="0 0 0.727452229299363", label="0.61",style=bold];
"DCT_S" -> "DMBT1_M" [color="0 0 0.7268789808917198", label="0.61",style=bold];
"LMOD2_S" -> "NELL1_E" [color="0 0 0.7263057324840765", label="0.62",style=bold];
"LMOD2_S" -> "MUC2_S" [color="0 0 0.7257324840764331", label="0.62",style=bold];
"ZNF716_S" -> "MUC17_E" [color="0 0 0.7251592356687898", label="0.62",style=bold];
"KRT6C_S" -> "ADH7_S" [color="0 0 0.7245859872611465", label="0.63",style=bold];
"PGC_S" -> "GABRG2_S" [color="0 0 0.7240127388535031", label="0.63",style=bold];
"IRX4_S" -> "LOC100190940_E" [color="0 0 0.7234394904458599", label="0.63",style=bold];
"PGC_S" -> "RIMS1_E" [color="0 0 0.7228662420382166", label="0.63",style=bold];
"A2ML1_S" -> "LIPF_M" [color="0 0 0.7222929936305733", label="0.63",style=bold];
"DMBT1_S" -> "TRPM3_E" [color="0 0 0.7217197452229299", label="0.63",style=bold];
"CFHR1_S" -> "PCDHA1_S" [color="0 0 0.7211464968152866", label="0.64",style=bold];
"GABRG2_S" -> "MYO18B_E" [color="0 0 0.7205732484076433", label="0.64",style=bold];
"LIN28B_S" -> "MUC16_M" [color="0 0 0.72", label="0.64",style=bold];
"ZNF716_S" -> "SPAG17_M" [color="0 0 0.7194267515923567", label="0.64",style=bold];
"KRT24_S" -> "SLC12A1_E" [color="0 0 0.7188535031847134", label="0.65",style=bold];
"CFHR1_S" -> "MUC17_E" [color="0 0 0.71828025477707", label="0.65",style=bold];
"MYBPC1_S" -> "FBN3_M" [color="0 0 0.7177070063694267", label="0.65",style=bold];
"CFHR1_S" -> "NDST4_E" [color="0 0 0.7171337579617835", label="0.65",style=bold];
"DMBT1_S" -> "CFHR1_E" [color="0 0 0.7165605095541401", label="0.65",style=bold];
"ADH7_S" -> "PNLDC1_E" [color="0 0 0.7159872611464968", label="0.65",style=bold];
"PCDHA1_S" -> "CHRNA4_S" [color="0 0 0.7154140127388535", label="0.67",style=bold];
"TMPRSS11A_S" -> "NDST4_M" [color="0 0 0.7148407643312102", label="0.67",style=bold];
"LHCGR_S" -> "MYO18B_E" [color="0 0 0.7142675159235669", label="0.67",style=bold];
"CRCT1_S" -> "ZAN_E" [color="0 0 0.7136942675159236", label="0.68",style=bold];
"GABRA1_S" -> "DCT_E" [color="0 0 0.7131210191082803", label="0.68",style=bold];
"SLC1A6_S" -> "PNLDC1_E" [color="0 0 0.7125477707006369", label="0.68",style=bold];
"ADH7_S" -> "SBSN_M" [color="0 0 0.7119745222929936", label="0.68",style=bold];
"PNLDC1_S" -> "ADAMTS20_E" [color="0 0 0.7114012738853503", label="0.68",style=bold];
"ZAN_S" -> "UNC5D_E" [color="0 0 0.7108280254777071", label="0.69",style=bold];
"SERPINB13_S" -> "DPP6_M" [color="0 0 0.7102547770700637", label="0.69",style=bold];
"CA4_S" -> "LRRIQ1_S" [color="0 0 0.7096815286624204", label="0.69",style=bold];
"CRCT1_S" -> "MYO18B_M" [color="0 0 0.709108280254777", label="0.69",style=bold];
"LIPF_S" -> "MYO18B_E" [color="0 0 0.7085350318471337", label="0.69",style=bold];
"CRCT1_S" -> "MUC17_S" [color="0 0 0.7079617834394905", label="0.69",style=bold];
"CRCT1_S" -> "GPR98_S" [color="0 0 0.7073885350318472", label="0.69",style=bold];
"CRCT1_S" -> "KRT1_M" [color="0 0 0.7068152866242038", label="0.69",style=bold];
"KPRP_S" -> "KRT1_M" [color="0 0 0.7062420382165605", label="0.70",style=bold];
"PGC_S" -> "SOX1_S" [color="0 0 0.7056687898089172", label="0.70",style=bold];
"DCT_S" -> "DCT_E" [color="0 0 0.7050955414012738", label="0.70",style=bold];
"KRT24_S" -> "FBN3_S" [color="0 0 0.7045222929936306", label="0.70",style=bold];
"LHCGR_M" -> "MUC2_M" [color="0 0 0.7039490445859873", label="0.71",style=bold];
"HTR4_S" -> "MYO18B_E" [color="0 0 0.703375796178344", label="0.71",style=bold];
"PGC_S" -> "IRX4_E" [color="0 0 0.7028025477707006", label="0.71",style=bold];
"KRT1_S" -> "KRT78_M" [color="0 0 0.7022292993630573", label="0.72",style=bold];
"LOC100190940_S" -> "CFHR1_E" [color="0 0 0.701656050955414", label="0.72",style=bold];
"HTR4_S" -> "SBSN_E" [color="0 0 0.7010828025477707", label="0.72",style=bold];
"NDST4_S" -> "ADH7_M" [color="0 0 0.7005095541401274", label="0.72",style=bold];
"SERPINB11_S" -> "TRIML2_E" [color="0 0 0.6999363057324841", label="0.72",style=bold];
"KCNA10_S" -> "KPRP_S" [color="0 0 0.6993630573248408", label="0.72",style=bold];
"SERPINB11_S" -> "KRT1_E" [color="0 0 0.6987898089171974", label="0.73",style=bold];
"CFHR1_S" -> "CSMD1_M" [color="0 0 0.6982165605095542", label="0.73",style=bold];
"ADH7_S" -> "HTR4_E" [color="0 0 0.6976433121019108", label="0.73",style=bold];
"KRT78_S" -> "ADAMTS19_E" [color="0 0 0.6970700636942675", label="0.73",style=bold];
"KRT24_S" -> "CSMD1_E" [color="0 0 0.6964968152866242", label="0.74",style=bold];
"CA4_S" -> "MUC17_M" [color="0 0 0.6959235668789809", label="0.74",style=bold];
"SLC12A1_S" -> "FREM2_E" [color="0 0 0.6953503184713375", label="0.74",style=bold];
"SBSN_S" -> "CSMD1_S" [color="0 0 0.6947770700636943", label="0.75",style=bold];
"NELL1_S" -> "PCDHA1_E" [color="0 0 0.694203821656051", label="0.75",style=bold];
"SERPINB13_S" -> "FOXG1_E" [color="0 0 0.6936305732484076", label="0.75",style=bold];
"KRT78_S" -> "FBN3_S" [color="0 0 0.6930573248407643", label="0.76",style=bold];
"NDST4_S" -> "MUC16_S" [color="0 0 0.692484076433121", label="0.77",style=bold];
"PGC_S" -> "FOXG1_E" [color="0 0 0.6919108280254778", label="0.77",style=bold];
"CFHR1_S" -> "MUC4_E" [color="0 0 0.6913375796178344", label="0.77",style=bold];
"DSG1_S" -> "TRIML2_E" [color="0 0 0.6907643312101911", label="0.77",style=bold];
"KCNA10_S" -> "DSG1_M" [color="0 0 0.6901910828025477", label="0.77",style=bold];
"ZNF716_S" -> "RPL10L_E" [color="0 0 0.6896178343949044", label="0.78",style=bold];
"DUSP27_S" -> "NELL1_E" [color="0 0 0.6890445859872611", label="0.78",style=bold];
"ADH7_S" -> "MUC4_E" [color="0 0 0.6884713375796179", label="0.78",style=bold];
"LMOD2_S" -> "MUC2_M" [color="0 0 0.6878980891719745", label="0.78",style=bold];
"PGC_S" -> "SLC1A6_M" [color="0 0 0.6873248407643312", label="0.78",style=bold];
"LMOD2_S" -> "UNC5D_E" [color="0 0 0.6867515923566879", label="0.79",style=bold];
"ADH7_S" -> "KRT78_M" [color="0 0 0.6861783439490445", label="0.79",style=bold];
"DSG1_S" -> "HTR4_E" [color="0 0 0.6856050955414013", label="0.80",style=bold];
"KRT24_S" -> "ADAMTS20_E" [color="0 0 0.685031847133758", label="0.80",style=bold];
"LIN28B_S" -> "GPR98_S" [color="0 0 0.6844585987261147", label="0.80",style=bold];
"KCNV1_S" -> "CA4_S" [color="0 0 0.6838853503184713", label="0.80",style=bold];
"AKNAD1_S" -> "A2ML1_E" [color="0 0 0.683312101910828", label="0.81",style=bold];
"KPRP_S" -> "MUC16_S" [color="0 0 0.6827388535031848", label="0.81",style=bold];
"CA4_S" -> "MYO18B_S" [color="0 0 0.6821656050955414", label="0.81",style=bold];
"LRRC7_S" -> "SYT16_S" [color="0 0 0.6815923566878981", label="0.81",style=bold];
"CFHR1_S" -> "DPP6_S" [color="0 0 0.6810191082802548", label="0.81",style=bold];
"NDST4_S" -> "FREM2_E" [color="0 0 0.6804458598726115", label="0.82",style=bold];
"AKNAD1_S" -> "SERPINB11_M" [color="0 0 0.6798726114649681", label="0.82",style=bold];
"HTR4_S" -> "A2ML1_M" [color="0 0 0.6792993630573249", label="0.82",style=bold];
"BNC1_S" -> "GABRG2_M" [color="0 0 0.6787261146496815", label="0.82",style=bold];
"KRT78_S" -> "ADH7_M" [color="0 0 0.6781528662420382", label="0.82",style=bold];
"CFHR1_S" -> "LMOD2_E" [color="0 0 0.6775796178343949", label="0.82",style=bold];
"CA4_S" -> "MUC6_M" [color="0 0 0.6770063694267516", label="0.82",style=bold];
"A2ML1_S" -> "DSG1_M" [color="0 0 0.6764331210191082", label="0.82",style=bold];
"SERPINB13_S" -> "AKNAD1_M" [color="0 0 0.675859872611465", label="0.83",style=bold];
"SOX1_S" -> "GABRA1_S" [color="0 0 0.6752866242038217", label="0.83",style=bold];
"PNLDC1_S" -> "MYBPC1_E" [color="0 0 0.6747133757961783", label="0.83",style=bold];
"SBSN_S" -> "A2ML1_S" [color="0 0 0.674140127388535", label="0.83",style=bold];
"LOC100190940_S" -> "Tumor_Status" [color="0 0 0.6735668789808917", label="0.83",style=bold];
"KRT6C_S" -> "SYT16_M" [color="0 0 0.6729936305732485", label="0.84",style=bold];
"SERPINB11_S" -> "UNC5D_E" [color="0 0 0.6724203821656051", label="0.84",style=bold];
"CRCT1_S" -> "KRT78_E" [color="0 0 0.6718471337579618", label="0.84",style=bold];
"KCNA10_S" -> "FBN3_E" [color="0 0 0.6712738853503184", label="0.84",style=bold];
"HTR4_S" -> "CSMD3_E" [color="0 0 0.6707006369426751", label="0.84",style=bold];
"KRT6C_S" -> "TRIML2_E" [color="0 0 0.6701273885350318", label="0.84",style=bold];
"CFHR1_S" -> "TPTE_M" [color="0 0 0.6695541401273886", label="0.85",style=bold];
"PGC_S" -> "MYO18B_S" [color="0 0 0.6689808917197453", label="0.85",style=bold];
"BNC1_S" -> "TPTE_S" [color="0 0 0.6684076433121019", label="0.85",style=bold];
"LMOD2_S" -> "CWH43_E" [color="0 0 0.6678343949044586", label="0.85",style=bold];
"KCNA10_S" -> "PNLDC1_S" [color="0 0 0.6672611464968152", label="0.86",style=bold];
"CFHR1_S" -> "PNLDC1_S" [color="0 0 0.666687898089172", label="0.86",style=bold];
"KRT78_S" -> "CFHR1_E" [color="0 0 0.6661146496815287", label="0.86",style=bold];
"ZNF716_S" -> "ADAMTS19_E" [color="0 0 0.6655414012738854", label="0.86",style=bold];
"KRT78_S" -> "ADAMTS20_M" [color="0 0 0.664968152866242", label="0.86",style=bold];
"SERPINB11_S" -> "TMPRSS11A_M" [color="0 0 0.6643949044585987", label="0.86",style=bold];
"CFHR1_S" -> "ABCA13_M" [color="0 0 0.6638216560509554", label="0.87",style=bold];
"DCT_S" -> "MUC4_M" [color="0 0 0.663248407643312", label="0.87",style=bold];
"SERPINB11_S" -> "APOB_E" [color="0 0 0.6626751592356688", label="0.87",style=bold];
"CFHR1_S" -> "SERPINB11_S" [color="0 0 0.6621019108280255", label="0.87",style=bold];
"CFHR1_S" -> "PNLDC1_E" [color="0 0 0.6615286624203822", label="0.87",style=bold];
"CRCT1_S" -> "ZNF716_M" [color="0 0 0.6609554140127388", label="0.87",style=bold];
"AKNAD1_S" -> "MUC2_M" [color="0 0 0.6603821656050956", label="0.88",style=bold];
"SYT16_E" -> "SLC12A1_S" [color="0 0 0.6598089171974522", label="0.89",style=bold];
"HTR4_S" -> "DMBT1_M" [color="0 0 0.6592356687898089", label="0.89",style=bold];
"PGC_S" -> "TRIML2_M" [color="0 0 0.6586624203821656", label="0.89",style=bold];
"PGC_S" -> "LIPF_M" [color="0 0 0.6580891719745223", label="0.89",style=bold];
"GABRG2_S" -> "CFHR1_M" [color="0 0 0.6575159235668789", label="0.90",style=bold];
"LIPF_S" -> "MUC16_S" [color="0 0 0.6569426751592357", label="0.90",style=bold];
"ZNF716_S" -> "LMOD2_E" [color="0 0 0.6563694267515924", label="0.91",style=bold];
"CWH43_S" -> "MUC2_S" [color="0 0 0.655796178343949", label="0.91",style=bold];
"GABRG2_S" -> "UNC5D_S" [color="0 0 0.6552229299363057", label="0.91",style=bold];
"SBSN_S" -> "KCNA10_M" [color="0 0 0.6546496815286624", label="0.92",style=bold];
"LIN28B_S" -> "HTR4_E" [color="0 0 0.6540764331210192", label="0.92",style=bold];
"TMPRSS11A_S" -> "CNTNAP5_S" [color="0 0 0.6535031847133758", label="0.92",style=bold];
"CNTNAP4_S" -> "DUSP27_E" [color="0 0 0.6529299363057325", label="0.92",style=bold];
"ZNF716_S" -> "DPP6_E" [color="0 0 0.6523566878980892", label="0.92",style=bold];
"SOX1_S" -> "MUC5B_E" [color="0 0 0.6517834394904458", label="0.92",style=bold];
"ADH7_S" -> "LHCGR_M" [color="0 0 0.6512101910828025", label="0.92",style=bold];
"LMOD2_S" -> "SYT16_S" [color="0 0 0.6506369426751593", label="0.93",style=bold];
"KRT1_S" -> "IRX4_M" [color="0 0 0.650063694267516", label="0.93",style=bold];
"SERPINB13_S" -> "SYT16_E" [color="0 0 0.6494904458598726", label="0.93",style=bold];
"HTR4_S" -> "CA4_E" [color="0 0 0.6489171974522293", label="0.93",style=bold];
"ADH7_S" -> "RPL10L_S" [color="0 0 0.6483439490445859", label="0.93",style=bold];
"KRT6C_S" -> "CNTNAP5_S" [color="0 0 0.6477707006369426", label="0.93",style=bold];
"CA4_S" -> "CNTNAP4_E" [color="0 0 0.6471974522292994", label="0.93",style=bold];
"SERPINB13_S" -> "PAX1_E" [color="0 0 0.6466242038216561", label="0.94",style=bold];
"ZNF716_S" -> "BNC1_E" [color="0 0 0.6460509554140128", label="0.94",style=bold];
"NDST4_S" -> "MYO18B_E" [color="0 0 0.6454777070063694", label="0.95",style=bold];
"PNLDC1_S" -> "HTR4_M" [color="0 0 0.644904458598726", label="0.95",style=bold];
"DUSP27_S" -> "MUC6_E" [color="0 0 0.6443312101910827", label="0.96",style=bold];
"LIN28B_S" -> "KRT1_M" [color="0 0 0.6437579617834395", label="0.96",style=bold];
"CA4_S" -> "KCNA10_M" [color="0 0 0.6431847133757962", label="0.96",style=bold];
"HTR4_S" -> "GABRA1_S" [color="0 0 0.6426114649681529", label="0.96",style=bold];
"APOB_S" -> "ABCA13_M" [color="0 0 0.6420382165605095", label="0.96",style=bold];
"CNTNAP4_S" -> "TPTE_S" [color="0 0 0.6414649681528662", label="0.97",style=bold];
"KRT78_S" -> "TRIML2_S" [color="0 0 0.6408917197452229", label="0.97",style=bold];
"KRT78_S" -> "SLC12A1_E" [color="0 0 0.6403184713375796", label="0.97",style=bold];
"IRX4_S" -> "UNC5D_E" [color="0 0 0.6397452229299363", label="0.98",style=bold];
"PCDHA13_S" -> "DMBT1_S" [color="0 0 0.639171974522293", label="0.98",style=bold];
"SOX1_S" -> "LMOD2_M" [color="0 0 0.6385987261146497", label="0.98",style=bold];
"HTR4_S" -> "MUC17_E" [color="0 0 0.6380254777070063", label="0.98",style=bold];
"HTR4_S" -> "PCDHA1_M" [color="0 0 0.6374522292993631", label="0.98",style=bold];
"KCNA10_S" -> "NELL1_M" [color="0 0 0.6368789808917197", label="0.98",style=bold];
"KCNA10_S" -> "LOC100190940_E" [color="0 0 0.6363057324840764", label="0.99",style=bold];
"DPP6_S" -> "TRPM3_M" [color="0 0 0.6357324840764331", label="0.99",style=bold];
"ZNF716_S" -> "ZIC4_E" [color="0 0 0.6351592356687898", label="0.99",style=bold];
"SERPINB13_S" -> "CNTNAP5_M" [color="0 0 0.6345859872611466", label="0.99",style=bold];
"ABCA13_S" -> "APOB_E" [color="0 0 0.6340127388535032", label="0.99",style=bold];
"KRT78_S" -> "AKNAD1_E" [color="0 0 0.6334394904458598", label="0.99",style=bold];
"TMPRSS11A_S" -> "LIPF_M" [color="0 0 0.6328662420382165", label="1.00",style=bold];
"HTR4_S" -> "KRT1_S" [color="0 0 0.6322929936305732", label="1.00",style=bold];
"ZNF716_S" -> "PCDHA1_E" [color="0 0 0.63171974522293", label="1.00",style=bold];
"LMOD2_S" -> "KCNA10_E" [color="0 0 0.6311464968152867", label="1.01",style=bold];
"SERPINB13_S" -> "DPP6_S" [color="0 0 0.6305732484076433", label="1.01",style=bold];
"SERPINB13_S" -> "DCT_S" [color="0 0 0.63", label="1.01",style=bold];
"TMPRSS11A_S" -> "KRT6C_M" [color="0 0 0.6294267515923566", label="1.01",style=bold];
"KCNA10_S" -> "FLG_E" [color="0 0 0.6288535031847133", label="1.01",style=bold];
"HTR4_S" -> "LOC100190940_E" [color="0 0 0.6282802547770701", label="1.01",style=bold];
"SERPINB11_S" -> "PCDHA13_E" [color="0 0 0.6277070063694268", label="1.01",style=bold];
"FOXG1_S" -> "LMOD2_E" [color="0 0 0.6271337579617835", label="1.01",style=bold];
"SERPINB13_S" -> "NDST4_S" [color="0 0 0.6265605095541401", label="1.01",style=bold];
"DSG1_S" -> "ADH7_M" [color="0 0 0.6259872611464968", label="1.02",style=bold];
"DPP6_S" -> "ADAMTS19_E" [color="0 0 0.6254140127388534", label="1.02",style=bold];
"CWH43_S" -> "MUC4_S" [color="0 0 0.6248407643312102", label="1.02",style=bold];
"CRCT1_S" -> "KCNV1_E" [color="0 0 0.6242675159235669", label="1.02",style=bold];
"KRT1_S" -> "SOX1_E" [color="0 0 0.6236942675159236", label="1.03",style=bold];
"LIN28B_S" -> "KCNA10_M" [color="0 0 0.6231210191082803", label="1.03",style=bold];
"ADH7_S" -> "CNTNAP5_S" [color="0 0 0.6225477707006369", label="1.03",style=bold];
"CA4_S" -> "LOC100190940_E" [color="0 0 0.6219745222929935", label="1.03",style=bold];
"SERPINB13_S" -> "KCNA10_S" [color="0 0 0.6214012738853503", label="1.04",style=bold];
"KRT1_S" -> "DCT_E" [color="0 0 0.620828025477707", label="1.04",style=bold];
"SERPINB11_S" -> "CNTNAP4_M" [color="0 0 0.6202547770700637", label="1.04",style=bold];
"MYBPC1_S" -> "ZAN_M" [color="0 0 0.6196815286624204", label="1.05",style=bold];
"SERPINB11_S" -> "LHCGR_E" [color="0 0 0.619108280254777", label="1.05",style=bold];
"PNLDC1_S" -> "PNLDC1_M" [color="0 0 0.6185350318471338", label="1.05",style=bold];
"CRCT1_S" -> "MUC5B_S" [color="0 0 0.6179617834394904", label="1.05",style=bold];
"NDST4_S" -> "DUSP27_E" [color="0 0 0.6173885350318471", label="1.05",style=bold];
"KPRP_S" -> "CSMD1_E" [color="0 0 0.6168152866242038", label="1.05",style=bold];
"CWH43_S" -> "DUSP27_M" [color="0 0 0.6162420382165605", label="1.06",style=bold];
"MYBPC1_S" -> "MUC16_S" [color="0 0 0.6156687898089173", label="1.06",style=bold];
"MYBPC1_S" -> "DSG1_M" [color="0 0 0.6150955414012739", label="1.07",style=bold];
"MYBPC1_S" -> "PCDHA1_E" [color="0 0 0.6145222929936305", label="1.07",style=bold];
"LIN28B_S" -> "SLC1A6_S" [color="0 0 0.6139490445859872", label="1.07",style=bold];
"SBSN_S" -> "SOX1_S" [color="0 0 0.6133757961783439", label="1.08",style=bold];
"LOC100190940_S" -> "PCDHA13_E" [color="0 0 0.6128025477707006", label="1.08",style=bold];
"SERPINB11_S" -> "SPAG17_S" [color="0 0 0.6122292993630574", label="1.09",style=bold];
"FOXG1_S" -> "MUC16_S" [color="0 0 0.611656050955414", label="1.09",style=bold];
"SPAG17_S" -> "LRRC7_S" [color="0 0 0.6110828025477707", label="1.10",style=bold];
"NELL1_S" -> "SYT16_E" [color="0 0 0.6105095541401273", label="1.10",style=bold];
"CFHR1_S" -> "KCNA10_E" [color="0 0 0.609936305732484", label="1.10",style=bold];
"CRCT1_S" -> "IVL_S" [color="0 0 0.6093630573248408", label="1.11",style=bold];
"MYBPC1_S" -> "TPTE_S" [color="0 0 0.6087898089171975", label="1.11",style=bold];
"GABRA1_S" -> "FBN3_M" [color="0 0 0.6082165605095542", label="1.11",style=bold];
"PNLDC1_S" -> "UNC5D_E" [color="0 0 0.6076433121019108", label="1.11",style=bold];
"KRT78_S" -> "MUC4_S" [color="0 0 0.6070700636942675", label="1.11",style=bold];
"KRT78_S" -> "TMPRSS11A_M" [color="0 0 0.6064968152866241", label="1.12",style=bold];
"CFHR1_S" -> "CSMD1_E" [color="0 0 0.6059235668789809", label="1.12",style=bold];
"SERPINB11_S" -> "KCNA10_E" [color="0 0 0.6053503184713376", label="1.12",style=bold];
"MUC6_S" -> "CFHR1_E" [color="0 0 0.6047770700636943", label="1.13",style=bold];
"PNLIPRP3_S" -> "ADH7_M" [color="0 0 0.604203821656051", label="1.13",style=bold];
"CWH43_S" -> "MUC16_M" [color="0 0 0.6036305732484076", label="1.13",style=bold];
"AKNAD1_S" -> "MUC4_S" [color="0 0 0.6030573248407642", label="1.14",style=bold];
"SERPINB13_S" -> "MUC2_E" [color="0 0 0.602484076433121", label="1.14",style=bold];
"HTR4_S" -> "LIN28B_E" [color="0 0 0.6019108280254777", label="1.14",style=bold];
"LRRC7_S" -> "TRPM3_S" [color="0 0 0.6013375796178344", label="1.14",style=bold];
"PNLDC1_S" -> "ABCA13_M" [color="0 0 0.6007643312101911", label="1.14",style=bold];
"NELL1_S" -> "KCNV1_E" [color="0 0 0.6001910828025477", label="1.14",style=bold];
"TRIML2_S" -> "ADAMTS19_S" [color="0 0 0.5996178343949045", label="1.15",style=bold];
"KRT24_S" -> "PCDHA13_M" [color="0 0 0.5990445859872611", label="1.15",style=bold];
"SERPINB11_S" -> "NELL1_E" [color="0 0 0.5984713375796178", label="1.15",style=bold];
"ADH7_S" -> "DCT_M" [color="0 0 0.5978980891719745", label="1.15",style=bold];
"LHCGR_S" -> "DPP6_E" [color="0 0 0.5973248407643312", label="1.17",style=bold];
"NDST4_S" -> "MUC4_E" [color="0 0 0.596751592356688", label="1.17",style=bold];
"LOC100190940_S" -> "ZAN_M" [color="0 0 0.5961783439490446", label="1.17",style=bold];
"PNLDC1_S" -> "PCDHA13_E" [color="0 0 0.5956050955414013", label="1.17",style=bold];
"HTR4_S" -> "CFHR1_E" [color="0 0 0.5950318471337579", label="1.17",style=bold];
"SERPINB13_S" -> "KCNA10_E" [color="0 0 0.5944585987261146", label="1.17",style=bold];
"DCT_S" -> "TPTE_E" [color="0 0 0.5938853503184713", label="1.18",style=bold];
"SPAG17_S" -> "APOB_E" [color="0 0 0.5933121019108281", label="1.19",style=bold];
"SERPINB13_S" -> "LRRC7_M" [color="0 0 0.5927388535031848", label="1.19",style=bold];
"SBSN_S" -> "AKNAD1_E" [color="0 0 0.5921656050955414", label="1.20",style=bold];
"CA4_S" -> "TPTE_S" [color="0 0 0.591592356687898", label="1.20",style=bold];
"ADH7_S" -> "DSG1_S" [color="0 0 0.5910191082802547", label="1.20",style=bold];
"KCNA10_S" -> "PGC_E" [color="0 0 0.5904458598726114", label="1.21",style=bold];
"CA4_S" -> "CNTNAP5_S" [color="0 0 0.5898726114649682", label="1.21",style=bold];
"LMOD2_S" -> "KPRP_M" [color="0 0 0.5892993630573249", label="1.21",style=bold];
"A2ML1_S" -> "MYO18B_E" [color="0 0 0.5887261146496815", label="1.21",style=bold];
"CWH43_S" -> "MYO18B_M" [color="0 0 0.5881528662420382", label="1.21",style=bold];
"CFHR1_S" -> "SPRR2A_M" [color="0 0 0.5875796178343948", label="1.21",style=bold];
"SERPINB13_S" -> "SBSN_M" [color="0 0 0.5870063694267516", label="1.22",style=bold];
"MYBPC1_S" -> "UNC5D_E" [color="0 0 0.5864331210191083", label="1.22",style=bold];
"ZNF716_S" -> "GABRA1_E" [color="0 0 0.585859872611465", label="1.22",style=bold];
"CRCT1_S" -> "SPRR2A_E" [color="0 0 0.5852866242038217", label="1.23",style=bold];
"DCT_S" -> "CNTNAP5_M" [color="0 0 0.5847133757961783", label="1.23",style=bold];
"SERPINB13_S" -> "ADH7_S" [color="0 0 0.5841401273885349", label="1.23",style=bold];
"SBSN_S" -> "NDST4_M" [color="0 0 0.5835668789808917", label="1.23",style=bold];
"LHCGR_S" -> "TRPM3_M" [color="0 0 0.5829936305732484", label="1.23",style=bold];
"HTR4_S" -> "FREM2_M" [color="0 0 0.5824203821656051", label="1.23",style=bold];
"CFHR1_S" -> "HTR4_M" [color="0 0 0.5818471337579618", label="1.23",style=bold];
"FLG_S" -> "PCDHA1_E" [color="0 0 0.5812738853503184", label="1.24",style=bold];
"CWH43_S" -> "MUC16_S" [color="0 0 0.5807006369426752", label="1.24",style=bold];
"CWH43_S" -> "MUC17_E" [color="0 0 0.5801273885350318", label="1.24",style=bold];
"ZNF716_S" -> "GPR98_M" [color="0 0 0.5795541401273885", label="1.24",style=bold];
"KRT24_S" -> "TPTE_E" [color="0 0 0.5789808917197452", label="1.24",style=bold];
"DCT_S" -> "MUC5B_E" [color="0 0 0.5784076433121019", label="1.25",style=bold];
"SLC1A6_M" -> "CSMD3_S" [color="0 0 0.5778343949044586", label="1.25",style=bold];
"LIN28B_S" -> "PNLDC1_E" [color="0 0 0.5772611464968153", label="1.25",style=bold];
"SBSN_S" -> "PCDHA1_E" [color="0 0 0.576687898089172", label="1.25",style=bold];
"LMOD2_S" -> "PCDHA1_M" [color="0 0 0.5761146496815286", label="1.26",style=bold];
"ZIC4_S" -> "TRPM3_E" [color="0 0 0.5755414012738853", label="1.26",style=bold];
"CA4_S" -> "MUC16_M" [color="0 0 0.574968152866242", label="1.26",style=bold];
"PNLDC1_S" -> "SPRR2A_M" [color="0 0 0.5743949044585988", label="1.26",style=bold];
"SERPINB13_S" -> "CWH43_M" [color="0 0 0.5738216560509555", label="1.26",style=bold];
"PGC_S" -> "PNLIPRP3_M" [color="0 0 0.5732484076433121", label="1.26",style=bold];
"RPL10L_S" -> "TPTE_E" [color="0 0 0.5726751592356687", label="1.26",style=bold];
"LMOD2_S" -> "CNTNAP5_M" [color="0 0 0.5721019108280254", label="1.27",style=bold];
"CA4_S" -> "MYBPC1_E" [color="0 0 0.5715286624203821", label="1.27",style=bold];
"ADH7_S" -> "MUC5B_S" [color="0 0 0.5709554140127389", label="1.27",style=bold];
"CRCT1_S" -> "AKNAD1_M" [color="0 0 0.5703821656050956", label="1.27",style=bold];
"DUSP27_S" -> "LRRC7_M" [color="0 0 0.5698089171974522", label="1.27",style=bold];
"KPRP_S" -> "A2ML1_E" [color="0 0 0.5692356687898089", label="1.28",style=bold];
"NDST4_S" -> "SERPINB11_M" [color="0 0 0.5686624203821655", label="1.29",style=bold];
"KRT6C_S" -> "MUC4_E" [color="0 0 0.5680891719745222", label="1.29",style=bold];
"CHRNA4_S" -> "MUC4_M" [color="0 0 0.567515923566879", label="1.30",style=bold];
"TRPM3_S" -> "MUC6_S" [color="0 0 0.5669426751592357", label="1.30",style=bold];
"PNLDC1_S" -> "FLG_E" [color="0 0 0.5663694267515924", label="1.31",style=bold];
"PGC_S" -> "NELL1_E" [color="0 0 0.565796178343949", label="1.31",style=bold];
"LMOD2_S" -> "PNLDC1_E" [color="0 0 0.5652229299363057", label="1.31",style=bold];
"CWH43_S" -> "MYO18B_E" [color="0 0 0.5646496815286624", label="1.34",style=bold];
"KCNA10_S" -> "MYO18B_S" [color="0 0 0.5640764331210191", label="1.34",style=bold];
"PNLDC1_S" -> "LMOD2_E" [color="0 0 0.5635031847133758", label="1.34",style=bold];
"KCNA10_S" -> "TRPM3_M" [color="0 0 0.5629299363057325", label="1.35",style=bold];
"TMPRSS11A_S" -> "MYO18B_M" [color="0 0 0.5623566878980892", label="1.35",style=bold];
"DSG1_S" -> "RPL10L_M" [color="0 0 0.5617834394904458", label="1.35",style=bold];
"PGC_S" -> "PAX1_S" [color="0 0 0.5612101910828025", label="1.35",style=bold];
"SERPINB13_S" -> "GABRA1_E" [color="0 0 0.5606369426751592", label="1.36",style=bold];
"HTR4_S" -> "SERPINB11_M" [color="0 0 0.5600636942675159", label="1.36",style=bold];
"ZNF716_S" -> "SPAG17_E" [color="0 0 0.5594904458598726", label="1.36",style=bold];
"CNTNAP4_S" -> "LIPF_M" [color="0 0 0.5589171974522293", label="1.36",style=bold];
"SOX1_S" -> "ADH7_M" [color="0 0 0.558343949044586", label="1.36",style=bold];
"KRT78_S" -> "DCT_E" [color="0 0 0.5577707006369427", label="1.37",style=bold];
"SERPINB11_S" -> "IRX4_S" [color="0 0 0.5571974522292993", label="1.37",style=bold];
"CFHR1_S" -> "PGC_E" [color="0 0 0.556624203821656", label="1.37",style=bold];
"KRT6C_S" -> "MYO18B_E" [color="0 0 0.5560509554140127", label="1.37",style=bold];
"LIPF_S" -> "GABRG2_S" [color="0 0 0.5554777070063694", label="1.38",style=bold];
"PGC_S" -> "FREM2_S" [color="0 0 0.5549044585987262", label="1.39",style=bold];
"KCNA10_S" -> "SLC1A6_S" [color="0 0 0.5543312101910828", label="1.39",style=bold];
"TRIML2_S" -> "NDST4_E" [color="0 0 0.5537579617834394", label="1.39",style=bold];
"CHRNA4_S" -> "ABCA13_S" [color="0 0 0.5531847133757961", label="1.40",style=bold];
"KRT6C_S" -> "NELL1_E" [color="0 0 0.5526114649681528", label="1.40",style=bold];
"KRT1_S" -> "FOXG1_S" [color="0 0 0.5520382165605096", label="1.41",style=bold];
"CFHR1_S" -> "SLC1A6_M" [color="0 0 0.5514649681528663", label="1.41",style=bold];
"ZNF716_S" -> "SLC12A1_E" [color="0 0 0.5508917197452229", label="1.42",style=bold];
"ZNF716_S" -> "SLC1A6_E" [color="0 0 0.5503184713375796", label="1.42",style=bold];
"FBN3_S" -> "CSMD1_E" [color="0 0 0.5497452229299362", label="1.42",style=bold];
"PNLDC1_S" -> "Tumor_Status" [color="0 0 0.5491719745222929", label="1.43",style=bold];
"GABRA1_S" -> "LHCGR_E" [color="0 0 0.5485987261146497", label="1.45",style=bold];
"ZNF716_S" -> "DSG1_E" [color="0 0 0.5480254777070064", label="1.46",style=bold];
"BNC1_S" -> "MUC6_E" [color="0 0 0.5474522292993631", label="1.46",style=bold];
"FREM2_S" -> "ADAMTS19_S" [color="0 0 0.5468789808917197", label="1.46",style=bold];
"CWH43_S" -> "MYBPC1_M" [color="0 0 0.5463057324840764", label="1.47",style=bold];
"CRCT1_S" -> "MUC2_S" [color="0 0 0.545732484076433", label="1.47",style=bold];
"FOXG1_S" -> "PAX1_S" [color="0 0 0.5451592356687898", label="1.47",style=bold];
"PNLIPRP3_S" -> "PNLIPRP3_M" [color="0 0 0.5445859872611465", label="1.47",style=bold];
"LIN28B_S" -> "ADAMTS20_S" [color="0 0 0.5440127388535032", label="1.48",style=bold];
"KRT6C_S" -> "LRRIQ1_M" [color="0 0 0.5434394904458599", label="1.48",style=bold];
"KRT1_S" -> "TRIML2_S" [color="0 0 0.5428662420382165", label="1.48",style=bold];
"KRT24_S" -> "PNLIPRP3_M" [color="0 0 0.5422929936305732", label="1.49",style=bold];
"SBSN_S" -> "RPL10L_M" [color="0 0 0.5417197452229299", label="1.49",style=bold];
"SBSN_S" -> "KPRP_M" [color="0 0 0.5411464968152866", label="1.50",style=bold];
"TRIML2_S" -> "DUSP27_M" [color="0 0 0.5405732484076433", label="1.50",style=bold];
"CNTNAP4_S" -> "MUC4_E" [color="0 0 0.54", label="1.50",style=bold];
"IRX4_S" -> "CA4_M" [color="0 0 0.5394267515923566", label="1.51",style=bold];
"MYO18B_S" -> "RIMS1_E" [color="0 0 0.5388535031847134", label="1.52",style=bold];
"CA4_S" -> "FBN3_E" [color="0 0 0.53828025477707", label="1.52",style=bold];
"PAX1_S" -> "SLC1A6_S" [color="0 0 0.5377070063694267", label="1.52",style=bold];
"MYBPC1_S" -> "PCDHA13_E" [color="0 0 0.5371337579617834", label="1.52",style=bold];
"LMOD2_S" -> "CHRNA4_E" [color="0 0 0.5365605095541401", label="1.52",style=bold];
"LOC100190940_S" -> "FOXG1_S" [color="0 0 0.5359872611464969", label="1.53",style=bold];
"RPL10L_S" -> "FOXG1_E" [color="0 0 0.5354140127388535", label="1.53",style=bold];
"SOX1_S" -> "PNLDC1_E" [color="0 0 0.5348407643312102", label="1.53",style=bold];
"SBSN_S" -> "Tumor_Status" [color="0 0 0.5342675159235668", label="1.54",style=bold];
"CA4_S" -> "AKNAD1_E" [color="0 0 0.5336942675159235", label="1.54",style=bold];
"BNC1_S" -> "PCDHA1_E" [color="0 0 0.5331210191082802", label="1.54",style=bold];
"KCNA10_S" -> "KCNA10_E" [color="0 0 0.532547770700637", label="1.54",style=bold];
"CA4_S" -> "MUC17_S" [color="0 0 0.5319745222929937", label="1.55",style=bold];
"LMOD2_S" -> "MUC6_M" [color="0 0 0.5314012738853503", label="1.55",style=bold];
"FOXG1_S" -> "ABCA13_E" [color="0 0 0.5308280254777069", label="1.55",style=bold];
"PGC_S" -> "LRRIQ1_M" [color="0 0 0.5302547770700636", label="1.55",style=bold];
"CHRNA4_S" -> "LIPF_M" [color="0 0 0.5296815286624204", label="1.56",style=bold];
"ZNF716_S" -> "CSMD1_S" [color="0 0 0.5291082802547771", label="1.56",style=bold];
"KRT24_S" -> "FBN3_E" [color="0 0 0.5285350318471338", label="1.56",style=bold];
"KRT6C_S" -> "DCT_M" [color="0 0 0.5279617834394904", label="1.57",style=bold];
"SERPINB11_S" -> "MUC6_E" [color="0 0 0.5273885350318471", label="1.57",style=bold];
"GABRA1_S" -> "HTR4_E" [color="0 0 0.5268152866242037", label="1.57",style=bold];
"ZNF716_S" -> "NDST4_E" [color="0 0 0.5262420382165605", label="1.57",style=bold];
"KCNA10_S" -> "KRT24_E" [color="0 0 0.5256687898089172", label="1.57",style=bold];
"BNC1_E" -> "ADH7_E" [color="0 0 0.5250955414012739", label="1.58",style=bold];
"TMPRSS11A_S" -> "SLC12A1_E" [color="0 0 0.5245222929936306", label="1.58",style=bold];
"LIN28B_S" -> "PGC_M" [color="0 0 0.5239490445859872", label="1.58",style=bold];
"KPRP_S" -> "LMOD2_E" [color="0 0 0.5233757961783438", label="1.58",style=bold];
"SERPINB13_S" -> "CWH43_E" [color="0 0 0.5228025477707006", label="1.58",style=bold];
"LMOD2_S" -> "FLG_E" [color="0 0 0.5222292993630573", label="1.59",style=bold];
"CNTNAP5_S" -> "TMPRSS11A_M" [color="0 0 0.521656050955414", label="1.59",style=bold];
"HTR4_S" -> "SERPINB13_M" [color="0 0 0.5210828025477707", label="1.59",style=bold];
"LMOD2_S" -> "LHCGR_M" [color="0 0 0.5205095541401273", label="1.59",style=bold];
"ADAMTS19_S" -> "LOC100190940_E" [color="0 0 0.5199363057324841", label="1.60",style=bold];
"KPRP_S" -> "TRIML2_E" [color="0 0 0.5193630573248407", label="1.60",style=bold];
"LMOD2_S" -> "LIPF_M" [color="0 0 0.5187898089171974", label="1.61",style=bold];
"CNTNAP4_S" -> "LHCGR_E" [color="0 0 0.5182165605095541", label="1.61",style=bold];
"TMPRSS11A_S" -> "SYT16_M" [color="0 0 0.5176433121019108", label="1.61",style=bold];
"KCNA10_S" -> "LIPF_S" [color="0 0 0.5170700636942676", label="1.62",style=bold];
"LIPF_S" -> "KRT6C_M" [color="0 0 0.5164968152866242", label="1.62",style=bold];
"DSG1_S" -> "LIN28B_E" [color="0 0 0.5159235668789809", label="1.63",style=bold];
"ADH7_S" -> "ZAN_E" [color="0 0 0.5153503184713375", label="1.63",style=bold];
"SERPINB11_S" -> "TRIML2_M" [color="0 0 0.5147770700636942", label="1.63",style=bold];
"SYT16_S" -> "NDST4_E" [color="0 0 0.5142038216560509", label="1.63",style=bold];
"KRT24_S" -> "PCDHA13_E" [color="0 0 0.5136305732484077", label="1.63",style=bold];
"CFHR1_S" -> "MUC2_M" [color="0 0 0.5130573248407644", label="1.64",style=bold];
"MYBPC1_S" -> "MYBPC1_E" [color="0 0 0.512484076433121", label="1.64",style=bold];
"FOXG1_S" -> "LOC100190940_E" [color="0 0 0.5119108280254776", label="1.65",style=bold];
"HTR4_S" -> "APOB_S" [color="0 0 0.5113375796178343", label="1.65",style=bold];
"ZIC4_S" -> "TRIML2_E" [color="0 0 0.510764331210191", label="1.65",style=bold];
"ZNF716_S" -> "CSMD1_M" [color="0 0 0.5101910828025478", label="1.65",style=bold];
"DUSP27_S" -> "SBSN_M" [color="0 0 0.5096178343949045", label="1.67",style=bold];
"SERPINB11_S" -> "LRRIQ1_S" [color="0 0 0.5090445859872611", label="1.68",style=bold];
"ZNF716_S" -> "CNTNAP4_M" [color="0 0 0.5084713375796178", label="1.68",style=bold];
"SLC1A6_S" -> "CA4_E" [color="0 0 0.5078980891719744", label="1.68",style=bold];
"PNLDC1_S" -> "SERPINB13_M" [color="0 0 0.5073248407643312", label="1.68",style=bold];
"KRT1_S" -> "RIMS1_S" [color="0 0 0.5067515923566879", label="1.69",style=bold];
"PCDHA1_S" -> "GPR98_S" [color="0 0 0.5061783439490446", label="1.69",style=bold];
"SBSN_S" -> "ABCA13_S" [color="0 0 0.5056050955414013", label="1.69",style=bold];
"KCNA10_S" -> "MUC2_S" [color="0 0 0.5050318471337579", label="1.69",style=bold];
"KRT24_S" -> "AKNAD1_M" [color="0 0 0.5044585987261146", label="1.69",style=bold];
"PNLDC1_S" -> "SLC12A1_E" [color="0 0 0.5038853503184713", label="1.70",style=bold];
"MUC17_E" -> "NELL1_S" [color="0 0 0.503312101910828", label="1.71",style=bold];
"PGC_S" -> "CNTNAP4_M" [color="0 0 0.5027388535031847", label="1.71",style=bold];
"KCNA10_S" -> "FLG_M" [color="0 0 0.5021656050955414", label="1.72",style=bold];
"PCDHA1_S" -> "Tumor_Status" [color="0 0 0.5015923566878981", label="1.72",style=bold];
"DCT_S" -> "NDST4_E" [color="0 0 0.5010191082802548", label="1.72",style=bold];
"DUSP27_S" -> "Tumor_Status" [color="0 0 0.5004458598726114", label="1.72",style=bold];
"A2ML1_S" -> "SLC12A1_E" [color="0 0 0.49987261146496814", label="1.73",style=bold];
"DCT_S" -> "CNTNAP4_M" [color="0 0 0.4992993630573248", label="1.73",style=bold];
"AKNAD1_S" -> "MYBPC1_E" [color="0 0 0.49872611464968153", label="1.74",style=bold];
"KRT78_S" -> "DPP6_M" [color="0 0 0.4981528662420382", label="1.74",style=bold];
"KRT24_S" -> "MYBPC1_E" [color="0 0 0.49757961783439486", label="1.74",style=bold];
"ADH7_S" -> "APOB_S" [color="0 0 0.4970063694267516", label="1.74",style=bold];
"SERPINB11_S" -> "CFHR1_E" [color="0 0 0.49643312101910825", label="1.75",style=bold];
"A2ML1_S" -> "MUC5B_E" [color="0 0 0.49585987261146497", label="1.75",style=bold];
"SLC1A6_S" -> "SYT16_S" [color="0 0 0.49528662420382163", label="1.76",style=bold];
"ZNF716_S" -> "MYO18B_S" [color="0 0 0.4947133757961783", label="1.76",style=bold];
"PAX1_S" -> "TRIML2_M" [color="0 0 0.494140127388535", label="1.76",style=bold];
"RPL10L_S" -> "GABRG2_E" [color="0 0 0.4935668789808917", label="1.76",style=bold];
"KRT1_S" -> "ABCA13_S" [color="0 0 0.4929936305732484", label="1.77",style=bold];
"DPP6_S" -> "DPP6_E" [color="0 0 0.4924203821656051", label="1.77",style=bold];
"KRT78_S" -> "TRPM3_E" [color="0 0 0.49184713375796174", label="1.77",style=bold];
"IVL_S" -> "AKNAD1_E" [color="0 0 0.49127388535031846", label="1.77",style=bold];
"KRT78_S" -> "LRRC7_E" [color="0 0 0.49070063694267513", label="1.78",style=bold];
"KRT78_S" -> "NELL1_E" [color="0 0 0.4901273885350318", label="1.78",style=bold];
"SOX1_S" -> "MUC4_M" [color="0 0 0.4895541401273885", label="1.78",style=bold];
"CA4_S" -> "ABCA13_S" [color="0 0 0.4889808917197452", label="1.79",style=bold];
"UNC5D_S" -> "HTR4_M" [color="0 0 0.4884076433121019", label="1.79",style=bold];
"NDST4_S" -> "CSMD1_S" [color="0 0 0.48783439490445857", label="1.80",style=bold];
"NELL1_S" -> "PCDHA13_E" [color="0 0 0.48726114649681523", label="1.80",style=bold];
"A2ML1_S" -> "NELL1_E" [color="0 0 0.48668789808917196", label="1.80",style=bold];
"CHRNA4_S" -> "RIMS1_E" [color="0 0 0.4861146496815286", label="1.81",style=bold];
"IRX4_S" -> "CHRNA4_E" [color="0 0 0.48554140127388534", label="1.81",style=bold];
"CRCT1_S" -> "CNTNAP4_S" [color="0 0 0.484968152866242", label="1.82",style=bold];
"PGC_S" -> "ABCA13_M" [color="0 0 0.4843949044585987", label="1.82",style=bold];
"LOC100190940_S" -> "LMOD2_E" [color="0 0 0.4838216560509554", label="1.82",style=bold];
"KRT78_S" -> "ABCA13_M" [color="0 0 0.48324840764331206", label="1.82",style=bold];
"KRT1_S" -> "APOB_E" [color="0 0 0.4826751592356688", label="1.82",style=bold];
"KRT78_S" -> "RPL10L_S" [color="0 0 0.48210191082802545", label="1.82",style=bold];
"TRPM3_S" -> "ADAMTS20_S" [color="0 0 0.4815286624203821", label="1.82",style=bold];
"DSG1_S" -> "PNLDC1_E" [color="0 0 0.48095541401273884", label="1.83",style=bold];
"CRCT1_S" -> "UNC5D_S" [color="0 0 0.4803821656050955", label="1.83",style=bold];
"SOX1_S" -> "BNC1_S" [color="0 0 0.4798089171974522", label="1.83",style=bold];
"KRT24_S" -> "LIPF_E" [color="0 0 0.4792356687898089", label="1.84",style=bold];
"IRX4_S" -> "CSMD1_S" [color="0 0 0.47866242038216555", label="1.84",style=bold];
"HTR4_S" -> "KCNA10_E" [color="0 0 0.4780891719745223", label="1.85",style=bold];
"LOC100190940_S" -> "TRIML2_M" [color="0 0 0.47751592356687894", label="1.85",style=bold];
"SERPINB13_S" -> "TPTE_M" [color="0 0 0.47694267515923566", label="1.86",style=bold];
"SLC12A1_S" -> "KCNA10_M" [color="0 0 0.47636942675159233", label="1.86",style=bold];
"PCDHA13_S" -> "ZAN_E" [color="0 0 0.475796178343949", label="1.86",style=bold];
"ZNF716_S" -> "ABCA13_S" [color="0 0 0.4752229299363057", label="1.87",style=bold];
"RIMS1_S" -> "MYBPC1_S" [color="0 0 0.4746496815286624", label="1.88",style=bold];
"PGC_S" -> "GABRG2_M" [color="0 0 0.4740764331210191", label="1.88",style=bold];
"LOC100190940_S" -> "CA4_M" [color="0 0 0.47350318471337577", label="1.89",style=bold];
"KRT78_S" -> "MYBPC1_S" [color="0 0 0.47292993630573243", label="1.89",style=bold];
"SLC1A6_S" -> "RPL10L_S" [color="0 0 0.47235668789808916", label="1.90",style=bold];
"KRT78_S" -> "CNTNAP5_S" [color="0 0 0.4717834394904458", label="1.90",style=bold];
"KRT24_S" -> "DMBT1_E" [color="0 0 0.47121019108280254", label="1.91",style=bold];
"LIPF_S" -> "KRT1_M" [color="0 0 0.4706369426751592", label="1.91",style=bold];
"SERPINB13_S" -> "NDST4_M" [color="0 0 0.4700636942675159", label="1.91",style=bold];
"PGC_S" -> "IVL_M" [color="0 0 0.4694904458598726", label="1.91",style=bold];
"MUC2_S" -> "MUC17_S" [color="0 0 0.46891719745222926", label="1.92",style=bold];
"KPRP_S" -> "AKNAD1_E" [color="0 0 0.468343949044586", label="1.92",style=bold];
"KRT1_S" -> "LHCGR_S" [color="0 0 0.46777070063694265", label="1.93",style=bold];
"LOC100190940_S" -> "CFHR1_M" [color="0 0 0.4671974522292993", label="1.93",style=bold];
"LIPF_S" -> "PGC_M" [color="0 0 0.46662420382165604", label="1.93",style=bold];
"PGC_S" -> "FBN3_E" [color="0 0 0.4660509554140127", label="1.94",style=bold];
"PNLDC1_S" -> "DPP6_E" [color="0 0 0.4654777070063694", label="1.94",style=bold];
"PNLIPRP3_S" -> "GPR98_S" [color="0 0 0.4649044585987261", label="1.94",style=bold];
"CFHR1_S" -> "MUC6_M" [color="0 0 0.46433121019108275", label="1.94",style=bold];
"SERPINB13_S" -> "CFHR1_M" [color="0 0 0.4637579617834395", label="1.95",style=bold];
"ADAMTS20_S" -> "FBN3_E" [color="0 0 0.46318471337579614", label="1.95",style=bold];
"PGC_S" -> "KRT6C_M" [color="0 0 0.46261146496815286", label="1.95",style=bold];
"APOB_S" -> "ZAN_S" [color="0 0 0.46203821656050953", label="1.95",style=bold];
"DUSP27_S" -> "GABRG2_S" [color="0 0 0.4614649681528662", label="1.96",style=bold];
"PGC_S" -> "LRRC7_E" [color="0 0 0.4608917197452229", label="1.97",style=bold];
"SOX1_S" -> "MYBPC1_M" [color="0 0 0.4603184713375796", label="1.97",style=bold];
"LOC100190940_S" -> "CSMD3_S" [color="0 0 0.45974522292993625", label="1.97",style=bold];
"TRIML2_S" -> "LIN28B_E" [color="0 0 0.45917197452229297", label="1.98",style=bold];
"SERPINB11_S" -> "APOB_S" [color="0 0 0.45859872611464964", label="1.98",style=bold];
"CRCT1_S" -> "ZIC4_S" [color="0 0 0.45802547770700636", label="1.98",style=bold];
"KRT24_S" -> "A2ML1_M" [color="0 0 0.457452229299363", label="1.99",style=bold];
"KRT24_S" -> "LRRIQ1_S" [color="0 0 0.4568789808917197", label="2.00",style=bold];
"DMBT1_S" -> "CSMD1_S" [color="0 0 0.4563057324840764", label="2.00",style=bold];
"SLC1A6_S" -> "CFHR1_M" [color="0 0 0.4557324840764331", label="2.00",style=bold];
"ZIC4_S" -> "MUC16_S" [color="0 0 0.4551592356687898", label="2.00",style=bold];
"KRT1_S" -> "SERPINB11_M" [color="0 0 0.45458598726114646", label="2.01",style=bold];
"GABRA1_S" -> "FLG_S" [color="0 0 0.45401273885350313", label="2.01",style=bold];
"TMPRSS11A_S" -> "LOC100190940_E" [color="0 0 0.45343949044585985", label="2.01",style=bold];
"KRT24_S" -> "MUC2_S" [color="0 0 0.4528662420382165", label="2.02",style=bold];
"KRT78_S" -> "CHRNA4_E" [color="0 0 0.45229299363057324", label="2.02",style=bold];
"RPL10L_S" -> "RIMS1_E" [color="0 0 0.4517197452229299", label="2.03",style=bold];
"GABRA1_M" -> "SPAG17_M" [color="0 0 0.45114649681528657", label="2.03",style=bold];
"SERPINB13_S" -> "PCDHA13_E" [color="0 0 0.4505732484076433", label="2.05",style=bold];
"SERPINB13_S" -> "LRRIQ1_E" [color="0 0 0.44999999999999996", label="2.05",style=bold];
"KPRP_S" -> "UNC5D_E" [color="0 0 0.4494267515923567", label="2.05",style=bold];
"KRT78_S" -> "PAX1_S" [color="0 0 0.44885350318471334", label="2.06",style=bold];
"CFHR1_S" -> "LRRC7_E" [color="0 0 0.44828025477707", label="2.06",style=bold];
"TMPRSS11A_S" -> "CSMD1_S" [color="0 0 0.44770700636942673", label="2.06",style=bold];
"GABRA1_S" -> "TMPRSS11A_M" [color="0 0 0.4471337579617834", label="2.06",style=bold];
"HTR4_S" -> "ZAN_E" [color="0 0 0.4465605095541401", label="2.06",style=bold];
"HTR4_S" -> "TRPM3_M" [color="0 0 0.4459872611464968", label="2.06",style=bold];
"DUSP27_S" -> "MUC5B_S" [color="0 0 0.44541401273885345", label="2.07",style=bold];
"MYO18B_S" -> "LIPF_M" [color="0 0 0.44484076433121017", label="2.08",style=bold];
"SYT16_S" -> "SPAG17_E" [color="0 0 0.44426751592356684", label="2.08",style=bold];
"TMPRSS11A_S" -> "MUC6_E" [color="0 0 0.44369426751592356", label="2.08",style=bold];
"SERPINB11_S" -> "SLC1A6_E" [color="0 0 0.4431210191082802", label="2.09",style=bold];
"PNLDC1_S" -> "PGC_E" [color="0 0 0.4425477707006369", label="2.09",style=bold];
"GABRA1_S" -> "DSG1_M" [color="0 0 0.4419745222929936", label="2.10",style=bold];
"UNC5D_S" -> "MUC16_S" [color="0 0 0.4414012738853503", label="2.11",style=bold];
"PAX1_S" -> "PGC_M" [color="0 0 0.440828025477707", label="2.11",style=bold];
"GABRG2_S" -> "LIPF_M" [color="0 0 0.44025477707006366", label="2.11",style=bold];
"PCDHA1_S" -> "LRRIQ1_S" [color="0 0 0.43968152866242033", label="2.12",style=bold];
"PAX1_S" -> "SLC12A1_M" [color="0 0 0.43910828025477705", label="2.12",style=bold];
"DCT_S" -> "LOC100190940_E" [color="0 0 0.4385350318471337", label="2.12",style=bold];
"CA4_S" -> "DSG1_M" [color="0 0 0.43796178343949044", label="2.12",style=bold];
"SERPINB13_S" -> "PCDHA1_E" [color="0 0 0.4373885350318471", label="2.12",style=bold];
"NELL1_S" -> "CA4_S" [color="0 0 0.43681528662420377", label="2.12",style=bold];
"PAX1_S" -> "LMOD2_M" [color="0 0 0.4362420382165605", label="2.13",style=bold];
"PGC_S" -> "ABCA13_S" [color="0 0 0.43566878980891716", label="2.14",style=bold];
"PCDHA1_S" -> "DMBT1_E" [color="0 0 0.4350955414012739", label="2.14",style=bold];
"MUC5B_S" -> "TRPM3_S" [color="0 0 0.43452229299363054", label="2.14",style=bold];
"A2ML1_S" -> "SPAG17_S" [color="0 0 0.4339490445859872", label="2.14",style=bold];
"DCT_S" -> "MUC16_S" [color="0 0 0.43337579617834393", label="2.15",style=bold];
"SOX1_S" -> "APOB_E" [color="0 0 0.4328025477707006", label="2.16",style=bold];
"LOC100190940_S" -> "KRT1_M" [color="0 0 0.4322292993630573", label="2.16",style=bold];
"SBSN_S" -> "LHCGR_E" [color="0 0 0.431656050955414", label="2.16",style=bold];
"ZNF716_S" -> "MUC2_S" [color="0 0 0.43108280254777065", label="2.16",style=bold];
"SOX1_S" -> "DMBT1_E" [color="0 0 0.43050955414012737", label="2.16",style=bold];
"KPRP_S" -> "FBN3_E" [color="0 0 0.42993630573248404", label="2.16",style=bold];
"SLC1A6_S" -> "CHRNA4_S" [color="0 0 0.42936305732484076", label="2.16",style=bold];
"CFHR1_S" -> "RIMS1_S" [color="0 0 0.4287898089171974", label="2.16",style=bold];
"MUC17_S" -> "UNC5D_S" [color="0 0 0.4282165605095541", label="2.16",style=bold];
"DSG1_S" -> "SYT16_E" [color="0 0 0.4276433121019108", label="2.17",style=bold];
"CNTNAP4_M" -> "LIN28B_M" [color="0 0 0.4270700636942675", label="2.17",style=bold];
"ZIC4_M" -> "SOX1_E" [color="0 0 0.42649681528662414", label="2.18",style=bold];
"KRT6C_S" -> "ZAN_S" [color="0 0 0.42592356687898086", label="2.18",style=bold];
"GABRA1_S" -> "NELL1_S" [color="0 0 0.42535031847133753", label="2.18",style=bold];
"PNLDC1_S" -> "A2ML1_M" [color="0 0 0.42477707006369425", label="2.18",style=bold];
"RIMS1_S" -> "DPP6_E" [color="0 0 0.4242038216560509", label="2.19",style=bold];
"SBSN_S" -> "FBN3_E" [color="0 0 0.4236305732484076", label="2.20",style=bold];
"SLC1A6_S" -> "LIPF_S" [color="0 0 0.4230573248407643", label="2.21",style=bold];
"ZIC4_S" -> "LIN28B_E" [color="0 0 0.42248407643312097", label="2.21",style=bold];
"ADH7_S" -> "UNC5D_M" [color="0 0 0.4219108280254777", label="2.21",style=bold];
"SERPINB11_S" -> "FBN3_E" [color="0 0 0.42133757961783436", label="2.21",style=bold];
"KCNA10_S" -> "ADAMTS19_S" [color="0 0 0.420764331210191", label="2.22",style=bold];
"LOC100190940_S" -> "MUC2_E" [color="0 0 0.42019108280254774", label="2.22",style=bold];
"KRT6C_S" -> "PCDHA1_E" [color="0 0 0.4196178343949044", label="2.22",style=bold];
"TMPRSS11A_S" -> "MUC4_E" [color="0 0 0.41904458598726113", label="2.23",style=bold];
"RPL10L_S" -> "DMBT1_M" [color="0 0 0.4184713375796178", label="2.23",style=bold];
"KRT78_S" -> "ZIC4_M" [color="0 0 0.41789808917197446", label="2.24",style=bold];
"LOC100190940_S" -> "ABCA13_E" [color="0 0 0.4173248407643312", label="2.24",style=bold];
"APOB_S" -> "FOXG1_S" [color="0 0 0.41675159235668785", label="2.24",style=bold];
"IRX4_S" -> "AKNAD1_M" [color="0 0 0.41617834394904457", label="2.25",style=bold];
"DCT_S" -> "FLG_S" [color="0 0 0.41560509554140124", label="2.25",style=bold];
"ADAMTS20_S" -> "SYT16_S" [color="0 0 0.4150318471337579", label="2.26",style=bold];
"ADH7_S" -> "FBN3_M" [color="0 0 0.4144585987261146", label="2.27",style=bold];
"MUC5B_S" -> "A2ML1_M" [color="0 0 0.4138853503184713", label="2.27",style=bold];
"IRX4_S" -> "LHCGR_E" [color="0 0 0.413312101910828", label="2.27",style=bold];
"DPP6_S" -> "FREM2_S" [color="0 0 0.4127388535031847", label="2.30",style=bold];
"KPRP_S" -> "DSG1_M" [color="0 0 0.41216560509554134", label="2.30",style=bold];
"TPTE_S" -> "MUC4_E" [color="0 0 0.41159235668789806", label="2.31",style=bold];
"LIN28B_S" -> "TRIML2_S" [color="0 0 0.41101910828025473", label="2.31",style=bold];
"DSG1_S" -> "NELL1_E" [color="0 0 0.41044585987261145", label="2.32",style=bold];
"LHCGR_S" -> "DMBT1_M" [color="0 0 0.4098726114649681", label="2.32",style=bold];
"ADH7_S" -> "NELL1_E" [color="0 0 0.4092993630573248", label="2.32",style=bold];
"SBSN_S" -> "IRX4_E" [color="0 0 0.4087261146496815", label="2.33",style=bold];
"AKNAD1_S" -> "A2ML1_S" [color="0 0 0.40815286624203817", label="2.33",style=bold];
"TMPRSS11A_S" -> "ADH7_M" [color="0 0 0.4075796178343949", label="2.34",style=bold];
"IRX4_S" -> "DCT_E" [color="0 0 0.40700636942675156", label="2.34",style=bold];
"KCNV1_S" -> "MUC17_S" [color="0 0 0.4064331210191082", label="2.34",style=bold];
"LMOD2_S" -> "MYO18B_E" [color="0 0 0.40585987261146494", label="2.36",style=bold];
"ADAMTS19_S" -> "CSMD1_E" [color="0 0 0.4052866242038216", label="2.36",style=bold];
"LOC100190940_S" -> "CNTNAP4_M" [color="0 0 0.40471337579617833", label="2.37",style=bold];
"SLC1A6_S" -> "MUC4_E" [color="0 0 0.404140127388535", label="2.37",style=bold];
"RIMS1_S" -> "FREM2_S" [color="0 0 0.40356687898089166", label="2.37",style=bold];
"CHRNA4_S" -> "CSMD1_E" [color="0 0 0.4029936305732484", label="2.38",style=bold];
"ADAMTS20_S" -> "KRT6C_M" [color="0 0 0.40242038216560505", label="2.38",style=bold];
"LIPF_S" -> "MUC6_E" [color="0 0 0.40184713375796177", label="2.38",style=bold];
"IRX4_S" -> "KRT6C_M" [color="0 0 0.40127388535031844", label="2.39",style=bold];
"TMPRSS11A_S" -> "NELL1_E" [color="0 0 0.4007006369426751", label="2.39",style=bold];
"PCDHA1_S" -> "FOXG1_S" [color="0 0 0.4001273885350318", label="2.39",style=bold];
"PNLIPRP3_S" -> "FREM2_M" [color="0 0 0.39955414012738855", label="2.40",style=bold];
"TRIML2_S" -> "PGC_E" [color="0 0 0.39898089171974516", label="2.40",style=bold];
"DPP6_S" -> "FBN3_E" [color="0 0 0.3984076433121019", label="2.40",style=bold];
"TMPRSS11A_S" -> "PGC_E" [color="0 0 0.3978343949044586", label="2.40",style=bold];
"PGC_S" -> "NELL1_S" [color="0 0 0.3972611464968152", label="2.40",style=bold];
"TMPRSS11A_S" -> "MUC4_S" [color="0 0 0.39668789808917193", label="2.40",style=bold];
"CWH43_S" -> "CFHR1_M" [color="0 0 0.39611464968152865", label="2.40",style=bold];
"KPRP_S" -> "APOB_S" [color="0 0 0.39554140127388526", label="2.40",style=bold];
"MYBPC1_S" -> "PAX1_E" [color="0 0 0.394968152866242", label="2.40",style=bold];
"KRT1_S" -> "LHCGR_E" [color="0 0 0.3943949044585987", label="2.41",style=bold];
"BNC1_S" -> "CNTNAP5_S" [color="0 0 0.3938216560509554", label="2.42",style=bold];
"HTR4_S" -> "MYBPC1_E" [color="0 0 0.39324840764331204", label="2.42",style=bold];
"PGC_S" -> "SPAG17_E" [color="0 0 0.39267515923566876", label="2.43",style=bold];
"LIN28B_S" -> "LOC100190940_S" [color="0 0 0.3921019108280255", label="2.43",style=bold];
"SERPINB13_S" -> "GABRG2_M" [color="0 0 0.3915286624203821", label="2.44",style=bold];
"LMOD2_S" -> "TRPM3_S" [color="0 0 0.3909554140127388", label="2.44",style=bold];
"SERPINB13_S" -> "KRT1_E" [color="0 0 0.39038216560509553", label="2.45",style=bold];
"DCT_S" -> "DUSP27_E" [color="0 0 0.38980891719745214", label="2.46",style=bold];
"NDST4_S" -> "MUC6_S" [color="0 0 0.38923566878980886", label="2.46",style=bold];
"TMPRSS11A_S" -> "A2ML1_M" [color="0 0 0.3886624203821656", label="2.46",style=bold];
"BNC1_S" -> "GABRG2_E" [color="0 0 0.3880891719745223", label="2.46",style=bold];
"LOC100190940_S" -> "TMPRSS11A_M" [color="0 0 0.3875159235668789", label="2.47",style=bold];
"PNLDC1_S" -> "CNTNAP5_M" [color="0 0 0.38694267515923564", label="2.48",style=bold];
"CHRNA4_S" -> "ADH7_M" [color="0 0 0.38636942675159236", label="2.48",style=bold];
"KRT6C_S" -> "PNLDC1_E" [color="0 0 0.38579617834394897", label="2.48",style=bold];
"LIN28B_S" -> "ABCA13_S" [color="0 0 0.3852229299363057", label="2.50",style=bold];
"RPL10L_S" -> "LIPF_M" [color="0 0 0.3846496815286624", label="2.51",style=bold];
"ZIC4_M" -> "CA4_M" [color="0 0 0.384076433121019", label="2.52",style=bold];
"DMBT1_S" -> "APOB_E" [color="0 0 0.38350318471337574", label="2.53",style=bold];
"ADAMTS19_S" -> "GPR98_S" [color="0 0 0.38292993630573247", label="2.53",style=bold];
"ADAMTS19_S" -> "FREM2_E" [color="0 0 0.3823566878980892", label="2.53",style=bold];
"SYT16_S" -> "SERPINB13_M" [color="0 0 0.3817834394904458", label="2.53",style=bold];
"MUC6_S" -> "ZIC4_S" [color="0 0 0.3812101910828025", label="2.54",style=bold];
"LIN28B_S" -> "KRT1_S" [color="0 0 0.38063694267515924", label="2.56",style=bold];
"LMOD2_S" -> "DUSP27_E" [color="0 0 0.38006369426751585", label="2.56",style=bold];
"SYT16_S" -> "KCNA10_M" [color="0 0 0.37949044585987257", label="2.57",style=bold];
"KRT24_S" -> "MUC4_E" [color="0 0 0.3789171974522293", label="2.59",style=bold];
"CRCT1_S" -> "TMPRSS11A_S" [color="0 0 0.3783439490445859", label="2.59",style=bold];
"BNC1_S" -> "MUC2_M" [color="0 0 0.3777707006369426", label="2.60",style=bold];
"CRCT1_E" -> "SERPINB11_E" [color="0 0 0.37719745222929935", label="2.60",style=bold];
"NELL1_S" -> "ABCA13_E" [color="0 0 0.37662420382165596", label="2.61",style=bold];
"MYO18B_S" -> "GPR98_S" [color="0 0 0.3760509554140127", label="2.62",style=bold];
"LIN28B_S" -> "KPRP_M" [color="0 0 0.3754777070063694", label="2.63",style=bold];
"CFHR1_S" -> "CSMD1_S" [color="0 0 0.3749044585987261", label="2.63",style=bold];
"SERPINB13_S" -> "TMPRSS11A_M" [color="0 0 0.37433121019108273", label="2.64",style=bold];
"MUC2_S" -> "CFHR1_M" [color="0 0 0.37375796178343945", label="2.64",style=bold];
"KRT24_S" -> "GABRA1_S" [color="0 0 0.3731847133757962", label="2.66",style=bold];
"TMPRSS11A_S" -> "AKNAD1_E" [color="0 0 0.3726114649681528", label="2.67",style=bold];
"FBN3_S" -> "APOB_S" [color="0 0 0.3720382165605095", label="2.69",style=bold];
"KPRP_S" -> "Tumor_Status" [color="0 0 0.3714649681528662", label="2.69",style=bold];
"ADAMTS19_S" -> "MUC6_M" [color="0 0 0.37089171974522284", label="2.70",style=bold];
"KRT78_S" -> "MUC17_S" [color="0 0 0.37031847133757956", label="2.70",style=bold];
"PNLIPRP3_S" -> "KRT6C_M" [color="0 0 0.3697452229299363", label="2.72",style=bold];
"CFHR1_S" -> "LIPF_E" [color="0 0 0.369171974522293", label="2.73",style=bold];
"TRIML2_S" -> "CSMD1_S" [color="0 0 0.3685987261146496", label="2.73",style=bold];
"CRCT1_S" -> "DCT_E" [color="0 0 0.36802547770700633", label="2.74",style=bold];
"SYT16_S" -> "GABRA1_E" [color="0 0 0.36745222929936305", label="2.75",style=bold];
"CNTNAP4_S" -> "Tumor_Status" [color="0 0 0.36687898089171966", label="2.76",style=bold];
"DUSP27_S" -> "MYO18B_S" [color="0 0 0.3663057324840764", label="2.76",style=bold];
"LIN28B_S" -> "MUC2_S" [color="0 0 0.3657324840764331", label="2.77",style=bold];
"FOXG1_S" -> "DMBT1_M" [color="0 0 0.3651592356687897", label="2.77",style=bold];
"BNC1_S" -> "NELL1_M" [color="0 0 0.36458598726114644", label="2.77",style=bold];
"ABCA13_E" -> "MUC4_S" [color="0 0 0.36401273885350316", label="2.77",style=bold];
"TPTE_S" -> "LRRIQ1_E" [color="0 0 0.3634394904458599", label="2.77",style=bold];
"PGC_S" -> "PCDHA13_M" [color="0 0 0.3628662420382165", label="2.77",style=bold];
"FBN3_S" -> "KRT6C_S" [color="0 0 0.3622929936305732", label="2.78",style=bold];
"TRIML2_S" -> "IRX4_S" [color="0 0 0.36171974522292993", label="2.78",style=bold];
"KCNV1_S" -> "IRX4_S" [color="0 0 0.36114649681528654", label="2.79",style=bold];
"LMOD2_S" -> "NELL1_S" [color="0 0 0.36057324840764327", label="2.79",style=bold];
"TRIML2_S" -> "CSMD3_S" [color="0 0 0.36", label="2.80",style=bold];
"DUSP27_S" -> "MUC6_M" [color="0 0 0.3594267515923566", label="2.80",style=bold];
"KCNV1_S" -> "MUC5B_E" [color="0 0 0.3588535031847133", label="2.81",style=bold];
"AKNAD1_S" -> "DUSP27_S" [color="0 0 0.35828025477707004", label="2.81",style=bold];
"LMOD2_S" -> "SPRR2A_E" [color="0 0 0.35770700636942676", label="2.81",style=bold];
"IRX4_S" -> "MYBPC1_E" [color="0 0 0.35713375796178337", label="2.82",style=bold];
"DCT_S" -> "SLC12A1_S" [color="0 0 0.3565605095541401", label="2.85",style=bold];
"CHRNA4_S" -> "KRT6C_M" [color="0 0 0.3559872611464968", label="2.87",style=bold];
"MYBPC1_S" -> "CA4_E" [color="0 0 0.3554140127388534", label="2.88",style=bold];
"SPAG17_S" -> "DCT_S" [color="0 0 0.35484076433121015", label="2.89",style=bold];
"LRRC7_S" -> "HTR4_E" [color="0 0 0.35426751592356687", label="2.90",style=bold];
"KPRP_S" -> "NDST4_M" [color="0 0 0.3536942675159235", label="2.90",style=bold];
"ZNF716_S" -> "PNLIPRP3_S" [color="0 0 0.3531210191082802", label="2.91",style=bold];
"MYBPC1_S" -> "LRRIQ1_E" [color="0 0 0.3525477707006369", label="2.91",style=bold];
"PNLDC1_S" -> "DCT_E" [color="0 0 0.35197452229299364", label="2.91",style=bold];
"MUC4_S" -> "MYO18B_S" [color="0 0 0.35140127388535025", label="2.91",style=bold];
"SOX1_S" -> "PCDHA13_E" [color="0 0 0.350828025477707", label="2.91",style=bold];
"KRT78_S" -> "LRRIQ1_E" [color="0 0 0.3502547770700637", label="2.93",style=bold];
"CNTNAP5_S" -> "AKNAD1_M" [color="0 0 0.3496815286624203", label="2.94",style=bold];
"A2ML1_S" -> "TRPM3_S" [color="0 0 0.349108280254777", label="2.94",style=bold];
"ZNF716_E" -> "GABRA1_E" [color="0 0 0.34853503184713375", label="2.94",style=bold];
"CNTNAP4_S" -> "LMOD2_M" [color="0 0 0.34796178343949036", label="2.96",style=bold];
"NDST4_S" -> "BNC1_S" [color="0 0 0.3473885350318471", label="2.97",style=bold];
"CHRNA4_S" -> "KRT24_E" [color="0 0 0.3468152866242038", label="2.98",style=bold];
"CNTNAP4_M" -> "APOB_M" [color="0 0 0.3462420382165605", label="2.98",style=bold];
"SLC1A6_S" -> "MUC16_S" [color="0 0 0.34566878980891713", label="2.99",style=bold];
"KRT24_S" -> "HTR4_M" [color="0 0 0.34509554140127385", label="3.00",style=bold];
"TRPM3_S" -> "FLG_S" [color="0 0 0.3445222929936306", label="3.03",style=bold];
"KRT6C_E" -> "IVL_M" [color="0 0 0.3439490445859872", label="3.04",style=bold];
"RPL10L_S" -> "KCNA10_E" [color="0 0 0.3433757961783439", label="3.04",style=bold];
"FBN3_S" -> "RPL10L_S" [color="0 0 0.3428025477707006", label="3.04",style=bold];
"KCNV1_S" -> "MUC2_S" [color="0 0 0.34222929936305724", label="3.05",style=bold];
"KPRP_S" -> "LIPF_E" [color="0 0 0.34165605095541396", label="3.05",style=bold];
"KPRP_S" -> "ZAN_S" [color="0 0 0.3410828025477707", label="3.05",style=bold];
"PNLIPRP3_S" -> "DUSP27_M" [color="0 0 0.3405095541401273", label="3.05",style=bold];
"DUSP27_S" -> "MYO18B_M" [color="0 0 0.339936305732484", label="3.05",style=bold];
"KRT6C_S" -> "DSG1_E" [color="0 0 0.33936305732484073", label="3.06",style=bold];
"CRCT1_S" -> "KCNA10_M" [color="0 0 0.33878980891719745", label="3.06",style=bold];
"NELL1_S" -> "ABCA13_S" [color="0 0 0.33821656050955407", label="3.06",style=bold];
"SLC12A1_S" -> "ADAMTS19_E" [color="0 0 0.3376433121019108", label="3.06",style=bold];
"PCDHA13_S" -> "ADH7_M" [color="0 0 0.3370700636942675", label="3.07",style=bold];
"SERPINB13_S" -> "CSMD3_S" [color="0 0 0.3364968152866241", label="3.08",style=bold];
"CNTNAP4_S" -> "NELL1_S" [color="0 0 0.33592356687898084", label="3.09",style=bold];
"DCT_S" -> "APOB_M" [color="0 0 0.33535031847133756", label="3.09",style=bold];
"LIN28B_S" -> "MUC2_M" [color="0 0 0.33477707006369417", label="3.10",style=bold];
"PNLIPRP3_S" -> "MUC5B_M" [color="0 0 0.3342038216560509", label="3.10",style=bold];
"KPRP_S" -> "NELL1_S" [color="0 0 0.3336305732484076", label="3.11",style=bold];
"KRT6C_S" -> "MUC6_E" [color="0 0 0.33305732484076433", label="3.12",style=bold];
"ZAN_S" -> "ABCA13_S" [color="0 0 0.33248407643312095", label="3.13",style=bold];
"ADH7_S" -> "ZAN_M" [color="0 0 0.33191082802547767", label="3.13",style=bold];
"CNTNAP4_S" -> "IVL_S" [color="0 0 0.3313375796178344", label="3.14",style=bold];
"KCNV1_S" -> "CNTNAP4_E" [color="0 0 0.330764331210191", label="3.15",style=bold];
"BNC1_M" -> "ADAMTS19_M" [color="0 0 0.3301910828025477", label="3.16",style=bold];
"TRIML2_S" -> "KCNA10_E" [color="0 0 0.32961783439490444", label="3.17",style=bold];
"DPP6_S" -> "FBN3_S" [color="0 0 0.32904458598726105", label="3.18",style=bold];
"KRT6C_S" -> "PGC_E" [color="0 0 0.3284713375796178", label="3.21",style=bold];
"MYO18B_S" -> "DSG1_S" [color="0 0 0.3278980891719745", label="3.21",style=bold];
"HTR4_S" -> "TPTE_E" [color="0 0 0.3273248407643312", label="3.21",style=bold];
"FBN3_S" -> "MUC4_M" [color="0 0 0.3267515923566878", label="3.22",style=bold];
"DUSP27_S" -> "RPL10L_M" [color="0 0 0.32617834394904455", label="3.23",style=bold];
"PAX1_S" -> "KRT1_M" [color="0 0 0.32560509554140127", label="3.23",style=bold];
"CHRNA4_S" -> "FREM2_E" [color="0 0 0.3250318471337579", label="3.24",style=bold];
"SERPINB13_S" -> "DCT_E" [color="0 0 0.3244585987261146", label="3.25",style=bold];
"CRCT1_S" -> "PCDHA1_S" [color="0 0 0.3238853503184713", label="3.26",style=bold];
"FLG_E" -> "PGC_M" [color="0 0 0.32331210191082793", label="3.26",style=bold];
"SBSN_S" -> "PCDHA1_S" [color="0 0 0.32273885350318465", label="3.28",style=bold];
"PNLIPRP3_S" -> "TPTE_E" [color="0 0 0.3221656050955414", label="3.29",style=bold];
"MUC2_S" -> "APOB_M" [color="0 0 0.3215923566878981", label="3.32",style=bold];
"LOC100190940_S" -> "BNC1_S" [color="0 0 0.3210191082802547", label="3.32",style=bold];
"KCNV1_S" -> "FLG_S" [color="0 0 0.3204458598726114", label="3.32",style=bold];
"AKNAD1_S" -> "PCDHA1_S" [color="0 0 0.31987261146496815", label="3.32",style=bold];
"KRT78_S" -> "KCNA10_M" [color="0 0 0.31929936305732476", label="3.35",style=bold];
"LIN28B_E" -> "RPL10L_E" [color="0 0 0.3187261146496815", label="3.35",style=bold];
"KPRP_S" -> "PCDHA1_S" [color="0 0 0.3181528662420382", label="3.37",style=bold];
"SLC1A6_E" -> "DPP6_E" [color="0 0 0.3175796178343948", label="3.38",style=bold];
"CSMD1_S" -> "DUSP27_S" [color="0 0 0.31700636942675153", label="3.38",style=bold];
"ADH7_S" -> "CNTNAP4_E" [color="0 0 0.31643312101910825", label="3.40",style=bold];
"PNLIPRP3_S" -> "LIPF_M" [color="0 0 0.315859872611465", label="3.40",style=bold];
"FBN3_S" -> "DMBT1_E" [color="0 0 0.3152866242038216", label="3.41",style=bold];
"IVL_S" -> "RIMS1_E" [color="0 0 0.3147133757961783", label="3.43",style=bold];
"PAX1_S" -> "SYT16_M" [color="0 0 0.31414012738853503", label="3.44",style=bold];
"UNC5D_S" -> "KRT6C_M" [color="0 0 0.31356687898089164", label="3.44",style=bold];
"A2ML1_S" -> "LRRIQ1_E" [color="0 0 0.31299363057324836", label="3.44",style=bold];
"RIMS1_S" -> "IVL_S" [color="0 0 0.3124203821656051", label="3.44",style=bold];
"KRT1_M" -> "DCT_M" [color="0 0 0.3118471337579617", label="3.44",style=bold];
"ZAN_E" -> "CSMD3_E" [color="0 0 0.3112738853503184", label="3.46",style=bold];
"MUC2_M" -> "KRT78_M" [color="0 0 0.31070063694267513", label="3.47",style=bold];
"DMBT1_S" -> "DMBT1_M" [color="0 0 0.31012738853503174", label="3.49",style=bold];
"KCNV1_S" -> "SBSN_S" [color="0 0 0.30955414012738847", label="3.49",style=bold];
"APOB_S" -> "Tumor_Status" [color="0 0 0.3089808917197452", label="3.50",style=bold];
"NELL1_S" -> "LRRIQ1_E" [color="0 0 0.3084076433121019", label="3.50",style=bold];
"BNC1_M" -> "MUC6_S" [color="0 0 0.3078343949044585", label="3.50",style=bold];
"ADAMTS19_S" -> "RIMS1_E" [color="0 0 0.30726114649681524", label="3.51",style=bold];
"LRRIQ1_S" -> "RIMS1_E" [color="0 0 0.30668789808917196", label="3.52",style=bold];
"KPRP_S" -> "TMPRSS11A_S" [color="0 0 0.3061146496815286", label="3.52",style=bold];
"IVL_S" -> "APOB_E" [color="0 0 0.3055414012738853", label="3.53",style=bold];
"MYBPC1_S" -> "DPP6_E" [color="0 0 0.304968152866242", label="3.53",style=bold];
"SOX1_S" -> "NDST4_M" [color="0 0 0.3043949044585986", label="3.54",style=bold];
"SPAG17_S" -> "FLG_S" [color="0 0 0.30382165605095535", label="3.55",style=bold];
"SERPINB11_S" -> "GABRG2_E" [color="0 0 0.30324840764331207", label="3.55",style=bold];
"KRT1_S" -> "CNTNAP4_S" [color="0 0 0.3026751592356688", label="3.55",style=bold];
"SERPINB11_S" -> "PNLDC1_E" [color="0 0 0.3021019108280254", label="3.55",style=bold];
"IRX4_S" -> "FLG_S" [color="0 0 0.3015286624203821", label="3.56",style=bold];
"DMBT1_S" -> "AKNAD1_M" [color="0 0 0.30095541401273884", label="3.56",style=bold];
"SPAG17_S" -> "MYBPC1_E" [color="0 0 0.30038216560509545", label="3.56",style=bold];
"PNLDC1_S" -> "HTR4_E" [color="0 0 0.2998089171974522", label="3.57",style=bold];
"KCNV1_S" -> "CFHR1_E" [color="0 0 0.2992356687898089", label="3.58",style=bold];
"HTR4_S" -> "LRRC7_S" [color="0 0 0.2986624203821655", label="3.59",style=bold];
"KPRP_S" -> "SYT16_S" [color="0 0 0.2980891719745222", label="3.59",style=bold];
"FBN3_S" -> "MUC6_E" [color="0 0 0.29751592356687895", label="3.59",style=bold];
"BNC1_S" -> "ADAMTS19_E" [color="0 0 0.29694267515923567", label="3.61",style=bold];
"AKNAD1_S" -> "DUSP27_E" [color="0 0 0.2963694267515923", label="3.61",style=bold];
"IVL_S" -> "TPTE_E" [color="0 0 0.295796178343949", label="3.62",style=bold];
"TRIML2_M" -> "PNLDC1_M" [color="0 0 0.2952229299363057", label="3.62",style=bold];
"RIMS1_S" -> "KRT6C_M" [color="0 0 0.29464968152866233", label="3.64",style=bold];
"PGC_S" -> "ADAMTS19_E" [color="0 0 0.29407643312101905", label="3.65",style=bold];
"NDST4_S" -> "PGC_E" [color="0 0 0.2935031847133758", label="3.69",style=bold];
"A2ML1_S" -> "MUC16_S" [color="0 0 0.2929299363057324", label="3.70",style=bold];
"GABRA1_S" -> "BNC1_S" [color="0 0 0.2923566878980891", label="3.70",style=bold];
"CFHR1_S" -> "MUC2_S" [color="0 0 0.29178343949044583", label="3.71",style=bold];
"LOC100190940_S" -> "FREM2_S" [color="0 0 0.29121019108280255", label="3.71",style=bold];
"DCT_S" -> "ADAMTS19_S" [color="0 0 0.29063694267515916", label="3.74",style=bold];
"TRPM3_S" -> "NELL1_E" [color="0 0 0.2900636942675159", label="3.75",style=bold];
"LRRIQ1_S" -> "FREM2_E" [color="0 0 0.2894904458598726", label="3.75",style=bold];
"CNTNAP5_S" -> "RIMS1_E" [color="0 0 0.2889171974522292", label="3.77",style=bold];
"FOXG1_S" -> "ZIC4_S" [color="0 0 0.28834394904458593", label="3.79",style=bold];
"TRPM3_S" -> "Tumor_Status" [color="0 0 0.28777070063694266", label="3.80",style=bold];
"SERPINB13_S" -> "AKNAD1_S" [color="0 0 0.28719745222929927", label="3.82",style=bold];
"ADH7_S" -> "TPTE_E" [color="0 0 0.286624203821656", label="3.83",style=bold];
"CRCT1_S" -> "LIPF_S" [color="0 0 0.2860509554140127", label="3.85",style=bold];
"CWH43_M" -> "PAX1_M" [color="0 0 0.28547770700636943", label="3.85",style=bold];
"CWH43_E" -> "CWH43_M" [color="0 0 0.28490445859872604", label="3.86",style=bold];
"NELL1_M" -> "PAX1_M" [color="0 0 0.28433121019108276", label="3.88",style=bold];
"DPP6_S" -> "APOB_E" [color="0 0 0.2837579617834395", label="3.89",style=bold];
"CWH43_S" -> "LRRIQ1_S" [color="0 0 0.2831847133757961", label="3.89",style=bold];
"GABRA1_S" -> "PCDHA1_E" [color="0 0 0.2826114649681528", label="3.94",style=bold];
"PNLIPRP3_S" -> "KRT24_M" [color="0 0 0.28203821656050954", label="3.94",style=bold];
"MUC5B_S" -> "ZIC4_E" [color="0 0 0.28146496815286615", label="3.95",style=bold];
"ADAMTS19_S" -> "PCDHA13_E" [color="0 0 0.28089171974522287", label="3.95",style=bold];
"DUSP27_S" -> "TRPM3_M" [color="0 0 0.2803184713375796", label="3.95",style=bold];
"IRX4_S" -> "KCNA10_M" [color="0 0 0.2797452229299362", label="3.97",style=bold];
"APOB_M" -> "NDST4_M" [color="0 0 0.2791719745222929", label="3.98",style=bold];
"LIN28B_S" -> "IRX4_S" [color="0 0 0.27859872611464964", label="4.00",style=bold];
"PCDHA13_S" -> "ZAN_S" [color="0 0 0.27802547770700636", label="4.00",style=bold];
"SOX1_S" -> "DMBT1_M" [color="0 0 0.277452229299363", label="4.01",style=bold];
"PGC_S" -> "PCDHA1_S" [color="0 0 0.2768789808917197", label="4.01",style=bold];
"SERPINB13_S" -> "KCNV1_M" [color="0 0 0.2763057324840764", label="4.02",style=bold];
"CWH43_S" -> "APOB_E" [color="0 0 0.275732484076433", label="4.03",style=bold];
"TRPM3_S" -> "KRT1_E" [color="0 0 0.27515923566878975", label="4.04",style=bold];
"KRT24_S" -> "UNC5D_S" [color="0 0 0.27458598726114647", label="4.06",style=bold];
"A2ML1_S" -> "CA4_S" [color="0 0 0.2740127388535031", label="4.07",style=bold];
"KRT1_M" -> "AKNAD1_M" [color="0 0 0.2734394904458598", label="4.08",style=bold];
"DPP6_S" -> "SERPINB13_M" [color="0 0 0.2728662420382165", label="4.12",style=bold];
"SLC1A6_S" -> "MUC17_S" [color="0 0 0.27229299363057324", label="4.14",style=bold];
"KRT24_S" -> "DCT_E" [color="0 0 0.27171974522292985", label="4.14",style=bold];
"PNLIPRP3_M" -> "SERPINB11_M" [color="0 0 0.2711464968152866", label="4.15",style=bold];
"MUC4_S" -> "SPAG17_M" [color="0 0 0.2705732484076433", label="4.16",style=bold];
"RIMS1_S" -> "SLC12A1_S" [color="0 0 0.2699999999999999", label="4.17",style=bold];
"GABRA1_S" -> "DMBT1_S" [color="0 0 0.26942675159235663", label="4.18",style=bold];
"CWH43_S" -> "DPP6_E" [color="0 0 0.26885350318471335", label="4.20",style=bold];
"LRRC7_S" -> "PCDHA1_M" [color="0 0 0.26828025477706996", label="4.20",style=bold];
"CWH43_S" -> "AKNAD1_E" [color="0 0 0.2677070063694267", label="4.23",style=bold];
"DCT_S" -> "RIMS1_S" [color="0 0 0.2671337579617834", label="4.26",style=bold];
"ADH7_S" -> "CSMD3_S" [color="0 0 0.2665605095541401", label="4.29",style=bold];
"SERPINB13_S" -> "SERPINB13_M" [color="0 0 0.26598726114649673", label="4.32",style=bold];
"PNLDC1_S" -> "CSMD1_S" [color="0 0 0.26541401273885346", label="4.33",style=bold];
"GABRA1_M" -> "MUC5B_S" [color="0 0 0.2648407643312102", label="4.33",style=bold];
"MUC4_S" -> "CHRNA4_E" [color="0 0 0.2642675159235668", label="4.34",style=bold];
"TRIML2_S" -> "MUC17_E" [color="0 0 0.2636942675159235", label="4.34",style=bold];
"CFHR1_S" -> "GABRA1_S" [color="0 0 0.26312101910828023", label="4.38",style=bold];
"ZIC4_S" -> "RPL10L_M" [color="0 0 0.26254777070063684", label="4.40",style=bold];
"MUC6_S" -> "ABCA13_S" [color="0 0 0.26197452229299356", label="4.44",style=bold];
"PCDHA13_S" -> "MYBPC1_E" [color="0 0 0.2614012738853503", label="4.46",style=bold];
"TRIML2_S" -> "ZAN_M" [color="0 0 0.260828025477707", label="4.46",style=bold];
"CNTNAP4_S" -> "MYBPC1_E" [color="0 0 0.2602547770700636", label="4.47",style=bold];
"KPRP_S" -> "MUC6_S" [color="0 0 0.25968152866242034", label="4.47",style=bold];
"DCT_S" -> "PCDHA13_E" [color="0 0 0.25910828025477706", label="4.50",style=bold];
"CA4_S" -> "FLG_S" [color="0 0 0.25853503184713367", label="4.51",style=bold];
"CSMD3_M" -> "LIN28B_M" [color="0 0 0.2579617834394904", label="4.51",style=bold];
"CWH43_M" -> "ADAMTS19_M" [color="0 0 0.2573885350318471", label="4.53",style=bold];
"SOX1_S" -> "LIPF_E" [color="0 0 0.2568152866242037", label="4.53",style=bold];
"FBN3_S" -> "PGC_E" [color="0 0 0.25624203821656044", label="4.57",style=bold];
"IVL_S" -> "ZAN_S" [color="0 0 0.25566878980891716", label="4.57",style=bold];
"RPL10L_S" -> "KCNA10_M" [color="0 0 0.2550955414012739", label="4.59",style=bold];
"SOX1_S" -> "ZAN_S" [color="0 0 0.2545222929936305", label="4.59",style=bold];
"TRIML2_S" -> "KRT1_E" [color="0 0 0.2539490445859872", label="4.59",style=bold];
"DCT_S" -> "MUC5B_S" [color="0 0 0.25337579617834394", label="4.60",style=bold];
"ADH7_M" -> "CA4_E" [color="0 0 0.25280254777070055", label="4.64",style=bold];
"KRT78_S" -> "APOB_E" [color="0 0 0.25222929936305727", label="4.67",style=bold];
"PNLIPRP3_E" -> "CSMD3_E" [color="0 0 0.251656050955414", label="4.68",style=bold];
"ZIC4_S" -> "TRIML2_M" [color="0 0 0.2510828025477706", label="4.68",style=bold];
"PNLIPRP3_S" -> "UNC5D_S" [color="0 0 0.2505095541401273", label="4.70",style=bold];
"SPAG17_S" -> "MUC5B_E" [color="0 0 0.24993630573248404", label="4.72",style=bold];
"TMPRSS11A_M" -> "PNLDC1_M" [color="0 0 0.24936305732484076", label="4.81",style=bold];
"CNTNAP5_S" -> "FREM2_E" [color="0 0 0.24878980891719737", label="4.82",style=bold];
"CA4_S" -> "TRPM3_S" [color="0 0 0.2482165605095541", label="4.82",style=bold];
"GPR98_S" -> "MUC16_S" [color="0 0 0.24764331210191082", label="4.87",style=bold];
"PNLIPRP3_E" -> "TRIML2_E" [color="0 0 0.24707006369426743", label="4.93",style=bold];
"ZAN_S" -> "CSMD1_E" [color="0 0 0.24649681528662415", label="4.94",style=bold];
"PCDHA13_S" -> "LHCGR_S" [color="0 0 0.24592356687898087", label="5.00",style=bold];
"LIN28B_S" -> "SPAG17_S" [color="0 0 0.24535031847133748", label="5.05",style=bold];
"DMBT1_S" -> "SBSN_M" [color="0 0 0.2447770700636942", label="5.08",style=bold];
"FBN3_S" -> "FREM2_S" [color="0 0 0.24420382165605092", label="5.09",style=bold];
"PCDHA1_M" -> "ADAMTS20_M" [color="0 0 0.24363057324840753", label="5.11",style=bold];
"PCDHA13_S" -> "FOXG1_S" [color="0 0 0.24305732484076426", label="5.11",style=bold];
"ZNF716_M" -> "FLG_M" [color="0 0 0.24248407643312098", label="5.12",style=bold];
"MUC5B_S" -> "MYO18B_S" [color="0 0 0.2419108280254777", label="5.12",style=bold];
"LHCGR_M" -> "SBSN_M" [color="0 0 0.2413375796178343", label="5.12",style=bold];
"IVL_E" -> "MUC6_E" [color="0 0 0.24076433121019103", label="5.22",style=bold];
"KPRP_S" -> "MUC17_S" [color="0 0 0.24019108280254775", label="5.22",style=bold];
"FBN3_S" -> "MUC5B_S" [color="0 0 0.23961783439490436", label="5.22",style=bold];
"MUC4_S" -> "GPR98_S" [color="0 0 0.23904458598726108", label="5.24",style=bold];
"FBN3_S" -> "TRPM3_S" [color="0 0 0.2384713375796178", label="5.24",style=bold];
"MUC2_S" -> "DPP6_S" [color="0 0 0.23789808917197441", label="5.32",style=bold];
"TPTE_S" -> "ADAMTS20_S" [color="0 0 0.23732484076433114", label="5.33",style=bold];
"CSMD3_M" -> "ZAN_M" [color="0 0 0.23675159235668786", label="5.36",style=bold];
"ADAMTS19_S" -> "PNLIPRP3_S" [color="0 0 0.23617834394904458", label="5.37",style=bold];
"TMPRSS11A_S" -> "FBN3_M" [color="0 0 0.2356050955414012", label="5.38",style=bold];
"AKNAD1_S" -> "FBN3_S" [color="0 0 0.2350318471337579", label="5.39",style=bold];
"SERPINB11_S" -> "SLC1A6_S" [color="0 0 0.23445859872611463", label="5.40",style=bold];
"BNC1_M" -> "ADAMTS20_M" [color="0 0 0.23388535031847124", label="5.42",style=bold];
"ZIC4_S" -> "APOB_E" [color="0 0 0.23331210191082796", label="5.48",style=bold];
"FBN3_S" -> "LRRC7_S" [color="0 0 0.23273885350318468", label="5.49",style=bold];
"FLG_M" -> "A2ML1_M" [color="0 0 0.2321656050955413", label="5.53",style=bold];
"DPP6_S" -> "SPAG17_S" [color="0 0 0.23159235668789802", label="5.68",style=bold];
"KPRP_S" -> "AKNAD1_S" [color="0 0 0.23101910828025474", label="5.71",style=bold];
"CHRNA4_M" -> "AKNAD1_S" [color="0 0 0.23044585987261146", label="5.71",style=bold];
"KRT1_S" -> "LIPF_M" [color="0 0 0.22987261146496807", label="5.73",style=bold];
"A2ML1_M" -> "CNTNAP5_S" [color="0 0 0.2292993630573248", label="5.74",style=bold];
"SLC12A1_S" -> "DUSP27_M" [color="0 0 0.2287261146496815", label="5.75",style=bold];
"LOC100190940_S" -> "MUC17_S" [color="0 0 0.22815286624203812", label="5.76",style=bold];
"SLC1A6_S" -> "APOB_E" [color="0 0 0.22757961783439484", label="5.77",style=bold];
"A2ML1_S" -> "PNLIPRP3_S" [color="0 0 0.22700636942675156", label="5.82",style=bold];
"FLG_S" -> "SERPINB11_M" [color="0 0 0.22643312101910817", label="5.84",style=bold];
"MUC2_S" -> "MUC5B_M" [color="0 0 0.2258598726114649", label="5.87",style=bold];
"UNC5D_S" -> "DMBT1_M" [color="0 0 0.22528662420382162", label="5.87",style=bold];
"MUC5B_S" -> "NELL1_E" [color="0 0 0.22471337579617834", label="5.87",style=bold];
"NELL1_S" -> "ZAN_E" [color="0 0 0.22414012738853495", label="5.90",style=bold];
"CRCT1_S" -> "NDST4_S" [color="0 0 0.22356687898089167", label="5.91",style=bold];
"KRT6C_E" -> "FLG_E" [color="0 0 0.2229936305732484", label="5.98",style=bold];
"IRX4_M" -> "BNC1_M" [color="0 0 0.222420382165605", label="6.05",style=bold];
"KCNV1_S" -> "KRT78_M" [color="0 0 0.22184713375796172", label="6.06",style=bold];
"FLG_M" -> "LRRC7_M" [color="0 0 0.22127388535031844", label="6.10",style=bold];
"CRCT1_S" -> "KCNV1_S" [color="0 0 0.22070063694267505", label="6.12",style=bold];
"MUC2_S" -> "ZAN_S" [color="0 0 0.22012738853503178", label="6.14",style=bold];
"MYO18B_M" -> "MUC16_S" [color="0 0 0.2195541401273885", label="6.17",style=bold];
"IVL_E" -> "PCDHA1_M" [color="0 0 0.21898089171974522", label="6.17",style=bold];
"ADAMTS20_S" -> "MUC4_S" [color="0 0 0.21840764331210183", label="6.19",style=bold];
"KCNV1_M" -> "GPR98_M" [color="0 0 0.21783439490445855", label="6.22",style=bold];
"TRPM3_E" -> "CSMD1_E" [color="0 0 0.21726114649681527", label="6.22",style=bold];
"MUC17_S" -> "SLC12A1_S" [color="0 0 0.21668789808917188", label="6.24",style=bold];
"AKNAD1_S" -> "MUC17_S" [color="0 0 0.2161146496815286", label="6.25",style=bold];
"RPL10L_S" -> "ZNF716_E" [color="0 0 0.21554140127388532", label="6.25",style=bold];
"APOB_S" -> "TPTE_S" [color="0 0 0.21496815286624193", label="6.25",style=bold];
"RIMS1_M" -> "KRT24_M" [color="0 0 0.21439490445859866", label="6.26",style=bold];
"LOC100190940_M" -> "LRRC7_E" [color="0 0 0.21382165605095538", label="6.31",style=bold];
"MYO18B_S" -> "UNC5D_E" [color="0 0 0.213248407643312", label="6.32",style=bold];
"SBSN_S" -> "FBN3_S" [color="0 0 0.2126751592356687", label="6.34",style=bold];
"NELL1_M" -> "KCNV1_M" [color="0 0 0.21210191082802543", label="6.35",style=bold];
"LOC100190940_M" -> "MUC17_S" [color="0 0 0.21152866242038215", label="6.35",style=bold];
"FREM2_M" -> "NDST4_M" [color="0 0 0.21095541401273876", label="6.37",style=bold];
"KCNV1_S" -> "TPTE_S" [color="0 0 0.21038216560509548", label="6.40",style=bold];
"GABRG2_S" -> "NELL1_E" [color="0 0 0.2098089171974522", label="6.40",style=bold];
"GABRA1_M" -> "ZAN_M" [color="0 0 0.20923566878980882", label="6.41",style=bold];
"PNLIPRP3_E" -> "CWH43_E" [color="0 0 0.20866242038216554", label="6.51",style=bold];
"MUC4_M" -> "AKNAD1_E" [color="0 0 0.20808917197452226", label="6.52",style=bold];
"LHCGR_M" -> "APOB_M" [color="0 0 0.20751592356687887", label="6.59",style=bold];
"A2ML1_E" -> "PCDHA1_M" [color="0 0 0.2069426751592356", label="6.60",style=bold];
"SOX1_S" -> "PAX1_S" [color="0 0 0.2063694267515923", label="6.61",style=bold];
"AKNAD1_S" -> "KRT1_S" [color="0 0 0.20579617834394903", label="6.65",style=bold];
"KRT6C_E" -> "BNC1_E" [color="0 0 0.20522292993630564", label="6.65",style=bold];
"SBSN_E" -> "CRCT1_E" [color="0 0 0.20464968152866236", label="6.66",style=bold];
"MUC2_S" -> "RIMS1_S" [color="0 0 0.20407643312101909", label="6.66",style=bold];
"DSG1_M" -> "Tumor_Status" [color="0 0 0.2035031847133757", label="6.73",style=bold];
"FREM2_S" -> "ABCA13_S" [color="0 0 0.20292993630573242", label="6.76",style=bold];
"CSMD3_S" -> "MUC4_S" [color="0 0 0.20235668789808914", label="6.78",style=bold];
"ZNF716_M" -> "MUC2_M" [color="0 0 0.20178343949044575", label="6.87",style=bold];
"ZNF716_M" -> "LHCGR_M" [color="0 0 0.20121019108280247", label="6.92",style=bold];
"SLC1A6_M" -> "MUC2_M" [color="0 0 0.2006369426751592", label="7.09",style=bold];
"IRX4_E" -> "FOXG1_E" [color="0 0 0.2000636942675159", label="7.12",style=bold];
"Tumor_Status" -> "HTR4_E" [color="0 0 0.19949044585987252", label="7.18",style=bold];
"CNTNAP5_M" -> "SBSN_M" [color="0 0 0.19891719745222924", label="7.19",style=bold];
"DSG1_E" -> "LOC100190940_E" [color="0 0 0.19834394904458597", label="7.22",style=bold];
"FREM2_M" -> "SPAG17_M" [color="0 0 0.19777070063694258", label="7.24",style=bold];
"SERPINB13_E" -> "TRIML2_E" [color="0 0 0.1971974522292993", label="7.26",style=bold];
"FREM2_S" -> "MYO18B_S" [color="0 0 0.19662420382165602", label="7.28",style=bold];
"KCNV1_E" -> "CNTNAP5_E" [color="0 0 0.19605095541401263", label="7.33",style=bold];
"FBN3_M" -> "KRT78_M" [color="0 0 0.19547770700636935", label="7.35",style=bold];
"DMBT1_S" -> "MUC4_S" [color="0 0 0.19490445859872607", label="7.36",style=bold];
"PCDHA13_S" -> "APOB_S" [color="0 0 0.1943312101910828", label="7.38",style=bold];
"LIN28B_S" -> "TMPRSS11A_S" [color="0 0 0.1937579617834394", label="7.46",style=bold];
"BNC1_E" -> "FLG_E" [color="0 0 0.19318471337579612", label="7.50",style=bold];
"ZIC4_M" -> "LOC100190940_M" [color="0 0 0.19261146496815285", label="7.54",style=bold];
"NELL1_M" -> "LRRIQ1_M" [color="0 0 0.19203821656050946", label="7.63",style=bold];
"A2ML1_E" -> "BNC1_E" [color="0 0 0.19146496815286618", label="7.66",style=bold];
"SBSN_E" -> "PCDHA1_M" [color="0 0 0.1908917197452229", label="7.68",style=bold];
"SERPINB13_E" -> "FLG_E" [color="0 0 0.1903184713375795", label="7.73",style=bold];
"A2ML1_S" -> "SLC1A6_S" [color="0 0 0.18974522292993623", label="7.75",style=bold];
"LHCGR_M" -> "LRRC7_M" [color="0 0 0.18917197452229295", label="7.78",style=bold];
"UNC5D_M" -> "DSG1_E" [color="0 0 0.18859872611464967", label="7.79",style=bold];
"SLC12A1_S" -> "LRRIQ1_S" [color="0 0 0.18802547770700628", label="7.88",style=bold];
"MUC2_S" -> "DCT_S" [color="0 0 0.187452229299363", label="7.93",style=bold];
"SLC1A6_E" -> "TRIML2_E" [color="0 0 0.18687898089171973", label="8.02",style=bold];
"GABRA1_M" -> "PNLDC1_E" [color="0 0 0.18630573248407634", label="8.06",style=bold];
"LRRIQ1_S" -> "MYBPC1_M" [color="0 0 0.18573248407643306", label="8.15",style=bold];
"KRT24_S" -> "CSMD3_S" [color="0 0 0.18515923566878978", label="8.19",style=bold];
"A2ML1_S" -> "PCDHA13_S" [color="0 0 0.1845859872611464", label="8.22",style=bold];
"SERPINB13_S" -> "GABRA1_S" [color="0 0 0.1840127388535031", label="8.33",style=bold];
"MYO18B_S" -> "MUC16_S" [color="0 0 0.18343949044585983", label="8.38",style=bold];
"PCDHA1_E" -> "APOB_E" [color="0 0 0.18286624203821655", label="8.38",style=bold];
"RIMS1_S" -> "CSMD3_S" [color="0 0 0.18229299363057316", label="8.42",style=bold];
"DSG1_E" -> "APOB_S" [color="0 0 0.18171974522292988", label="8.43",style=bold];
"APOB_S" -> "ABCA13_S" [color="0 0 0.1811464968152866", label="8.46",style=bold];
"DPP6_E" -> "TRPM3_E" [color="0 0 0.18057324840764322", label="8.50",style=bold];
"CSMD3_E" -> "SYT16_E" [color="0 0 0.17999999999999994", label="8.58",style=bold];
"TRPM3_E" -> "KCNV1_E" [color="0 0 0.17942675159235666", label="8.61",style=bold];
"TRPM3_E" -> "MUC4_M" [color="0 0 0.17885350318471327", label="8.63",style=bold];
"CFHR1_S" -> "PAX1_S" [color="0 0 0.17828025477707", label="8.86",style=bold];
"GABRA1_E" -> "LMOD2_E" [color="0 0 0.1777070063694267", label="8.87",style=bold];
"PCDHA1_M" -> "ZIC4_M" [color="0 0 0.17713375796178332", label="8.96",style=bold];
"MUC2_M" -> "DUSP27_M" [color="0 0 0.17656050955414004", label="8.96",style=bold];
"SPRR2A_M" -> "SLC12A1_M" [color="0 0 0.17598726114649677", label="8.98",style=bold];
"MUC2_M" -> "FLG_S" [color="0 0 0.1754140127388535", label="9.08",style=bold];
"CFHR1_S" -> "LRRC7_S" [color="0 0 0.1748407643312101", label="9.09",style=bold];
"ADAMTS20_M" -> "GABRA1_M" [color="0 0 0.17426751592356682", label="9.12",style=bold];
"RIMS1_E" -> "CNTNAP4_E" [color="0 0 0.17369426751592354", label="9.16",style=bold];
"ABCA13_S" -> "LRRIQ1_S" [color="0 0 0.17312101910828015", label="9.30",style=bold];
"KCNV1_S" -> "ZAN_S" [color="0 0 0.17254777070063687", label="9.33",style=bold];
"MUC17_S" -> "FREM2_S" [color="0 0 0.1719745222929936", label="9.40",style=bold];
"FREM2_E" -> "GPR98_S" [color="0 0 0.1714012738853502", label="9.42",style=bold];
"DPP6_S" -> "DMBT1_M" [color="0 0 0.17082802547770692", label="9.45",style=bold];
"KCNV1_M" -> "CRCT1_M" [color="0 0 0.17025477707006365", label="9.52",style=bold];
"TPTE_M" -> "GABRG2_M" [color="0 0 0.16968152866242037", label="9.55",style=bold];
"CSMD1_E" -> "TPTE_E" [color="0 0 0.16910828025477698", label="9.56",style=bold];
"PGC_E" -> "KCNA10_E" [color="0 0 0.1685350318471337", label="9.60",style=bold];
"GABRA1_E" -> "SYT16_E" [color="0 0 0.16796178343949042", label="9.61",style=bold];
"KRT78_M" -> "DCT_M" [color="0 0 0.16738853503184703", label="9.68",style=bold];
"SLC12A1_M" -> "TRPM3_M" [color="0 0 0.16681528662420375", label="9.71",style=bold];
"ABCA13_M" -> "ABCA13_E" [color="0 0 0.16624203821656047", label="9.89",style=bold];
"ZNF716_M" -> "DUSP27_M" [color="0 0 0.16566878980891708", label="9.93",style=bold];
"SPAG17_E" -> "PNLDC1_E" [color="0 0 0.1650955414012738", label="9.96",style=bold];
"GPR98_M" -> "LMOD2_M" [color="0 0 0.16452229299363053", label="10.07",style=bold];
"BNC1_M" -> "IRX4_E" [color="0 0 0.16394904458598725", label="10.23",style=bold];
"MYBPC1_E" -> "MYO18B_E" [color="0 0 0.16337579617834386", label="10.37",style=bold];
"MUC6_E" -> "CSMD3_S" [color="0 0 0.16280254777070058", label="10.46",style=bold];
"LRRC7_E" -> "LHCGR_E" [color="0 0 0.1622292993630573", label="10.61",style=bold];
"ZNF716_E" -> "PCDHA1_E" [color="0 0 0.1616560509554139", label="10.66",style=bold];
"ZAN_S" -> "ADH7_M" [color="0 0 0.16108280254777063", label="10.68",style=bold];
"APOB_M" -> "MUC2_M" [color="0 0 0.16050955414012735", label="10.72",style=bold];
"MUC4_S" -> "CFHR1_E" [color="0 0 0.15993630573248396", label="10.79",style=bold];
"MUC5B_M" -> "MUC6_M" [color="0 0 0.15936305732484068", label="10.87",style=bold];
"ZNF716_M" -> "KRT24_M" [color="0 0 0.1587898089171974", label="10.87",style=bold];
"SERPINB13_E" -> "PNLIPRP3_E" [color="0 0 0.15821656050955413", label="10.88",style=bold];
"APOB_E" -> "CA4_E" [color="0 0 0.15764331210191074", label="10.92",style=bold];
"KRT1_M" -> "FLG_M" [color="0 0 0.15707006369426746", label="10.93",style=bold];
"ADH7_E" -> "MYBPC1_E" [color="0 0 0.15649681528662418", label="11.00",style=bold];
"CA4_M" -> "GPR98_M" [color="0 0 0.1559235668789808", label="11.10",style=bold];
"SLC1A6_M" -> "SYT16_M" [color="0 0 0.1553503184713375", label="11.17",style=bold];
"SLC12A1_E" -> "LMOD2_E" [color="0 0 0.15477707006369423", label="11.25",style=bold];
"RIMS1_M" -> "FREM2_M" [color="0 0 0.15420382165605084", label="11.32",style=bold];
"PGC_E" -> "MYBPC1_E" [color="0 0 0.15363057324840756", label="11.46",style=bold];
"CSMD1_M" -> "ZNF716_E" [color="0 0 0.1530573248407643", label="11.50",style=bold];
"KPRP_M" -> "CA4_E" [color="0 0 0.152484076433121", label="11.60",style=bold];
"PNLIPRP3_E" -> "PCDHA1_M" [color="0 0 0.15191082802547762", label="11.62",style=bold];
"TMPRSS11A_E" -> "KRT78_E" [color="0 0 0.15133757961783434", label="11.67",style=bold];
"GPR98_M" -> "MUC16_S" [color="0 0 0.15076433121019106", label="11.76",style=bold];
"CHRNA4_M" -> "FREM2_E" [color="0 0 0.15019108280254767", label="11.79",style=bold];
"GABRA1_E" -> "HTR4_E" [color="0 0 0.1496178343949044", label="11.80",style=bold];
"ADAMTS19_M" -> "RIMS1_S" [color="0 0 0.1490445859872611", label="11.85",style=bold];
"SLC1A6_E" -> "CNTNAP5_E" [color="0 0 0.14847133757961772", label="11.90",style=bold];
"DSG1_E" -> "LOC100190940_M" [color="0 0 0.14789808917197445", label="12.05",style=bold];
"CSMD1_E" -> "MYO18B_E" [color="0 0 0.14732484076433117", label="12.12",style=bold];
"NELL1_M" -> "CSMD1_S" [color="0 0 0.14675159235668778", label="12.15",style=bold];
"KRT24_M" -> "MYBPC1_M" [color="0 0 0.1461783439490445", label="12.31",style=bold];
"CHRNA4_E" -> "FBN3_E" [color="0 0 0.14560509554140122", label="12.39",style=bold];
"CNTNAP4_E" -> "CSMD1_E" [color="0 0 0.14503184713375794", label="12.48",style=bold];
"KRT6C_E" -> "A2ML1_E" [color="0 0 0.14445859872611455", label="12.49",style=bold];
"A2ML1_E" -> "TMPRSS11A_E" [color="0 0 0.14388535031847127", label="12.67",style=bold];
"CHRNA4_M" -> "CRCT1_M" [color="0 0 0.143312101910828", label="12.84",style=bold];
"PNLIPRP3_E" -> "TMPRSS11A_M" [color="0 0 0.1427388535031846", label="13.01",style=bold];
"KRT1_M" -> "TRPM3_M" [color="0 0 0.14216560509554133", label="13.09",style=bold];
"KRT6C_E" -> "KRT1_E" [color="0 0 0.14159235668789805", label="13.18",style=bold];
"CA4_M" -> "SPAG17_M" [color="0 0 0.14101910828025466", label="13.22",style=bold];
"SERPINB13_E" -> "FOXG1_E" [color="0 0 0.14044585987261138", label="13.32",style=bold];
"KRT6C_E" -> "SBSN_E" [color="0 0 0.1398726114649681", label="13.38",style=bold];
"ZNF716_M" -> "CNTNAP5_M" [color="0 0 0.13929936305732482", label="13.41",style=bold];
"BNC1_M" -> "MUC5B_E" [color="0 0 0.13872611464968143", label="13.42",style=bold];
"DSG1_M" -> "LIPF_M" [color="0 0 0.13815286624203815", label="13.47",style=bold];
"DCT_M" -> "A2ML1_M" [color="0 0 0.13757961783439487", label="13.50",style=bold];
"SPAG17_E" -> "GPR98_E" [color="0 0 0.13700636942675148", label="13.52",style=bold];
"TRPM3_E" -> "GABRA1_E" [color="0 0 0.1364331210191082", label="13.76",style=bold];
"ADH7_M" -> "LIPF_M" [color="0 0 0.13585987261146493", label="13.80",style=bold];
"IVL_E" -> "CWH43_E" [color="0 0 0.13528662420382154", label="13.80",style=bold];
"TMPRSS11A_M" -> "PCDHA13_E" [color="0 0 0.13471337579617826", label="13.88",style=bold];
"DMBT1_M" -> "MUC17_M" [color="0 0 0.13414012738853498", label="13.94",style=bold];
"KPRP_E" -> "KRT1_E" [color="0 0 0.1335668789808917", label="13.95",style=bold];
"LIN28B_E" -> "LOC100190940_E" [color="0 0 0.1329936305732483", label="13.99",style=bold];
"LOC100190940_M" -> "PNLDC1_E" [color="0 0 0.13242038216560503", label="14.00",style=bold];
"LOC100190940_M" -> "SPAG17_M" [color="0 0 0.13184713375796175", label="14.09",style=bold];
"SLC12A1_E" -> "DUSP27_E" [color="0 0 0.13127388535031836", label="14.17",style=bold];
"TMPRSS11A_E" -> "ADH7_E" [color="0 0 0.13070063694267509", label="14.24",style=bold];
"CNTNAP5_E" -> "PCDHA1_E" [color="0 0 0.1301273885350318", label="14.27",style=bold];
"DMBT1_E" -> "LRRC7_E" [color="0 0 0.12955414012738842", label="14.31",style=bold];
"DUSP27_M" -> "PGC_M" [color="0 0 0.12898089171974514", label="14.32",style=bold];
"SLC1A6_E" -> "CSMD3_E" [color="0 0 0.12840764331210186", label="14.32",style=bold];
"MUC17_E" -> "KCNA10_E" [color="0 0 0.12783439490445858", label="14.41",style=bold];
"KCNV1_E" -> "ZNF716_E" [color="0 0 0.1272611464968152", label="14.43",style=bold];
"CNTNAP5_M" -> "RIMS1_M" [color="0 0 0.1266878980891719", label="14.43",style=bold];
"DCT_M" -> "SYT16_M" [color="0 0 0.12611464968152863", label="14.49",style=bold];
"LRRC7_E" -> "TRPM3_E" [color="0 0 0.12554140127388524", label="14.52",style=bold];
"IVL_E" -> "SBSN_E" [color="0 0 0.12496815286624197", label="14.59",style=bold];
"IVL_E" -> "KRT78_E" [color="0 0 0.12439490445859869", label="14.60",style=bold];
"KPRP_E" -> "PNLIPRP3_E" [color="0 0 0.1238216560509553", label="14.69",style=bold];
"LHCGR_M" -> "TRPM3_M" [color="0 0 0.12324840764331202", label="14.74",style=bold];
"AKNAD1_E" -> "SLC12A1_E" [color="0 0 0.12267515923566874", label="14.85",style=bold];
"KRT6C_M" -> "AKNAD1_M" [color="0 0 0.12210191082802546", label="14.91",style=bold];
"PCDHA1_M" -> "CWH43_M" [color="0 0 0.12152866242038207", label="14.95",style=bold];
"NDST4_E" -> "ADAMTS19_E" [color="0 0 0.1209554140127388", label="14.95",style=bold];
"FLG_E" -> "KRT6C_M" [color="0 0 0.12038216560509551", label="15.03",style=bold];
"BNC1_M" -> "PGC_E" [color="0 0 0.11980891719745212", label="15.26",style=bold];
"BNC1_M" -> "CSMD1_E" [color="0 0 0.11923566878980885", label="15.31",style=bold];
"NELL1_E" -> "RIMS1_E" [color="0 0 0.11866242038216557", label="15.49",style=bold];
"TRPM3_E" -> "UNC5D_E" [color="0 0 0.11808917197452218", label="15.50",style=bold];
"SBSN_E" -> "BNC1_E" [color="0 0 0.1175159235668789", label="15.54",style=bold];
"SPRR2A_E" -> "ABCA13_E" [color="0 0 0.11694267515923562", label="15.54",style=bold];
"GABRA1_M" -> "LIN28B_M" [color="0 0 0.11636942675159223", label="15.57",style=bold];
"ZNF716_M" -> "APOB_M" [color="0 0 0.11579617834394895", label="15.57",style=bold];
"PCDHA1_M" -> "PAX1_M" [color="0 0 0.11522292993630567", label="15.64",style=bold];
"KPRP_M" -> "DUSP27_M" [color="0 0 0.1146496815286624", label="15.66",style=bold];
"BNC1_M" -> "CRCT1_M" [color="0 0 0.114076433121019", label="15.73",style=bold];
"IVL_M" -> "DMBT1_E" [color="0 0 0.11350318471337573", label="15.74",style=bold];
"MUC6_E" -> "MUC6_M" [color="0 0 0.11292993630573245", label="15.76",style=bold];
"SBSN_M" -> "KRT1_M" [color="0 0 0.11235668789808906", label="15.90",style=bold];
"KPRP_M" -> "SOX1_M" [color="0 0 0.11178343949044578", label="15.92",style=bold];
"FOXG1_E" -> "DSG1_M" [color="0 0 0.1112101910828025", label="15.97",style=bold];
"ADH7_M" -> "KRT24_M" [color="0 0 0.11063694267515911", label="16.06",style=bold];
"DCT_E" -> "SLC12A1_E" [color="0 0 0.11006369426751583", label="16.06",style=bold];
"RIMS1_E" -> "NDST4_E" [color="0 0 0.10949044585987255", label="16.17",style=bold];
"SLC1A6_E" -> "SLC12A1_E" [color="0 0 0.10891719745222928", label="16.22",style=bold];
"GPR98_E" -> "NELL1_E" [color="0 0 0.10834394904458589", label="16.27",style=bold];
"MYO18B_M" -> "AKNAD1_M" [color="0 0 0.1077707006369426", label="16.30",style=bold];
"SPAG17_E" -> "LRRIQ1_E" [color="0 0 0.10719745222929933", label="16.38",style=bold];
"KPRP_M" -> "FREM2_M" [color="0 0 0.10662420382165594", label="16.54",style=bold];
"SERPINB13_M" -> "SERPINB11_M" [color="0 0 0.10605095541401266", label="16.85",style=bold];
"GABRA1_M" -> "GABRG2_E" [color="0 0 0.10547770700636938", label="16.89",style=bold];
"NELL1_E" -> "MYO18B_E" [color="0 0 0.10490445859872599", label="16.92",style=bold];
"BNC1_E" -> "IRX4_M" [color="0 0 0.10433121019108271", label="16.95",style=bold];
"KCNV1_M" -> "LRRIQ1_M" [color="0 0 0.10375796178343943", label="16.95",style=bold];
"UNC5D_M" -> "CNTNAP4_M" [color="0 0 0.10318471337579616", label="17.20",style=bold];
"MUC2_E" -> "MUC4_E" [color="0 0 0.10261146496815277", label="17.21",style=bold];
"TPTE_M" -> "NELL1_M" [color="0 0 0.10203821656050949", label="17.33",style=bold];
"CWH43_M" -> "GPR98_M" [color="0 0 0.10146496815286621", label="17.41",style=bold];
"DCT_M" -> "SLC12A1_M" [color="0 0 0.10089171974522282", label="17.47",style=bold];
"LHCGR_M" -> "SYT16_M" [color="0 0 0.10031847133757954", label="17.77",style=bold];
"SERPINB13_M" -> "A2ML1_M" [color="0 0 0.09974522292993626", label="17.85",style=bold];
"FREM2_M" -> "PCDHA13_M" [color="0 0 0.09917197452229287", label="17.88",style=bold];
"LRRC7_E" -> "AKNAD1_E" [color="0 0 0.09859872611464959", label="17.88",style=bold];
"IRX4_M" -> "CHRNA4_E" [color="0 0 0.09802547770700631", label="17.88",style=bold];
"LHCGR_E" -> "DCT_E" [color="0 0 0.09745222929936304", label="17.92",style=bold];
"ADAMTS19_M" -> "FREM2_M" [color="0 0 0.09687898089171965", label="18.07",style=bold];
"RIMS1_M" -> "LRRC7_M" [color="0 0 0.09630573248407637", label="18.07",style=bold];
"PGC_M" -> "MUC17_M" [color="0 0 0.09573248407643309", label="18.18",style=bold];
"LIPF_M" -> "PNLDC1_M" [color="0 0 0.0951592356687897", label="18.32",style=bold];
"ADAMTS20_E" -> "SLC1A6_E" [color="0 0 0.09458598726114642", label="18.63",style=bold];
"BNC1_M" -> "FOXG1_M" [color="0 0 0.09401273885350314", label="18.69",style=bold];
"PCDHA1_E" -> "PCDHA13_E" [color="0 0 0.09343949044585975", label="18.82",style=bold];
"IVL_E" -> "SLC1A6_M" [color="0 0 0.09286624203821647", label="18.88",style=bold];
"LRRC7_M" -> "SERPINB13_M" [color="0 0 0.0922929936305732", label="19.05",style=bold];
"LIN28B_E" -> "SLC1A6_E" [color="0 0 0.09171974522292992", label="19.16",style=bold];
"FLG_M" -> "DCT_M" [color="0 0 0.09114649681528653", label="19.18",style=bold];
"DPP6_E" -> "CFHR1_E" [color="0 0 0.09057324840764325", label="19.44",style=bold];
"RIMS1_M" -> "FBN3_E" [color="0 0 0.08999999999999997", label="19.69",style=bold];
"FOXG1_M" -> "ADAMTS20_M" [color="0 0 0.08942675159235658", label="19.83",style=bold];
"BNC1_M" -> "LIPF_M" [color="0 0 0.0888535031847133", label="20.08",style=bold];
"SLC1A6_E" -> "FBN3_E" [color="0 0 0.08828025477707002", label="20.12",style=bold];
"LRRIQ1_E" -> "Tumor_Status" [color="0 0 0.08770700636942663", label="20.15",style=bold];
"MUC5B_E" -> "MUC5B_M" [color="0 0 0.08713375796178335", label="20.21",style=bold];
"BNC1_M" -> "LOC100190940_E" [color="0 0 0.08656050955414007", label="20.31",style=bold];
"TMPRSS11A_E" -> "MUC4_E" [color="0 0 0.0859872611464968", label="20.36",style=bold];
"ZAN_E" -> "PAX1_E" [color="0 0 0.0854140127388534", label="20.42",style=bold];
"LRRC7_M" -> "ABCA13_M" [color="0 0 0.08484076433121013", label="20.60",style=bold];
"FBN3_M" -> "MYO18B_M" [color="0 0 0.08426751592356685", label="20.67",style=bold];
"CRCT1_E" -> "SPRR2A_E" [color="0 0 0.08369426751592346", label="20.77",style=bold];
"SPAG17_M" -> "SPAG17_E" [color="0 0 0.08312101910828018", label="20.78",style=bold];
"MUC5B_E" -> "MUC2_E" [color="0 0 0.0825477707006369", label="20.83",style=bold];
"LHCGR_M" -> "LIN28B_E" [color="0 0 0.08197452229299351", label="20.99",style=bold];
"BNC1_E" -> "BNC1_M" [color="0 0 0.08140127388535023", label="21.19",style=bold];
"KRT6C_E" -> "TMPRSS11A_E" [color="0 0 0.08082802547770696", label="21.19",style=bold];
"SOX1_E" -> "CFHR1_E" [color="0 0 0.08025477707006357", label="21.30",style=bold];
"KRT1_M" -> "SERPINB11_M" [color="0 0 0.07968152866242029", label="21.37",style=bold];
"CHRNA4_M" -> "CA4_M" [color="0 0 0.07910828025477701", label="21.55",style=bold];
"DCT_M" -> "CFHR1_M" [color="0 0 0.07853503184713373", label="21.62",style=bold];
"CSMD3_M" -> "KCNV1_M" [color="0 0 0.07796178343949034", label="21.62",style=bold];
"NDST4_M" -> "HTR4_M" [color="0 0 0.07738853503184706", label="21.66",style=bold];
"IVL_E" -> "A2ML1_E" [color="0 0 0.07681528662420378", label="21.73",style=bold];
"CSMD1_M" -> "ZNF716_M" [color="0 0 0.07624203821656039", label="21.74",style=bold];
"SLC1A6_M" -> "CSMD1_M" [color="0 0 0.07566878980891711", label="21.77",style=bold];
"MUC17_E" -> "DUSP27_E" [color="0 0 0.07509554140127384", label="21.97",style=bold];
"ADH7_E" -> "ABCA13_E" [color="0 0 0.07452229299363045", label="22.68",style=bold];
"PAX1_M" -> "GABRA1_M" [color="0 0 0.07394904458598717", label="22.93",style=bold];
"SLC12A1_M" -> "LMOD2_M" [color="0 0 0.07337579617834389", label="22.94",style=bold];
"ZNF716_M" -> "SERPINB13_M" [color="0 0 0.07280254777070061", label="22.99",style=bold];
"LRRC7_M" -> "SLC12A1_M" [color="0 0 0.07222929936305722", label="23.04",style=bold];
"IVL_E" -> "ZAN_E" [color="0 0 0.07165605095541394", label="23.08",style=bold];
"MUC2_M" -> "ZAN_M" [color="0 0 0.07108280254777066", label="23.20",style=bold];
"SYT16_M" -> "TRIML2_M" [color="0 0 0.07050955414012727", label="23.21",style=bold];
"ZAN_E" -> "AKNAD1_E" [color="0 0 0.069936305732484", label="23.63",style=bold];
"BNC1_M" -> "CWH43_M" [color="0 0 0.06936305732484072", label="23.64",style=bold];
"FLG_M" -> "ABCA13_M" [color="0 0 0.06878980891719733", label="23.70",style=bold];
"KRT1_M" -> "PGC_M" [color="0 0 0.06821656050955405", label="23.87",style=bold];
"SLC12A1_M" -> "PNLIPRP3_M" [color="0 0 0.06764331210191077", label="23.87",style=bold];
"PNLIPRP3_M" -> "TMPRSS11A_M" [color="0 0 0.06707006369426749", label="23.92",style=bold];
"DPP6_M" -> "ZNF716_M" [color="0 0 0.0664968152866241", label="24.06",style=bold];
"CNTNAP5_E" -> "GABRG2_E" [color="0 0 0.06592356687898082", label="24.12",style=bold];
"MUC4_M" -> "DMBT1_M" [color="0 0 0.06535031847133754", label="24.14",style=bold];
"MUC6_E" -> "MUC5B_E" [color="0 0 0.06477707006369415", label="24.16",style=bold];
"DPP6_M" -> "CSMD1_M" [color="0 0 0.06420382165605087", label="24.20",style=bold];
"GABRA1_M" -> "MYO18B_E" [color="0 0 0.0636305732484076", label="24.20",style=bold];
"DSG1_E" -> "MUC17_E" [color="0 0 0.0630573248407642", label="24.58",style=bold];
"PCDHA1_M" -> "DPP6_M" [color="0 0 0.06248407643312093", label="24.62",style=bold];
"IVL_E" -> "CRCT1_E" [color="0 0 0.06191082802547765", label="24.75",style=bold];
"LIN28B_E" -> "TPTE_E" [color="0 0 0.06133757961783437", label="24.80",style=bold];
"ADAMTS20_E" -> "LIN28B_E" [color="0 0 0.06076433121019098", label="24.85",style=bold];
"IVL_M" -> "PNLIPRP3_M" [color="0 0 0.0601910828025477", label="24.91",style=bold];
"DPP6_M" -> "RIMS1_M" [color="0 0 0.05961783439490442", label="25.22",style=bold];
"LRRC7_M" -> "NDST4_M" [color="0 0 0.05904458598726103", label="25.25",style=bold];
"SYT16_M" -> "LMOD2_M" [color="0 0 0.058471337579617755", label="25.36",style=bold];
"CHRNA4_M" -> "MUC5B_M" [color="0 0 0.057898089171974476", label="25.43",style=bold];
"MUC16_E" -> "MUC16_M" [color="0 0 0.057324840764331086", label="25.54",style=bold];
"TMPRSS11A_M" -> "CFHR1_M" [color="0 0 0.05675159235668781", label="25.83",style=bold];
"ADH7_E" -> "SERPINB11_E" [color="0 0 0.05617834394904453", label="25.85",style=bold];
"APOB_E" -> "HTR4_E" [color="0 0 0.05560509554140125", label="26.46",style=bold];
"ADAMTS19_M" -> "ADAMTS19_E" [color="0 0 0.05503184713375786", label="26.49",style=bold];
"LOC100190940_M" -> "KCNV1_M" [color="0 0 0.05445859872611458", label="26.50",style=bold];
"NDST4_E" -> "UNC5D_E" [color="0 0 0.0538853503184713", label="26.78",style=bold];
"DMBT1_E" -> "ADH7_M" [color="0 0 0.053312101910827914", label="26.81",style=bold];
"SOX1_M" -> "BNC1_M" [color="0 0 0.052738853503184635", label="26.86",style=bold];
"KCNV1_M" -> "ADAMTS19_M" [color="0 0 0.052165605095541356", label="26.95",style=bold];
"MUC2_M" -> "SBSN_M" [color="0 0 0.05159235668789797", label="27.03",style=bold];
"KCNV1_M" -> "CA4_M" [color="0 0 0.05101910828025469", label="27.17",style=bold];
"MUC2_M" -> "MUC6_M" [color="0 0 0.05044585987261141", label="27.35",style=bold];
"DMBT1_M" -> "APOB_E" [color="0 0 0.04987261146496802", label="27.60",style=bold];
"GABRA1_E" -> "NDST4_E" [color="0 0 0.04929936305732474", label="28.12",style=bold];
"KPRP_M" -> "FLG_M" [color="0 0 0.04872611464968146", label="28.16",style=bold];
"MUC17_E" -> "MUC2_E" [color="0 0 0.048152866242038184", label="28.17",style=bold];
"RIMS1_E" -> "LHCGR_E" [color="0 0 0.047579617834394794", label="28.62",style=bold];
"LRRIQ1_M" -> "LRRIQ1_E" [color="0 0 0.047006369426751515", label="28.67",style=bold];
"PCDHA1_M" -> "TPTE_M" [color="0 0 0.04643312101910824", label="28.98",style=bold];
"KRT78_M" -> "KRT6C_M" [color="0 0 0.04585987261146485", label="29.05",style=bold];
"APOB_E" -> "PCDHA13_E" [color="0 0 0.04528662420382157", label="29.35",style=bold];
"DPP6_M" -> "CNTNAP5_M" [color="0 0 0.04471337579617829", label="29.41",style=bold];
"CSMD1_M" -> "GABRA1_M" [color="0 0 0.0441401273885349", label="29.43",style=bold];
"FBN3_E" -> "FREM2_E" [color="0 0 0.04356687898089162", label="29.49",style=bold];
"ZNF716_M" -> "UNC5D_M" [color="0 0 0.04299363057324834", label="29.53",style=bold];
"CHRNA4_M" -> "PCDHA1_M" [color="0 0 0.042420382165605064", label="29.66",style=bold];
"RIMS1_E" -> "FREM2_E" [color="0 0 0.041847133757961674", label="29.69",style=bold];
"ZNF716_M" -> "CNTNAP4_M" [color="0 0 0.041273885350318396", label="29.74",style=bold];
"DPP6_M" -> "ADAMTS20_E" [color="0 0 0.04070063694267512", label="29.86",style=bold];
"FLG_M" -> "TRIML2_M" [color="0 0 0.04012738853503173", label="30.08",style=bold];
"KRT6C_E" -> "SPRR2A_E" [color="0 0 0.03955414012738845", label="30.28",style=bold];
"CFHR1_M" -> "DSG1_M" [color="0 0 0.03898089171974517", label="30.30",style=bold];
"MUC2_M" -> "PNLDC1_M" [color="0 0 0.03840764331210178", label="30.85",style=bold];
"LIPF_E" -> "NELL1_E" [color="0 0 0.0378343949044585", label="31.00",style=bold];
"SLC1A6_E" -> "ZAN_E" [color="0 0 0.03726114649681522", label="31.50",style=bold];
"LHCGR_M" -> "UNC5D_M" [color="0 0 0.036687898089171944", label="32.01",style=bold];
"TRPM3_E" -> "DCT_E" [color="0 0 0.036114649681528554", label="32.10",style=bold];
"SLC1A6_E" -> "LRRC7_E" [color="0 0 0.035541401273885276", label="32.30",style=bold];
"MUC17_E" -> "MUC17_M" [color="0 0 0.034968152866242", label="34.72",style=bold];
"ADH7_M" -> "KCNA10_M" [color="0 0 0.03439490445859861", label="35.61",style=bold];
"SBSN_M" -> "FBN3_M" [color="0 0 0.03382165605095533", label="35.94",style=bold];
"CSMD1_M" -> "KPRP_M" [color="0 0 0.03324840764331205", label="36.37",style=bold];
"LRRC7_E" -> "DPP6_E" [color="0 0 0.03267515923566866", label="36.45",style=bold];
"KCNA10_M" -> "DMBT1_M" [color="0 0 0.03210191082802538", label="36.92",style=bold];
"UNC5D_M" -> "KRT6C_M" [color="0 0 0.0315286624203821", label="37.00",style=bold];
"SBSN_M" -> "KCNA10_M" [color="0 0 0.030955414012738824", label="37.02",style=bold];
"KRT6C_E" -> "PGC_E" [color="0 0 0.030382165605095435", label="37.25",style=bold];
"SERPINB13_E" -> "ADAMTS20_E" [color="0 0 0.029808917197452156", label="37.30",style=bold];
"PGC_E" -> "DMBT1_E" [color="0 0 0.029235668789808877", label="37.60",style=bold];
"CNTNAP5_E" -> "NELL1_E" [color="0 0 0.028662420382165488", label="38.53",style=bold];
"IRX4_M" -> "LOC100190940_M" [color="0 0 0.02808917197452221", label="38.70",style=bold];
"LHCGR_M" -> "RPL10L_M" [color="0 0 0.02751592356687893", label="38.81",style=bold];
"RIMS1_E" -> "APOB_E" [color="0 0 0.02694267515923554", label="38.88",style=bold];
"PNLIPRP3_E" -> "PAX1_E" [color="0 0 0.026369426751592262", label="39.00",style=bold];
"GABRA1_M" -> "GABRG2_M" [color="0 0 0.025796178343948983", label="39.10",style=bold];
"ADAMTS20_E" -> "GPR98_E" [color="0 0 0.025222929936305705", label="39.61",style=bold];
"A2ML1_M" -> "TMPRSS11A_M" [color="0 0 0.024649681528662315", label="42.09",style=bold];
"PAX1_M" -> "MYBPC1_M" [color="0 0 0.024076433121019036", label="42.28",style=bold];
"CNTNAP5_M" -> "LHCGR_M" [color="0 0 0.023503184713375758", label="42.55",style=bold];
"IRX4_E" -> "ADH7_M" [color="0 0 0.022929936305732368", label="42.89",style=bold];
"MUC4_M" -> "FBN3_M" [color="0 0 0.02235668789808909", label="42.96",style=bold];
"MUC2_M" -> "MUC5B_M" [color="0 0 0.02178343949044581", label="43.13",style=bold];
"LHCGR_M" -> "SPRR2A_M" [color="0 0 0.02121019108280242", label="43.49",style=bold];
"LIN28B_E" -> "KCNV1_E" [color="0 0 0.020636942675159142", label="43.75",style=bold];
"DPP6_E" -> "RIMS1_E" [color="0 0 0.020063694267515864", label="45.06",style=bold];
"KRT78_M" -> "DSG1_M" [color="0 0 0.019490445859872474", label="45.72",style=bold];
"LIN28B_E" -> "CNTNAP4_E" [color="0 0 0.018917197452229195", label="46.27",style=bold];
"ADAMTS20_E" -> "ZIC4_E" [color="0 0 0.018343949044585917", label="46.66",style=bold];
"KRT1_M" -> "KRT78_M" [color="0 0 0.017770700636942638", label="46.96",style=bold];
"A2ML1_E" -> "SPRR2A_M" [color="0 0 0.017197452229299248", label="47.14",style=bold];
"DMBT1_E" -> "MUC17_E" [color="0 0 0.01662420382165597", label="47.94",style=bold];
"CSMD3_M" -> "HTR4_M" [color="0 0 0.01605095541401269", label="48.26",style=bold];
"SOX1_M" -> "FOXG1_M" [color="0 0 0.015477707006369301", label="50.92",style=bold];
"RIMS1_M" -> "CSMD3_M" [color="0 0 0.014904458598726023", label="51.10",style=bold];
"TPTE_M" -> "RPL10L_M" [color="0 0 0.014331210191082744", label="52.68",style=bold];
"BNC1_M" -> "SOX1_E" [color="0 0 0.013757961783439354", label="55.69",style=bold];
"KRT1_M" -> "MYO18B_M" [color="0 0 0.013184713375796075", label="56.17",style=bold];
"SERPINB13_E" -> "IRX4_E" [color="0 0 0.012611464968152797", label="59.10",style=bold];
"DPP6_M" -> "TPTE_M" [color="0 0 0.012038216560509518", label="61.22",style=bold];
"SERPINB13_E" -> "SPAG17_E" [color="0 0 0.011464968152866128", label="64.37",style=bold];
"MUC2_M" -> "MUC4_M" [color="0 0 0.01089171974522285", label="67.19",style=bold];
"PGC_E" -> "MUC6_E" [color="0 0 0.010318471337579571", label="67.31",style=bold];
"CWH43_M" -> "NELL1_M" [color="0 0 0.009745222929936181", label="70.79",style=bold];
"ADAMTS20_M" -> "ZIC4_M" [color="0 0 0.009171974522292903", label="70.98",style=bold];
"PCDHA1_M" -> "PCDHA13_M" [color="0 0 0.008598726114649624", label="76.83",style=bold];
"GABRA1_M" -> "CSMD3_M" [color="0 0 0.008025477707006234", label="78.15",style=bold];
"TMPRSS11A_E" -> "KRT24_E" [color="0 0 0.007452229299362956", label="81.56",style=bold];
"ZNF716_M" -> "MUC16_M" [color="0 0 0.006878980891719677", label="85.92",style=bold];
"SOX1_M" -> "IRX4_M" [color="0 0 0.006305732484076398", label="93.75",style=bold];
"IVL_E" -> "DSG1_E" [color="0 0 0.005732484076433009", label="97.79",style=bold];
"PGC_E" -> "LIPF_E" [color="0 0 0.00515923566878973", label="99.67",style=bold];
"PCDHA1_M" -> "SOX1_M" [color="0 0 0.004585987261146451", label="100.35",style=bold];
"IVL_M" -> "KPRP_M" [color="0 0 0.004012738853503062", label="107.58",style=bold];
"SPRR2A_M" -> "IVL_M" [color="0 0 0.003439490445859783", label="127.72",style=bold];
"SLC1A6_M" -> "DPP6_M" [color="0 0 0.0028662420382165044", label="139.69",style=bold];
"IVL_M" -> "KRT1_M" [color="0 0 0.0022929936305731147", label="140.65",style=bold];
"SLC1A6_M" -> "CHRNA4_M" [color="0 0 0.001719745222929836", label="159.82",style=bold];
"IVL_E" -> "KPRP_E" [color="0 0 0.0011464968152865573", label="162.75",style=bold];
"IVL_E" -> "KRT6C_E" [color="0 0 0.0005732484076432787", label="175.61",style=bold];
"SERPINB13_E" -> "IVL_E" [color="0 0 0.0", label="203.22",style=bold];
}
```
