## Supplementary material for "New analysis framework incorporating mixed mutual information and scalable Bayesian networks for multimodal high dimensional genomic and epigenomic cancer data": Suppl Table 12

```
digraph G{
 ratio=fill;
 node [shape=box, style=rounded];
 edge [arrowhead=none];
"A2ML1_S";
"A2ML1_E";
"A2ML1_M";
"ABCA12_S";
"ABCA12_E";
"ABCA12_M";
"ABCA13_S";
"ABCA13_E";
"ABCA13_M";
"ACTN2_S";
"ACTN2_E";
"ACTN2_M";
"ADAMTS20_S";
"ADAMTS20_E";
"ADAMTS20_M";
"AGBL1_S";
"AGBL1_E";
"AGBL1_M";
"APOB_S";
"APOB_E";
"APOB_M";
"C20orf114_S";
"C20orf114_E";
"C20orf114_M";
"CACNA1B_S";
"CACNA1B_E";
"CACNA1B_M";
"CNTN1_S";
"CNTN1_E";
"CNTN1_M";
"CNTNAP2_S";
"CNTNAP2_E";
"CNTNAP2_M";
"COL11A1_S";
"COL11A1_E";
"COL11A1_M";
"COL2A1_S";
"COL2A1_E";
"COL2A1_M";
"CPA2_S";
"CPA2_E";
"CPA2_M";
"CR2_S";
"CR2_E";
"CR2_M";
"CSMD3_S";
"CSMD3_E";
"CSMD3_M";
"CTSE_S";
"CTSE_E";
"CTSE_M";
"CWH43_S";
"CWH43_E";
"CWH43_M";
"DCDC1_S";
"DCDC1_E";
"DCDC1_M";
"DMBT1_S";
"DMBT1_E";
"DMBT1_M";
"DPP6_S";
"DPP6_E";
"DPP6_M";
"DSG1_S";
"DSG1_E";
"DSG1_M";
"EGF_S";
"EGF_E";
"EGF_M";
"FAM83C_S";
"FAM83C_E";
"FAM83C_M";
"FBN3_S";
"FBN3_E";
"FBN3_M";
"FER1L6_S";
"FER1L6_E";
"FER1L6_M";
"FLG_S";
"FLG_E";
"FLG_M";
"FREM2_S";
"FREM2_E";
"FREM2_M";
"FRMPD2_S";
"FRMPD2_E";
"FRMPD2_M";
"FSTL5_S";
"FSTL5_E";
"FSTL5_M";
"GLRA1_S";
"GLRA1_E";
"GLRA1_M";
"GPR87_S";
"GPR87_E";
"GPR87_M";
"GPR98_S";
"GPR98_E";
"GPR98_M";
"IRX6_S";
"IRX6_E";
"IRX6_M";
"KCNA1_S";
"KCNA1_E";
"KCNA1_M";
"KLK5_S";
"KLK5_E";
"KLK5_M";
"KRT13_S";
"KRT13_E";
"KRT13_M";
"KRT16_S";
"KRT16_E";
"KRT16_M";
"KRT6B_S";
"KRT6B_E";
"KRT6B_M";
"KRT6C_S";
"KRT6C_E";
"KRT6C_M";
"LPA_S";
"LPA_E";
"LPA_M";
"LRRC7_S";
"LRRC7_E";
"LRRC7_M";
"MDGA2_S";
"MDGA2_E";
"MDGA2_M";
"MUC15_S";
"MUC15_E";
"MUC15_M";
"MUC16_S";
"MUC16_E";
"MUC16_M";
"MUC17_S";
"MUC17_E";
"MUC17_M";
"MUC2_S";
"MUC2_E";
"MUC2_M";
"MUC4_S";
"MUC4_E";
"MUC4_M";
"MUC5B_S";
"MUC5B_E";
"MUC5B_M";
"MUC6_S";
"MUC6_E";
"MUC6_M";
"MYH4_S";
"MYH4_E";
"MYH4_M";
"NETO1_S";
"NETO1_E";
"NETO1_M";
"OGDHL_S";
"OGDHL_E";
"OGDHL_M";
"PCDH10_S";
"PCDH10_E";
"PCDH10_M";
"PCDHA11_S";
"PCDHA11_E";
"PCDHA11_M";
"PCDHA12_S";
"PCDHA12_E";
"PCDHA12_M";
"PCK1_S";
"PCK1_E";
"PCK1_M";
"PCLO_S";
"PCLO_E";
"PCLO_M";
"PGC_S";
"PGC_E";
"PGC_M";
"PGLYRP3_S";
"PGLYRP3_E";
"PGLYRP3_M";
"PKHD1L1_S";
"PKHD1L1_E";
"PKHD1L1_M";
"PNLIPRP2_S";
"PNLIPRP2_E";
"PNLIPRP2_M";
"PTPRZ1_S";
"PTPRZ1_E";
"PTPRZ1_M";
"RFX6_S";
"RFX6_E";
"RFX6_M";
"RIMS2_S";
"RIMS2_E";
"RIMS2_M";
"RP1_S";
"RP1_E";
"RP1_M";
"S100A7_S";
"S100A7_E";
"S100A7_M";
"SERPINB13_S";
"SERPINB13_E";
"SERPINB13_M";
"SERPINB3_S";
"SERPINB3_E";
"SERPINB3_M";
"SERPINB4_S";
"SERPINB4_E";
"SERPINB4_M";
"SLC13A2_S";
"SLC13A2_E";
"SLC13A2_M";
"SLC26A3_S";
"SLC26A3_E";
"SLC26A3_M";
"SLC6A19_S";
"SLC6A19_E";
"SLC6A19_M";
"SLC9A11_S";
"SLC9A11_E";
"SLC9A11_M";
"SLC9A4_S";
"SLC9A4_E";
"SLC9A4_M";
"SLCO1B1_S";
"SLCO1B1_E";
"SLCO1B1_M";
"SLCO1B3_S";
"SLCO1B3_E";
"SLCO1B3_M";
"SLITRK1_S";
"SLITRK1_E";
"SLITRK1_M";
"TGM5_S";
"TGM5_E";
"TGM5_M";
"THBS4_S";
"THBS4_E";
"THBS4_M";
"ZIC4_S";
"ZIC4_E";
"ZIC4_M";
"Survival";
"TGM5_S" -> "TGM5_E" [color="0 0 0.9", label="0.00",style=bold];
"PGC_S" -> "OGDHL_M" [color="0 0 0.8994382022471911", label="0.00",style=bold];
"KRT16_S" -> "LPA_E" [color="0 0 0.898876404494382", label="0.01",style=bold];
"SERPINB3_S" -> "DSG1_E" [color="0 0 0.898314606741573", label="0.01",style=bold];
"A2ML1_S" -> "PCLO_M" [color="0 0 0.8977528089887641", label="0.01",style=bold];
"PNLIPRP2_S" -> "APOB_S" [color="0 0 0.8971910112359551", label="0.01",style=bold];
"PGC_S" -> "KCNA1_M" [color="0 0 0.8966292134831461", label="0.01",style=bold];
"SERPINB3_S" -> "DSG1_M" [color="0 0 0.8960674157303371", label="0.02",style=bold];
"SLC13A2_S" -> "TGM5_E" [color="0 0 0.8955056179775281", label="0.02",style=bold];
"SLC13A2_S" -> "MUC5B_E" [color="0 0 0.8949438202247191", label="0.02",style=bold];
"MUC15_S" -> "ABCA12_M" [color="0 0 0.8943820224719101", label="0.03",style=bold];
"PGLYRP3_S" -> "KRT6C_M" [color="0 0 0.8938202247191012", label="0.03",style=bold];
"SLC26A3_S" -> "S100A7_M" [color="0 0 0.8932584269662922", label="0.03",style=bold];
"MUC2_S" -> "RFX6_S" [color="0 0 0.8926966292134831", label="0.04",style=bold];
"GPR87_S" -> "MYH4_S" [color="0 0 0.8921348314606742", label="0.04",style=bold];
"KRT6C_S" -> "APOB_E" [color="0 0 0.8915730337078652", label="0.04",style=bold];
"S100A7_S" -> "MUC2_M" [color="0 0 0.8910112359550562", label="0.04",style=bold];
"TGM5_S" -> "CR2_E" [color="0 0 0.8904494382022472", label="0.04",style=bold];
"CPA2_S" -> "CNTN1_E" [color="0 0 0.8898876404494382", label="0.04",style=bold];
"OGDHL_S" -> "MUC6_E" [color="0 0 0.8893258426966293", label="0.04",style=bold];
"CWH43_S" -> "MUC4_E" [color="0 0 0.8887640449438202", label="0.04",style=bold];
"SERPINB3_S" -> "ADAMTS20_S" [color="0 0 0.8882022471910113", label="0.04",style=bold];
"TGM5_S" -> "CTSE_E" [color="0 0 0.8876404494382023", label="0.04",style=bold];
"PCDHA12_S" -> "RFX6_M" [color="0 0 0.8870786516853932", label="0.04",style=bold];
"SERPINB3_S" -> "CR2_E" [color="0 0 0.8865168539325843", label="0.04",style=bold];
"GLRA1_S" -> "PKHD1L1_E" [color="0 0 0.8859550561797753", label="0.04",style=bold];
"CPA2_S" -> "RIMS2_M" [color="0 0 0.8853932584269664", label="0.05",style=bold];
"CTSE_S" -> "SLC6A19_M" [color="0 0 0.8848314606741573", label="0.05",style=bold];
"GLRA1_S" -> "THBS4_E" [color="0 0 0.8842696629213483", label="0.05",style=bold];
"PGLYRP3_S" -> "MYH4_E" [color="0 0 0.8837078651685394", label="0.05",style=bold];
"DSG1_S" -> "ABCA12_S" [color="0 0 0.8831460674157303", label="0.05",style=bold];
"SLC13A2_S" -> "C20orf114_M" [color="0 0 0.8825842696629214", label="0.06",style=bold];
"LRRC7_S" -> "MUC16_M" [color="0 0 0.8820224719101124", label="0.06",style=bold];
"C20orf114_S" -> "PNLIPRP2_E" [color="0 0 0.8814606741573034", label="0.06",style=bold];
"FRMPD2_S" -> "SERPINB4_M" [color="0 0 0.8808988764044944", label="0.06",style=bold];
"KRT16_S" -> "SLCO1B1_M" [color="0 0 0.8803370786516854", label="0.06",style=bold];
"KRT6B_S" -> "EGF_E" [color="0 0 0.8797752808988765", label="0.06",style=bold];
"TGM5_S" -> "CTSE_M" [color="0 0 0.8792134831460674", label="0.06",style=bold];
"C20orf114_S" -> "KRT6B_M" [color="0 0 0.8786516853932584", label="0.06",style=bold];
"CPA2_S" -> "DSG1_M" [color="0 0 0.8780898876404495", label="0.06",style=bold];
"CPA2_S" -> "SLC9A4_M" [color="0 0 0.8775280898876404", label="0.06",style=bold];
"C20orf114_S" -> "MUC6_E" [color="0 0 0.8769662921348315", label="0.06",style=bold];
"KRT6B_S" -> "SLCO1B3_E" [color="0 0 0.8764044943820225", label="0.06",style=bold];
"KRT16_S" -> "SLC13A2_M" [color="0 0 0.8758426966292135", label="0.06",style=bold];
"CWH43_S" -> "MUC5B_S" [color="0 0 0.8752808988764045", label="0.07",style=bold];
"KRT6B_S" -> "MDGA2_S" [color="0 0 0.8747191011235955", label="0.07",style=bold];
"MUC15_S" -> "MUC2_M" [color="0 0 0.8741573033707866", label="0.07",style=bold];
"DSG1_S" -> "KRT13_M" [color="0 0 0.8735955056179775", label="0.08",style=bold];
"CPA2_S" -> "FER1L6_E" [color="0 0 0.8730337078651685", label="0.08",style=bold];
"FAM83C_S" -> "MUC15_M" [color="0 0 0.8724719101123596", label="0.08",style=bold];
"SLC13A2_S" -> "SLCO1B1_E" [color="0 0 0.8719101123595506", label="0.09",style=bold];
"SERPINB13_S" -> "NETO1_E" [color="0 0 0.8713483146067416", label="0.09",style=bold];
"PGC_S" -> "MUC6_M" [color="0 0 0.8707865168539326", label="0.09",style=bold];
"CPA2_S" -> "MUC15_E" [color="0 0 0.8702247191011236", label="0.09",style=bold];
"GPR87_S" -> "PCK1_M" [color="0 0 0.8696629213483146", label="0.09",style=bold];
"C20orf114_S" -> "SLCO1B1_M" [color="0 0 0.8691011235955056", label="0.10",style=bold];
"PGC_S" -> "SLCO1B1_M" [color="0 0 0.8685393258426967", label="0.10",style=bold];
"GLRA1_S" -> "PCDHA11_E" [color="0 0 0.8679775280898877", label="0.10",style=bold];
"CNTN1_S" -> "SLCO1B1_M" [color="0 0 0.8674157303370786", label="0.10",style=bold];
"CTSE_S" -> "GPR87_M" [color="0 0 0.8668539325842697", label="0.10",style=bold];
"GLRA1_S" -> "SLC9A11_E" [color="0 0 0.8662921348314607", label="0.10",style=bold];
"GPR87_S" -> "OGDHL_E" [color="0 0 0.8657303370786517", label="0.11",style=bold];
"CTSE_S" -> "MYH4_M" [color="0 0 0.8651685393258427", label="0.11",style=bold];
"KLK5_S" -> "SLC6A19_M" [color="0 0 0.8646067415730337", label="0.11",style=bold];
"CTSE_S" -> "KRT6B_M" [color="0 0 0.8640449438202248", label="0.11",style=bold];
"GLRA1_S" -> "CPA2_M" [color="0 0 0.8634831460674157", label="0.11",style=bold];
"SLC26A3_S" -> "RP1_E" [color="0 0 0.8629213483146068", label="0.11",style=bold];
"PGLYRP3_S" -> "MUC16_E" [color="0 0 0.8623595505617978", label="0.11",style=bold];
"PNLIPRP2_S" -> "PCDHA11_E" [color="0 0 0.8617977528089887", label="0.11",style=bold];
"CWH43_S" -> "PCDHA12_S" [color="0 0 0.8612359550561798", label="0.11",style=bold];
"KLK5_S" -> "KRT13_M" [color="0 0 0.8606741573033708", label="0.11",style=bold];
"PGC_S" -> "APOB_M" [color="0 0 0.8601123595505619", label="0.11",style=bold];
"KRT16_S" -> "COL2A1_S" [color="0 0 0.8595505617977528", label="0.11",style=bold];
"PGLYRP3_S" -> "ABCA13_M" [color="0 0 0.8589887640449438", label="0.11",style=bold];
"C20orf114_S" -> "SLC6A19_E" [color="0 0 0.8584269662921349", label="0.11",style=bold];
"CWH43_S" -> "TGM5_M" [color="0 0 0.8578651685393258", label="0.11",style=bold];
"SERPINB3_S" -> "KCNA1_E" [color="0 0 0.8573033707865169", label="0.11",style=bold];
"KCNA1_S" -> "CTSE_S" [color="0 0 0.8567415730337079", label="0.12",style=bold];
"CNTN1_S" -> "RIMS2_S" [color="0 0 0.8561797752808988", label="0.12",style=bold];
"SLC9A11_S" -> "THBS4_S" [color="0 0 0.8556179775280899", label="0.12",style=bold];
"SLC6A19_S" -> "MUC5B_S" [color="0 0 0.8550561797752809", label="0.12",style=bold];
"SLITRK1_S" -> "ACTN2_E" [color="0 0 0.854494382022472", label="0.12",style=bold];
"PNLIPRP2_S" -> "ZIC4_S" [color="0 0 0.8539325842696629", label="0.12",style=bold];
"SERPINB3_S" -> "FAM83C_M" [color="0 0 0.8533707865168539", label="0.12",style=bold];
"FER1L6_S" -> "SLC9A11_M" [color="0 0 0.852808988764045", label="0.12",style=bold];
"SLC13A2_S" -> "ACTN2_M" [color="0 0 0.8522471910112359", label="0.12",style=bold];
"NETO1_S" -> "RFX6_E" [color="0 0 0.851685393258427", label="0.12",style=bold];
"KRT6B_S" -> "EGF_M" [color="0 0 0.851123595505618", label="0.13",style=bold];
"FER1L6_S" -> "IRX6_S" [color="0 0 0.850561797752809", label="0.13",style=bold];
"PGLYRP3_S" -> "PCDHA12_E" [color="0 0 0.85", label="0.13",style=bold];
"KRT16_S" -> "FREM2_S" [color="0 0 0.849438202247191", label="0.13",style=bold];
"GLRA1_S" -> "FLG_S" [color="0 0 0.8488764044943821", label="0.13",style=bold];
"GLRA1_S" -> "MUC15_M" [color="0 0 0.848314606741573", label="0.13",style=bold];
"GLRA1_S" -> "FAM83C_M" [color="0 0 0.847752808988764", label="0.13",style=bold];
"FRMPD2_S" -> "FBN3_M" [color="0 0 0.8471910112359551", label="0.14",style=bold];
"GPR87_S" -> "FRMPD2_M" [color="0 0 0.8466292134831461", label="0.14",style=bold];
"FAM83C_S" -> "GPR87_M" [color="0 0 0.8460674157303371", label="0.14",style=bold];
"C20orf114_S" -> "EGF_S" [color="0 0 0.8455056179775281", label="0.14",style=bold];
"COL2A1_S" -> "GPR98_E" [color="0 0 0.8449438202247191", label="0.15",style=bold];
"MDGA2_S" -> "GLRA1_E" [color="0 0 0.8443820224719101", label="0.15",style=bold];
"AGBL1_S" -> "FBN3_M" [color="0 0 0.8438202247191011", label="0.15",style=bold];
"SERPINB13_S" -> "FRMPD2_E" [color="0 0 0.8432584269662922", label="0.15",style=bold];
"KRT13_S" -> "EGF_M" [color="0 0 0.8426966292134832", label="0.15",style=bold];
"CPA2_S" -> "CPA2_M" [color="0 0 0.8421348314606741", label="0.15",style=bold];
"PGC_S" -> "CTSE_E" [color="0 0 0.8415730337078652", label="0.15",style=bold];
"FAM83C_S" -> "LRRC7_E" [color="0 0 0.8410112359550562", label="0.15",style=bold];
"C20orf114_S" -> "MUC5B_E" [color="0 0 0.8404494382022472", label="0.15",style=bold];
"PGC_S" -> "NETO1_M" [color="0 0 0.8398876404494382", label="0.15",style=bold];
"COL2A1_S" -> "SLC9A4_E" [color="0 0 0.8393258426966292", label="0.16",style=bold];
"EGF_S" -> "GPR87_M" [color="0 0 0.8387640449438203", label="0.16",style=bold];
"KRT13_S" -> "MUC6_S" [color="0 0 0.8382022471910112", label="0.16",style=bold];
"PGC_S" -> "PCK1_M" [color="0 0 0.8376404494382023", label="0.17",style=bold];
"KLK5_S" -> "DSG1_E" [color="0 0 0.8370786516853933", label="0.17",style=bold];
"SLC13A2_S" -> "ABCA13_M" [color="0 0 0.8365168539325842", label="0.17",style=bold];
"FAM83C_S" -> "CNTN1_S" [color="0 0 0.8359550561797753", label="0.17",style=bold];
"PGC_S" -> "KRT16_M" [color="0 0 0.8353932584269663", label="0.17",style=bold];
"SERPINB13_S" -> "ADAMTS20_S" [color="0 0 0.8348314606741574", label="0.17",style=bold];
"SLC9A4_S" -> "CACNA1B_E" [color="0 0 0.8342696629213483", label="0.17",style=bold];
"PGC_S" -> "EGF_E" [color="0 0 0.8337078651685393", label="0.17",style=bold];
"SERPINB3_S" -> "KRT6B_M" [color="0 0 0.8331460674157304", label="0.17",style=bold];
"SLC26A3_S" -> "MUC17_S" [color="0 0 0.8325842696629213", label="0.18",style=bold];
"TGM5_S" -> "KRT16_M" [color="0 0 0.8320224719101124", label="0.18",style=bold];
"MUC15_S" -> "OGDHL_E" [color="0 0 0.8314606741573034", label="0.18",style=bold];
"SERPINB4_S" -> "CNTN1_S" [color="0 0 0.8308988764044944", label="0.19",style=bold];
"FAM83C_S" -> "GLRA1_E" [color="0 0 0.8303370786516854", label="0.19",style=bold];
"KRT13_S" -> "SLC9A4_E" [color="0 0 0.8297752808988764", label="0.19",style=bold];
"S100A7_S" -> "A2ML1_E" [color="0 0 0.8292134831460675", label="0.19",style=bold];
"A2ML1_S" -> "SLC26A3_E" [color="0 0 0.8286516853932584", label="0.20",style=bold];
"KRT6B_S" -> "AGBL1_M" [color="0 0 0.8280898876404494", label="0.20",style=bold];
"KRT16_S" -> "ABCA12_E" [color="0 0 0.8275280898876405", label="0.20",style=bold];
"AGBL1_S" -> "APOB_E" [color="0 0 0.8269662921348315", label="0.21",style=bold];
"C20orf114_S" -> "SERPINB4_M" [color="0 0 0.8264044943820225", label="0.22",style=bold];
"DSG1_S" -> "FBN3_M" [color="0 0 0.8258426966292135", label="0.22",style=bold];
"SERPINB13_S" -> "SLC6A19_M" [color="0 0 0.8252808988764045", label="0.22",style=bold];
"KRT6C_S" -> "ADAMTS20_E" [color="0 0 0.8247191011235955", label="0.22",style=bold];
"S100A7_S" -> "SLC9A11_S" [color="0 0 0.8241573033707865", label="0.22",style=bold];
"S100A7_S" -> "SERPINB13_S" [color="0 0 0.8235955056179776", label="0.22",style=bold];
"S100A7_S" -> "MUC17_E" [color="0 0 0.8230337078651686", label="0.22",style=bold];
"S100A7_S" -> "MYH4_E" [color="0 0 0.8224719101123595", label="0.22",style=bold];
"SERPINB3_S" -> "CPA2_E" [color="0 0 0.8219101123595506", label="0.22",style=bold];
"FAM83C_S" -> "FLG_S" [color="0 0 0.8213483146067416", label="0.23",style=bold];
"CWH43_S" -> "CWH43_M" [color="0 0 0.8207865168539326", label="0.23",style=bold];
"SERPINB3_S" -> "ABCA12_S" [color="0 0 0.8202247191011236", label="0.23",style=bold];
"PGLYRP3_S" -> "COL11A1_E" [color="0 0 0.8196629213483146", label="0.23",style=bold];
"GLRA1_S" -> "S100A7_M" [color="0 0 0.8191011235955057", label="0.23",style=bold];
"PGC_S" -> "FRMPD2_M" [color="0 0 0.8185393258426966", label="0.24",style=bold];
"PNLIPRP2_S" -> "ACTN2_E" [color="0 0 0.8179775280898877", label="0.24",style=bold];
"CWH43_S" -> "SLC9A11_M" [color="0 0 0.8174157303370787", label="0.24",style=bold];
"MUC15_S" -> "MUC16_S" [color="0 0 0.8168539325842696", label="0.24",style=bold];
"CWH43_S" -> "SERPINB3_M" [color="0 0 0.8162921348314607", label="0.25",style=bold];
"PGC_S" -> "MUC17_M" [color="0 0 0.8157303370786517", label="0.25",style=bold];
"KRT6B_S" -> "FER1L6_E" [color="0 0 0.8151685393258428", label="0.25",style=bold];
"A2ML1_S" -> "FREM2_E" [color="0 0 0.8146067415730337", label="0.25",style=bold];
"SERPINB3_S" -> "SERPINB3_M" [color="0 0 0.8140449438202247", label="0.26",style=bold];
"CTSE_S" -> "FREM2_M" [color="0 0 0.8134831460674158", label="0.26",style=bold];
"SERPINB3_S" -> "MYH4_M" [color="0 0 0.8129213483146067", label="0.26",style=bold];
"SERPINB4_S" -> "CTSE_E" [color="0 0 0.8123595505617978", label="0.26",style=bold];
"FER1L6_S" -> "FREM2_E" [color="0 0 0.8117977528089888", label="0.27",style=bold];
"GPR87_S" -> "APOB_M" [color="0 0 0.8112359550561798", label="0.27",style=bold];
"GLRA1_S" -> "SERPINB13_M" [color="0 0 0.8106741573033708", label="0.28",style=bold];
"KRT6B_S" -> "MUC4_E" [color="0 0 0.8101123595505618", label="0.28",style=bold];
"KRT13_S" -> "CSMD3_E" [color="0 0 0.8095505617977529", label="0.28",style=bold];
"THBS4_S" -> "CSMD3_S" [color="0 0 0.8089887640449438", label="0.28",style=bold];
"CNTN1_S" -> "SLCO1B3_S" [color="0 0 0.8084269662921348", label="0.28",style=bold];
"MUC2_S" -> "CSMD3_S" [color="0 0 0.8078651685393259", label="0.29",style=bold];
"KLK5_S" -> "SLC26A3_E" [color="0 0 0.8073033707865169", label="0.29",style=bold];
"GPR87_S" -> "CTSE_E" [color="0 0 0.8067415730337079", label="0.29",style=bold];
"GPR87_S" -> "SLC13A2_E" [color="0 0 0.8061797752808989", label="0.29",style=bold];
"S100A7_S" -> "RIMS2_E" [color="0 0 0.80561797752809", label="0.29",style=bold];
"KRT6B_S" -> "RIMS2_M" [color="0 0 0.8050561797752809", label="0.29",style=bold];
"S100A7_S" -> "CWH43_M" [color="0 0 0.8044943820224719", label="0.29",style=bold];
"PNLIPRP2_S" -> "CPA2_M" [color="0 0 0.803932584269663", label="0.29",style=bold];
"SLC26A3_S" -> "CPA2_M" [color="0 0 0.803370786516854", label="0.29",style=bold];
"TGM5_S" -> "COL11A1_E" [color="0 0 0.8028089887640449", label="0.29",style=bold];
"SERPINB13_S" -> "DCDC1_M" [color="0 0 0.802247191011236", label="0.29",style=bold];
"CPA2_S" -> "RFX6_M" [color="0 0 0.801685393258427", label="0.29",style=bold];
"CPA2_S" -> "PNLIPRP2_M" [color="0 0 0.801123595505618", label="0.29",style=bold];
"FAM83C_S" -> "MUC2_E" [color="0 0 0.800561797752809", label="0.29",style=bold];
"GPR87_S" -> "SLCO1B1_M" [color="0 0 0.8", label="0.29",style=bold];
"GPR87_S" -> "EGF_M" [color="0 0 0.799438202247191", label="0.29",style=bold];
"CTSE_S" -> "MUC6_E" [color="0 0 0.798876404494382", label="0.29",style=bold];
"SLC13A2_S" -> "RP1_E" [color="0 0 0.7983146067415731", label="0.29",style=bold];
"PGC_S" -> "RIMS2_M" [color="0 0 0.7977528089887641", label="0.29",style=bold];
"KRT6C_S" -> "RP1_E" [color="0 0 0.797191011235955", label="0.29",style=bold];
"KRT16_S" -> "KCNA1_S" [color="0 0 0.7966292134831461", label="0.29",style=bold];
"SERPINB13_S" -> "SLCO1B3_M" [color="0 0 0.7960674157303371", label="0.29",style=bold];
"SLC13A2_S" -> "A2ML1_E" [color="0 0 0.7955056179775281", label="0.29",style=bold];
"CNTN1_S" -> "RP1_E" [color="0 0 0.7949438202247191", label="0.29",style=bold];
"SLC6A19_S" -> "AGBL1_S" [color="0 0 0.7943820224719101", label="0.30",style=bold];
"RFX6_S" -> "MDGA2_E" [color="0 0 0.7938202247191011", label="0.30",style=bold];
"SERPINB4_S" -> "ZIC4_E" [color="0 0 0.7932584269662921", label="0.31",style=bold];
"KRT6C_S" -> "FRMPD2_M" [color="0 0 0.7926966292134832", label="0.31",style=bold];
"KCNA1_S" -> "MYH4_S" [color="0 0 0.7921348314606742", label="0.31",style=bold];
"SERPINB4_S" -> "APOB_S" [color="0 0 0.7915730337078651", label="0.32",style=bold];
"EGF_S" -> "PKHD1L1_M" [color="0 0 0.7910112359550562", label="0.32",style=bold];
"PGLYRP3_S" -> "RFX6_E" [color="0 0 0.7904494382022472", label="0.32",style=bold];
"S100A7_S" -> "PCDHA11_S" [color="0 0 0.7898876404494382", label="0.32",style=bold];
"SERPINB13_S" -> "SLC9A11_M" [color="0 0 0.7893258426966292", label="0.32",style=bold];
"GLRA1_S" -> "MUC2_E" [color="0 0 0.7887640449438202", label="0.32",style=bold];
"SERPINB13_S" -> "PCLO_S" [color="0 0 0.7882022471910113", label="0.32",style=bold];
"A2ML1_S" -> "CR2_S" [color="0 0 0.7876404494382022", label="0.33",style=bold];
"PCK1_S" -> "PKHD1L1_S" [color="0 0 0.7870786516853933", label="0.33",style=bold];
"DCDC1_S" -> "CPA2_M" [color="0 0 0.7865168539325843", label="0.33",style=bold];
"PGC_S" -> "ACTN2_E" [color="0 0 0.7859550561797752", label="0.33",style=bold];
"KLK5_S" -> "SLCO1B3_M" [color="0 0 0.7853932584269663", label="0.33",style=bold];
"FAM83C_S" -> "MUC16_E" [color="0 0 0.7848314606741573", label="0.33",style=bold];
"FRMPD2_S" -> "DCDC1_M" [color="0 0 0.7842696629213484", label="0.34",style=bold];
"SLC9A4_S" -> "FREM2_E" [color="0 0 0.7837078651685393", label="0.34",style=bold];
"DSG1_S" -> "OGDHL_S" [color="0 0 0.7831460674157303", label="0.34",style=bold];
"SLC9A4_S" -> "PKHD1L1_M" [color="0 0 0.7825842696629214", label="0.34",style=bold];
"GLRA1_S" -> "APOB_E" [color="0 0 0.7820224719101123", label="0.35",style=bold];
"PGC_S" -> "SLCO1B3_E" [color="0 0 0.7814606741573034", label="0.35",style=bold];
"SLC9A11_S" -> "FREM2_E" [color="0 0 0.7808988764044944", label="0.35",style=bold];
"SERPINB3_S" -> "FREM2_M" [color="0 0 0.7803370786516854", label="0.35",style=bold];
"MUC15_S" -> "CNTNAP2_M" [color="0 0 0.7797752808988764", label="0.35",style=bold];
"RIMS2_M" -> "CNTN1_M" [color="0 0 0.7792134831460674", label="0.35",style=bold];
"SERPINB3_M" -> "CSMD3_S" [color="0 0 0.7786516853932585", label="0.35",style=bold];
"KRT6B_S" -> "TGM5_M" [color="0 0 0.7780898876404494", label="0.36",style=bold];
"KRT6B_S" -> "GPR87_E" [color="0 0 0.7775280898876404", label="0.36",style=bold];
"C20orf114_S" -> "ABCA12_E" [color="0 0 0.7769662921348315", label="0.36",style=bold];
"KRT16_S" -> "COL2A1_M" [color="0 0 0.7764044943820225", label="0.36",style=bold];
"MUC15_S" -> "DPP6_E" [color="0 0 0.7758426966292135", label="0.36",style=bold];
"KRT16_S" -> "ABCA12_M" [color="0 0 0.7752808988764045", label="0.37",style=bold];
"GPR87_S" -> "FLG_S" [color="0 0 0.7747191011235955", label="0.37",style=bold];
"SERPINB3_S" -> "ABCA12_M" [color="0 0 0.7741573033707865", label="0.37",style=bold];
"PGC_S" -> "MUC16_E" [color="0 0 0.7735955056179775", label="0.38",style=bold];
"SERPINB4_S" -> "MUC2_S" [color="0 0 0.7730337078651686", label="0.38",style=bold];
"SLC9A4_S" -> "RP1_S" [color="0 0 0.7724719101123596", label="0.39",style=bold];
"SERPINB3_S" -> "AGBL1_E" [color="0 0 0.7719101123595505", label="0.39",style=bold];
"C20orf114_S" -> "CNTNAP2_E" [color="0 0 0.7713483146067416", label="0.39",style=bold];
"GPR87_S" -> "SLC6A19_E" [color="0 0 0.7707865168539326", label="0.39",style=bold];
"SLC9A11_S" -> "DCDC1_M" [color="0 0 0.7702247191011236", label="0.39",style=bold];
"ACTN2_S" -> "DCDC1_M" [color="0 0 0.7696629213483146", label="0.40",style=bold];
"KRT13_S" -> "ZIC4_E" [color="0 0 0.7691011235955056", label="0.40",style=bold];
"EGF_S" -> "KRT13_M" [color="0 0 0.7685393258426967", label="0.40",style=bold];
"S100A7_S" -> "FRMPD2_S" [color="0 0 0.7679775280898876", label="0.41",style=bold];
"PGC_S" -> "SLC26A3_E" [color="0 0 0.7674157303370787", label="0.41",style=bold];
"CPA2_S" -> "SLC6A19_M" [color="0 0 0.7668539325842697", label="0.41",style=bold];
"S100A7_S" -> "GLRA1_S" [color="0 0 0.7662921348314606", label="0.41",style=bold];
"S100A7_S" -> "MDGA2_S" [color="0 0 0.7657303370786517", label="0.41",style=bold];
"PGC_S" -> "MUC6_E" [color="0 0 0.7651685393258427", label="0.41",style=bold];
"KRT6C_S" -> "CSMD3_S" [color="0 0 0.7646067415730338", label="0.41",style=bold];
"SLCO1B3_S" -> "SLC26A3_M" [color="0 0 0.7640449438202247", label="0.41",style=bold];
"PGC_S" -> "KRT6B_E" [color="0 0 0.7634831460674157", label="0.41",style=bold];
"PNLIPRP2_S" -> "ACTN2_M" [color="0 0 0.7629213483146068", label="0.41",style=bold];
"CNTN1_S" -> "RP1_S" [color="0 0 0.7623595505617977", label="0.41",style=bold];
"PGC_S" -> "SERPINB3_M" [color="0 0 0.7617977528089888", label="0.41",style=bold];
"KLK5_S" -> "KRT6C_E" [color="0 0 0.7612359550561798", label="0.41",style=bold];
"PNLIPRP2_S" -> "TGM5_M" [color="0 0 0.7606741573033708", label="0.42",style=bold];
"CPA2_S" -> "FAM83C_M" [color="0 0 0.7601123595505618", label="0.42",style=bold];
"PGLYRP3_S" -> "SLCO1B1_M" [color="0 0 0.7595505617977528", label="0.42",style=bold];
"PNLIPRP2_S" -> "KRT6C_M" [color="0 0 0.7589887640449439", label="0.43",style=bold];
"PGLYRP3_S" -> "CR2_E" [color="0 0 0.7584269662921348", label="0.43",style=bold];
"PNLIPRP2_S" -> "SLC26A3_E" [color="0 0 0.7578651685393258", label="0.43",style=bold];
"C20orf114_S" -> "SLC9A4_S" [color="0 0 0.7573033707865169", label="0.43",style=bold];
"EGF_S" -> "OGDHL_E" [color="0 0 0.7567415730337079", label="0.43",style=bold];
"SLCO1B3_S" -> "MUC16_S" [color="0 0 0.7561797752808989", label="0.44",style=bold];
"PGC_S" -> "CPA2_E" [color="0 0 0.7556179775280899", label="0.44",style=bold];
"CWH43_S" -> "MUC15_M" [color="0 0 0.755056179775281", label="0.44",style=bold];
"CPA2_S" -> "MUC4_M" [color="0 0 0.7544943820224719", label="0.44",style=bold];
"GLRA1_S" -> "PTPRZ1_M" [color="0 0 0.7539325842696629", label="0.44",style=bold];
"A2ML1_S" -> "RP1_E" [color="0 0 0.753370786516854", label="0.44",style=bold];
"DSG1_S" -> "MDGA2_S" [color="0 0 0.752808988764045", label="0.44",style=bold];
"SERPINB3_S" -> "CR2_M" [color="0 0 0.7522471910112359", label="0.44",style=bold];
"PGC_S" -> "SLITRK1_E" [color="0 0 0.751685393258427", label="0.45",style=bold];
"SLC13A2_S" -> "MUC6_M" [color="0 0 0.751123595505618", label="0.45",style=bold];
"GLRA1_S" -> "THBS4_M" [color="0 0 0.750561797752809", label="0.45",style=bold];
"SLC9A11_S" -> "LRRC7_E" [color="0 0 0.75", label="0.45",style=bold];
"PNLIPRP2_S" -> "COL11A1_M" [color="0 0 0.749438202247191", label="0.46",style=bold];
"CTSE_S" -> "SLITRK1_E" [color="0 0 0.7488764044943821", label="0.46",style=bold];
"C20orf114_S" -> "PCDH10_E" [color="0 0 0.748314606741573", label="0.46",style=bold];
"SERPINB3_S" -> "MUC16_E" [color="0 0 0.7477528089887641", label="0.46",style=bold];
"COL2A1_S" -> "S100A7_E" [color="0 0 0.7471910112359551", label="0.46",style=bold];
"CPA2_S" -> "AGBL1_M" [color="0 0 0.746629213483146", label="0.46",style=bold];
"A2ML1_S" -> "MUC16_M" [color="0 0 0.7460674157303371", label="0.46",style=bold];
"AGBL1_S" -> "FSTL5_M" [color="0 0 0.7455056179775281", label="0.46",style=bold];
"SERPINB3_S" -> "S100A7_E" [color="0 0 0.7449438202247192", label="0.46",style=bold];
"PCDHA11_S" -> "SLCO1B1_M" [color="0 0 0.7443820224719101", label="0.48",style=bold];
"A2ML1_S" -> "PCDHA11_E" [color="0 0 0.7438202247191011", label="0.48",style=bold];
"DSG1_S" -> "CACNA1B_E" [color="0 0 0.7432584269662921", label="0.48",style=bold];
"PGLYRP3_S" -> "PCDHA12_S" [color="0 0 0.7426966292134831", label="0.49",style=bold];
"KRT13_S" -> "FRMPD2_S" [color="0 0 0.7421348314606742", label="0.49",style=bold];
"CPA2_S" -> "SLCO1B1_M" [color="0 0 0.7415730337078652", label="0.49",style=bold];
"SERPINB3_S" -> "IRX6_E" [color="0 0 0.7410112359550562", label="0.50",style=bold];
"LPA_S" -> "FAM83C_M" [color="0 0 0.7404494382022472", label="0.50",style=bold];
"EGF_S" -> "GPR98_S" [color="0 0 0.7398876404494382", label="0.50",style=bold];
"AGBL1_S" -> "CR2_E" [color="0 0 0.7393258426966292", label="0.50",style=bold];
"KRT13_S" -> "SERPINB13_E" [color="0 0 0.7387640449438202", label="0.51",style=bold];
"SERPINB3_S" -> "CWH43_M" [color="0 0 0.7382022471910112", label="0.51",style=bold];
"GLRA1_S" -> "COL11A1_E" [color="0 0 0.7376404494382023", label="0.51",style=bold];
"SERPINB3_S" -> "DCDC1_M" [color="0 0 0.7370786516853933", label="0.51",style=bold];
"PNLIPRP2_S" -> "ABCA13_M" [color="0 0 0.7365168539325843", label="0.51",style=bold];
"GPR87_S" -> "DCDC1_M" [color="0 0 0.7359550561797753", label="0.51",style=bold];
"GPR87_S" -> "FER1L6_M" [color="0 0 0.7353932584269662", label="0.51",style=bold];
"PGC_S" -> "KLK5_E" [color="0 0 0.7348314606741573", label="0.51",style=bold];
"CTSE_S" -> "SLC9A4_M" [color="0 0 0.7342696629213483", label="0.51",style=bold];
"COL2A1_S" -> "MUC17_S" [color="0 0 0.7337078651685394", label="0.51",style=bold];
"C20orf114_S" -> "KRT13_M" [color="0 0 0.7331460674157304", label="0.51",style=bold];
"MUC15_S" -> "DPP6_M" [color="0 0 0.7325842696629213", label="0.51",style=bold];
"SERPINB3_S" -> "FAM83C_E" [color="0 0 0.7320224719101124", label="0.52",style=bold];
"CWH43_S" -> "SERPINB13_E" [color="0 0 0.7314606741573033", label="0.53",style=bold];
"DSG1_S" -> "CPA2_E" [color="0 0 0.7308988764044944", label="0.54",style=bold];
"PNLIPRP2_S" -> "PCK1_E" [color="0 0 0.7303370786516854", label="0.54",style=bold];
"COL2A1_S" -> "SLC13A2_M" [color="0 0 0.7297752808988764", label="0.54",style=bold];
"IRX6_S" -> "PGLYRP3_M" [color="0 0 0.7292134831460675", label="0.54",style=bold];
"PCK1_S" -> "MUC17_S" [color="0 0 0.7286516853932584", label="0.54",style=bold];
"KRT16_S" -> "MUC16_E" [color="0 0 0.7280898876404495", label="0.55",style=bold];
"GPR87_S" -> "S100A7_M" [color="0 0 0.7275280898876404", label="0.55",style=bold];
"KCNA1_S" -> "SLCO1B3_M" [color="0 0 0.7269662921348314", label="0.55",style=bold];
"SERPINB3_S" -> "SLCO1B3_E" [color="0 0 0.7264044943820225", label="0.55",style=bold];
"CNTN1_S" -> "FRMPD2_M" [color="0 0 0.7258426966292135", label="0.55",style=bold];
"SERPINB13_S" -> "RFX6_E" [color="0 0 0.7252808988764046", label="0.56",style=bold];
"C20orf114_S" -> "KRT6C_M" [color="0 0 0.7247191011235955", label="0.56",style=bold];
"A2ML1_S" -> "RIMS2_M" [color="0 0 0.7241573033707865", label="0.56",style=bold];
"EGF_S" -> "TGM5_S" [color="0 0 0.7235955056179775", label="0.56",style=bold];
"C20orf114_S" -> "MYH4_E" [color="0 0 0.7230337078651685", label="0.57",style=bold];
"PGC_S" -> "PGC_E" [color="0 0 0.7224719101123596", label="0.57",style=bold];
"SLCO1B3_S" -> "RIMS2_S" [color="0 0 0.7219101123595506", label="0.57",style=bold];
"CPA2_S" -> "ABCA12_E" [color="0 0 0.7213483146067416", label="0.57",style=bold];
"SERPINB4_S" -> "KRT6C_E" [color="0 0 0.7207865168539326", label="0.57",style=bold];
"KRT6B_S" -> "DSG1_M" [color="0 0 0.7202247191011236", label="0.58",style=bold];
"CPA2_S" -> "ABCA12_M" [color="0 0 0.7196629213483146", label="0.58",style=bold];
"CWH43_S" -> "FRMPD2_M" [color="0 0 0.7191011235955056", label="0.58",style=bold];
"COL2A1_S" -> "SERPINB4_M" [color="0 0 0.7185393258426966", label="0.58",style=bold];
"C20orf114_S" -> "DPP6_S" [color="0 0 0.7179775280898877", label="0.58",style=bold];
"CTSE_S" -> "DMBT1_E" [color="0 0 0.7174157303370786", label="0.58",style=bold];
"ACTN2_S" -> "ABCA13_E" [color="0 0 0.7168539325842697", label="0.58",style=bold];
"GLRA1_S" -> "PGLYRP3_M" [color="0 0 0.7162921348314607", label="0.59",style=bold];
"SLC26A3_S" -> "MUC6_M" [color="0 0 0.7157303370786516", label="0.59",style=bold];
"A2ML1_S" -> "TGM5_M" [color="0 0 0.7151685393258427", label="0.59",style=bold];
"C20orf114_S" -> "FER1L6_M" [color="0 0 0.7146067415730337", label="0.59",style=bold];
"SERPINB3_S" -> "PNLIPRP2_M" [color="0 0 0.7140449438202248", label="0.59",style=bold];
"KRT6B_S" -> "MUC2_E" [color="0 0 0.7134831460674157", label="0.60",style=bold];
"KLK5_S" -> "AGBL1_M" [color="0 0 0.7129213483146067", label="0.60",style=bold];
"MYH4_S" -> "CSMD3_S" [color="0 0 0.7123595505617978", label="0.60",style=bold];
"PCDHA12_S" -> "FREM2_M" [color="0 0 0.7117977528089887", label="0.60",style=bold];
"KRT13_S" -> "IRX6_M" [color="0 0 0.7112359550561798", label="0.60",style=bold];
"KRT13_S" -> "SERPINB4_M" [color="0 0 0.7106741573033708", label="0.61",style=bold];
"KRT6C_S" -> "MUC5B_E" [color="0 0 0.7101123595505618", label="0.61",style=bold];
"SLC13A2_S" -> "GPR98_M" [color="0 0 0.7095505617977528", label="0.61",style=bold];
"CPA2_S" -> "TGM5_E" [color="0 0 0.7089887640449438", label="0.61",style=bold];
"KRT6B_S" -> "SLC13A2_E" [color="0 0 0.7084269662921349", label="0.61",style=bold];
"COL2A1_S" -> "CACNA1B_E" [color="0 0 0.7078651685393258", label="0.62",style=bold];
"PCDHA12_S" -> "MYH4_S" [color="0 0 0.7073033707865168", label="0.64",style=bold];
"CTSE_S" -> "FLG_S" [color="0 0 0.7067415730337079", label="0.64",style=bold];
"KRT6B_S" -> "Survival" [color="0 0 0.7061797752808989", label="0.64",style=bold];
"EGF_S" -> "SERPINB13_M" [color="0 0 0.7056179775280899", label="0.65",style=bold];
"KLK5_S" -> "EGF_M" [color="0 0 0.7050561797752809", label="0.66",style=bold];
"SLC13A2_S" -> "MUC2_E" [color="0 0 0.7044943820224719", label="0.66",style=bold];
"FER1L6_S" -> "MDGA2_S" [color="0 0 0.7039325842696629", label="0.66",style=bold];
"DSG1_S" -> "FER1L6_E" [color="0 0 0.7033707865168539", label="0.66",style=bold];
"CPA2_S" -> "CWH43_E" [color="0 0 0.702808988764045", label="0.66",style=bold];
"GLRA1_S" -> "CNTNAP2_E" [color="0 0 0.702247191011236", label="0.66",style=bold];
"SERPINB3_S" -> "CPA2_M" [color="0 0 0.7016853932584269", label="0.66",style=bold];
"PNLIPRP2_S" -> "KRT16_M" [color="0 0 0.701123595505618", label="0.68",style=bold];
"KLK5_S" -> "FSTL5_S" [color="0 0 0.700561797752809", label="0.68",style=bold];
"PCDH10_S" -> "SERPINB3_M" [color="0 0 0.7", label="0.68",style=bold];
"KLK5_S" -> "CNTN1_S" [color="0 0 0.699438202247191", label="0.69",style=bold];
"SLCO1B3_S" -> "MUC6_S" [color="0 0 0.698876404494382", label="0.69",style=bold];
"PGC_S" -> "DPP6_E" [color="0 0 0.6983146067415731", label="0.69",style=bold];
"CPA2_S" -> "EGF_M" [color="0 0 0.697752808988764", label="0.69",style=bold];
"PNLIPRP2_S" -> "DCDC1_M" [color="0 0 0.6971910112359551", label="0.69",style=bold];
"CPA2_S" -> "DCDC1_E" [color="0 0 0.6966292134831461", label="0.69",style=bold];
"COL2A1_S" -> "GLRA1_E" [color="0 0 0.696067415730337", label="0.70",style=bold];
"PGC_S" -> "C20orf114_M" [color="0 0 0.6955056179775281", label="0.70",style=bold];
"FER1L6_S" -> "ABCA12_S" [color="0 0 0.6949438202247191", label="0.71",style=bold];
"PNLIPRP2_S" -> "AGBL1_M" [color="0 0 0.6943820224719102", label="0.71",style=bold];
"PGC_S" -> "SLC6A19_M" [color="0 0 0.6938202247191011", label="0.71",style=bold];
"MDGA2_S" -> "ABCA12_S" [color="0 0 0.6932584269662921", label="0.71",style=bold];
"SLC6A19_S" -> "LPA_E" [color="0 0 0.6926966292134832", label="0.71",style=bold];
"MUC15_S" -> "SLCO1B1_M" [color="0 0 0.6921348314606741", label="0.72",style=bold];
"OGDHL_S" -> "MUC4_M" [color="0 0 0.6915730337078652", label="0.72",style=bold];
"LRRC7_S" -> "RIMS2_E" [color="0 0 0.6910112359550562", label="0.72",style=bold];
"KRT16_S" -> "ADAMTS20_S" [color="0 0 0.6904494382022472", label="0.73",style=bold];
"CTSE_S" -> "SLC9A11_E" [color="0 0 0.6898876404494382", label="0.73",style=bold];
"CTSE_S" -> "MUC6_S" [color="0 0 0.6893258426966292", label="0.74",style=bold];
"DCDC1_S" -> "ADAMTS20_E" [color="0 0 0.6887640449438203", label="0.74",style=bold];
"ACTN2_S" -> "MUC4_E" [color="0 0 0.6882022471910112", label="0.74",style=bold];
"COL2A1_S" -> "TGM5_M" [color="0 0 0.6876404494382022", label="0.74",style=bold];
"NETO1_S" -> "PTPRZ1_E" [color="0 0 0.6870786516853933", label="0.74",style=bold];
"AGBL1_S" -> "PNLIPRP2_E" [color="0 0 0.6865168539325843", label="0.74",style=bold];
"CPA2_S" -> "FAM83C_E" [color="0 0 0.6859550561797753", label="0.75",style=bold];
"PGC_S" -> "DCDC1_E" [color="0 0 0.6853932584269663", label="0.75",style=bold];
"SLC9A11_S" -> "SERPINB3_M" [color="0 0 0.6848314606741573", label="0.75",style=bold];
"SERPINB13_S" -> "MDGA2_E" [color="0 0 0.6842696629213483", label="0.75",style=bold];
"GLRA1_S" -> "CWH43_M" [color="0 0 0.6837078651685393", label="0.75",style=bold];
"NETO1_S" -> "CACNA1B_E" [color="0 0 0.6831460674157304", label="0.75",style=bold];
"S100A7_S" -> "KRT6C_S" [color="0 0 0.6825842696629214", label="0.75",style=bold];
"KRT6B_S" -> "PGC_E" [color="0 0 0.6820224719101123", label="0.76",style=bold];
"IRX6_S" -> "PNLIPRP2_M" [color="0 0 0.6814606741573034", label="0.76",style=bold];
"PNLIPRP2_S" -> "NETO1_E" [color="0 0 0.6808988764044944", label="0.77",style=bold];
"SERPINB13_S" -> "SLC13A2_E" [color="0 0 0.6803370786516854", label="0.77",style=bold];
"FLG_S" -> "ACTN2_S" [color="0 0 0.6797752808988764", label="0.77",style=bold];
"CPA2_S" -> "FBN3_M" [color="0 0 0.6792134831460674", label="0.77",style=bold];
"SERPINB4_S" -> "SLC26A3_E" [color="0 0 0.6786516853932585", label="0.77",style=bold];
"PGC_S" -> "FLG_M" [color="0 0 0.6780898876404494", label="0.77",style=bold];
"PGLYRP3_S" -> "FBN3_E" [color="0 0 0.6775280898876405", label="0.77",style=bold];
"SERPINB13_S" -> "KRT16_M" [color="0 0 0.6769662921348315", label="0.78",style=bold];
"KRT16_S" -> "SLC6A19_E" [color="0 0 0.6764044943820224", label="0.78",style=bold];
"FRMPD2_S" -> "SLCO1B3_M" [color="0 0 0.6758426966292135", label="0.78",style=bold];
"PGC_S" -> "CNTNAP2_E" [color="0 0 0.6752808988764045", label="0.78",style=bold];
"KRT6C_S" -> "SERPINB3_M" [color="0 0 0.6747191011235956", label="0.78",style=bold];
"SLC26A3_S" -> "FRMPD2_M" [color="0 0 0.6741573033707865", label="0.79",style=bold];
"TGM5_S" -> "DPP6_E" [color="0 0 0.6735955056179775", label="0.79",style=bold];
"DSG1_S" -> "DCDC1_E" [color="0 0 0.6730337078651685", label="0.80",style=bold];
"PGC_S" -> "CR2_E" [color="0 0 0.6724719101123595", label="0.80",style=bold];
"SERPINB3_S" -> "KRT16_M" [color="0 0 0.6719101123595506", label="0.80",style=bold];
"KRT16_S" -> "CPA2_M" [color="0 0 0.6713483146067416", label="0.80",style=bold];
"PGC_S" -> "DCDC1_M" [color="0 0 0.6707865168539326", label="0.80",style=bold];
"C20orf114_S" -> "NETO1_E" [color="0 0 0.6702247191011236", label="0.80",style=bold];
"SERPINB4_S" -> "SLC26A3_M" [color="0 0 0.6696629213483146", label="0.80",style=bold];
"GLRA1_S" -> "RP1_E" [color="0 0 0.6691011235955056", label="0.80",style=bold];
"KRT6B_S" -> "SLCO1B1_M" [color="0 0 0.6685393258426966", label="0.80",style=bold];
"PGLYRP3_S" -> "DMBT1_M" [color="0 0 0.6679775280898876", label="0.80",style=bold];
"RFX6_S" -> "NETO1_E" [color="0 0 0.6674157303370787", label="0.81",style=bold];
"KRT13_S" -> "FSTL5_E" [color="0 0 0.6668539325842697", label="0.81",style=bold];
"CTSE_S" -> "MUC16_S" [color="0 0 0.6662921348314607", label="0.81",style=bold];
"SERPINB4_S" -> "MUC16_E" [color="0 0 0.6657303370786517", label="0.82",style=bold];
"FAM83C_S" -> "MUC6_S" [color="0 0 0.6651685393258426", label="0.82",style=bold];
"KLK5_S" -> "SLC26A3_S" [color="0 0 0.6646067415730337", label="0.82",style=bold];
"SLC13A2_S" -> "KRT13_S" [color="0 0 0.6640449438202247", label="0.84",style=bold];
"KRT6C_S" -> "KRT16_S" [color="0 0 0.6634831460674158", label="0.84",style=bold];
"CPA2_S" -> "ACTN2_E" [color="0 0 0.6629213483146068", label="0.85",style=bold];
"CPA2_S" -> "FRMPD2_M" [color="0 0 0.6623595505617977", label="0.85",style=bold];
"S100A7_S" -> "IRX6_E" [color="0 0 0.6617977528089888", label="0.85",style=bold];
"GLRA1_S" -> "MUC4_M" [color="0 0 0.6612359550561797", label="0.85",style=bold];
"GPR87_S" -> "SLC9A4_M" [color="0 0 0.6606741573033708", label="0.85",style=bold];
"SERPINB3_S" -> "KLK5_M" [color="0 0 0.6601123595505618", label="0.85",style=bold];
"SERPINB3_S" -> "MUC17_M" [color="0 0 0.6595505617977528", label="0.85",style=bold];
"KLK5_S" -> "PCK1_E" [color="0 0 0.6589887640449439", label="0.85",style=bold];
"FAM83C_E" -> "KLK5_E" [color="0 0 0.6584269662921348", label="0.85",style=bold];
"SLC13A2_S" -> "S100A7_M" [color="0 0 0.6578651685393259", label="0.85",style=bold];
"SERPINB3_S" -> "APOB_M" [color="0 0 0.6573033707865168", label="0.86",style=bold];
"KRT16_S" -> "DCDC1_M" [color="0 0 0.6567415730337078", label="0.86",style=bold];
"CPA2_S" -> "TGM5_M" [color="0 0 0.6561797752808989", label="0.86",style=bold];
"GPR87_S" -> "THBS4_E" [color="0 0 0.6556179775280899", label="0.86",style=bold];
"SERPINB3_S" -> "PNLIPRP2_E" [color="0 0 0.655056179775281", label="0.86",style=bold];
"ZIC4_S" -> "FER1L6_E" [color="0 0 0.6544943820224719", label="0.87",style=bold];
"IRX6_S" -> "DMBT1_E" [color="0 0 0.6539325842696629", label="0.87",style=bold];
"DCDC1_S" -> "Survival" [color="0 0 0.6533707865168539", label="0.87",style=bold];
"SLC13A2_S" -> "FRMPD2_M" [color="0 0 0.6528089887640449", label="0.88",style=bold];
"PCDH10_S" -> "RFX6_M" [color="0 0 0.652247191011236", label="0.88",style=bold];
"SLC26A3_S" -> "SLCO1B1_E" [color="0 0 0.651685393258427", label="0.89",style=bold];
"SERPINB3_S" -> "S100A7_M" [color="0 0 0.651123595505618", label="0.89",style=bold];
"PGC_S" -> "FER1L6_E" [color="0 0 0.650561797752809", label="0.89",style=bold];
"A2ML1_S" -> "MUC6_E" [color="0 0 0.65", label="0.89",style=bold];
"IRX6_S" -> "CPA2_M" [color="0 0 0.649438202247191", label="0.90",style=bold];
"ZIC4_S" -> "PCLO_S" [color="0 0 0.648876404494382", label="0.90",style=bold];
"KCNA1_S" -> "FRMPD2_S" [color="0 0 0.648314606741573", label="0.90",style=bold];
"SLC9A4_S" -> "COL11A1_S" [color="0 0 0.6477528089887641", label="0.90",style=bold];
"PGC_S" -> "FAM83C_M" [color="0 0 0.6471910112359551", label="0.91",style=bold];
"KRT16_S" -> "IRX6_E" [color="0 0 0.6466292134831461", label="0.91",style=bold];
"C20orf114_S" -> "RFX6_E" [color="0 0 0.6460674157303371", label="0.91",style=bold];
"PNLIPRP2_S" -> "SLC6A19_M" [color="0 0 0.645505617977528", label="0.92",style=bold];
"SERPINB4_S" -> "FREM2_E" [color="0 0 0.6449438202247191", label="0.92",style=bold];
"KRT6B_S" -> "RIMS2_E" [color="0 0 0.6443820224719101", label="0.92",style=bold];
"DCDC1_S" -> "PCK1_E" [color="0 0 0.6438202247191012", label="0.92",style=bold];
"A2ML1_S" -> "SERPINB4_S" [color="0 0 0.6432584269662922", label="0.92",style=bold];
"IRX6_S" -> "FER1L6_M" [color="0 0 0.6426966292134831", label="0.93",style=bold];
"KLK5_S" -> "PTPRZ1_E" [color="0 0 0.6421348314606742", label="0.93",style=bold];
"PGC_S" -> "GLRA1_E" [color="0 0 0.6415730337078651", label="0.94",style=bold];
"SLC13A2_S" -> "RIMS2_E" [color="0 0 0.6410112359550562", label="0.94",style=bold];
"EGF_S" -> "NETO1_E" [color="0 0 0.6404494382022472", label="0.95",style=bold];
"SLC9A4_S" -> "ACTN2_E" [color="0 0 0.6398876404494382", label="0.95",style=bold];
"KRT13_S" -> "SERPINB3_M" [color="0 0 0.6393258426966293", label="0.95",style=bold];
"PNLIPRP2_S" -> "PNLIPRP2_M" [color="0 0 0.6387640449438202", label="0.95",style=bold];
"KRT13_S" -> "COL11A1_E" [color="0 0 0.6382022471910113", label="0.95",style=bold];
"C20orf114_S" -> "PCLO_M" [color="0 0 0.6376404494382022", label="0.95",style=bold];
"PGC_S" -> "KCNA1_E" [color="0 0 0.6370786516853932", label="0.95",style=bold];
"GLRA1_S" -> "PCDHA11_M" [color="0 0 0.6365168539325843", label="0.95",style=bold];
"PNLIPRP2_S" -> "CPA2_E" [color="0 0 0.6359550561797753", label="0.96",style=bold];
"SLC26A3_S" -> "COL11A1_E" [color="0 0 0.6353932584269664", label="0.96",style=bold];
"SLC26A3_S" -> "GPR98_S" [color="0 0 0.6348314606741573", label="0.96",style=bold];
"SERPINB13_S" -> "CPA2_E" [color="0 0 0.6342696629213482", label="0.97",style=bold];
"SLCO1B1_S" -> "CPA2_E" [color="0 0 0.6337078651685393", label="0.97",style=bold];
"A2ML1_S" -> "FSTL5_E" [color="0 0 0.6331460674157303", label="0.97",style=bold];
"DMBT1_S" -> "DCDC1_S" [color="0 0 0.6325842696629214", label="0.97",style=bold];
"KRT6C_S" -> "DMBT1_E" [color="0 0 0.6320224719101124", label="0.97",style=bold];
"KRT13_S" -> "GLRA1_E" [color="0 0 0.6314606741573034", label="0.98",style=bold];
"ACTN2_S" -> "DPP6_E" [color="0 0 0.6308988764044944", label="0.98",style=bold];
"CPA2_S" -> "MUC6_M" [color="0 0 0.6303370786516853", label="0.98",style=bold];
"CPA2_S" -> "CACNA1B_E" [color="0 0 0.6297752808988764", label="0.98",style=bold];
"PGC_S" -> "SLC9A4_M" [color="0 0 0.6292134831460674", label="0.98",style=bold];
"DSG1_S" -> "RP1_E" [color="0 0 0.6286516853932584", label="0.98",style=bold];
"CNTN1_S" -> "SLC6A19_M" [color="0 0 0.6280898876404495", label="0.98",style=bold];
"SLC9A4_S" -> "EGF_S" [color="0 0 0.6275280898876405", label="0.98",style=bold];
"GLRA1_S" -> "FRMPD2_E" [color="0 0 0.6269662921348315", label="0.99",style=bold];
"EGF_S" -> "KRT6B_M" [color="0 0 0.6264044943820224", label="0.99",style=bold];
"PGC_S" -> "SERPINB4_E" [color="0 0 0.6258426966292134", label="0.99",style=bold];
"PNLIPRP2_S" -> "SERPINB3_M" [color="0 0 0.6252808988764045", label="1.00",style=bold];
"LRRC7_S" -> "DMBT1_M" [color="0 0 0.6247191011235955", label="1.00",style=bold];
"PCK1_S" -> "CPA2_M" [color="0 0 0.6241573033707866", label="1.00",style=bold];
"KLK5_S" -> "ABCA12_M" [color="0 0 0.6235955056179776", label="1.00",style=bold];
"PTPRZ1_S" -> "TGM5_M" [color="0 0 0.6230337078651685", label="1.00",style=bold];
"FBN3_S" -> "FREM2_E" [color="0 0 0.6224719101123595", label="1.00",style=bold];
"PCDH10_S" -> "RP1_S" [color="0 0 0.6219101123595505", label="1.01",style=bold];
"TGM5_S" -> "NETO1_E" [color="0 0 0.6213483146067416", label="1.01",style=bold];
"GPR87_S" -> "KRT6C_M" [color="0 0 0.6207865168539326", label="1.01",style=bold];
"SLC26A3_S" -> "CACNA1B_M" [color="0 0 0.6202247191011236", label="1.01",style=bold];
"GLRA1_S" -> "RFX6_E" [color="0 0 0.6196629213483146", label="1.01",style=bold];
"CPA2_S" -> "FSTL5_E" [color="0 0 0.6191011235955056", label="1.01",style=bold];
"FAM83C_S" -> "KCNA1_S" [color="0 0 0.6185393258426966", label="1.01",style=bold];
"SERPINB3_S" -> "TGM5_M" [color="0 0 0.6179775280898876", label="1.02",style=bold];
"C20orf114_S" -> "COL11A1_E" [color="0 0 0.6174157303370786", label="1.02",style=bold];
"KRT6C_S" -> "AGBL1_S" [color="0 0 0.6168539325842697", label="1.02",style=bold];
"PNLIPRP2_S" -> "SLC26A3_M" [color="0 0 0.6162921348314607", label="1.02",style=bold];
"PGLYRP3_S" -> "SLC9A4_E" [color="0 0 0.6157303370786517", label="1.03",style=bold];
"SLCO1B1_S" -> "MUC5B_E" [color="0 0 0.6151685393258427", label="1.03",style=bold];
"PNLIPRP2_S" -> "MUC4_S" [color="0 0 0.6146067415730336", label="1.03",style=bold];
"DPP6_S" -> "SLCO1B1_S" [color="0 0 0.6140449438202247", label="1.03",style=bold];
"PNLIPRP2_S" -> "SLC9A11_M" [color="0 0 0.6134831460674157", label="1.03",style=bold];
"NETO1_S" -> "APOB_E" [color="0 0 0.6129213483146068", label="1.04",style=bold];
"FAM83C_S" -> "APOB_E" [color="0 0 0.6123595505617978", label="1.04",style=bold];
"KRT13_S" -> "PCDH10_E" [color="0 0 0.6117977528089887", label="1.05",style=bold];
"PNLIPRP2_S" -> "CSMD3_S" [color="0 0 0.6112359550561798", label="1.05",style=bold];
"GLRA1_S" -> "SLC9A4_E" [color="0 0 0.6106741573033707", label="1.05",style=bold];
"GLRA1_S" -> "PNLIPRP2_E" [color="0 0 0.6101123595505618", label="1.05",style=bold];
"CPA2_S" -> "CSMD3_E" [color="0 0 0.6095505617977528", label="1.06",style=bold];
"MUC2_S" -> "A2ML1_E" [color="0 0 0.6089887640449438", label="1.06",style=bold];
"CPA2_S" -> "MUC2_E" [color="0 0 0.6084269662921349", label="1.06",style=bold];
"FRMPD2_S" -> "SLCO1B1_M" [color="0 0 0.6078651685393258", label="1.06",style=bold];
"KLK5_S" -> "MUC17_S" [color="0 0 0.6073033707865169", label="1.07",style=bold];
"KRT16_S" -> "DMBT1_E" [color="0 0 0.6067415730337078", label="1.07",style=bold];
"DSG1_S" -> "CNTN1_E" [color="0 0 0.6061797752808988", label="1.07",style=bold];
"PCDHA11_S" -> "FER1L6_S" [color="0 0 0.6056179775280899", label="1.07",style=bold];
"SLC26A3_S" -> "SLC9A11_E" [color="0 0 0.6050561797752809", label="1.07",style=bold];
"SLCO1B1_S" -> "ADAMTS20_S" [color="0 0 0.604494382022472", label="1.08",style=bold];
"THBS4_S" -> "SLC6A19_E" [color="0 0 0.6039325842696629", label="1.08",style=bold];
"PCDHA11_S" -> "KCNA1_S" [color="0 0 0.6033707865168539", label="1.08",style=bold];
"SLC13A2_S" -> "KRT13_E" [color="0 0 0.6028089887640449", label="1.08",style=bold];
"SLC6A19_S" -> "S100A7_M" [color="0 0 0.6022471910112359", label="1.09",style=bold];
"KRT6C_S" -> "PCK1_E" [color="0 0 0.601685393258427", label="1.09",style=bold];
"SLC13A2_S" -> "PKHD1L1_M" [color="0 0 0.601123595505618", label="1.09",style=bold];
"MUC6_S" -> "SLITRK1_E" [color="0 0 0.600561797752809", label="1.09",style=bold];
"FER1L6_S" -> "TGM5_M" [color="0 0 0.6", label="1.10",style=bold];
"KLK5_S" -> "ABCA13_M" [color="0 0 0.599438202247191", label="1.10",style=bold];
"SLC6A19_S" -> "MUC4_M" [color="0 0 0.598876404494382", label="1.10",style=bold];
"FAM83C_S" -> "Survival" [color="0 0 0.598314606741573", label="1.10",style=bold];
"SERPINB3_S" -> "PTPRZ1_M" [color="0 0 0.597752808988764", label="1.10",style=bold];
"PGC_S" -> "DSG1_M" [color="0 0 0.5971910112359551", label="1.10",style=bold];
"KRT6C_S" -> "GPR98_S" [color="0 0 0.5966292134831461", label="1.10",style=bold];
"CPA2_S" -> "RP1_M" [color="0 0 0.5960674157303371", label="1.10",style=bold];
"PGC_S" -> "PNLIPRP2_M" [color="0 0 0.5955056179775281", label="1.10",style=bold];
"PGC_S" -> "FER1L6_M" [color="0 0 0.594943820224719", label="1.10",style=bold];
"CR2_S" -> "PKHD1L1_E" [color="0 0 0.5943820224719101", label="1.10",style=bold];
"TGM5_S" -> "MYH4_M" [color="0 0 0.5938202247191011", label="1.10",style=bold];
"CTSE_S" -> "MUC4_E" [color="0 0 0.5932584269662922", label="1.11",style=bold];
"SLC13A2_S" -> "SLCO1B1_M" [color="0 0 0.5926966292134832", label="1.11",style=bold];
"SLC6A19_S" -> "FSTL5_M" [color="0 0 0.5921348314606741", label="1.12",style=bold];
"FAM83C_S" -> "PKHD1L1_M" [color="0 0 0.5915730337078652", label="1.12",style=bold];
"SERPINB3_S" -> "AGBL1_S" [color="0 0 0.5910112359550561", label="1.12",style=bold];
"DSG1_S" -> "SLC9A4_E" [color="0 0 0.5904494382022472", label="1.13",style=bold];
"GLRA1_S" -> "SLC13A2_E" [color="0 0 0.5898876404494382", label="1.13",style=bold];
"MUC17_S" -> "Survival" [color="0 0 0.5893258426966292", label="1.13",style=bold];
"EGF_S" -> "PGC_M" [color="0 0 0.5887640449438203", label="1.13",style=bold];
"GPR87_S" -> "PTPRZ1_S" [color="0 0 0.5882022471910112", label="1.13",style=bold];
"SLC13A2_S" -> "APOB_M" [color="0 0 0.5876404494382023", label="1.13",style=bold];
"GPR87_S" -> "KLK5_M" [color="0 0 0.5870786516853932", label="1.13",style=bold];
"KRT13_S" -> "ADAMTS20_S" [color="0 0 0.5865168539325842", label="1.13",style=bold];
"PGC_S" -> "MUC15_E" [color="0 0 0.5859550561797753", label="1.13",style=bold];
"TGM5_S" -> "CACNA1B_E" [color="0 0 0.5853932584269663", label="1.13",style=bold];
"SLC26A3_S" -> "PNLIPRP2_M" [color="0 0 0.5848314606741574", label="1.13",style=bold];
"PGC_S" -> "RP1_E" [color="0 0 0.5842696629213483", label="1.13",style=bold];
"GLRA1_S" -> "AGBL1_E" [color="0 0 0.5837078651685393", label="1.14",style=bold];
"SLCO1B3_S" -> "MUC5B_E" [color="0 0 0.5831460674157303", label="1.14",style=bold];
"LPA_S" -> "SLCO1B3_S" [color="0 0 0.5825842696629213", label="1.15",style=bold];
"PGC_S" -> "KLK5_M" [color="0 0 0.5820224719101124", label="1.15",style=bold];
"PGC_S" -> "PCDHA12_M" [color="0 0 0.5814606741573034", label="1.15",style=bold];
"DSG1_S" -> "MUC5B_E" [color="0 0 0.5808988764044944", label="1.15",style=bold];
"SERPINB13_S" -> "SLCO1B1_M" [color="0 0 0.5803370786516854", label="1.16",style=bold];
"LPA_S" -> "AGBL1_S" [color="0 0 0.5797752808988764", label="1.16",style=bold];
"DCDC1_S" -> "MUC5B_S" [color="0 0 0.5792134831460674", label="1.16",style=bold];
"KRT13_S" -> "GPR98_S" [color="0 0 0.5786516853932584", label="1.16",style=bold];
"KLK5_S" -> "MUC2_E" [color="0 0 0.5780898876404494", label="1.16",style=bold];
"TGM5_S" -> "PNLIPRP2_M" [color="0 0 0.5775280898876405", label="1.17",style=bold];
"FER1L6_S" -> "SLCO1B1_S" [color="0 0 0.5769662921348315", label="1.17",style=bold];
"CPA2_S" -> "FLG_M" [color="0 0 0.5764044943820225", label="1.17",style=bold];
"DSG1_S" -> "KRT13_S" [color="0 0 0.5758426966292135", label="1.17",style=bold];
"AGBL1_S" -> "MUC17_M" [color="0 0 0.5752808988764044", label="1.18",style=bold];
"PGC_S" -> "SLC13A2_M" [color="0 0 0.5747191011235955", label="1.18",style=bold];
"KRT16_S" -> "AGBL1_M" [color="0 0 0.5741573033707865", label="1.18",style=bold];
"MUC15_S" -> "LRRC7_E" [color="0 0 0.5735955056179776", label="1.18",style=bold];
"CNTN1_S" -> "FREM2_M" [color="0 0 0.5730337078651686", label="1.18",style=bold];
"SLCO1B3_S" -> "PCLO_E" [color="0 0 0.5724719101123595", label="1.19",style=bold];
"FBN3_S" -> "SLC6A19_E" [color="0 0 0.5719101123595506", label="1.19",style=bold];
"KRT6C_S" -> "MUC15_S" [color="0 0 0.5713483146067415", label="1.19",style=bold];
"PNLIPRP2_S" -> "DMBT1_M" [color="0 0 0.5707865168539326", label="1.19",style=bold];
"EGF_S" -> "PNLIPRP2_M" [color="0 0 0.5702247191011236", label="1.19",style=bold];
"KRT6B_S" -> "PNLIPRP2_E" [color="0 0 0.5696629213483146", label="1.20",style=bold];
"KRT16_S" -> "FER1L6_S" [color="0 0 0.5691011235955057", label="1.20",style=bold];
"PNLIPRP2_S" -> "SLC9A4_M" [color="0 0 0.5685393258426966", label="1.20",style=bold];
"CWH43_S" -> "DCDC1_M" [color="0 0 0.5679775280898876", label="1.20",style=bold];
"CPA2_S" -> "PGC_M" [color="0 0 0.5674157303370786", label="1.20",style=bold];
"SERPINB4_S" -> "RP1_E" [color="0 0 0.5668539325842696", label="1.20",style=bold];
"PGC_S" -> "MUC4_E" [color="0 0 0.5662921348314607", label="1.20",style=bold];
"PGLYRP3_S" -> "PCDH10_S" [color="0 0 0.5657303370786517", label="1.20",style=bold];
"SLCO1B3_S" -> "FSTL5_S" [color="0 0 0.5651685393258428", label="1.21",style=bold];
"FRMPD2_S" -> "FSTL5_E" [color="0 0 0.5646067415730337", label="1.21",style=bold];
"PGLYRP3_S" -> "PGLYRP3_M" [color="0 0 0.5640449438202246", label="1.21",style=bold];
"GPR87_S" -> "RIMS2_M" [color="0 0 0.5634831460674157", label="1.22",style=bold];
"FRMPD2_S" -> "APOB_S" [color="0 0 0.5629213483146067", label="1.22",style=bold];
"MYH4_S" -> "CNTNAP2_E" [color="0 0 0.5623595505617978", label="1.22",style=bold];
"FBN3_S" -> "PKHD1L1_S" [color="0 0 0.5617977528089888", label="1.22",style=bold];
"PGLYRP3_S" -> "RIMS2_S" [color="0 0 0.5612359550561798", label="1.23",style=bold];
"SERPINB4_S" -> "SLC6A19_E" [color="0 0 0.5606741573033708", label="1.23",style=bold];
"CNTNAP2_S" -> "MUC4_M" [color="0 0 0.5601123595505617", label="1.24",style=bold];
"TGM5_S" -> "MUC5B_E" [color="0 0 0.5595505617977528", label="1.24",style=bold];
"EGF_S" -> "KRT6C_M" [color="0 0 0.5589887640449438", label="1.24",style=bold];
"KLK5_S" -> "GLRA1_E" [color="0 0 0.5584269662921348", label="1.24",style=bold];
"KLK5_S" -> "DMBT1_S" [color="0 0 0.5578651685393259", label="1.25",style=bold];
"A2ML1_S" -> "COL2A1_S" [color="0 0 0.5573033707865169", label="1.25",style=bold];
"COL2A1_S" -> "SLC26A3_M" [color="0 0 0.5567415730337079", label="1.25",style=bold];
"KLK5_S" -> "MYH4_M" [color="0 0 0.5561797752808988", label="1.25",style=bold];
"DSG1_S" -> "FRMPD2_S" [color="0 0 0.5556179775280898", label="1.25",style=bold];
"SERPINB3_S" -> "FLG_E" [color="0 0 0.5550561797752809", label="1.26",style=bold];
"SERPINB3_S" -> "KLK5_E" [color="0 0 0.5544943820224719", label="1.27",style=bold];
"SLC13A2_S" -> "PTPRZ1_S" [color="0 0 0.553932584269663", label="1.27",style=bold];
"KCNA1_S" -> "SERPINB4_E" [color="0 0 0.553370786516854", label="1.27",style=bold];
"CNTN1_S" -> "CNTNAP2_E" [color="0 0 0.5528089887640449", label="1.28",style=bold];
"GLRA1_S" -> "CWH43_E" [color="0 0 0.5522471910112359", label="1.28",style=bold];
"KLK5_S" -> "CSMD3_E" [color="0 0 0.5516853932584269", label="1.28",style=bold];
"C20orf114_S" -> "SLCO1B1_S" [color="0 0 0.551123595505618", label="1.28",style=bold];
"FBN3_S" -> "FLG_S" [color="0 0 0.550561797752809", label="1.29",style=bold];
"PNLIPRP2_S" -> "MUC2_E" [color="0 0 0.55", label="1.29",style=bold];
"DSG1_S" -> "LPA_E" [color="0 0 0.549438202247191", label="1.29",style=bold];
"SLC13A2_S" -> "MUC6_E" [color="0 0 0.548876404494382", label="1.30",style=bold];
"MUC15_S" -> "MUC6_E" [color="0 0 0.548314606741573", label="1.30",style=bold];
"KRT6B_S" -> "SLC6A19_E" [color="0 0 0.547752808988764", label="1.30",style=bold];
"C20orf114_S" -> "TGM5_M" [color="0 0 0.547191011235955", label="1.30",style=bold];
"FRMPD2_S" -> "A2ML1_M" [color="0 0 0.5466292134831461", label="1.30",style=bold];
"KRT6B_S" -> "FLG_E" [color="0 0 0.5460674157303371", label="1.31",style=bold];
"KLK5_S" -> "SLC9A4_E" [color="0 0 0.5455056179775281", label="1.31",style=bold];
"CPA2_S" -> "EGF_E" [color="0 0 0.5449438202247191", label="1.31",style=bold];
"C20orf114_S" -> "Survival" [color="0 0 0.54438202247191", label="1.32",style=bold];
"KLK5_S" -> "SERPINB3_M" [color="0 0 0.5438202247191011", label="1.33",style=bold];
"THBS4_S" -> "C20orf114_E" [color="0 0 0.5432584269662921", label="1.33",style=bold];
"PGC_S" -> "IRX6_E" [color="0 0 0.5426966292134832", label="1.33",style=bold];
"SLC9A11_S" -> "FAM83C_M" [color="0 0 0.5421348314606742", label="1.33",style=bold];
"TGM5_S" -> "A2ML1_M" [color="0 0 0.5415730337078651", label="1.33",style=bold];
"TGM5_S" -> "MUC6_S" [color="0 0 0.5410112359550562", label="1.33",style=bold];
"FAM83C_S" -> "APOB_M" [color="0 0 0.5404494382022471", label="1.33",style=bold];
"GPR87_S" -> "PGLYRP3_M" [color="0 0 0.5398876404494382", label="1.34",style=bold];
"FBN3_S" -> "FREM2_S" [color="0 0 0.5393258426966292", label="1.35",style=bold];
"DSG1_S" -> "FER1L6_S" [color="0 0 0.5387640449438202", label="1.35",style=bold];
"MUC2_S" -> "IRX6_S" [color="0 0 0.5382022471910113", label="1.35",style=bold];
"FRMPD2_S" -> "PTPRZ1_S" [color="0 0 0.5376404494382022", label="1.35",style=bold];
"LPA_S" -> "APOB_E" [color="0 0 0.5370786516853933", label="1.36",style=bold];
"KLK5_S" -> "PCLO_S" [color="0 0 0.5365168539325842", label="1.36",style=bold];
"KRT16_S" -> "PGLYRP3_M" [color="0 0 0.5359550561797752", label="1.36",style=bold];
"MDGA2_S" -> "ZIC4_E" [color="0 0 0.5353932584269663", label="1.36",style=bold];
"CTSE_S" -> "CTSE_E" [color="0 0 0.5348314606741573", label="1.37",style=bold];
"PKHD1L1_S" -> "FER1L6_M" [color="0 0 0.5342696629213484", label="1.37",style=bold];
"ADAMTS20_S" -> "AGBL1_S" [color="0 0 0.5337078651685393", label="1.37",style=bold];
"SLC26A3_S" -> "LRRC7_E" [color="0 0 0.5331460674157303", label="1.37",style=bold];
"SERPINB13_S" -> "LRRC7_M" [color="0 0 0.5325842696629213", label="1.38",style=bold];
"SERPINB3_S" -> "MUC15_E" [color="0 0 0.5320224719101123", label="1.38",style=bold];
"KRT6C_S" -> "CPA2_E" [color="0 0 0.5314606741573034", label="1.38",style=bold];
"DPP6_S" -> "LRRC7_E" [color="0 0 0.5308988764044944", label="1.38",style=bold];
"A2ML1_S" -> "PCDHA12_S" [color="0 0 0.5303370786516854", label="1.39",style=bold];
"MUC15_S" -> "LRRC7_M" [color="0 0 0.5297752808988764", label="1.39",style=bold];
"PCDHA11_S" -> "SLC13A2_S" [color="0 0 0.5292134831460674", label="1.40",style=bold];
"SERPINB3_E" -> "SLC9A4_E" [color="0 0 0.5286516853932584", label="1.40",style=bold];
"KRT6C_S" -> "FREM2_S" [color="0 0 0.5280898876404494", label="1.40",style=bold];
"CR2_S" -> "SERPINB4_M" [color="0 0 0.5275280898876404", label="1.41",style=bold];
"FBN3_S" -> "CNTNAP2_E" [color="0 0 0.5269662921348315", label="1.42",style=bold];
"SLC9A4_S" -> "MUC16_S" [color="0 0 0.5264044943820225", label="1.42",style=bold];
"KRT16_S" -> "LPA_M" [color="0 0 0.5258426966292135", label="1.42",style=bold];
"PGC_S" -> "Survival" [color="0 0 0.5252808988764045", label="1.43",style=bold];
"GPR87_S" -> "FBN3_E" [color="0 0 0.5247191011235954", label="1.43",style=bold];
"KRT16_S" -> "FBN3_M" [color="0 0 0.5241573033707865", label="1.43",style=bold];
"CPA2_S" -> "PCK1_M" [color="0 0 0.5235955056179775", label="1.43",style=bold];
"DCDC1_S" -> "COL2A1_M" [color="0 0 0.5230337078651686", label="1.43",style=bold];
"PKHD1L1_E" -> "DMBT1_S" [color="0 0 0.5224719101123596", label="1.43",style=bold];
"PTPRZ1_S" -> "DCDC1_M" [color="0 0 0.5219101123595505", label="1.43",style=bold];
"CR2_S" -> "OGDHL_E" [color="0 0 0.5213483146067416", label="1.44",style=bold];
"KRT6C_S" -> "MDGA2_M" [color="0 0 0.5207865168539325", label="1.44",style=bold];
"SERPINB13_S" -> "ADAMTS20_E" [color="0 0 0.5202247191011236", label="1.45",style=bold];
"CNTN1_S" -> "PCK1_E" [color="0 0 0.5196629213483146", label="1.45",style=bold];
"SERPINB3_S" -> "MYH4_E" [color="0 0 0.5191011235955056", label="1.45",style=bold];
"GLRA1_S" -> "PCDH10_E" [color="0 0 0.5185393258426967", label="1.46",style=bold];
"IRX6_S" -> "DSG1_M" [color="0 0 0.5179775280898876", label="1.46",style=bold];
"SLC26A3_S" -> "MDGA2_E" [color="0 0 0.5174157303370787", label="1.46",style=bold];
"CPA2_S" -> "LRRC7_E" [color="0 0 0.5168539325842696", label="1.47",style=bold];
"CSMD3_M" -> "PCDHA12_M" [color="0 0 0.5162921348314606", label="1.47",style=bold];
"SERPINB3_S" -> "RIMS2_S" [color="0 0 0.5157303370786517", label="1.47",style=bold];
"SLC13A2_S" -> "APOB_S" [color="0 0 0.5151685393258427", label="1.47",style=bold];
"IRX6_S" -> "SERPINB4_M" [color="0 0 0.5146067415730338", label="1.47",style=bold];
"SERPINB13_S" -> "SERPINB3_M" [color="0 0 0.5140449438202247", label="1.47",style=bold];
"CNTN1_S" -> "AGBL1_M" [color="0 0 0.5134831460674157", label="1.48",style=bold];
"SLITRK1_S" -> "THBS4_M" [color="0 0 0.5129213483146067", label="1.48",style=bold];
"FREM2_S" -> "KRT6C_M" [color="0 0 0.5123595505617977", label="1.49",style=bold];
"MYH4_S" -> "DPP6_E" [color="0 0 0.5117977528089888", label="1.49",style=bold];
"PNLIPRP2_S" -> "GLRA1_S" [color="0 0 0.5112359550561798", label="1.49",style=bold];
"PCDHA11_S" -> "APOB_S" [color="0 0 0.5106741573033708", label="1.50",style=bold];
"FAM83C_S" -> "CACNA1B_M" [color="0 0 0.5101123595505618", label="1.50",style=bold];
"GLRA1_S" -> "A2ML1_M" [color="0 0 0.5095505617977528", label="1.50",style=bold];
"APOB_M" -> "DPP6_M" [color="0 0 0.5089887640449438", label="1.51",style=bold];
"OGDHL_S" -> "SLCO1B1_M" [color="0 0 0.5084269662921348", label="1.51",style=bold];
"KCNA1_S" -> "CPA2_M" [color="0 0 0.5078651685393258", label="1.51",style=bold];
"PCDHA12_S" -> "SLC9A11_E" [color="0 0 0.5073033707865169", label="1.52",style=bold];
"C20orf114_S" -> "DSG1_E" [color="0 0 0.5067415730337079", label="1.52",style=bold];
"CR2_S" -> "AGBL1_M" [color="0 0 0.5061797752808989", label="1.52",style=bold];
"KRT16_S" -> "FRMPD2_M" [color="0 0 0.5056179775280899", label="1.52",style=bold];
"C20orf114_S" -> "DMBT1_M" [color="0 0 0.5050561797752808", label="1.52",style=bold];
"PGC_S" -> "FREM2_S" [color="0 0 0.5044943820224719", label="1.53",style=bold];
"PNLIPRP2_S" -> "MUC6_M" [color="0 0 0.5039325842696629", label="1.53",style=bold];
"SERPINB13_S" -> "FER1L6_S" [color="0 0 0.503370786516854", label="1.53",style=bold];
"SLC13A2_S" -> "RIMS2_M" [color="0 0 0.502808988764045", label="1.54",style=bold];
"PCDH10_S" -> "MUC2_E" [color="0 0 0.5022471910112359", label="1.54",style=bold];
"FBN3_S" -> "MUC5B_S" [color="0 0 0.5016853932584269", label="1.54",style=bold];
"SLC26A3_S" -> "MUC15_M" [color="0 0 0.5011235955056179", label="1.54",style=bold];
"LPA_S" -> "SLC6A19_M" [color="0 0 0.500561797752809", label="1.55",style=bold];
"FAM83C_S" -> "PCDH10_E" [color="0 0 0.5", label="1.55",style=bold];
"ADAMTS20_S" -> "SLC13A2_M" [color="0 0 0.499438202247191", label="1.55",style=bold];
"ADAMTS20_S" -> "DCDC1_M" [color="0 0 0.49887640449438203", label="1.55",style=bold];
"CNTN1_S" -> "FER1L6_S" [color="0 0 0.498314606741573", label="1.55",style=bold];
"SLC6A19_S" -> "SERPINB4_M" [color="0 0 0.497752808988764", label="1.56",style=bold];
"SLC26A3_S" -> "FBN3_M" [color="0 0 0.49719101123595505", label="1.56",style=bold];
"GLRA1_S" -> "CSMD3_S" [color="0 0 0.49662921348314604", label="1.57",style=bold];
"SLC13A2_S" -> "SLC9A4_M" [color="0 0 0.4960674157303371", label="1.58",style=bold];
"DPP6_S" -> "RP1_E" [color="0 0 0.49550561797752807", label="1.59",style=bold];
"DSG1_S" -> "KRT6B_M" [color="0 0 0.4949438202247191", label="1.59",style=bold];
"CWH43_S" -> "FBN3_E" [color="0 0 0.4943820224719101", label="1.59",style=bold];
"MUC2_S" -> "LRRC7_M" [color="0 0 0.4938202247191011", label="1.60",style=bold];
"RFX6_S" -> "RP1_E" [color="0 0 0.49325842696629213", label="1.60",style=bold];
"PGLYRP3_S" -> "SLCO1B1_E" [color="0 0 0.4926966292134831", label="1.60",style=bold];
"MUC15_S" -> "ADAMTS20_E" [color="0 0 0.49213483146067416", label="1.60",style=bold];
"COL2A1_S" -> "THBS4_E" [color="0 0 0.49157303370786515", label="1.60",style=bold];
"PNLIPRP2_S" -> "SLC6A19_E" [color="0 0 0.4910112359550562", label="1.61",style=bold];
"SLCO1B3_S" -> "MUC6_E" [color="0 0 0.4904494382022472", label="1.61",style=bold];
"LPA_S" -> "SLC9A4_E" [color="0 0 0.48988764044943817", label="1.61",style=bold];
"SERPINB3_S" -> "MUC6_M" [color="0 0 0.4893258426966292", label="1.62",style=bold];
"NETO1_S" -> "FREM2_M" [color="0 0 0.4887640449438202", label="1.62",style=bold];
"SERPINB3_S" -> "SLC26A3_M" [color="0 0 0.48820224719101124", label="1.62",style=bold];
"DSG1_S" -> "CWH43_S" [color="0 0 0.48764044943820223", label="1.62",style=bold];
"GPR87_S" -> "MYH4_E" [color="0 0 0.48707865168539327", label="1.62",style=bold];
"PGLYRP3_S" -> "RP1_E" [color="0 0 0.48651685393258426", label="1.62",style=bold];
"SERPINB3_S" -> "FSTL5_M" [color="0 0 0.48595505617977525", label="1.63",style=bold];
"SLC6A19_S" -> "COL11A1_M" [color="0 0 0.4853932584269663", label="1.63",style=bold];
"PGC_S" -> "DMBT1_M" [color="0 0 0.4848314606741573", label="1.64",style=bold];
"PNLIPRP2_S" -> "SLC13A2_E" [color="0 0 0.4842696629213483", label="1.64",style=bold];
"THBS4_S" -> "RP1_S" [color="0 0 0.4837078651685393", label="1.64",style=bold];
"TGM5_S" -> "CPA2_E" [color="0 0 0.4831460674157303", label="1.64",style=bold];
"PGLYRP3_S" -> "PKHD1L1_E" [color="0 0 0.48258426966292134", label="1.65",style=bold];
"ABCA12_S" -> "RP1_S" [color="0 0 0.4820224719101123", label="1.66",style=bold];
"C20orf114_S" -> "LPA_E" [color="0 0 0.48146067415730337", label="1.66",style=bold];
"SLC13A2_S" -> "NETO1_E" [color="0 0 0.48089887640449436", label="1.66",style=bold];
"GPR87_E" -> "KRT6B_M" [color="0 0 0.4803370786516854", label="1.67",style=bold];
"MDGA2_S" -> "MUC5B_E" [color="0 0 0.4797752808988764", label="1.67",style=bold];
"PGLYRP3_S" -> "MUC5B_E" [color="0 0 0.4792134831460674", label="1.67",style=bold];
"SLCO1B1_S" -> "ADAMTS20_E" [color="0 0 0.4786516853932584", label="1.68",style=bold];
"KLK5_S" -> "A2ML1_S" [color="0 0 0.4780898876404494", label="1.69",style=bold];
"KRT6B_S" -> "SLC26A3_S" [color="0 0 0.47752808988764045", label="1.69",style=bold];
"PGC_S" -> "GPR87_E" [color="0 0 0.47696629213483144", label="1.71",style=bold];
"SLCO1B1_S" -> "PTPRZ1_M" [color="0 0 0.4764044943820225", label="1.71",style=bold];
"SERPINB4_S" -> "DMBT1_E" [color="0 0 0.47584269662921347", label="1.71",style=bold];
"FER1L6_S" -> "MUC6_E" [color="0 0 0.47528089887640446", label="1.71",style=bold];
"GLRA1_S" -> "SERPINB13_S" [color="0 0 0.4747191011235955", label="1.71",style=bold];
"SERPINB13_S" -> "PKHD1L1_M" [color="0 0 0.4741573033707865", label="1.71",style=bold];
"PGC_S" -> "FLG_S" [color="0 0 0.47359550561797753", label="1.72",style=bold];
"MUC15_S" -> "FRMPD2_E" [color="0 0 0.4730337078651685", label="1.73",style=bold];
"KRT16_S" -> "RP1_E" [color="0 0 0.47247191011235956", label="1.73",style=bold];
"GLRA1_S" -> "ADAMTS20_E" [color="0 0 0.47191011235955055", label="1.73",style=bold];
"SLITRK1_S" -> "MUC6_E" [color="0 0 0.47134831460674154", label="1.73",style=bold];
"SLC6A19_S" -> "MYH4_M" [color="0 0 0.4707865168539326", label="1.73",style=bold];
"SERPINB4_S" -> "KRT13_E" [color="0 0 0.47022471910112357", label="1.73",style=bold];
"CWH43_S" -> "SLC13A2_E" [color="0 0 0.4696629213483146", label="1.74",style=bold];
"SERPINB3_S" -> "PGLYRP3_M" [color="0 0 0.4691011235955056", label="1.74",style=bold];
"PCLO_S" -> "DCDC1_M" [color="0 0 0.46853932584269664", label="1.74",style=bold];
"PNLIPRP2_S" -> "SERPINB13_M" [color="0 0 0.46797752808988763", label="1.75",style=bold];
"PGC_S" -> "MUC2_E" [color="0 0 0.4674157303370786", label="1.75",style=bold];
"CWH43_S" -> "S100A7_M" [color="0 0 0.46685393258426966", label="1.76",style=bold];
"SERPINB13_S" -> "AGBL1_M" [color="0 0 0.46629213483146065", label="1.76",style=bold];
"KRT13_S" -> "MUC6_E" [color="0 0 0.4657303370786517", label="1.76",style=bold];
"ADAMTS20_S" -> "IRX6_M" [color="0 0 0.4651685393258427", label="1.76",style=bold];
"ADAMTS20_S" -> "MUC2_S" [color="0 0 0.46460674157303367", label="1.77",style=bold];
"SERPINB4_S" -> "CR2_S" [color="0 0 0.4640449438202247", label="1.77",style=bold];
"C20orf114_S" -> "CNTNAP2_S" [color="0 0 0.4634831460674157", label="1.77",style=bold];
"SLC26A3_S" -> "PGC_M" [color="0 0 0.46292134831460674", label="1.78",style=bold];
"KRT13_S" -> "PCLO_M" [color="0 0 0.4623595505617977", label="1.78",style=bold];
"KLK5_S" -> "KRT16_E" [color="0 0 0.46179775280898877", label="1.79",style=bold];
"PGC_S" -> "LPA_M" [color="0 0 0.46123595505617976", label="1.80",style=bold];
"DSG1_S" -> "CNTNAP2_E" [color="0 0 0.46067415730337075", label="1.80",style=bold];
"KRT16_S" -> "A2ML1_M" [color="0 0 0.4601123595505618", label="1.80",style=bold];
"SERPINB4_S" -> "NETO1_M" [color="0 0 0.4595505617977528", label="1.80",style=bold];
"SLC9A11_S" -> "RFX6_S" [color="0 0 0.4589887640449438", label="1.80",style=bold];
"FAM83C_S" -> "PCLO_S" [color="0 0 0.4584269662921348", label="1.82",style=bold];
"NETO1_S" -> "DPP6_E" [color="0 0 0.45786516853932585", label="1.82",style=bold];
"COL11A1_E" -> "Survival" [color="0 0 0.45730337078651684", label="1.83",style=bold];
"PGLYRP3_S" -> "Survival" [color="0 0 0.4567415730337078", label="1.83",style=bold];
"SLC6A19_S" -> "PGC_E" [color="0 0 0.45617977528089887", label="1.83",style=bold];
"SLCO1B3_S" -> "FSTL5_M" [color="0 0 0.45561797752808986", label="1.84",style=bold];
"SLC9A11_S" -> "MUC16_S" [color="0 0 0.4550561797752809", label="1.84",style=bold];
"CR2_S" -> "THBS4_S" [color="0 0 0.4544943820224719", label="1.84",style=bold];
"SERPINB13_S" -> "MUC4_M" [color="0 0 0.45393258426966293", label="1.84",style=bold];
"CACNA1B_S" -> "FRMPD2_E" [color="0 0 0.4533707865168539", label="1.85",style=bold];
"CPA2_S" -> "AGBL1_E" [color="0 0 0.4528089887640449", label="1.85",style=bold];
"COL2A1_S" -> "APOB_E" [color="0 0 0.45224719101123595", label="1.86",style=bold];
"MUC15_S" -> "MUC6_S" [color="0 0 0.45168539325842694", label="1.87",style=bold];
"PGLYRP3_S" -> "MUC2_E" [color="0 0 0.451123595505618", label="1.87",style=bold];
"C20orf114_S" -> "SLC26A3_E" [color="0 0 0.45056179775280897", label="1.87",style=bold];
"MUC17_E" -> "PNLIPRP2_E" [color="0 0 0.44999999999999996", label="1.88",style=bold];
"KLK5_S" -> "MDGA2_M" [color="0 0 0.449438202247191", label="1.88",style=bold];
"LPA_S" -> "MUC4_E" [color="0 0 0.448876404494382", label="1.88",style=bold];
"SLC13A2_S" -> "SLC9A4_S" [color="0 0 0.44831460674157303", label="1.88",style=bold];
"LRRC7_S" -> "MUC5B_E" [color="0 0 0.447752808988764", label="1.89",style=bold];
"KRT6C_S" -> "SERPINB4_S" [color="0 0 0.44719101123595506", label="1.89",style=bold];
"KRT16_S" -> "A2ML1_E" [color="0 0 0.44662921348314605", label="1.90",style=bold];
"TGM5_S" -> "CPA2_M" [color="0 0 0.44606741573033704", label="1.90",style=bold];
"ADAMTS20_S" -> "FREM2_E" [color="0 0 0.4455056179775281", label="1.90",style=bold];
"DMBT1_S" -> "MUC5B_S" [color="0 0 0.44494382022471907", label="1.92",style=bold];
"PNLIPRP2_S" -> "MUC15_M" [color="0 0 0.4443820224719101", label="1.92",style=bold];
"SLCO1B3_S" -> "KLK5_M" [color="0 0 0.4438202247191011", label="1.92",style=bold];
"FAM83C_S" -> "FRMPD2_M" [color="0 0 0.44325842696629214", label="1.93",style=bold];
"GLRA1_S" -> "CACNA1B_S" [color="0 0 0.44269662921348313", label="1.93",style=bold];
"PGLYRP3_S" -> "SERPINB4_M" [color="0 0 0.4421348314606741", label="1.93",style=bold];
"KCNA1_S" -> "Survival" [color="0 0 0.44157303370786516", label="1.93",style=bold];
"MUC6_S" -> "FREM2_E" [color="0 0 0.44101123595505615", label="1.94",style=bold];
"C20orf114_S" -> "SLC13A2_M" [color="0 0 0.4404494382022472", label="1.94",style=bold];
"NETO1_S" -> "PGLYRP3_E" [color="0 0 0.4398876404494382", label="1.95",style=bold];
"SLC9A4_S" -> "COL11A1_E" [color="0 0 0.4393258426966292", label="1.95",style=bold];
"DCDC1_S" -> "FREM2_E" [color="0 0 0.4387640449438202", label="1.96",style=bold];
"SLC9A4_S" -> "Survival" [color="0 0 0.4382022471910112", label="1.96",style=bold];
"PGLYRP3_S" -> "RP1_M" [color="0 0 0.43764044943820224", label="1.96",style=bold];
"ABCA12_S" -> "SLC9A4_M" [color="0 0 0.4370786516853932", label="1.96",style=bold];
"RFX6_S" -> "KRT16_M" [color="0 0 0.43651685393258427", label="1.97",style=bold];
"GPR87_M" -> "DCDC1_M" [color="0 0 0.43595505617977526", label="1.97",style=bold];
"PNLIPRP2_S" -> "SLC9A11_S" [color="0 0 0.4353932584269663", label="1.97",style=bold];
"SERPINB4_S" -> "PKHD1L1_E" [color="0 0 0.4348314606741573", label="1.98",style=bold];
"FLG_E" -> "MUC6_S" [color="0 0 0.4342696629213483", label="1.98",style=bold];
"GPR87_S" -> "PNLIPRP2_S" [color="0 0 0.4337078651685393", label="1.98",style=bold];
"OGDHL_M" -> "ABCA13_S" [color="0 0 0.4331460674157303", label="1.98",style=bold];
"PCDHA12_S" -> "FREM2_E" [color="0 0 0.43258426966292135", label="1.99",style=bold];
"A2ML1_S" -> "KCNA1_S" [color="0 0 0.43202247191011234", label="1.99",style=bold];
"ACTN2_S" -> "RIMS2_E" [color="0 0 0.4314606741573033", label="1.99",style=bold];
"SLC9A4_S" -> "MUC17_M" [color="0 0 0.43089887640449437", label="2.00",style=bold];
"MUC15_S" -> "SLITRK1_S" [color="0 0 0.43033707865168536", label="2.00",style=bold];
"SLC9A11_S" -> "ACTN2_E" [color="0 0 0.4297752808988764", label="2.01",style=bold];
"SLITRK1_S" -> "DCDC1_E" [color="0 0 0.4292134831460674", label="2.03",style=bold];
"SLITRK1_M" -> "CSMD3_M" [color="0 0 0.42865168539325843", label="2.03",style=bold];
"CACNA1B_E" -> "ZIC4_E" [color="0 0 0.4280898876404494", label="2.03",style=bold];
"KLK5_S" -> "LPA_M" [color="0 0 0.4275280898876404", label="2.03",style=bold];
"MUC15_S" -> "SLC9A11_M" [color="0 0 0.42696629213483145", label="2.03",style=bold];
"KRT16_S" -> "CNTN1_S" [color="0 0 0.42640449438202244", label="2.03",style=bold];
"SLC13A2_S" -> "PCK1_M" [color="0 0 0.4258426966292135", label="2.04",style=bold];
"KLK5_S" -> "FREM2_E" [color="0 0 0.42528089887640447", label="2.04",style=bold];
"PGC_S" -> "RP1_S" [color="0 0 0.4247191011235955", label="2.04",style=bold];
"SLC13A2_S" -> "COL11A1_E" [color="0 0 0.4241573033707865", label="2.05",style=bold];
"NETO1_E" -> "CACNA1B_S" [color="0 0 0.4235955056179775", label="2.05",style=bold];
"A2ML1_S" -> "GLRA1_E" [color="0 0 0.42303370786516853", label="2.06",style=bold];
"ABCA13_S" -> "PCDH10_E" [color="0 0 0.4224719101123595", label="2.06",style=bold];
"ABCA13_S" -> "APOB_S" [color="0 0 0.42191011235955056", label="2.06",style=bold];
"APOB_S" -> "LPA_S" [color="0 0 0.42134831460674155", label="2.07",style=bold];
"PGLYRP3_S" -> "CTSE_E" [color="0 0 0.4207865168539326", label="2.07",style=bold];
"SERPINB13_S" -> "S100A7_E" [color="0 0 0.4202247191011236", label="2.07",style=bold];
"MUC17_S" -> "NETO1_S" [color="0 0 0.41966292134831457", label="2.07",style=bold];
"OGDHL_S" -> "KLK5_E" [color="0 0 0.4191011235955056", label="2.08",style=bold];
"PGC_S" -> "SLC9A4_E" [color="0 0 0.4185393258426966", label="2.08",style=bold];
"PGC_S" -> "LPA_E" [color="0 0 0.41797752808988764", label="2.08",style=bold];
"PNLIPRP2_S" -> "FREM2_E" [color="0 0 0.4174157303370786", label="2.08",style=bold];
"SERPINB4_S" -> "TGM5_M" [color="0 0 0.4168539325842696", label="2.08",style=bold];
"COL2A1_S" -> "DCDC1_M" [color="0 0 0.41629213483146066", label="2.09",style=bold];
"CTSE_S" -> "CNTNAP2_S" [color="0 0 0.41573033707865165", label="2.09",style=bold];
"SERPINB13_S" -> "NETO1_M" [color="0 0 0.4151685393258427", label="2.09",style=bold];
"ACTN2_S" -> "PCDHA12_E" [color="0 0 0.4146067415730337", label="2.10",style=bold];
"KCNA1_S" -> "MUC17_S" [color="0 0 0.4140449438202247", label="2.12",style=bold];
"FAM83C_S" -> "ACTN2_E" [color="0 0 0.4134831460674157", label="2.12",style=bold];
"EGF_S" -> "COL11A1_E" [color="0 0 0.4129213483146067", label="2.13",style=bold];
"IRX6_M" -> "THBS4_M" [color="0 0 0.41235955056179774", label="2.13",style=bold];
"SLCO1B3_S" -> "A2ML1_M" [color="0 0 0.4117977528089887", label="2.13",style=bold];
"GPR98_E" -> "PCLO_E" [color="0 0 0.41123595505617977", label="2.13",style=bold];
"KRT16_S" -> "KRT13_M" [color="0 0 0.41067415730337076", label="2.13",style=bold];
"SLC6A19_S" -> "ABCA13_M" [color="0 0 0.4101123595505618", label="2.15",style=bold];
"KRT6C_E" -> "KLK5_E" [color="0 0 0.4095505617977528", label="2.15",style=bold];
"A2ML1_S" -> "MUC5B_E" [color="0 0 0.4089887640449438", label="2.16",style=bold];
"SERPINB4_S" -> "NETO1_E" [color="0 0 0.4084269662921348", label="2.16",style=bold];
"KCNA1_S" -> "SLC9A11_E" [color="0 0 0.4078651685393258", label="2.16",style=bold];
"CNTN1_S" -> "FSTL5_M" [color="0 0 0.40730337078651685", label="2.16",style=bold];
"FAM83C_S" -> "KRT6C_E" [color="0 0 0.40674157303370784", label="2.16",style=bold];
"APOB_S" -> "SLC26A3_E" [color="0 0 0.4061797752808989", label="2.17",style=bold];
"KLK5_S" -> "C20orf114_M" [color="0 0 0.40561797752808987", label="2.17",style=bold];
"PGC_S" -> "KRT6C_E" [color="0 0 0.40505617977528086", label="2.18",style=bold];
"GLRA1_S" -> "MUC5B_S" [color="0 0 0.4044943820224719", label="2.18",style=bold];
"TGM5_S" -> "MUC5B_S" [color="0 0 0.4039325842696629", label="2.18",style=bold];
"SERPINB3_S" -> "GPR98_S" [color="0 0 0.40337078651685393", label="2.18",style=bold];
"DSG1_S" -> "COL11A1_E" [color="0 0 0.4028089887640449", label="2.18",style=bold];
"SLC9A11_S" -> "SLC6A19_M" [color="0 0 0.40224719101123596", label="2.18",style=bold];
"C20orf114_S" -> "A2ML1_S" [color="0 0 0.40168539325842695", label="2.19",style=bold];
"DMBT1_S" -> "IRX6_S" [color="0 0 0.40112359550561794", label="2.19",style=bold];
"TGM5_S" -> "RP1_S" [color="0 0 0.400561797752809", label="2.19",style=bold];
"KRT16_S" -> "FREM2_M" [color="0 0 0.4", label="2.19",style=bold];
"KRT6B_S" -> "MYH4_E" [color="0 0 0.39943820224719095", label="2.20",style=bold];
"KRT13_S" -> "ABCA13_E" [color="0 0 0.398876404494382", label="2.21",style=bold];
"SLC13A2_S" -> "COL2A1_M" [color="0 0 0.39831460674157304", label="2.22",style=bold];
"PGC_S" -> "ACTN2_S" [color="0 0 0.397752808988764", label="2.22",style=bold];
"MUC15_S" -> "ZIC4_E" [color="0 0 0.397191011235955", label="2.22",style=bold];
"FAM83C_S" -> "CSMD3_S" [color="0 0 0.39662921348314606", label="2.22",style=bold];
"SLC26A3_S" -> "DPP6_E" [color="0 0 0.3960674157303371", label="2.23",style=bold];
"C20orf114_S" -> "GLRA1_E" [color="0 0 0.39550561797752803", label="2.23",style=bold];
"KRT6C_S" -> "MUC5B_S" [color="0 0 0.3949438202247191", label="2.23",style=bold];
"DSG1_S" -> "DSG1_M" [color="0 0 0.3943820224719101", label="2.25",style=bold];
"MYH4_S" -> "FRMPD2_M" [color="0 0 0.39382022471910105", label="2.25",style=bold];
"TGM5_S" -> "LRRC7_E" [color="0 0 0.3932584269662921", label="2.25",style=bold];
"LPA_S" -> "Survival" [color="0 0 0.39269662921348314", label="2.25",style=bold];
"SLC26A3_S" -> "MUC16_E" [color="0 0 0.3921348314606742", label="2.26",style=bold];
"MUC15_S" -> "MDGA2_S" [color="0 0 0.3915730337078651", label="2.27",style=bold];
"SERPINB3_S" -> "SLCO1B1_S" [color="0 0 0.39101123595505616", label="2.28",style=bold];
"KRT6B_S" -> "PGC_M" [color="0 0 0.3904494382022472", label="2.28",style=bold];
"Survival" -> "PCLO_S" [color="0 0 0.38988764044943813", label="2.28",style=bold];
"ADAMTS20_S" -> "ABCA12_S" [color="0 0 0.3893258426966292", label="2.28",style=bold];
"PNLIPRP2_S" -> "ADAMTS20_E" [color="0 0 0.3887640449438202", label="2.28",style=bold];
"PNLIPRP2_S" -> "GPR98_S" [color="0 0 0.38820224719101126", label="2.29",style=bold];
"RFX6_S" -> "DMBT1_E" [color="0 0 0.3876404494382022", label="2.29",style=bold];
"GPR87_S" -> "PCDHA11_S" [color="0 0 0.38707865168539324", label="2.29",style=bold];
"PGLYRP3_S" -> "THBS4_M" [color="0 0 0.3865168539325843", label="2.29",style=bold];
"SLCO1B1_S" -> "PKHD1L1_S" [color="0 0 0.3859550561797752", label="2.29",style=bold];
"SERPINB13_S" -> "PCDHA11_M" [color="0 0 0.38539325842696626", label="2.31",style=bold];
"KRT13_S" -> "FREM2_S" [color="0 0 0.3848314606741573", label="2.31",style=bold];
"LRRC7_S" -> "THBS4_S" [color="0 0 0.38426966292134834", label="2.33",style=bold];
"MUC15_S" -> "SERPINB4_E" [color="0 0 0.3837078651685393", label="2.33",style=bold];
"S100A7_E" -> "LRRC7_E" [color="0 0 0.3831460674157303", label="2.33",style=bold];
"FREM2_S" -> "MUC6_E" [color="0 0 0.38258426966292136", label="2.33",style=bold];
"GPR87_S" -> "RIMS2_S" [color="0 0 0.3820224719101123", label="2.34",style=bold];
"KRT13_S" -> "MDGA2_M" [color="0 0 0.38146067415730334", label="2.34",style=bold];
"GLRA1_M" -> "MDGA2_M" [color="0 0 0.3808988764044944", label="2.35",style=bold];
"OGDHL_S" -> "SLC9A4_S" [color="0 0 0.3803370786516854", label="2.36",style=bold];
"RP1_S" -> "MUC5B_S" [color="0 0 0.37977528089887636", label="2.36",style=bold];
"DCDC1_S" -> "EGF_M" [color="0 0 0.3792134831460674", label="2.37",style=bold];
"PNLIPRP2_S" -> "FSTL5_E" [color="0 0 0.37865168539325844", label="2.38",style=bold];
"PGC_S" -> "EGF_M" [color="0 0 0.3780898876404494", label="2.38",style=bold];
"SLC9A4_S" -> "MUC6_S" [color="0 0 0.3775280898876404", label="2.38",style=bold];
"DSG1_S" -> "CNTNAP2_S" [color="0 0 0.37696629213483146", label="2.38",style=bold];
"PCDH10_S" -> "SLC9A11_E" [color="0 0 0.3764044943820225", label="2.39",style=bold];
"PTPRZ1_S" -> "PTPRZ1_M" [color="0 0 0.37584269662921344", label="2.39",style=bold];
"SERPINB4_S" -> "SLCO1B1_M" [color="0 0 0.3752808988764045", label="2.41",style=bold];
"SERPINB3_S" -> "DMBT1_M" [color="0 0 0.3747191011235955", label="2.41",style=bold];
"KCNA1_S" -> "KLK5_E" [color="0 0 0.37415730337078645", label="2.41",style=bold];
"KLK5_S" -> "SLC13A2_E" [color="0 0 0.3735955056179775", label="2.41",style=bold];
"CTSE_S" -> "RFX6_E" [color="0 0 0.37303370786516854", label="2.42",style=bold];
"AGBL1_S" -> "FREM2_S" [color="0 0 0.37247191011235947", label="2.42",style=bold];
"THBS4_S" -> "MUC5B_S" [color="0 0 0.3719101123595505", label="2.43",style=bold];
"KRT6B_S" -> "S100A7_M" [color="0 0 0.37134831460674156", label="2.43",style=bold];
"FAM83C_S" -> "FER1L6_S" [color="0 0 0.3707865168539326", label="2.44",style=bold];
"FRMPD2_S" -> "PGLYRP3_M" [color="0 0 0.37022471910112353", label="2.46",style=bold];
"KLK5_S" -> "S100A7_M" [color="0 0 0.3696629213483146", label="2.47",style=bold];
"LPA_S" -> "MUC4_M" [color="0 0 0.3691011235955056", label="2.47",style=bold];
"SERPINB3_S" -> "ABCA13_S" [color="0 0 0.36853932584269655", label="2.47",style=bold];
"KRT6B_S" -> "PNLIPRP2_S" [color="0 0 0.3679775280898876", label="2.48",style=bold];
"C20orf114_S" -> "CR2_M" [color="0 0 0.36741573033707864", label="2.48",style=bold];
"KCNA1_S" -> "SLC9A11_S" [color="0 0 0.3668539325842697", label="2.48",style=bold];
"S100A7_S" -> "CNTN1_S" [color="0 0 0.3662921348314606", label="2.48",style=bold];
"PGC_S" -> "MDGA2_E" [color="0 0 0.36573033707865166", label="2.49",style=bold];
"SLCO1B1_S" -> "FBN3_M" [color="0 0 0.3651685393258427", label="2.50",style=bold];
"SLC13A2_S" -> "FREM2_E" [color="0 0 0.36460674157303363", label="2.52",style=bold];
"KRT13_S" -> "MUC15_M" [color="0 0 0.3640449438202247", label="2.52",style=bold];
"SLC6A19_S" -> "C20orf114_E" [color="0 0 0.3634831460674157", label="2.53",style=bold];
"CTSE_S" -> "ZIC4_S" [color="0 0 0.36292134831460676", label="2.53",style=bold];
"MYH4_S" -> "DSG1_M" [color="0 0 0.3623595505617977", label="2.54",style=bold];
"SLC9A4_S" -> "SLITRK1_S" [color="0 0 0.36179775280898874", label="2.54",style=bold];
"NETO1_S" -> "FREM2_E" [color="0 0 0.3612359550561798", label="2.54",style=bold];
"ABCA13_S" -> "MUC16_S" [color="0 0 0.3606741573033707", label="2.55",style=bold];
"SERPINB3_E" -> "ABCA13_E" [color="0 0 0.36011235955056176", label="2.56",style=bold];
"FAM83C_S" -> "MUC5B_M" [color="0 0 0.3595505617977528", label="2.57",style=bold];
"C20orf114_S" -> "S100A7_S" [color="0 0 0.35898876404494384", label="2.57",style=bold];
"CWH43_S" -> "SLC26A3_M" [color="0 0 0.3584269662921348", label="2.58",style=bold];
"COL11A1_M" -> "CNTN1_M" [color="0 0 0.3578651685393258", label="2.58",style=bold];
"FAM83C_S" -> "OGDHL_E" [color="0 0 0.35730337078651686", label="2.58",style=bold];
"KCNA1_S" -> "CACNA1B_E" [color="0 0 0.3567415730337078", label="2.58",style=bold];
"MUC15_S" -> "SERPINB3_M" [color="0 0 0.35617977528089884", label="2.59",style=bold];
"CWH43_S" -> "EGF_M" [color="0 0 0.3556179775280899", label="2.59",style=bold];
"PNLIPRP2_S" -> "A2ML1_S" [color="0 0 0.3550561797752809", label="2.59",style=bold];
"ZIC4_S" -> "DSG1_E" [color="0 0 0.35449438202247185", label="2.59",style=bold];
"MDGA2_S" -> "CNTNAP2_S" [color="0 0 0.3539325842696629", label="2.60",style=bold];
"TGM5_S" -> "ACTN2_E" [color="0 0 0.35337078651685394", label="2.60",style=bold];
"CR2_S" -> "SLC26A3_E" [color="0 0 0.3528089887640449", label="2.60",style=bold];
"RP1_S" -> "CTSE_S" [color="0 0 0.3522471910112359", label="2.61",style=bold];
"MUC16_S" -> "CR2_E" [color="0 0 0.35168539325842696", label="2.61",style=bold];
"ACTN2_S" -> "SLC13A2_M" [color="0 0 0.351123595505618", label="2.61",style=bold];
"CTSE_S" -> "KCNA1_E" [color="0 0 0.35056179775280893", label="2.62",style=bold];
"MUC15_S" -> "SLC13A2_E" [color="0 0 0.35", label="2.62",style=bold];
"RIMS2_S" -> "DCDC1_S" [color="0 0 0.349438202247191", label="2.64",style=bold];
"SERPINB4_S" -> "MDGA2_S" [color="0 0 0.34887640449438195", label="2.64",style=bold];
"DCDC1_S" -> "PNLIPRP2_M" [color="0 0 0.348314606741573", label="2.65",style=bold];
"LPA_S" -> "MUC6_E" [color="0 0 0.34775280898876404", label="2.65",style=bold];
"GPR87_S" -> "PKHD1L1_S" [color="0 0 0.3471910112359551", label="2.67",style=bold];
"SERPINB3_S" -> "GPR98_M" [color="0 0 0.346629213483146", label="2.68",style=bold];
"AGBL1_S" -> "ACTN2_E" [color="0 0 0.34606741573033706", label="2.68",style=bold];
"GLRA1_S" -> "NETO1_M" [color="0 0 0.3455056179775281", label="2.68",style=bold];
"SLC9A4_S" -> "MUC5B_S" [color="0 0 0.34494382022471903", label="2.70",style=bold];
"ZIC4_M" -> "SERPINB4_M" [color="0 0 0.3443820224719101", label="2.70",style=bold];
"MUC15_S" -> "CSMD3_E" [color="0 0 0.3438202247191011", label="2.71",style=bold];
"PGC_S" -> "SLCO1B1_E" [color="0 0 0.34325842696629216", label="2.71",style=bold];
"MUC6_S" -> "SLC9A11_M" [color="0 0 0.3426966292134831", label="2.71",style=bold];
"MUC2_M" -> "PKHD1L1_E" [color="0 0 0.34213483146067414", label="2.72",style=bold];
"DCDC1_S" -> "SLCO1B1_E" [color="0 0 0.3415730337078652", label="2.76",style=bold];
"ZIC4_S" -> "DCDC1_S" [color="0 0 0.3410112359550561", label="2.76",style=bold];
"DCDC1_S" -> "OGDHL_E" [color="0 0 0.34044943820224716", label="2.77",style=bold];
"SLC26A3_S" -> "KLK5_E" [color="0 0 0.3398876404494382", label="2.78",style=bold];
"ABCA13_S" -> "SLC9A4_S" [color="0 0 0.33932584269662913", label="2.79",style=bold];
"CNTNAP2_M" -> "PTPRZ1_M" [color="0 0 0.3387640449438202", label="2.81",style=bold];
"EGF_S" -> "KRT16_M" [color="0 0 0.3382022471910112", label="2.81",style=bold];
"KRT16_M" -> "KRT13_M" [color="0 0 0.33764044943820226", label="2.82",style=bold];
"PNLIPRP2_S" -> "SLCO1B3_S" [color="0 0 0.3370786516853932", label="2.82",style=bold];
"MUC6_S" -> "MUC4_S" [color="0 0 0.33651685393258424", label="2.82",style=bold];
"APOB_S" -> "SERPINB13_E" [color="0 0 0.3359550561797753", label="2.84",style=bold];
"KCNA1_S" -> "MUC6_S" [color="0 0 0.3353932584269662", label="2.84",style=bold];
"DSG1_S" -> "THBS4_E" [color="0 0 0.33483146067415726", label="2.86",style=bold];
"SLC9A4_S" -> "PNLIPRP2_M" [color="0 0 0.3342696629213483", label="2.87",style=bold];
"PNLIPRP2_S" -> "LPA_M" [color="0 0 0.33370786516853934", label="2.87",style=bold];
"KRT16_S" -> "CR2_S" [color="0 0 0.3331460674157303", label="2.87",style=bold];
"GLRA1_S" -> "KRT13_E" [color="0 0 0.3325842696629213", label="2.87",style=bold];
"IRX6_S" -> "GPR87_M" [color="0 0 0.33202247191011236", label="2.90",style=bold];
"KRT13_S" -> "RFX6_S" [color="0 0 0.3314606741573033", label="2.91",style=bold];
"MUC15_S" -> "MUC2_E" [color="0 0 0.33089887640449434", label="2.91",style=bold];
"FRMPD2_S" -> "SLC13A2_M" [color="0 0 0.3303370786516854", label="2.91",style=bold];
"DCDC1_S" -> "CSMD3_S" [color="0 0 0.3297752808988764", label="2.92",style=bold];
"THBS4_M" -> "PCDHA12_M" [color="0 0 0.32921348314606735", label="2.93",style=bold];
"KRT6C_E" -> "PCDHA11_S" [color="0 0 0.3286516853932584", label="2.93",style=bold];
"ACTN2_S" -> "FBN3_E" [color="0 0 0.32808988764044944", label="2.93",style=bold];
"PCK1_S" -> "DSG1_M" [color="0 0 0.3275280898876404", label="2.94",style=bold];
"GPR87_S" -> "ZIC4_S" [color="0 0 0.3269662921348314", label="2.94",style=bold];
"SLCO1B3_S" -> "Survival" [color="0 0 0.32640449438202246", label="2.95",style=bold];
"MDGA2_S" -> "ABCA12_M" [color="0 0 0.3258426966292135", label="2.95",style=bold];
"CWH43_S" -> "FRMPD2_E" [color="0 0 0.32528089887640443", label="2.95",style=bold];
"FRMPD2_S" -> "DPP6_S" [color="0 0 0.3247191011235955", label="2.95",style=bold];
"CWH43_S" -> "FSTL5_E" [color="0 0 0.3241573033707865", label="2.97",style=bold];
"FAM83C_S" -> "FRMPD2_S" [color="0 0 0.32359550561797745", label="2.98",style=bold];
"SLC9A11_S" -> "MUC4_M" [color="0 0 0.3230337078651685", label="2.98",style=bold];
"SERPINB4_S" -> "PCDHA12_S" [color="0 0 0.32247191011235954", label="2.98",style=bold];
"APOB_S" -> "ACTN2_S" [color="0 0 0.3219101123595506", label="3.00",style=bold];
"FRMPD2_S" -> "ABCA13_M" [color="0 0 0.3213483146067415", label="3.01",style=bold];
"CWH43_S" -> "PKHD1L1_E" [color="0 0 0.32078651685393256", label="3.01",style=bold];
"KRT13_S" -> "SLC9A11_M" [color="0 0 0.3202247191011236", label="3.03",style=bold];
"OGDHL_S" -> "RP1_M" [color="0 0 0.31966292134831453", label="3.03",style=bold];
"MUC6_S" -> "ACTN2_S" [color="0 0 0.3191011235955056", label="3.05",style=bold];
"SLCO1B3_S" -> "APOB_E" [color="0 0 0.3185393258426966", label="3.05",style=bold];
"PCK1_S" -> "SERPINB4_M" [color="0 0 0.31797752808988766", label="3.06",style=bold];
"THBS4_S" -> "SERPINB4_M" [color="0 0 0.3174157303370786", label="3.08",style=bold];
"SLCO1B3_S" -> "GPR98_S" [color="0 0 0.31685393258426964", label="3.08",style=bold];
"GPR87_S" -> "CNTNAP2_S" [color="0 0 0.3162921348314607", label="3.09",style=bold];
"FAM83C_S" -> "ZIC4_S" [color="0 0 0.3157303370786516", label="3.10",style=bold];
"SLC13A2_S" -> "MUC6_S" [color="0 0 0.31516853932584266", label="3.13",style=bold];
"SERPINB4_S" -> "FREM2_S" [color="0 0 0.3146067415730337", label="3.14",style=bold];
"PCLO_S" -> "FSTL5_S" [color="0 0 0.31404494382022474", label="3.14",style=bold];
"FBN3_S" -> "RP1_M" [color="0 0 0.3134831460674157", label="3.14",style=bold];
"KRT13_S" -> "FER1L6_S" [color="0 0 0.3129213483146067", label="3.14",style=bold];
"PCDHA12_S" -> "SERPINB3_M" [color="0 0 0.31235955056179776", label="3.15",style=bold];
"CSMD3_S" -> "DCDC1_M" [color="0 0 0.3117977528089887", label="3.16",style=bold];
"DPP6_S" -> "NETO1_E" [color="0 0 0.31123595505617974", label="3.16",style=bold];
"SLCO1B1_S" -> "FREM2_S" [color="0 0 0.3106741573033708", label="3.17",style=bold];
"KRT6B_S" -> "S100A7_E" [color="0 0 0.3101123595505618", label="3.22",style=bold];
"SLCO1B1_M" -> "SLCO1B3_M" [color="0 0 0.30955056179775275", label="3.22",style=bold];
"FER1L6_S" -> "AGBL1_E" [color="0 0 0.3089887640449438", label="3.22",style=bold];
"OGDHL_S" -> "SLC26A3_E" [color="0 0 0.30842696629213484", label="3.23",style=bold];
"MUC2_S" -> "APOB_E" [color="0 0 0.3078651685393258", label="3.23",style=bold];
"SLCO1B3_S" -> "PKHD1L1_M" [color="0 0 0.3073033707865168", label="3.24",style=bold];
"KCNA1_M" -> "CNTNAP2_S" [color="0 0 0.30674157303370786", label="3.25",style=bold];
"S100A7_S" -> "DSG1_S" [color="0 0 0.3061797752808988", label="3.25",style=bold];
"SLITRK1_S" -> "A2ML1_M" [color="0 0 0.30561797752808983", label="3.27",style=bold];
"AGBL1_S" -> "PGLYRP3_M" [color="0 0 0.3050561797752809", label="3.28",style=bold];
"PCK1_S" -> "SLC9A4_M" [color="0 0 0.3044943820224719", label="3.28",style=bold];
"SLITRK1_S" -> "KRT13_M" [color="0 0 0.30393258426966285", label="3.30",style=bold];
"MUC2_S" -> "THBS4_S" [color="0 0 0.3033707865168539", label="3.31",style=bold];
"KCNA1_M" -> "CNTN1_M" [color="0 0 0.30280898876404494", label="3.32",style=bold];
"PCDHA12_S" -> "MUC5B_S" [color="0 0 0.30224719101123587", label="3.32",style=bold];
"COL11A1_S" -> "APOB_S" [color="0 0 0.3016853932584269", label="3.36",style=bold];
"S100A7_M" -> "PGC_M" [color="0 0 0.30112359550561796", label="3.36",style=bold];
"FAM83C_S" -> "RIMS2_S" [color="0 0 0.300561797752809", label="3.37",style=bold];
"FRMPD2_S" -> "EGF_M" [color="0 0 0.29999999999999993", label="3.38",style=bold];
"CNTN1_S" -> "COL2A1_S" [color="0 0 0.299438202247191", label="3.38",style=bold];
"SLITRK1_S" -> "APOB_S" [color="0 0 0.298876404494382", label="3.39",style=bold];
"GPR87_E" -> "TGM5_E" [color="0 0 0.29831460674157295", label="3.39",style=bold];
"FAM83C_S" -> "SLCO1B1_S" [color="0 0 0.297752808988764", label="3.39",style=bold];
"FREM2_M" -> "PCDHA12_M" [color="0 0 0.29719101123595504", label="3.41",style=bold];
"PTPRZ1_S" -> "SERPINB3_M" [color="0 0 0.2966292134831461", label="3.42",style=bold];
"KRT16_E" -> "SLC9A4_E" [color="0 0 0.296067415730337", label="3.43",style=bold];
"SERPINB4_S" -> "LPA_S" [color="0 0 0.29550561797752806", label="3.43",style=bold];
"PNLIPRP2_S" -> "SLC9A4_S" [color="0 0 0.2949438202247191", label="3.44",style=bold];
"SLCO1B1_S" -> "SLC6A19_S" [color="0 0 0.29438202247191003", label="3.44",style=bold];
"S100A7_S" -> "SLC26A3_S" [color="0 0 0.2938202247191011", label="3.45",style=bold];
"SERPINB4_E" -> "ABCA13_E" [color="0 0 0.2932584269662921", label="3.45",style=bold];
"KRT6C_S" -> "IRX6_S" [color="0 0 0.29269662921348316", label="3.46",style=bold];
"SERPINB4_S" -> "SLC13A2_E" [color="0 0 0.2921348314606741", label="3.47",style=bold];
"FREM2_S" -> "CR2_E" [color="0 0 0.29157303370786514", label="3.47",style=bold];
"SLCO1B1_S" -> "NETO1_E" [color="0 0 0.2910112359550562", label="3.48",style=bold];
"KLK5_S" -> "ABCA13_S" [color="0 0 0.2904494382022471", label="3.49",style=bold];
"SERPINB13_S" -> "ABCA12_S" [color="0 0 0.28988764044943816", label="3.49",style=bold];
"PGC_S" -> "ABCA12_S" [color="0 0 0.2893258426966292", label="3.49",style=bold];
"PCDHA12_S" -> "SLCO1B3_M" [color="0 0 0.28876404494382024", label="3.50",style=bold];
"FAM83C_S" -> "KLK5_S" [color="0 0 0.2882022471910112", label="3.52",style=bold];
"SLC26A3_S" -> "GPR98_M" [color="0 0 0.2876404494382022", label="3.53",style=bold];
"MYH4_S" -> "CPA2_M" [color="0 0 0.28707865168539326", label="3.53",style=bold];
"CR2_S" -> "SLCO1B1_M" [color="0 0 0.2865168539325842", label="3.53",style=bold];
"ZIC4_S" -> "ABCA12_M" [color="0 0 0.28595505617977524", label="3.54",style=bold];
"PKHD1L1_E" -> "FLG_E" [color="0 0 0.2853932584269663", label="3.55",style=bold];
"ADAMTS20_S" -> "KLK5_M" [color="0 0 0.2848314606741573", label="3.56",style=bold];
"KCNA1_S" -> "CNTNAP2_S" [color="0 0 0.28426966292134825", label="3.57",style=bold];
"SLC9A11_S" -> "ACTN2_S" [color="0 0 0.2837078651685393", label="3.59",style=bold];
"FRMPD2_S" -> "COL11A1_E" [color="0 0 0.28314606741573034", label="3.61",style=bold];
"SLC13A2_S" -> "COL2A1_E" [color="0 0 0.2825842696629213", label="3.62",style=bold];
"ACTN2_S" -> "RP1_E" [color="0 0 0.2820224719101123", label="3.66",style=bold];
"KRT6C_S" -> "APOB_S" [color="0 0 0.28146067415730336", label="3.66",style=bold];
"SERPINB13_S" -> "COL11A1_S" [color="0 0 0.2808988764044944", label="3.67",style=bold];
"S100A7_S" -> "PNLIPRP2_S" [color="0 0 0.28033707865168533", label="3.74",style=bold];
"GPR87_S" -> "RP1_S" [color="0 0 0.2797752808988764", label="3.74",style=bold];
"FLG_S" -> "AGBL1_E" [color="0 0 0.2792134831460674", label="3.74",style=bold];
"CNTNAP2_S" -> "SLC13A2_E" [color="0 0 0.27865168539325835", label="3.75",style=bold];
"PCDHA11_S" -> "GPR98_S" [color="0 0 0.2780898876404494", label="3.75",style=bold];
"APOB_M" -> "PCK1_M" [color="0 0 0.27752808988764044", label="3.76",style=bold];
"KCNA1_S" -> "MUC5B_S" [color="0 0 0.2769662921348315", label="3.76",style=bold];
"FER1L6_S" -> "APOB_S" [color="0 0 0.2764044943820224", label="3.76",style=bold];
"KRT13_S" -> "ABCA12_S" [color="0 0 0.27584269662921346", label="3.76",style=bold];
"COL11A1_S" -> "RFX6_E" [color="0 0 0.2752808988764045", label="3.77",style=bold];
"GPR87_M" -> "LPA_E" [color="0 0 0.27471910112359543", label="3.77",style=bold];
"PCDH10_S" -> "KRT13_S" [color="0 0 0.2741573033707865", label="3.78",style=bold];
"OGDHL_M" -> "COL2A1_E" [color="0 0 0.2735955056179775", label="3.79",style=bold];
"AGBL1_M" -> "LRRC7_M" [color="0 0 0.27303370786516845", label="3.79",style=bold];
"FBN3_S" -> "SLC13A2_E" [color="0 0 0.2724719101123595", label="3.79",style=bold];
"SLC9A4_S" -> "GPR87_M" [color="0 0 0.27191011235955054", label="3.79",style=bold];
"FLG_M" -> "LRRC7_M" [color="0 0 0.2713483146067416", label="3.80",style=bold];
"SLC9A11_S" -> "NETO1_E" [color="0 0 0.2707865168539325", label="3.82",style=bold];
"CR2_S" -> "FREM2_E" [color="0 0 0.27022471910112356", label="3.83",style=bold];
"KRT13_S" -> "CWH43_S" [color="0 0 0.2696629213483146", label="3.85",style=bold];
"OGDHL_S" -> "PCK1_S" [color="0 0 0.26910112359550553", label="3.85",style=bold];
"IRX6_S" -> "IRX6_M" [color="0 0 0.2685393258426966", label="3.85",style=bold];
"PGC_S" -> "FAM83C_E" [color="0 0 0.2679775280898876", label="3.87",style=bold];
"FBN3_S" -> "TGM5_S" [color="0 0 0.26741573033707866", label="3.87",style=bold];
"SLC9A11_S" -> "SLC26A3_E" [color="0 0 0.2668539325842696", label="3.88",style=bold];
"KRT6C_S" -> "MUC4_S" [color="0 0 0.26629213483146064", label="3.89",style=bold];
"EGF_S" -> "S100A7_M" [color="0 0 0.2657303370786517", label="3.90",style=bold];
"SERPINB13_S" -> "MDGA2_M" [color="0 0 0.2651685393258426", label="3.93",style=bold];
"CPA2_S" -> "RFX6_S" [color="0 0 0.26460674157303365", label="3.97",style=bold];
"KLK5_S" -> "PCDH10_S" [color="0 0 0.2640449438202247", label="3.98",style=bold];
"ABCA12_S" -> "FSTL5_S" [color="0 0 0.26348314606741574", label="3.98",style=bold];
"CACNA1B_M" -> "KLK5_M" [color="0 0 0.2629213483146067", label="3.99",style=bold];
"CWH43_S" -> "GLRA1_E" [color="0 0 0.2623595505617977", label="3.99",style=bold];
"ABCA12_M" -> "EGF_M" [color="0 0 0.26179775280898876", label="3.99",style=bold];
"CNTN1_S" -> "GLRA1_E" [color="0 0 0.2612359550561797", label="3.99",style=bold];
"GPR87_S" -> "CR2_S" [color="0 0 0.26067415730337073", label="4.00",style=bold];
"DSG1_S" -> "NETO1_E" [color="0 0 0.2601123595505618", label="4.00",style=bold];
"CNTNAP2_M" -> "MUC5B_M" [color="0 0 0.2595505617977528", label="4.02",style=bold];
"KRT6C_S" -> "CNTNAP2_S" [color="0 0 0.25898876404494375", label="4.03",style=bold];
"PTPRZ1_E" -> "SLCO1B1_E" [color="0 0 0.2584269662921348", label="4.03",style=bold];
"NETO1_S" -> "SLC9A4_E" [color="0 0 0.25786516853932584", label="4.04",style=bold];
"KRT6C_E" -> "PCDH10_S" [color="0 0 0.25730337078651677", label="4.05",style=bold];
"GPR87_S" -> "NETO1_S" [color="0 0 0.2567415730337078", label="4.10",style=bold];
"MUC5B_S" -> "COL11A1_E" [color="0 0 0.25617977528089886", label="4.11",style=bold];
"ADAMTS20_S" -> "APOB_E" [color="0 0 0.2556179775280899", label="4.12",style=bold];
"FAM83C_E" -> "ABCA13_E" [color="0 0 0.25505617977528083", label="4.12",style=bold];
"CR2_S" -> "PCDHA12_S" [color="0 0 0.2544943820224719", label="4.17",style=bold];
"GLRA1_S" -> "TGM5_S" [color="0 0 0.2539325842696629", label="4.17",style=bold];
"S100A7_S" -> "CPA2_S" [color="0 0 0.25337078651685385", label="4.17",style=bold];
"FBN3_S" -> "DPP6_S" [color="0 0 0.2528089887640449", label="4.17",style=bold];
"CR2_S" -> "PTPRZ1_S" [color="0 0 0.25224719101123594", label="4.20",style=bold];
"PCK1_S" -> "ZIC4_S" [color="0 0 0.251685393258427", label="4.20",style=bold];
"SLC6A19_M" -> "LPA_E" [color="0 0 0.2511235955056179", label="4.20",style=bold];
"SLC26A3_E" -> "MUC5B_E" [color="0 0 0.25056179775280896", label="4.21",style=bold];
"APOB_S" -> "EGF_S" [color="0 0 0.25", label="4.22",style=bold];
"ABCA12_S" -> "SLC6A19_M" [color="0 0 0.24943820224719093", label="4.23",style=bold];
"SLC9A11_S" -> "KRT16_E" [color="0 0 0.24887640449438198", label="4.23",style=bold];
"GPR87_S" -> "SLC13A2_M" [color="0 0 0.24831460674157302", label="4.24",style=bold];
"SLCO1B1_S" -> "MUC17_S" [color="0 0 0.24775280898876406", label="4.24",style=bold];
"OGDHL_E" -> "SLCO1B3_E" [color="0 0 0.247191011235955", label="4.25",style=bold];
"SLC13A2_S" -> "ACTN2_E" [color="0 0 0.24662921348314604", label="4.27",style=bold];
"DMBT1_S" -> "APOB_S" [color="0 0 0.24606741573033708", label="4.31",style=bold];
"CNTN1_S" -> "MUC5B_E" [color="0 0 0.245505617977528", label="4.32",style=bold];
"SLC6A19_E" -> "PKHD1L1_S" [color="0 0 0.24494382022471906", label="4.32",style=bold];
"DMBT1_S" -> "PTPRZ1_E" [color="0 0 0.2443820224719101", label="4.33",style=bold];
"LRRC7_S" -> "SLCO1B3_S" [color="0 0 0.24382022471910114", label="4.34",style=bold];
"SLC6A19_S" -> "RP1_S" [color="0 0 0.24325842696629207", label="4.34",style=bold];
"CNTN1_S" -> "RP1_M" [color="0 0 0.24269662921348312", label="4.35",style=bold];
"GPR87_S" -> "FAM83C_S" [color="0 0 0.24213483146067416", label="4.35",style=bold];
"OGDHL_M" -> "GPR98_M" [color="0 0 0.2415730337078651", label="4.35",style=bold];
"SERPINB4_E" -> "KRT16_E" [color="0 0 0.24101123595505614", label="4.36",style=bold];
"DPP6_S" -> "Survival" [color="0 0 0.24044943820224718", label="4.37",style=bold];
"FER1L6_S" -> "KCNA1_S" [color="0 0 0.2398876404494381", label="4.37",style=bold];
"SLC9A4_S" -> "MUC4_S" [color="0 0 0.23932584269662915", label="4.37",style=bold];
"GPR98_S" -> "MUC17_S" [color="0 0 0.2387640449438202", label="4.39",style=bold];
"LRRC7_M" -> "PKHD1L1_M" [color="0 0 0.23820224719101124", label="4.41",style=bold];
"SERPINB3_E" -> "PGC_E" [color="0 0 0.23764044943820217", label="4.41",style=bold];
"ABCA13_S" -> "SLCO1B1_E" [color="0 0 0.23707865168539322", label="4.42",style=bold];
"SLC6A19_E" -> "RFX6_E" [color="0 0 0.23651685393258426", label="4.44",style=bold];
"SLC9A11_S" -> "CTSE_E" [color="0 0 0.2359550561797752", label="4.44",style=bold];
"PCK1_M" -> "SLC26A3_M" [color="0 0 0.23539325842696623", label="4.45",style=bold];
"GPR87_S" -> "MUC4_M" [color="0 0 0.23483146067415728", label="4.46",style=bold];
"OGDHL_S" -> "CSMD3_S" [color="0 0 0.23426966292134832", label="4.48",style=bold];
"CACNA1B_M" -> "PKHD1L1_E" [color="0 0 0.23370786516853925", label="4.51",style=bold];
"KRT16_E" -> "KRT6B_E" [color="0 0 0.2331460674157303", label="4.51",style=bold];
"CR2_S" -> "SLC9A11_E" [color="0 0 0.23258426966292134", label="4.53",style=bold];
"SLITRK1_M" -> "THBS4_M" [color="0 0 0.23202247191011227", label="4.53",style=bold];
"KRT16_E" -> "FAM83C_E" [color="0 0 0.23146067415730331", label="4.53",style=bold];
"EGF_S" -> "MYH4_S" [color="0 0 0.23089887640449436", label="4.53",style=bold];
"ZIC4_S" -> "LRRC7_M" [color="0 0 0.2303370786516854", label="4.55",style=bold];
"PNLIPRP2_S" -> "MDGA2_M" [color="0 0 0.22977528089887633", label="4.56",style=bold];
"RFX6_M" -> "APOB_M" [color="0 0 0.22921348314606738", label="4.59",style=bold];
"CPA2_S" -> "SLCO1B3_S" [color="0 0 0.22865168539325842", label="4.60",style=bold];
"COL11A1_S" -> "DCDC1_M" [color="0 0 0.22808988764044935", label="4.63",style=bold];
"PCLO_S" -> "SERPINB13_M" [color="0 0 0.2275280898876404", label="4.69",style=bold];
"TGM5_S" -> "S100A7_E" [color="0 0 0.22696629213483144", label="4.73",style=bold];
"KLK5_E" -> "FLG_S" [color="0 0 0.22640449438202248", label="4.73",style=bold];
"GPR87_S" -> "PGLYRP3_S" [color="0 0 0.2258426966292134", label="4.74",style=bold];
"DMBT1_S" -> "COL11A1_S" [color="0 0 0.22528089887640446", label="4.75",style=bold];
"PKHD1L1_S" -> "NETO1_M" [color="0 0 0.2247191011235955", label="4.76",style=bold];
"KRT13_S" -> "DMBT1_S" [color="0 0 0.22415730337078643", label="4.76",style=bold];
"RIMS2_M" -> "IRX6_M" [color="0 0 0.22359550561797747", label="4.77",style=bold];
"TGM5_M" -> "KLK5_M" [color="0 0 0.22303370786516852", label="4.77",style=bold];
"FRMPD2_S" -> "FSTL5_M" [color="0 0 0.22247191011235956", label="4.79",style=bold];
"APOB_S" -> "RFX6_S" [color="0 0 0.2219101123595505", label="4.80",style=bold];
"EGF_S" -> "ABCA12_S" [color="0 0 0.22134831460674154", label="4.81",style=bold];
"IRX6_S" -> "CR2_E" [color="0 0 0.22078651685393258", label="4.82",style=bold];
"DMBT1_S" -> "MUC16_S" [color="0 0 0.2202247191011235", label="4.82",style=bold];
"COL2A1_S" -> "PKHD1L1_E" [color="0 0 0.21966292134831455", label="4.83",style=bold];
"ADAMTS20_M" -> "SERPINB4_M" [color="0 0 0.2191011235955056", label="4.84",style=bold];
"APOB_M" -> "SLC6A19_M" [color="0 0 0.21853932584269664", label="4.88",style=bold];
"PCK1_M" -> "MUC15_M" [color="0 0 0.21797752808988757", label="4.94",style=bold];
"CR2_M" -> "RP1_E" [color="0 0 0.21741573033707862", label="4.97",style=bold];
"SLC13A2_S" -> "CNTNAP2_S" [color="0 0 0.21685393258426966", label="4.99",style=bold];
"KRT6C_E" -> "SERPINB3_E" [color="0 0 0.2162921348314606", label="5.00",style=bold];
"A2ML1_S" -> "OGDHL_S" [color="0 0 0.21573033707865163", label="5.03",style=bold];
"COL2A1_M" -> "MUC16_S" [color="0 0 0.21516853932584268", label="5.07",style=bold];
"KRT13_E" -> "MUC6_M" [color="0 0 0.21460674157303372", label="5.10",style=bold];
"FRMPD2_S" -> "MUC17_S" [color="0 0 0.21404494382022465", label="5.12",style=bold];
"KLK5_S" -> "MYH4_S" [color="0 0 0.2134831460674157", label="5.12",style=bold];
"ABCA13_S" -> "SLCO1B3_S" [color="0 0 0.21292134831460674", label="5.13",style=bold];
"KRT13_S" -> "LPA_S" [color="0 0 0.21235955056179767", label="5.17",style=bold];
"KCNA1_S" -> "MUC15_S" [color="0 0 0.21179775280898872", label="5.19",style=bold];
"CTSE_S" -> "RIMS2_S" [color="0 0 0.21123595505617976", label="5.20",style=bold];
"SLC6A19_S" -> "ABCA12_E" [color="0 0 0.2106741573033708", label="5.21",style=bold];
"SLC6A19_S" -> "CNTN1_E" [color="0 0 0.21011235955056173", label="5.24",style=bold];
"RFX6_M" -> "PCLO_E" [color="0 0 0.20955056179775278", label="5.31",style=bold];
"DPP6_M" -> "MUC5B_M" [color="0 0 0.20898876404494382", label="5.32",style=bold];
"SERPINB3_M" -> "FER1L6_M" [color="0 0 0.20842696629213475", label="5.36",style=bold];
"PCDH10_S" -> "GLRA1_E" [color="0 0 0.2078651685393258", label="5.37",style=bold];
"APOB_S" -> "ADAMTS20_E" [color="0 0 0.20730337078651684", label="5.37",style=bold];
"SLC26A3_E" -> "SLITRK1_E" [color="0 0 0.20674157303370788", label="5.41",style=bold];
"GPR98_E" -> "ABCA13_E" [color="0 0 0.2061797752808988", label="5.42",style=bold];
"COL11A1_S" -> "DPP6_E" [color="0 0 0.20561797752808986", label="5.42",style=bold];
"SERPINB4_S" -> "FLG_S" [color="0 0 0.2050561797752809", label="5.43",style=bold];
"DPP6_S" -> "APOB_E" [color="0 0 0.20449438202247183", label="5.45",style=bold];
"LRRC7_S" -> "SLITRK1_S" [color="0 0 0.20393258426966288", label="5.46",style=bold];
"MUC5B_M" -> "MUC6_M" [color="0 0 0.20337078651685392", label="5.50",style=bold];
"AGBL1_M" -> "LPA_M" [color="0 0 0.20280898876404485", label="5.55",style=bold];
"APOB_M" -> "PKHD1L1_M" [color="0 0 0.2022471910112359", label="5.55",style=bold];
"ZIC4_S" -> "MDGA2_M" [color="0 0 0.20168539325842694", label="5.55",style=bold];
"CR2_S" -> "PCDHA11_S" [color="0 0 0.20112359550561798", label="5.56",style=bold];
"SLC6A19_S" -> "CNTNAP2_S" [color="0 0 0.2005617977528089", label="5.57",style=bold];
"A2ML1_E" -> "ABCA13_E" [color="0 0 0.19999999999999996", label="5.59",style=bold];
"SLC13A2_S" -> "PCDH10_S" [color="0 0 0.199438202247191", label="5.60",style=bold];
"ABCA13_S" -> "KLK5_E" [color="0 0 0.19887640449438193", label="5.61",style=bold];
"DPP6_S" -> "SLC9A11_M" [color="0 0 0.19831460674157297", label="5.63",style=bold];
"MUC5B_E" -> "FER1L6_E" [color="0 0 0.19775280898876402", label="5.63",style=bold];
"PNLIPRP2_S" -> "PCK1_S" [color="0 0 0.19719101123595506", label="5.66",style=bold];
"ZIC4_S" -> "MUC2_E" [color="0 0 0.196629213483146", label="5.67",style=bold];
"SLITRK1_M" -> "PCDHA12_M" [color="0 0 0.19606741573033704", label="5.70",style=bold];
"SLC9A11_M" -> "OGDHL_E" [color="0 0 0.19550561797752808", label="5.72",style=bold];
"PNLIPRP2_M" -> "MUC17_M" [color="0 0 0.194943820224719", label="5.75",style=bold];
"ADAMTS20_E" -> "ZIC4_E" [color="0 0 0.19438202247191005", label="5.78",style=bold];
"AGBL1_S" -> "DPP6_E" [color="0 0 0.1938202247191011", label="5.83",style=bold];
"FREM2_S" -> "SLC26A3_E" [color="0 0 0.19325842696629214", label="5.88",style=bold];
"DPP6_M" -> "ABCA13_M" [color="0 0 0.19269662921348307", label="5.91",style=bold];
"ABCA13_S" -> "ACTN2_E" [color="0 0 0.19213483146067412", label="5.96",style=bold];
"PCDHA12_S" -> "PGLYRP3_E" [color="0 0 0.19157303370786516", label="6.00",style=bold];
"OGDHL_S" -> "KRT6C_S" [color="0 0 0.1910112359550561", label="6.01",style=bold];
"RIMS2_M" -> "ZIC4_M" [color="0 0 0.19044943820224713", label="6.02",style=bold];
"APOB_S" -> "RIMS2_S" [color="0 0 0.18988764044943818", label="6.10",style=bold];
"MUC4_S" -> "RIMS2_S" [color="0 0 0.18932584269662922", label="6.11",style=bold];
"ACTN2_S" -> "FREM2_E" [color="0 0 0.18876404494382015", label="6.12",style=bold];
"TGM5_S" -> "PGC_E" [color="0 0 0.1882022471910112", label="6.17",style=bold];
"SLC9A11_S" -> "CACNA1B_S" [color="0 0 0.18764044943820224", label="6.19",style=bold];
"PCDHA12_S" -> "PCLO_S" [color="0 0 0.18707865168539317", label="6.21",style=bold];
"TGM5_S" -> "PCK1_S" [color="0 0 0.18651685393258421", label="6.21",style=bold];
"A2ML1_E" -> "EGF_E" [color="0 0 0.18595505617977526", label="6.22",style=bold];
"FLG_M" -> "CSMD3_S" [color="0 0 0.1853932584269663", label="6.30",style=bold];
"RIMS2_E" -> "NETO1_E" [color="0 0 0.18483146067415723", label="6.32",style=bold];
"PCLO_E" -> "MUC15_E" [color="0 0 0.18426966292134828", label="6.34",style=bold];
"FER1L6_S" -> "RIMS2_S" [color="0 0 0.18370786516853932", label="6.38",style=bold];
"PTPRZ1_S" -> "APOB_S" [color="0 0 0.18314606741573025", label="6.38",style=bold];
"CTSE_M" -> "KRT13_M" [color="0 0 0.1825842696629213", label="6.41",style=bold];
"ACTN2_E" -> "KCNA1_E" [color="0 0 0.18202247191011234", label="6.48",style=bold];
"PGLYRP3_S" -> "ABCA12_S" [color="0 0 0.18146067415730338", label="6.49",style=bold];
"PTPRZ1_M" -> "DSG1_M" [color="0 0 0.1808988764044943", label="6.51",style=bold];
"MYH4_M" -> "MUC4_S" [color="0 0 0.18033707865168536", label="6.55",style=bold];
"SLCO1B1_E" -> "DCDC1_M" [color="0 0 0.1797752808988764", label="6.61",style=bold];
"RFX6_M" -> "RP1_E" [color="0 0 0.17921348314606733", label="6.62",style=bold];
"MUC4_M" -> "PCLO_S" [color="0 0 0.17865168539325837", label="6.65",style=bold];
"PCDHA12_S" -> "COL11A1_S" [color="0 0 0.17808988764044942", label="6.72",style=bold];
"LPA_M" -> "ABCA13_M" [color="0 0 0.17752808988764046", label="6.73",style=bold];
"MUC4_M" -> "KRT16_M" [color="0 0 0.1769662921348314", label="6.74",style=bold];
"MYH4_S" -> "CACNA1B_S" [color="0 0 0.17640449438202244", label="6.75",style=bold];
"DMBT1_E" -> "CACNA1B_E" [color="0 0 0.17584269662921348", label="6.76",style=bold];
"SERPINB4_S" -> "MUC17_S" [color="0 0 0.1752808988764044", label="6.79",style=bold];
"PCLO_M" -> "CSMD3_M" [color="0 0 0.17471910112359545", label="6.84",style=bold];
"LRRC7_M" -> "PCLO_M" [color="0 0 0.1741573033707865", label="6.85",style=bold];
"CACNA1B_M" -> "THBS4_E" [color="0 0 0.17359550561797754", label="6.85",style=bold];
"FREM2_M" -> "SLC9A11_E" [color="0 0 0.17303370786516847", label="6.87",style=bold];
"FAM83C_M" -> "GPR98_S" [color="0 0 0.17247191011235952", label="6.91",style=bold];
"SLC6A19_S" -> "DCDC1_S" [color="0 0 0.17191011235955056", label="6.94",style=bold];
"MUC4_M" -> "DSG1_M" [color="0 0 0.1713483146067415", label="6.97",style=bold];
"PCDHA11_E" -> "TGM5_M" [color="0 0 0.17078651685393254", label="7.00",style=bold];
"MDGA2_S" -> "FER1L6_E" [color="0 0 0.17022471910112358", label="7.04",style=bold];
"DCDC1_S" -> "SLC9A11_M" [color="0 0 0.1696629213483145", label="7.11",style=bold];
"SLITRK1_M" -> "Survival" [color="0 0 0.16910112359550555", label="7.12",style=bold];
"SLC9A4_S" -> "DMBT1_S" [color="0 0 0.1685393258426966", label="7.16",style=bold];
"DPP6_E" -> "LRRC7_E" [color="0 0 0.16797752808988764", label="7.17",style=bold];
"ABCA12_S" -> "CSMD3_S" [color="0 0 0.16741573033707857", label="7.18",style=bold];
"PCLO_M" -> "GPR98_M" [color="0 0 0.16685393258426962", label="7.22",style=bold];
"SERPINB13_E" -> "TGM5_E" [color="0 0 0.16629213483146066", label="7.25",style=bold];
"MUC16_E" -> "KRT6C_E" [color="0 0 0.1657303370786516", label="7.25",style=bold];
"SLITRK1_S" -> "FRMPD2_E" [color="0 0 0.16516853932584263", label="7.26",style=bold];
"KLK5_M" -> "KRT6C_M" [color="0 0 0.16460674157303368", label="7.29",style=bold];
"SLC6A19_M" -> "GPR87_M" [color="0 0 0.16404494382022472", label="7.34",style=bold];
"PGLYRP3_E" -> "KRT13_E" [color="0 0 0.16348314606741565", label="7.40",style=bold];
"AGBL1_M" -> "THBS4_E" [color="0 0 0.1629213483146067", label="7.46",style=bold];
"DPP6_M" -> "ABCA12_M" [color="0 0 0.16235955056179774", label="7.49",style=bold];
"MUC2_S" -> "MUC16_S" [color="0 0 0.16179775280898867", label="7.51",style=bold];
"PNLIPRP2_E" -> "MUC16_S" [color="0 0 0.16123595505617971", label="7.52",style=bold];
"LRRC7_M" -> "SLCO1B3_M" [color="0 0 0.16067415730337076", label="7.53",style=bold];
"FAM83C_S" -> "AGBL1_S" [color="0 0 0.1601123595505618", label="7.54",style=bold];
"PCLO_M" -> "PCLO_E" [color="0 0 0.15955056179775273", label="7.56",style=bold];
"COL11A1_S" -> "FBN3_S" [color="0 0 0.15898876404494378", label="7.63",style=bold];
"GPR87_S" -> "FBN3_S" [color="0 0 0.15842696629213482", label="7.70",style=bold];
"FLG_E" -> "PCDH10_E" [color="0 0 0.15786516853932575", label="7.74",style=bold];
"CTSE_S" -> "PCK1_S" [color="0 0 0.1573033707865168", label="7.75",style=bold];
"FER1L6_M" -> "PKHD1L1_M" [color="0 0 0.15674157303370784", label="7.76",style=bold];
"PKHD1L1_E" -> "CSMD3_E" [color="0 0 0.15617977528089888", label="7.77",style=bold];
"PCK1_M" -> "MUC5B_M" [color="0 0 0.1556179775280898", label="7.78",style=bold];
"PCDH10_S" -> "KRT13_E" [color="0 0 0.15505617977528086", label="7.82",style=bold];
"RIMS2_E" -> "CACNA1B_E" [color="0 0 0.1544943820224719", label="7.83",style=bold];
"THBS4_E" -> "IRX6_M" [color="0 0 0.15393258426966283", label="7.86",style=bold];
"OGDHL_M" -> "IRX6_M" [color="0 0 0.15337078651685387", label="7.87",style=bold];
"FAM83C_E" -> "SLC26A3_E" [color="0 0 0.15280898876404492", label="7.92",style=bold];
"FRMPD2_M" -> "FER1L6_M" [color="0 0 0.15224719101123596", label="7.93",style=bold];
"NETO1_S" -> "ABCA12_E" [color="0 0 0.1516853932584269", label="7.95",style=bold];
"FREM2_M" -> "CPA2_M" [color="0 0 0.15112359550561794", label="7.96",style=bold];
"LRRC7_M" -> "C20orf114_M" [color="0 0 0.15056179775280898", label="7.99",style=bold];
"ACTN2_M" -> "ADAMTS20_E" [color="0 0 0.1499999999999999", label="7.99",style=bold];
"FLG_M" -> "ABCA13_M" [color="0 0 0.14943820224719095", label="8.05",style=bold];
"DCDC1_S" -> "GPR98_S" [color="0 0 0.148876404494382", label="8.10",style=bold];
"KLK5_S" -> "OGDHL_S" [color="0 0 0.14831460674157304", label="8.11",style=bold];
"PKHD1L1_M" -> "EGF_M" [color="0 0 0.14775280898876397", label="8.12",style=bold];
"FBN3_M" -> "MUC17_S" [color="0 0 0.14719101123595502", label="8.13",style=bold];
"KRT16_E" -> "A2ML1_E" [color="0 0 0.14662921348314606", label="8.14",style=bold];
"PKHD1L1_M" -> "FREM2_M" [color="0 0 0.146067415730337", label="8.16",style=bold];
"MUC17_M" -> "SLC9A4_M" [color="0 0 0.14550561797752803", label="8.27",style=bold];
"PTPRZ1_E" -> "PTPRZ1_M" [color="0 0 0.14494382022471908", label="8.30",style=bold];
"S100A7_E" -> "KLK5_E" [color="0 0 0.14438202247191012", label="8.32",style=bold];
"SLCO1B1_S" -> "DCDC1_S" [color="0 0 0.14382022471910105", label="8.34",style=bold];
"A2ML1_M" -> "SLC26A3_M" [color="0 0 0.1432584269662921", label="8.44",style=bold];
"RIMS2_M" -> "COL2A1_E" [color="0 0 0.14269662921348314", label="8.46",style=bold];
"C20orf114_M" -> "PNLIPRP2_M" [color="0 0 0.14213483146067407", label="8.51",style=bold];
"ABCA12_S" -> "CR2_E" [color="0 0 0.14157303370786511", label="8.51",style=bold];
"KRT6C_S" -> "FBN3_S" [color="0 0 0.14101123595505616", label="8.55",style=bold];
"GLRA1_E" -> "AGBL1_E" [color="0 0 0.1404494382022472", label="8.62",style=bold];
"KLK5_M" -> "PNLIPRP2_M" [color="0 0 0.13988764044943813", label="8.62",style=bold];
"GPR87_E" -> "COL11A1_E" [color="0 0 0.13932584269662918", label="8.76",style=bold];
"DMBT1_M" -> "FER1L6_M" [color="0 0 0.13876404494382022", label="8.77",style=bold];
"A2ML1_E" -> "NETO1_E" [color="0 0 0.13820224719101115", label="8.84",style=bold];
"THBS4_S" -> "CACNA1B_S" [color="0 0 0.1376404494382022", label="8.91",style=bold];
"THBS4_M" -> "GPR98_M" [color="0 0 0.13707865168539324", label="8.96",style=bold];
"OGDHL_S" -> "CR2_S" [color="0 0 0.13651685393258417", label="9.02",style=bold];
"KRT13_E" -> "PCK1_E" [color="0 0 0.1359550561797752", label="9.02",style=bold];
"PTPRZ1_E" -> "FBN3_E" [color="0 0 0.13539325842696626", label="9.09",style=bold];
"CWH43_M" -> "THBS4_M" [color="0 0 0.1348314606741573", label="9.12",style=bold];
"CNTN1_M" -> "CACNA1B_E" [color="0 0 0.13426966292134823", label="9.13",style=bold];
"COL11A1_M" -> "FSTL5_M" [color="0 0 0.13370786516853927", label="9.18",style=bold];
"THBS4_E" -> "ACTN2_E" [color="0 0 0.13314606741573032", label="9.25",style=bold];
"PCLO_E" -> "FREM2_E" [color="0 0 0.13258426966292125", label="9.27",style=bold];
"PGC_M" -> "SLC9A4_M" [color="0 0 0.1320224719101123", label="9.28",style=bold];
"ACTN2_E" -> "MDGA2_E" [color="0 0 0.13146067415730334", label="9.29",style=bold];
"SERPINB4_E" -> "RIMS2_M" [color="0 0 0.13089887640449438", label="9.42",style=bold];
"KRT16_E" -> "ABCA12_E" [color="0 0 0.1303370786516853", label="9.46",style=bold];
"PKHD1L1_E" -> "RFX6_E" [color="0 0 0.12977528089887636", label="9.61",style=bold];
"KCNA1_M" -> "COL2A1_M" [color="0 0 0.1292134831460674", label="9.73",style=bold];
"MUC4_S" -> "GPR98_S" [color="0 0 0.12865168539325833", label="9.94",style=bold];
"PCLO_E" -> "OGDHL_E" [color="0 0 0.12808988764044937", label="9.99",style=bold];
"PGLYRP3_E" -> "EGF_E" [color="0 0 0.12752808988764042", label="10.02",style=bold];
"FLG_E" -> "S100A7_M" [color="0 0 0.12696629213483146", label="10.05",style=bold];
"SLITRK1_S" -> "SLCO1B1_E" [color="0 0 0.1264044943820224", label="10.09",style=bold];
"FAM83C_M" -> "KLK5_M" [color="0 0 0.12584269662921344", label="10.10",style=bold];
"TGM5_E" -> "IRX6_E" [color="0 0 0.12528089887640448", label="10.11",style=bold];
"AGBL1_E" -> "FRMPD2_E" [color="0 0 0.12471910112359541", label="10.14",style=bold];
"CSMD3_M" -> "PTPRZ1_M" [color="0 0 0.12415730337078645", label="10.17",style=bold];
"LRRC7_S" -> "PKHD1L1_S" [color="0 0 0.1235955056179775", label="10.20",style=bold];
"MUC5B_S" -> "CR2_E" [color="0 0 0.12303370786516854", label="10.25",style=bold];
"MUC2_S" -> "ABCA12_S" [color="0 0 0.12247191011235947", label="10.30",style=bold];
"SLC26A3_E" -> "RFX6_E" [color="0 0 0.12191011235955052", label="10.35",style=bold];
"FAM83C_M" -> "GPR87_M" [color="0 0 0.12134831460674156", label="10.44",style=bold];
"C20orf114_M" -> "PGC_M" [color="0 0 0.12078651685393249", label="10.70",style=bold];
"CSMD3_S" -> "MUC4_E" [color="0 0 0.12022471910112353", label="10.80",style=bold];
"KCNA1_M" -> "SLC26A3_M" [color="0 0 0.11966292134831458", label="10.83",style=bold];
"KRT6C_E" -> "A2ML1_E" [color="0 0 0.11910112359550562", label="11.08",style=bold];
"EGF_M" -> "DSG1_M" [color="0 0 0.11853932584269655", label="11.18",style=bold];
"MUC17_E" -> "PCK1_E" [color="0 0 0.1179775280898876", label="11.21",style=bold];
"SLC6A19_E" -> "SLC6A19_M" [color="0 0 0.11741573033707864", label="11.21",style=bold];
"MUC17_M" -> "SLC26A3_M" [color="0 0 0.11685393258426957", label="11.22",style=bold];
"FAM83C_S" -> "MUC2_S" [color="0 0 0.11629213483146061", label="11.23",style=bold];
"CTSE_M" -> "SLC9A11_E" [color="0 0 0.11573033707865166", label="11.35",style=bold];
"KRT6C_E" -> "GPR87_E" [color="0 0 0.1151685393258427", label="11.47",style=bold];
"ACTN2_M" -> "CPA2_M" [color="0 0 0.11460674157303363", label="11.53",style=bold];
"MUC2_M" -> "FLG_S" [color="0 0 0.11404494382022468", label="11.54",style=bold];
"FRMPD2_S" -> "ABCA13_S" [color="0 0 0.11348314606741572", label="11.59",style=bold];
"FSTL5_E" -> "CNTNAP2_E" [color="0 0 0.11292134831460665", label="11.60",style=bold];
"THBS4_E" -> "KCNA1_E" [color="0 0 0.1123595505617977", label="11.61",style=bold];
"IRX6_S" -> "LRRC7_S" [color="0 0 0.11179775280898874", label="11.63",style=bold];
"SERPINB3_M" -> "SLCO1B1_M" [color="0 0 0.11123595505617978", label="11.63",style=bold];
"PCDH10_M" -> "NETO1_M" [color="0 0 0.11067415730337071", label="11.67",style=bold];
"SLC26A3_M" -> "ABCA12_M" [color="0 0 0.11011235955056176", label="11.77",style=bold];
"A2ML1_M" -> "KRT16_M" [color="0 0 0.1095505617977528", label="11.78",style=bold];
"SLC9A4_E" -> "FER1L6_E" [color="0 0 0.10898876404494373", label="11.84",style=bold];
"SLC9A4_E" -> "CPA2_E" [color="0 0 0.10842696629213477", label="12.02",style=bold];
"AGBL1_M" -> "PKHD1L1_E" [color="0 0 0.10786516853932582", label="12.11",style=bold];
"KRT6B_E" -> "MYH4_M" [color="0 0 0.10730337078651686", label="12.20",style=bold];
"CR2_S" -> "FBN3_S" [color="0 0 0.10674157303370779", label="12.20",style=bold];
"PGLYRP3_E" -> "MUC15_E" [color="0 0 0.10617977528089884", label="12.20",style=bold];
"KRT16_E" -> "MUC6_E" [color="0 0 0.10561797752808988", label="12.26",style=bold];
"FAM83C_M" -> "KRT13_M" [color="0 0 0.10505617977528081", label="12.31",style=bold];
"TGM5_M" -> "MUC15_M" [color="0 0 0.10449438202247185", label="12.33",style=bold];
"GPR98_E" -> "EGF_E" [color="0 0 0.1039325842696629", label="12.36",style=bold];
"CR2_S" -> "PCDH10_S" [color="0 0 0.10337078651685383", label="12.41",style=bold];
"MYH4_M" -> "MUC15_M" [color="0 0 0.10280898876404487", label="12.42",style=bold];
"SLCO1B1_M" -> "AGBL1_M" [color="0 0 0.10224719101123592", label="12.50",style=bold];
"PKHD1L1_E" -> "THBS4_E" [color="0 0 0.10168539325842696", label="12.76",style=bold];
"C20orf114_M" -> "KRT16_M" [color="0 0 0.10112359550561789", label="12.85",style=bold];
"GLRA1_M" -> "CSMD3_M" [color="0 0 0.10056179775280893", label="12.89",style=bold];
"PGC_E" -> "SLC6A19_E" [color="0 0 0.09999999999999998", label="12.97",style=bold];
"SERPINB3_E" -> "SERPINB13_E" [color="0 0 0.09943820224719091", label="13.21",style=bold];
"SLITRK1_E" -> "LRRC7_E" [color="0 0 0.09887640449438195", label="13.23",style=bold];
"RIMS2_E" -> "IRX6_E" [color="0 0 0.098314606741573", label="13.29",style=bold];
"MYH4_M" -> "CPA2_M" [color="0 0 0.09775280898876404", label="13.39",style=bold];
"A2ML1_M" -> "ABCA12_M" [color="0 0 0.09719101123595497", label="13.53",style=bold];
"SLC13A2_M" -> "MUC17_M" [color="0 0 0.09662921348314601", label="13.55",style=bold];
"FREM2_E" -> "DMBT1_M" [color="0 0 0.09606741573033706", label="13.71",style=bold];
"GPR87_E" -> "ABCA12_E" [color="0 0 0.09550561797752799", label="13.91",style=bold];
"SLITRK1_E" -> "GLRA1_E" [color="0 0 0.09494382022471903", label="13.95",style=bold];
"EGF_E" -> "RP1_E" [color="0 0 0.09438202247191008", label="14.11",style=bold];
"MUC2_E" -> "PCK1_E" [color="0 0 0.09382022471910112", label="14.14",style=bold];
"KRT16_E" -> "GPR87_E" [color="0 0 0.09325842696629205", label="14.26",style=bold];
"PGLYRP3_M" -> "PGLYRP3_E" [color="0 0 0.0926966292134831", label="14.32",style=bold];
"MUC15_E" -> "S100A7_E" [color="0 0 0.09213483146067414", label="14.54",style=bold];
"SLC6A19_M" -> "FAM83C_M" [color="0 0 0.09157303370786507", label="14.59",style=bold];
"PCK1_M" -> "FRMPD2_M" [color="0 0 0.09101123595505611", label="14.72",style=bold];
"RFX6_M" -> "FREM2_M" [color="0 0 0.09044943820224716", label="14.76",style=bold];
"CTSE_E" -> "DMBT1_E" [color="0 0 0.0898876404494382", label="14.76",style=bold];
"CTSE_M" -> "PGC_M" [color="0 0 0.08932584269662913", label="14.80",style=bold];
"ADAMTS20_E" -> "FSTL5_E" [color="0 0 0.08876404494382018", label="14.89",style=bold];
"KCNA1_M" -> "GPR98_E" [color="0 0 0.08820224719101122", label="15.07",style=bold];
"FLG_M" -> "FREM2_M" [color="0 0 0.08764044943820215", label="15.30",style=bold];
"DMBT1_E" -> "COL11A1_E" [color="0 0 0.0870786516853932", label="15.39",style=bold];
"ZIC4_M" -> "FREM2_E" [color="0 0 0.08651685393258424", label="15.41",style=bold];
"OGDHL_M" -> "OGDHL_E" [color="0 0 0.08595505617977528", label="15.70",style=bold];
"SERPINB3_M" -> "RIMS2_M" [color="0 0 0.08539325842696621", label="15.71",style=bold];
"CTSE_M" -> "MUC5B_E" [color="0 0 0.08483146067415726", label="15.86",style=bold];
"SLCO1B3_E" -> "SLCO1B3_M" [color="0 0 0.0842696629213483", label="15.96",style=bold];
"CR2_M" -> "OGDHL_M" [color="0 0 0.08370786516853923", label="16.04",style=bold];
"DSG1_E" -> "FBN3_E" [color="0 0 0.08314606741573027", label="16.07",style=bold];
"MUC6_E" -> "MUC5B_E" [color="0 0 0.08258426966292132", label="16.16",style=bold];
"PCDH10_E" -> "ACTN2_E" [color="0 0 0.08202247191011236", label="16.21",style=bold];
"TGM5_M" -> "SLC9A11_M" [color="0 0 0.08146067415730329", label="16.26",style=bold];
"ADAMTS20_M" -> "KCNA1_M" [color="0 0 0.08089887640449434", label="16.30",style=bold];
"NETO1_E" -> "COL11A1_E" [color="0 0 0.08033707865168538", label="16.30",style=bold];
"PCDH10_E" -> "DPP6_E" [color="0 0 0.07977528089887631", label="16.51",style=bold];
"MUC2_M" -> "MUC6_M" [color="0 0 0.07921348314606735", label="16.62",style=bold];
"SERPINB13_E" -> "MUC6_E" [color="0 0 0.0786516853932584", label="16.66",style=bold];
"FAM83C_M" -> "FBN3_M" [color="0 0 0.07808988764044944", label="16.84",style=bold];
"FAM83C_E" -> "MUC15_E" [color="0 0 0.07752808988764037", label="16.91",style=bold];
"GPR87_E" -> "SLC26A3_E" [color="0 0 0.07696629213483142", label="17.01",style=bold];
"CWH43_E" -> "COL2A1_E" [color="0 0 0.07640449438202246", label="17.06",style=bold];
"CTSE_E" -> "FER1L6_E" [color="0 0 0.07584269662921339", label="17.19",style=bold];
"CACNA1B_M" -> "CR2_M" [color="0 0 0.07528089887640443", label="17.26",style=bold];
"FRMPD2_M" -> "RP1_M" [color="0 0 0.07471910112359548", label="17.65",style=bold];
"MUC4_M" -> "PNLIPRP2_M" [color="0 0 0.07415730337078652", label="17.66",style=bold];
"SERPINB3_E" -> "DSG1_E" [color="0 0 0.07359550561797745", label="18.21",style=bold];
"MUC17_E" -> "MUC17_M" [color="0 0 0.0730337078651685", label="18.28",style=bold];
"GPR87_E" -> "RIMS2_M" [color="0 0 0.07247191011235954", label="18.30",style=bold];
"MUC2_M" -> "SLC6A19_M" [color="0 0 0.07191011235955047", label="18.31",style=bold];
"SLC13A2_E" -> "CNTNAP2_E" [color="0 0 0.07134831460674151", label="18.37",style=bold];
"SLC26A3_E" -> "MYH4_E" [color="0 0 0.07078651685393256", label="18.50",style=bold];
"AGBL1_M" -> "PCK1_M" [color="0 0 0.07022471910112349", label="18.76",style=bold];
"MUC2_E" -> "MUC4_E" [color="0 0 0.06966292134831453", label="18.78",style=bold];
"ADAMTS20_E" -> "MDGA2_E" [color="0 0 0.06910112359550558", label="19.08",style=bold];
"SLC6A19_M" -> "KRT6B_M" [color="0 0 0.06853932584269662", label="19.15",style=bold];
"GPR98_E" -> "ADAMTS20_E" [color="0 0 0.06797752808988755", label="19.50",style=bold];
"PCK1_M" -> "MUC2_M" [color="0 0 0.0674157303370786", label="19.51",style=bold];
"SLC26A3_E" -> "PNLIPRP2_E" [color="0 0 0.06685393258426964", label="19.61",style=bold];
"KCNA1_M" -> "CTSE_E" [color="0 0 0.06629213483146057", label="19.63",style=bold];
"SERPINB3_E" -> "MUC4_E" [color="0 0 0.06573033707865161", label="19.66",style=bold];
"ZIC4_M" -> "SLITRK1_M" [color="0 0 0.06516853932584266", label="19.76",style=bold];
"SERPINB13_E" -> "GPR98_E" [color="0 0 0.0646067415730337", label="19.80",style=bold];
"GPR87_M" -> "MYH4_M" [color="0 0 0.06404494382022463", label="19.88",style=bold];
"CTSE_E" -> "SLC26A3_E" [color="0 0 0.06348314606741567", label="20.05",style=bold];
"ZIC4_M" -> "PCDH10_M" [color="0 0 0.06292134831460672", label="20.05",style=bold];
"SERPINB3_M" -> "AGBL1_M" [color="0 0 0.06235955056179765", label="20.38",style=bold];
"MDGA2_M" -> "LRRC7_M" [color="0 0 0.06179775280898869", label="20.49",style=bold];
"SERPINB4_M" -> "CNTNAP2_E" [color="0 0 0.061235955056179736", label="20.54",style=bold];
"MUC16_E" -> "MUC16_M" [color="0 0 0.06067415730337078", label="20.64",style=bold];
"OGDHL_M" -> "COL2A1_M" [color="0 0 0.06011235955056171", label="20.67",style=bold];
"PGLYRP3_M" -> "MUC6_E" [color="0 0 0.059550561797752755", label="20.68",style=bold];
"AGBL1_M" -> "MYH4_M" [color="0 0 0.0589887640449438", label="20.77",style=bold];
"SERPINB13_E" -> "FAM83C_E" [color="0 0 0.05842696629213473", label="21.37",style=bold];
"ZIC4_M" -> "COL11A1_M" [color="0 0 0.05786516853932577", label="21.48",style=bold];
"MUC17_E" -> "DMBT1_E" [color="0 0 0.057303370786516816", label="21.53",style=bold];
"RFX6_M" -> "DCDC1_M" [color="0 0 0.05674157303370786", label="21.58",style=bold];
"ACTN2_E" -> "DPP6_E" [color="0 0 0.05617977528089879", label="21.67",style=bold];
"THBS4_E" -> "APOB_E" [color="0 0 0.055617977528089835", label="21.69",style=bold];
"LRRC7_M" -> "A2ML1_M" [color="0 0 0.05505617977528088", label="21.72",style=bold];
"PCDH10_M" -> "CWH43_M" [color="0 0 0.05449438202247181", label="21.86",style=bold];
"FBN3_M" -> "SLC13A2_M" [color="0 0 0.05393258426966285", label="22.05",style=bold];
"C20orf114_M" -> "KRT6B_M" [color="0 0 0.053370786516853896", label="22.22",style=bold];
"KRT6B_E" -> "RP1_E" [color="0 0 0.05280898876404494", label="22.24",style=bold];
"FRMPD2_M" -> "EGF_M" [color="0 0 0.05224719101123587", label="22.26",style=bold];
"SLC6A19_E" -> "SLC13A2_E" [color="0 0 0.051685393258426915", label="23.07",style=bold];
"FLG_E" -> "RIMS2_E" [color="0 0 0.05112359550561796", label="23.51",style=bold];
"SLC26A3_E" -> "SLC13A2_E" [color="0 0 0.05056179775280889", label="23.53",style=bold];
"CNTN1_E" -> "PTPRZ1_E" [color="0 0 0.04999999999999993", label="23.56",style=bold];
"PGLYRP3_M" -> "FLG_M" [color="0 0 0.04943820224719098", label="23.74",style=bold];
"MUC17_E" -> "RIMS2_E" [color="0 0 0.04887640449438202", label="23.90",style=bold];
"CNTNAP2_M" -> "FBN3_E" [color="0 0 0.04831460674157295", label="24.79",style=bold];
"CACNA1B_M" -> "SERPINB13_M" [color="0 0 0.047752808988763995", label="24.94",style=bold];
"DPP6_E" -> "FSTL5_E" [color="0 0 0.04719101123595504", label="25.16",style=bold];
"FLG_M" -> "LPA_M" [color="0 0 0.04662921348314597", label="25.38",style=bold];
"ADAMTS20_M" -> "NETO1_M" [color="0 0 0.046067415730337014", label="25.39",style=bold];
"SERPINB3_M" -> "SERPINB4_M" [color="0 0 0.04550561797752806", label="25.69",style=bold];
"KRT16_E" -> "GPR87_M" [color="0 0 0.0449438202247191", label="25.99",style=bold];
"APOB_M" -> "CNTNAP2_M" [color="0 0 0.04438202247191003", label="26.06",style=bold];
"SLITRK1_M" -> "GLRA1_M" [color="0 0 0.043820224719101075", label="26.11",style=bold];
"CTSE_E" -> "MUC17_E" [color="0 0 0.04325842696629212", label="26.21",style=bold];
"KRT6C_E" -> "KRT6B_E" [color="0 0 0.04269662921348305", label="26.65",style=bold];
"CACNA1B_M" -> "FBN3_M" [color="0 0 0.042134831460674094", label="26.79",style=bold];
"C20orf114_M" -> "CTSE_M" [color="0 0 0.04157303370786514", label="27.13",style=bold];
"ADAMTS20_E" -> "SLCO1B1_E" [color="0 0 0.04101123595505618", label="27.64",style=bold];
"SLC26A3_E" -> "MUC2_E" [color="0 0 0.04044943820224711", label="27.85",style=bold];
"MUC17_E" -> "SLC6A19_E" [color="0 0 0.039887640449438155", label="28.49",style=bold];
"LPA_M" -> "LPA_E" [color="0 0 0.0393258426966292", label="29.01",style=bold];
"RIMS2_M" -> "ACTN2_M" [color="0 0 0.03876404494382013", label="29.07",style=bold];
"KRT16_M" -> "FAM83C_M" [color="0 0 0.038202247191011174", label="29.13",style=bold];
"DPP6_M" -> "ACTN2_M" [color="0 0 0.03764044943820222", label="29.16",style=bold];
"TGM5_M" -> "SERPINB3_M" [color="0 0 0.03707865168539326", label="29.22",style=bold];
"MDGA2_E" -> "SLITRK1_E" [color="0 0 0.03651685393258419", label="29.57",style=bold];
"TGM5_M" -> "FRMPD2_M" [color="0 0 0.035955056179775235", label="29.62",style=bold];
"KRT6B_M" -> "KRT6C_M" [color="0 0 0.03539325842696628", label="29.82",style=bold];
"PGLYRP3_M" -> "SLC13A2_M" [color="0 0 0.03483146067415721", label="30.26",style=bold];
"PCK1_M" -> "C20orf114_M" [color="0 0 0.034269662921348254", label="30.43",style=bold];
"PGC_E" -> "SLC9A4_E" [color="0 0 0.0337078651685393", label="30.79",style=bold];
"MYH4_E" -> "CPA2_E" [color="0 0 0.03314606741573023", label="30.85",style=bold];
"OGDHL_M" -> "FSTL5_M" [color="0 0 0.03258426966292127", label="31.29",style=bold];
"A2ML1_E" -> "CNTN1_E" [color="0 0 0.032022471910112316", label="31.43",style=bold];
"MUC5B_E" -> "MUC2_E" [color="0 0 0.03146067415730336", label="31.82",style=bold];
"SLC6A19_M" -> "SLC9A4_M" [color="0 0 0.03089887640449429", label="31.82",style=bold];
"ADAMTS20_E" -> "DCDC1_E" [color="0 0 0.030337078651685334", label="32.02",style=bold];
"SERPINB13_E" -> "AGBL1_E" [color="0 0 0.029775280898876377", label="32.34",style=bold];
"DPP6_M" -> "CACNA1B_M" [color="0 0 0.02921348314606731", label="32.39",style=bold];
"KRT6B_E" -> "PTPRZ1_E" [color="0 0 0.028651685393258353", label="32.42",style=bold];
"CNTNAP2_M" -> "RP1_M" [color="0 0 0.028089887640449396", label="33.00",style=bold];
"KRT6C_E" -> "KRT13_E" [color="0 0 0.02752808988764044", label="33.19",style=bold];
"KRT6C_E" -> "SERPINB13_E" [color="0 0 0.02696629213483137", label="33.34",style=bold];
"SLC26A3_E" -> "APOB_E" [color="0 0 0.026404494382022414", label="33.45",style=bold];
"MUC2_M" -> "CACNA1B_M" [color="0 0 0.025842696629213457", label="33.88",style=bold];
"AGBL1_M" -> "FLG_M" [color="0 0 0.02528089887640439", label="34.02",style=bold];
"DPP6_M" -> "MDGA2_M" [color="0 0 0.024719101123595433", label="34.10",style=bold];
"FAM83C_E" -> "DSG1_E" [color="0 0 0.024157303370786476", label="34.99",style=bold];
"PCK1_M" -> "A2ML1_M" [color="0 0 0.02359550561797752", label="35.13",style=bold];
"SERPINB4_E" -> "S100A7_E" [color="0 0 0.02303370786516845", label="35.40",style=bold];
"RIMS2_M" -> "RFX6_M" [color="0 0 0.022471910112359494", label="35.92",style=bold];
"RIMS2_M" -> "CWH43_M" [color="0 0 0.021910112359550538", label="36.93",style=bold];
"KRT6C_E" -> "KRT16_E" [color="0 0 0.02134831460674147", label="38.22",style=bold];
"SLC26A3_E" -> "MUC17_E" [color="0 0 0.020786516853932513", label="40.44",style=bold];
"PCK1_M" -> "S100A7_M" [color="0 0 0.020224719101123556", label="40.63",style=bold];
"DPP6_M" -> "GLRA1_M" [color="0 0 0.0196629213483146", label="40.75",style=bold];
"RIMS2_M" -> "OGDHL_M" [color="0 0 0.01910112359550553", label="40.92",style=bold];
"PCDH10_E" -> "CNTN1_E" [color="0 0 0.018539325842696575", label="40.93",style=bold];
"SLITRK1_M" -> "PCDH10_M" [color="0 0 0.017977528089887618", label="42.12",style=bold];
"CTSE_M" -> "SLC9A11_M" [color="0 0 0.01741573033707855", label="42.65",style=bold];
"MDGA2_E" -> "CSMD3_E" [color="0 0 0.016853932584269593", label="42.98",style=bold];
"SERPINB3_M" -> "SERPINB13_M" [color="0 0 0.016292134831460636", label="43.14",style=bold];
"PGC_E" -> "CTSE_E" [color="0 0 0.01573033707865168", label="44.33",style=bold];
"CNTNAP2_M" -> "DPP6_M" [color="0 0 0.015168539325842612", label="44.34",style=bold];
"MUC5B_M" -> "MUC2_M" [color="0 0 0.014606741573033655", label="45.88",style=bold];
"THBS4_E" -> "PCDH10_E" [color="0 0 0.014044943820224698", label="49.53",style=bold];
"AGBL1_M" -> "MUC16_M" [color="0 0 0.01348314606741563", label="52.15",style=bold];
"CSMD3_M" -> "COL11A1_M" [color="0 0 0.012921348314606673", label="53.08",style=bold];
"SLC6A19_M" -> "DMBT1_M" [color="0 0 0.012359550561797716", label="53.19",style=bold];
"OGDHL_M" -> "PCLO_M" [color="0 0 0.01179775280898876", label="53.90",style=bold];
"IRX6_M" -> "ZIC4_M" [color="0 0 0.011235955056179692", label="55.03",style=bold];
"SLCO1B1_E" -> "SLCO1B3_E" [color="0 0 0.010674157303370735", label="56.14",style=bold];
"SERPINB13_E" -> "PGLYRP3_E" [color="0 0 0.010112359550561778", label="56.14",style=bold];
"RIMS2_M" -> "CNTNAP2_M" [color="0 0 0.00955056179775271", label="58.40",style=bold];
"MUC6_E" -> "PGC_E" [color="0 0 0.008988764044943753", label="59.76",style=bold];
"AGBL1_M" -> "APOB_M" [color="0 0 0.008426966292134797", label="60.33",style=bold];
"PKHD1L1_E" -> "CR2_E" [color="0 0 0.00786516853932584", label="61.22",style=bold];
"CACNA1B_M" -> "MUC4_M" [color="0 0 0.007303370786516772", label="62.15",style=bold];
"KRT6C_E" -> "FLG_E" [color="0 0 0.006741573033707815", label="62.36",style=bold];
"PGC_E" -> "C20orf114_E" [color="0 0 0.006179775280898858", label="63.07",style=bold];
"KRT6C_E" -> "SERPINB4_E" [color="0 0 0.00561797752808979", label="65.44",style=bold];
"RFX6_M" -> "CR2_M" [color="0 0 0.0050561797752808335", label="65.49",style=bold];
"SERPINB13_E" -> "CWH43_E" [color="0 0 0.004494382022471877", label="67.34",style=bold];
"KCNA1_M" -> "SLITRK1_M" [color="0 0 0.00393258426966292", label="67.57",style=bold];
"CTSE_E" -> "CTSE_M" [color="0 0 0.003370786516853852", label="72.20",style=bold];
"SERPINB4_E" -> "SERPINB3_E" [color="0 0 0.002808988764044895", label="72.75",style=bold];
"S100A7_M" -> "PGLYRP3_M" [color="0 0 0.0022471910112359383", label="74.86",style=bold];
"ZIC4_M" -> "ADAMTS20_M" [color="0 0 0.0016853932584268705", label="96.13",style=bold];
"RIMS2_M" -> "KCNA1_M" [color="0 0 0.0011235955056179137", label="96.43",style=bold];
"PCDHA11_E" -> "PCDHA12_E" [color="0 0 0.0005617977528089568", label="102.46",style=bold];
"PCDHA12_M" -> "PCDHA11_M" [color="0 0 0.0", label="190.45",style=bold];
}
```
